# The crisprVerse: a comprehensive Bioconductor ecosystem for the design of CRISPR guide RNAs across nucleases and technologies

**DOI:** 10.1101/2022.04.21.488824

**Authors:** Luke Hoberecht, Pirunthan Perampalam, Aaron Lun, Jean-Philippe Fortin

**Affiliations:** Genentech Research and Early Development, Genentech, Inc., 1 DNA Way, South San Francisco, CA, 94080, USA; ProCogia Inc. under contract to Hoffmann-La Roche Limited

**Keywords:** CRISPR, Nuclease, gRNA, functional genomics, Sequence design, Bioconductor, Base Editor, Cas9, Cas12a, Cas13d, crisprVerse

## Abstract

The success of CRISPR-mediated gene perturbation studies is highly dependent on the quality of gRNAs, and several tools have been developed to enable optimal gRNA design. However, these tools are not all adaptable to the latest CRISPR modalities or nucleases, nor do they offer comprehensive annotation methods for advanced CRISPR applications. Here, we present a new ecosystem of R packages, called *crisprVerse*, that enables efficient gRNA design and annotation for a multitude of CRISPR technologies. This includes CRISPR knockout (CRISPRko), CRISPR activation (CRISPRa), CRISPR interference (CRISPRi), CRISPR base editing (CRISPRbe) and CRISPR knockdown (CRISPRkd). The core package, *crisprDesign*, offers a comprehensive, user-friendly, and unified interface to add on- and off-target annotations via several alignment methods, rich gene and SNP annotations, and a dozen on- and off-target activity scores. These functionalities are enabled for any RNA- or DNA-targeting nucleases, including Cas9, Cas12, and Cas13. We illustrate the general applicability of our tools by designing optimal gRNAs for three case studies: tiling CRISPRbe library for *BRCA1* using the base editor BE4max, tiling RNA-targeting libraries for *CD46* and *CD55* using CasRx, and activation of *MMP7* using CRISPRa. The *crisprVerse* ecosystem is open-source and deployed through the Bioconductor project to facilitate its use by the CRISPR community (https://github.com/crisprVerse).

## 1 Main

The performance of CRISPR-based experiments depends critically on the choice of the guide RNAs (gRNAs) used to guide the CRISPR nuclease to the target site. Variable gRNA on-target activity, as well as unintended off-targeting effects, can lead to inconsistent phenotypic readouts in screening experiments. For the purpose of analyzing pooled screens, many approaches attempt to model gRNA quality in the generation of gene-level scores to improve statistical inference [Meyers et al., 2017, Kim and Hart, 2021, Dempster et al., 2021, Allen et al., 2019, Li et al., 2015]. However, suboptimal gRNA design is only partially mitigated by analysis strategies that sacrifice statistical power for robustness to suboptimal guides. One way to increase the signal-to-noise ratio in screening experiments is to enrich gRNA libraries for gRNAs that have high predicted on-target activity. Predicting on-target activity from the spacer sequence is an extensive area of research, and several algorithms leveraging experimental data have been developed for different nucleases and contexts [Doench et al., 2016b, 2014, 2016a, Wang et al., 2019b, Kim et al., 2018, Hart et al., 2017a, Moreno-Mateos et al., 2015, Horlbeck et al., 2016].

In addition to its sequence, the genomic context of the on- and off-target sites for each gRNA is another important consideration for gRNA design. For example, designing gRNAs that uniquely map to the genome can be challenging, especially for genes sharing high homology with other genomic loci, either in coding or non-coding regions [Fortin et al., 2019]. Furthermore, knowing whether or not an off-target is located in the coding region of another gene can rule out the use of a given gRNA. Finally, genetic variation, such as single-nucleotide polymorphisms (SNPs) and small indels, can have a direct impact on gRNA binding activity and on-target specificity by altering complementarity between spacer sequences and the host cell genomic DNA [Scott and Zhang, 2017, Lessard et al., 2017, Canver et al., 2017, Wang et al., 2018].

The rapid increase of CRISPR-based applications and technologies poses another challenge to gRNA library design. A large variety of nucleases are now available and routinely used, including engineered nucleases that recognize a larger set of PAM sequences [Hu et al., 2018, Nishimasu et al., 2018, Walton et al., 2020, DeWeirdt et al., 2020] and novel classes of nucleases such as the RNA-targeting Cas13 family [Shmakov et al., 2015, Abudayyeh et al., 2016, Konermann et al., 2018]. Each nuclease comes with its own set of gRNA design rules and constraints. In addition, these nucleases can also be mixed and matched with different types of CRISPR applications, increasing the complexity of gRNA design. As an example, CRISPR base editing (CRISPRbe) [Gaudelli et al., 2017, Komor et al., 2016], which requires additional gRNA design functionalities to capture the editing window and prediction of editing outcomes, can be combined with the Cas13 family to perform RNA editing [Cox et al., 2017]. Finally, emerging screening modalities, such as optical pooled CRISPR screening [Feldman et al., 2019] and gRNA pairing, require additional specialized gRNA design considerations.

Given the complexity, heterogeneity, and fast growth of the aforementioned CRISPR modalities and applications, it is paramount to develop and maintain adaptable, modular, and robust software for gRNA design. This ensures that the scientific community can efficiently design first-class CRISPR reagents in a timely manner for both well-established and emerging technologies. An ideal gRNA design framework has the following qualities: (1) it offers multiple cutting-edge methods for on-target scoring and off-target prediction based on gRNA sequences, (2) it provides comprehensive gRNA annotation to enable consideration of the genomic context for all gRNA on-target and off-target sites, (3) it already supports (or can be easily extended to) newer CRISPR technologies, including an arbitrary combination of nucleases and modalities, and (4) it easily scales for designing large-scale gRNA libraries for different screening platforms. While a multitude of web applications and command line interfaces has been developed to enable gene- or other target-specific gRNA design [Heigwer et al., 2014, Moreno-Mateos et al., 2015, Perez et al., 2017, Bae et al., 2014, Montague et al., 2014, Concordet and Haeussler, 2018, Stemmer et al., 2015, McKenna and Shendure, 2018, Heigwer et al., 2016, Bhagwat et al., 2020, Zhu et al., 2014], none of the existing tools completely satisfies the requirements listed above.

In this work, we describe a modular ecosystem of R packages, called *crisprVerse*, that enable the design of CRISPR gRNAs across a variety of nucleases, genomes, and applications. The *crisprBowtie* and *crisprBwa* packages provide comprehensive on-target and off-target search for reference genomes, transcriptomes, or any custom sequences. The *crisprScore* package provides a harmonized framework to access a large array of R- and Python-based gRNA scoring algorithms developed by the CRISPR community, for both on-target and off-target scoring. The *crisprBase* package implements functionalities to describe and represent DNA- and RNA-targeting CRISPR nucleases, nickases and base editors, as well as genomic arithmetic rules that are specific to CRISPR design. The package *crisprDesign* provides a user-friendly package to design and annotate gRNAs in one place, including gene and TSS annotation, search for SNP overlap, addition of evolutionary conservation scores, characterization of edited alleles for base editors, sequence-based design rules, and library design functionalities such as ranking and platform-specific considerations. Finally, the package *crisprViz* allows users to visualize gRNAs within genomic tracks, with the option of embedding additional genomic annotations such as SNPs, repeat elements, or chromatin accessibility data. The *crisprVerse* ecosystem currently supports five different CRISPR modalities: CRISPR knockout (CRISPRko), CRISPR activation (CRISPRa), CRISPR interference (CRISPRi), CRISPR base editing (CRISPRbe) and CRISPR knockdown (CRISPRkd) using Cas13.

We illustrate the rich functionalities of our ecosystem through three case studies: designing gRNAs to edit *BRCA1* using the base editor BE4max, designing gRNAs to knock down *CD55* and *CD46* using CasRx, and designing optimal gRNAs to activate *MMP7* through CRISPRa for different wildtype and engineered nucleases.

We also show that our default gRNA ranking criteria yield optimal gRNAs by reanalyzing five genome-wide fitness screening datasets. Our R packages are open-source and deployed through the Bioconductor project [Gentleman et al., 2004, Huber et al., 2015]. This makes our tools fully interoperable with other packages, and facilitates long-term development and maintenance of our ecosystem. Source code, tutorials, and extensive documentation are provided on our website: https://github.com/crisprVerse.

## 2 Results

### 2.1 *crisprBase* as a core infrastructure package to represent CRISPR nucleases and base editors

The *crisprBase* package implements a common framework in the *crisprVerse* ecosystem for representing and manipulating nucleases and base editors through a set of classes and CRISPR-specific genome arithmetic functions. The *CrisprNuclease* class provides a general representation of a CRISPR nuclease, encoding all of the information necessary to perform gRNA design and other analyses involving CRISPR technologies. This includes the PAM side with respect to the protospacer sequence, recognized PAM sequences with optional tolerance weights, and the relative cut site. Specific *CrisprNuclease* instances can be easily created to represent a diversity of wildtype and engineered CRISPR nucleases (Figure 1). We also implement a *BaseEditor* subclass that provides additional base editing information such as the editing strand and a matrix of editing probabilities for possible nucleotide substitutions.

**Figure 1.**
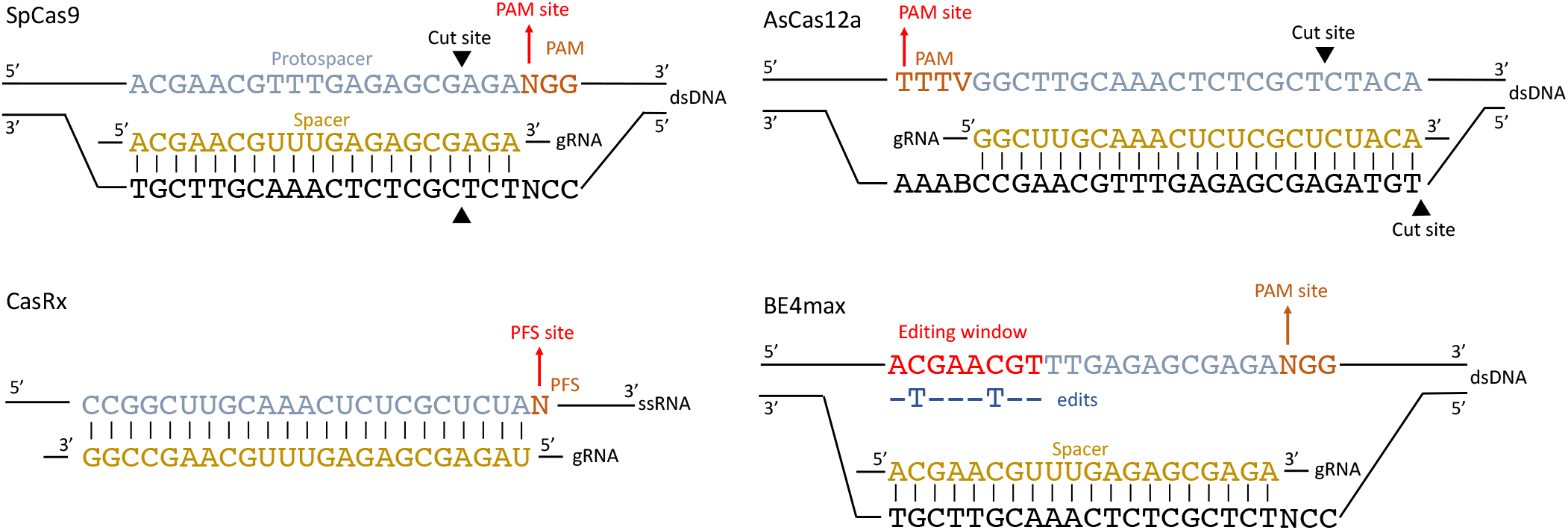
Examples of DNA- and RNA-targeting nucleases represented in *crisprBase*. gRNA spacer sequences are shown in yellow. Target DNA/RNA protospacer sequences are shown in blue. Protospacer adjacent motifs (PAMs) and protospacer flanking sequences (PFSs) are shown in orange. Nuclease-specific cutting sites are represented by black triangles. For the C to T base editor BE4max, on-target editing happens on the DNA strand containing the protospacer sequence. The editing window varies by base editor. The first nucleotide of the PAM/PFS is used as the representative coordinate of a given target sequence.

### 2.2 *crisprDesign* : a comprehensive tool to perform complex gRNA design

*crisprDesign* offers a comprehensive suite of methods to design and annotate gRNAs (see Table 1) and represents the core package of the *crisprVerse* ecosystem. For users, the package provides a centralized and streamlined end-to-end workflow for gRNA design, alleviating the burden of using different tools at different stages of the design process. For developers, *crisprDesign* is built on top of a modular package ecosystem that implements the gRNA design tasks (see Table 4 in the Methods section), allowing the same code to be easily re-used outside of CRISPR applications and gRNA design.

**Table 1.**
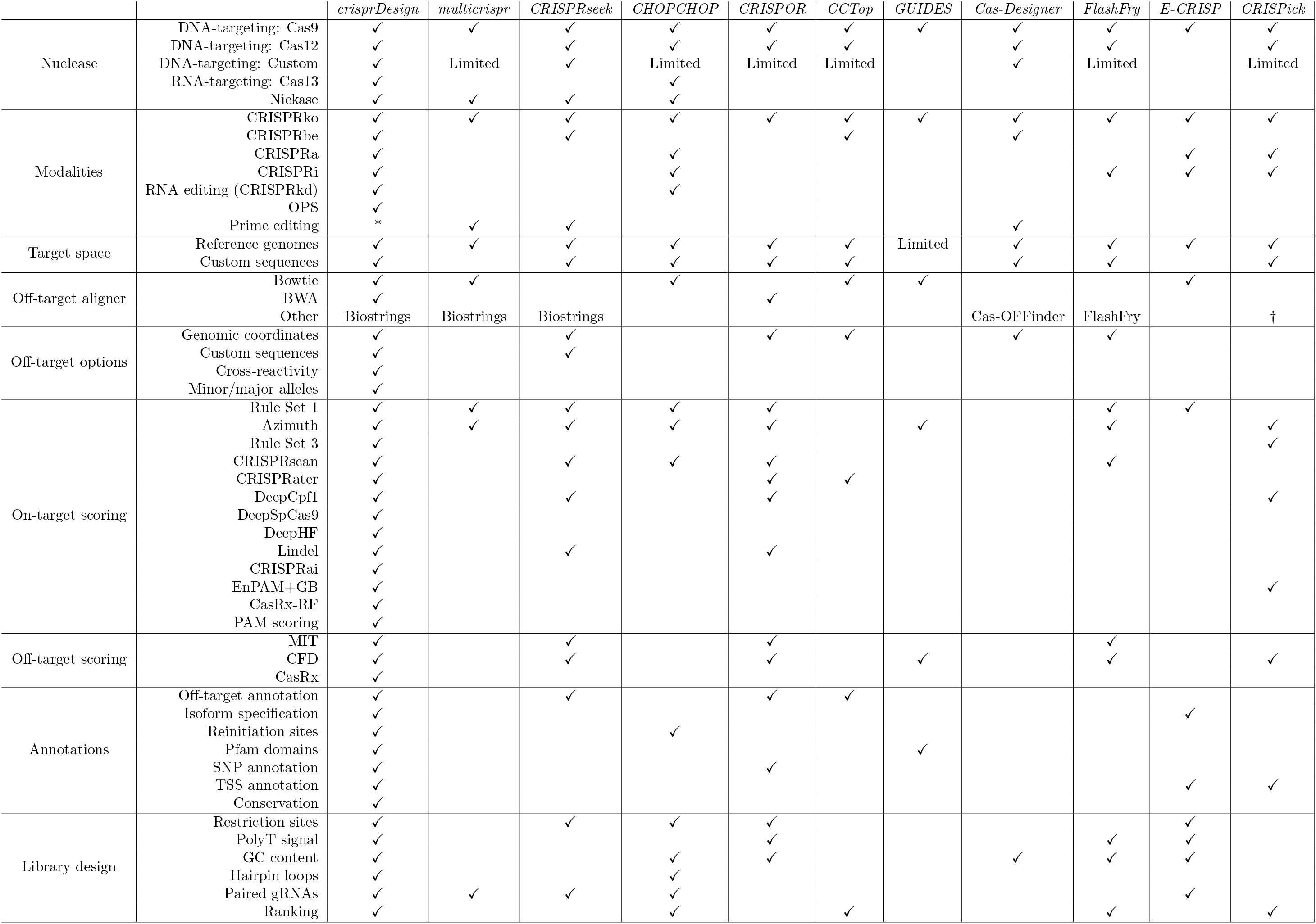
gRNA design functionalities implemented in *crisprVerse* and commonly-used gRNA design tools. Check marks indicate which functionalities are present in each tool at time of publication. *\** In progress. † Information could not be found. See the Methods section for a detailed description of the criteria used for assessing feature availability.

Table 1 includes a comparison to ten commonly-used gRNA design softwares: *multicrispr* [Bhagwat et al., 2020], *CRISPRseek* [Zhu et al., 2014], *CHOPCHOP* [Montague et al., 2014], *CRISPOR* [Concordet and Haeussler, 2018], *CCTop* [Stemmer et al., 2015], *GUIDES* [Meier et al., 2017], *Cas-Designer* [Park et al., 2015], *FlashFry* [McKenna and Shendure, 2018], *E-CRISP* [Heigwer et al., 2014] and CRISPick; see the Methods section for a detailed description of the criteria used for benchmarking. While several of the features implemented in *crisprDesign* are also available in other tools, *crisprDesign* provides the most complete gRNA design solution across nucleases and modalities. Unlike *crisprDesign*, many of the other tools do not provide informative on- and off-target annotations, limiting their use for optimal gRNA selection. In the following sections, we describe each of the gRNA design components and functionalities that are available in *crisprDesign*.

### 2.3 Representation of gRNAs using the *GuideSet* container

The genomic coordinates of gRNA protospacer sequences in a target genome can be represented using genomic ranges. The Bioconductor project [Gentleman et al., 2004, Huber et al., 2015] provides a robust and well-developed core data structure, called *GRanges* [Lawrence et al., 2013, Aboyoun et al., 2012]), to efficiently represent genomic intervals. We provide in *crisprDesign* an extension of the *GRanges* class to represent and annotate gRNA sequences: the *GuideSet* container. Briefly, the container extends the *GRanges* object to store additional project-specific metadata information, such as the CRISPR nuclease employed and target mRNA or DNA sequences (if different from a reference genome), as well as rich gRNA-level annotation columns such as on- and off-target alignments tables and gene context annotations. In Figure 2, we show an example of a *GuideSet* storing information about gRNAs targeting the coding sequence of *KRAS* using SpCas9.

**Figure 2.**
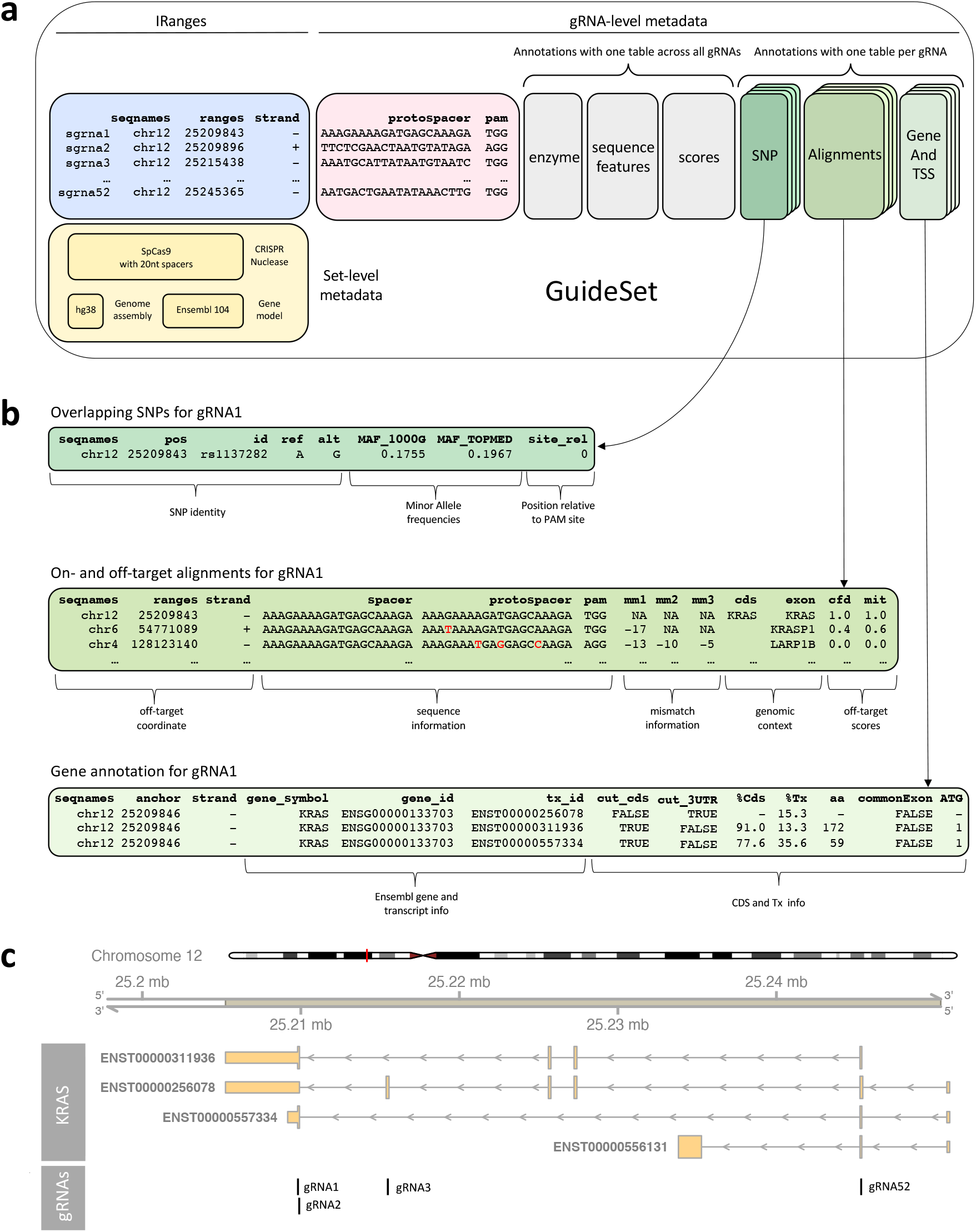
Example of a GuideSet container for gRNAs targeting *KRAS* using SpCas9. **a** The blue box stores the genomic coordinates in GRCh38 to represent the target protospacer sequences using a GRanges object. By convention, we use the first nucleotide of the PAM sequence (in the 5′ to 3′ direction) as the representative genomic coordinate of protospacer sequences. The pink box stores sequence information of the protospacers and PAMs. The yellow box represents global metadata used for creating the GuideSet, including a formal *CrisprNuclease* object, the reference genome of the target protospacers, and gene model used for annotation. The grey boxes are examples of optional gRNA-level metadata columns that store information about enzyme restriction sites, spacer sequence features such as GC content, and on- and off-target scores. The green boxes represent optional per-gRNA annotations for SNP overlap, on- and off-target alignments, and gene context; each annotation stores a detailed table (2 dimensions) for each gRNA (3rd dimension). **b** Selected annotations for gRNA 1 corresponding to the row highlighted in the green boxes of **a. c** The first genomic track represents the four annotated protein-coding isoforms of human gene *KRAS* in GRCh38 coordinates. The second track shows the 4 gRNAs shown in the blue box of **a**.

### 2.4 *crisprScore* implements state-of-the-art scoring methods

Predicting on-target binding and cutting efficiency of gRNAs is an extensive area of research. Many algorithms have been developed to tackle this problem, basing their prediction on a variety of features: sequence composition of the spacer sequence and flanking regions, including nucleotide content and melting temperature, cell type-specific chromatin accessibility data, and distance to transcription starting site (TSS). Unfortunately, the heterogeneity in the algorithm implementations hinders the practical use of those algorithms: some methods are implemented in Python 2 [Doench et al., 2016a, Kim et al., 2018, Horlbeck et al., 2016, Kim et al., 2019], in Python 3 [Chen et al., 2019, Wang et al., 2019b, DeWeirdt et al., 2020], or in R [Doench et al., 2014, Wessels et al., 2020, Moreno-Mateos et al., 2015, Labuhn et al., 2018]. In addition, the required inputs, data structures, and terminology are not consistent across software and algorithms, increasing the likelihood of user error. Finally, several of the algorithms are currently not bundled up into easy-to-use packages, limiting their accessibility and therefore their usage.

To resolve this, we created a general and harmonized framework for on-target and off-target prediction of gRNAs, implemented in our R package *crisprScore*. The philosophy behind *crisprScore* is to abstract away from the user the language, implementation, and complexity of the different algorithms used for prediction. It uses the Bioconductor package *basilisk* [Lun, 2021] to seamlessly integrate and manage incompatible Python modules in one user session. This enables *crisprScore* to centralize all Python-based scoring algorithms together with R-based prediction algorithms, reporting all scores in a single data frame for convenient inspection. By providing a harmonized user interface, our framework facilitates methods comparison.

We note that while the package provides a harmonized framework from a user perspective, it also allows each scoring algorithm to be implemented with its own sets of parameters and inputs. We have included as many methods as possible (Table 2), with the goal of democratizing the use of different scoring algorithms in an unbiased manner. Developers can easily contribute new methods to the *crisprScore* package as they become available.

**Table 2.**
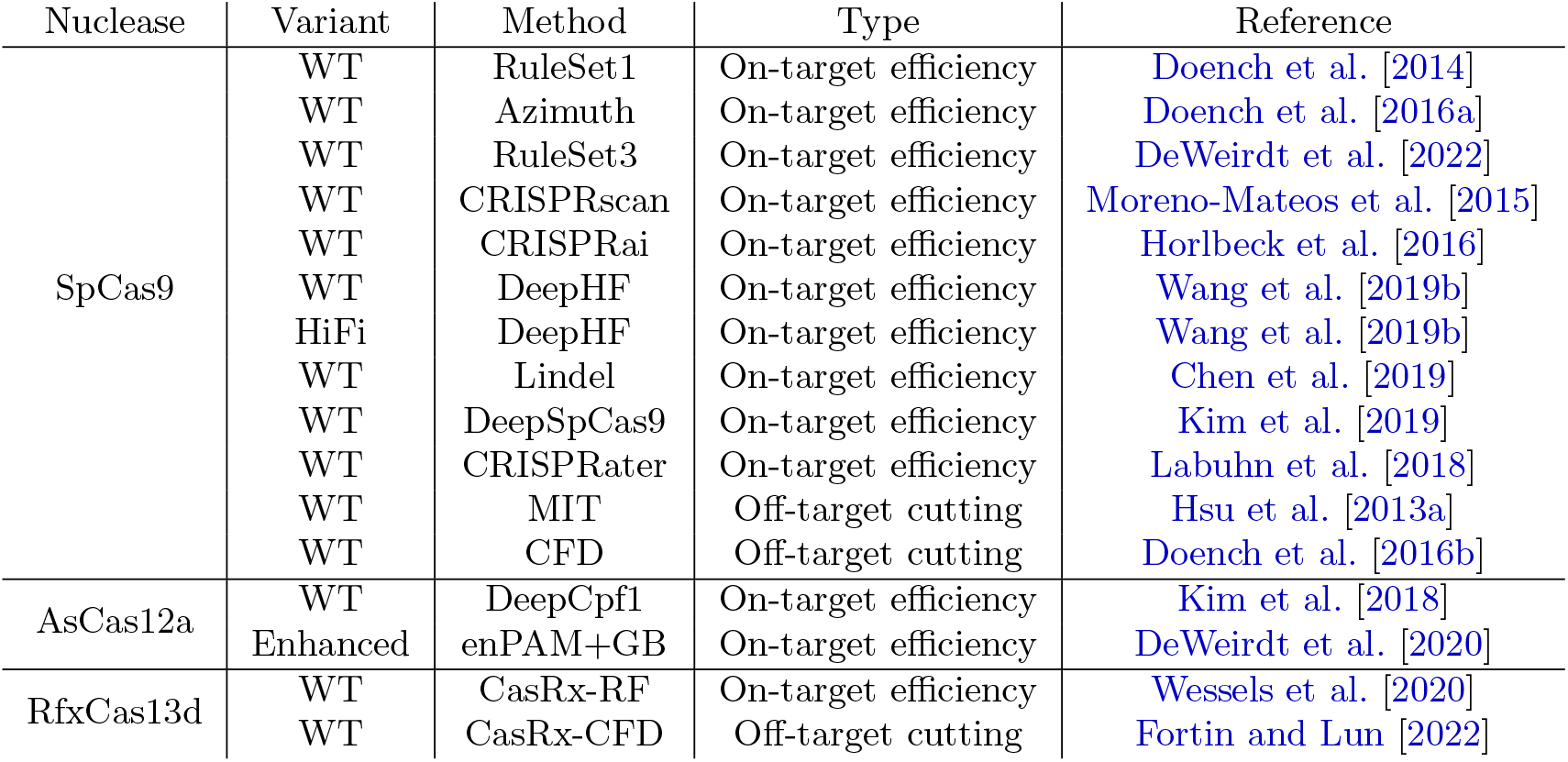
On-target and off-target scoring methods currently available in *crisprScore*.

### 2.5 *crisprDesign* enables fast characterization and annotation of off-targets

Off-targeting effects can occur when a spacer sequence maps with perfect or imperfect complementarity to a genomic locus other than the primary on-target. Given that nucleases can still bind and cut in the presence of nucleotide mismatches between spacer sequences and target DNA sequences [Fu et al., 2013, Hsu et al., 2013b, Pattanayak et al., 2013], it is paramount to obtain and characterize all putative mismatch-mediated off-targets.

The off-target functionalities in *crisprDesign* are divided into two parts: off-target search (alignment) and off-target characterization (genomic context and scoring). For the off-target search, we offer three different alignment methods: *Bowtie* [Langmead et al., 2009], the BWA-backtrack algorithm in *BWA* [Li and Durbin, 2009] and the Aho-Corasick exact string matching method implemented in *Biostrings* [Aho and Corasick, 1975, Pages et al., 2016]. We developed two independent R packages to implement the *Bowtie* and *BWA* alignment methods: *crisprBowtie* and *crisprBwa*. Notably, the packages were developed to work with any nucleases, and for both DNA and RNA target spaces (reference genomes and transcriptomes, respectively). While the maximum number of mismatches for *Bowtie* is limited to 3, there is no limit for *BWA*.

Given the short nature of gRNA spacer sequences, both *Bowtie* and *BWA* are ideal tools for off-target search and provide ultrafast results. On the other hand, the alignment method based on the Bioconductor package *Biostrings* does not need the creation of a genome index, and is particularly useful for off-target search in short custom sequences. All methods can be invoked via the *addSpacerAlignments* function, which returns the on- and off-target alignments as a *GRanges* object in the *GuideSet* metadata.

To add genomic context to the on- and off-targets, a *TxDb* object can be provided to the *addSpacerAlignments* function. The *TxDb* object is a standard Bioconductor object to store information about a gene model, and can easily be made from transcript annotations available as GFF3 or GTF files. Gene annotation columns are added to the off-target table for different contexts: 5’ UTRs, 3’ UTRs, CDS, exons, and introns. Finally, users can add the MIT and CDF off-target specificity scores [Hsu et al., 2013a, Doench et al., 2016a] implemented in *crisprScore* to characterize the likelihood of nuclease cleavage at the off-targets.

#### Comparison of the off-target alignment methods

We first compared the accuracy of the three alignment methods implemented in *crisprDesign*. Bhagwat et al. [2020] reported that *Bowtie* misses a large number of double-mismatch and triple-mismatch off-targets in comparison to the gold-standard complete string matching algorithm. To investigate this, we repeated the PAM-agnostic on- and off-target alignment of the 10 spacer sequences described in Bhagwat et al. [2020] to the GRCh38 reference genome using all three alignment methods. In contrast to Bhagwat et al. [2020], all three alignment methods implemented in *crisprDesign* return an identical list of off-targets (see Table S1). This indicates that, contrary to previous reports, both *BWA* and *Bowtie* provide a complete on- and off-target search. It appears that the missing off-targets in Bhagwat et al. [2020] are located on unlocalized and unplaced GRCh38 sequences.

Next, we evaluated the run times of four configurations offered in *crisprDesign* for alignment: *Bowtie, BWA*, an iterative version of *Bowtie* (bowtie-int) and an iterative version of *BWA* (bwa-int). We developed iterative versions of the *Bowtie* and *BWA* alignments to avoid situations where gRNAs are mapping to hundreds of loci in the genome, considerably slowing down the off-target search when a higher number of mismatches is allowed. The iterative strategy starts by aligning spacer sequences with no mismatches allowed. Then, it sequentially performs the alignment with a higher number of mismatches only for sequences that have a low number of off-targets at the previous step, thus avoiding the cost of extra searches for low-quality sequences that already have many off-targets. We performed the evaluation on three sets of gRNAs targeting the human genome, each with a different size (see Methods). For all three sets, the *Bowtie* and BWA gRNA alignments have comparable run times. (Figure S1). For *ZNF101*, which contains several non-specific gRNAs overlapping a repeat element, our iterative versions of the alignment methods shows substantial gain in speed.

Finally, we compared run times for designing SpCas9 gRNAs and performing a genome-wide off-target search for the following tools: *CCTop, CHOPCHOP, multicrispr, FlashFry*, and *crisprDesign*. Other tools were not included for reasons discussed in the Methods section. To perform the evaluation, we generated six random subsets of protein-coding exons located on chromosome 1 with the following sizes: 100, 200, 400, 800, 1600 and 3200 exons. For each tool and each subset, we ran the off-target alignment against the human reference genome (GRCh38 build) using a maximum of 2 mismatches. We included the alignment parameters used for each tool in the Methods section. Both *FlashFry* and the iterative *Bowtie* alignment implemented in *crisprDesign* show a substantial speed gain in comparison to other methods (Figure S2).

### 2.6 Accounting for human genetic variation by adding SNP annotation

Genetic variation, such as SNPs and small indels, can have a direct impact on gRNA binding productivity and on-target specificity by altering complementarity between spacer sequences and the target DNA or RNA [Scott and Zhang, 2017, Lessard et al., 2017, Canver et al., 2017, Wang et al., 2018]. In *crisprDesign*, users can apply the function *addSNPAnnotation* to annotate gRNAs for which the target protospacer sequence overlaps a SNP. This enables users to discard or flag undesirable gRNAs that are likely to have variable activity across different human genomes.

Given that the current human reference genome was built using only a small number of individuals, the allele represented in the human reference genome at a particular locus does not always correspond to the major allele in a population of interest. Inspired by the major-allele reference genome indices provided by the *Bowtie* team (see https://github.com/BenLangmead/bowtie-majref), we created two new human genomes to be used throughout our ecosystem that represent the major allele and the minor allele using dbSNP151 (see Methods).

Both genomes are available in Bioconductor as *BSgenome* packages. Both packages can be used in our ecosystem to improve gRNA design by designing gRNAs against either the minor or major allele genome, and searching for off-targets in both the major and minor allele genomes.

### 2.7 Comprehensive gene and functional annotations

The genomic context of the on-target sites is paramount for optimally selecting gRNAs in most, if not all, CRISPR applications. *crisprDesign* includes the *addGeneAnnotation* and *addTssAnnotation* functions, which report comprehensive transcript- and promoter-specific context for each gRNA target site, respectively. Users simply need to provide a standard Bioconductor *TxDb* object to specify which gene model should be used to annotate on- and off-targets.

For CRISPRko applications, *addGeneAnnotation* annotates which isoforms of a given gene are targeted. It also adds spatial information about the relative cut site within the coding sequence of each isoform, which has been shown to contribute to gRNA activity [Doench et al., 2016a]. Since translation reinitiation can result in residual protein expression [Smits et al., 2019], *addGeneAnnotation* reports whether or not the gRNA cut site precedes any downstream in-frame ATG sequences, following the rules of Cohen et al. [2019]. Additionally, to maximize gene knockout based on protein domains [He et al., 2019], we include Pfam domain annotation [Bateman et al., 2004] via the *biomarRt* package [Durinck et al., 2005]. For CRISPRa and CRISPRi applications, *addTssAnnotation* indicates which promoter regions are targeted by each gRNA, as well as the location of the target cut site relative to the TSS. This allows the user to easily select guides in the optimal targeting window.

To further put the gRNA targets into biological context, users can access thousands of genomic annotation datasets through the Bioconductor *AnnotationHub* resource. The resource includes common sources such as Ensembl, ENCODE, dbSNP and UCSC. Where appropriate, those annotations are in the *GenomicRanges* format, which make them directly compatible with the *GuideSet* object used to represent gRNAs in our ecosystem. By leveraging overlap operations on *GenomicRanges*, users can identify which gRNAs are present or absent in a given set of annotated features by using a few lines of code. For example, users can ask *AnnotationHub* whether a gRNA is targeting repeat elements to avoid cutting-induced toxicity, or whether a gRNA targets the region upstream of an annotated Cap Analysis of Gene Expression (CAGE) peak for CRISPRa applications. Additionally, the *rtracklayer* Bioconductor package [Lawrence et al., 2009] provides functionalities to easily read genome annotations that are stored in the commonly-used WIG, BED, bigWig and bedGraph formats. Utilizing *rtracklayer, crisprDesign* provides the function *addConservationScores* to annotate gRNAs with evolutionary conservation scores obtained from the UCSC genome browser (see Methods).

### 2.8 Advanced functionalities for designing screening libraries

Efficient cleavage can be disrupted by certain features of the gRNA sequence, such as very low or high percent GC content [Chen et al., 2018, Doench et al., 2014, Wang et al., 2014], homopolymers of four nucleotides or longer [Gilbert et al., 2014, Veeneman et al., 2020], and self-complementarity conducive to hairpin formation [Thyme et al., 2016, Labun et al., 2016]. When gRNAs are expressed from a U6 promoter, thymine homopolymers (TTTT) are particularly undesirable as they signal transcription termination. The *addSequenceFeatures* function flags all gRNAs that contain such undesirable sequence features. Another consideration in designing gRNA libraries is to exclude spacer sequences that are not compatible with the oligonucleotide cloning strategy. gRNAs that contain restriction sites of the enzymes used to clone the spacer sequences into a lentiviral vector should be excluded. The *addRestrictionEnzymes* function flags all gRNAs that contain restriction enzyme recognition motifs.

Optical pooled screening (OPS) is a promising novel screening modality that combines image-based *in situ* sequencing of gRNAs and optical phenotyping on the same physical wells [Feldman et al., 2019]. This enables linking genomic perturbations with high-content imaging at large scale. In such experiments, gRNA spacer sequences are partially sequenced. This translates to additional gRNA design constraints to ensure sufficient dissimilarity of the truncated spacer sequences. *crisprDesign* contains a suite of design functions that take into account OPS constraints, while ensuring that the final OPS library is enriched for gRNAs with best predicted activity.

To assist with the design of complex libraries, we developed the package *crisprViz* to visualize gRNAs. The package uses the Bioconductor package *Gviz* [Hahne and Ivanek, 2016] to offer a flexible and integrated visualization of gRNAs along genomic coordinates. Users can visually inspect gRNAs within a genomic track with the option of adding annotation tracks such as transcript models, SNP annotations, repeat elements, and nucleotide sequences.

### 2.9 Functional annotations in *crisprDesign* improve gRNA selection

We illustrate how functional annotations implemented in *crisprDesign* can improve gRNA selection by focusing on two functionalities: *addConservationScores* and *addGeneAnnotation*. We assessed the *addConservationScores* function using the large-scale CRISPRko fitness screening dataset from Project Achilles [Meyers et al., 2017]. We obtained normalized log fold changes (LFCs) measuring gRNA dropout over time (see Methods). In fitness screens, gRNAs targeting essential genes should deplete over time, and are therefore expected to have negative LFCs. Therefore, gRNAs targeting essential genes can be used to investigate determinants of gRNA activity. We downloaded basewise phyloP scores [Pollard et al., 2010] from the UCSC genome browser. Scores were calculated from a phylogenic alignment of 30 vertebrate species (see Methods). Positive and negative scores represent evolutionary conservation and acceleration, respectively. In Figure S3a, we show the correlation between LFCs and conservation scores obtained using different window sizes, for gRNAs targeting a reference set of essential genes [Hart et al., 2014]. The data suggest an optimal window of 18 nucleotides around the cut site, which is our recommended window size in *crisprDesign*. In Figure S3b, we present LFC distributions of gRNAs targeting essential genes, split by the sign of the gRNA conservation score. gRNAs targeting conserved regions (positive score, red line) show greater activity than less conserved regions (negative score, black line) as observed by greater gRNA dropout. This is in line with previous results [Schoonenberg et al., 2018, DeWeirdt et al., 2022, Veeneman et al., 2020]. gRNAs targeting non-essential genes serve as negative controls and show no dropout, irrespective of the conservation scores, as expected (Figure S3c).

Next, we sought to understand how gRNA position within the CDS of the target gene impacts gRNA activity. Given that most gRNAs in Project Achilles were located in the first 50% of the target CDS by design, we obtained a different screening dataset; we downloaded data from a genome-wide fitness screen performed in HCT116 cells (Hart2015 dataset, see Methods). We used the *addGeneAnnotation* function in *crisprDesign* to annotate gRNAs with a position percentage of the target CDS. We used the Ensembl canonical transcript of the target gene as the representative CDS. In Figure S3d, we show the relationship between LFCs and % CDS for gRNAs targeting essential genes. gRNAs located beyond the first 85% of the CDS (to the right of the vertical line) show a progressive decline in activity. The results agree with the litterature [Doench et al., 2016a]. gRNAs targeting non-essential genes serve as negative controls and behave as expected (Figure S3e).

Based on these results, both functional annotations help selecting more active gRNAs; we recommend in *crisprDesign* to prioritize gRNAs with positive conservation scores, and located in the first 85% of the target CDS. Those recommended parameters are implemented as the default parameters in the *crisprDesign* gRNA ranking procedure discussed next.

## 3 Case studies

### 3.1 gRNA ranking from crisprDesign returns optimized gRNAs

To complement gRNA annotation and assist in library design, *crisprDesign* provides a gRNA ranking function called *rankSpacers*. The function implements our recommended ranking parameters for the nucleases SpCas9, enCas12a, and CasRx, effectively enabling library design automation across targets. It is designed to optimize both on-target activity and minimize off-targeting effects, and includes the functional annotations described in the previous section. Details are provided in the Methods section.

We compared our default gRNA ranking procedure to other tools listed in Table 1 that provide gRNA rankings: *CHOPCHOP, CCTop, FlashFry* and *CRISPick*. To perform the evaluation, we designed and ranked SpCas9 gRNAs for all human protein-coding genes (Ensembl release 104) using each tool separately (see Methods). Next, we obtained and processed 5 human genome-wide fitness screen datasets from published studies (Table 3), each performed using a different gRNA library. For each dataset and gRNA, a LFC between later and earlier time point samples was calculated to quantify gRNA dropout over time.

**Table 3.** Genome-wide human CRISPRko screen datasets used for comparing SpCas9 gRNA rankings.

| Dataset | gRNA library | Cell Line | Number of gRNAs | Reference |
| --- | --- | --- | --- | --- |
| Achilles | Avana | (many) | 67,816 | Meyers et al. [2017] |
| Hart2015 | TKOv1 | HCT116 | 164,576 | Hart et al. [2015] |
| Hart2017 | TKOv3 | HAP1 | 81,967 | Hart et al. [2017b] |
| Wang2015 | Sabatini | K562 | 166, 855 | Wang et al. [2015] |
| Tzelepis2016 | Yusa | HL60 | 85,192 | Tzelepis et al. [2016] |

gRNAs targeting essential genes are expected to drop out and can be used for benchmarking purposes. To investigate the relationship between gRNA activity and gRNA ranking, we considered for each gRNA library the subset of gRNAs targeting a common reference set of essential genes [Hart et al., 2014]. For each gene and tool, we identified the top 15 ranked gRNAs based on the tool-specific in silico ranking. In Figure 3a, we show the distributions of LFCs in the Hart2015 dataset based on two groups: red lines show the distributions of the top 15 ranked gRNAs across genes, and green lines show the distributions of remaining gRNAs. Top ranked gRNAs from *CRISPick* and *crisprDesign* show greater activity than lower ranked gRNAs, as indicated by a negative shift in the red distributions with respect to the green distributions.

**Figure 3.**
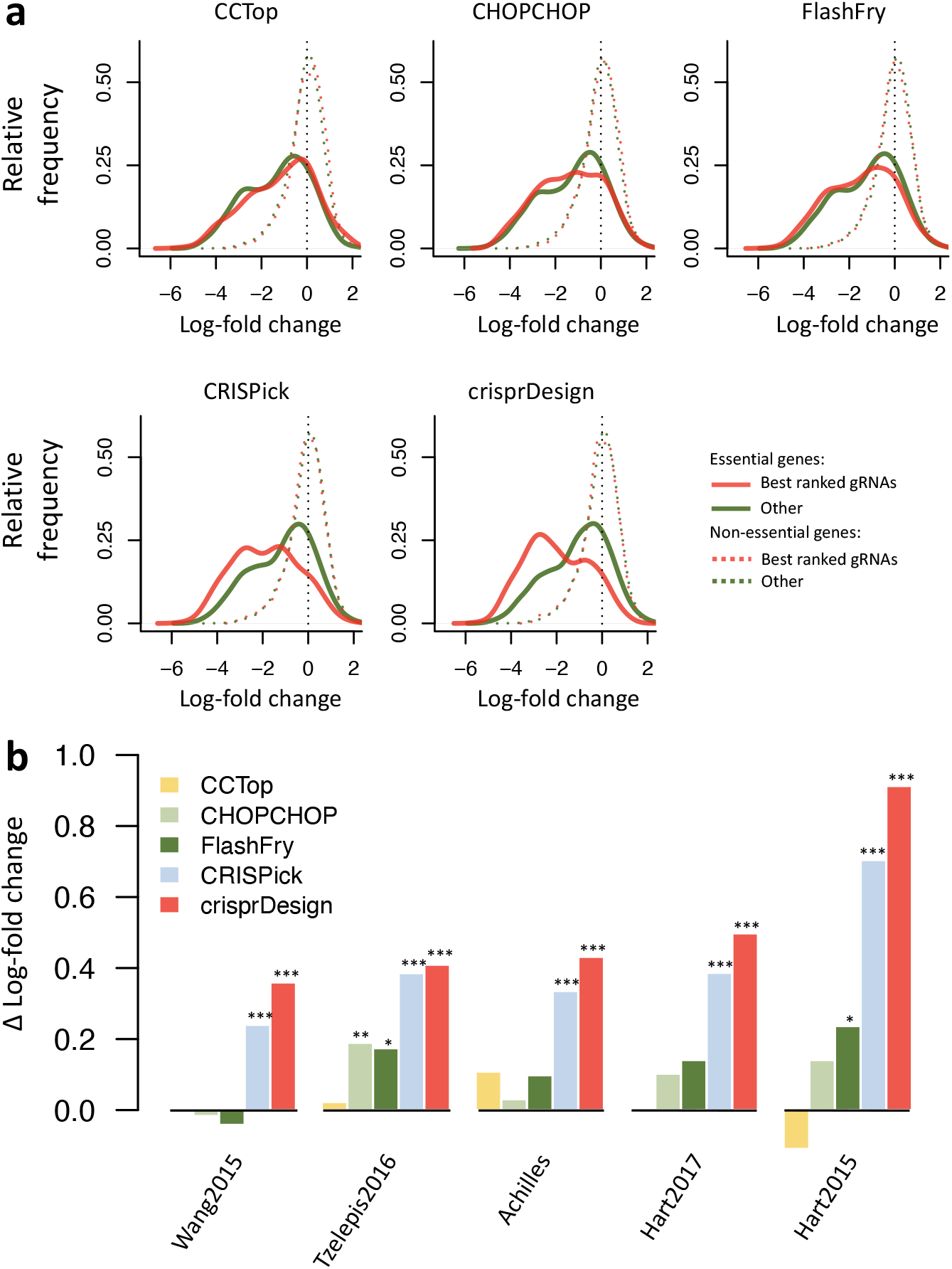
Comparison of CRISPRko Cas9 gRNA rankings for protein-coding human genes. We designed and ranked gRNAs targeting all protein-coding human genes (Ensembl release 104) using tools that provide gRNA rankings: *CCTop, CHOPCHOP, FlashFry, CRISPick* and *crisprDesign*. To compare gRNA ranking performance across tools, we obtained gRNA LFCs from 5 genome-wide CRISPRko fitness screening datasets, listed in Table 3. In these fitness screens, active gRNAs targeting essential genes are expected to drop out and show negative LFCs. To investigate the relationship between gRNA activity and gRNA ranking, we considered for each gRNA library the subset of gRNAs targeting a common reference set of essential genes [Hart et al., 2014]. For each gene and tool, we identified the top 15 ranked gRNAs based on the tool-specific in silico ranking. **a** LFC distributions in the Hart2015 dataset for gRNAs targeting essential genes (solid lines) and gRNAs targeting non-essential genes (dotted lines). Red lines show the distributions of the top 15 ranked gRNAs across genes, and green lines show the distributions of remaining gRNAs. For essential genes, top ranked gRNAs from *CRISPick* and *crisprDesign* show greater activity than lower ranked gRNAs (red distributions are negatively skewed). As expected, there are no differences for gRNAs targeting non-essential genes. **b** We repeated the analysis described in **a** for each dataset. We summarized the performance of the top ranked gRNAs by calculating the difference in means between the green and red distributions (Δ LFC), for essential genes only. A higher Δ LFC indicates better performance. For each method and dataset, a t-test was performed to quantify the difference in LFCs between the top ranked gRNAs and the remaining gRNAs. Corresponding p-values are reported above the bars (*\**: p-value *<* 0.05; *\*\**: p-value *<* 0.01; *\* * \**: p-value *<* 0.001.)

We repeated the analysis for each dataset, and summarized the performance of the top ranked gRNAs at discriminating active gRNAs by calculating the difference in means between the green and red distributions (Δ LFC). Results are shown in Figure 3b. Higher Δ LFCs indicate better performance, and results indicate that both *CRISPick* and *crisprDesign* perform well across all datasets.

### 3.2 Designing gRNAs targeting *BRCA1* for the base editor BE4max

CRISPR base editors are deaminases fused to CRISPR nickases to introduce mutations at loci targeted by the gRNAs without introducing double-stranded breaks (DSBs) [Gaudelli et al., 2017, Komor et al., 2016]. A recent study showed high heterogeneity and complexity of the editing outcomes across eight popular base editors, motivating the need of robust but flexible software to design gRNAs for base editing applications [Arbab et al., 2020]. In particular, this includes functionalities for listing and characterizing potential edited alleles introduced by gRNAs to inform the phenotypic readouts created by those gRNAs.

To illustrate the functionalities of our ecosystem for designing base editor gRNAs, we designed and characterized all possible gRNAs targeting the coding sequence of *BRCA1* for the cytidine base editor BE4max [Koblan et al., 2018]. The design workflow is shown in Figure 4.

**Figure 4.**
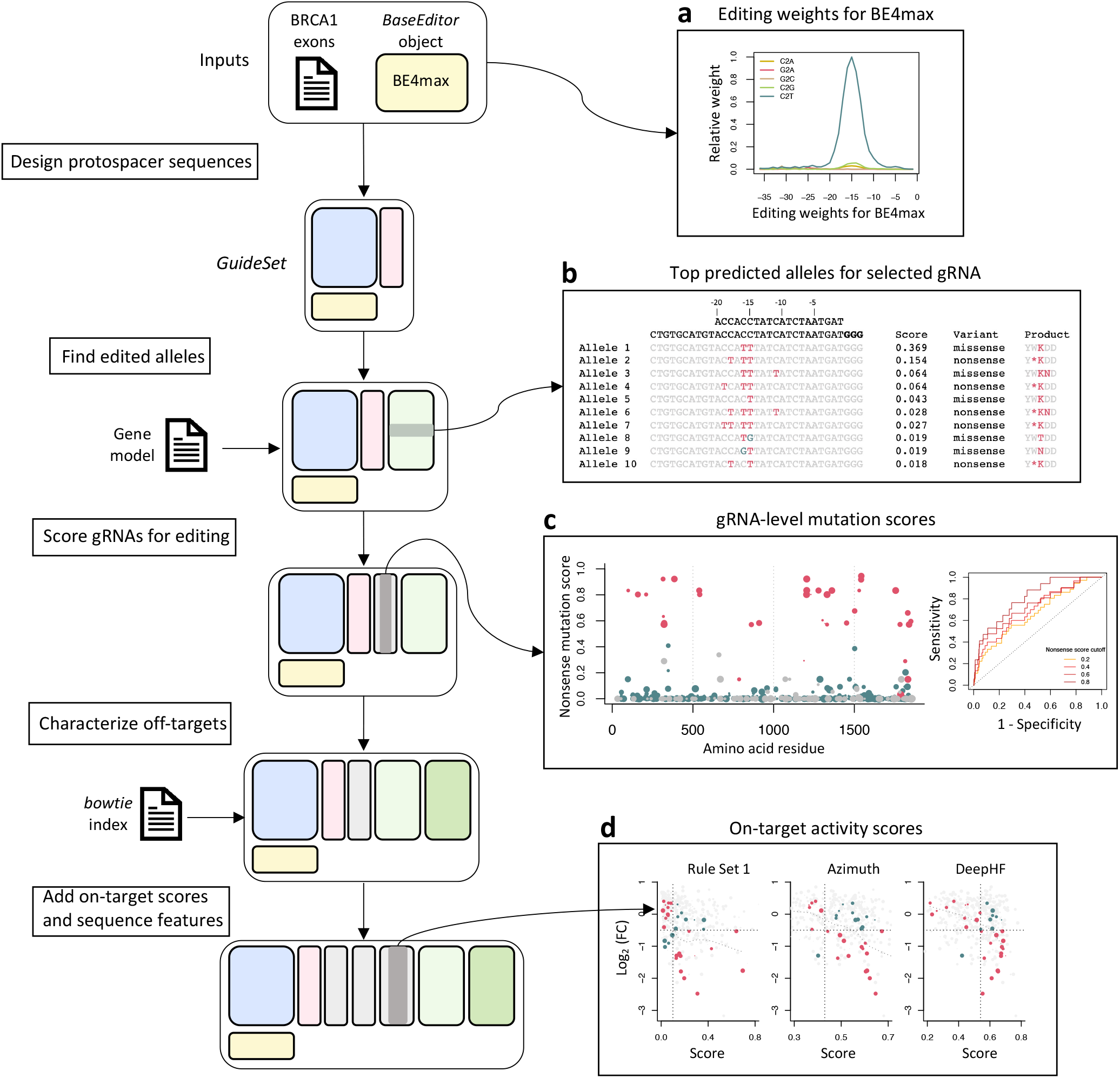
*crisprDesign* workflow to design gRNAs tiling *BRCA1* using the base editor BE4max. On the left: schematic showing the major steps involved in designing BE4max gRNAs targeting *BRCA1*. Two inputs are required: DNA sequences of *BRCA1* exons and a *BaseEditor* object from *crisprBase*. **a** Editing weights for the BE4max base editor from *crisprBase*. **b** 10 top predicted edited alleles for one selected gRNA as returned by *crisprDesign*. The wildtype allele and the protospacer sequence are positioned at the top of the first column, with the PAM sequence highlighted in bold. Edited nucleotides are highlighted in red (C to T) and blue (C to G). Editing scores, variant annotations, and protein product of the edited alleles are also shown. **c** On the left, gRNA-level nonsense mutation score as calculated by *crisprDesign*. Colors represent variant classification: nonsense in red, missense in blue, silent in grey. The size of the dot is proportional to the on-target efficiency *DeepHF* score. On the right, ROC curves for classifying gRNA mutation type (nonsense or not) based on gRNA dropout from the *BRCA1* BE4max dataset (see Methods). Different thresholds of the nonsense score were used to label a gRNA as nonsense or not. **d** Relationships between gRNA dropout from the *BRCA1* BE4max dataset and several on-target activity scores. gRNAs that are not predicted to induce a nonsense mutation are colored in grey, and the size of the dots is proportional to the magnitude of the mutation score. The horizontal dotted lines at -0.5 represent a cutoff to classify a gRNA as active or not. For each method, a score cutoff was determined to classify active versus non-active gRNAs (vertical dotted line). Red and blue dots correspond to gRNAs that are correctly and incorrectly classified, respectively.

The first step consisted of designing all possible guides targeting *BRCA1* using the *findSpacers* function in *crisprDesign*. The BE4max *BaseEditor* object from *crisprBase* was used to store nucleotide- and position-specific editing probabilities (see Figure 4a), which inform the editing window of interest for each of the gRNA targets. Next, using the function *getEditedAlleles*, we generated and stored all possible editing events at each gRNA (see Figure 4b). The function also adds a score for each edited allele that quantifies the likelihood of editing to occur based on the editing probabilities stored in the *BaseEditor* object (see Methods). In addition, each edited allele is annotated for its predicted functional consequence: silent, missense, or nonsense mutation.

In case several mutations occur in a given edited allele, the most consequential mutation is used to label the allele (nonsense over missense, and missense over silent). For each gRNA, and for each mutation type, we then generated a gRNA-level score by aggregating the likelihood scores across all possible alleles (see Methods). The score represents the likelihood of a gRNA to induce a given mutation type (see Figure 4c, left plot).

To show how our gRNA annotations can be used to understand the phenotypic effects observed in screening data, we obtained data from a negative selection pooled screen performed in MelJuSo using a base editing library tiling the *BRCA1* gene [Hanna et al., 2021]. Given that loss-of-function mutations in *BRCA1* reduce cell fitness [Findlay et al., 2018], gRNAs introducing nonsense mutations are expected to drop out. We created Receiver

Operating Characteristic (ROC) curves to measure how well gRNA dropout can separate positive controls from other gRNAs. We used LFCs in gRNA abundance between the later time point and the plasmid DNA (pDNA) library as a measure of gRNA dropout (see Methods). We used several thresholds of the nonsense mutation score to label gRNAs as positive controls or not. We observed that gRNA dropout in the screen can separate positive controls well from all other gRNAs, and that performance is improved when using positive controls defined by higher nonsense mutation scores (Figure 4c).

We also characterized gRNAs for off-targeting effects using *crisprBowtie*, added sequence features using *crisprDesign*, and added on-target scores using *crisprScore*. We asked whether or not the magnitude of gRNA dropout in the screen associates with predicted on-target activity for the SpCas9 nuclease. In Figure 4d, we show gRNA dropout as a function of different predicted gRNA efficacy scores: Rule Set 1, Azimuth, and DeepHF. gRNAs predicted to induce nonsense mutations are shown in red, and grey otherwise. Despite the fact that each algorithm was trained on data using a SpCas9 nuclease with intact endonuclease activity, gRNA dropout and predicted gRNA efficacy correlate for all methods (*r* = *−*0.30 for Rule Set 1, *r* = *−*0.20 for Azimuth, and *r* = *−*0.17 for DeepHF). Overall, the different functionalities implemented in our ecosystem provides a set of informative annotations for base editor gRNAs and facilitate the interpretation of experimental data obtained from base editor screens.

### 3.3 Annotating and scoring gRNAs for gene knockdown using CasRx

One of the challenges in designing gRNAs specific to RNA-targeting nucleases is to enable on-target and off-target characterization to be performed in a transcriptome space, as opposed to a reference genome. This requires strand-specific functionalities, transcriptome-specific alignment indexes, as well as additional gene annotation functionalities to capture isoform-specific targeting.

Here, we describe a workflow for designing gRNAs targeting *CD46* and *CD55* using the RNA-targeting nuclease CasRx (RfxCas13d) [Konermann et al., 2018] (Figure 5). The workflow takes into consideration the aforementioned issues. To validate our design process, we obtained CasRx pooled screening data performed in HEK 293 cells with gRNA libraries tiling the human genes *CD46* and *CD55* from Wessels et al. [2020]. Since both genes encode for cell-surface proteins, the authors used fluorescence-activated cell sorting (FACS) to sort cells with high and low expression. Their data can be used to investigate gRNA knockdown efficacy based on the change in relative abundance of high- and low-expressing cells for each targeted gene (see Methods).

**Figure 5.**
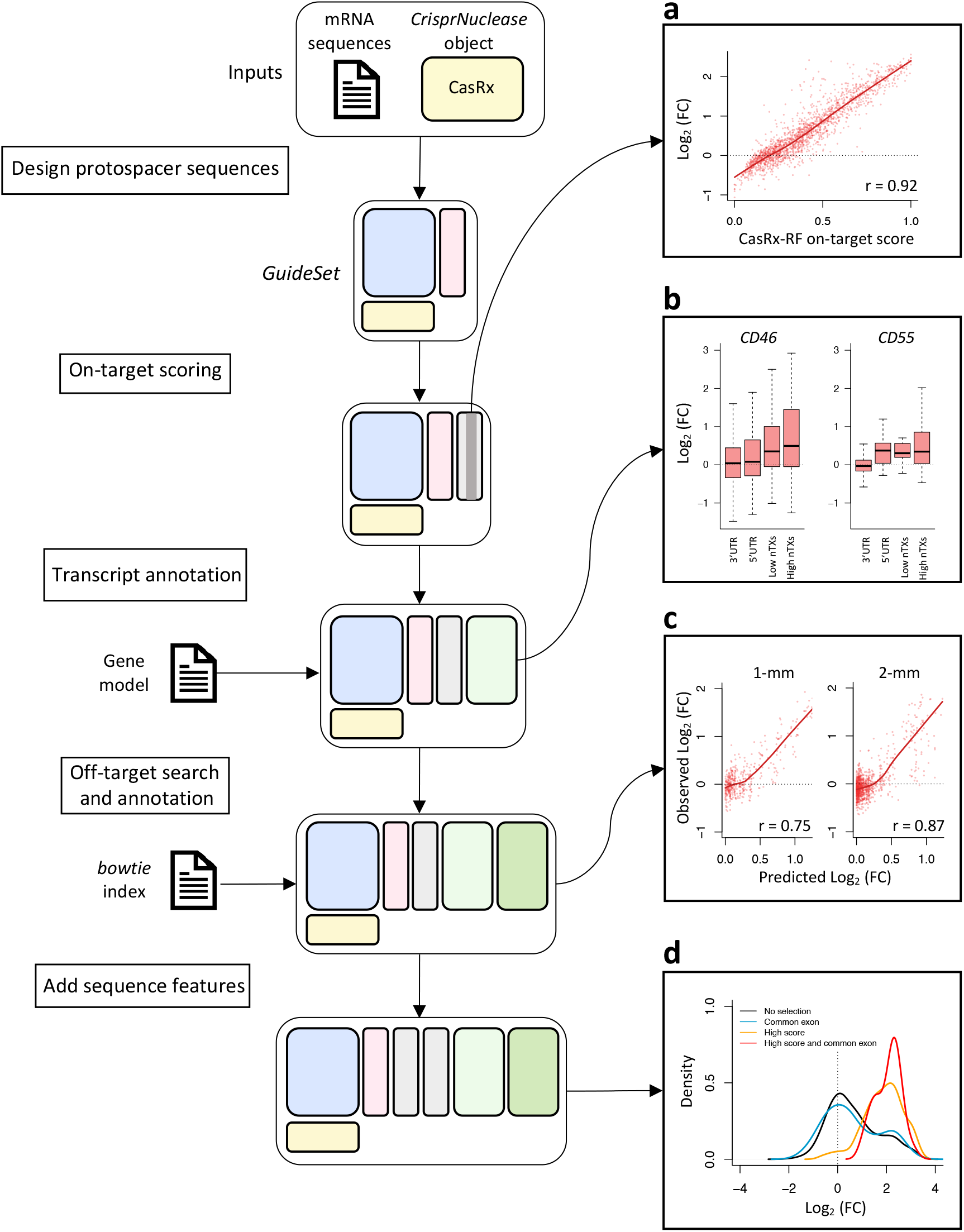
*crisprDesign* workflow to design gRNAs tiling *CD55* and *CD46* using CasRx. On the left: schematic showing the major steps involved in designing CasRx gRNAs targeting *CD55* and *CD46*. Two inputs are required: mRNA sequences of *CD55* and *CD46* and a *CrisprNuclease* object from *crisprBase*. **a** Relationship between on-target CasRx-RF score calculated in *crisprScore* and LFCs from the pooled FACS tiling CasRx screening data (see Methods). A higher LFC indicates higher gRNA activity. **b** Relationship between LFCs from the CasRx screening data and gRNA context for *CD46* and *CD55* : gRNAs targeting 5′ UTR and 3′ UTR for the canonical transcript, and guides targeting a low and high number of isoforms for each of the genes. gRNAs targeting more isoforms show higher enrichment in the screening data. The full isoform annotation is stored in the *GuideSet* objects. **c** Left: relationship between observed LFCs of on-target gRNAs in the *CD55* screen and predicted LFCs of single-mismatch gRNAs using the off-target CFD-CasRx score implemented in *crisprScore* (see Methods). Right: same as left, but for double-mismatch gRNAs. **D** gRNAs selected in the *CD46* screen for high on-target activity (CasRx-RF score) and targeting a common exon across all protein-coding isoforms enrich for high gRNA activity.

We first extracted mRNA sequences of both genes using the function *getMrnaSequence* from *crisprDesign*. The mRNA sequences, together with the *CrisprNuclease* object CasRx from *crisprBase*, served as inputs to create a *Guideset*. Next, we predicted on-target activity of the gRNAs using our implementation of the CasRx-RF method [Wessels et al., 2020] available in *crisprScore* (see Methods). The normalized LFCs in the screen correlate well with the CasRx-RF score (Figure 5a). We then added a transcript annotation to each gRNA using an Ensembl *TxDb* object as input. This adds a list of targeted isoforms to each gRNA, as well as transcript context (CDS, 5′UTR, or 3′UTR). We observed in the screen that gRNAs targeting a higher number of isoforms, and gRNAs located in CDS, lead to higher activity (Figure 5b, and Figure S4).

We performed an off-target search using *crisprBowtie* to the human transcriptome by providing a *Bowtie* index built on mRNA sequences. We extended the CFD off-target scoring algorithm implemented in *crisprScore* to work with CasRx by estimating mismatch tolerance weights on the GFP tiling screen data from Wessels et al. [2020] (see Methods). The off-target CFD-CasRx score performs well at predicting gRNA activity of single-mismatch and double-mismatch gRNAs in the *CD55* screen (Figure 5c, and see Methods).

Finally, we added sequence features, and ranked gRNAs for targeting *CD55* and *CD46* based on (1) high on-target score, (2) low number of off-targets, and (3) high number of targeted isoforms. If we select gRNAs that target a common exon and that have high on-target score, we enrich for highly active gRNAs in the screening data (Figure 5d).

### 3.4 Designing optimal gRNAs to activate *MMP7* using CRISPRa using different nucleases

Designing gRNAs for either CRISPRa and CRISPRi applications requires additional considerations. This includes choosing an optimal target region based on chromatin accessibility data and TSS data, and selecting gRNAs based on their positioning with respect to the TSS.

To demonstrate the utility of our ecosystem functionalities for CRISPRa and CRISPRi, we designed gRNAs for CRISPRa using the human gene *MMP7* as an example target (Figure 6). CRISPRi is discussed at the end of this section. One CRISPRa-specific design consideration is the limited number of candidate gRNAs available for a given gene due to the narrow window of optimal activation. Engineered nucleases with less constrained PAM sequences can improve CRISPRa applicability by expanding the set of candidate gRNAs. To investigate this, we designed gRNAs for the promoter region of *MMP7* using four nucleases in parallel: SpCas9, AsCas12a, and the more PAM-flexible versions SpGCas9 [Walton et al., 2020] and enAsCas12a [DeWeirdt et al., 2020].

**Figure 6.**
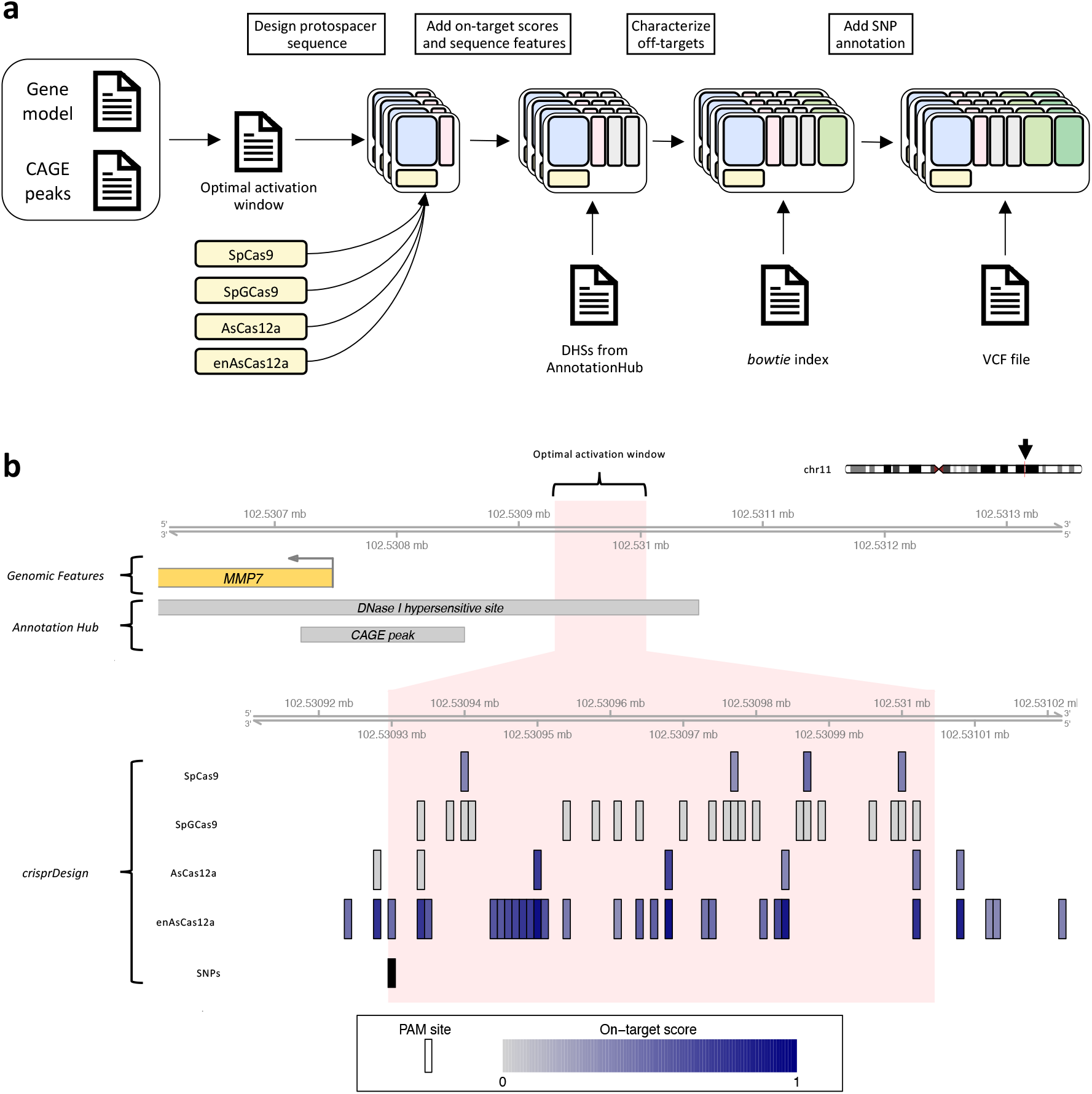
Design of CRISPRa gRNAs for human gene *MMP7* for different CRISPR nucleases. **a** Schematic showing the steps involved in designing CRISPRa gRNAs targeting the promoter region of *MMP7*. A gene model and a list of CAGE peaks are used to define the optimal window for gene activation. A *GuideSet* is created separately for each CRISPR nuclease. DNase I hypersensitive site (DHS) information is obtained from *AnnotationHub* and added to the gRNA annotation. **b** The top track shows the promoter region of human gene *MMP7* on chromosome 11, including part of the 5′ UTR of *MMP7* (yellow). The DHS and CAGE peak grey boxes were obtained using *AnnotationHub* (see Methods). The light pink region corresponds to the optimal region of activation based on Sanson et al. [2018], corresponding to a region [75,150]bp upstream of the 5′ end of the CAGE peak. For each of the four selected nucleases, all canonical PAM sites located within the optimal region are shown. PAM sites are colored by their on-target score: DeepHF for SpCas9, DeepCpf1 for AsCas12a, and enPAM+GB for enAsCas12a. No on-target scoring algorithm was available at time of publication for SpGCas9. The last track corresponds to common SNPs obtained from dbSNP151.

The first step of the gRNA design was to specify the target region for *MMP7*. We used *AnnotationHub* to find CAGE peaks in the promoter region of *MMP7* to specify the TSS position. We used the CAGE data to identify TSSs instead of RefSeq or Ensembl as the former provides more accurate annotations for designing CRISPRi and CRISPRa gRNAs [Radzisheuskaya et al., 2016]. The 5′ end of the CAGE peak was used as the TSS to define the coordinates of the optimal window of activation (75 and 150 nucleotides upstream of the TSS, as recommended by Sanson et al. [2018]).

Next, we designed all possible gRNAs for the four nucleases using the *findSpacers* function in *crisprDesign*, and stored the gRNAs in four separate *GuideSet* containers. We annotated each *GuideSet* for overlap with DNase I hypersensitivity sites (DHS) from consolidated epigenomes from the Roadmap Epigenomics Project [Kundaje et al., 2015] using *AnnotationHub*. Open-chromatin regions are favorable for the binding of the catalytically inactive Cas9 (dCas9) used in both CRISPRa and CRISPRi [Kuscu et al., 2014, Wu et al., 2014]. We then added sequence features using *crisprDesign*, on-target scores using *crisprScore*, and off-target sites using *crisprBowtie* for each nuclease. Finally, we added overlapping SNPs information using the *addSNPAnnotation* function and using dbSNP151. The end-to-end workflow is presented in Figure 6a.

The designed gRNAs are presented in Figure 6b. With *crisprDesign*, it is straightforward to select candidate gRNAs in the most promising genomic regions - in this case, lying inside both the annotated DHS and the optimal activation window for MMP7. One can immediately appreciate that both nuclease variants (SpGCas9 and enAsCas12a) yield substantially more available gRNAs in the optimal window activation. In particular, enAsCas12a offers several gRNAs with high predicted on-target activity, making it a better candidate for gene activation of *MMP7*. One SNP was also found in the region of interest, and overlapping one gRNA for enAsCas12a that should be avoided. Altogether, our ecosystem provides an easy and comprehensive workflow to enable users to design optimal gRNAs for CRISPRa across nucleases.

Designing gRNAs for CRISPRi applications using *crisprDesign* is nearly identical, with the exception that the preferred target region for interference is located downstream of the TSS. The CRISPRai scoring algorithm from Horlbeck et al. [2016], available through *crisprScore*, can be used to select optimal gRNAs for each TSS separately, taking into account both gRNA positioning and sequence content to maximize on-target inhibition.

For both CRISPRa and CRISPRi, our gRNA design workflow is also applicable to non-coding regulatory elements, for instance long non-coding RNAs (lncRNAs) as it was done in Liu et al. [2017]. Overall, *crisprDesign* provides end-to-end functionalities that are well-suited for a large array CRISPRa and CRISPRi applications.

## 4 Discussion

In this work, we introduced a new suite of R packages to perform comprehensive end-to-end gRNA design for a multitude of CRISPR technologies and applications. Our ecosystem, named the *crisprVerse*, enables users to perform gRNA design for diverse nucleases such as PAM-free nucleases and RNA-targeting nucleases, and for several applications beyond CRISPRko such as RNA and DNA base editing and CRISPRa/i. All design functionalities are available from a core package, *crisprDesign*. This eliminates the need to use multiple tools to obtain the necessary information for selecting optimal gRNAs, which is both time consuming and error prone.

We demonstrated the diversity of our framework by applying it in three case studies involving different CRISPR technologies with their own specific design considerations. We were able to show that creating rich gRNA annotations can help investigate gRNA variability and biases observed in experimental data generated from newer CRISPR technologies. To do so, we obtained public pooled screening data from two published studies, a tiling base editor screen of *BRCA1*, and a tiling CasRx screen of *CD46* and *CD55*, and show how some of the gRNA features derived from *crisprDesign* can explain some of the variability in gRNA activity observed in both screens. We also showed that our default gRNA ranking criteria implemented in *crisprDesign* yield optimal gRNAs by reanalyzing five genome-wide fitness screening datasets.

The modular architecture of the *crisprVerse* enables nucleases, base editors, scoring methods and annotations to be combined depending on the needs of the user. As a result, our design framework can easily adapt to new CRISPR technologies by swapping out the necessary components. For instance, a recent study has shown that the resolution of base editor screens can be greatly increased by combining existing base editors with PAM-extended Cas9 variants [Sangree et al., 2021], while another study shows that RNA-targeting Cas13 nucleases can be combined with deaminases to form RNA base editors [Cox et al., 2017]. Both applications can be readily supported by our ecosystem without the need for further development.

Our ecosystem is completely implemented within the Bioconductor project, which provides robust and feature-rich data structures, high-quality documentation and workflows, and seamless interoperability between packages. Data structures defined in *crisprBase* can be reused to facilitate the analysis of CRISPR-based editing events in other packages, such as *ampliCan* [Labun et al., 2019], *GUIDEseq* [Zhu et al., 2017] and *CrisprRVariants* [Lindsay et al., 2016]. *GuideSet* gRNA containers can be integrated with packages that provide analysis workflows for pooled screening data [Wang et al., 2019a, Imkeller et al., 2020, Bainer et al., 2021] to investigate biases and filter out undesirable gRNAs. Finally, the *crisprBowtie* and *crisprBwa* packages provide general functions that can be used to map any short sequences, including small-hairpin RNAs and short-interfering RNAs. We are continuously extending our suite of tools to make available the latest developments for gRNA design, such as prime editing [Anzalone et al., 2019] and combinatorial libraries [Replogle et al., 2020]

## 5 Methods

### Reference genomes, gene models, and genome indexes

The FASTA file for the human reference genome (GRCh38.p13 assembly) was obtained from UCSC to build *Bowtie* and *BWA* indexes via the *Rbowtie* (v1.37) [Hahne et al., 2012] and *Rbwa* (v1.1) R packages, respectively. The packages use Bowtie v1.3 and BWA Release 0.7.17, respectively. The gene model used throughout the manuscript was obtained from Ensembl (release 104) using the R package *GenomicFeatures* (v1.49.6). Common SNPs were obtained from NCBI dbSNP build 151 (https://ftp.ncbi.nlm.nih.gov/snp/).

### CAGE peak and DNAse I hypersensitivity data

RIKEN/ENCODE CAGE peaks were obtained from *AnnotationHub* (v3.5) using accession number AH5084 [Djebali et al., 2012]. Genomic coordinates were lifted over from hg19 to hg38 using the R package *rtracklayer* (v1.57). DNAse I hypersensitive sites were obtained from *AnnotationHub* using accession number AH30743. The narrow DNase peaks were obtained using MACS2 on consolidated epigenomes from the Roadmap Epigenomics Project (`E116-DNase.macs2.narrowPeak.gz`) [Kundaje et al., 2015]. Genomic coordinates were lifted over from hg19 to hg38 using the R package *rtracklayer*.

### On-target scoring

We implemented several commonly-used algorithms for Cas9, Cas12 and Cas13 nucleases in *crisprScore*. For predicting on-target activity of the wildtype SpCas9 nuclease, we implemented the popular Rule Set 1 [Doench et al., 2014] and *Azimuth* algorithms [Doench et al., 2016a] (iteration of the popular Rule Set 2 algorithm by the same authors), and the sequence-only Rule Set 3 [DeWeirdt et al., 2022]. The package also provides the deep learning-based algorithms *DeepWT* and *DeepHF*, developed to predict cutting efficiency of the wildtype SpCas9 and SpCas9-High Fidelity (SpCas9-HF1) nucleases, respectively [Wang et al., 2019b], and the *DeepSp-Cas9* algorithm [Kim et al., 2019]. We also included the *CRISPRscan* algorithm [Moreno-Mateos et al., 2015] for predicting on-target activity of SpCas9 gRNAs expressed from a T7 promoter, as well as the *CRISPRater* algorithm [Labuhn et al., 2018]. For the wildtype AsCas12a, *crisprScore* offers the deep-learning based prediction method DeepCpf1 [Kim et al., 2018]. For the enhanced AsCas12a (enAsCas12a), *crisprScore* offers the *enPAM+GB* algorithm [DeWeirdt et al., 2020]. For CasRx (RfxCas13d), we adapted the code from the random forest model developed in Wessels et al. [2020]; we referred to the method as *CasRx-RF*.

For predicting gRNA activity for CRISPRa and CRISPRi, we implemented the prediction method used to design the commonly-used Weissman CRISPRa and CRISPRi v2 genome-wide libraries for human and mouse [Horlbeck et al., 2016]. This method predicts CRISPRa (or CRISPRi) gRNA activity based on the distance to the transcription starting site (TSS), spacer sequence-derived features, as well as chromatin accessibility data and nucleosome positioning using DNase-Seq, MNase-Seq, and FAIRE-Seq data. The chromatin data in hg38 coordinates are available on Zenodo (DOI: 10.5281/zenodo.6716721).

The function *addCompositeScores* from *crisprDesign* creates an aggregate score from a specified list of on-target scoring methods. It takes the average of the specified scores after performing a rank transformation. More specifically, consider *s*_*ij*_ to be the score value for gRNA *i* and method *j*. The composite score *S*_*i*_ for gRNA *i* is

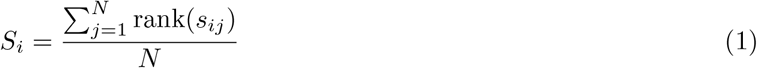

where *N* is the total number of user-specified on-target scoring methods, and rank(*s*_*ij*_) is the ranked score within method *j*. Importantly, if the number of missing values varies across on-target scoring methods, we ensure that the scale of the rank-transformed values are comparable across methods by simply scaling the ranks so that highest ranked value is equal across all methods. Missing values are uncommon but can happen when designing gRNAs targeting custom sequences. Indeed, several scoring algorithms require nucleotide context around the protospacer sequences, and this is not possible for gRNAs located near the end of the user-provided custom sequences.

#### On-target prediction of frameshift-causing indels using *Lindel*

In *crisprScore*, we implemented *Lindel* [Chen et al., 2019], a logistic regression model that was trained to use local sequence context to predict the distribution of mutational outcomes for CRISPR/Cas9. The *Lindel* final score reported in *crisprScore* is the proportion of “frameshifting” indels, that is the frequency of indels predicted to introduce frameshift mutations. By chance, assuming a random distribution of indel lengths, gRNAs should have a frameshifting proportion of 0.66. A *Lindel* score higher than 0.66 indicates that a given gRNA is more likely to cause a frameshift mutation than by chance.

### Off-target scoring of individual off-targets

The exact formula that we use to calculate the CFD score in *crisprScore* is

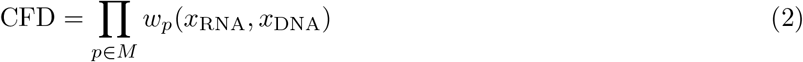

where *M* is the set of positions for which there is a mismatch between the gRNA spacer sequence and the off-target sequence. *w*_*p*_(*x*_RNA_, *x*_DNA_) is an experimentally-derived mismatch tolerance weight at position *p* depending on the RNA nucleotide *x*_RNA_ and the DNA nucleotide *x*_DNA_ (see Doench et al. [2016b] for more details).

The exact formula that we use to calculate the MIT score in *crisprScore* was obtained from the MIT design website (crispr.mit.edu):

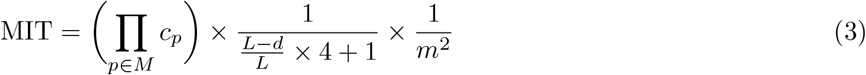

where *M* is the set of positions for which there is a mismatch between the gRNA spacer sequence and the off-target sequence, *c*_*p*_ is an experimentally-derived mismatch tolerance weight at position *p, d* is the average distance between mismatches, *m* is the total number of mismatches, and *L* is the spacer length. The spacer length used in Hsu et al. [2013a] is 19. As the number of mismatches increases, the cutting likelihood decreases.

### Composite off-target score for gRNA specificity

To create a gRNA-level composite specificity score, individual off-target cutting scores are aggregated using the following inverse summation formula:

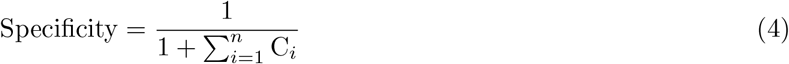

where C_*i*_ is the cutting likelihood score (either using the MIT or the CFD method) for the *i*^th^ putative off-target. A higher composite score indicates higher specificity, which decreases with more off-targets and/or a greater likelihood of cleavage at each off-target. A gRNA with no putative off-targets have a composite score of 1. A gRNA with 2 on-targets, that is a gRNA targeting two genomic loci with perfect complementarity, will have a composite score of 0.5.

### Evolutionary conservation scores

The function *addConservationScores* in *crisprDesign* annotates gRNAs with evolutionary conservation scores. It requires bigWig files containing basewise conservation scores, which can be easily obtained from the UCSC genome browser database [Karolchik et al., 2003] at the following link: https://hgdownload.soe.ucsc.edu/downloads.html. The gRNA score is calculated as the average conservation score of a region centered around the predicted cut site of the gRNA. By default, the width of the region is 18 nucleotides, but can be changed by users. For our analysis of human protein-coding genes, we used the phyloP score from an alignment of 29 genome sequences to the human genome available at https://hgdownload.soe.ucsc.edu/goldenPath/hg38/phyloP30way/. Positive phyloP scores indicate conserved regions, while negative scores indicate evolution faster than expected under neutral drift.

### Base editing scoring

The behavior of a base editor can be quantified in a 3-dimensional array of editing probabilities. Let *p* be the genomic position relative to the PAM site; let *nuc*_*u*_ be the original nucleotide; and let *nuc*_*e*_ be the edited nucleotide. Denote *q*(*p, nuc*_*u*_, *nuc*_*e*_) as the probability that *nuc*_*u*_ is edited to *nuc*_*e*_ at position *p*. Experimental editing weights can be used, possibly after some adequate transformation, to obtain those probabilities.

To score the likelihood of each edited allele, we assume independence of editing events with respect to nucleotide position. Specifically, consider a wildtype allele 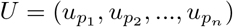 and an edited allele 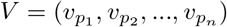, where 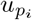 and 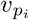 are the nucleotides at position *p*_*i*_ relative to the PAM site for the wildtype and edited allele, respectively. The parameter *n* is chosen by the user, and should be large enough so that all nucleotides within the editing window of the chosen base editor are represented. We calculate the editing score for the edited allele *V* (with respect to the wildtype allele *U*) as follows:

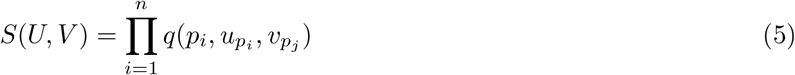

For a given edited allele *V*, we classify the functional consequence of editing as either a silent, missense, or nonsense mutation. We use *f* (*V*) to label the mutation. In case an edited allele results in more than one mutation, we choose the most consequential mutation as the label (nonsense over missense, and missense over silent). For a given gRNA targeting the wildtype allele *U*, and the set of all possible edited alleles *V*_*j*_, we calculate an aggregated score for each mutation type by summing the editing scores across alleles for each mutation type. For instance, the aggregated score for silent mutations is calculated as follows:

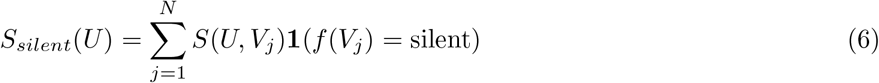

where *N* is the total number of possible edited alleles *V*_*j*_.

### Creation of major and minor allele human genomes

We built major and minor allele genomes for the hg38 build using common SNPs from the dbSNP151 RefSNP database. The “common” category is based on germline origin and a minor allele frequency (MAF) of *>*= 0.01 in at least one major population, with at least two unrelated individuals having the minor allele. See the dbSNP website https://www.ncbi.nlm.nih.gov/variation/docs/human_variation_vcf/ for more information. We excluded indels, and only considered SNPs that have MAF greater than 1% in the 1000 Genomes Project population. We then injected major alleles and minor alleles into the reference genome hg38 sequence to create “major allele” and “minor allele” genomes, respectively. Both resulting genomes are provided as standard

FASTA files. We generated *Bowtie* and *BWA* indexes for the two genomes. All results files are available on Zenodo (DOI: 10.5281/zenodo.6862556). The two allele genomes are also available from Bioconductor via their respective packages:

- `BSgenome.Hsapiens.UCSC.hg38.dbSNP151.major` [Fortin, 2021a]
- `BSgenome.Hsapiens.UCSC.hg38.dbSNP151.minor` [Fortin, 2021b]

### Base editing pooled screen data analysis

Fitness screen data in the MelJuSo cell line using a gRNA library tiling *BRCA1* were obtained from the supplementary material of Hanna et al. [2021]. We normalized the raw counts by scaling by the total number of reads, and log_2_-transformed the data. We filtered out low-abundance gRNAs that were further than 3 standard deviations below the mean in the plasmid (pDNA) sample. From the later timepoint samples, we subtracted from the pDNA sample log counts to obtain LFCs, and averaged the LFCs across replicates. We filtered out gRNAs targeting multiple loci, and gRNAs with off-targets (with up to 2 mismatches) located in genes other than *BRCA1*.

### CasRx pooled screen data analysis

CasRx FACS pooled screening data tiling *CD55, CD46* and *GFP* were obtained from Wessels et al. [2020], including processed and normalized LFCs for each gRNA https://gitlab.com/sanjanalab/cas13. We redesigned all possible gRNAs targeting any of the isoforms of *CD55* and *CD46* using *crisprDesign*, and considered only gRNAs also present in the pooled screening data for downstream analyses. We annotated all gRNAs with gene information (Ensembl release 104) and obtained off-targets with up to 3 mismatches for all gRNAs using *crisprBowtie*. We obtained CasRx-RF on-target activity scores using *crisprScore*. The transcripts annotated as canonical by Ensembl (ENST00000367042 for *CD46*, and ENST00000367064 for *CD55*) were used to visualize LFCs.

For each gRNA, we quantified the abundance of its target gene by summing transcript per million (TPM) counts in HEK-293 cells for all transcripts targeted by the gRNA. Transcript-level RNA quantification for HEK-293 cells was obtained from the Protein Atlas web portal https://www.proteinatlas.org, on March 5 2022. Data are based on The Human Protein Atlas version 21.0 and Ensembl version 103. We averaged TPM counts across the two replicates.

We used the single-mismatch (SM) gRNA constructs from the GFP tiling screen to estimate position-dependent probabilities of mismatch tolerance by the CasRx nuclease. To do so, we first calculated differences in LFC (ΔLFC) between SM gRNAs and their corresponding perfect-match (PM) gRNAs. We then fitted a LOESS curve with respect to the nucleotide position to obtain an average ΔLFC at each spacer position (Figure S5a). We transformed the LOESS fitted values to a scale between 0 and 1 to represent them as percentages of activity with respect to the median activity of the PM gRNAs tiling GFP (Figure S5b). Given the sparsity of the data, specifying a nucleotide-specific weight at each position was not possible. We adapted in *crisprScore* the CFD off-targeting scoring method to CasRx by using those probabilities as scoring weights. The corresponding scoring algorithm is named CFD-CasRx.

To evaluate the performance of the CFD-CasRx score on an independent dataset, we calculated CFD-CasRx off-target scores on all SM and double-mismatch (DM) gRNAs included in the *CD55* tiling screen. To predict LFCs of the DM gRNAs, we multiplied their respective PM gRNA LFCs with the CFD-CasRx on-target scores.

### Evaluation of the off-target alignment methods within *crisprDesign*

For comparing runtimes of the off-target alignment methods, the following sets of gRNAs were chosen: (1) gRNAs targeting the coding sequence of *KRAS*, for a total of 52 gRNAs; (2) gRNAs targeting the coding sequence of *EGFR*, for a total of 645 gRNAs, and (3) gRNAs targeting the coding sequence of *ZNF101*, for a total of 152 gRNAs. The *KRAS* and *EGFR* cases represent small- and medium-sized sets of gRNAs. For

*ZNF101*, a few gRNAs overlap a repeat element, and therefore have a high number of on- and off-targets. Alignment was performed to the GRCh38.p13 genome. The *Bowtie* and *Biostrings* alignment methods were evaluated using 0 to 3 mismatches, and the BWA alignment methods were evaluated using 0 to 5 mismatches.

Run times were collected on a Macbook Pro with an Intel Core i7 CPU (2.6GHz, 6 cores, 16 GB memory).

### Comparison of off-target alignments across tools

We compared computing times for designing SpCas9 gRNAs and performing a genome-wide off-target search for the following tools: *CCTop, CHOPCHOP, multicrispr, FlashFry*, and *crisprDesign*. The following tools were excluded from the comparison: *CRISPick* as it does not provide a standalone software; *CRISPRseek* as we were not able to complete the search within a reasonable time; *Cas-Designer* due to its requirement for specialized software that we were not able to install on our machines; *E-CRISP* as it was not possible to run their command line interface on customs DNA sequences or exons.

To perform the comparison, we generated six random subsets of protein-coding exons located on chr1 with the following sizes:100, 200, 400, 800, 1600 and 3200 exons. Off-target alignment was performed against the human reference genome (GRCh38 build) using a maximum of 2 mismatches for all methods. Run times were collected on a Macbook Pro with an Intel Core i7 CPU (2.6GHz, 6 cores, 16 GB memory). For each tool, parameters optimized for speed were chosen based on available documentation. In particular, the following parameters were used. For *CCTop*, we used `--totalMM 2 --coreMM 2 --maxOT 100000`. For *CHOPCHOP*, we used: `--fasta -G HG38 -t WHOLE -v 2`. For *FlashFry*, we used: `--maximumOffTargets 100000 --forceLinear -` `-maxMismatch 2`. For *multicrispr*, we used Bowtie with 2 mismatches, with no on-target scoring. For *crisprDesign*, we used the function *addSpacerAlignmentsIterative* with the *Bowtie* and *BWA* aligners with 2 mismatches.

### Processing of genome-wide screen datasets

#### Achilles dataset

CRISPRko fitness screening gRNA-level LFCs from Project Achilles (22Q2 release) were downloaded from the DepMap portal https://depmap.org/portal/download/all/. Processed LFCs representing changes in gRNA abundances between the last time point of the fitness screen and the plasmid DNA were available for 957 human cell lines. Reference lists of essential and non-essential genes were downloaded from Hart et al. [2014]. For each cell line, we first centered LFCs using the median value of the set of non-essential genes, and then scaled LFCs using the median value of the set of essential genes. This enables normalized LFCs to be comparable across cell lines. For each gRNA, we then summarized gRNA activity by averaging LFCs across cell lines.

#### Hart2015 dataset

Processed data from a genome-wide screen performed in HCT116 cells using the Toronto Knockout v1 (TKOv1) library [Hart et al., 2015] were downloaded from http://tko.ccbr.utoronto.ca/. We computed LFCs between Day 18 and Day 0.

#### Hart2017 dataset

Processed data from a genome-wide screen performed in HAP1 cells using the Toronto Knockout v3 (TKOv3) library [Hart et al., 2017b] were downloaded from http://tko.ccbr.utoronto.ca/. We obtained available LFCs between Day 18 and Day 0.

#### Wang2015 dataset

Processed data from a genome-wide screen performed in K562 cells were obtained from the supplementary material of Wang et al. [2015]. LFCs were calculated between the final and initial timepoints.

#### Tzelepis2016 dataset

Processed LFCs from a genome-wide screen performed in HL60 cells were obtained from the supplementary material of Tzelepis et al. [2016].

LFCs for the Hart2015, Hart2017, Wang2015 and Tzelepis2016 were further standardized using the approach used for the Achilles dataset, with the same sets of non-essential and essential genes. For each dataset, gRNAs were mapped to the set of human protein-coding genes found in the Ensembl release 104, and unmapped gRNAs were filtered out. Given that gRNAs with multiple on- and off-targets can confound the analysis of fitness screens [Fortin et al., 2019], we removed gRNAs that map to multiple loci in the GRCh38 genome, as well as gRNAs with 1- and 2-mismatch off-targets located in coding regions other than the intended target. The final numbers of gRNAs further considered for analysis are presented in Table 3.

### Default gRNA rankings implemented in *crisprDesign*

For each nuclease, we rank gRNAs based on several rounds of priority. For SpCas9, gRNAs with unique target sequences and without one- or two-mismatch off-targets located in coding regions are placed into the first round. Then, gRNAs with a small number of one- or two-mismatch off-targets (less than 5) are placed into the second round. Remaining gRNAs are placed into the third round. Finally, any gRNAs overlapping a common SNP (human only), containing a polyT stretch, or with extreme GC content (below 20% or above 80%) are placed into the fourth round. For CRISPRko applications, within each round of selection, gRNAs targeting the first 85% of the coding sequence of the canonical Ensembl isoform, as well as gRNAs targeting conserved regions (phyloP conservation score greater than 0), are prioritized first. gRNAs with the same priority are then ranked by a composite on-target activity rank to further prioritize active gRNAs. Based on the consistently reliable performance performance and generalization of the methods *DeepHF* and *DeepSpCas9* shown in Konstantakos et al. [2022], Wang et al. [2019b], Kim et al. [2019], the composite on-target activity rank is calculated by taking the average rank across the *DeepHF* and *DeepSpCas9* scores. For CRISPRa and CRISPRi applications, the CRISPRai on-target score is used instead of the composite score.

The process is identical for enAsCas12a, with the exception that the *enPAM+GB* method is used as the composite score given that it is the only method available for the enAsCas12a nuclease. For CasRx, gRNAs targeting at least 75% of the isoforms of a given gene, with no one- or two-mismatch off-targets, are placed into the first round. gRNAs targeting at least 50% of the isoforms of a given gene, with no one- or two-mismatch off-targets, are placed into the second round, and remaining gRNAs are placed into the third round. Finally, any gRNAs containing a polyT stretch, or with extreme GC content (below 20% or above 80%) are placed into the fourth round. Within each round of selection, gRNAs are further ranked by the *CasRxRF* on-target score, using the canonical Ensembl isoform for scoring.

### Generation of gRNA rankings from other tools

In addition to *crisprDesign*, we designed and ranked SpCas9 gRNAs for all human protein-coding (Ensembl release 104) using four additional tools. For *CHOPCHOP* (v3), we used the command line interface (CLI) available at https://bitbucket.org/valenlab/chopchop with default parameters. For *CCTop* (v1.0.0), we used the CLI available at https://bitbucket.org/juanlmateo/cctop_standalone with default parameters. For *FlashFry* (v1.15), we used the CLI available at https://github.com/mckennalab/FlashFry with default parameters. For *CRISPick*, due to the lack of a CLI, we submitted batch query jobs through the portal https://portals.broadinstitute.org/gppx/crispick/public (accessed on July 27 2022) with default parameters for the Hsu (2013) tracrRNA sequence using the Rule Set 3.

### Criteria used to compare feature availability across gRNA design tools

The following gRNA design tools were used for comparison in Table 1: *multicrispr* (v1.7.0), *CRISPRseek* (v1.37.2), *CHOPCHOP* (v3), *CRISPOR* (website v5.01), *CCTop* (v1.0.0), *Guides* (v1.0), *Cas-Designer* (v3.0), *FlashFry* (v1.15), *E-CRISP* (v5.4) and *CRISPick* (no version, accessed on July 27 2022). The criteria listed below were used for assessing feature availability.

#### Nuclease section

a check mark indicates support for the corresponding nuclease, and *Limited* indicates that only a subset of custom nucleases are available. *Modalities* section: a check mark indicates that the software offers at least one specific functionality for that modality. *Target space section*: for the *Reference genomes* row, a check mark indicates that the software supports gRNA design against reference genomes; for this row, *Limited* indicates that the versions of the reference genomes are outdated. For the *Custom sequences* row, a check mark indicates that the software supports the design of gRNAs targeting custom DNA sequences.

The *Off-target aligner* section indicates which alignment methods are available in each tool. The *Off-target options* section describes which off-target alignment functionalities are implemented: genomic coordinates of the off-targets are available to the user (*Genomic coordinates* row), off-target alignment to custom sequences (*Custom sequences* row), concurrent off-target alignment to multiple organisms (*cross-reactivity* row), and alignment to major or minor allele genomes (*Minor/major alleles* row). The *On-target* and *Off-target* scoring sections indicate which scoring methods are implemented in the software.

The *Annotations* section indicates whether or not users have access to several annotations in the gRNA outputs. *Off-target annotation* refers to gene context annotation of the off-targets; *Isoform specification* refers to information about which gene isoforms are targeted by a given gRNA; *Reinitiation sites* refers to gRNAs annotated as being upstream of potential reinitiation sites; *Pfam domains* refers to information about which Pfam domains are targeted by a given gRNA; *SNP annotation* refers to an annotation of gRNAs overlapping common SNPs; *TSS annotation* refers to whether or not gRNAs are annotated to fall into the promoter region of knows TSSs; *Conservation* refers to evolutionary conservation annotation.

The *Library design* section indicates which library design features are available in each of the tools. *Restriction sites* indicates whether or not gRNAs can be filtered for restriction sites of common enzymes. *PolyT signal* indicates if PolyT stretch filtering is available. *GC content* indicates filtering based on percentage GC content. *Hairpin loops* indicates filtering based on potential self-complementarity. *Paired gRNAs* indicates whether or not design of paired gRNAs is enabled. *Ranking* indicates if the software returns a gRNA rank for user selection.

### Figure generation

All figures were made in R (4.2.1), with the exception of the following figures that were made in Microsoft PowerPoint (v16.64): Figure 1, Figure 2a-b, and the workflow diagrams of Figure 4, Figure 5 and Figure 6. Figure 2c and Figure 6b were made using the R package *Gviz* (v1.41.1). Figure 3, Figure 4, Figure 5, Figure S1, Figure S3, Figure S4 and Figure S5 were made using base plotting functions in R. Reproducible code to generate all figures can be found in our [GitHub manuscript repository](https://github.com/crisprVerse/crisprVersePaper).

## Supporting information

Tutorial1_Installation

Tutorial16_Validating_Libraries

Tutorial11_PairedDesign

Tutorial13_Optical_Pooled_Screening

Tutoria15_Major_and_Minor_Alleles

Tutorial12_CustomSequences

Tutorial14_Mapping_Across_Species

Tutorial5_CRISPRko_Design_Cas9

Tutorial6_CRISPRko_Design_Cas12a

Tutorial7_CRISPRko_Design_CasRx

Tutorial10_CRISPRi_Design

Tutorial8_CRISPRbe_Design

Tutorial9_CRISPRa_Design

Tutorial2_Building_Genome_Index

Tutorial4_Building_Gene_Annotation

Tutorial17_Building_gRNA_Database

Tutorial3_Specifying_CRISPR_Nuclease

Vignette_crisprBase

Vignette_crisprBowtie

Vignette_crisprBwa

Vignette_crisprDesign

Vignette_crisprDesignData

Vignette_crisprDesignScore

Vignette_crisprDesignVerse

Vignette_crisprViz

## Data availability

Chromatin accessibility data from Horlbeck et al. [2016] necessary for the CRISPRai on-target algorithm are available on Zenodo (DOI: 10.5281/zenodo.6716721). Fasta files, *Bowtie* indexes and *BWA* indexes for the major and minor alleles of hg38 using dbSNP151 are available on Zenodo (DOI: 10.5281/zenodo.6862556). We precomputed and fully annotated gRNAs for human and mouse protein-coding genes using *crisprDesign* for the following nucleases: SpCas9, enAsCas12a, and CasRx. Ensembl release 104 and Ensembl release 102 were used to define genes for human and mouse, respectively. Separate datasets were generated for the CRISPRko, CRISPRa, CRISPRi, and CRISPRkd modalities. All files are available on Zenodo (DOI: 10.5281/zenodo.7042164).

## Software and code availability

All crisprVerse packages are open-source and available on GitHub (Table 4). At time of publication, all packages were accepted at Bioconductor and available on the development branch of Bioconductor. Because of its size, the data package *crisprDesignData* is hosted on GitHub only. Reproducible code of all analyses can be found at https://github.com/crisprVerse/crisprDesignPaper. A list of extensive tutorials can be found at https://github.com/crisprVerse/Tutorials.

**Table 4.** R packages in the crisprVerse ecosystem.

| R package | Description |
| --- | --- |
| crisprVerse | Easily install of the crisprVerse ecosystem |
| crisprDesign | Core package for gRNA design |
| crisprBase | Nuclease specification and gRNA arithmetics |
| crisprBowtie | gRNA spacer alignment with *Bowtie* |
| crisprBwa | gRNA spacer alignment with *BWA* |
| crisprScore | On- and off-target scoring algorithms for gRNAs |
| crisprViz | Visualization of gRNAs using genomic tracks |
| Rbwa | R wrapper for *BWA* aligner |
| crisprScoreData | Pre-trained machine learning models for *crisprScore* |
| crisprDesignData | Pre-computed data for the crisprVerse ecosystem (human and mouse) |

The analyses included in this paper were produced using the following package versions: *crisprDesign* (v0.99.134), *crisprScore* (v1.1.14), *crisprScoreData* (v1.1.3), *crisprBowtie* (v1.1.1), *crisprBase* v(1.1.5), *crisprVerse* v(0.99.8), *crisprBwa* (v1.1.3), *Rbwa* (v1.1.0), *crisprDesignData* (0.99.17), *crisprViz* (0.99.18)

We also offer a Docker container encapsulating the latest crisprVerse ecosystem on our [DockerHub page](https://hub.docker.com/repository/docker/fortin946/crisprverse). Documentation about the installation and usage of the container can be found [here](https://github.com/crisprVerse/Docker).

## Abbreviations

ABE: adenine base editor
BWA: Burrows-Wheeler Aligner
CAGE: Cap Analysis of Gene Expression
CBE: cytosine base editor
CCLE: cancer cell line encyclopedia
CDS: coding DNA sequence
CFD: cutting frequency determination
CLI: command line interface
CRISPR: clustered regularly interspaced short palindromic repeats
CRISPRa: CRISPR activation
CRISPRbe: CRISPR base editing
CRISPRi: CRISPR interference
DHS: DNAse I hypersensitive site
DSB: DNA double-strand break
FACS: fluorescence-activated cell sorting
LFC: log-fold change
lncRNAs: long non-coding RNAs
MAF: minor allele frequency
OPS: optical pooled screening
PAM: protospacer adjacent motif
PE: prime editing
pegRNA: prime editing guide RNA
PFS: protospacer flanking sequence
RNAi: RNA interference
gRNA: single-guide RNA
SNP: single-nucleotide polymorphism
TSS: transcription starting site

## Competing interests

The authors declare that they have no competing interests.

## Authors contributions

JPF led the software development and supervised the work. JPF conceptualized and wrote the manuscript, with contributions and input from all authors. LH and JPF developed the R packages, with contributions from PP and AL. All authors read and approved the final manuscript.

## Acknowledgements

We thank Benjamin Haley, Mike Costa, Amy Heidersbach, Kristel Dorighi, Scott Martin, Rena Yang, Allison Vuong, Oleg Mayba, Sandra Melo Carlos, and Russell Xie for sharing their expertise with us and guiding the development of our software ecosystem. We also thank William Forrest, Maggie Crow, Hector Corrada Bravo, Michael Lawrence, and Benjamin Haley for providing invaluable feedback on the manuscript and software. We thank Nitesh Turaga, Lori Shepherd, Marcel Ramos, Helena Crowell and Kayla Morrell who kindly and thoroughly reviewed our R packages as part of the Bioconductor submission process. Finally, we want to thank the referees for their invaluable input and suggestions.

## Supplementary Tables and Figures

**Supplementary Table S1.**
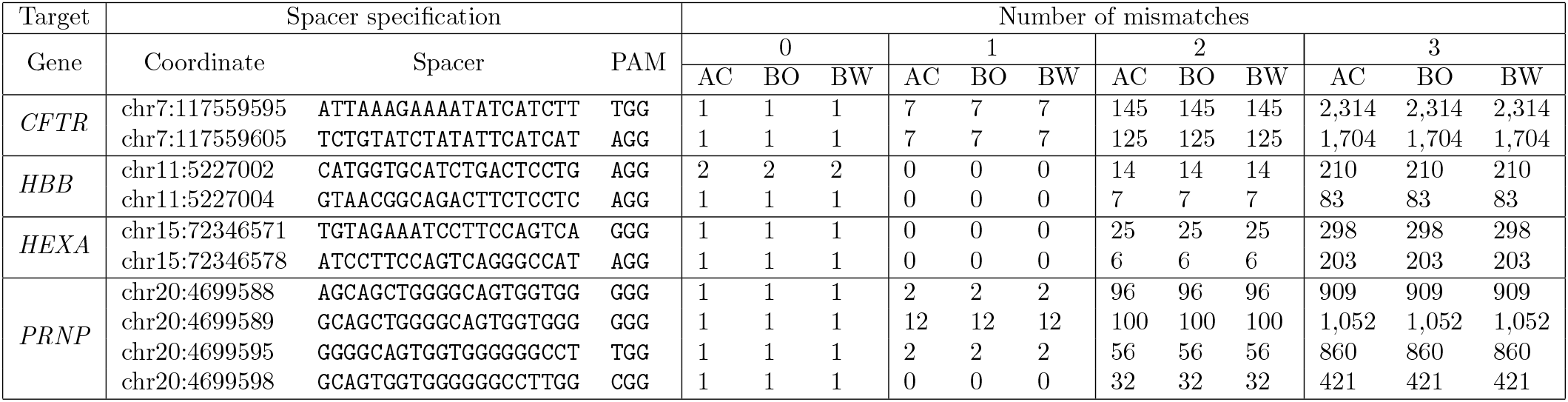
Table of on- and off-target alignments in the GRCh38.p13 for the 10 spacer sequences reported in Bhagwat et al. [2020] using a PAM-agnostic approach. Number of mismatches between 0 and 3 were considered for 3 different aligners: Aho-Corasick exact string matching as implemented in *Biostrings* (AC), *Bowtie* aligner via the *crisprBowtie* package (BO), and *BWA* aligner via the *crisprBwa* package (BW). All 3 alignment methods agree.

**Supplementary Figure S1.**
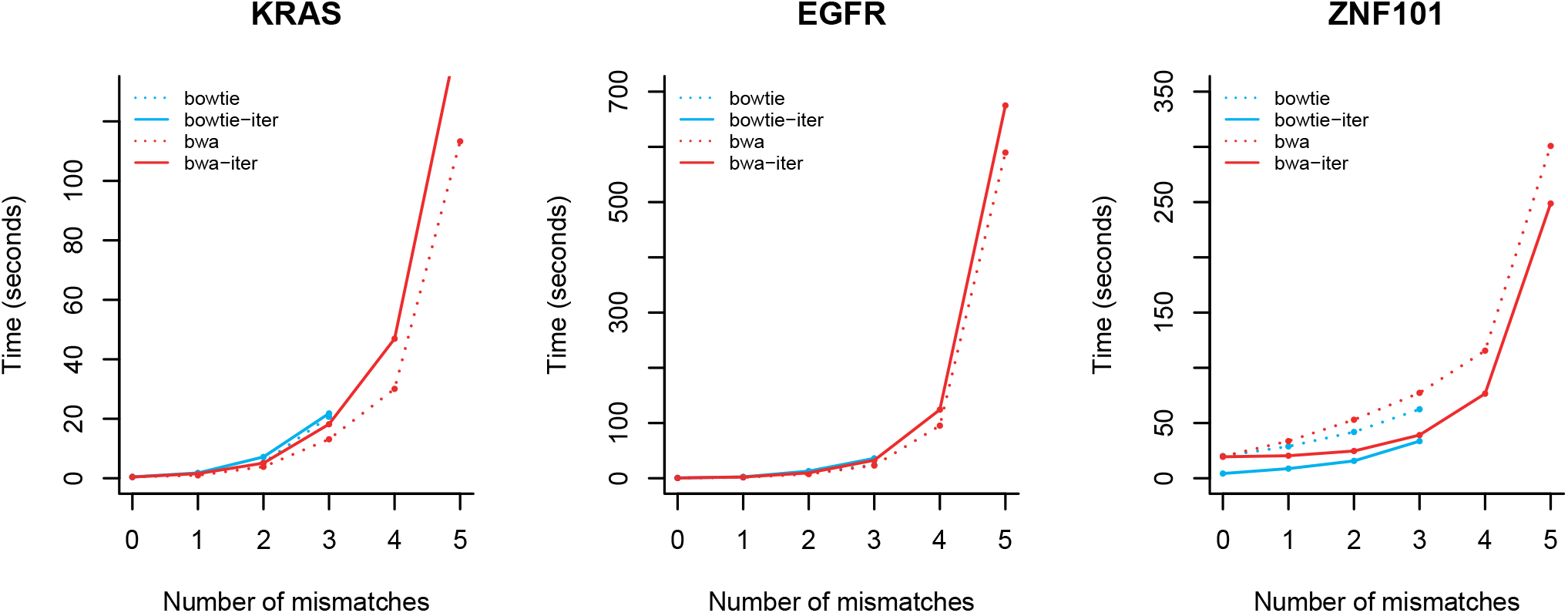
Comparison of computing times for off-target alignment methods implemented in *crisprDesign*. We compare computing time for the four different off-target methods available via the *addSpacerAligmments* function in *crisprDesign*: *Bowtie*, via the *crisprBowtie* package (`Bowtie`), *BWA*, via the *crisprBwa* package (`bwa`), and an iterative version of both algorithms to diminish the impact of highly non-specific gRNAs on computing time (`bowtie-iter` and `bwa-iter`).

**Supplementary Figure S2.**
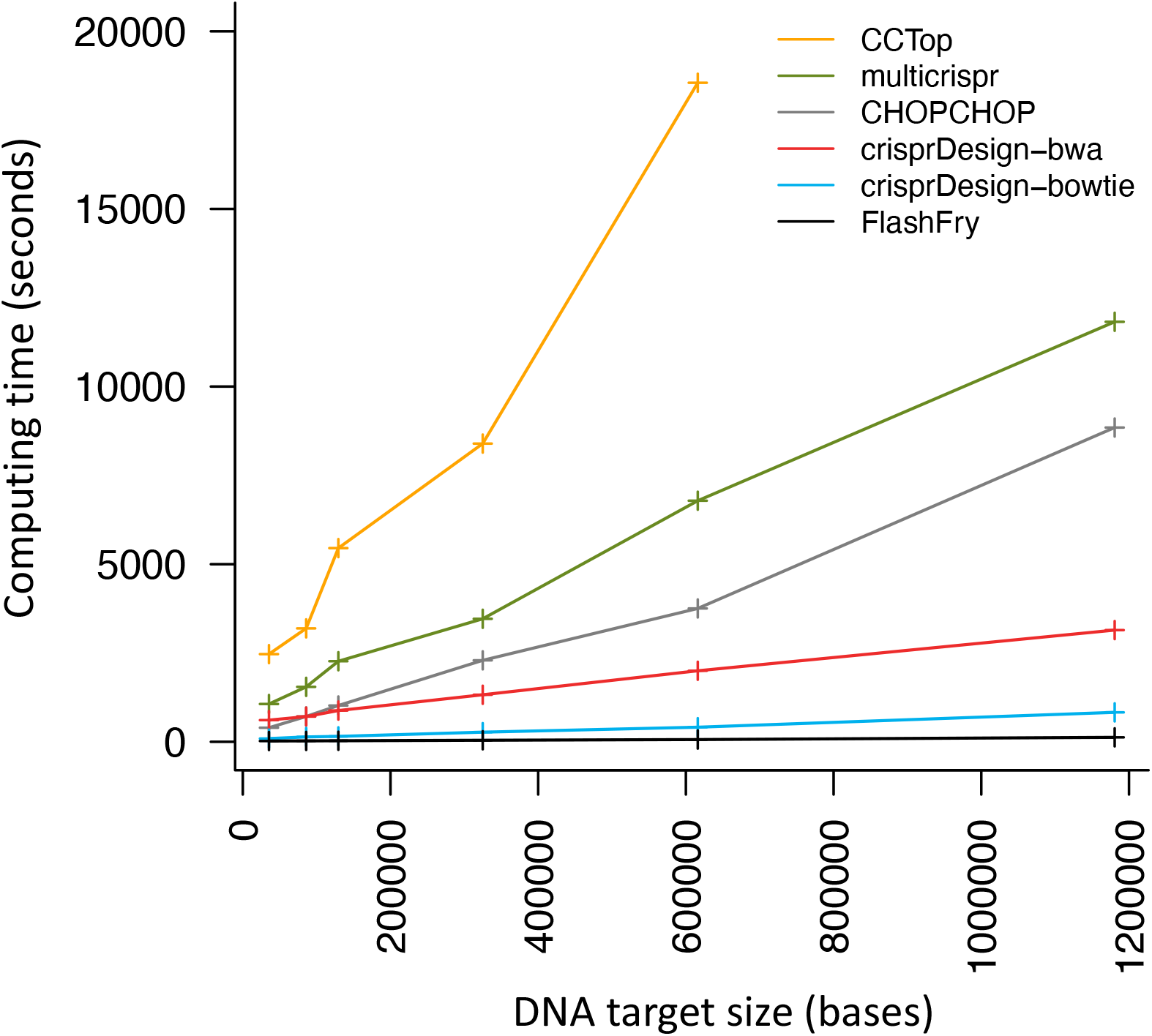
Comparison of computing times for subsets of human protein-coding exons. We compared computing times across tools to design gRNAs and perform a genome-wide off-target search in the human genome. Six random subsets of protein-coding exons located on chr1 were used to perform the comparison. The sizes of the subsets were 100, 200, 400, 800, 1600 and 3200 exons. The x-axis shows the total size in nucleotides of the DNA target space formed by each subset, and the y-axis shows computing times in seconds. Details about the alignment parameters for each method can be found in the Methods section.

**Supplementary Figure S3.**
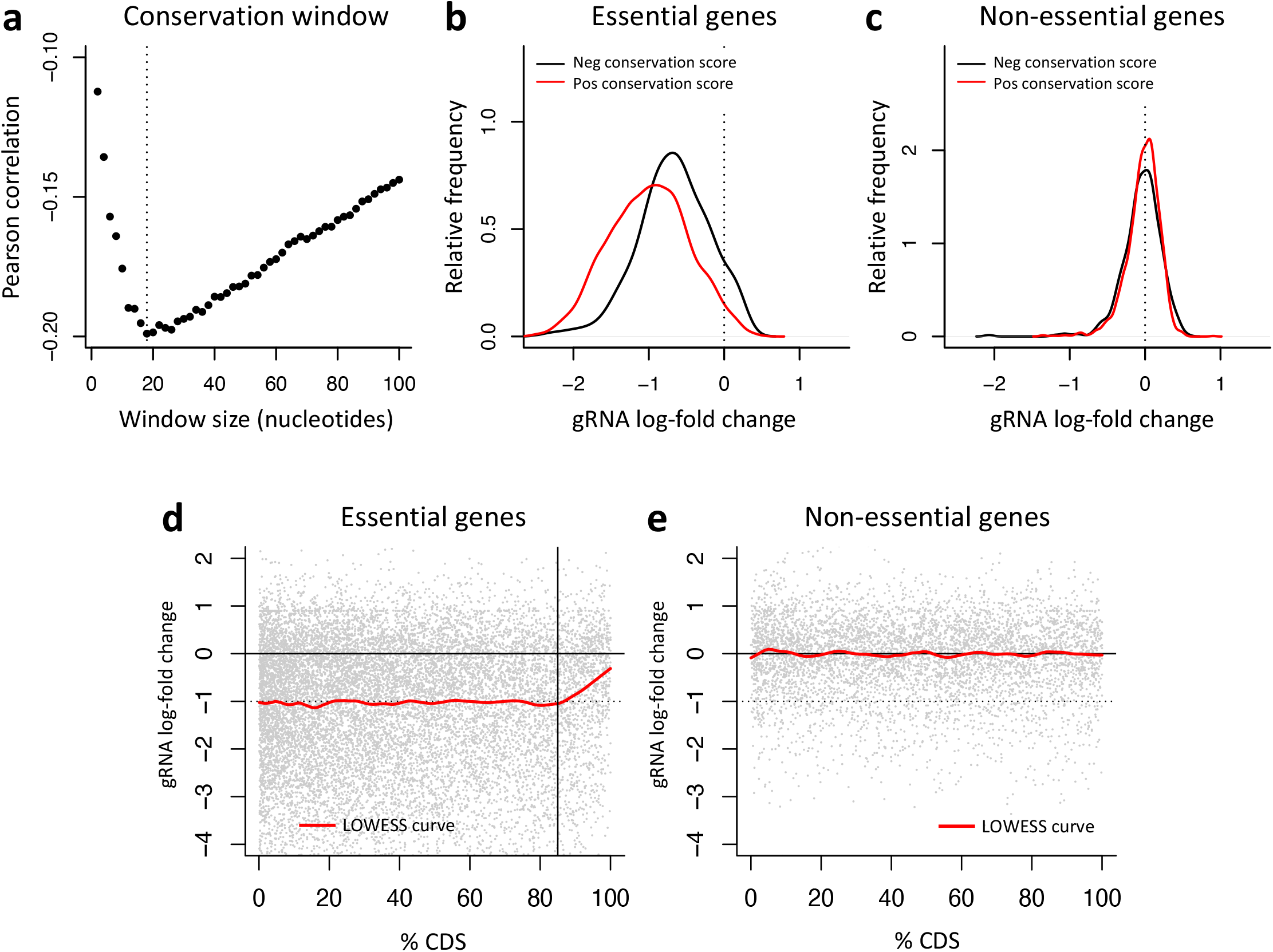
Influence of evolutionary conservation and gene target position on gRNA activity. **a-c** We annotated each gRNA present in Project Achilles with a conservation score using the function *addConservationScores* implemented in *crisprDesign* (see Methods). The gRNA conservation score is taken as the average DNA conservation score across nucleotides in a user-specified window around the gRNA cut site. In **a**, we show the correlation between observed gRNA activity and the conservation scores for different window sizes for essential genes. The data suggest an optimal window of 18 nucleotides around the cut site. **b** Distributions of the observed gRNA log-fold changes (LFCs) based on whether or not gRNAs are targeting regions of high conservation (positive gRNA conservation score) or regions of low conservation (negative gRNA conservation score), for gRNAs targeting essential genes. **c** Same as **b**, but for gRNAs targeting non-essential genes. **d** Relationship between gRNA activity and gRNA position within the target coding sequence (CDS) for gRNAs targeting essential genes in the Hart2015 dataset. The Ensembl canonical transcript was used as the target CDS for each gene. The red curve represents a LOWESS trend. gRNAs located beyond the first 85% of the CDS (to the right of the the vertical line) show a progressive decline in activity. **e** Same as **d**, but for gRNAs targeting non-essential genes.

**Supplementary Figure S4.**
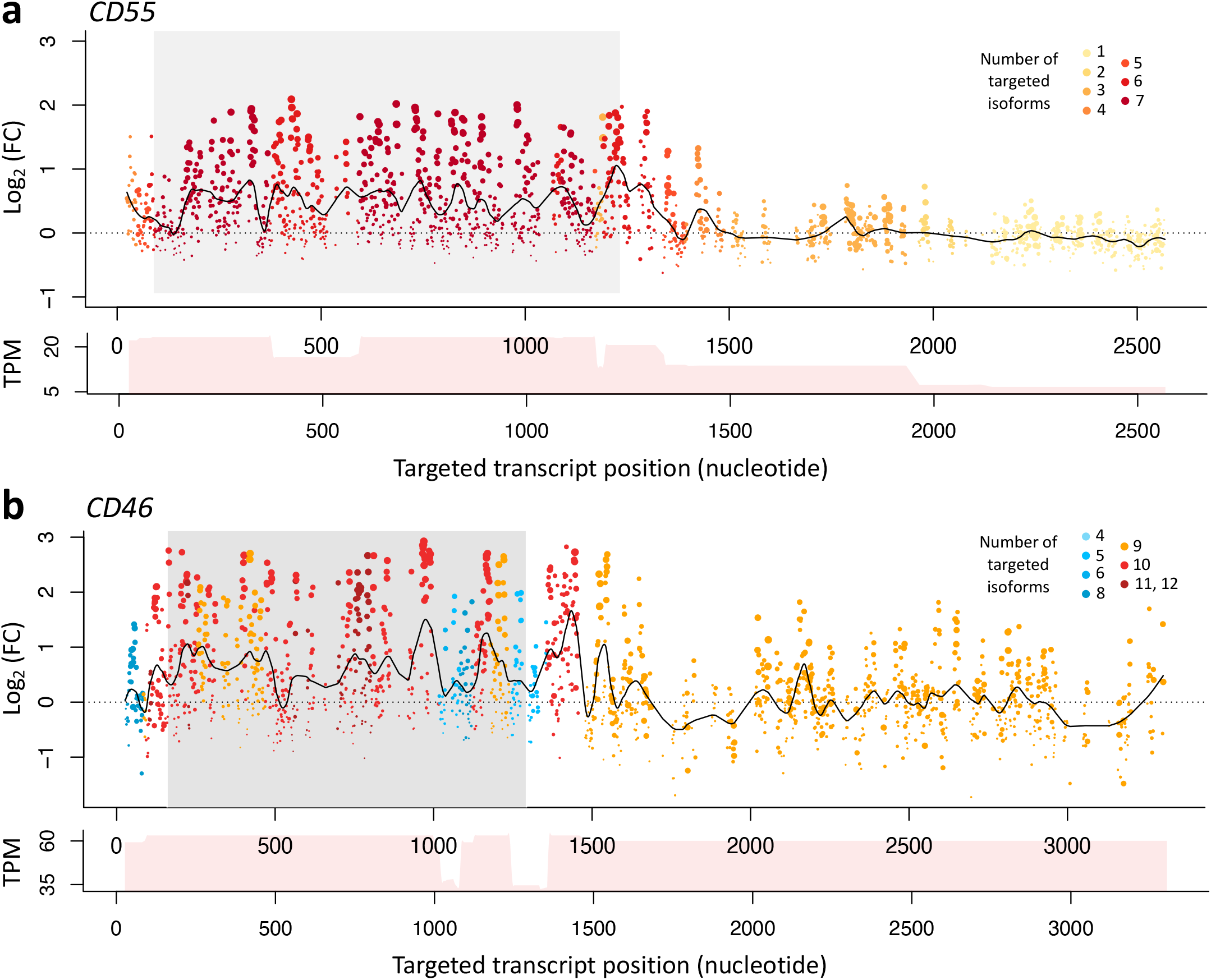
CasRx tiling screens of *CD55* and *CD46*. Pooled FACS tiling screening data of genes *CD55* and *CD46* performed in HEK 294 cells using CasRx (*Rfx*Cas13d). Processed and normalized log_2_ fold changes were obtained from Wessels et al. [2020]. Both screens are represented using the canonical Ensembl isoforms. We remapped and reannotated all gRNA sequences using *crisprDesign*; isoform annotation, on-target activity score using CasRx-RF as implemented in *crisprScore*, and off-target alignments were added to each gRNA. The color of the dots indicates the number of isoforms targeted by each gRNA. The size of the dots is proportional to the on-target activity score. The coding sequence (CDS) is highlighted in grey. LOESS regression curves are shown as solid lines. For both genes, transcript per million (TPM) counts in HEK 293 cells summed across all isoforms overlapping a given nucleotide position are shown below the log-fold change panels.

**Supplementary Figure S5.**
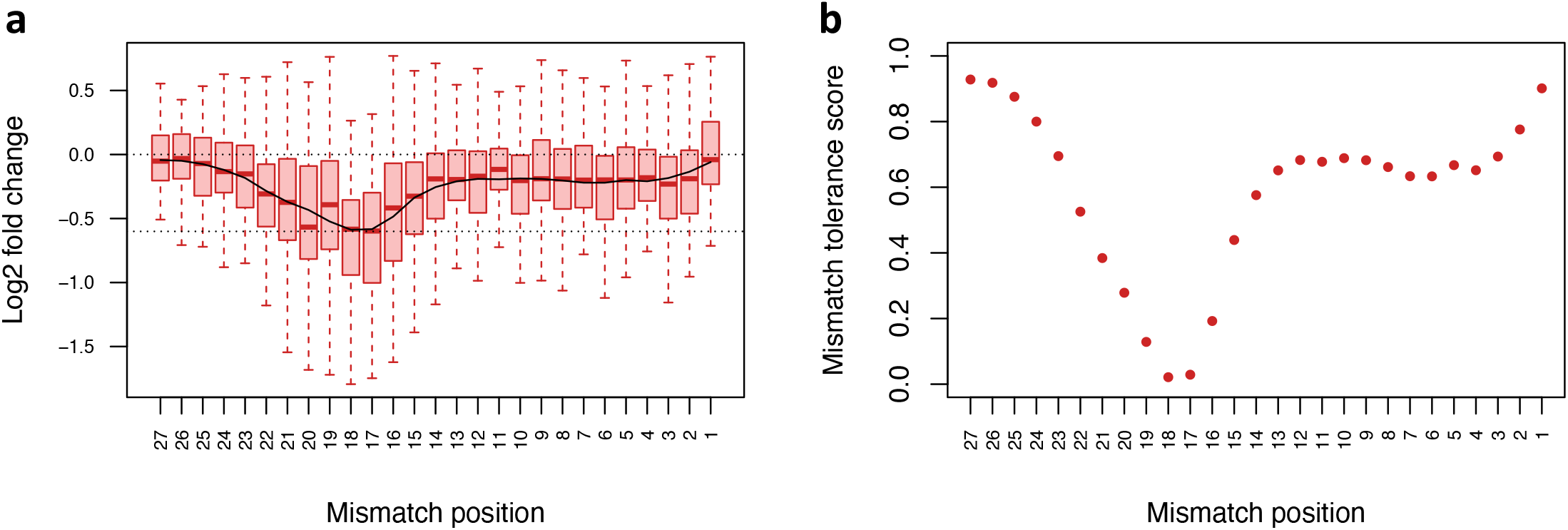
Probability weights used for off-target scoring of CasRx gRNAs. **a** Boxplots of the differences in log2 fold change (ΔLFC) between single-mismatch (SM) gRNAs and their corresponding perfect-match (PM) gRNAs in the GFP tiling screen. X-axis represents the mismatch position within the spacer sequence, with 1 being the position next to the direct repeat. The smooth curve was obtained using LOESS regression. The dotted line represents the average log-fold change of all PM gRNAs after multiplying by -1. **b** CasRx mismatch tolerance probabilities estimated from (a) and used in the CFD scoring method in *crisprScore*.

## Notes

https://github.com/crisprVerse

### Competing Interest Statement

The authors have declared no competing interest.

### Summary of Updates

- Added crisprViz and crisprVerse packages - Added tutorials - Added an analysis of gRNA ranking - Added conservation scores

## References

Patrick Aboyoun, Herve Pages, and Michael Lawrence. GenomicRanges: Representation and manipulation of genomic intervals, 2012.

Omar O Abudayyeh, Jonathan S Gootenberg, Silvana Konermann, Julia Joung, Ian M Slaymaker, David BT Cox, Sergey Shmakov, Kira S Makarova, Ekaterina Semenova, Leonid Minakhin, et al. C2c2 is a single-component programmable rna-guided rna-targeting crispr effector. Science, 353(6299), 2016.

Alfred V Aho and Margaret J Corasick. Efficient string matching: an aid to bibliographic search. Communications of the ACM, 18(6):333–340, 1975.

Felicity Allen, Fiona Behan, Anton Khodak, Francesco Iorio, Kosuke Yusa, Mathew Garnett, and Leopold Parts. Jacks: joint analysis of crispr/cas9 knockout screens. Genome research, 29(3):464–471, 2019.

Andrew V Anzalone, Peyton B Randolph, Jessie R Davis, Alexander A Sousa, Luke W Koblan, Jonathan M Levy, Peter J Chen, Christopher Wilson, Gregory A Newby, Aditya Raguram, et al. Search-and-replace genome editing without double-strand breaks or donor dna. Nature, 576(7785):149–157, 2019.

Mandana Arbab, Max W Shen, Beverly Mok, Christopher Wilson, Żaneta Matuszek, Christopher A Cassa, and David R Liu. Determinants of base editing outcomes from target library analysis and machine learning. Cell, 182(2):463–480, 2020.

Sangsu Bae, Jeongbin Park, and Jin-Soo Kim. Cas-offinder: a fast and versatile algorithm that searches for potential off-target sites of cas9 rna-guided endonucleases. Bioinformatics, 30(10):1473–1475, 2014.

Russell Bainer, Dariusz Ratman, Peter Haverty, and Steve Lianoglou. gCrisprTools: Suite of Functions for Pooled Crispr Screen QC and Analysis, 2021. R package version 2.0.0.

Alex Bateman, Lachlan Coin, Richard Durbin, Robert D Finn, Volker Hollich, Sam Griffiths-Jones, Ajay Khanna, Mhairi Marshall, Simon Moxon, Erik LL Sonnhammer, et al. The pfam protein families database. Nucleic acids research, 32(Suppl 1):D138–D141, 2004.

Aditya M Bhagwat, Johannes Graumann, Rene Wiegandt, Mette Bentsen, Jordan Welker, Carsten Kuenne, Jens Preussner, Thomas Braun, and Mario Looso. multicrispr: grna design for prime editing and parallel targeting of thousands of targets. Life science alliance, 3(11), 2020.

Matthew C Canver, Samuel Lessard, Luca Pinello, Yuxuan Wu, Yann Ilboudo, Emily N Stern, Austen J Needleman, Frédéric Galactéros, Carlo Brugnara, Abdullah Kutlar, et al. Variant-aware saturating mutagenesis using multiple cas9 nucleases identifies regulatory elements at trait-associated loci. Nature genetics, 49(4):625, 2017.

Chen-Hao Chen, Tengfei Xiao, Han Xu, Peng Jiang, Clifford A Meyer, Wei Li, Myles Brown, and X Shirley Liu. Improved design and analysis of crispr knockout screens. Bioinformatics, 2018.

Wei Chen, Aaron McKenna, Jacob Schreiber, Maximilian Haeussler, Yi Yin, Vikram Agarwal, William Stafford Noble, and Jay Shendure. Massively parallel profiling and predictive modeling of the outcomes of crispr/cas9-mediated double-strand break repair. Nucleic acids research, 47(15):7989–8003, 2019.

Sarit Cohen, Lior Kramarski, Shahar Levi, Noa Deshe, Oshrit Ben David, and Eyal Arbely. Nonsense mutation-dependent reinitiation of translation in mammalian cells. Nucleic acids research, 47(12):6330–6338, 2019.

Jean-Paul Concordet and Maximilian Haeussler. Crispor: intuitive guide selection for crispr/cas9 genome editing experiments and screens. Nucleic acids research, 46(W1):W242–W245, 2018.

David BT Cox, Jonathan S Gootenberg, Omar O Abudayyeh, Brian Franklin, Max J Kellner, Julia Joung, and Feng Zhang. Rna editing with crispr-cas13. Science, 358(6366):1019–1027, 2017.

Joshua M Dempster, Isabella Boyle, Francisca Vazquez, David E Root, Jesse S Boehm, William C Hahn, Aviad Tsherniak, and James M McFarland. Chronos: a cell population dynamics model of crispr experiments that improves inference of gene fitness effects. Genome biology, 22(1):1–23, 2021.

Peter C DeWeirdt, Kendall R Sanson, Annabel K Sangree, Mudra Hegde, Ruth E Hanna, Marissa N Feeley, Audrey L Griffith, Teng Teng, Samantha M Borys, Christine Strand, et al. Optimization of ascas12a for combinatorial genetic screens in human cells. Nature Biotechnology, pages 1–11, 2020.

Peter C. DeWeirdt, Abby V. McGee, Fengyi Zheng, Ifunanya Nwolah, Mudra Hegde, and John G. Doench. Accounting for small variations in the tracrrna sequence improves sgrna activity predictions for crispr screening. Nature Communications, 13(1):5255, 2022. doi:10.1038/s41467-022-33024-2. URL https://doi.org/10.1038/s41467-022-33024-2.

Sarah Djebali, Carrie A Davis, Angelika Merkel, Alex Dobin, Timo Lassmann, Ali Mortazavi, Andrea Tanzer, Julien Lagarde, Wei Lin, Felix Schlesinger, et al. Landscape of transcription in human cells. Nature, 489 (7414):101–108, 2012.

John G Doench, Ella Hartenian, Daniel B Graham, Zuzana Tothova, Mudra Hegde, Ian Smith, Meagan Sullender, Benjamin L Ebert, Ramnik J Xavier, and David E Root. Rational design of highly active sgrnas for crispr-cas9–mediated gene inactivation. Nature biotechnology, 32(12):1262, 2014.

John G Doench, Nicolo Fusi, Meagan Sullender, Mudra Hegde, Emma W Vaimberg, Katherine F Donovan, Ian Smith, Zuzana Tothova, Craig Wilen, Robert Orchard, et al. Optimized sgrna design to maximize activity and minimize off-target effects of crispr-cas9. Nature biotechnology, 34(2):184, 2016a.

John G Doench, Nicolo Fusi, Meagan Sullender, Mudra Hegde, Emma W Vaimberg, Katherine F Donovan, Ian Smith, Zuzana Tothova, Craig Wilen, Robert Orchard, et al. Optimized sgrna design to maximize activity and minimize off-target effects of crispr-cas9. Nature biotechnology, 34(2):184, 2016b.

Steffen Durinck, Yves Moreau, Arek Kasprzyk, Sean Davis, Bart De Moor, Alvis Brazma, and Wolfgang Huber. Biomart and bioconductor: a powerful link between biological databases and microarray data analysis. Bioinformatics, 21(16):3439–3440, 2005.

David Feldman, Avtar Singh, Jonathan L Schmid-Burgk, Rebecca J Carlson, Anja Mezger, Anthony J Garrity, Feng Zhang, and Paul C Blainey. Optical pooled screens in human cells. Cell, 179(3):787–799, 2019.

Gregory M Findlay, Riza M Daza, Beth Martin, Melissa D Zhang, Anh P Leith, Molly Gasperini, Joseph D Janizek, Xingfan Huang, Lea M Starita, and Jay Shendure. Accurate classification of brca1 variants with saturation genome editing. Nature, 562(7726):217–222, 2018.

Jean-Philippe Fortin. BSgenome.Hsapiens.UCSC.hg38.dbSNP151.major: Full genome sequences for Homo sapiens (UCSC version hg38, based on GRCh38.p12) with injected major alleles (dbSNP151), 2021a. R package version 0.0.9999.

Jean-Philippe Fortin. BSgenome.Hsapiens.UCSC.hg38.dbSNP151.minor: Full genome sequences for Homo sapiens (UCSC version hg38, based on GRCh38.p12) with injected minor alleles (dbSNP151), 2021b. R package version 0.0.9999.

Jean-Philippe Fortin and Aaron Lun. crisprScore: On-Target and Off-Target Scoring Algorithms for CRISPR gRNAs, 2022. URL https://github.com/Jfortin1/crisprScore/issues. R package version 0.99.16.

Jean-Philippe Fortin, Jenille Tan, Karen E Gascoigne, Peter M Haverty, William F Forrest, Michael R Costa, and Scott E Martin. Multiple-gene targeting and mismatch tolerance can confound analysis of genome-wide pooled crispr screens. Genome biology, 20(1):21, 2019.

Yanfang Fu, Jennifer A Foden, Cyd Khayter, Morgan L Maeder, Deepak Reyon, J Keith Joung, and Jeffry D Sander. High-frequency off-target mutagenesis induced by crispr-cas nucleases in human cells. Nature biotechnology, 31(9):822–826, 2013.

Nicole M Gaudelli, Alexis C Komor, Holly A Rees, Michael S Packer, Ahmed H Badran, David I Bryson, and David R Liu. Programmable base editing of a* t to g* c in genomic dna without dna cleavage. Nature, 551 (7681):464–471, 2017.

Robert C Gentleman, Vincent J Carey, Douglas M Bates, Ben Bolstad, Marcel Dettling, Sandrine Dudoit, Byron Ellis, Laurent Gautier, Yongchao Ge, Jeff Gentry, et al. Bioconductor: open software development for computational biology and bioinformatics. Genome biology, 5(10):1–16, 2004.

Luke A Gilbert, Max A Horlbeck, Britt Adamson, Jacqueline E Villalta, Yuwen Chen, Evan H Whitehead, Carla Guimaraes, Barbara Panning, Hidde L Ploegh, Michael C Bassik, Lei S Qi, Martin Kampmann, and Weissman Jonathan S. Genome-scale crispr-mediated control of gene repression and activation. Cell, 159(3):647–661, Oct 2014.

F Hahne, A Lerch, and MB Stadler. Rbowtie: A r wrapper for bowtie and splicemap short read aligners. URL http://bioconductor.org/packages/release/bioc/html/Rbowtie.html, 2012.

Florian Hahne and Robert Ivanek. Visualizing genomic data using gviz and bioconductor. In Statistical genomics, pages 335–351. Springer, 2016.

Ruth E Hanna, Mudra Hegde, Christian R Fagre, Peter C DeWeirdt, Annabel K Sangree, Zsofia Szegletes, Audrey Griffith, Marissa N Feeley, Kendall R Sanson, Yossef Baidi, et al. Massively parallel assessment of human variants with base editor screens. Cell, 184(4):1064–1080, 2021.

Traver Hart, Kevin R Brown, Fabrice Sircoulomb, Robert Rottapel, and Jason Moffat. Measuring error rates in genomic perturbation screens: gold standards for human functional genomics. Molecular systems biology, 10(7):733, 2014.

Traver Hart, Megha Chandrashekhar, Michael Aregger, Zachary Steinhart, Kevin R Brown, Graham MacLeod, Monika Mis, Michal Zimmermann, Amelie Fradet-Turcotte, Song Sun, et al. High-resolution crispr screens reveal fitness genes and genotype-specific cancer liabilities. Cell, 163(6):1515–1526, 2015.

Traver Hart, Amy Hin Yan Tong, Katie Chan, Jolanda Van Leeuwen, Ashwin Seetharaman, Michael Aregger, Megha Chandrashekhar, Nicole Hustedt, Sahil Seth, Avery Noonan, et al. Evaluation and design of genome-wide crispr/spcas9 knockout screens. G3: Genes, Genomes, Genetics, 7(8):2719–2727, 2017a.

Traver Hart, Amy Hin Yan Tong, Katie Chan, Jolanda Van Leeuwen, Ashwin Seetharaman, Michael Aregger, Megha Chandrashekhar, Nicole Hustedt, Sahil Seth, Avery Noonan, et al. Evaluation and design of genome-wide crispr/spcas9 knockout screens. G3: Genes, Genomes, Genetics, 7(8):2719–2727, 2017b.

Wei He, Liang Zhang, Oscar D Villarreal, Rongjie Fu, Ella Bedford, Jingzhuang Dou, Anish Y Patel, Mark T Bedford, Xiaobing Shi, Taiping Chen, et al. De novo identification of essential protein domains from crispr-cas9 tiling-sgrna knockout screens. Nature communications, 10(1):1–10, 2019.

Florian Heigwer, Grainne Kerr, and Michael Boutros. E-crisp: fast crispr target site identification. Nature methods, 11(2):122–123, 2014.

Florian Heigwer, Tianzuo Zhan, Marco Breinig, Jan Winter, Dirk Brügemann, Svenja Leible, and Michael Boutros. Crispr library designer (cld): software for multispecies design of single guide rna libraries. Genome biology, 17(1):1–10, 2016.

Max A Horlbeck, Luke A Gilbert, Jacqueline E Villalta, Britt Adamson, Ryan A Pak, Yuwen Chen, Alexander P Fields, Chong Yon Park, Jacob E Corn, Martin Kampmann, et al. Compact and highly active next-generation libraries for crispr-mediated gene repression and activation. Elife, 5:e19760, 2016.

Patrick D Hsu, David A Scott, Joshua A Weinstein, F Ann Ran, Silvana Konermann, Vineeta Agarwala, Yinqing Li, Eli J Fine, Xuebing Wu, Ophir Shalem, et al. Dna targeting specificity of rna-guided cas9 nucleases. Nature biotechnology, 31(9):827, 2013a.

Patrick D Hsu, David A Scott, Joshua A Weinstein, F Ann Ran, Silvana Konermann, Vineeta Agarwala, Yinqing Li, Eli J Fine, Xuebing Wu, Ophir Shalem, et al. Dna targeting specificity of rna-guided cas9 nucleases. Nature biotechnology, 31(9):827–832, 2013b.

Johnny H Hu, Shannon M Miller, Maarten H Geurts, Weixin Tang, Liwei Chen, Ning Sun, Christina M Zeina, Xue Gao, Holly A Rees, Zhi Lin, et al. Evolved cas9 variants with broad pam compatibility and high dna specificity. Nature, 556(7699):57–63, 2018.

Wolfgang Huber, Vincent J Carey, Robert Gentleman, Simon Anders, Marc Carlson, Benilton S Carvalho, Hector Corrada Bravo, Sean Davis, Laurent Gatto, Thomas Girke, et al. Orchestrating high-throughput genomic analysis with bioconductor. Nature methods, 12(2):115–121, 2015.

Katharina Imkeller, Giulia Ambrosi, Michael Boutros, and Wolfgang Huber. gscreend: modelling asymmetric count ratios in crispr screens to decrease experiment size and improve phenotype detection. Genome biology, 21(1):1–13, 2020.

Donna Karolchik, Robert Baertsch, Mark Diekhans, Terrence S Furey, Angie Hinrichs, YT Lu, Krishna M Roskin, Matt Schwartz, Charles W Sugnet, Daryl J Thomas, et al. The ucsc genome browser database. Nucleic acids research, 31(1):51–54, 2003.

Eiru Kim and Traver Hart. Improved analysis of crispr fitness screens and reduced off-target effects with the bagel2 gene essentiality classifier. Genome medicine, 13(1):1–11, 2021.

Hui Kwon Kim, Seonwoo Min, Myungjae Song, Soobin Jung, Jae Woo Choi, Younggwang Kim, Sangeun Lee, Sungroh Yoon, and Hyongbum Henry Kim. Deep learning improves prediction of crispr–cpf1 guide rna activity. Nature biotechnology, 36(3):239, 2018.

Hui Kwon Kim, Younggwang Kim, Sungtae Lee, Seonwoo Min, Jung Yoon Bae, Jae Woo Choi, Jinman Park, Dongmin Jung, Sungroh Yoon, and Hyongbum Henry Kim. Spcas9 activity prediction by deepspcas9, a deep learning–based model with high generalization performance. Science advances, 5(11):eaax9249, 2019.

Luke W Koblan, Jordan L Doman, Christopher Wilson, Jonathan M Levy, Tristan Tay, Gregory A Newby, Juan Pablo Maianti, Aditya Raguram, and David R Liu. Improving cytidine and adenine base editors by expression optimization and ancestral reconstruction. Nature biotechnology, 36(9):843–846, 2018.

Alexis C Komor, Yongjoo B Kim, Michael S Packer, John A Zuris, and David R Liu. Programmable editing of a target base in genomic dna without double-stranded dna cleavage. Nature, 533(7603):420–424, 2016.

Silvana Konermann, Peter Lotfy, Nicholas J Brideau, Jennifer Oki, Maxim N Shokhirev, and Patrick D Hsu. Transcriptome engineering with rna-targeting type vi-d crispr effectors. Cell, 173(3):665–676, 2018.

Vasileios Konstantakos, Anastasios Nentidis, Anastasia Krithara, and Georgios Paliouras. Crispr–cas9 grna efficiency prediction: an overview of predictive tools and the role of deep learning. Nucleic Acids Research, 50(7):3616–3637, 2022.

Anshul Kundaje, Wouter Meuleman, Jason Ernst, Misha Bilenky, Angela Yen, Alireza Heravi-Moussavi, Pouya Kheradpour, Zhizhuo Zhang, Jianrong Wang, Michael J Ziller, et al. Integrative analysis of 111 reference human epigenomes. Nature, 518(7539):317–330, 2015.

Cem Kuscu, Sevki Arslan, Ritambhara Singh, Jeremy Thorpe, and Mazhar Adli. Genome-wide analysis reveals characteristics of off-target sites bound by the cas9 endonuclease. Nature biotechnology, 32(7):677–683, 2014.

Maurice Labuhn, Felix F Adams, Michelle Ng, Sabine Knoess, Axel Schambach, Emmanuelle M Charpentier, Adrian Schwarzer, Juan L Mateo, Jan-Henning Klusmann, and Dirk Heckl. Refined sgrna efficacy prediction improves large-and small-scale crispr–cas9 applications. Nucleic acids research, 46(3):1375–1385, 2018.

Kornel Labun, Tessa G Montague, James A Gagnon, Summer B Thyme, and Eivind Valen. Chopchop v2: a web tool for the next generation of crispr genome engineering. Nucleic acids research, 44(W1):W272–W276, 2016.

Kornel Labun, Xiaoge Guo, Alejandro Chavez, George Church, James A Gagnon, and Eivind Valen. Accurate analysis of genuine crispr editing events with amplican. Genome research, 29(5):843–847, 2019.

Ben Langmead, Cole Trapnell, Mihai Pop, and Steven L Salzberg. Ultrafast and memory-efficient alignment of short dna sequences to the human genome. Genome biology, 10(3):R25, 2009.

Michael Lawrence, Robert Gentleman, and Vincent Carey. rtracklayer: an r package for interfacing with genome browsers. Bioinformatics, 25(14):1841–1842, 2009.

Michael Lawrence, Wolfgang Huber, Hervé Pages, Patrick Aboyoun, Marc Carlson, Robert Gentleman, Martin T Morgan, and Vincent J Carey. Software for computing and annotating genomic ranges. PLoS computational biology, 9(8):e1003118, 2013.

Samuel Lessard, Laurent Francioli, Jessica Alfoldi, Jean-Claude Tardif, Patrick T Ellinor, Daniel G MacArthur, Guillaume Lettre, Stuart H Orkin, and Matthew C Canver. Human genetic variation alters crispr-cas9 on-and off-targeting specificity at therapeutically implicated loci. Proceedings of the National Academy of Sciences, page 201714640, 2017.

Heng Li and Richard Durbin. Fast and accurate short read alignment with burrows–wheeler transform. bioinformatics, 25(14):1754–1760, 2009.

Wei Li, Johannes Köster, Han Xu, Chen-Hao Chen, Tengfei Xiao, Jun S Liu, Myles Brown, and X Shirley Liu. Quality control, modeling, and visualization of crispr screens with mageck-vispr. Genome biology, 16(1):1–13, 2015.

Helen Lindsay, Alexa Burger, Berthin Biyong, Anastasia Felker, Christopher Hess, Jonas Zaugg, Elena Chiavacci, Carolin Anders, Martin Jinek, Christian Mosimann, et al. Crisprvariants charts the mutation spectrum of genome engineering experiments. Nature biotechnology, 34(7):701–702, 2016.

S John Liu, Max A Horlbeck, Seung Woo Cho, Harjus S Birk, Martina Malatesta, Daniel He, Frank J Attenello, Jacqueline E Villalta, Min Y Cho, Yuwen Chen, et al. Crispri-based genome-scale identification of functional long noncoding rna loci in human cells. Science, 355(6320):eaah7111, 2017.

Aaron Lun. basilisk: Freezing Python Dependencies Inside Bioconductor Packages, 2021. R package version 1.3.5.

Aaron McKenna and Jay Shendure. Flashfry: a fast and flexible tool for large-scale crispr target design. BMC biology, 16(1):1–6, 2018.

Joshua A Meier, Feng Zhang, and Neville E Sanjana. Guides: sgrna design for loss-of-function screens. Nature methods, 14(9):831–832, 2017.

Robin M Meyers, Jordan G Bryan, James M McFarland, Barbara A Weir, Ann E Sizemore, Han Xu, Neekesh V Dharia, Phillip G Montgomery, Glenn S Cowley, Sasha Pantel, et al. Computational correction of copy number effect improves specificity of crispr–cas9 essentiality screens in cancer cells. Nature genetics, 49(12):1779–1784, 2017.

Tessa G Montague, José M Cruz, James A Gagnon, George M Church, and Eivind Valen. Chopchop: a crispr/cas9 and talen web tool for genome editing. Nucleic acids research, 42(W1):W401–W407, 2014.

Miguel A Moreno-Mateos, Charles E Vejnar, Jean-Denis Beaudoin, Juan P Fernandez, Emily K Mis, Mustafa K Khokha, and Antonio J Giraldez. Crisprscan: designing highly efficient sgrnas for crispr-cas9 targeting in vivo. Nature methods, 12(10):982–988, 2015.

Hiroshi Nishimasu, Xi Shi, Soh Ishiguro, Linyi Gao, Seiichi Hirano, Sae Okazaki, Taichi Noda, Omar O Abudayyeh, Jonathan S Gootenberg, Hideto Mori, et al. Engineered crispr-cas9 nuclease with expanded targeting space. Science, 361(6408):1259–1262, 2018.

Hervé Pages, Patrick Aboyoun, Robert Gentleman, and Saikat DebRoy. Biostrings: String objects representing biological sequences, and matching algorithms. R package version, 2(0):10–18129, 2016.

Jeongbin Park, Sangsu Bae, and Jin-Soo Kim. Cas-designer: a web-based tool for choice of crispr-cas9 target sites. Bioinformatics, 31(24):4014–4016, 2015.

Vikram Pattanayak, Steven Lin, John P Guilinger, Enbo Ma, Jennifer A Doudna, and David R Liu. High-throughput profiling of off-target dna cleavage reveals rna-programmed cas9 nuclease specificity. Nature biotechnology, 31(9):839–843, 2013.

Alexendar R Perez, Yuri Pritykin, Joana A Vidigal, Sagar Chhangawala, Lee Zamparo, Christina S Leslie, and Andrea Ventura. Guidescan software for improved single and paired crispr guide rna design. Nature biotechnology, 35(4):347–349, 2017.

Katherine S Pollard, Melissa J Hubisz, Kate R Rosenbloom, and Adam Siepel. Detection of nonneutral substitution rates on mammalian phylogenies. Genome research, 20(1):110–121, 2010.

Aliaksandra Radzisheuskaya, Daria Shlyueva, Iris Müller, and Kristian Helin. Optimizing sgrna position markedly improves the efficiency of crispr/dcas9-mediated transcriptional repression. Nucleic acids research, 44(18):e141–e141, 2016.

Joseph M Replogle, Thomas M Norman, Albert Xu, Jeffrey A Hussmann, Jin Chen, J Zachery Cogan, Elliott J Meer, Jessica M Terry, Daniel P Riordan, Niranjan Srinivas, et al. Combinatorial single-cell crispr screens by direct guide rna capture and targeted sequencing. Nature biotechnology, 38(8):954–961, 2020.

Annabel K Sangree, Audrey L Griffith, Zsofia M Szegletesh, Priyanka Roy, Peter C DeWeirdt, Mudra Hegde, Abby V McGee, Ruth E Hanna, and John G Doench. Benchmarking of spcas9 variants enables deeper base editor screens of brca1 and bcl2. bioRxiv, 2021.

Kendall R Sanson, Ruth E Hanna, Mudra Hegde, Katherine F Donovan, Christine Strand, Meagan E Sullender, Emma W Vaimberg, Amy Goodale, David E Root, Federica Piccioni, et al. Optimized libraries for crispr-cas9 genetic screens with multiple modalities. Nature communications, 9(1):1–15, 2018.

Vivien AC Schoonenberg, Mitchel A Cole, Qiuming Yao, Claudio Macias-Treviño, Falak Sher, Patrick G Schupp, Matthew C Canver, Takahiro Maeda, Luca Pinello, and Daniel E Bauer. Crispro: identification of functional protein coding sequences based on genome editing dense mutagenesis. Genome biology, 19(1):1–19, 2018.

David A Scott and Feng Zhang. Implications of human genetic variation in crispr-based therapeutic genome editing. Nature medicine, 23(9):1095, 2017.

Sergey Shmakov, Omar O Abudayyeh, Kira S Makarova, Yuri I Wolf, Jonathan S Gootenberg, Ekaterina Semenova, Leonid Minakhin, Julia Joung, Silvana Konermann, Konstantin Severinov, et al. Discovery and functional characterization of diverse class 2 crispr-cas systems. Molecular cell, 60(3):385–397, 2015.

Arne H Smits, Frederik Ziebell, Gerard Joberty, Nico Zinn, William F Mueller, Sandra Clauder-Münster, Dirk Eberhard, Maria Fälth Savitski, Paola Grandi, Petra Jakob, et al. Biological plasticity rescues target activity in crispr knock outs. Nature methods, 16(11):1087–1093, 2019.

Manuel Stemmer, Thomas Thumberger, Maria del Sol Keyer, Joachim Wittbrodt, and Juan L Mateo. Cctop: an intuitive, flexible and reliable crispr/cas9 target prediction tool. PloS one, 10(4):e0124633, 2015.

Summer B Thyme, Laila Akhmetova, Tessa G Montague, Eivind Valen, and Alexander F Schier. Internal guide rna interactions interfere with cas9-mediated cleavage. Nature communications, 7:11750, 2016.

Konstantinos Tzelepis, Hiroko Koike-Yusa, Etienne De Braekeleer, Yilong Li, Emmanouil Metzakopian, Oliver M Dovey, Annalisa Mupo, Vera Grinkevich, Meng Li, Milena Mazan, et al. A crispr dropout screen identifies genetic vulnerabilities and therapeutic targets in acute myeloid leukemia. Cell reports, 17(4):1193–1205, 2016.

Brendan Veeneman, Ying Gao, Joy Grant, David Fruhling, James Ahn, Benedikt Bosbach, Jadwiga Bienkowska, Maximillian Follettie, Kim Arndt, Jeremy Myers, and Wenyan Zhong. Pincer: improved crispr/cas9 screening by efficient cleavage at conserved residues. Nucleic acids research, 48(17):9462–9477, 2020.

Russell T Walton, Kathleen A Christie, Madelynn N Whittaker, and Benjamin P Kleinstiver. Unconstrained genome targeting with near-pamless engineered crispr-cas9 variants. Science, 368(6488):290–296, 2020.

Binbin Wang, Mei Wang, Wubing Zhang, Tengfei Xiao, Chen-Hao Chen, Alexander Wu, Feizhen Wu, Nicole Traugh, Xiaoqing Wang, Ziyi Li, et al. Integrative analysis of pooled crispr genetic screens using mageckflute. Nature protocols, 14(3):756–780, 2019a.

Daqi Wang, Chengdong Zhang, Bei Wang, Bin Li, Qiang Wang, Dong Liu, Hongyan Wang, Yan Zhou, Leming Shi, Feng Lan, et al. Optimized crispr guide rna design for two high-fidelity cas9 variants by deep learning. Nature communications, 10(1):1–14, 2019b.

Guanqun Wang, Meijie Du, Jianbin Wang, and Ting F Zhu. Genetic variation may confound analysis of crispr-cas9 off-target mutations. Cell discovery, 4(1):18, 2018.

Tim Wang, Jenny J Wei, David M Sabatini, and Eric S Lander. Genetic screens in human cells using the crispr/cas9 system. Science, 343(6166):80–84, Jan 2014.

Tim Wang, Kivanç Birsoy, Nicholas W Hughes, Kevin M Krupczak, Yorick Post, Jenny J Wei, Eric S Lander, and David M Sabatini. Identification and characterization of essential genes in the human genome. Science, 350(6264):1096–1101, 2015.

Hans-Hermann Wessels, Alejandro Méndez-Mancilla, Xinyi Guo, Mateusz Legut, Zharko Daniloski, and Neville E Sanjana. Massively parallel cas13 screens reveal principles for guide rna design. Nature biotechnology, 38(6):722–727, 2020.

Xuebing Wu, David A Scott, Andrea J Kriz, Anthony C Chiu, Patrick D Hsu, Daniel B Dadon, Albert W Cheng, Alexandro E Trevino, Silvana Konermann, Sidi Chen, et al. Genome-wide binding of the crispr endonuclease cas9 in mammalian cells. Nature biotechnology, 32(7):670–676, 2014.

Lihua J Zhu, Benjamin R Holmes, Neil Aronin, and Michael H Brodsky. Crisprseek: a bioconductor package to identify target-specific guide rnas for crispr-cas9 genome-editing systems. PloS one, 9(9):e108424, 2014.

Lihua Julie Zhu, Michael Lawrence, Ankit Gupta, Alper Kucukural, Manuel Garber, Scot A Wolfe, et al. Guideseq: a bioconductor package to analyze guide-seq datasets for crispr-cas nucleases. BMC genomics, 18 (1):1–10, 2017.

