## Supplementary material for "The crisprVerse: a comprehensive Bioconductor ecosystem for the design of CRISPR guide RNAs across nucleases and technologies": Tutorial1_Installation

### Installing the crisprVerse and packages necessary for the tutorials

#### Installation

We show in this tutorial how to install the crisprVerse packages, as well as other packages necessary for some of the crisprVerse tutorials.

##### Requirements

The crisprVerse is supported for macOS, Linux and Windows machines. It requires R version >=4.2.1. Some of the third-party functionalities are not available for Windows machines (BWA alignment, and some of the scoring functions). To download and install R, follow the instructions on the R-project website.

##### Bioconductor versions

The Bioconductor project has 2 concurrent branches: `release` and `devel`. Currently (August 2022), the release branch is `3.15`, and the devel branch is `3.16`. Release versions are created twice a year. See the Bioconductor install page for more information regarding Bioconductor versions.

The crisprVerse ecosystem is currently available on the Bioconductor devel branch (`3.16`). Earlier versions of some of our packages are available on the release branch, but we do not recommend using the release branch as most of the functionalities described in the tutorials require updated functionalities only available on the devel branch.

##### Installing the core crisprVerse packages

Type in the following commands in an R session to install the core crisprVerse packages from the Bioconductor devel branch:

```
if (!requireNamespace("BiocManager", quietly = TRUE))
    install.packages("BiocManager")

BiocManager::install(version="devel")
BiocManager::install("crisprVerse")
```

This will install the following packages:

- crisprBase to specify and manipulate CRISPR nucleases.
- crisprBowtie to perform gRNA spacer sequence alignment with Bowtie.
- crisprScore to annotate gRNAs with on-target and off-target scores.
- crisprDesign to design and manipulate gRNAs with `GuideSet` objects.
- crisprScoreData to use pre-trained models for the `crisprScore` package.

The following command will load all of those packages in an R session:

```
library(crisprVerse)
```

You can check that all crisprVerse packages are up-to-date with `crisprVerse_update()`:

```
crisprVerse_update()
```

#### Installing data packages

The following genome data packages from Bioconductor are required for several of the tutorials:

```
if (!requireNamespace("BiocManager", quietly = TRUE))
    install.packages("BiocManager")

BiocManager::install(version="devel")
BiocManager::install("BSgenome.Mmusculus.UCSC.mm10")
BiocManager::install("BSgenome.Hsapiens.UCSC.hg38")
BiocManager::install("BSgenome.Hsapiens.UCSC.hg38.dbSNP151.major")
BiocManager::install("BSgenome.Hsapiens.UCSC.hg38.dbSNP151.minor")
```

The crisrpDesignData package is also required for most of the tutorials and can be installed directly from our GitHub page using the `devtools` package:

```
if (!requireNamespace("devtools", quietly = TRUE))
    install.packages("devtools")

devtools::install.packages("crisprVerse/crisprDesignData")
```

#### Installing optional packages

For maxOS and Linux users, the crisprBwa can be installed from Bioconductor using the following:

```
if (!requireNamespace("BiocManager", quietly = TRUE))
    install.packages("BiocManager")

BiocManager::install(version="devel")
BiocManager::install("crisprBwa")
```

The crisprViz package is currently under review at Bioconductor, but can be installed directly from GitHub:

```
if (!requireNamespace("devtools", quietly = TRUE))
    install.packages("devtools")

devtools::install.packages("crisprVerse/crisprViz")
```

### Reproducibility

```
sessionInfo()
```

```
## R version 4.2.1 (2022-06-23)
## Platform: x86_64-apple-darwin17.0 (64-bit)
## Running under: macOS Catalina 10.15.7
##
## Matrix products: default
## BLAS:   /Library/Frameworks/R.framework/Versions/4.2/Resources/lib/libRblas.0.dylib
## LAPACK: /Library/Frameworks/R.framework/Versions/4.2/Resources/lib/libRlapack.dylib
##
## locale:
## [1] en_US.UTF-8/en_US.UTF-8/en_US.UTF-8/C/en_US.UTF-8/en_US.UTF-8
##
```

```
## attached base packages:
## [1] stats     graphics  grDevices utils     datasets  methods   base
##
## other attached packages:
## [1] rmarkdown_2.15.2
##
## loaded via a namespace (and not attached):
##  [1] compiler_4.2.1  magrittr_2.0.3  fastmap_1.1.0   cli_3.3.0
##  [5] tools_4.2.1     htmltools_0.5.3 yaml_2.3.5      stringi_1.7.8
##  [9] knitr_1.40      stringr_1.4.1   xfun_0.32       digest_0.6.29
## [13] rlang_1.0.4     evaluate_0.16
```

### References
