## Supplementary material for "The crisprVerse: a comprehensive Bioconductor ecosystem for the design of CRISPR guide RNAs across nucleases and technologies": Tutorial16_Validating_Libraries

### Validating existing gRNA libraries

#### Introduction

In this vignette, we characterize a small mouse CRISPR knockout (CRISPRko) library that was designed to target tumor suppressors. The library was obtained from Addgene, and is stored in the folder `extdata` in the current directory.

#### Loading necessary packages

```
library(crisprDesign)
library(crisprBowtie)
library(crisprBase)
library(readxl)
library(BSgenome.Mmusculus.UCSC.mm10)
bsgenome <- BSgenome.Mmusculus.UCSC.mm10
```

We also load `crisprDesignData`, which is a data package containing already-processed Ensembl objects for gene annotation of human and mouse gRNAs:

```
library(crisprDesignData)
```

#### Reading in data

```
data <- read_excel("extdata/mtsg-grnas-readcounts.xlsx")
data <- as.data.frame(data)[,1:2]
colnames(data) <- c("ID", "spacer_20mer")

# Getting genes names:
data$gene_symbol <- sapply(strsplit(data$ID, split="_"), function(x)x[[1]])
head(data)
```

```
##             ID         spacer_20mer gene_symbol
## 1   Fat1_sg1 GGGCAGTGTTTCAAAATCCA        Fat1
## 2   Fat1_sg2 GGAACACGAGCCGTCAGCGG        Fat1
## 3   Fat1_sg3 GGATTTCTGTTCTGCATCAA        Fat1
## 4   Fat1_sg4 GGTCCCATCTGTTGCCTCCA        Fat1
## 5   Fat1_sg5 GTTTGGAGATCCACTCGATA        Fat1
## 6 Arid1b_sg1 GTACCCAGTGCAAGCTACAG      Arid1b
```

#### Building a `GuideSet` object

We first define the nuclease for the analysis. We here use the standard wildtype Cas9 (SpCas9) from the `crisprBase` package:

```
data(SpCas9, package="crisprBase")
crisprNuclease <- SpCas9
crisprNuclease
```

```
## Class: CrisprNuclease
##   Name: SpCas9
##   Target type: DNA
##   Metadata: list of length 1
##   PAMs: NGG, NAG, NGA
##   Weights: 1, 0.2593, 0.0694
##   Spacer length: 20
##   PAM side: 3prime
##     Distance from PAM: 0
##   Prototype protospacers: 5'--SSSSSSSSSSSSSSSSSSSS[NGG]--3', 5'--SSSSSSSSSSSSSSSSSSSS[NAG]--3', 5'--S
```

The default length of the spacer sequences is 20nt. This can be changed to a different length if needed, for instance 19nt:

```
# Not run
spacerLength(SpCas9) <- 19
```

We next need to define a bowtie index that we will use for alignment:

```
bowtie_index <- "/Users/fortinj2/crisprIndices/bowtie/mm10/mm10"
```

For instructions on how to build a Bowtie index from a given reference genome, see the genome index tutorial or the crisprBowtie page .

We first map the gRNAs to the reference genome with perfect match to obtain genomic coordinates of those gRNAs:

```
spacers <- unique(data$spacer_20mer)
aln <- runCrisprBowtie(spacers,
                       crisprNuclease=crisprNuclease,
                       bowtie_index=bowtie_index,
                       n_mismatches=0)
```

```
## [runCrisprBowtie] Searching for SpCas9 protospacers
```

```
head(aln)
```

```
##                  spacer          protospacer pam                chr  pam_site
## 1 GAAAACAGCCAAGGTTTGTA GAAAACAGCCAAGGTTTGTA CGG               chr8 106659780
## 2 GAAAACCCTGAAGTGCCCAC GAAAACCCTGAAGTGCCCAC GGG              chr17  29099403
## 3 GAAAACCCTGAAGTGCCCAC GAAAACCCTGAAGTGCCCAC GGG chr17_JH584267_alt   1798300
## 4 GAAAAGGGAAGACCAGCCCC GAAAAGGGAAGACCAGCCCC TGG               chr3 152219735
## 5 GAACCGACAAACAGTCCTGG GAACCGACAAACAGTCCTGG AGG              chr14  31255196
## 6 GAACCTAGATTTTGAGACAG GAACCTAGATTTTGAGACAG GGG              chr17  33952329
##   strand n_mismatches canonical
## 1      +            0      TRUE
## 2      +            0      TRUE
## 3      +            0      TRUE
## 4      +            0      TRUE
## 5      +            0      TRUE
## 6      +            0      TRUE
```

`n_mismatches=0` specifies that we require a perfect match between spacer and protospacer sequences (on-targets).

Non-targeting controls should not have any alignments to the genome, and some guides might have multiple alignments if they were not designed carefully. For such guides, that's OK, we can pick up pick one genomic coordinate for now, and the multiple alignments annotation will be handled later on.

We keep only alignments to the standard chromosomes:

```
chrs <- paste0("chr",c(1:22, "X", "Y"))
aln <- aln[aln$chr %in% chrs,,drop=FALSE]
```

We add the genomic coordinates to the data.frame:

```
wh <- match(data$spacer_20mer, aln$spacer)
data$chr <- aln$chr[wh]
data$pam_site <- aln$pam_site[wh]
data$pam <- aln$pam[wh]
data$strand <- aln$strand[wh]
head(data)
```

```
##            ID         spacer_20mer gene_symbol   chr pam_site pam strand
## 1   Fat1_sg1 GGGCAGTGTTTCAAAATCCA        Fat1  chr8 45023166 AGG      -
## 2   Fat1_sg2 GGAACACGAGCCGTCAGCGG        Fat1  chr8 45024178 TGG      +
## 3   Fat1_sg3 GGATTTCTGTTCTGCATCAA        Fat1  chr8 45013029 GGG      -
## 4   Fat1_sg4 GGTCCCATCTGTTGCCTCCA        Fat1  chr8 45013065 CGG      -
## 5   Fat1_sg5 GTTTGGAGATCCACTCGATA        Fat1  chr8 45010453 TGG      -
## 6 Arid1b_sg1 GTACCCAGTGCAAGCTACAG      Arid1b chr17  5291008 CGG      +
```

We can now build a proper `GuideSet` object in `crisprDesign` that will allow us to do (more) sophisticated analyses.

We need to filter out first guides that don't have a match to the genome:

```
data <- data[!is.na(data$pam_site),,drop=FALSE]
```

Finally, we create unique ids to identify the spacer sequences:

```
ids <- paste0("gRNA_", seq_len(nrow(data)))
head(ids)
```

```
## [1] "gRNA_1" "gRNA_2" "gRNA_3" "gRNA_4" "gRNA_5" "gRNA_6"
```

We are now ready to build the `GuideSet` object using the constructor function `GuideSet` from `crisprDesign`:

```
gs <- GuideSet(ids=ids,
               protospacers=data$spacer_20mer,
               pams=data$pam,
               pam_site=data$pam_site,
               seqnames=data$chr,
               strand=data$strand,
               CrisprNuclease=crisprNuclease,
               bsgenome=bsgenome)
gs$gene_symbol <- data$gene_symbol
```

The `GuideSet` object, and crisprDesign, provide rich functionalities to annotate and manipulate gRNAs. See the CRISPRko design tutorial to get an overview of the functionalities. For the rest of this tutorial, we only focus on characterizing the off-targets.

#### Off-target characterization

Having a `GuideSet` object, it is now a piece of cake to characterize the off-targets. We characterize off-targets using the bowtie aligner, with up to 3 mismatches between the spacer (gRNA) and protospacer (target DNA) sequences. The function `addSpacerAlignments` accomplishes that.

It has an optional argument `txObject` that can be used to provide gene model data to put the off-targets in a gene model context. We made such objects available for human and mouse in the crisprDesignData package (see `txdb_human` and `txdb_mouse`).

```
data(txdb_mouse, package="crisprDesignData")
txObject <- txdb_mouse
gs <- addSpacerAlignments(gs,
                          txObject=txObject,
                          aligner="bowtie",
                          aligner_index=bowtie_index,
                          bsgenome=bsgenome,
                          n_mismatches=2)
```

```
## [runCrisprBowtie] Using BSgenome.Mmusculus.UCSC.mm10
## [runCrisprBowtie] Searching for SpCas9 protospacers
```

The alignments are stored in a metadata column called `alignments`. See `?getSpacerAlignments` for more details about what the different columns are.

As an example, we can access the on- and off-target alignments of the first gRNA using the following commands:

```
aln <- gs$alignments[[1]]
aln
```

```
## GRanges object with 2 ranges and 14 metadata columns:
##          seqnames    ranges strand |               spacer          protospacer
##             <Rle> <IRanges>  <Rle> |       <DNAStringSet>       <DNAStringSet>
##   gRNA_1     chr8  45023166      - | GGGCAGTGTTTCAAAATCCA GGGCAGTGTTTCAAAATCCA
##   gRNA_1    chr12  80656576      - | GGGCAGTGTTTCAAAATCCA AGGCAGGGTTTCAAAATCCA
##                    pam  pam_site n_mismatches canonical  cut_site         cds
##          <DNAStringSet> <numeric>    <integer> <logical> <numeric> <character>
##   gRNA_1            AGG  45023166            0      TRUE  45023169        Fat1
##   gRNA_1            AGG  80656576            2      TRUE  80656579        <NA>
##             fiveUTRs   threeUTRs       exons     introns  intergenic
##          <character> <character> <character> <character> <character>
##   gRNA_1        <NA>        <NA>        Fat1        <NA>        <NA>
##   gRNA_1        <NA>        <NA>        <NA>     Slc39a9        <NA>
##          intergenic_distance
##                    <integer>
##   gRNA_1                <NA>
##   gRNA_1                <NA>
##   -------
##   seqinfo: 22 sequences (1 circular) from mm10 genome
```

We can also add CFD and MIT scores to the off-targets to characterize the likelihood of SpCas9 to cut at the off-targets:

```
gs <- addOffTargetScores(gs)
```

The scores range from 0 to 1, and a higher score indicates a higher probability of the off-target to occur.

### Session Info

```
sessionInfo()
```

```
## R version 4.2.1 (2022-06-23)
## Platform: x86_64-apple-darwin17.0 (64-bit)
## Running under: macOS Catalina 10.15.7
##
## Matrix products: default
## BLAS:   /Library/Frameworks/R.framework/Versions/4.2/Resources/lib/libRblas.0.dylib
## LAPACK: /Library/Frameworks/R.framework/Versions/4.2/Resources/lib/libRlapack.dylib
##
## locale:
## [1] en_US.UTF-8/en_US.UTF-8/en_US.UTF-8/C/en_US.UTF-8/en_US.UTF-8
##
## attached base packages:
## [1] stats4    stats     graphics  grDevices utils     datasets  methods
## [8] base
##
## other attached packages:
##  [1] readxl_1.4.1
##  [2] BSgenome.Hsapiens.UCSC.hg38.dbSNP151.minor_0.0.9999
##  [3] BSgenome.Hsapiens.UCSC.hg38.dbSNP151.major_0.0.9999
##  [4] BSgenome.Mmusculus.UCSC.mm10_1.4.3
##  [5] BSgenome.Hsapiens.UCSC.hg38_1.4.4
##  [6] BSgenome_1.65.2
##  [7] rtracklayer_1.57.0
##  [8] Biostrings_2.65.2
##  [9] XVector_0.37.0
## [10] GenomicRanges_1.49.1
## [11] GenomeInfoDb_1.33.5
## [12] IRanges_2.31.2
## [13] S4Vectors_0.35.1
## [14] crisprDesignData_0.99.17
## [15] crisprDesign_0.99.133
## [16] crisprScore_1.1.14
## [17] crisprScoreData_1.1.3
## [18] ExperimentHub_2.5.0
## [19] AnnotationHub_3.5.0
## [20] BiocFileCache_2.5.0
## [21] dbplyr_2.2.1
## [22] BiocGenerics_0.43.1
## [23] crisprBowtie_1.1.1
## [24] crisprBase_1.1.5
## [25] crisprVerse_0.99.8
## [26] rmarkdown_2.15.2
##
## loaded via a namespace (and not attached):
##  [1] rjson_0.2.21                  ellipsis_0.3.2
##  [3] Rbowtie_1.37.0                bit64_4.0.5
##  [5] lubridate_1.8.0               interactiveDisplayBase_1.35.0
##  [7] AnnotationDbi_1.59.1          fansi_1.0.3
##  [9] xml2_1.3.3                    codetools_0.2-18
## [11] cachem_1.0.6                  knitr_1.40
```

```
## [13] jsonlite_1.8.0            Rsamtools_2.13.4
## [15] png_0.1-7                 shiny_1.7.2
## [17] BiocManager_1.30.18       readr_2.1.2
## [19] compiler_4.2.1            httr_1.4.4
## [21] basilisk_1.9.2            assertthat_0.2.1
## [23] Matrix_1.4-1              fastmap_1.1.0
## [25] cli_3.3.0                 later_1.3.0
## [27] htmltools_0.5.3           prettyunits_1.1.1
## [29] tools_4.2.1               glue_1.6.2
## [31] GenomeInfoDbData_1.2.8    dplyr_1.0.9
## [33] rappdirs_0.3.3            tinytex_0.41
## [35] Rcpp_1.0.9                Biobase_2.57.1
## [37] cellranger_1.1.0          vctrs_0.4.1
## [39] crisprBwa_1.1.3           xfun_0.32
## [41] stringr_1.4.1             mime_0.12
## [43] lifecycle_1.0.1           restfulr_0.0.15
## [45] XML_3.99-0.10             zlibbioc_1.43.0
## [47] basilisk.utils_1.9.1      vroom_1.5.7
## [49] VariantAnnotation_1.43.3  hms_1.1.2
## [51] promises_1.2.0.1          MatrixGenerics_1.9.1
## [53] parallel_4.2.1            SummarizedExperiment_1.27.1
## [55] RMariaDB_1.2.2            yaml_2.3.5
## [57] curl_4.3.2                memoise_2.0.1
## [59] reticulate_1.25           biomaRt_2.53.2
## [61] stringi_1.7.8             RSQLite_2.2.16
## [63] BiocVersion_3.16.0        highr_0.9
## [65] BiocIO_1.7.1              randomForest_4.7-1.1
## [67] GenomicFeatures_1.49.6    filelock_1.0.2
## [69] BiocParallel_1.31.12      rlang_1.0.4
## [71] pkgconfig_2.0.3           matrixStats_0.62.0
## [73] bitops_1.0-7              evaluate_0.16
## [75] lattice_0.20-45           purrr_0.3.4
## [77] GenomicAlignments_1.33.1  bit_4.0.4
## [79] tidyselect_1.1.2          magrittr_2.0.3
## [81] R6_2.5.1                  generics_0.1.3
## [83] DelayedArray_0.23.1       DBI_1.1.3
## [85] pillar_1.8.1              KEGGREST_1.37.3
## [87] RCurl_1.98-1.8            tibble_3.1.8
## [89] dir.expiry_1.5.0          crayon_1.5.1
## [91] utf8_1.2.2                tzdb_0.3.0
## [93] progress_1.2.2            grid_4.2.1
## [95] blob_1.2.3                digest_0.6.29
## [97] xtable_1.8-4              httpuv_1.6.5
## [99] Rbwa_1.1.0
```
