## Supplementary material for "The crisprVerse: a comprehensive Bioconductor ecosystem for the design of CRISPR guide RNAs across nucleases and technologies": Tutorial11_PairedDesign

### Using crisprDesign to design paired gRNAs

#### Introduction

In this tutorial, we illustrate the main functionalities of `crisprDesign` for designing pairs of gRNAs.

#### Getting started

#### Installation

See the Installation tutorial to learn how to install the packages necessary for this tutorial: `crisprDesign`, `crisprDesignData`

##### Terminology

See the CRISPRko Cas9 design tutorial to get familiar with the terminology used throughout this tutorial.

##### Paired gRNA design overview

There are several applications that require the design of gRNA pairs:

1. Double nicking with CRISPR/Cas9 (Ran et al. et al. 2013)
2. Dual-promoter screening systems (Han et al. 2017)
3. Multiplexing gRNAs with enAsCas12a (DeWeirdt et al. et al. 2021)
4. Nanopore Cas9-targeted sequencing (nCATS) (Gilpatrick et al. 2020)

The `crisprDesign` package provides an infrastructure to store an annotate gRNA pairs via the `PaireGuideSet` object, which behaves very similarly to the `GuideSet` object used for unpaired gRNAs. We designed the functionalities for paired gRNAs with the aforementioned applications in mind.

In this tutorial, we will go through a simple example to illustrate the general concept behind paired gRNA design with `crisprDesign`.

#### A simple example: deleting a KRAS exon with a pair of gRNAs

We will show here how to design an optimal pair of Cas9 gRNAs flanking the second exon of the human gene KRAS (ENSG00000133703), with the goal of creating a deletion that will excise the exon.

We first start by loading the necessary packages:

```
library(crisprDesign)
library(crisprDesignData)
library(crisprBase)
library(BSgenome.Hsapiens.UCSC.hg38)
```

We will be designing gRNAs for the SpCas9 nuclease, which can be loaded from We load the `crisprBase` package (see the `crisprBase` vignette for instructions on how to create or load alternative nucleases):

```
data(SpCas9, package="crisprBase")
```

Let's get the genomic coordinates of the second exon. First, we obtain from `crisprDesignData` a `GRangesList` object that defines the genomic coordinates (hg38 genome) of human protein-coding genes:

```
data(txdb_human, package="crisprDesignData")
```

We then get the exonic coordinates of the canonical transcript ENST00000311936 using the function `queryTxObject` from `crisprDesign`:

```
exons <- queryTxObject(txObject=txdb_human,
                       featureType="exons",
                       queryColumn="tx_id",
                       queryValue="ENST00000311936")
exons
```

```
## GRanges object with 5 ranges and 14 metadata columns:
##            seqnames            ranges strand |           tx_id         gene_id
##               <Rle>         <IRanges>  <Rle> |     <character>     <character>
##   region_1    chr12 25250751-25250929      - | ENST00000311936 ENSG00000133703
##   region_2    chr12 25245274-25245395      - | ENST00000311936 ENSG00000133703
##   region_3    chr12 25227234-25227412      - | ENST00000311936 ENSG00000133703
##   region_4    chr12 25225614-25225773      - | ENST00000311936 ENSG00000133703
##   region_5    chr12 25205246-25209911      - | ENST00000311936 ENSG00000133703
##                 protein_id        tx_type gene_symbol         exon_id exon_rank
##                <character>    <character> <character>     <character> <integer>
##   region_1            <NA> protein_coding        KRAS ENSE00003903543         1
##   region_2 ENSP00000256078 protein_coding        KRAS ENSE00000936617         2
##   region_3 ENSP00000256078 protein_coding        KRAS ENSE00001719809         3
##   region_4 ENSP00000256078 protein_coding        KRAS ENSE00001644818         4
##   region_5 ENSP00000308495 protein_coding        KRAS ENSE00002456976         5
##            cds_start   cds_end  tx_start    tx_end   cds_len exon_start
##            <integer> <integer> <integer> <integer> <integer>  <integer>
##   region_1      <NA>      <NA>  25205246  25250929       567       <NA>
##   region_2  25245274  25245384  25205246  25250929       567       <NA>
##   region_3  25227234  25227412  25205246  25250929       567       <NA>
##   region_4  25225614  25225773  25205246  25250929       567       <NA>
##   region_5  25209795  25209911  25205246  25250929       567       <NA>
##             exon_end
##            <integer>
##   region_1      <NA>
##   region_2      <NA>
##   region_3      <NA>
##   region_4      <NA>
##   region_5      <NA>
##   -------
##   seqinfo: 25 sequences (1 circular) from hg38 genome
```

Finally, we select the second exon:

```
exon <- exons[exons$exon_rank==2]
names(exon) <- "exon_kras"
exon
```

```
## GRanges object with 1 range and 14 metadata columns:
##             seqnames            ranges strand |           tx_id         gene_id
##                <Rle>         <IRanges>  <Rle> |     <character>     <character>
```

```
##   exon_kras    chr12 25245274-25245395      - | ENST00000311936 ENSG00000133703
##                  protein_id        tx_type gene_symbol         exon_id
##                 <character>    <character> <character>     <character>
##   exon_kras ENSP00000256078 protein_coding        KRAS ENSE00000936617
##             exon_rank cds_start   cds_end  tx_start    tx_end   cds_len
##             <integer> <integer> <integer> <integer> <integer> <integer>
##   exon_kras         2  25245274  25245384  25205246  25250929       567
##             exon_start  exon_end
##              <integer> <integer>
##   exon_kras       <NA>      <NA>
##   -------
##   seqinfo: 25 sequences (1 circular) from hg38 genome
```

The exon is on chr12, and spans the region 25245274-25245395 (122 nucleotides in length). We aim to design gRNAs pairs for which one gRNA is located upstream of the exon, and another located downstream of the exon. To be able to find good gRNA candddates, let's define those regions to have 100 nucleotides on each side:

```
library(IRanges)
regionUpstream   <- IRanges::flank(exon, width=100, start=FALSE)
regionDownstream <- IRanges::flank(exon, width=100, start=TRUE)
names(regionUpstream) <- "upstreamTarget"
names(regionDownstream) <- "downstreamTarget"
```

Similar to the `findSpacers` function in `crisprDesign`, we will need to specify a `BSgenome` object containing the reference genome DNA sequences:

```
bsgenome <- BSgenome.Hsapiens.UCSC.hg38
```

We are now ready to find all candidate gRNA pairs:

```
pairs <- findSpacerPairs(x1=regionUpstream,
                         x2=regionDownstream,
                         bsgenome=bsgenome,
                         crisprNuclease=SpCas9)
```

The `x1` and `x2` arguments specify the genomic regions in which gRNAs at position 1 and position 2 should be targeting, respectively. The function finds all possible pair combinations between spacers found in the region specified by `x1` and spacers found in the region s pecified by `x2`. Let' first name our pairs:

```
names(pairs) <- paste0("pair_", seq_along(pairs))
```

Let's see what the results look like:

```
head(pairs, n=3)
```

```
## PairedGuideSet object with 3 pairs and 4 metadata columns:
##                     first           second | pamOrientation pamDistance
##                 <GuideSet>       <GuideSet> |    <character>   <numeric>
##   pair_1 chr12:25245201:+ chr12:25245397:- |             in         196
##   pair_2 chr12:25245215:- chr12:25245397:- |            rev         182
##   pair_3 chr12:25245233:+ chr12:25245397:- |             in         164
##          spacerDistance cutLength
##               <integer> <numeric>
##   pair_1            198       202
##   pair_2            163       182
##   pair_3            166       170
```

The returned object is a `PairedGuideSet`, which can be though of a list of two `GuideSet` objects. The first and second `GuideSet` store information about gRNAs at position 1 and position 2, respectively. They can be accessed using the `first` and `second` functions:

```
grnas1 <- first(pairs)
head(grnas1, n=3)
```

```
## GuideSet object with 3 ranges and 5 metadata columns:
##             seqnames    ranges strand |          protospacer            pam
##                <Rle> <IRanges>  <Rle> |       <DNAStringSet> <DNAStringSet>
##   spacer_1     chr12  25245201      + | GTAATAAGTACTCATGAAAA            TGG
##   spacer_2     chr12  25245215      - | CCATTCTTTGATACAGATAA            AGG
##   spacer_3     chr12  25245233      + | CCTTTATCTGTATCAAAGAA            TGG
##             pam_site  cut_site         region
##            <numeric> <numeric>    <character>
##   spacer_1  25245201  25245198 upstreamTarget
##   spacer_2  25245215  25245218 upstreamTarget
##   spacer_3  25245233  25245230 upstreamTarget
##   -------
##   seqinfo: 640 sequences (1 circular) from hg38 genome
##   crisprNuclease: SpCas9
```

and

```
grnas2 <- second(pairs)
head(grnas2, n=3)
```

```
## GuideSet object with 3 ranges and 5 metadata columns:
##             seqnames    ranges strand |          protospacer            pam
##                <Rle> <IRanges>  <Rle> |       <DNAStringSet> <DNAStringSet>
##   spacer_1     chr12  25245397      - | TTTTCATTATTTTTATTATA            AGG
##   spacer_1     chr12  25245397      - | TTTTCATTATTTTTATTATA            AGG
##   spacer_1     chr12  25245397      - | TTTTCATTATTTTTATTATA            AGG
##             pam_site  cut_site           region
##            <numeric> <numeric>      <character>
##   spacer_1  25245397  25245400 downstreamTarget
##   spacer_1  25245397  25245400 downstreamTarget
##   spacer_1  25245397  25245400 downstreamTarget
##   -------
##   seqinfo: 640 sequences (1 circular) from hg38 genome
##   crisprNuclease: SpCas9
```

The `pamOrientation` function returns the PAM orientation of the pairs:

```
head(pamOrientation(pairs))
```

```
## [1] "in"  "rev" "in"  "rev" "rev" "fwd"
```

and takes 4 different values: `in` (for PAM-in configuration), `out` (for PAM-out configuration), `fwd` (both gRNAs target the forward strand), and `rev` (both gRNAs target the reverse strand); see figure below for an illustration of the PAM orientations for the SpCas9 nuclease. The importance of the PAM orientation is application-specific. For Nanopore Cas9-targeted sequencing, PAM-in configuration is preferred. For double nicking with CRISPR/Cas9, PAM-out configuration is preferred. For applications using a dual-promoter system, no configuration is preferred.

The function `pamDistance` returns the distance between the PAM sites of the two gRNAs. The function `cutLength` returns the distance between the cut sites of the two gRNAs, and the function `spacerDistance` returns the distance between the two spacer sequences of the gRNAs.

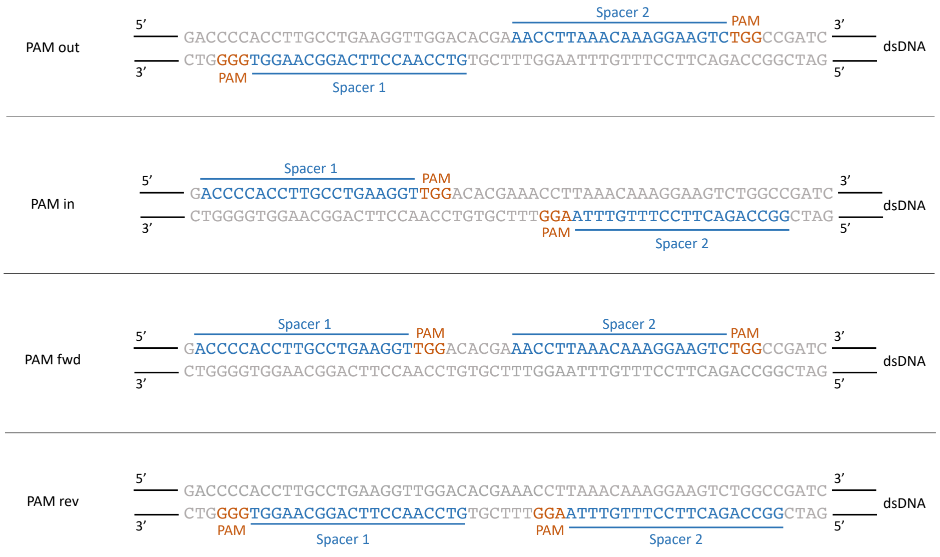


Figure 1: Different PAM orientations for Cas9 paired gRNAs

Most functionalities available for designing single gRNAs (`GuideSet` annotation functions described in this tutorial) work similarly for `PairedGuideSet` objects. This includes:

- `addSequenceFeatures`
- `addSpacerAlignments`
- `addGeneAnnotation`
- `addTssAnnotation`
- `addOnTargetScores`
- `addOffTargetScores`
- `addPamScores`
- `addSNPAnnotation`
- `addRestrictionEnzymes`
- `addCompositeScores`
- `addConservationScores`

Each function adds an annotation to the first and second `GuideSet` objects stored in the `PairedGuideSet`. Let's look at an example using `addSequenceFeatures`:

```
pairs <- addSequenceFeatures(pairs)
```

and let's look at the `GuideSet` in the first position:

```
head(first(pairs), n=3)
```

```
## GuideSet object with 3 ranges and 12 metadata columns:
##            seqnames    ranges strand |          protospacer            pam
##               <Rle> <IRanges>  <Rle> |       <DNAStringSet> <DNAStringSet>
##   spacer_1    chr12  25245201      + | GTAATAAGTACTCATGAAAA            TGG
##   spacer_2    chr12  25245215      - | CCATTCTTTGATACAGATAA            AGG
##   spacer_3    chr12  25245233      + | CCTTTATCTGTATCAAAGAA            TGG
##             pam_site  cut_site         region          coordID percentGC
##            <numeric> <numeric>    <character>      <character> <numeric>
##   spacer_1  25245201  25245198 upstreamTarget chr12_25245201_+        25
##   spacer_2  25245215  25245218 upstreamTarget chr12_25245215_-        30
```

```
##   spacer_3  25245233  25245230 upstreamTarget chr12_25245233_+        30
##               polyA     polyC     polyG     polyT startingGGGGG
##           <logical> <logical> <logical> <logical>     <logical>
##   spacer_1      TRUE     FALSE     FALSE     FALSE         FALSE
##   spacer_2     FALSE     FALSE     FALSE     FALSE         FALSE
##   spacer_3     FALSE     FALSE     FALSE     FALSE         FALSE
##   -------
##   seqinfo: 640 sequences (1 circular) from hg38 genome
##   crisprNuclease: SpCas9
```

This comes in handy to filter out pairs with unwanted sgRNA characteristics, e.g. sgRNA with polyT stretches:

```
good1 <- !first(pairs)$polyT
good2 <- !second(pairs)$polyT
pairs <- pairs[good1 & good2]
```

To select the final candidate pairs to excise the KRAS exon, we will filter out pairs with low predicted on-target activity using the DeepHF on-target activity score. We first add the score:

```
pairs <- addOnTargetScores(pairs, methods="deephf")
```

```
## [addOnTargetScores] Adding deephf scores.

## snapshotDate(): 2022-08-23

## see ?crisprScoreData and browseVignettes('crisprScoreData') for documentation

## loading from cache
```

and only keep pairs for which both gRNAs have a score greater than 0.5:

```
good1 <- first(pairs)$score_deephf>=0.5
good2 <- second(pairs)$score_deephf>=0.5
pairs <- pairs[good1 & good2]
```

This leaves us with 2 candidate pairs:

```
pairs
```

```
## PairedGuideSet object with 2 pairs and 4 metadata columns:
##                      first            second | pamOrientation pamDistance
##                 <GuideSet>       <GuideSet> |    <character>   <numeric>
##   pair_14 chr12:25245239:- chr12:25245472:- |            rev         233
##   pair_19 chr12:25245239:- chr12:25245475:- |            rev         236
##           spacerDistance cutLength
##                <integer> <numeric>
##   pair_14            214       233
##   pair_19            217       236
```

Finally, let's check for off-targets. We need to specify the path of the bowtie index that was generated from the human reference genome:

```
bowtie_index <- "/Users/fortinj2/crisprIndices/bowtie/hg38/hg38"
```

For instructions on how to build a Bowtie index from a given reference genome, see the genome index tutorial or the crisprBowtie page.

We are now ready to search for off-targets with up to 3 mismatches:

```
pairs <- addSpacerAlignments(pairs,
                             txObject=txdb_human,
                             aligner_index=bowtie_index,
```

```r
                              bsgenome=bsgenome,
                              n_mismatches=3)
```

```
## [runCrisprBowtie] Using BSgenome.Hsapiens.UCSC.hg38
## [runCrisprBowtie] Searching for SpCas9 protospacers
```

We are in luck, none of the spacer sequences has an off-target in the coding region of other genes:

```r
good1 <- first(pairs)$n1_c==0 & first(pairs)$n2_c==0 & first(pairs)$n3_c==0
good2 <- second(pairs)$n1_c==0 & second(pairs)$n2_c==0 & second(pairs)$n3_c==0
pairs <- pairs[good1 & good2]
pairs
```

```
## PairedGuideSet object with 2 pairs and 4 metadata columns:
##                       first            second | pamOrientation pamDistance
##                  <GuideSet>        <GuideSet> |    <character>   <numeric>
##   pair_14 chr12:25245239:- chr12:25245472:- |            rev         233
##   pair_19 chr12:25245239:- chr12:25245475:- |            rev         236
##           spacerDistance cutLength
##                <integer> <numeric>
##   pair_14            214       233
##   pair_19            217       236
```

One can get the spacer sequences using the `spacers` accessor function as usual:

```r
spacers(pairs)
```

```
## DataFrame with 2 rows and 2 columns
##                   first               second
##          <DNAStringSet>       <DNAStringSet>
## 1 AATATGCATATTACTGGTGC TTTGTATTAAAAGGTACTGG
## 2 AATATGCATATTACTGGTGC GAGTTTGTATTAAAAGGTAC
```

### Session Info

```r
sessionInfo()
```

```
## R version 4.2.1 (2022-06-23)
## Platform: x86_64-apple-darwin17.0 (64-bit)
## Running under: macOS Catalina 10.15.7
##
## Matrix products: default
## BLAS:   /Library/Frameworks/R.framework/Versions/4.2/Resources/lib/libRblas.0.dylib
## LAPACK: /Library/Frameworks/R.framework/Versions/4.2/Resources/lib/libRlapack.dylib
##
## locale:
## [1] en_US.UTF-8/en_US.UTF-8/en_US.UTF-8/C/en_US.UTF-8/en_US.UTF-8
##
## attached base packages:
## [1] stats4    stats     graphics  grDevices utils     datasets  methods
## [8] base
##
## other attached packages:
##  [1] BSgenome.Hsapiens.UCSC.hg38.dbSNP151.minor_0.0.9999
##  [2] BSgenome.Hsapiens.UCSC.hg38.dbSNP151.major_0.0.9999
##  [3] BSgenome.Mmusculus.UCSC.mm10_1.4.3
```

```
##  [4] BSgenome.Hsapiens.UCSC.hg38_1.4.4
##  [5] BSgenome_1.65.2
##  [6] rtracklayer_1.57.0
##  [7] Biostrings_2.65.2
##  [8] XVector_0.37.0
##  [9] GenomicRanges_1.49.1
## [10] GenomeInfoDb_1.33.5
## [11] IRanges_2.31.2
## [12] S4Vectors_0.35.1
## [13] crisprDesignData_0.99.17
## [14] crisprDesign_0.99.133
## [15] crisprScore_1.1.14
## [16] crisprScoreData_1.1.3
## [17] ExperimentHub_2.5.0
## [18] AnnotationHub_3.5.0
## [19] BiocFileCache_2.5.0
## [20] dbplyr_2.2.1
## [21] BiocGenerics_0.43.1
## [22] crisprBowtie_1.1.1
## [23] crisprBase_1.1.5
## [24] crisprVerse_0.99.8
## [25] rmarkdown_2.15.2
##
## loaded via a namespace (and not attached):
##  [1] rjson_0.2.21                  ellipsis_0.3.2
##  [3] Rbowtie_1.37.0                bit64_4.0.5
##  [5] lubridate_1.8.0               interactiveDisplayBase_1.35.0
##  [7] AnnotationDbi_1.59.1          fansi_1.0.3
##  [9] xml2_1.3.3                    codetools_0.2-18
## [11] cachem_1.0.6                  knitr_1.40
## [13] jsonlite_1.8.0                Rsamtools_2.13.4
## [15] png_0.1-7                     shiny_1.7.2
## [17] BiocManager_1.30.18           readr_2.1.2
## [19] compiler_4.2.1                httr_1.4.4
## [21] basilisk_1.9.2                assertthat_0.2.1
## [23] Matrix_1.4-1                  fastmap_1.1.0
## [25] cli_3.3.0                     later_1.3.0
## [27] htmltools_0.5.3               prettyunits_1.1.1
## [29] tools_4.2.1                   glue_1.6.2
## [31] GenomeInfoDbData_1.2.8        dplyr_1.0.9
## [33] rappdirs_0.3.3                tinytex_0.41
## [35] Rcpp_1.0.9                    Biobase_2.57.1
## [37] vctrs_0.4.1                   crisprBwa_1.1.3
## [39] xfun_0.32                     stringr_1.4.1
## [41] mime_0.12                     lifecycle_1.0.1
## [43] restfulr_0.0.15               XML_3.99-0.10
## [45] zlibbioc_1.43.0               basilisk.utils_1.9.1
## [47] vroom_1.5.7                   VariantAnnotation_1.43.3
## [49] hms_1.1.2                     promises_1.2.0.1
## [51] MatrixGenerics_1.9.1          parallel_4.2.1
## [53] SummarizedExperiment_1.27.1   RMariaDB_1.2.2
## [55] yaml_2.3.5                    curl_4.3.2
## [57] memoise_2.0.1                 reticulate_1.25
## [59] biomaRt_2.53.2                stringi_1.7.8
```

```
## [61] RSQLite_2.2.16             BiocVersion_3.16.0
## [63] highr_0.9                   BiocIO_1.7.1
## [65] randomForest_4.7-1.1        GenomicFeatures_1.49.6
## [67] filelock_1.0.2              BiocParallel_1.31.12
## [69] rlang_1.0.4                 pkgconfig_2.0.3
## [71] matrixStats_0.62.0          bitops_1.0-7
## [73] evaluate_0.16               lattice_0.20-45
## [75] purrr_0.3.4                 GenomicAlignments_1.33.1
## [77] bit_4.0.4                   tidyselect_1.1.2
## [79] magrittr_2.0.3              R6_2.5.1
## [81] generics_0.1.3              DelayedArray_0.23.1
## [83] DBI_1.1.3                   pillar_1.8.1
## [85] KEGGREST_1.37.3             RCurl_1.98-1.8
## [87] tibble_3.1.8                dir.expiry_1.5.0
## [89] crayon_1.5.1                utf8_1.2.2
## [91] tzdb_0.3.0                  progress_1.2.2
## [93] grid_4.2.1                  blob_1.2.3
## [95] digest_0.6.29               xtable_1.8-4
## [97] httpuv_1.6.5                Rbwa_1.1.0
```

### References


DeWeirdt, Peter C, Kendall R Sanson, Annabel K Sangree, Mudra Hegde, Ruth E Hanna, Marissa N Feeley, Audrey L Griffith, et al., et al. 2021. "Optimization of AsCas12a for Combinatorial Genetic Screens in Human Cells." *Nature Biotechnology* 39 (1): 94–104.

Gilpatrick, Timothy, Isac Lee, James E Graham, Etienne Raimondeau, Rebecca Bowen, Andrew Heron, Bradley Downs, Saraswati Sukumar, Fritz J Sedlazeck, and Winston Timp. 2020. "Targeted Nanopore Sequencing with Cas9-Guided Adapter Ligation." *Nature Biotechnology* 38 (4): 433–38.

Han, Kyuho, Edwin E Jeng, Gaelen T Hess, David W Morgens, Amy Li, and Michael C Bassik. 2017. "Synergistic Drug Combinations for Cancer Identified in a CRISPR Screen for Pairwise Genetic Interactions." *Nature Biotechnology* 35 (5): 463–74.

Ran, F Ann, Patrick D Hsu, Chie-Yu Lin, Jonathan S Gootenberg, Silvana Konermann, Alexandro E Trevino, David A Scott, et al., et al. 2013. "Double Nicking by RNA-Guided CRISPR Cas9 for Enhanced Genome Editing Specificity." *Cell* 154 (6): 1380–89.
