## Supplementary material for "The crisprVerse: a comprehensive Bioconductor ecosystem for the design of CRISPR guide RNAs across nucleases and technologies": Tutorial13_Optical_Pooled_Screening

### Using crisprDesign to design gRNAs for optical pooled screening (OPS)

#### Introduction

Optical pooled screening (OPS) combines image-based sequencing (in situ sequencing) of gRNAs and optical phenotyping on the same physical wells (Feldman et al. 2019). In such experiments, guide RNA (gRNA) spacer sequences are partially sequenced from the 5-prime end; the length of these truncated sequences, or barcodes, which corresponds to the number of sequencing cycles, is fixed and chosen by the experimentalist. From a gRNA design perspective, additional constraints are needed to ensure sufficient dissimilarity between the truncated barcodes for their identification during the analysis.

This tutorial will demonstrate how to design gRNAs for use in optical pooled screens, with emphasis on the constraints described above. Common gRNA design steps that are not specific to OPS are omitted in this tutorial (e.g. off-target search, or on-target activity prediction) here. Users can peruse through the list of available tutorials for more information regarding application-specific gRNA design rules.

#### Installation

See the Installation tutorial to learn how to install the packages necessary for this tutorial: `crisprDesign`, `crisprDesignData`

#### Terminology

See the CRISPRko design vignette to get familiar with the terminology used throughout this tutorial.

#### Design for optical pooled screening (OPS)

To illustrate the functionalities of `crisprDesign` for designing OPS libraries, we will design a small CRISPRko OPS library targeting 3 genes of the human RAS family: KRAS, HRAS, and NRAS. We will use the SpCas9 nuclease.

We will design gRNAs for an experiment that uses 8 in situ sequencing cycles:

```
n_cycles=8
```

##### Loading packages

Before we start, we first load the necessary packages for this tutorial:

```
library(crisprBase)
library(crisprDesign)
library(crisprDesignData)
library(BSgenome.Hsapiens.UCSC.hg38)
```

#### Creating the GuideSet

We begin by loading the SpCas9 `CrisprNuclease` object from the `crisprBase` package

```
data(SpCas9, package="crisprBase")
```

as well as data containing gene regions for the human genome:

```
data(txdb_human, package="crisprDesignData")
```

For more information on `txdb_human` and how to create similar gene annotation objects, see the Building a gene annotation object tutorial.

Next, we find the CDS coordinates for our genes using the `queryTxObject` function:

```
target_genes <- c("KRAS", "HRAS", "NRAS")
target_regions <- queryTxObject(txdb_human,
                                featureType="cds",
                                queryColumn="gene_symbol",
                                queryValue=target_genes)
```

then build our `GuideSet` with the `findSpacers` function:

```
gs <- findSpacers(target_regions,
                  crisprNuclease=SpCas9,
                  bsgenome=BSgenome.Hsapiens.UCSC.hg38)
```

As we will want to distinguish which gene each spacer targets, we will add `gene_symbol` and `gene_id` columns from `target_regions`.

```
gene_info <- target_regions[gs$region]
gs$gene_symbol <- gene_info$gene_symbol
gs$gene_id <- gene_info$gene_id
```

#### Adding OPS barcodes

We can add our OPS barcodes to the GuideSet with the `addOpsBarcodes` function. This function extracts the `n_cycles` nucleotides from the 5-prime end of our spacers and stores them in the `opsBarcode` column:

```
gs <- addOpsBarcodes(gs,
                     n_cycles=n_cycles)
head(gs$opsBarcode)
```

```
## DNAStringSet object of length 6:
##     width seq                                               names
## [1]     8 CCATGTGT                                          spacer_1
## [2]     8 TTGTATGG                                          spacer_2
## [3]     8 CCTTGTTA                                          spacer_3
## [4]     8 GATGGGAC                                          spacer_4
## [5]     8 TGATGGGA                                          spacer_5
## [6]     8 AGCAGTGA                                          spacer_6
```

#### Barcode distance matrix

We can pass our barcodes to the function `getBarcodeDistanceMatrix` to calculate the nucleotide distance between them. The `dist_method` argument determines the type of distance to calculate: `"hamming"`, which only considers substitutions (default) or `"levenstein"`, which also allows for insertions and deletions.

As a brief demonstration, let's look at the distances between the first few barcodes in our `GuideSet`. We set the `binarize` argument (more on this parameter later) to `FALSE` to show distances:

```r
barcodes <- gs$opsBarcode
dist <- getBarcodeDistanceMatrix(barcodes[1:5],
                                 binnarize=FALSE)
dist
```

```
## 5 x 5 sparse Matrix of class "dsCMatrix"
##          CCATGTGT TTGTATGG CCTTGTTA GATGGGAC TGATGGGA
## CCATGTGT        .        5        3        7        4
## TTGTATGG        5        .        6        8        5
## CCTTGTTA        3        6        .        6        5
## GATGGGAC        7        8        6        .        6
## TGATGGGA        4        5        5        6        .
```

Note that the output is a sparse matrix, so the barcodes along the diagonal (i.e., compared against themselves) return `.`, or a distance of zero. To compare one set of barcodes against another, we can pass the other set to the `targetBarcodes` argument (the former barcode set being passed to the `queryBarcodes` argument, which is compared against itself when `targetBarcodes` is `NULL`):

```r
dist <- getBarcodeDistanceMatrix(barcodes[1:5],
                                 targetBarcodes=barcodes[6:10],
                                 binnarize=FALSE)
dist
```

```
## 5 x 5 sparse Matrix of class "dgCMatrix"
##          AGCAGTGA CAGCAGTG AACTCAAC AAACTCAA TGCTGTTG
## CCATGTGT        5        7        7        7        5
## TTGTATGG        6        5        7        8        4
## CCTTGTTA        5        6        7        7        4
## GATGGGAC        7        6        5        6        7
## TGATGGGA        4        7        7        6        4
```

The question we are interested in with respect to barcode distances is whether this distance is sufficiently dissimilar for accurate identification of spacers during sequencing. This minimum distance edit (`min_dist_edit`) relies on the accuracy of various steps in the experiment. Suppose, as a conservative estimate, that we can expect no more than two edits per barcode in our example. A `min_dist_edit` of `3` should suffice. Setting the `binnarize` argument to `TRUE`, and passing our minimum distance edit value to `min_dist_edit` will binarize the output, flagging barcodes (with a value of `1`) that are too similar and should not both be included in our library:

```r
dist <- getBarcodeDistanceMatrix(barcodes[1:5],
                                 barcodes[6:10],
                                 binnarize=TRUE,
                                 min_dist_edit=3)
dist
```

```
## 5 x 5 sparse Matrix of class "dtCMatrix"
##          AGCAGTGA CAGCAGTG AACTCAAC AAACTCAA TGCTGTTG
## CCATGTGT        .        .        .        .        .
## TTGTATGG        .        .        .        .        .
## CCTTGTTA        .        .        .        .        .
## GATGGGAC        .        .        .        .        .
## TGATGGGA        .        .        .        .        .
```

Using this function with large sets of barcodes can be taxing on memory. To manage this, it is recommended to set `splitByChunks=TRUE` and specify the number of chunks with `n_chunks` (see `?getBarcodeDistanceMatrix`).

#### Designing OPS libraries

The `designOpsLibrary` function allows users to perform a complete end-to-end OPS library design. We will design our library with 4 gRNAs per gene using the `n_guides` and `gene_field` (to identify gRNAs by gene target) parameters. We will also use the same distance method and minimum distance edit parameters as in the example above.

NOTE: it is advised to first complete other steps in gRNA design (annotating, filtering, and ranking gRNAs in the `GuideSet`) prior to using this function; this will ensure the library contains the best gRNAs. As this example did not rank gRNAs, we are notified that rankings are assigned by the order in which gRNAs appear in our input.

```
df <- data.frame(ID=names(gs),
                 spacer=gs$protospacer,
                 opsBarcode=gs$opsBarcode,
                 gene_symbol=gs$gene_symbol)
opsLibrary <- designOpsLibrary(df,
                               n_guides=4,
                               gene_field="gene_symbol",
                               min_dist_edit=5,
                               dist_method="hamming")
```

```
## Since 'rank' column is not provided, using default order has ranking.
```

```
opsLibrary
```

```
##                   ID               spacer opsBarcode gene_symbol rank
## HRAS      spacer_73 ACTTGCAGCTCATGCAGCCG   ACTTGCAG        HRAS   10
## HRAS1     spacer_76 CTGAACCCTCCTGATGAGAG   CTGAACCC        HRAS   13
## HRAS2     spacer_79 CAGCCGGGGCCACTCTCATC   CAGCCGGG        HRAS   16
## HRAS3    spacer_131 TGGGTCACATGGGTCCCGGG   TGGGTCAC        HRAS   68
## KRAS     spacer_531 AAAGAAAAGATGAGCAAAGA   AAAGAAAA        KRAS    1
## KRAS1    spacer_533 TTCTCGAACTAATGTATAGA   TTCTCGAA        KRAS    3
## KRAS2    spacer_539 GGAGGATGCTTTTTATACAT   GGAGGATG        KRAS    9
## KRAS3    spacer_564 AACTCTTTTAATTTGTTCTC   AACTCTTT        KRAS   34
## NRAS       spacer_1 CCATGTGTGGTGATGTAACA   CCATGTGT        NRAS    1
## spacer_2   spacer_2 TTGTATGGGATTGCCATGTG   TTGTATGG        NRAS    2
## spacer_4   spacer_4 GATGGGACTCAGGGTTGTAT   GATGGGAC        NRAS    4
## NRAS1      spacer_6 AGCAGTGATGATGGGACTCA   AGCAGTGA        NRAS    6
```

#### Adding gRNAs to an existing OPS library

Suppose we later wish to add another gene target to our library, but also want to retain the gRNAs that are currently in our library. We can append these additional gRNAs with the `updateOpsLibrary` function. This function has the same parameters as `designOpsLibrary`, with an additional `opsLibrary` argument to which we pass our original OPS library.

To demonstrate, we will add the MRAS gene to our library. We first construct the `GuideSet` for MRAS:

```
target_region <- queryTxObject(txdb_human,
                               featureType="cds",
                               queryColumn="gene_symbol",
                               queryValue="MRAS")
gs_mras <- findSpacers(target_region,
                       crisprNuclease=SpCas9,
                       bsgenome=BSgenome.Hsapiens.UCSC.hg38)
gs_mras$gene_symbol <- "MRAS"
```

```r
gs_mras$gene_id <- "ENSG00000158186"
```

then add barcodes and construct the data.frame:

```r
## add OPS barcodes
gs_mras <- addOpsBarcodes(gs_mras,
                          n_cycles=n_cycles)

## construct data.frame
df_mras <- data.frame(ID=names(gs_mras),
                      spacer=gs_mras$protospacer,
                      opsBarcode=gs_mras$opsBarcode,
                      gene_symbol=gs_mras$gene_symbol)
```

which we then pass with our other parameters to updateOpsLibrary:

```r
opsLibrary <- updateOpsLibrary(opsLibrary,
                               df_mras,
                               n_guides=4,
                               gene_field="gene_symbol",
                               min_dist_edit=5,
                               dist_method="hamming")
```

```
## Since 'rank' column is not provided, using default order has ranking.
```

```r
opsLibrary
```

```
##                  ID               spacer opsBarcode gene_symbol rank
## HRAS      spacer_73 ACTTGCAGCTCATGCAGCCG   ACTTGCAG        HRAS   10
## HRAS1     spacer_76 CTGAACCCTCCTGATGAGAG   CTGAACCC        HRAS   13
## HRAS2     spacer_79 CAGCCGGGGCCACTCTCATC   CAGCCGGG        HRAS   16
## HRAS3    spacer_131 TGGGTCACATGGGTCCCGGG   TGGGTCAC        HRAS   68
## KRAS     spacer_531 AAAGAAAAGATGAGCAAAGA   AAAGAAAA        KRAS    1
## KRAS1    spacer_533 TTCTCGAACTAATGTATAGA   TTCTCGAA        KRAS    3
## KRAS2    spacer_539 GGAGGATGCTTTTTATACAT   GGAGGATG        KRAS    9
## KRAS3    spacer_564 AACTCTTTTAATTTGTTCTC   AACTCTTT        KRAS   34
## MRAS       spacer_4 GGGGAGGTTGTCACTGGGGA   GGGGAGGT        MRAS    4
## MRAS1     spacer_19 CCACCACCAGCTTGTATGTG   CCACCACC        MRAS   19
## MRAS2     spacer_34 CCCCACATACAAGCTGGTGG   CCCCACAT        MRAS   34
## MRAS3     spacer_37 CACATACAAGCTGGTGGTGG   CACATACA        MRAS   37
## NRAS       spacer_1 CCATGTGTGGTGATGTAACA   CCATGTGT        NRAS    1
## spacer_2   spacer_2 TTGTATGGGATTGCCATGTG   TTGTATGG        NRAS    2
## spacer_4   spacer_4 GATGGGACTCAGGGTTGTAT   GATGGGAC        NRAS    4
## NRAS1      spacer_6 AGCAGTGATGATGGGACTCA   AGCAGTGA        NRAS    6
```

### Session Info

```r
sessionInfo()
```

```
## R version 4.2.1 (2022-06-23)
## Platform: x86_64-apple-darwin17.0 (64-bit)
## Running under: macOS Catalina 10.15.7
##
## Matrix products: default
## BLAS:   /Library/Frameworks/R.framework/Versions/4.2/Resources/lib/libRblas.0.dylib
```

```
## LAPACK: /Library/Frameworks/R.framework/Versions/4.2/Resources/lib/libRlapack.dylib
##
## locale:
## [1] en_US.UTF-8/en_US.UTF-8/en_US.UTF-8/C/en_US.UTF-8/en_US.UTF-8
##
## attached base packages:
## [1] stats4    stats     graphics  grDevices utils     datasets  methods
## [8] base
##
## other attached packages:
##  [1] BSgenome.Hsapiens.UCSC.hg38.dbSNP151.minor_0.0.9999
##  [2] BSgenome.Hsapiens.UCSC.hg38.dbSNP151.major_0.0.9999
##  [3] BSgenome.Mmusculus.UCSC.mm10_1.4.3
##  [4] BSgenome.Hsapiens.UCSC.hg38_1.4.4
##  [5] BSgenome_1.65.2
##  [6] rtracklayer_1.57.0
##  [7] Biostrings_2.65.2
##  [8] XVector_0.37.0
##  [9] GenomicRanges_1.49.1
## [10] GenomeInfoDb_1.33.5
## [11] IRanges_2.31.2
## [12] S4Vectors_0.35.1
## [13] crisprDesignData_0.99.17
## [14] crisprDesign_0.99.133
## [15] crisprScore_1.1.14
## [16] crisprScoreData_1.1.3
## [17] ExperimentHub_2.5.0
## [18] AnnotationHub_3.5.0
## [19] BiocFileCache_2.5.0
## [20] dbplyr_2.2.1
## [21] BiocGenerics_0.43.1
## [22] crisprBowtie_1.1.1
## [23] crisprBase_1.1.5
## [24] crisprVerse_0.99.8
## [25] rmarkdown_2.15.2
##
## loaded via a namespace (and not attached):
##  [1] rjson_0.2.21                 ellipsis_0.3.2
##  [3] Rbowtie_1.37.0               bit64_4.0.5
##  [5] lubridate_1.8.0              interactiveDisplayBase_1.35.0
##  [7] AnnotationDbi_1.59.1         fansi_1.0.3
##  [9] xml2_1.3.3                   codetools_0.2-18
## [11] cachem_1.0.6                 knitr_1.40
## [13] jsonlite_1.8.0               Rsamtools_2.13.4
## [15] png_0.1-7                    shiny_1.7.2
## [17] BiocManager_1.30.18          readr_2.1.2
## [19] compiler_4.2.1               httr_1.4.4
## [21] basilisk_1.9.2               assertthat_0.2.1
## [23] Matrix_1.4-1                 fastmap_1.1.0
## [25] cli_3.3.0                    later_1.3.0
## [27] htmltools_0.5.3              prettyunits_1.1.1
## [29] tools_4.2.1                  glue_1.6.2
## [31] GenomeInfoDbData_1.2.8       dplyr_1.0.9
## [33] rappdirs_0.3.3               tinytex_0.41
```

```
## [35] Rcpp_1.0.9                    Biobase_2.57.1
## [37] vctrs_0.4.1                   crisprBwa_1.1.3
## [39] xfun_0.32                     stringr_1.4.1
## [41] mime_0.12                     lifecycle_1.0.1
## [43] restfulr_0.0.15               XML_3.99-0.10
## [45] zlibbioc_1.43.0               basilisk.utils_1.9.1
## [47] vroom_1.5.7                   VariantAnnotation_1.43.3
## [49] hms_1.1.2                     promises_1.2.0.1
## [51] MatrixGenerics_1.9.1          parallel_4.2.1
## [53] SummarizedExperiment_1.27.1   RMariaDB_1.2.2
## [55] yaml_2.3.5                    curl_4.3.2
## [57] memoise_2.0.1                 reticulate_1.25
## [59] biomaRt_2.53.2                stringi_1.7.8
## [61] RSQLite_2.2.16                BiocVersion_3.16.0
## [63] highr_0.9                     BiocIO_1.7.1
## [65] randomForest_4.7-1.1          GenomicFeatures_1.49.6
## [67] filelock_1.0.2                BiocParallel_1.31.12
## [69] rlang_1.0.4                   pkgconfig_2.0.3
## [71] matrixStats_0.62.0            bitops_1.0-7
## [73] evaluate_0.16                 lattice_0.20-45
## [75] purrr_0.3.4                   GenomicAlignments_1.33.1
## [77] bit_4.0.4                     tidyselect_1.1.2
## [79] magrittr_2.0.3                R6_2.5.1
## [81] generics_0.1.3                DelayedArray_0.23.1
## [83] DBI_1.1.3                     pillar_1.8.1
## [85] KEGGREST_1.37.3               RCurl_1.98-1.8
## [87] tibble_3.1.8                  dir.expiry_1.5.0
## [89] crayon_1.5.1                  utf8_1.2.2
## [91] tzdb_0.3.0                    progress_1.2.2
## [93] grid_4.2.1                    blob_1.2.3
## [95] digest_0.6.29                 xtable_1.8-4
## [97] httpuv_1.6.5                  Rbwa_1.1.0
```

### References


Feldman, David, Avtar Singh, Jonathan L Schmid-Burgk, Rebecca J Carlson, Anja Mezger, Anthony J Garrity, Feng Zhang, and Paul C Blainey. 2019. "Optical Pooled Screens in Human Cells." *Cell* 179 (3): 787–99.
