## Supplementary material for "The crisprVerse: a comprehensive Bioconductor ecosystem for the design of CRISPR guide RNAs across nucleases and technologies": Tutoria15_Major_and_Minor_Alleles

### Using crisprDesign to design gRNAs with minor and major alleles

#### Introduction

Genetic variants such as single nucleotide polymorphisms (SNPs) can be problematic in guide RNA (gRNA) design, as different alleles can result in unintended gRNA:DNA mismatches for on-targets that reduce gRNA efficacy. To circumvent this, it is advisable to generally avoid targeting sequences that contain variants. However, this may not always be possible, due to a small target window and/or few target options, or desirable, if, for example, a CRISPR application intends to target a pathogenic variant.

Functions in `crisprDesign` are well equipped to handle these cases. gRNAs overlapping SNPs can be identified with the `addSNPAnnotation` function, as documented in the CRISPRko design with Cas9 tutorial. Should the user wish to target region despite (or because of) the presence of variants, the user only needs to take care in the choice of `BSgenome` when constructing the `GuideSet` (an alternative option is to supply a custom target sequence; see the Working with a custom sequence tutorial for more information).

This tutorial covers use cases for `BSgenome` objects that store variants of the reference human genome (hg38) injected with major and minor alleles. It assumes the reader is familiar with constructing and using gene annotation objects (see the Building a gene annotation object tutorial) and `GuideSet` objects (see the CRISPRko design with Cas9 tutorial) so that the content may focus on the utility of the `BSgenome` variants discussed herein. Please consult the applicable tutorials if necessary.

Finally, it goes without saying that the user should be knowledgeable of the sequence(s), including possible variations in such, he or she is designing gRNAs for.

#### Installation

See the Installation tutorial to learn how to install the crisprVerse packages necessary for this tutorial.

#### Terminology

See the CRISPRko design tutorial to get familiar with the terminology used throughout this tutorial.

#### gRNA design

##### Loading packages

We first load the necessary packages for this tutorial:

```
library(crisprBase)
library(crisprDesign)
library(crisprDesignData)
library(BSgenome.Hsapiens.UCSC.hg38)
library(BSgenome.Hsapiens.UCSC.hg38.dbSNP151.major)
library(BSgenome.Hsapiens.UCSC.hg38.dbSNP151.minor)
```

We will also load the gene annotation model `txdb_human` from the `crisprDesignData` package:

```
data(txdb_human, package="crisprDesignData")
```

The `BSgenome.Hsapiens.UCSC.hg38.dbSNP151.major` and `BSgenome.Hsapiens.UCSC.hg38.dbSNP151.minor` packages are `BSgenome` packages that contain the major and minor alleles of the human reference genome hg38 based dbSNP151.For more information, type the following:

```
help(BSgenome.Hsapiens.UCSC.hg38.dbSNP151.major)
help(BSgenome.Hsapiens.UCSC.hg38.dbSNP151.minor)
```

#### Designing gRNAs for human (hg38) with injected major alleles

It is worth noting that the human reference genome sequence (GRCh38.p12), does not give the major allele (i.e. most common allele in a population) at all nucleotide locations. Indeed, given that it was historically constructed from a small set of human genomes, it contains minor alleles that were common across this set of human genomes.

For example, in the coding region (CDS) of the human gene SMC3, the reference genome contains the minor allele of the SNP rs2419565; the reference allele (A) frequency is 0.00577, and the alternative allele (G) frequency is 0.99423 as indicated in the table here here).

Designing gRNAs targeting the CDS of SMC3 with the SpCas9 nuclease returns one gRNA that overlaps this SNP. Below, we first construct a `GuideSet` object using the reference `BSgenome` object:

```
smc3 <- queryTxObject(txdb_human,
                      featureType="cds",
                      queryColumn="gene_symbol",
                      queryValue="SMC3")
gs_reference <- findSpacers(smc3,
                            crisprNuclease=SpCas9,
                            bsgenome=BSgenome.Hsapiens.UCSC.hg38)
```

and a `GuideSet` object with the `BSgenome` object that contains the major alleles:

```
gs_major <- findSpacers(smc3,
                        crisprNuclease=SpCas9,
                        bsgenome=BSgenome.Hsapiens.UCSC.hg38.dbSNP151.major)
```

Let's compare the protospacer sequences from both objects:

```
protospacers(gs_reference["spacer_199"])
## DNAStringSet object of length 1:
##     width seq                                               names
## [1]    20 TAGGGGTTACAAATCAATCA                              spacer_199
protospacers(gs_major["spacer_199"])
## DNAStringSet object of length 1:
##     width seq                                               names
## [1]    20 TAGGGGTTACAAATCGATCA                              spacer_199
```

The variant occurs in the seed sequence of this gRNA, 5 bases upstream of the `pam_site`, so a gRNA:DNA mismatch at this location is likely detrimental to its efficacy. Also, as this major allele occurs at >99% frequency, it may be more beneficial to design gRNAs in this example using the major allele genome contained in `BSgenome.Hsapiens.UCSC.hg38.dbSNP151.major`.

#### Designing gRNAs for human (hg38) with injected minor alleles

It may be desirable, in some applications, to target a genic sequence that contains a minor allele (i.e. less common allele) rather than the major or reference allele. For example, if a particular minor allele is pathogenic and the host cell has a single copy of that allele, the user may want to target that pathogenic variant and disrupt its behavior while leaving the other copy undisturbed.

As an example, using `BSgenome.Hsapiens.UCSC.hg38.dbSNP151.minor`, we can target the pathogenic minor allele (rs398122995) located in the human gene ALMS1:

```
alms1 <- queryTxObject(txdb_human, 'cds', 'gene_symbol', 'ALMS1')
gs_minor <- findSpacers(alms1,
                        crisprNuclease=SpCas9,
                        bsgenome=BSgenome.Hsapiens.UCSC.hg38.dbSNP151.minor)
gs_minor <- unique(gs_minor)
```

We also include, for comparison, the resulting `GuideSet` using the reference genome sequence:

```
gs_reference <- findSpacers(alms1,
                            crisprNuclease=SpCas9,
                            bsgenome=BSgenome.Hsapiens.UCSC.hg38)
gs_reference <- unique(gs_reference)
```

and compare the two versions of the gRNA:

```
gs_reference["spacer_615"]
```

```
## GuideSet object with 1 range and 5 metadata columns:
##              seqnames    ranges strand |          protospacer             pam
##                 <Rle> <IRanges>  <Rle> |       <DNAStringSet> <DNAStringSet>
##   spacer_615     chr2  73448425      - | TACTCTCTGGTAACTCTTGT            TGG
##               pam_site  cut_site      region
##              <numeric> <numeric> <character>
##   spacer_615  73448425  73448428    region_7
##   -------
##   seqinfo: 640 sequences (1 circular) from hg38 genome
##   crisprNuclease: SpCas9
```

```
gs_minor["spacer_612"]
```

```
## GuideSet object with 1 range and 5 metadata columns:
##              seqnames    ranges strand |          protospacer             pam
##                 <Rle> <IRanges>  <Rle> |       <DNAStringSet> <DNAStringSet>
##   spacer_612     chr2  73448425      - | TACTCTCTGGTAACTCTTGC            TGG
##               pam_site  cut_site      region
##              <numeric> <numeric> <character>
##   spacer_612  73448425  73448428    region_7
##   -------
##   seqinfo: 595 sequences (1 circular) from hg38 genome
##   crisprNuclease: SpCas9
```

The variant occurs 1 base upstream of the `pam_site`, and likely influences gRNA activity, that is, we can design a gRNA that targets the minor allele and has a much lower affinity for the reference, or major allele.

Note that while the two `GuideSet` objects differ only by their `BSgenome` object, we need to provide different indices to access protospacers at equivalent `pam_sites`. This is due to variants in one `BSgenome` (in this case theone with minor alleles) eliminating PAM sequences, that is, one of the Gs in NGG is changed to another base such that SpCas9 does not recognize it. This, where permissible, is also an effective way of ensuring gRNAs only target a specific sequence if that sequence contains the desired variant.
