## Supplementary material for "The crisprVerse: a comprehensive Bioconductor ecosystem for the design of CRISPR guide RNAs across nucleases and technologies": Tutorial12_CustomSequences

### Using crisprDesign to design gRNAs for custom sequences

#### Introduction

In this tutorial, we illustrate the main functionalities of `crisprDesign` for designing gRNAs for custom sequences. To design gRNAs for targets located in an organism genome, see the introductory CRISPRko tutorial.

#### Installation

See the Installation tutorial to learn how to install the packages necessary for this tutorial: `crisprDesign`, `crisprDesignData`

#### Use case: designing gRNAs against EGFP

Suppose we are engineering a human cell line to express the enhanced green fluorescent protein (EGFP) marker, and that we want to design gRNAs that knockout EGFP as experimental controls. Such control gRNAs should target EGFP with (1) high efficiency, and (2) should be specific to EGFP, that is, should not target the cell genome (human genome in this case). Supposed also that the cell line is also stably expressing SpCas9.

##### Loading necessary packages

We first start by loading the necessary packages:

```
library(Biostrings)
library(crisprBase)
library(crisprDesign)
library(crisprDesignData)
library(BSgenome.Hsapiens.UCSC.hg38)
```

##### Obtaining the DNA sequence

In the folder `data`, we have included a fasta file containing the DNA sequence of the EGFP marker. The sequence was obtained from the SnapGene website

We can read in the fasta file using the `readDNAStringSet` function from the package `Biostrings`:

```
dna <- Biostrings::readDNAStringSet("data/egfp.fa")
names(dna) <- "EGFP"
dna
```

```
## DNAStringSet object of length 1:
##     width seq                                               names
## [1]   720 ATGGTGAGCAAGGGCGAGGAGCT...GCATGGACGAGCTGTACAAGTAA EGFP
```

This could also be simply constructed from a regular string:

```
dna <- "ATGGTGAGCAAGGGCGAGGAGCTGTTCACCGGGGTGGTGCCCATCCTGGTCGAGCTGGACGGCGACGTAAACGGCCACAAGTTCAGCGTGTCCGG[illegible]
dna <- DNAStringSet(dna)
names(dna) <- "EGFP"
```

(Note that the function also accepts a simple string, which would be internally converted into a `DNAStringSet`). This is the custom sequence input that we will use to design gRNAs.

##### Constructing the `GuideSet` object:

Next, we design all possible SpCas9 gRNAs targeting EGFP. First, we load the SpCas9 object from the `crisprBase` package:

```
data(SpCas9, package="crisprBase")
```

and we design gRNAs using the function `findSpacers` from `crisprDesign`:

```
gs <- findSpacers(dna,
                  crisprNuclease=SpCas9)
head(gs)
```

```
## GuideSet object with 6 ranges and 5 metadata columns:
##           seqnames    ranges strand |          protospacer             pam
##              <Rle> <IRanges>  <Rle> |       <DNAStringSet> <DNAStringSet>
##   spacer_1     EGFP        30      + | AAGGGCGAGGAGCTGTTCAC             CGG
##   spacer_2     EGFP        31      + | AGGGCGAGGAGCTGTTCACC             GGG
##   spacer_3     EGFP        31      - | GACCAGGATGGGCACCACCC             CGG
##   spacer_4     EGFP        32      + | GGGCGAGGAGCTGTTCACCG             GGG
##   spacer_5     EGFP        35      + | CGAGGAGCTGTTCACCGGGG             TGG
##   spacer_6     EGFP        42      - | CCGTCCAGCTCGACCAGGAT             GGG
##            pam_site  cut_site      region
##           <numeric> <numeric> <character>
##   spacer_1        30        27        EGFP
##   spacer_2        31        28        EGFP
##   spacer_3        31        34        EGFP
##   spacer_4        32        29        EGFP
##   spacer_5        35        32        EGFP
##   spacer_6        42        45        EGFP
##   -------
##   seqinfo: 1 sequence from custom genome
##   crisprNuclease: SpCas9
```

The resulting output is a regular `GuideSet` object, and all functionalities described in the introductory CRISPRko tutorial can be applied here as well.

There are a few key differences to note with respect to a `GuideSet` object constructed using a reference genome. First, the name of the input DNA sequence (EGFP) is used as the chromosome name stored in the `seqnames` field. Second, the `pam_site` and `cut_site` coordinates are all relative to the first nucleotide of the custom DNA sequence. Finally, the `GuideSet` object stores the input sequence, which can be accessed using the function `customSequences`:

```
customSequences(gs)
```

```
## DNAStringSet object of length 1:
##     width seq                                               names
## [1]   720 ATGGTGAGCAAGGGCGAGGAGCT...GCATGGACGAGCTGTACAAGTAA EGFP
```

#### Finding off-targets in the human genome to find gRNAs specific to EGFP

Now that we have designed all possible gRNAs targeting EGFP, we will filter out gRNAs that have on- and off-targets located in the human genome. We will use the bowtie aligner to find targets, so we need to first specify the path of a bowtie index constructed on the human genome:

```
# Path of the hg38 bowtie index on my personal laptop:
bowtie_index <- "/Users/fortinj2/crisprIndices/bowtie/hg38/hg38"
```

For instructions on how to build a Bowtie index from a given reference genome, see the genome index tutorial.

To annotate off-targets with genomic context, for instance to know whether or not they are located in coding regions, we will also need a gene model object. We will use the gene model object `txdb_human` from `crisprDesignData`, which contains genomic coordinates of all human protein-coding genes. See the crisprDesignData package for more details.

```
data(txdb_human, package="crisprDesignData")
```

We are now ready to find all on- and off-targets using the `addSpacerAlignments` function from `crisprDesign`:

```
gs <- addSpacerAlignments(gs,
                          aligner="bowtie",
                          aligner_index=bowtie_index,
                          bsgenome=BSgenome.Hsapiens.UCSC.hg38,
                          n_mismatches=3,
                          txObject=txdb_human)
```

```
## [runCrisprBowtie] Using BSgenome.Hsapiens.UCSC.hg38
## [runCrisprBowtie] Searching for SpCas9 protospacers
```

```
gs
```

```
## GuideSet object with 119 ranges and 14 metadata columns:
##               seqnames    ranges strand |          protospacer           pam
##                  <Rle> <IRanges>  <Rle> |       <DNAStringSet> <DNAStringSet>
##     spacer_1      EGFP        30      + | AAGGGCGAGGAGCTGTTCAC           CGG
##     spacer_2      EGFP        31      + | AGGGCGAGGAGCTGTTCACC           GGG
##     spacer_3      EGFP        31      - | GACCAGGATGGGCACCACCC           CGG
##     spacer_4      EGFP        32      + | GGGCGAGGAGCTGTTCACCG           GGG
##     spacer_5      EGFP        35      + | CGAGGAGCTGTTCACCGGGG           TGG
##          ...       ...       ...    ... .                  ...           ...
##   spacer_115      EGFP       684      + | CTGGAGTTCGTGACCGCCGC           CGG
##   spacer_116      EGFP       685      + | TGGAGTTCGTGACCGCCGCC           GGG
##   spacer_117      EGFP       685      - | GTCCATGCCGAGAGTGATCC           CGG
##   spacer_118      EGFP       696      + | ACCGCCGCCGGGATCACTCT           CGG
##   spacer_119      EGFP       701      + | CGCCGGGATCACTCTCGGCA           TGG
##                pam_site  cut_site      region        n0        n1        n2
##               <numeric> <numeric> <character> <numeric> <numeric> <numeric>
##     spacer_1         30        27        EGFP         0         0         1
##     spacer_2         31        28        EGFP         0         0         2
##     spacer_3         31        34        EGFP         0         0         1
##     spacer_4         32        29        EGFP         0         0         0
##     spacer_5         35        32        EGFP         0         0         0
##          ...        ...       ...         ...       ...       ...       ...
##   spacer_115        684       681        EGFP         0         0         1
##   spacer_116        685       682        EGFP         0         0         0
##   spacer_117        685       688        EGFP         0         0         0
##   spacer_118        696       693        EGFP         0         0         0
```

```
##   spacer_119       701       698        EGFP         0         0         0
##                     n3      n0_c      n1_c      n2_c      n3_c
##              <numeric> <numeric> <numeric> <numeric> <numeric>
##     spacer_1         7         0         0         0         2
##     spacer_2         6         0         0         0         0
##     spacer_3        23         0         0         0         3
##     spacer_4         6         0         0         0         1
##     spacer_5         4         0         0         0         2
##          ...       ...       ...       ...       ...       ...
##   spacer_115         5         0         0         0         0
##   spacer_116         2         0         0         0         0
##   spacer_117         2         0         0         0         0
##   spacer_118         2         0         0         0         1
##   spacer_119         0         0         0         0         0
##                                                        alignments
##                                                     <GRangesList>
##     spacer_1   chr4:151052480:-,chr1:29350153:-,chrX:44232571:-,...
##     spacer_2  chr6:115711229:+,chr6:52987132:+,chr3:186748384:+,...
##     spacer_3 chr6:149167095:+,chr17:37844933:-,chr17:82019964:-,...
##     spacer_4 chr17:82484917:+,chr11:35064010:-,chr18:48539310:-,...
##     spacer_5    chr4:426282:-,chr19:16847037:+,chr19:14471687:+,...
##          ...                                                    ...
##   spacer_115  chr2:142494626:+,chr1:54281906:+,chr2:117131280:-,...
##   spacer_116                    chr4:139453357:-,chr17:82078529:+
##   spacer_117                      chr1:53536150:+,chr7:24540197:+
##   spacer_118                     chr18:77345044:+,chr3:51662810:-
##   spacer_119
##   -------
##   seqinfo: 1 sequence from custom genome
##   crisprNuclease: SpCas9
```

#### Predicting on-target activity

We also want to make sure to filter out gRNAs that are predicted to have poor on-target activity. To do so, we annotate gRNAs with the DeepHF on-target activity score:

```
gs <- addOnTargetScores(gs, methods="deephf")
```

```
## [addOnTargetScores] Adding deephf scores.

## snapshotDate(): 2022-08-23

## see ?crisprScoreData and browseVignettes('crisprScoreData') for documentation

## loading from cache
```

Finally, we characterize the spacer sequences using the `addSequenceFeatures` function from `crisprDesign`:

```
gs <- addSequenceFeatures(gs)
```

#### Final selection

For our use case, we will only retain gRNAs that do not map to the human genome (`n0=0`), don't have any 1 or 2-mismatch off-targets (`n1=0` and `n2=0`), and do not have 3-mismatch off-targets located in coding regions (`n3_c=0`):

```
gs <- gs[gs$n0==0 & gs$n1==0 & gs$n2==0 & gs$n3_c==0]
```

We also remove gRNAs that contain polyT sequences

```
gs <- gs[!gs$polyT,]
```

and only keep gRNAs that don't have extreme GC content:

```
gs <- gs[gs$percentGC>=20 & gs$percentGC<=80]
```

Finally, we rank gRNAs from the highest to the lowest on-target activity score:

```
gs <- gs[order(-gs$score_deephf)]
head(gs)
```

```
## GuideSet object with 6 ranges and 21 metadata columns:
##               seqnames    ranges strand |          protospacer           pam
##                  <Rle> <IRanges>  <Rle> |       <DNAStringSet> <DNAStringSet>
##    spacer_64      EGFP       359      + | GAAGTTCGAGGGCGACACCC           TGG
##   spacer_114      EGFP       682      - | CATGCCGAGAGTGATCCCGG           CGG
##    spacer_77      EGFP       446      - | CGGCCATGATATAGACGTTG           TGG
##    spacer_16      EGFP        98      + | CAAGTTCAGCGTGTCCGGCG           AGG
##    spacer_45      EGFP       229      - | GTCGTGCTGCTTCATGTGGT           CGG
##   spacer_107      EGFP       635      - | TGTGATCGCGCTTCTCGTTG           GGG
##                pam_site  cut_site      region        n0        n1        n2
##               <numeric> <numeric> <character> <numeric> <numeric> <numeric>
##    spacer_64        359       356        EGFP         0         0         0
##   spacer_114        682       685        EGFP         0         0         0
##    spacer_77        446       449        EGFP         0         0         0
##    spacer_16         98        95        EGFP         0         0         0
##    spacer_45        229       232        EGFP         0         0         0
##   spacer_107        635       638        EGFP         0         0         0
##                      n3      n0_c      n1_c      n2_c      n3_c
##               <numeric> <numeric> <numeric> <numeric> <numeric>
##    spacer_64          0         0         0         0         0
##   spacer_114          0         0         0         0         0
##    spacer_77          1         0         0         0         0
##    spacer_16          3         0         0         0         0
##    spacer_45          9         0         0         0         0
##   spacer_107          2         0         0         0         0
##                                                 alignments score_deephf
##                                              <GRangesList>    <numeric>
##    spacer_64                                                   0.716188
##   spacer_114                                                   0.700199
##    spacer_77                                   chr3:140247159:+     0.686111
##    spacer_16    chr5:132092368:+,chr6:42782483:+,chrX:97074134:+     0.670439
##    spacer_45 chr7:87066799:+,chr4:89369669:-,chrX:82007103:+,...     0.664180
##   spacer_107                   chr2:128980117:+,chr1:53729319:-     0.654151
##               percentGC     polyA     polyC     polyG     polyT startingGGGGG
##               <numeric> <logical> <logical> <logical> <logical>     <logical>
##    spacer_64         65     FALSE     FALSE     FALSE     FALSE         FALSE
##   spacer_114         65     FALSE     FALSE     FALSE     FALSE         FALSE
##    spacer_77         50     FALSE     FALSE     FALSE     FALSE         FALSE
##    spacer_16         65     FALSE     FALSE     FALSE     FALSE         FALSE
##    spacer_45         55     FALSE     FALSE     FALSE     FALSE         FALSE
##   spacer_107         55     FALSE     FALSE     FALSE     FALSE         FALSE
##   -------
##   seqinfo: 1 sequence from custom genome
```

```
##   crisprNuclease: SpCas9
```

Users can select the top gRNAs as their control gRNAs.

### Session Info

```
sessionInfo()
```

```
## R version 4.2.1 (2022-06-23)
## Platform: x86_64-apple-darwin17.0 (64-bit)
## Running under: macOS Catalina 10.15.7
##
## Matrix products: default
## BLAS:   /Library/Frameworks/R.framework/Versions/4.2/Resources/lib/libRblas.0.dylib
## LAPACK: /Library/Frameworks/R.framework/Versions/4.2/Resources/lib/libRlapack.dylib
##
## locale:
## [1] en_US.UTF-8/en_US.UTF-8/en_US.UTF-8/C/en_US.UTF-8/en_US.UTF-8
##
## attached base packages:
## [1] stats4    stats     graphics  grDevices utils     datasets  methods
## [8] base
##
## other attached packages:
##  [1] BSgenome.Mmusculus.UCSC.mm10_1.4.3 BSgenome.Hsapiens.UCSC.hg38_1.4.4
##  [3] BSgenome_1.65.2                    rtracklayer_1.57.0
##  [5] Biostrings_2.65.2                  XVector_0.37.0
##  [7] GenomicRanges_1.49.1               GenomeInfoDb_1.33.5
##  [9] IRanges_2.31.2                     S4Vectors_0.35.1
## [11] crisprDesignData_0.99.17           crisprDesign_0.99.133
## [13] crisprScore_1.1.14                 crisprScoreData_1.1.3
## [15] ExperimentHub_2.5.0                AnnotationHub_3.5.0
## [17] BiocFileCache_2.5.0                dbplyr_2.2.1
## [19] BiocGenerics_0.43.1                crisprBowtie_1.1.1
## [21] crisprBase_1.1.5                   crisprVerse_0.99.8
## [23] rmarkdown_2.15.2
##
## loaded via a namespace (and not attached):
##  [1] rjson_0.2.21                  ellipsis_0.3.2
##  [3] Rbowtie_1.37.0                bit64_4.0.5
##  [5] lubridate_1.8.0               interactiveDisplayBase_1.35.0
##  [7] AnnotationDbi_1.59.1          fansi_1.0.3
##  [9] xml2_1.3.3                    codetools_0.2-18
## [11] cachem_1.0.6                  knitr_1.40
## [13] jsonlite_1.8.0                Rsamtools_2.13.4
## [15] png_0.1-7                     shiny_1.7.2
## [17] BiocManager_1.30.18           readr_2.1.2
## [19] compiler_4.2.1                httr_1.4.4
## [21] basilisk_1.9.2                assertthat_0.2.1
## [23] Matrix_1.4-1                  fastmap_1.1.0
## [25] cli_3.3.0                     later_1.3.0
## [27] htmltools_0.5.3               prettyunits_1.1.1
## [29] tools_4.2.1                   glue_1.6.2
## [31] GenomeInfoDbData_1.2.8        dplyr_1.0.9
```

```
## [33] rappdirs_0.3.3              tinytex_0.41
## [35] Rcpp_1.0.9                  Biobase_2.57.1
## [37] vctrs_0.4.1                 crisprBwa_1.1.3
## [39] xfun_0.32                   stringr_1.4.1
## [41] mime_0.12                   lifecycle_1.0.1
## [43] restfulr_0.0.15             XML_3.99-0.10
## [45] zlibbioc_1.43.0             basilisk.utils_1.9.1
## [47] vroom_1.5.7                 VariantAnnotation_1.43.3
## [49] hms_1.1.2                   promises_1.2.0.1
## [51] MatrixGenerics_1.9.1        parallel_4.2.1
## [53] SummarizedExperiment_1.27.1 RMariaDB_1.2.2
## [55] yaml_2.3.5                  curl_4.3.2
## [57] memoise_2.0.1               reticulate_1.25
## [59] biomaRt_2.53.2              stringi_1.7.8
## [61] RSQLite_2.2.16              BiocVersion_3.16.0
## [63] highr_0.9                   BiocIO_1.7.1
## [65] randomForest_4.7-1.1        GenomicFeatures_1.49.6
## [67] filelock_1.0.2              BiocParallel_1.31.12
## [69] rlang_1.0.4                 pkgconfig_2.0.3
## [71] matrixStats_0.62.0          bitops_1.0-7
## [73] evaluate_0.16               lattice_0.20-45
## [75] purrr_0.3.4                 GenomicAlignments_1.33.1
## [77] bit_4.0.4                   tidyselect_1.1.2
## [79] magrittr_2.0.3              R6_2.5.1
## [81] generics_0.1.3              DelayedArray_0.23.1
## [83] DBI_1.1.3                   pillar_1.8.1
## [85] KEGGREST_1.37.3             RCurl_1.98-1.8
## [87] tibble_3.1.8                dir.expiry_1.5.0
## [89] crayon_1.5.1                utf8_1.2.2
## [91] tzdb_0.3.0                  progress_1.2.2
## [93] grid_4.2.1                  blob_1.2.3
## [95] digest_0.6.29               xtable_1.8-4
## [97] httpuv_1.6.5                Rbwa_1.1.0
```

### References
