## Supplementary material for "The crisprVerse: a comprehensive Bioconductor ecosystem for the design of CRISPR guide RNAs across nucleases and technologies": Tutorial14_Mapping_Across_Species

### Using crisprDesign to design gRNAs that map across species

#### Introduction

This tutorial describes how to design guide RNAs (gRNAs) that target homologous genes across multiple species using functions from the crisprDesign package. This strategy can be applied to any two (or more) species for which the genome sequence and gene model annotation is available.

#### Installation

See the Installation tutorial to learn how to install the packages necessary for this tutorial: `crisprDesign`, `crisprDesignData`

#### Terminology

See the CRISPRko design tutorial to get familiar with the terminology used throughout this tutorial.

#### Mapping gRNAs across species

##### Loading packages

We first load the necessary packages for this tutorial:

```
library(crisprBase)
library(crisprDesign)
library(crisprDesignData)
library(BSgenome.Hsapiens.UCSC.hg38)
library(BSgenome.Mmusculus.UCSC.mm10)
```

##### Creating the GuideSet

In this tutorial, we will design gRNAs using the SpCas9 nuclease that target both the human KRAS gene and its mouse ortholog Kras. There are multiple ways to go about this, which we describe in the following sections.

We first create a `GuideSet` object containing gRNAs targeting the coding sequence (CDS) of human KRAS. To do so, we start by loading the SpCas9 `CrisprNuclease` object from the `crisprBase` package:

```
data(SpCas9, package="crisprBase")
```

and then load data containing gene regions for the human genome from the `crisprDesignData` package, `txdb_human` (we will also load a similar object for the mouse genome, `txdb_mouse`):

```
data(txdb_human, package="crisprDesignData")
data(txdb_mouse, package="crisprDesignData")
```

For more information on `txdb_human` and `txdb_mouse` and how to create similar gene annotation objects, see the Building a gene annotation object tutorial tutorial.

Next, we find the coordinates for the CDS of KRAS using the `queryTxObject` function:

```
kras_human <- queryTxObject(txdb_human,
                            featureType="cds",
                            queryColumn="gene_symbol",
                            queryValue="KRAS")
```

and build our `GuideSet` object with the `findSpacers` function:

```
gs_human <- findSpacers(kras_human,
                        crisprNuclease=SpCas9,
                        bsgenome=BSgenome.Hsapiens.UCSC.hg38)
```

#### Mapping gRNAs across species via `intersect`

As a first strategy to find gRNAs that target both species, we first create a similar `GuideSet` targeting the mouse ortholog Kras:

```
kras_mouse <- queryTxObject(txdb_mouse,
                            featureType="cds",
                            queryColumn="gene_symbol",
                            queryValue="Kras")
gs_mouse <- findSpacers(kras_mouse,
                        crisprNuclease=SpCas9,
                        bsgenome=BSgenome.Mmusculus.UCSC.mm10)
```

Then, we find the common spacers between the two `GuideSet` objects using `intersect`

```
common_spacers <- intersect(spacers(gs_human),
                            spacers(gs_mouse))
length(common_spacers)
```

```
## [1] 18
```

There are 18 spacers that target KRAS in both species. We can filter each `GuideSet` object for this common spacer set:

```
results_human <- gs_human[spacers(gs_human) %in% common_spacers]
results_mouse <- gs_mouse[spacers(gs_mouse) %in% common_spacers]
```

Let's look at the results:

```
results_human
```

```
## GuideSet object with 33 ranges and 5 metadata columns:
##              seqnames    ranges strand |          protospacer             pam
##                 <Rle> <IRanges>  <Rle> |       <DNAStringSet> <DNAStringSet>
##     spacer_1    chr12  25209843      - | AAAGAAAAGATGAGCAAAGA             TGG
##     spacer_2    chr12  25209843      - | AAAGAAAAGATGAGCAAAGA             TGG
##     spacer_6    chr12  25215477      - | AGCAAAGAAGAAAAGACTCC             TGG
##     spacer_7    chr12  25215477      + | TTTTTAATTTTCACACAGCC             AGG
##     spacer_9    chr12  25215535      - | GGAGGATGCTTTTTATACAT             TGG
##          ...      ...       ...    ... .                  ...             ...
##    spacer_73    chr12  25227373      + | GTCGAGAATATCCAAGAGAC             AGG
##    spacer_92    chr12  25245330      + | CTGAATTAGCTGTATCGTCA             AGG
##    spacer_93    chr12  25245330      + | CTGAATTAGCTGTATCGTCA             AGG
```

```
##   spacer_94    chr12  25245330      + | CTGAATTAGCTGTATCGTCA            AGG
##   spacer_95    chr12  25245330      + | CTGAATTAGCTGTATCGTCA            AGG
##             pam_site  cut_site      region
##            <numeric> <numeric> <character>
##    spacer_1  25209843  25209846    region_8
##    spacer_2  25209843  25209846   region_10
##    spacer_6  25215477  25215480    region_4
##    spacer_7  25215477  25215474    region_4
##    spacer_9  25215535  25215538    region_4
##         ...       ...       ...         ...
##   spacer_73  25227373  25227370    region_6
##   spacer_92  25245330  25245327    region_1
##   spacer_93  25245330  25245327    region_5
##   spacer_94  25245330  25245327    region_9
##   spacer_95  25245330  25245327   region_11
##   -------
##   seqinfo: 640 sequences (1 circular) from hg38 genome
##   crisprNuclease: SpCas9
```

```
results_mouse
```

```
## GuideSet object with 32 ranges and 5 metadata columns:
##             seqnames    ranges strand |          protospacer            pam
##                <Rle> <IRanges>  <Rle> |       <DNAStringSet> <DNAStringSet>
##     spacer_7     chr6 145220656      - | AAAGAAAAGATGAGCAAAGA            TGG
##     spacer_8     chr6 145220656      - | AAAGAAAAGATGAGCAAAGA            TGG
##    spacer_14     chr6 145225114      - | AGCAAAGAAGAAAAGACTCC            TGG
##    spacer_15     chr6 145225114      + | TTTTTAATTTTCACACAGCC            AGG
##    spacer_17     chr6 145225172      - | GGAGGATGCTTTTTATACAT            TGG
##          ...      ...       ...    ... .                  ...            ...
##    spacer_88     chr6 145234387      + | GTCGAGAATATCCAAGAGAC            AGG
##    spacer_89     chr6 145234387      + | GTCGAGAATATCCAAGAGAC            AGG
##   spacer_108     chr6 145246751      + | CTGAATTAGCTGTATCGTCA            AGG
##   spacer_109     chr6 145246751      + | CTGAATTAGCTGTATCGTCA            AGG
##   spacer_110     chr6 145246751      + | CTGAATTAGCTGTATCGTCA            AGG
##              pam_site  cut_site      region
##             <numeric> <numeric> <character>
##     spacer_7 145220656 145220659    region_4
##     spacer_8 145220656 145220659    region_6
##    spacer_14 145225114 145225117   region_10
##    spacer_15 145225114 145225111   region_10
##    spacer_17 145225172 145225175   region_10
##          ...       ...       ...         ...
##    spacer_88 145234387 145234384    region_2
##    spacer_89 145234387 145234384    region_8
##   spacer_108 145246751 145246748    region_1
##   spacer_109 145246751 145246748    region_5
##   spacer_110 145246751 145246748    region_7
##   -------
##   seqinfo: 239 sequences (1 circular) from mm10 genome
##   crisprNuclease: SpCas9
```

This simple approach, however, has some drawbacks. It requires gRNAs to have perfect sequence matching, which, while perhaps acceptable for targets having many gRNA choices, may be too restrictive for those applications that have fewer choices and may need tolerate mismatches in the target genes. Also, and more notably, we now have multiple `GuideSet` objects to maintain in the process of selecting candidate gRNAs (see CRISPRko design with Cas9)–essentially twice the work.

#### Mapping gRNAs across species via `addSpacerAlignments`

To avoid the drawbacks of the above strategy, we can use the `addSpacerAlignments` function on our human KRAS `GuideSet` to append alignment annotation of the **mouse** genome.

For this example, we will use the bowtie aligner, and weneed to specify a bowtie index for the mouse genome:

```
# Path of the mm10 bowtie index on my personal laptop:
bowtie_index_mouse <- "/Users/fortinj2/crisprIndices/bowtie/mm10/mm10"
```

For instructions on how to build a Bowtie index from a given reference genome, see the genome index tutorial.

We will also search up to 1 mismatch and pass the gene model object `txdb_mouse` to the `txObject` argument, so the alignments will be annotated with genomic context and we can determine which of our spacers map to the CDS of Kras.

As we will also want to search for off-targets in the human genome in a later step, we can ensure these results are not overwritten by setting the `colname` argument to a non-default value, such as `alignments_mouse`.

```
results_human <- addSpacerAlignments(gs_human,
                                     aligner="bowtie",
                                     aligner_index=bowtie_index_mouse,
                                     bsgenome=BSgenome.Mmusculus.UCSC.mm10,
                                     txObject=txdb_mouse,
                                     colname="alignments_mouse",
                                     n_mismatches=1)
```

```
## [runCrisprBowtie] Using BSgenome.Mmusculus.UCSC.mm10
## [runCrisprBowtie] Searching for SpCas9 protospacers
```

```
results_human
```

```
## GuideSet object with 115 ranges and 10 metadata columns:
##               seqnames    ranges strand |          protospacer            pam
##                  <Rle> <IRanges>  <Rle> |       <DNAStringSet> <DNAStringSet>
##     spacer_1     chr12  25209843      - | AAAGAAAAGATGAGCAAAGA            TGG
##     spacer_2     chr12  25209843      - | AAAGAAAAGATGAGCAAAGA            TGG
##     spacer_3     chr12  25209896      + | TTCTCGAACTAATGTATAGA            AGG
##     spacer_4     chr12  25209896      + | TTCTCGAACTAATGTATAGA            AGG
##     spacer_5     chr12  25215438      - | AAATGCATTATAATGTAATC            TGG
##          ...       ...       ...    ... .                  ...            ...
##   spacer_111     chr12  25245358      - | GAATATAAACTTGTGGTAGT            TGG
##   spacer_112     chr12  25245365      - | AATGACTGAATATAAACTTG            TGG
##   spacer_113     chr12  25245365      - | AATGACTGAATATAAACTTG            TGG
##   spacer_114     chr12  25245365      - | AATGACTGAATATAAACTTG            TGG
##   spacer_115     chr12  25245365      - | AATGACTGAATATAAACTTG            TGG
##                 pam_site  cut_site      region        n0        n1      n0_c
##                <numeric> <numeric> <character> <numeric> <numeric> <numeric>
##     spacer_1  25209843  25209846    region_8         1         0         1
##     spacer_2  25209843  25209846   region_10         1         0         1
##     spacer_3  25209896  25209893    region_8         0         1         0
##     spacer_4  25209896  25209893   region_10         0         1         0
##     spacer_5  25215438  25215441    region_4         0         1         0
##          ...       ...       ...         ...       ...       ...       ...
##   spacer_111  25245358  25245361   region_11         0         0         0
```

```
##   spacer_112  25245365  25245368    region_1         0          1          0
##   spacer_113  25245365  25245368    region_5         0          1          0
##   spacer_114  25245365  25245368    region_9         0          1          0
##   spacer_115  25245365  25245368   region_11         0          1          0
##                    n1_c alignments_mouse
##               <numeric>    <GRangesList>
##     spacer_1          0 chr6:145220656:-
##     spacer_2          0 chr6:145220656:-
##     spacer_3          1 chr6:145220709:+
##     spacer_4          1 chr6:145220709:+
##     spacer_5          1 chr6:145225075:-
##          ...        ...              ...
##   spacer_111          0
##   spacer_112          1 chr6:145246786:-
##   spacer_113          1 chr6:145246786:-
##   spacer_114          1 chr6:145246786:-
##   spacer_115          1 chr6:145246786:-
##   -------
##   seqinfo: 640 sequences (1 circular) from hg38 genome
##   crisprNuclease: SpCas9
```

Our results are stored in the `alignments_mouse` column. We can access these alignments with the `alignments` function and by specifying the `columnName`:

```
alignments(results_human, columnName="alignments_mouse")
```

```
## GRanges object with 95 ranges and 14 metadata columns:
##               seqnames    ranges strand |               spacer
##                  <Rle> <IRanges>  <Rle> |       <DNAStringSet>
##     spacer_1      chr6 145220656      - | AAAGAAAAGATGAGCAAAGA
##     spacer_2      chr6 145220656      - | AAAGAAAAGATGAGCAAAGA
##     spacer_3      chr6 145220709      + | TTCTCGAACTAATGTATAGA
##     spacer_4      chr6 145220709      + | TTCTCGAACTAATGTATAGA
##     spacer_5      chr6 145225075      - | AAATGCATTATAATGTAATC
##          ...       ...       ...    ... .                  ...
##   spacer_107      chr6 145246773      - | AAACTTGTGGTAGTTGGAGC
##   spacer_112      chr6 145246786      - | AATGACTGAATATAAACTTG
##   spacer_113      chr6 145246786      - | AATGACTGAATATAAACTTG
##   spacer_114      chr6 145246786      - | AATGACTGAATATAAACTTG
##   spacer_115      chr6 145246786      - | AATGACTGAATATAAACTTG
##                        protospacer            pam  pam_site n_mismatches
##                     <DNAStringSet> <DNAStringSet> <numeric>    <integer>
##     spacer_1 AAAGAAAAGATGAGCAAAGA            TGG 145220656            0
##     spacer_2 AAAGAAAAGATGAGCAAAGA            TGG 145220656            0
##     spacer_3 TTCTCGGACTAATGTATAGA            AGG 145220709            1
##     spacer_4 TTCTCGGACTAATGTATAGA            AGG 145220709            1
##     spacer_5 AAATGCGTTATAATGTAATC            TGG 145225075            1
##          ...                  ...            ...       ...          ...
##   spacer_107 AAACTTGTGGTGGTTGGAGC            TGG 145246773            1
##   spacer_112 AATGACTGAGTATAAACTTG            TGG 145246786            1
##   spacer_113 AATGACTGAGTATAAACTTG            TGG 145246786            1
##   spacer_114 AATGACTGAGTATAAACTTG            TGG 145246786            1
##   spacer_115 AATGACTGAGTATAAACTTG            TGG 145246786            1
##               canonical  cut_site         cds     fiveUTRs    threeUTRs
##               <logical> <numeric> <character> <character> <character>
```

```
##     spacer_1      TRUE 145220659        Kras        <NA>        Kras
##     spacer_2      TRUE 145220659        Kras        <NA>        Kras
##     spacer_3      TRUE 145220706        Kras        <NA>        Kras
##     spacer_4      TRUE 145220706        Kras        <NA>        Kras
##     spacer_5      TRUE 145225078        Kras        <NA>        <NA>
##          ...       ...       ...         ...         ...         ...
##   spacer_107      TRUE 145246776        Kras        <NA>        <NA>
##   spacer_112      TRUE 145246789        Kras        <NA>        <NA>
##   spacer_113      TRUE 145246789        Kras        <NA>        <NA>
##   spacer_114      TRUE 145246789        Kras        <NA>        <NA>
##   spacer_115      TRUE 145246789        Kras        <NA>        <NA>
##                      exons     introns  intergenic intergenic_distance
##                <character> <character> <character>           <integer>
##     spacer_1         Kras        <NA>        <NA>                <NA>
##     spacer_2         Kras        <NA>        <NA>                <NA>
##     spacer_3         Kras        <NA>        <NA>                <NA>
##     spacer_4         Kras        <NA>        <NA>                <NA>
##     spacer_5         Kras        Kras        <NA>                <NA>
##          ...          ...         ...         ...                 ...
##   spacer_107         Kras        <NA>        <NA>                <NA>
##   spacer_112         Kras        <NA>        <NA>                <NA>
##   spacer_113         Kras        <NA>        <NA>                <NA>
##   spacer_114         Kras        <NA>        <NA>                <NA>
##   spacer_115         Kras        <NA>        <NA>                <NA>
##   -------
##   seqinfo: 22 sequences (1 circular) from mm10 genome
```

With these data, we can filter our gRNAs for those that target both orthologs (and we have off-target annotation for the mouse genome).

```
aln <- alignments(results_human, columnName="alignments_mouse")
cds_targets <- aln$cds
aln <- aln[!is.na(cds_targets) & cds_targets == "Kras"]
targets_Kras <- unique(names(aln))
results_human <- results_human[targets_Kras]
```

Adding alignments for the human genome (or any other genome) will overwrite the summary columns in `results_human` (`n0`, `n0_c`, `n1`, and `n1_c`) unless we set `addSummary=FALSE` in `addSpacerAlignments`. We should also take care to ensure the column name for our alignments annotation remains unique so it will not be overwritten. Here, we add alignment annotation for the human genome, but overwrite the mouse alignment summary columns (see the warning message below).

```
# Path of the hg38 bowtie index on my personal laptop:
bowtie_index_human <- "/Users/fortinj2/crisprIndices/bowtie/hg38/hg38"

results_human <- addSpacerAlignments(results_human,
                                     aligner="bowtie",
                                     aligner_index=bowtie_index_human,
                                     bsgenome=BSgenome.Hsapiens.UCSC.hg38,
                                     txObject=txdb_human,
                                     colname="alignments_human",
                                     n_mismatches=1)
```

```
## [runCrisprBowtie] Using BSgenome.Hsapiens.UCSC.hg38
## [runCrisprBowtie] Searching for SpCas9 protospacers
```

```
## Warning in .addAlignmentsSummary(guideSet = object, aln = aln, addSummary =
## addSummary, : Overwriting existing alignments summary. To avoid overwriting, set
## addSummary=FALSE.
```

```
results_human
```

```
## GuideSet object with 89 ranges and 11 metadata columns:
##              seqnames    ranges strand |          protospacer            pam
##                 <Rle> <IRanges>  <Rle> |       <DNAStringSet> <DNAStringSet>
##     spacer_1    chr12  25209843      - | AAAGAAAAGATGAGCAAAGA            TGG
##     spacer_2    chr12  25209843      - | AAAGAAAAGATGAGCAAAGA            TGG
##     spacer_3    chr12  25209896      + | TTCTCGAACTAATGTATAGA            AGG
##     spacer_4    chr12  25209896      + | TTCTCGAACTAATGTATAGA            AGG
##     spacer_5    chr12  25215438      - | AAATGCATTATAATGTAATC            TGG
##          ...      ...       ...    ... .                  ...            ...
##   spacer_107    chr12  25245352      - | AAACTTGTGGTAGTTGGAGC            TGG
##   spacer_112    chr12  25245365      - | AATGACTGAATATAAACTTG            TGG
##   spacer_113    chr12  25245365      - | AATGACTGAATATAAACTTG            TGG
##   spacer_114    chr12  25245365      - | AATGACTGAATATAAACTTG            TGG
##   spacer_115    chr12  25245365      - | AATGACTGAATATAAACTTG            TGG
##               pam_site  cut_site      region        n0        n1      n0_c
##              <numeric> <numeric> <character> <numeric> <numeric> <numeric>
##     spacer_1  25209843  25209846    region_8         1         2         1
##     spacer_2  25209843  25209846   region_10         1         2         1
##     spacer_3  25209896  25209893    region_8         1         1         1
##     spacer_4  25209896  25209893   region_10         1         1         1
##     spacer_5  25215438  25215441    region_4         1         0         1
##          ...       ...       ...         ...       ...       ...       ...
##   spacer_107  25245352  25245355   region_11         1         1         1
##   spacer_112  25245365  25245368    region_1         2         0         1
##   spacer_113  25245365  25245368    region_5         2         0         1
##   spacer_114  25245365  25245368    region_9         2         0         1
##   spacer_115  25245365  25245368   region_11         2         0         1
##                   n1_c alignments_mouse
##              <numeric>    <GRangesList>
##     spacer_1         0 chr6:145220656:-
##     spacer_2         0 chr6:145220656:-
##     spacer_3         0 chr6:145220709:+
##     spacer_4         0 chr6:145220709:+
##     spacer_5         0 chr6:145225075:-
##          ...       ...              ...
##   spacer_107         0 chr6:145246773:-
##   spacer_112         0 chr6:145246786:-
##   spacer_113         0 chr6:145246786:-
##   spacer_114         0 chr6:145246786:-
##   spacer_115         0 chr6:145246786:-
##                                             alignments_human
##                                                <GRangesList>
##     spacer_1 chr12:25209843:-,chr6:54771089:+,chr5:4348033:+
##     spacer_2 chr12:25209843:-,chr6:54771089:+,chr5:4348033:+
##     spacer_3               chr12:25209896:+,chr6:54771050:-
##     spacer_4               chr12:25209896:+,chr6:54771050:-
##     spacer_5                               chr12:25215438:-
##          ...                                            ...
##   spacer_107               chr12:25245352:-,chr6:54770615:+
```

```
##   spacer_112               chr12:25245365:-,chr6:54770602:+
##   spacer_113               chr12:25245365:-,chr6:54770602:+
##   spacer_114               chr12:25245365:-,chr6:54770602:+
##   spacer_115               chr12:25245365:-,chr6:54770602:+
##   -------
##   seqinfo: 640 sequences (1 circular) from hg38 genome
##   crisprNuclease: SpCas9
```

### Session Info

```
sessionInfo()
```

```
## R version 4.2.1 (2022-06-23)
## Platform: x86_64-apple-darwin17.0 (64-bit)
## Running under: macOS Catalina 10.15.7
##
## Matrix products: default
## BLAS:   /Library/Frameworks/R.framework/Versions/4.2/Resources/lib/libRblas.0.dylib
## LAPACK: /Library/Frameworks/R.framework/Versions/4.2/Resources/lib/libRlapack.dylib
##
## locale:
## [1] en_US.UTF-8/en_US.UTF-8/en_US.UTF-8/C/en_US.UTF-8/en_US.UTF-8
##
## attached base packages:
## [1] stats4    stats     graphics  grDevices utils     datasets  methods
## [8] base
##
## other attached packages:
##  [1] BSgenome.Mmusculus.UCSC.mm10_1.4.3 BSgenome.Hsapiens.UCSC.hg38_1.4.4
##  [3] BSgenome_1.65.2                    rtracklayer_1.57.0
##  [5] Biostrings_2.65.2                  XVector_0.37.0
##  [7] GenomicRanges_1.49.1               GenomeInfoDb_1.33.5
##  [9] IRanges_2.31.2                     S4Vectors_0.35.1
## [11] crisprDesignData_0.99.17           crisprDesign_0.99.133
## [13] crisprScore_1.1.14                 crisprScoreData_1.1.3
## [15] ExperimentHub_2.5.0                AnnotationHub_3.5.0
## [17] BiocFileCache_2.5.0                dbplyr_2.2.1
## [19] BiocGenerics_0.43.1                crisprBowtie_1.1.1
## [21] crisprBase_1.1.5                   crisprVerse_0.99.8
## [23] rmarkdown_2.15.2
##
## loaded via a namespace (and not attached):
##  [1] rjson_0.2.21                ellipsis_0.3.2
##  [3] Rbowtie_1.37.0              bit64_4.0.5
##  [5] lubridate_1.8.0             interactiveDisplayBase_1.35.0
##  [7] AnnotationDbi_1.59.1        fansi_1.0.3
##  [9] xml2_1.3.3                  codetools_0.2-18
## [11] cachem_1.0.6                knitr_1.40
## [13] jsonlite_1.8.0              Rsamtools_2.13.4
## [15] png_0.1-7                   shiny_1.7.2
## [17] BiocManager_1.30.18         readr_2.1.2
## [19] compiler_4.2.1              httr_1.4.4
## [21] basilisk_1.9.2              assertthat_0.2.1
```

```
## [23] Matrix_1.4-1                 fastmap_1.1.0
## [25] cli_3.3.0                    later_1.3.0
## [27] htmltools_0.5.3              prettyunits_1.1.1
## [29] tools_4.2.1                  glue_1.6.2
## [31] GenomeInfoDbData_1.2.8       dplyr_1.0.9
## [33] rappdirs_0.3.3               tinytex_0.41
## [35] Rcpp_1.0.9                   Biobase_2.57.1
## [37] vctrs_0.4.1                  crisprBwa_1.1.3
## [39] xfun_0.32                    stringr_1.4.1
## [41] mime_0.12                    lifecycle_1.0.1
## [43] restfulr_0.0.15              XML_3.99-0.10
## [45] zlibbioc_1.43.0              basilisk.utils_1.9.1
## [47] vroom_1.5.7                  VariantAnnotation_1.43.3
## [49] hms_1.1.2                    promises_1.2.0.1
## [51] MatrixGenerics_1.9.1         parallel_4.2.1
## [53] SummarizedExperiment_1.27.1  RMariaDB_1.2.2
## [55] yaml_2.3.5                   curl_4.3.2
## [57] memoise_2.0.1                reticulate_1.25
## [59] biomaRt_2.53.2               stringi_1.7.8
## [61] RSQLite_2.2.16               BiocVersion_3.16.0
## [63] highr_0.9                    BiocIO_1.7.1
## [65] randomForest_4.7-1.1         GenomicFeatures_1.49.6
## [67] filelock_1.0.2               BiocParallel_1.31.12
## [69] rlang_1.0.4                  pkgconfig_2.0.3
## [71] matrixStats_0.62.0           bitops_1.0-7
## [73] evaluate_0.16                lattice_0.20-45
## [75] purrr_0.3.4                  GenomicAlignments_1.33.1
## [77] bit_4.0.4                    tidyselect_1.1.2
## [79] magrittr_2.0.3               R6_2.5.1
## [81] generics_0.1.3               DelayedArray_0.23.1
## [83] DBI_1.1.3                    pillar_1.8.1
## [85] KEGGREST_1.37.3              RCurl_1.98-1.8
## [87] tibble_3.1.8                 dir.expiry_1.5.0
## [89] crayon_1.5.1                 utf8_1.2.2
## [91] tzdb_0.3.0                   progress_1.2.2
## [93] grid_4.2.1                   blob_1.2.3
## [95] digest_0.6.29                xtable_1.8-4
## [97] httpuv_1.6.5                 Rbwa_1.1.0
```
