## Supplementary material for "The crisprVerse: a comprehensive Bioconductor ecosystem for the design of CRISPR guide RNAs across nucleases and technologies": Tutorial5_CRISPRko_Design_Cas9

### Design gRNAs for CRISPRko with the SpCas9 nuclease

#### Introduction

In this tutorial, we illustrate the main functionalities of crisprDesign, the central package of the crisprVerse ecosystem, by designing CRISPR/Cas9 gRNAs targeting the coding sequence of the human gene KRAS. Most steps described in the tutorial are applicable to any genomic target.

#### Some terminology before we get started

Before we start designing gRNAs, we first introduce some terminology that will be useful throughout this and subsequent tutorials. CRISPR nucleases require two binding components for cleavage. First, the nuclease needs to recognize a constant nucleotide motif in the target DNA called the protospacer adjacent motif (PAM) sequence. Second, the gRNA, which guides the nuclease to the target sequence, needs to bind to a complementary sequence adjacent to the PAM sequence, called the **protospacer** sequence. The latter can be thought of as a variable binding motif that can be specified by designing corresponding gRNA sequences.

The **spacer** sequence is used in the gRNA construct to guide the CRISPR nuclease to the target **protospacer** sequence in the host genome. While a gRNA spacer sequence may not always uniquely target the host genome (i.e. it may map to multiple protospacers in the host genome), we can, for a given reference genome, uniquely identify a protospacer sequence with a combination of 3 attributes:

- `chr`: chromosome name
- `strand`: forward (+) or reverse (-)
- `pam_site`: genomic coordinate of the first nucleotide of the nuclease-specific PAM sequence; for SpCas9 this is the "N" in the NGG PAM sequence

For CRISPRko applications, we use an additional genomic coordinate, called `cut_site`, to represent where the double-stranded break (DSB) occurs. For SpCas9, the cut site (blunt-ended dsDNA break) is located 4nt upstream of the pam_site (PAM-proximal editing).

#### Installation

See the Installation tutorial to learn how to install the packages necessary for this tutorial: `crisprDesign`, `crisprDesignData`

#### End-to-end gRNA design workflow

We first start by loading the crisprVerse packages needed for this tutorial:

```
library(crisprBase)
library(crisprDesign)
library(crisprDesignData)
```

We will also load the `BSgenome` package containing DNA sequences for the hg38 genome:

```r
library(BSgenome.Hsapiens.UCSC.hg38)
```

#### Nuclease specification

We first load the `SpCas9` nuclease object from the `crisprBase` package:

```r
data(SpCas9, package="crisprBase")
SpCas9
```

```
## Class: CrisprNuclease
##   Name: SpCas9
##   Target type: DNA
##   Metadata: list of length 1
##   PAMs: NGG, NAG, NGA
##   Weights: 1, 0.2593, 0.0694
##   Spacer length: 20
##   PAM side: 3prime
##     Distance from PAM: 0
##   Prototype protospacers: 5'--SSSSSSSSSSSSSSSSSSSS[NGG]--3', 5'--SSSSSSSSSSSSSSSSSSSS[NAG]--3', 5'--[illegible]
```

To learn how to specify a custom nuclease, see the nuclease tutorial.

The three motifs (NGG, NAG and NGA) represent the recognized PAM sequences by SpCas9, and the weights indicate a recognition score. The canonical PAM sequence NGG is fully recognized (weight of 1), while the two non-canonical PAM sequences NAG and NGA are much less tolerated.

The spacer sequence is located on the 5-prime end with respect to the PAM sequence, and the default spacer sequence length is 20 nucleotides. If necessary, one can change the spacer length using the function `spacerLength` from `crisprBase`. We can inspect the protospacer construct by using `prototypeSequence`:

```r
prototypeSequence(SpCas9)
```

```
## [1] "5'--SSSSSSSSSSSSSSSSSSSS[NGG]--3'"
```

#### Specification of the target DNA sequence (KRAS CDS)

Since we aim to design gRNAs that knock out the human KRAS gene, we first need to retrieve the DNA sequence of the coding region (CDS) of KRAS. W e show in the gene annotation tutorial how to build convenient gene model objects that allows to quickly access gene-specific sequences. Here, we obtain from `crisprDesignData` a `GRangesList` object that defines the genomic coordinates (in hg38 coordinates) of coding genes in the human genome:

```r
data(txdb_human, package="crisprDesignData")
```

The `queryTxObject` function allows us to query this object for a specific gene and feature. Here, we obtain a `GRanges` object containing the CDS coordinates of KRAS:

```r
gr <- queryTxObject(txObject=txdb_human,
                    featureType="cds",
                    queryColumn="gene_symbol",
                    queryValue="KRAS")
```

To simplify our design, we will only consider exons that constitute the primary transcript of KRAS (transcript ID ENST00000311936).

```r
gr <- gr[gr$tx_id == "ENST00000311936"]
```

Optionally, we could also adjust the arguments in our call to `queryTxObject` to retrieve those transcript-specific coordinates:

```r
gr <- queryTxObject(txObject=txObject,
                    featureType="cds",
                    queryColumn="tx_id",
                    queryValue="ENST00000311936")
```

#### Finding spacer sequences targeting KRAS

`findSpacers` is the main function of `crisprDesign` for obtaining all possible spacer sequences that target protospacers located in our target DNA sequence(s). If a `GRanges` object is provided as input, a `BSgenome` object (an object that contains sequences of a reference genome) must be provided as well:

```r
bsgenome <- BSgenome.Hsapiens.UCSC.hg38
guideSet <- findSpacers(gr,
                        bsgenome=bsgenome,
                        crisprNuclease=SpCas9)
guideSet
```

```
## GuideSet object with 45 ranges and 5 metadata columns:
##              seqnames    ranges strand |          protospacer            pam
##                 <Rle> <IRanges>  <Rle> |       <DNAStringSet> <DNAStringSet>
##    spacer_1    chr12  25209843      - | AAAGAAAAAGATGAGCAAAGA            TGG
##    spacer_2    chr12  25209896      + | TTCTCGAACTAATGTATAGA            AGG
##    spacer_3    chr12  25225615      - | AACATCAGCAAAGACAAGAC            AGG
##    spacer_4    chr12  25225644      + | TTTGCTGATGTTTCAATAAA            AGG
##    spacer_5    chr12  25225653      - | CAGGACTTAGCAAGAAGTTA            TGG
##         ...      ...       ...    ... .                  ...            ...
##   spacer_41    chr12  25245343      - | GTAGTTGGAGCTGGTGGCGT            AGG
##   spacer_42    chr12  25245349      - | CTTGTGGTAGTTGGAGCTGG            TGG
##   spacer_43    chr12  25245352      - | AAACTTGTGGTAGTTGGAGC            TGG
##   spacer_44    chr12  25245358      - | GAATATAAACTTGTGGTAGT            TGG
##   spacer_45    chr12  25245365      - | AATGACTGAATATAAACTTG            TGG
##              pam_site  cut_site      region
##             <numeric> <numeric> <character>
##    spacer_1  25209843  25209846    region_8
##    spacer_2  25209896  25209893    region_8
##    spacer_3  25225615  25225618    region_7
##    spacer_4  25225644  25225641    region_7
##    spacer_5  25225653  25225656    region_7
##         ...       ...       ...         ...
##   spacer_41  25245343  25245346    region_5
##   spacer_42  25245349  25245352    region_5
##   spacer_43  25245352  25245355    region_5
##   spacer_44  25245358  25245361    region_5
##   spacer_45  25245365  25245368    region_5
##   -------
##   seqinfo: 640 sequences (1 circular) from hg38 genome
##   crisprNuclease: SpCas9
```

This function returns a `GuideSet` object that stores the genomic coordinates (PAM sites) for all spacer sequences found in the regions provided by `gr`. The `GuideSet` object is an extension of a `GenomicRanges` object that stores additional information about gRNAs.

There are several accessor functions we can use to extract information about the spacer sequences in `guideSet`, and here are a few examples with their corresponding outputs:

```
spacers(guideSet)
```

```
## DNAStringSet object of length 45:
##      width seq                                                  names
##  [1]    20 AAAGAAAAGATGAGCAAAGA                                 spacer_1
##  [2]    20 TTCTCGAACTAATGTATAGA                                 spacer_2
##  [3]    20 AACATCAGCAAAGACAAGAC                                 spacer_3
##  [4]    20 TTTGCTGATGTTTCAATAAA                                 spacer_4
##  [5]    20 CAGGACTTAGCAAGAAGTTA                                 spacer_5
##  ...   ... ...
## [41]    20 GTAGTTGGAGCTGGTGGCGT                                 spacer_41
## [42]    20 CTTGTGGTAGTTGGAGCTGG                                 spacer_42
## [43]    20 AAACTTGTGGTAGTTGGAGC                                 spacer_43
## [44]    20 GAATATAAACTTGTGGTAGT                                 spacer_44
## [45]    20 AATGACTGAATATAAACTTG                                 spacer_45
```

```
protospacers(guideSet)
```

```
## DNAStringSet object of length 45:
##      width seq                                                  names
##  [1]    20 AAAGAAAAGATGAGCAAAGA                                 spacer_1
##  [2]    20 TTCTCGAACTAATGTATAGA                                 spacer_2
##  [3]    20 AACATCAGCAAAGACAAGAC                                 spacer_3
##  [4]    20 TTTGCTGATGTTTCAATAAA                                 spacer_4
##  [5]    20 CAGGACTTAGCAAGAAGTTA                                 spacer_5
##  ...   ... ...
## [41]    20 GTAGTTGGAGCTGGTGGCGT                                 spacer_41
## [42]    20 CTTGTGGTAGTTGGAGCTGG                                 spacer_42
## [43]    20 AAACTTGTGGTAGTTGGAGC                                 spacer_43
## [44]    20 GAATATAAACTTGTGGTAGT                                 spacer_44
## [45]    20 AATGACTGAATATAAACTTG                                 spacer_45
```

```
pams(guideSet)
```

```
## DNAStringSet object of length 45:
##      width seq                                                  names
##  [1]     3 TGG                                                  spacer_1
##  [2]     3 AGG                                                  spacer_2
##  [3]     3 AGG                                                  spacer_3
##  [4]     3 AGG                                                  spacer_4
##  [5]     3 TGG                                                  spacer_5
##  ...   ... ...
## [41]     3 AGG                                                  spacer_41
## [42]     3 TGG                                                  spacer_42
## [43]     3 TGG                                                  spacer_43
## [44]     3 TGG                                                  spacer_44
## [45]     3 TGG                                                  spacer_45
```

```
head(pamSites(guideSet))
```

```
## spacer_1 spacer_2 spacer_3 spacer_4 spacer_5 spacer_6
## 25209843 25209896 25225615 25225644 25225653 25225672
```

```
head(cutSites(guideSet))
```

```
## spacer_1 spacer_2 spacer_3 spacer_4 spacer_5 spacer_6
## 25209846 25209893 25225618 25225641 25225656 25225675
```

#### Characterizing gRNA spacer sequences

There are specific spacer sequence features, independent of the genomic context of the protospacer sequence, that can reduce or even eliminate gRNA activity:

- **Poly-T stretches**: four or more consecutive T nucleotides in the spacer sequence may act as a transcriptional termination signal for the U6 promoter.
- **Self-complementarity**: complementary sites with the gRNA backbone can compete with the targeted genomic sequence.
- **Percent GC**: gRNAs with GC content between 20% and 80% are preferred.

Use the function `addSequenceFeatures` to evaluate the spacer sequences with respect to these characteristics and add the results to the `GuideSet` object:

```
guideSet <- addSequenceFeatures(guideSet)
head(guideSet)
```

```
## GuideSet object with 6 ranges and 11 metadata columns:
##            seqnames    ranges strand |          protospacer            pam
##               <Rle> <IRanges>  <Rle> |       <DNAStringSet> <DNAStringSet>
##   spacer_1    chr12  25209843      - | AAAGAAAAGATGAGCAAAGA            TGG
##   spacer_2    chr12  25209896      + | TTCTCGAACTAATGTATAGA            AGG
##   spacer_3    chr12  25225615      - | AACATCAGCAAAGACAAGAC            AGG
##   spacer_4    chr12  25225644      + | TTTGCTGATGTTTCAATAAA            AGG
##   spacer_5    chr12  25225653      - | CAGGACTTAGCAAGAAGTTA            TGG
##   spacer_6    chr12  25225672      - | AGTAGACACAAAACAGGCTC            AGG
##             pam_site  cut_site      region percentGC     polyA     polyC
##            <numeric> <numeric> <character> <numeric> <logical> <logical>
##   spacer_1  25209843  25209846    region_8        30      TRUE     FALSE
##   spacer_2  25209896  25209893    region_8        30     FALSE     FALSE
##   spacer_3  25225615  25225618    region_7        40     FALSE     FALSE
##   spacer_4  25225644  25225641    region_7        25     FALSE     FALSE
##   spacer_5  25225653  25225656    region_7        40     FALSE     FALSE
##   spacer_6  25225672  25225675    region_7        45      TRUE     FALSE
##                polyG     polyT startingGGGGG
##            <logical> <logical>     <logical>
##   spacer_1     FALSE     FALSE         FALSE
##   spacer_2     FALSE     FALSE         FALSE
##   spacer_3     FALSE     FALSE         FALSE
##   spacer_4     FALSE     FALSE         FALSE
##   spacer_5     FALSE     FALSE         FALSE
##   spacer_6     FALSE     FALSE         FALSE
##   -------
##   seqinfo: 640 sequences (1 circular) from hg38 genome
##   crisprNuclease: SpCas9
```

#### Off-target search with bowtie

In order to select gRNAs that are most specific to our target of interest, it is important to avoid gRNAs that target additional loci in the genome with either perfect sequence complementarity (multiple on-targets), or imperfect complementarity through tolerated mismatches (off-targets). As the SpCas9 nuclease can tolerate mismatches between the gRNA spacer sequence (RNA) and the protospacer sequence (DNA), it is necessary to characterize off-targets to minimize the introduction of double-stranded breaks (DSBs) beyond our intended target.

The `addSpacerAlignments` function appends a list of putative on- and off-targets to a `GuideSet` object using one of three methods. The first method uses the fast aligner bowtie [@langmead2009bowtie] via the crisprBowtie package to map spacer sequences to a specified reference genome. This can be done by specifying aligner="bowtie and providing a path to a bowtie index file to aligner_index in addSpacerAlignments.

We can control the alignment parameters and output with several function arguments.

- n_mismatches sets the maximum number of permitted gRNA:DNA mismatches (up to 3 mismatches).
- n_max_alignments specifies the maximum number of alignments for a given gRNA spacer sequence (1000 by default).
- all_alignments, when set to TRUE, overrules the n_max_alignments and returns all possible alignments.
- canonical filters out protospacer sequences that do not have a canonical PAM sequence when TRUE.

Let's search for on- and off-targets having up to 2 mismatches using bowtie. To use bowtie, we need to specify a bowtie index for the human genome:

```
# Path of the hg38 bowtie index on my personal laptop:
bowtie_index <- "/Users/fortinj2/crisprIndices/bowtie/hg38/hg38"
```

For instructions on how to build a Bowtie index from a given reference genome, see the genome index tutorial or the crisprBowtie page .

We will also specify the gene model object txdb_human from crisprDesignData described above for txObject argument, which is needed for the function to annotate genomic alignments with genic context. This is useful for identifying potentially more problematic off-targets, such as those located in the CDS of another gene, for instance.

```
guideSet <- addSpacerAlignments(guideSet,
                                aligner="bowtie",
                                aligner_index=bowtie_index,
                                bsgenome=BSgenome.Hsapiens.UCSC.hg38,
                                n_mismatches=2,
                                txObject=txdb_human)
```

```
## [runCrisprBowtie] Using BSgenome.Hsapiens.UCSC.hg38
## [runCrisprBowtie] Searching for SpCas9 protospacers
```

```
guideSet
```

```
## GuideSet object with 45 ranges and 18 metadata columns:
##              seqnames    ranges strand |          protospacer            pam
##                 <Rle> <IRanges>  <Rle> |       <DNAStringSet> <DNAStringSet>
##    spacer_1     chr12  25209843      - | AAAGAAAAGATGAGCAAAGA            TGG
##    spacer_2     chr12  25209896      + | TTCTCGAACTAATGTATAGA            AGG
##    spacer_3     chr12  25225615      - | AACATCAGCAAAGACAAGAC            AGG
##    spacer_4     chr12  25225644      + | TTTGCTGATGTTTCAATAAA            AGG
##    spacer_5     chr12  25225653      - | CAGGACTTAGCAAGAAGTTA            TGG
##         ...       ...       ...    ... .                  ...            ...
##   spacer_41     chr12  25245343      - | GTAGTTGGAGCTGGTGGCGT            AGG
##   spacer_42     chr12  25245349      - | CTTGTGGTAGTTGGAGCTGG            TGG
##   spacer_43     chr12  25245352      - | AAACTTGTGGTAGTTGGAGC            TGG
##   spacer_44     chr12  25245358      - | GAATATAAACTTGTGGTAGT            TGG
##   spacer_45     chr12  25245365      - | AATGACTGAATATAAACTTG            TGG
##               pam_site  cut_site      region percentGC     polyA     polyC
##              <numeric> <numeric> <character> <numeric> <logical> <logical>
##    spacer_1   25209843  25209846    region_8        30      TRUE     FALSE
##    spacer_2   25209896  25209893    region_8        30     FALSE     FALSE
##    spacer_3   25225615  25225618    region_7        40     FALSE     FALSE
##    spacer_4   25225644  25225641    region_7        25     FALSE     FALSE
##    spacer_5   25225653  25225656    region_7        40     FALSE     FALSE
```

```
##          ...       ...       ...         ...       ...       ...       ...
##   spacer_41  25245343  25245346    region_5        60     FALSE     FALSE
##   spacer_42  25245349  25245352    region_5        55     FALSE     FALSE
##   spacer_43  25245352  25245355    region_5        45     FALSE     FALSE
##   spacer_44  25245358  25245361    region_5        30     FALSE     FALSE
##   spacer_45  25245365  25245368    region_5        25     FALSE     FALSE
##                 polyG     polyT startingGGGGG        n0        n1        n2
##             <logical> <logical>     <logical> <numeric> <numeric> <numeric>
##    spacer_1     FALSE     FALSE         FALSE         1         2        19
##    spacer_2     FALSE     FALSE         FALSE         1         1         0
##    spacer_3     FALSE     FALSE         FALSE         1         0         1
##    spacer_4     FALSE     FALSE         FALSE         1         0         3
##    spacer_5     FALSE     FALSE         FALSE         1         1         0
##         ...       ...       ...           ...       ...       ...       ...
##   spacer_41     FALSE     FALSE         FALSE         1         0         1
##   spacer_42     FALSE     FALSE         FALSE         1         1         0
##   spacer_43     FALSE     FALSE         FALSE         1         1         1
##   spacer_44     FALSE     FALSE         FALSE         1         1         0
##   spacer_45     FALSE     FALSE         FALSE         2         0         0
##                  n0_c      n1_c      n2_c
##             <numeric> <numeric> <numeric>
##    spacer_1         1         0         0
##    spacer_2         1         0         0
##    spacer_3         1         0         0
##    spacer_4         1         0         0
##    spacer_5         1         0         0
##         ...       ...       ...       ...
##   spacer_41         1         0         0
##   spacer_42         1         0         0
##   spacer_43         1         0         1
##   spacer_44         1         0         0
##   spacer_45         1         0         0
##                                                         alignments
##                                                      <GRangesList>
##    spacer_1   chr12:25209843:-,chr6:54771089:+,chr5:4348033:+,...
##    spacer_2                      chr12:25209896:+,chr6:54771050:-
##    spacer_3                     chr12:25225615:-,chr12:88104409:-
##    spacer_4 chr12:25225644:+,chr5:59533197:+,chr9:132980355:-,...
##    spacer_5                      chr12:25225653:-,chr6:54771002:+
##         ...                                                   ...
##   spacer_41                     chr12:25245343:-,chr9:121850427:+
##   spacer_42                      chr12:25245349:-,chr6:54770618:+
##   spacer_43     chr12:25245352:-,chr6:54770615:+,chr1:114716128:-
##   spacer_44                      chr12:25245358:-,chr6:54770609:+
##   spacer_45                      chr12:25245365:-,chr6:54770602:+
##   -------
##   seqinfo: 640 sequences (1 circular) from hg38 genome
##   crisprNuclease: SpCas9
```

Several columns were added to the `GuideSet` object that summarize the number of on- and off-targets for each spacer sequence, and take genomic context into account:

- **n0, n1, n2, n3**: specify the number of alignments with 0, 1, 2 and 3 mismatches, respectively.
- **n0_c, n1_c, n2_c, n3_c**: specify the number of alignments in a coding region, with 0, 1, 2 and 3 mismatches, respectively.

- **n0__p, n1__p, n2__p, n3__p**: specify the number of alignments in a promoter region of a coding gene, with 0, 1, 2 and 3 mismatches, respectively.

Our `guideSet` now has columns of the first two categories, up to 2 mismatches (the value passed to `n_mismatches`); had we also supplied a `GRanges` of TSS coordinates to the `tssObject` argument, our `guideSet` would include columns in the last category.

To inspect individual on- and off-targets and their context, one can use the `alignments` function, which returns a table of all genomic alignments stored in the `GuideSet` object:

```
alignments(guideSet)
```

```
## GRanges object with 149 ranges and 14 metadata columns:
##             seqnames    ranges strand |               spacer
##                <Rle> <IRanges>  <Rle> |       <DNAStringSet>
##    spacer_1    chr12  25209843      - | AAAGAAAAGATGAGCAAAGA
##    spacer_1     chr6  54771089      + | AAAGAAAAGATGAGCAAAGA
##    spacer_1     chr5   4348033      + | AAAGAAAAGATGAGCAAAGA
##    spacer_1    chr10  35808237      - | AAAGAAAAGATGAGCAAAGA
##    spacer_1    chr20  55997350      + | AAAGAAAAGATGAGCAAAGA
##         ...      ...       ...    ... .                  ...
##   spacer_43     chr1 114716128      - | AAACTTGTGGTAGTTGGAGC
##   spacer_44    chr12  25245358      - | GAATATAAACTTGTGGTAGT
##   spacer_44     chr6  54770609      + | GAATATAAACTTGTGGTAGT
##   spacer_45    chr12  25245365      - | AATGACTGAATATAAACTTG
##   spacer_45     chr6  54770602      + | AATGACTGAATATAAACTTG
##                      protospacer            pam  pam_site n_mismatches
##                   <DNAStringSet> <DNAStringSet> <numeric>    <integer>
##    spacer_1 AAAGAAAAGATGAGCAAAGA            TGG  25209843            0
##    spacer_1 AAATAAAAGATGAGCAAAGA            TGG  54771089            1
##    spacer_1 AGAGAAAAGATGAGCAAAGA            TGG   4348033            1
##    spacer_1 AAAGAAAAAAGGAGCAAAGA            AGG  35808237            2
##    spacer_1 AAAGAAAAAAGGAGCAAAGA            AGG  55997350            2
##         ...                  ...            ...       ...          ...
##   spacer_43 AAACTGGTGGTGGTTGGAGC            AGG 114716128            2
##   spacer_44 GAATATAAACTTGTGGTAGT            TGG  25245358            0
##   spacer_44 GAATATAAACTTGCGGTAGT            TGG  54770609            1
##   spacer_45 AATGACTGAATATAAACTTG            TGG  25245365            0
##   spacer_45 AATGACTGAATATAAACTTG            CGG  54770602            0
##             canonical  cut_site         cds   fiveUTRs   threeUTRs       exons
##             <logical> <numeric> <character> <character> <character> <character>
##    spacer_1      TRUE  25209846        KRAS        <NA>        KRAS        KRAS
##    spacer_1      TRUE  54771086        <NA>        <NA>        <NA>      KRASP1
##    spacer_1      TRUE   4348030        <NA>        <NA>        <NA>        <NA>
##    spacer_1      TRUE  35808240        <NA>        <NA>        <NA>        <NA>
##    spacer_1      TRUE  55997347        <NA>        <NA>        <NA>        <NA>
##         ...       ...       ...         ...         ...         ...         ...
##   spacer_43      TRUE 114716131        NRAS        <NA>        <NA>        NRAS
##   spacer_44      TRUE  25245361        KRAS        <NA>        <NA>        KRAS
##   spacer_44      TRUE  54770606        <NA>        <NA>        <NA>      KRASP1
##   spacer_45      TRUE  25245368        KRAS        <NA>        <NA>        KRAS
##   spacer_45      TRUE  54770599        <NA>        <NA>        <NA>      KRASP1
##                 introns  intergenic intergenic_distance
##             <character> <character>           <integer>
##    spacer_1        <NA>        <NA>                <NA>
##    spacer_1        <NA>        <NA>                <NA>
```

```
##    spacer_1        <NA>                              88819
##    spacer_1        <NA>       PCAT5                   7319
##    spacer_1        <NA>       CBLN4                      9
##         ...         ...         ...                    ...
##   spacer_43        <NA>        <NA>                   <NA>
##   spacer_44        <NA>        <NA>                   <NA>
##   spacer_44        <NA>        <NA>                   <NA>
##   spacer_45        <NA>        <NA>                   <NA>
##   spacer_45        <NA>        <NA>                   <NA>
##   -------
##   seqinfo: 25 sequences (1 circular) from hg38 genome
```

Similarly, the functions `onTargets` and `offTargets` return on-target alignments (no mismatches) and off-target alignments (having at least one mismatch), respectively. See `?addSpacerAlignments` for more details about the different options.

We note that gRNAs that align to hundreds of different locations are highly unspecific and undesirable. This can also cause `addSpacerAlignments` to be slow. The function `addSpacerAlignmentsIterative` is an iterative version of `addSpacerAlignments` that curtails alignment searches for gRNAs having more hits than the user-defined threshold. See `?addSpacerAlignmentsIterative` for more information.

#### Removing repeat elements

Many promiscuous protospacer sequences occur in repeats or low-complexity DNA sequences (regions identified by RepeatMasker). These sequences are usually not of interest due to their low specificity, and can be easily removed with `removeRepeats`:

```
data("gr.repeats.hg38", package="crisprDesignData")
guideSet <- removeRepeats(guideSet,
                          gr.repeats=gr.repeats.hg38)
```

#### Off-target scoring (MIT and CFD specificity scores)

After retrieving a list of putative off-targets and on-targets for a given spacer sequence, we can use `addOffTargetScores` to predict the likelihood of the nuclease to cut at the off-target locations based on mismatch tolerance

```
guideSet <- addOffTargetScores(guideSet)
guideSet
```

```
## GuideSet object with 45 ranges and 21 metadata columns:
##              seqnames    ranges strand |          protospacer            pam
##                 <Rle> <IRanges>  <Rle> |       <DNAStringSet> <DNAStringSet>
##    spacer_1     chr12  25209843      - | AAAGAAAAGATGAGCAAAGA            TGG
##    spacer_2     chr12  25209896      + | TTCTCGAACTAATGTATAGA            AGG
##    spacer_3     chr12  25225615      - | AACATCAGCAAAGACAAGAC            AGG
##    spacer_4     chr12  25225644      + | TTTGCTGATGTTTCAATAAA            AGG
##    spacer_5     chr12  25225653      - | CAGGACTTAGCAAGAAGTTA            TGG
##         ...       ...       ...    ... .                  ...            ...
##   spacer_41     chr12  25245343      - | GTAGTTGGAGCTGGTGGCGT            AGG
##   spacer_42     chr12  25245349      - | CTTGTGGTAGTTGGAGCTGG            TGG
##   spacer_43     chr12  25245352      - | AAACTTGTGGTAGTTGGAGC            TGG
##   spacer_44     chr12  25245358      - | GAATATAAACTTGTGGTAGT            TGG
##   spacer_45     chr12  25245365      - | AATGACTGAATATAAACTTG            TGG
##               pam_site  cut_site      region percentGC     polyA     polyC
##              <numeric> <numeric> <character> <numeric> <logical> <logical>
```

```
##    spacer_1  25209843  25209846    region_8        30      TRUE     FALSE
##    spacer_2  25209896  25209893    region_8        30     FALSE     FALSE
##    spacer_3  25225615  25225618    region_7        40     FALSE     FALSE
##    spacer_4  25225644  25225641    region_7        25     FALSE     FALSE
##    spacer_5  25225653  25225656    region_7        40     FALSE     FALSE
##         ...       ...       ...         ...       ...       ...       ...
##   spacer_41  25245343  25245346    region_5        60     FALSE     FALSE
##   spacer_42  25245349  25245352    region_5        55     FALSE     FALSE
##   spacer_43  25245352  25245355    region_5        45     FALSE     FALSE
##   spacer_44  25245358  25245361    region_5        30     FALSE     FALSE
##   spacer_45  25245365  25245368    region_5        25     FALSE     FALSE
##                 polyG     polyT startingGGGGG        n0        n1        n2
##             <logical> <logical>     <logical> <numeric> <numeric> <numeric>
##    spacer_1     FALSE     FALSE         FALSE         1         2        19
##    spacer_2     FALSE     FALSE         FALSE         1         1         0
##    spacer_3     FALSE     FALSE         FALSE         1         0         1
##    spacer_4     FALSE     FALSE         FALSE         1         0         3
##    spacer_5     FALSE     FALSE         FALSE         1         1         0
##         ...       ...       ...           ...       ...       ...       ...
##   spacer_41     FALSE     FALSE         FALSE         1         0         1
##   spacer_42     FALSE     FALSE         FALSE         1         1         0
##   spacer_43     FALSE     FALSE         FALSE         1         1         1
##   spacer_44     FALSE     FALSE         FALSE         1         1         0
##   spacer_45     FALSE     FALSE         FALSE         2         0         0
##                  n0_c      n1_c      n2_c
##             <numeric> <numeric> <numeric>
##    spacer_1         1         0         0
##    spacer_2         1         0         0
##    spacer_3         1         0         0
##    spacer_4         1         0         0
##    spacer_5         1         0         0
##         ...       ...       ...       ...
##   spacer_41         1         0         0
##   spacer_42         1         0         0
##   spacer_43         1         0         1
##   spacer_44         1         0         0
##   spacer_45         1         0         0
##                                                       alignments inRepeats
##                                                    <GRangesList> <logical>
##    spacer_1   chr12:25209843:-,chr6:54771089:+,chr5:4348033:+,...     FALSE
##    spacer_2                     chr12:25209896:+,chr6:54771050:-     FALSE
##    spacer_3                    chr12:25225615:-,chr12:88104409:-     FALSE
##    spacer_4 chr12:25225644:+,chr5:59533197:+,chr9:132980355:-,...     FALSE
##    spacer_5                     chr12:25225653:-,chr6:54771002:+     FALSE
##         ...                                                   ...       ...
##   spacer_41                    chr12:25245343:-,chr9:121850427:+     FALSE
##   spacer_42                     chr12:25245349:-,chr6:54770618:+     FALSE
##   spacer_43     chr12:25245352:-,chr6:54770615:+,chr1:114716128:-     FALSE
##   spacer_44                     chr12:25245358:-,chr6:54770609:+     FALSE
##   spacer_45                     chr12:25245365:-,chr6:54770602:+     FALSE
##             score_cfd score_mit
##             <numeric> <numeric>
##    spacer_1  0.176077  0.418801
##    spacer_2  0.500000  0.577367
```

```
##    spacer_3  0.518519  0.976087
##    spacer_4  0.530345  0.987166
##    spacer_5  0.606061  0.716846
##         ...       ...       ...
##   spacer_41  0.928339  0.999684
##   spacer_42  0.500000  0.547046
##   spacer_43  0.414474  0.612726
##   spacer_44  0.777778  0.759301
##   spacer_45  0.500000  0.500000
##   -------
##   seqinfo: 640 sequences (1 circular) from hg38 genome
##   crisprNuclease: SpCas9
```

Note that this will only work after calling `addSpacerAlignments`, as it requires a list of off-targets for each gRNA entry.

#### On-target scoring (gRNA efficiency)

`addOnTargetScores` adds scores from on-target efficiency algorithms specified by the `methods` argument (or all available methods if `NULL`) available in the R package crisprScore and appends them to the `GuideSet`. Here' let's add the DeepHF and DeepSpCas9 scores:

```
guideSet <- addOnTargetScores(guideSet,
                              methods=c("deephf", "deepspcas9"))
guideSet
```

```
## GuideSet object with 45 ranges and 23 metadata columns:
##               seqnames    ranges strand |          protospacer             pam
##                  <Rle> <IRanges>  <Rle> |       <DNAStringSet> <DNAStringSet>
##    spacer_1    chr12  25209843      - | AAAGAAAAGATGAGCAAAGA             TGG
##    spacer_2    chr12  25209896      + | TTCTCGAACTAATGTATAGA             AGG
##    spacer_3    chr12  25225615      - | AACATCAGCAAAGACAAGAC             AGG
##    spacer_4    chr12  25225644      + | TTTGCTGATGTTTCAATAAA             AGG
##    spacer_5    chr12  25225653      - | CAGGACTTAGCAAGAAGTTA             TGG
##         ...      ...       ...    ... .                  ...             ...
##   spacer_41    chr12  25245343      - | GTAGTTGGAGCTGGTGGCGT             AGG
##   spacer_42    chr12  25245349      - | CTTGTGGTAGTTGGAGCTGG             TGG
##   spacer_43    chr12  25245352      - | AAACTTGTGGTAGTTGGAGC             TGG
##   spacer_44    chr12  25245358      - | GAATATAAACTTGTGGTAGT             TGG
##   spacer_45    chr12  25245365      - | AATGACTGAATATAAACTTG             TGG
##                pam_site  cut_site      region percentGC     polyA     polyC
##               <numeric> <numeric> <character> <numeric> <logical> <logical>
##    spacer_1  25209843  25209846    region_8        30      TRUE     FALSE
##    spacer_2  25209896  25209893    region_8        30     FALSE     FALSE
##    spacer_3  25225615  25225618    region_7        40     FALSE     FALSE
##    spacer_4  25225644  25225641    region_7        25     FALSE     FALSE
##    spacer_5  25225653  25225656    region_7        40     FALSE     FALSE
##         ...       ...       ...         ...       ...       ...       ...
##   spacer_41  25245343  25245346    region_5        60     FALSE     FALSE
##   spacer_42  25245349  25245352    region_5        55     FALSE     FALSE
##   spacer_43  25245352  25245355    region_5        45     FALSE     FALSE
##   spacer_44  25245358  25245361    region_5        30     FALSE     FALSE
##   spacer_45  25245365  25245368    region_5        25     FALSE     FALSE
##                  polyG     polyT startingGGGGG        n0        n1        n2
##               <logical> <logical>     <logical> <numeric> <numeric> <numeric>
```

```
##    spacer_1     FALSE     FALSE         FALSE         1         2        19
##    spacer_2     FALSE     FALSE         FALSE         1         1         0
##    spacer_3     FALSE     FALSE         FALSE         1         0         1
##    spacer_4     FALSE     FALSE         FALSE         1         0         3
##    spacer_5     FALSE     FALSE         FALSE         1         1         0
##         ...       ...       ...           ...       ...       ...       ...
##   spacer_41     FALSE     FALSE         FALSE         1         0         1
##   spacer_42     FALSE     FALSE         FALSE         1         1         0
##   spacer_43     FALSE     FALSE         FALSE         1         1         1
##   spacer_44     FALSE     FALSE         FALSE         1         1         0
##   spacer_45     FALSE     FALSE         FALSE         2         0         0
##                  n0_c      n1_c      n2_c
##             <numeric> <numeric> <numeric>
##    spacer_1         1         0         0
##    spacer_2         1         0         0
##    spacer_3         1         0         0
##    spacer_4         1         0         0
##    spacer_5         1         0         0
##         ...       ...       ...       ...
##   spacer_41         1         0         0
##   spacer_42         1         0         0
##   spacer_43         1         0         1
##   spacer_44         1         0         0
##   spacer_45         1         0         0
##                                                       alignments inRepeats
##                                                    <GRangesList> <logical>
##    spacer_1   chr12:25209843:-,chr6:54771089:+,chr5:4348033:+,...     FALSE
##    spacer_2                       chr12:25209896:+,chr6:54771050:-     FALSE
##    spacer_3                      chr12:25225615:-,chr12:88104409:-     FALSE
##    spacer_4 chr12:25225644:+,chr5:59533197:+,chr9:132980355:-,...     FALSE
##    spacer_5                       chr12:25225653:-,chr6:54771002:+     FALSE
##         ...                                                    ...       ...
##   spacer_41                      chr12:25245343:-,chr9:121850427:+     FALSE
##   spacer_42                       chr12:25245349:-,chr6:54770618:+     FALSE
##   spacer_43     chr12:25245352:-,chr6:54770615:+,chr1:114716128:-     FALSE
##   spacer_44                       chr12:25245358:-,chr6:54770609:+     FALSE
##   spacer_45                       chr12:25245365:-,chr6:54770602:+     FALSE
##             score_cfd score_mit score_deephf score_deepspcas9
##             <numeric> <numeric>    <numeric>        <numeric>
##    spacer_1  0.176077  0.418801     0.450868        0.4272767
##    spacer_2  0.500000  0.577367     0.428607        0.2041316
##    spacer_3  0.518519  0.976087     0.613590        0.5043279
##    spacer_4  0.530345  0.987166     0.182062        0.0782121
##    spacer_5  0.606061  0.716846     0.514199        0.3894395
##         ...       ...       ...          ...              ...
##   spacer_41  0.928339  0.999684     0.692967         0.585668
##   spacer_42  0.500000  0.547046     0.644286         0.525602
##   spacer_43  0.414474  0.612726     0.439317         0.365770
##   spacer_44  0.777778  0.759301     0.433265         0.255677
##   spacer_45  0.500000  0.500000     0.671397         0.627091
##   -------
##   seqinfo: 640 sequences (1 circular) from hg38 genome
##   crisprNuclease: SpCas9
```

See the crisprScore page for a full description of the different scores.

#### Restriction enzymes

Since the gRNA library synthesis process usually involves restriction enzymes, it is often necessary to remove gRNAs that contain restriction sites of specific enzymes. The function `addRestrictionEnzymes` allows the user to flag gRNAs containing restriction sites for a user-defined set of enzymes.

```
guideSet <- addRestrictionEnzymes(guideSet)
```

By default (that is, when `includeDefault` is `TRUE`), the function adds annotation for the following commonly used enzymes: EcoRI, KpnI, BsmBI, BsaI, BbsI, PacI, ISceI and MluI. Additional enzymes can be included by name via `enzymeNames`, and custom restriction sites can be defined using the `patterns` argument. It also accepts arguments to specify the nucleotide sequence that flanks the spacer sequence on the 5' end (`flanking5`) and on the 3' end (`flanking3`) in the lentiviral cassette used for gRNA delivery. The function effectively searches for restriction sites in the full sequence: `[flanking5][spacer][flanking3]`.

One can use the `enzymeAnnotation` function to retrieve the added annotation:

```
head(enzymeAnnotation(guideSet))
```

```
## DataFrame with 6 rows and 7 columns
##                EcoRI      KpnI     BsmBI      BsaI      BbsI      PacI      MluI
##            <logical> <logical> <logical> <logical> <logical> <logical> <logical>
## spacer_1       FALSE     FALSE     FALSE     FALSE     FALSE     FALSE     FALSE
## spacer_2       FALSE     FALSE     FALSE     FALSE     FALSE     FALSE     FALSE
## spacer_3       FALSE     FALSE     FALSE     FALSE     FALSE     FALSE     FALSE
## spacer_4       FALSE     FALSE     FALSE     FALSE     FALSE     FALSE     FALSE
## spacer_5       FALSE     FALSE     FALSE     FALSE     FALSE     FALSE     FALSE
## spacer_6       FALSE     FALSE     FALSE     FALSE     FALSE     FALSE     FALSE
```

#### Gene annotation

The function `addGeneAnnotation` adds transcript- and gene-level context to gRNAs from a `TxDb`-like object:

```
guideSet <- addGeneAnnotation(guideSet,
                              txObject=txdb_human)
```

The gene annotation can be retrieved using the function `geneAnnotation`:

```
geneAnnotation(guideSet)
```

```
## DataFrame with 113 rows and 23 columns
##                  chr anchor_site   strand gene_symbol         gene_id
##             <factor>   <integer> <factor> <character>     <character>
## spacer_1       chr12    25209846        -        KRAS ENSG00000133703
## spacer_1       chr12    25209846        -        KRAS ENSG00000133703
## spacer_1       chr12    25209846        -        KRAS ENSG00000133703
## spacer_2       chr12    25209893        +        KRAS ENSG00000133703
## spacer_2       chr12    25209893        +        KRAS ENSG00000133703
## ...              ...         ...      ...         ...             ...
## spacer_44      chr12    25245361        -        KRAS ENSG00000133703
## spacer_45      chr12    25245368        -        KRAS ENSG00000133703
## spacer_45      chr12    25245368        -        KRAS ENSG00000133703
## spacer_45      chr12    25245368        -        KRAS ENSG00000133703
## spacer_45      chr12    25245368        -        KRAS ENSG00000133703
##                      tx_id      protein_id   cut_cds cut_fiveUTRs cut_threeUTRs
##                <character>     <character> <logical>    <logical>     <logical>
```

```
## spacer_1  ENST00000256078              NA     FALSE        FALSE          TRUE
## spacer_1  ENST00000311936 ENSP00000308495      TRUE        FALSE         FALSE
## spacer_1  ENST00000557334 ENSP00000452512      TRUE        FALSE         FALSE
## spacer_2  ENST00000256078              NA     FALSE        FALSE          TRUE
## spacer_2  ENST00000311936 ENSP00000308495      TRUE        FALSE         FALSE
## ...                   ...             ...       ...          ...           ...
## spacer_44 ENST00000556131 ENSP00000256078      TRUE        FALSE         FALSE
## spacer_45 ENST00000256078 ENSP00000256078      TRUE        FALSE         FALSE
## spacer_45 ENST00000311936 ENSP00000256078      TRUE        FALSE         FALSE
## spacer_45 ENST00000557334 ENSP00000256078      TRUE        FALSE         FALSE
## spacer_45 ENST00000556131 ENSP00000256078      TRUE        FALSE         FALSE
##           cut_introns percentCDS aminoAcidIndex downtreamATG percentTx
##             <logical>  <numeric>      <numeric>    <numeric> <numeric>
## spacer_1        FALSE         NA             NA           NA      15.3
## spacer_1        FALSE       91.0            172            1      13.3
## spacer_1        FALSE       77.6             59            1      35.6
## spacer_2        FALSE         NA             NA           NA      14.4
## spacer_2        FALSE       82.7            157            2      12.4
## ...               ...        ...            ...          ...       ...
## spacer_44       FALSE       18.2              8            1      11.9
## spacer_45       FALSE        3.0              6            2       3.8
## spacer_45       FALSE        3.0              6            2       3.9
## spacer_45       FALSE        7.5              6            2      20.3
## spacer_45       FALSE       12.9              6            1      11.4
##           nIsoforms totalIsoforms percentIsoforms isCommonExon nCodingIsoforms
##           <integer>     <numeric>       <numeric>    <logical>       <integer>
## spacer_1          3             4              75        FALSE               3
## spacer_1          3             4              75        FALSE               3
## spacer_1          3             4              75        FALSE               3
## spacer_2          3             4              75        FALSE               3
## spacer_2          3             4              75        FALSE               3
## ...             ...           ...             ...          ...             ...
## spacer_44         4             4             100         TRUE               4
## spacer_45         4             4             100         TRUE               4
## spacer_45         4             4             100         TRUE               4
## spacer_45         4             4             100         TRUE               4
## spacer_45         4             4             100         TRUE               4
##           totalCodingIsoforms percentCodingIsoforms isCommonCodingExon
##                     <numeric>             <numeric>          <logical>
## spacer_1                    4                    75              FALSE
## spacer_1                    4                    75              FALSE
## spacer_1                    4                    75              FALSE
## spacer_2                    4                    75              FALSE
## spacer_2                    4                    75              FALSE
## ...                       ...                   ...                ...
## spacer_44                   4                   100               TRUE
## spacer_45                   4                   100               TRUE
## spacer_45                   4                   100               TRUE
## spacer_45                   4                   100               TRUE
## spacer_45                   4                   100               TRUE
```

It provides a great deal of information in describing the genomic location of the protospacer sequences.

- Ensembl ID columns are provided for all applicable levels: `gene_id`, `tx_id`, `protein_id`, `exon_id`.
- `exon_rank` gives the order of the exon for the transcript; for example "2" indicates it is the second

exon (from the 5' end) in the mature transcript.

- several columns describe for which gene the the guide sequence overlaps the indicated transcript segment: `cut_cds`, `cut_fiveUTRs`, `cut_threeUTRs`, `cut_introns`.
- `percentCDS` and `percentTx` give the location of the `cut_site` within the CDS of the transcript and the entire transcript, respectively, as a percent from the 5' end to the 3' end.
- `aminoAcidIndex` gives the number of the specific amino acid in the protein where the cut is predicted to occur.
- `downstreamATG` shows how many in-frame ATGs are downstream of the `cut_site` (and upstream from the defined percent transcript cutoff, `met_cutoff`), indicating a potential alternative translation initiation site that may preserve protein function.
- isoform coverage is described by four columns:
  - `nIsoforms` gives the number of isoforms of the target gene (from `gene_id`) that overlap with the protospacer sequence.
  - `totalIsoforms` is the number of isoforms for the target gene.
  - `percentIsoforms` calculates the percentage of isoforms for the target gene that overlap with the protospacer sequence (`100*nIsoforms/totalIsoforms`).
  - `isCommonExon` identifies protospacer sequences that overlap with all isoforms for the target gene.
- isoform coverage when exclusively considering the CDS of the target gene is similarly described by the `nCodingIsoforms`, `totalCodingIsoforms`, `percentCodingIsoforms`, and `isCommonCodingExon` columns.
- `pfam` gives the ID of Pfam domain(s) overlapping the protospacer sequence.

#### TSS annotation

Similarly, one might want to know which protospacer sequences are located within promoter regions of known genes:

```
data(tssObjectExample, package="crisprDesign")
guideSet <- addTssAnnotation(guideSet,
                             tssObject=tssObjectExample)
tssAnnotation(guideSet)
```

```
## DataFrame with 0 rows and 11 columns
```

Not surprisingly, as our `GuideSet` targets the CDS of KRAS, none of our guides overlap a gene promoter region.

#### SNP annotation

Common single-nucleotide polymorphisms (SNPs) can change the on-target and off-target properties of gRNAs by altering the binding. The function `addSNPAnnotation` annotates gRNAs with respect to a reference database of SNPs (stored in a VCF file), specified by the `vcf` argument.

VCF files for common SNPs (dbSNPs) can be downloaded from NCBI on the dbSNP website. We will use one of those files, after having downloaded it to our local machine.

```
vcf <- "/Users/fortinj2/crisprIndices/snps/dbsnp151.grch38/00-common_all_snps_only.vcf.gz"
guideSet <- addSNPAnnotation(guideSet, vcf=vcf)
snps(guideSet)
```

```
## DataFrame with 1 row and 9 columns
##                    rs   rs_site rs_site_rel     allele_ref   allele_minor
##           <character> <integer>   <numeric> <DNAStringSet> <DNAStringSet>
## spacer_1    rs1137282  25209843           0              A              G
##           MAF_1000G MAF_TOPMED        type    length
##           <numeric>  <numeric> <character> <integer>
## spacer_1     0.1755    0.19671         snp         1
```

The rs_site_rel gives the relative position of the SNP with respect to the pam_site. allele_ref and allele_minor report the nucleotide of the reference and minor alleles, respectively. MAF_1000G and MAF_TOPMED report the minor allele frequency (MAF) in the 1000Genomes and TOPMED populations, respectively.

#### Filtering and ranking gRNAs

Once gRNAs are fully annotated, it is easy to filter out any unwanted gRNAs since GuideSet objects can be subsetted like regular vectors in R.

As an example, suppose that we only want to keep gRNAs that have percent GC between 20% and 80% and that do not contain a polyT stretch. This can be achieved using the following lines:

```
guideSet <- guideSet[guideSet$percentGC>=20]
guideSet <- guideSet[guideSet$percentGC<=80]
guideSet <- guideSet[!guideSet$polyT]
```

Similarly, it is easy to rank gRNAs based on a set of criteria using the regular order function.

For instance, let's sort gRNAs by the DeepHF on-target score:

```
# Creating an ordering index based on the DeepHF score:
# Using the negative values to make sure higher scores are ranked first:
o <- order(-guideSet$score_deephf)
# Ordering the GuideSet:
guideSet <- guideSet[o]
head(guideSet)
```

```
## GuideSet object with 6 ranges and 28 metadata columns:
##            seqnames    ranges strand |          protospacer           pam
##               <Rle> <IRanges>  <Rle> |       <DNAStringSet> <DNAStringSet>
##   spacer_29    chr12  25227322      - | AAGAGGAGTACAGTGCAATG           AGG
##   spacer_28    chr12  25227321      - | AGAGGAGTACAGTGCAATGA           GGG
##   spacer_41    chr12  25245343      - | GTAGTTGGAGCTGGTGGCGT           AGG
##   spacer_23    chr12  25227300      - | GGACCAGTACATGAGGACTG           GGG
##   spacer_24    chr12  25227301      - | GGGACCAGTACATGAGGACT           GGG
##   spacer_25    chr12  25227302      - | AGGGACCAGTACATGAGGAC           TGG
##             pam_site  cut_site      region percentGC     polyA     polyC
##            <numeric> <numeric> <character> <numeric> <logical> <logical>
##   spacer_29  25227322  25227325    region_6        45     FALSE     FALSE
##   spacer_28  25227321  25227324    region_6        45     FALSE     FALSE
##   spacer_41  25245343  25245346    region_5        60     FALSE     FALSE
##   spacer_23  25227300  25227303    region_6        55     FALSE     FALSE
##   spacer_24  25227301  25227304    region_6        55     FALSE     FALSE
##   spacer_25  25227302  25227305    region_6        55     FALSE     FALSE
##                polyG     polyT startingGGGGG        n0        n1        n2
##            <logical> <logical>     <logical> <numeric> <numeric> <numeric>
##   spacer_29     FALSE     FALSE         FALSE         1         0         1
##   spacer_28     FALSE     FALSE         FALSE         1         0         1
##   spacer_41     FALSE     FALSE         FALSE         1         0         1
##   spacer_23     FALSE     FALSE         FALSE         1         0         0
##   spacer_24     FALSE     FALSE         FALSE         1         1         1
##   spacer_25     FALSE     FALSE         FALSE         1         0         2
##                 n0_c      n1_c      n2_c
##            <numeric> <numeric> <numeric>
##   spacer_29         1         0         0
##   spacer_28         1         0         0
```

```
##   spacer_41         1         0         0
##   spacer_23         1         0         0
##   spacer_24         1         0         0
##   spacer_25         1         0         1
##                                                       alignments inRepeats
##                                                    <GRangesList> <logical>
##   spacer_29                   chr12:25227322:-,chr6:54770784:+     FALSE
##   spacer_28                  chr12:25227321:-,chr16:70072284:-     FALSE
##   spacer_41                  chr12:25245343:-,chr9:121850427:+     FALSE
##   spacer_23                                   chr12:25227300:-     FALSE
##   spacer_24 chr12:25227301:-,chr6:54770804:+,chr12:131824051:+     FALSE
##   spacer_25  chr12:25227302:-,chr1:114713868:-,chr6:54770803:+     FALSE
##             score_cfd score_mit score_deephf score_deepspcas9
##             <numeric> <numeric>    <numeric>        <numeric>
##   spacer_29  0.555556  1.000000     0.712546         0.761093
##   spacer_28  0.796078  1.000000     0.693555         0.777314
##   spacer_41  0.928339  0.999684     0.692967         0.585668
##   spacer_23  1.000000  1.000000     0.686824         0.710386
##   spacer_24  0.459184  0.621749     0.672358         0.604710
##   spacer_25  0.429499  0.954044     0.672062         0.444870
##                  enzymeAnnotation
##              <SplitDataFrameList>
##   spacer_29 FALSE:FALSE:FALSE:...
##   spacer_28 FALSE:FALSE:FALSE:...
##   spacer_41 FALSE:FALSE:FALSE:...
##   spacer_23 FALSE:FALSE:FALSE:...
##   spacer_24 FALSE:FALSE:FALSE:...
##   spacer_25 FALSE:FALSE:FALSE:...
##                                                             geneAnnotation
##                                                       <SplitDataFrameList>
##   spacer_29                     chr12:25227325:-:...,chr12:25227325:-:...,...
##   spacer_28                     chr12:25227324:-:...,chr12:25227324:-:...,...
##   spacer_41 chr12:25245346:-:...,chr12:25245346:-:...,chr12:25245346:-:...,...
##   spacer_23                     chr12:25227303:-:...,chr12:25227303:-:...,...
##   spacer_24                     chr12:25227304:-:...,chr12:25227304:-:...,...
##   spacer_25                     chr12:25227305:-:...,chr12:25227305:-:...,...
##                   tssAnnotation    hasSNP                 snps
##             <SplitDataFrameList> <logical> <SplitDataFrameList>
##   spacer_29              :...,...     FALSE             :...,...
##   spacer_28              :...,...     FALSE             :...,...
##   spacer_41              :...,...     FALSE             :...,...
##   spacer_23              :...,...     FALSE             :...,...
##   spacer_24              :...,...     FALSE             :...,...
##   spacer_25              :...,...     FALSE             :...,...
##   -------
##   seqinfo: 640 sequences (1 circular) from hg38 genome
##   crisprNuclease: SpCas9
```

One can also sort gRNAs using several annotation columns. For instance, let's sort gRNAs using the DeepHF score, but also by prioritizing first gRNAs that have no 1-mismatch off-targets in coding regions:

```
o <- order(guideSet$n1_c, -guideSet$score_deephf)
# Ordering the GuideSet:
guideSet <- guideSet[o]
head(guideSet)
```

```
## GuideSet object with 6 ranges and 28 metadata columns:
##             seqnames    ranges strand |          protospacer            pam
##                <Rle> <IRanges>  <Rle> |       <DNAStringSet> <DNAStringSet>
##   spacer_29    chr12  25227322      - | AAGAGGAGTACAGTGCAATG            AGG
##   spacer_28    chr12  25227321      - | AGAGGAGTACAGTGCAATGA            GGG
##   spacer_41    chr12  25245343      - | GTAGTTGGAGCTGGTGGCGT            AGG
##   spacer_23    chr12  25227300      - | GGACCAGTACATGAGGACTG            GGG
##   spacer_24    chr12  25227301      - | GGGACCAGTACATGAGGACT            GGG
##   spacer_25    chr12  25227302      - | AGGGACCAGTACATGAGGAC            TGG
##              pam_site  cut_site      region percentGC     polyA     polyC
##             <numeric> <numeric> <character> <numeric> <logical> <logical>
##   spacer_29  25227322  25227325    region_6        45     FALSE     FALSE
##   spacer_28  25227321  25227324    region_6        45     FALSE     FALSE
##   spacer_41  25245343  25245346    region_5        60     FALSE     FALSE
##   spacer_23  25227300  25227303    region_6        55     FALSE     FALSE
##   spacer_24  25227301  25227304    region_6        55     FALSE     FALSE
##   spacer_25  25227302  25227305    region_6        55     FALSE     FALSE
##                 polyG     polyT startingGGGGG        n0        n1        n2
##             <logical> <logical>     <logical> <numeric> <numeric> <numeric>
##   spacer_29     FALSE     FALSE         FALSE         1         0         1
##   spacer_28     FALSE     FALSE         FALSE         1         0         1
##   spacer_41     FALSE     FALSE         FALSE         1         0         1
##   spacer_23     FALSE     FALSE         FALSE         1         0         0
##   spacer_24     FALSE     FALSE         FALSE         1         1         1
##   spacer_25     FALSE     FALSE         FALSE         1         0         2
##                  n0_c      n1_c      n2_c
##             <numeric> <numeric> <numeric>
##   spacer_29         1         0         0
##   spacer_28         1         0         0
##   spacer_41         1         0         0
##   spacer_23         1         0         0
##   spacer_24         1         0         0
##   spacer_25         1         0         1
##                                                      alignments inRepeats
##                                                   <GRangesList> <logical>
##   spacer_29                   chr12:25227322:-,chr6:54770784:+     FALSE
##   spacer_28                  chr12:25227321:-,chr16:70072284:-     FALSE
##   spacer_41                  chr12:25245343:-,chr9:121850427:+     FALSE
##   spacer_23                                   chr12:25227300:-     FALSE
##   spacer_24 chr12:25227301:-,chr6:54770804:+,chr12:131824051:+     FALSE
##   spacer_25  chr12:25227302:-,chr1:114713868:-,chr6:54770803:+     FALSE
##             score_cfd score_mit score_deephf score_deepspcas9
##             <numeric> <numeric>    <numeric>        <numeric>
##   spacer_29  0.555556  1.000000     0.712546         0.761093
##   spacer_28  0.796078  1.000000     0.693555         0.777314
##   spacer_41  0.928339  0.999684     0.692967         0.585668
##   spacer_23  1.000000  1.000000     0.686824         0.710386
##   spacer_24  0.459184  0.621749     0.672358         0.604710
##   spacer_25  0.429499  0.954044     0.672062         0.444870
##                 enzymeAnnotation
##              <SplitDataFrameList>
##   spacer_29 FALSE:FALSE:FALSE:...
##   spacer_28 FALSE:FALSE:FALSE:...
##   spacer_41 FALSE:FALSE:FALSE:...
```

```
##   spacer_23 FALSE:FALSE:FALSE:...
##   spacer_24 FALSE:FALSE:FALSE:...
##   spacer_25 FALSE:FALSE:FALSE:...
##                                                                   geneAnnotation
##                                                             <SplitDataFrameList>
##   spacer_29                    chr12:25227325:-:...,chr12:25227325:-:...,...
##   spacer_28                    chr12:25227324:-:...,chr12:25227324:-:...,...
##   spacer_41 chr12:25245346:-:...,chr12:25245346:-:...,chr12:25245346:-:...,...
##   spacer_23                    chr12:25227303:-:...,chr12:25227303:-:...,...
##   spacer_24                    chr12:25227304:-:...,chr12:25227304:-:...,...
##   spacer_25                    chr12:25227305:-:...,chr12:25227305:-:...,...
##                    tssAnnotation    hasSNP                 snps
##             <SplitDataFrameList> <logical> <SplitDataFrameList>
##   spacer_29             :...,...     FALSE             :...,...
##   spacer_28             :...,...     FALSE             :...,...
##   spacer_41             :...,...     FALSE             :...,...
##   spacer_23             :...,...     FALSE             :...,...
##   spacer_24             :...,...     FALSE             :...,...
##   spacer_25             :...,...     FALSE             :...,...
##   -------
##   seqinfo: 640 sequences (1 circular) from hg38 genome
##   crisprNuclease: SpCas9
```

The `rankSpacers` function is a convenience function that implements our recommended rankings for the SpCas9, enAsCas12a and CasRx nucleases. For a detailed description of our recommended rankings, see the documentation of `rankSpacers` by typing `?rankSpacers`.

If an Ensembl transcript ID is provided, the ranking function will also take into account the position of the gRNA within the target CDS of the transcript ID in the ranking procedure. Our recommendation is to specify the Ensembl canonical transcript as the representative transcript for the gene. In our example, ENST00000311936 is the canonical transcript for KRAS:

```
tx_id <- "ENST00000311936"
guideSet <- rankSpacers(guideSet,
                        tx_id=tx_id)
head(guideSet)
```

```
## GuideSet object with 6 ranges and 36 metadata columns:
##             seqnames    ranges strand |          protospacer            pam
##                <Rle> <IRanges>  <Rle> |       <DNAStringSet> <DNAStringSet>
##   spacer_41    chr12  25245343      - | GTAGTTGGAGCTGGTGGCGT            AGG
##   spacer_42    chr12  25245349      - | CTTGTGGTAGTTGGAGCTGG            TGG
##   spacer_37    chr12  25245275      - | CGAATATGATCCAACAATAG            AGG
##   spacer_40    chr12  25245330      + | CTGAATTAGCTGTATCGTCA            AGG
##   spacer_44    chr12  25245358      - | GAATATAAACTTGTGGTAGT            TGG
##   spacer_38    chr12  25245283      + | ACAAGATTTACCTCTATTGT            TGG
##              pam_site  cut_site      region percentGC     polyA     polyC
##             <numeric> <numeric> <character> <numeric> <logical> <logical>
##   spacer_41  25245343  25245346    region_5        60     FALSE     FALSE
##   spacer_42  25245349  25245352    region_5        55     FALSE     FALSE
##   spacer_37  25245275  25245278    region_5        35     FALSE     FALSE
##   spacer_40  25245330  25245327    region_5        40     FALSE     FALSE
##   spacer_44  25245358  25245361    region_5        30     FALSE     FALSE
##   spacer_38  25245283  25245280    region_5        30     FALSE     FALSE
##                 polyG     polyT startingGGGGG        n0        n1        n2
##             <logical> <logical>     <logical> <numeric> <numeric> <numeric>
```

```
## spacer_41     FALSE     FALSE         FALSE         1         0         1
## spacer_42     FALSE     FALSE         FALSE         1         1         0
## spacer_37     FALSE     FALSE         FALSE         1         0         0
## spacer_40     FALSE     FALSE         FALSE         1         0         0
## spacer_44     FALSE     FALSE         FALSE         1         1         0
## spacer_38     FALSE     FALSE         FALSE         1         0         1
##                  n0_c      n1_c      n2_c                    alignments
##             <numeric> <numeric> <numeric>                 <GRangesList>
## spacer_41           1         0         0 chr12:25245343:-,chr9:121850427:+
## spacer_42           1         0         0  chr12:25245349:-,chr6:54770618:+
## spacer_37           1         0         0                  chr12:25245275:-
## spacer_40           1         0         0                  chr12:25245330:+
## spacer_44           1         0         0  chr12:25245358:-,chr6:54770609:+
## spacer_38           1         0         0  chr12:25245283:+,chr8:38993077:+
##             inRepeats score_cfd score_mit score_deephf score_deepspcas9
##             <logical> <numeric> <numeric>    <numeric>        <numeric>
## spacer_41     FALSE  0.928339  0.999684     0.692967         0.585668
## spacer_42     FALSE  0.500000  0.547046     0.644286         0.525602
## spacer_37     FALSE  1.000000  1.000000     0.609294         0.465393
## spacer_40     FALSE  1.000000  1.000000     0.551647         0.390722
## spacer_44     FALSE  0.777778  0.759301     0.433265         0.255677
## spacer_38     FALSE  0.819820  0.998398     0.414292         0.172458
##                  enzymeAnnotation
##              <SplitDataFrameList>
## spacer_41 FALSE:FALSE:FALSE:...
## spacer_42 FALSE:FALSE:FALSE:...
## spacer_37 FALSE:FALSE:FALSE:...
## spacer_40 FALSE:FALSE:FALSE:...
## spacer_44 FALSE:FALSE:FALSE:...
## spacer_38 FALSE:FALSE:FALSE:...
##                                                          geneAnnotation
##                                                    <SplitDataFrameList>
## spacer_41 chr12:25245346:-:...,chr12:25245346:-:...,chr12:25245346:-:...,...
## spacer_42 chr12:25245352:-:...,chr12:25245352:-:...,chr12:25245352:-:...,...
## spacer_37 chr12:25245278:-:...,chr12:25245278:-:...,chr12:25245278:-:...,...
## spacer_40 chr12:25245327:+:...,chr12:25245327:+:...,chr12:25245327:+:...,...
## spacer_44 chr12:25245361:-:...,chr12:25245361:-:...,chr12:25245361:-:...,...
## spacer_38 chr12:25245280:+:...,chr12:25245280:+:...,chr12:25245280:+:...,...
##                 tssAnnotation    hasSNP                 snps percentCDS
##             <SplitDataFrameList> <logical> <SplitDataFrameList>  <numeric>
## spacer_41             :...,...     FALSE             :...,...        6.9
## spacer_42             :...,...     FALSE             :...,...        5.8
## spacer_37             :...,...     FALSE             :...,...       18.9
## spacer_40             :...,...     FALSE             :...,...       10.2
## spacer_44             :...,...     FALSE             :...,...        4.2
## spacer_38             :...,...     FALSE             :...,...       18.5
##             percentCodingIsoforms isCommonCodingExon score_cds score_exon
##                         <numeric>          <logical> <numeric>  <numeric>
## spacer_41                     100               TRUE         1          1
## spacer_42                     100               TRUE         1          1
## spacer_37                     100               TRUE         1          1
## spacer_40                     100               TRUE         1          1
## spacer_44                     100               TRUE         1          1
## spacer_38                     100               TRUE         1          1
```

```
##                 round score_composite      rank
##             <numeric>       <numeric> <integer>
##   spacer_41         1            41.5         1
##   spacer_42         1            33.0         2
##   spacer_37         1            27.5         3
##   spacer_40         1            20.5         4
##   spacer_44         1            12.5         5
##   spacer_38         1             9.0         6
##   -------
##   seqinfo: 640 sequences (1 circular) from hg38 genome
##   crisprNuclease: SpCas9
```

### Session Info

```
sessionInfo()
```

```
## R version 4.2.1 (2022-06-23)
## Platform: x86_64-apple-darwin17.0 (64-bit)
## Running under: macOS Catalina 10.15.7
##
## Matrix products: default
## BLAS:   /Library/Frameworks/R.framework/Versions/4.2/Resources/lib/libRblas.0.dylib
## LAPACK: /Library/Frameworks/R.framework/Versions/4.2/Resources/lib/libRlapack.dylib
##
## locale:
## [1] en_US.UTF-8/en_US.UTF-8/en_US.UTF-8/C/en_US.UTF-8/en_US.UTF-8
##
## attached base packages:
## [1] stats4    stats     graphics  grDevices utils     datasets  methods
## [8] base
##
## other attached packages:
##  [1] BSgenome.Hsapiens.UCSC.hg38_1.4.4 BSgenome_1.65.2
##  [3] rtracklayer_1.57.0                Biostrings_2.65.2
##  [5] XVector_0.37.0                    GenomicRanges_1.49.1
##  [7] GenomeInfoDb_1.33.5               IRanges_2.31.2
##  [9] S4Vectors_0.35.1                  crisprDesignData_0.99.17
## [11] crisprDesign_0.99.133             crisprScore_1.1.14
## [13] crisprScoreData_1.1.3             ExperimentHub_2.5.0
## [15] AnnotationHub_3.5.0               BiocFileCache_2.5.0
## [17] dbplyr_2.2.1                      BiocGenerics_0.43.1
## [19] crisprBowtie_1.1.1                crisprBase_1.1.5
## [21] crisprVerse_0.99.8                rmarkdown_2.15.2
##
## loaded via a namespace (and not attached):
##  [1] rjson_0.2.21                  ellipsis_0.3.2
##  [3] Rbowtie_1.37.0                bit64_4.0.5
##  [5] lubridate_1.8.0               interactiveDisplayBase_1.35.0
##  [7] AnnotationDbi_1.59.1          fansi_1.0.3
##  [9] xml2_1.3.3                    codetools_0.2-18
## [11] cachem_1.0.6                  knitr_1.40
## [13] jsonlite_1.8.0                Rsamtools_2.13.4
## [15] png_0.1-7                     shiny_1.7.2
```

```
## [17] BiocManager_1.30.18           readr_2.1.2
## [19] compiler_4.2.1                httr_1.4.4
## [21] basilisk_1.9.2                assertthat_0.2.1
## [23] Matrix_1.4-1                  fastmap_1.1.0
## [25] cli_3.3.0                     later_1.3.0
## [27] htmltools_0.5.3               prettyunits_1.1.1
## [29] tools_4.2.1                   glue_1.6.2
## [31] GenomeInfoDbData_1.2.8        dplyr_1.0.9
## [33] rappdirs_0.3.3                tinytex_0.41
## [35] Rcpp_1.0.9                    Biobase_2.57.1
## [37] vctrs_0.4.1                   crisprBwa_1.1.3
## [39] xfun_0.32                     stringr_1.4.1
## [41] mime_0.12                     lifecycle_1.0.1
## [43] restfulr_0.0.15               XML_3.99-0.10
## [45] zlibbioc_1.43.0               basilisk.utils_1.9.1
## [47] vroom_1.5.7                   VariantAnnotation_1.43.3
## [49] hms_1.1.2                     promises_1.2.0.1
## [51] MatrixGenerics_1.9.1          parallel_4.2.1
## [53] SummarizedExperiment_1.27.1   RMariaDB_1.2.2
## [55] yaml_2.3.5                    curl_4.3.2
## [57] memoise_2.0.1                 reticulate_1.25
## [59] biomaRt_2.53.2                stringi_1.7.8
## [61] RSQLite_2.2.16                BiocVersion_3.16.0
## [63] highr_0.9                     BiocIO_1.7.1
## [65] randomForest_4.7-1.1          GenomicFeatures_1.49.6
## [67] filelock_1.0.2                BiocParallel_1.31.12
## [69] rlang_1.0.4                   pkgconfig_2.0.3
## [71] matrixStats_0.62.0            bitops_1.0-7
## [73] evaluate_0.16                 lattice_0.20-45
## [75] purrr_0.3.4                   GenomicAlignments_1.33.1
## [77] bit_4.0.4                     tidyselect_1.1.2
## [79] magrittr_2.0.3                R6_2.5.1
## [81] generics_0.1.3                DelayedArray_0.23.1
## [83] DBI_1.1.3                     pillar_1.8.1
## [85] KEGGREST_1.37.3               RCurl_1.98-1.8
## [87] tibble_3.1.8                  dir.expiry_1.5.0
## [89] crayon_1.5.1                  utf8_1.2.2
## [91] tzdb_0.3.0                    progress_1.2.2
## [93] grid_4.2.1                    blob_1.2.3
## [95] digest_0.6.29                 xtable_1.8-4
## [97] httpuv_1.6.5                  Rbwa_1.1.0
```
