## Supplementary material for "The crisprVerse: a comprehensive Bioconductor ecosystem for the design of CRISPR guide RNAs across nucleases and technologies": Tutorial6_CRISPRko_Design_Cas12a

### Design gRNAs for CRISPRko with the AsCas12a nuclease

#### Introduction

In this tutorial, we design CRISPR/Cas12a gRNAs targeting the coding sequence of the human gene KRAS. In particular, we use the AsCas12a nuclease. The tutorial is very similar to the gRNA design for Cas9.

#### Terminology

Before we start designing gRNAs, we first introduce some terminology that will be useful throughout this and subsequent tutorials. CRISPR nucleases require two binding components for cleavage. First, the nuclease needs to recognize a constant nucleotide motif in the target DNA called the protospacer adjacent motif (PAM) sequence. Second, the gRNA, which guides the nuclease to the target sequence, needs to bind to a complementary sequence adjacent to the PAM sequence, called the **protospacer** sequence. The latter can be thought of as a variable binding motif that can be specified by designing corresponding gRNA sequences.

The **spacer** sequence is used in the gRNA construct to guide the CRISPR nuclease to the target **protospacer** sequence in the host genome. While a gRNA spacer sequence may not always uniquely target the host genome (i.e. it may map to multiple protospacers in the host genome), we can, for a given reference genome, uniquely identify a protospacer sequence with a combination of 3 attributes:

- `chr`: chromosome name
- `strand`: forward (+) or reverse (-)
- `pam_site`: genomic coordinate of the first nucleotide of the nuclease-specific PAM sequence; for AsCas12a this is the first "T" in the TTTV PAM sequence

For CRISPRko applications, we use an additional genomic coordinate, called `cut_site`, to represent where the double-stranded break (DSB) occurs. For enAsCas12a, the 5nt 5' overhang dsDNA break will cause a cut 19nt after the PAM sequence on the targeted strand, and 23nt after the PAM sequence on the opposite strand (PAM-distal editing).

#### Installation

See the Installation tutorial to learn how to install the packages necessary for this tutorial: `crisprDesign`, `crisprDesignData`

#### End-to-end gRNA design workflow

We first start by loading the crisprVerse packages needed for this tutorial:

```
library(crisprBase)
library(crisprDesign)
library(crisprDesignData)
```

We will also load the `BSgenome` package containing DNA sequences for the hg38 genome:

```
library(BSgenome.Hsapiens.UCSC.hg38)
```

#### Nuclease specification

We first load the AsCas12a nuclease object from the crisprBase package:

```
data(AsCas12a, package="crisprBase")
AsCas12a
```

```
## Class: CrisprNuclease
##   Name: AsCas12a
##   Target type: DNA
##   Metadata: list of length 1
##   PAMs: TTTV
##   Weights: 1
##   Spacer length: 23
##   PAM side: 5prime
##     Distance from PAM: 0
##   Prototype protospacers: 5'--[TTTV]SSSSSSSSSSSSSSSSSSSSSSS--3'
```

To learn how to specify a custom nuclease, see the nuclease tutorial.

The motif (TTTV) represents the recognized PAM sequences by AsCas12a, and the weights indicate a recognition score. The single canonical PAM sequence for AsCas12a has a weight of 1.

The spacer sequence is located on the 3-prime end with respect to the PAM sequence, and the default spacer sequence length is 23 nucleotides. If necessary, one can change the spacer length using the function spacerLength from crisprBase. We can inspect the protospacer construct by using prototypeSequence:

```
prototypeSequence(AsCas12a)
```

```
## [1] "5'--[TTTV]SSSSSSSSSSSSSSSSSSSSSSS--3'"
```

#### Specification of the target DNA sequence (KRAS CDS)

Since we aim to design gRNAs that knock out the human KRAS gene, we first need to retrieve the DNA sequence of the coding region (CDS) of KRAS. W e show in the gene annotation tutorial how to build convenient gene model objects that allows to quickly access gene-specific sequences. Here, we obtain from crisprDesignData a GRangesList object that defines the genomic coordinates (in hg38 coordinates) of coding genes in the human genome:

```
data(txdb_human, package="crisprDesignData")
```

The queryTxObject function allows us to query this object for a specific gene and feature. Here, we obtain a GRanges object containing the CDS coordinates of KRAS:

```
gr <- queryTxObject(txObject=txdb_human,
                    featureType="cds",
                    queryColumn="gene_symbol",
                    queryValue="KRAS")
```

To simplify our design, we will only consider exons that constitute the primary transcript of KRAS (transcript ID ENST00000311936).

```
gr <- gr[gr$tx_id == "ENST00000311936"]
```

Optionally, we could also adjust the arguments in our call to queryTxObject to retrieve those transcript-specific coordinates:

```r
gr <- queryTxObject(txObject=txObject,
                    featureType="cds",
                    queryColumn="tx_id",
                    queryValue="ENST00000311936")
```

#### Finding spacer sequences targeting KRAS

`findSpacers` is the main function of `crisprDesign` for obtaining all possible spacer sequences that target protospacers located in our target DNA sequence(s). If a `GRanges` object is provided as input, a `BSgenome` object (an object that contains sequences of a reference genome) must be provided as well:

```r
bsgenome <- BSgenome.Hsapiens.UCSC.hg38
guideSet <- findSpacers(gr,
                        bsgenome=bsgenome,
                        crisprNuclease=AsCas12a)
guideSet
```

```
## GuideSet object with 34 ranges and 5 metadata columns:
##              seqnames    ranges strand |             protospacer           pam
##                 <Rle> <IRanges>  <Rle> |          <DNAStringSet> <DNAStringSet>
##    spacer_1     chr12  25209794      + | CATAATTACACACTTTGTCTTTG          TTTA
##    spacer_2     chr12  25209811      + | TCTTTGACTTCTTTTTCTTCTTT          TTTG
##    spacer_3     chr12  25209817      + | ACTTCTTTTTCTTCTTTTTACCA          TTTG
##    spacer_4     chr12  25209828      + | TTCTTTTTACCATCTTTGCTCAT          TTTC
##    spacer_5     chr12  25209837      + | CCATCTTTGCTCATCTTTTCTTT          TTTA
##         ...       ...       ...    ... .                     ...           ...
##   spacer_30     chr12  25227376      + | TCCATCAATTACTACTTGCTTCC          TTTC
##   spacer_31     chr12  25227427      - | TCCCTTCTCAGGATTCCTACAGG          TTTC
##   spacer_32     chr12  25245269      + | CCTCTATTGTTGGATCATATTCG          TTTA
##   spacer_33     chr12  25245303      - | TGGACGAATATGATCCAACAATA          TTTG
##   spacer_34     chr12  25245406      - | TTATAAGGCCTGCTGAAAATGAC          TTTA
##               pam_site  cut_site      region
##              <numeric> <numeric> <character>
##    spacer_1   25209794  25209818    region_8
##    spacer_2   25209811  25209835    region_8
##    spacer_3   25209817  25209841    region_8
##    spacer_4   25209828  25209852    region_8
##    spacer_5   25209837  25209861    region_8
##         ...        ...       ...         ...
##   spacer_30   25227376  25227400    region_6
##   spacer_31   25227427  25227403    region_6
##   spacer_32   25245269  25245293    region_5
##   spacer_33   25245303  25245279    region_5
##   spacer_34   25245406  25245382    region_5
##   -------
##   seqinfo: 640 sequences (1 circular) from hg38 genome
##   crisprNuclease: AsCas12a
```

This function returns a `GuideSet` object that stores the genomic coordinates (PAM sites) for all spacer sequences found in the regions provided by `gr`. The `GuideSet` object is an extension of a `GenomicRanges` object that stores additional information about gRNAs.

There are several accessor functions we can use to extract information about the spacer sequences in `guideSet`, and here are a few examples with their corresponding outputs:

```r
spacers(guideSet)
```

```
## DNAStringSet object of length 34:
##      width seq                                               names
##  [1]    23 CATAATTACACACTTTGTCTTTG                           spacer_1
##  [2]    23 TCTTTGACTTCTTTTTCTTCTTT                           spacer_2
##  [3]    23 ACTTCTTTTTCTTCTTTTTACCA                           spacer_3
##  [4]    23 TTCTTTTTACCATCTTTGCTCAT                           spacer_4
##  [5]    23 CCATCTTTGCTCATCTTTTCTTT                           spacer_5
##  ...   ... ...
## [30]    23 TCCATCAATTACTACTTGCTTCC                           spacer_30
## [31]    23 TCCCTTCTCAGGATTCCTACAGG                           spacer_31
## [32]    23 CCTCTATTGTTGGATCATATTCG                           spacer_32
## [33]    23 TGGACGAATATGATCCAACAATA                           spacer_33
## [34]    23 TTATAAGGCCTGCTGAAAATGAC                           spacer_34
```

```r
protospacers(guideSet)
```

```
## DNAStringSet object of length 34:
##      width seq                                               names
##  [1]    23 CATAATTACACACTTTGTCTTTG                           spacer_1
##  [2]    23 TCTTTGACTTCTTTTTCTTCTTT                           spacer_2
##  [3]    23 ACTTCTTTTTCTTCTTTTTACCA                           spacer_3
##  [4]    23 TTCTTTTTACCATCTTTGCTCAT                           spacer_4
##  [5]    23 CCATCTTTGCTCATCTTTTCTTT                           spacer_5
##  ...   ... ...
## [30]    23 TCCATCAATTACTACTTGCTTCC                           spacer_30
## [31]    23 TCCCTTCTCAGGATTCCTACAGG                           spacer_31
## [32]    23 CCTCTATTGTTGGATCATATTCG                           spacer_32
## [33]    23 TGGACGAATATGATCCAACAATA                           spacer_33
## [34]    23 TTATAAGGCCTGCTGAAAATGAC                           spacer_34
```

```r
pams(guideSet)
```

```
## DNAStringSet object of length 34:
##      width seq                                               names
##  [1]     4 TTTA                                              spacer_1
##  [2]     4 TTTG                                              spacer_2
##  [3]     4 TTTG                                              spacer_3
##  [4]     4 TTTC                                              spacer_4
##  [5]     4 TTTA                                              spacer_5
##  ...   ... ...
## [30]     4 TTTC                                              spacer_30
## [31]     4 TTTC                                              spacer_31
## [32]     4 TTTA                                              spacer_32
## [33]     4 TTTG                                              spacer_33
## [34]     4 TTTA                                              spacer_34
```

```r
head(pamSites(guideSet))
```

```
## spacer_1 spacer_2 spacer_3 spacer_4 spacer_5 spacer_6
## 25209794 25209811 25209817 25209828 25209837 25209846
```

```r
head(cutSites(guideSet))
```

```
## spacer_1 spacer_2 spacer_3 spacer_4 spacer_5 spacer_6
## 25209818 25209835 25209841 25209852 25209861 25209870
```

#### Characterizing gRNA spacer sequences

There are specific spacer sequence features, independent of the genomic context of the protospacer sequence, that can reduce or even eliminate gRNA activity:

- **Poly-T stretches**: four or more consecutive T nucleotides in the spacer sequence may act as a transcriptional termination signal for the U6 promoter.
- **Self-complementarity**: complementary sites with the gRNA backbone can compete with the targeted genomic sequence.
- **Percent GC**: gRNAs with GC content between 20% and 80% are preferred.

Use the function `addSequenceFeatures` to evaluate the spacer sequences with respect to these characteristics and add the results to the `GuideSet` object:

```
guideSet <- addSequenceFeatures(guideSet)
head(guideSet)
```

```
## GuideSet object with 6 ranges and 11 metadata columns:
##            seqnames    ranges strand |            protospacer            pam
##               <Rle> <IRanges>  <Rle> |         <DNAStringSet> <DNAStringSet>
##   spacer_1    chr12  25209794      + | CATAATTACACACTTTGTCTTTG           TTTA
##   spacer_2    chr12  25209811      + | TCTTTGACTTCTTTTTCTTCTTT           TTTG
##   spacer_3    chr12  25209817      + | ACTTCTTTTTCTTCTTTTTACCA           TTTG
##   spacer_4    chr12  25209828      + | TTCTTTTTACCATCTTTGCTCAT           TTTC
##   spacer_5    chr12  25209837      + | CCATCTTTGCTCATCTTTTCTTT           TTTA
##   spacer_6    chr12  25209846      + | CTCATCTTTTCTTTATGTTTTCG           TTTG
##             pam_site  cut_site      region percentGC     polyA     polyC
##            <numeric> <numeric> <character> <numeric> <logical> <logical>
##   spacer_1  25209794  25209818    region_8      30.4     FALSE     FALSE
##   spacer_2  25209811  25209835    region_8      26.1     FALSE     FALSE
##   spacer_3  25209817  25209841    region_8      26.1     FALSE     FALSE
##   spacer_4  25209828  25209852    region_8      30.4     FALSE     FALSE
##   spacer_5  25209837  25209861    region_8      34.8     FALSE     FALSE
##   spacer_6  25209846  25209870    region_8      30.4     FALSE     FALSE
##                polyG     polyT startingGGGGG
##            <logical> <logical>     <logical>
##   spacer_1     FALSE     FALSE         FALSE
##   spacer_2     FALSE      TRUE         FALSE
##   spacer_3     FALSE      TRUE         FALSE
##   spacer_4     FALSE      TRUE         FALSE
##   spacer_5     FALSE      TRUE         FALSE
##   spacer_6     FALSE      TRUE         FALSE
##   -------
##   seqinfo: 640 sequences (1 circular) from hg38 genome
##   crisprNuclease: AsCas12a
```

```
guideSet <- addSpacerAlignments(guideSet,
                                aligner="bowtie",
                                aligner_index=bowtie_index,
                                bsgenome=BSgenome.Hsapiens.UCSC.hg38,
                                n_mismatches=2,
                                txObject=txdb_human)
```

```
## [runCrisprBowtie] Using BSgenome.Hsapiens.UCSC.hg38
## [runCrisprBowtie] Searching for AsCas12a protospacers
```

```
guideSet
```

```
## GuideSet object with 34 ranges and 18 metadata columns:
##               seqnames    ranges strand |            protospacer            pam
##                  <Rle> <IRanges>  <Rle> |         <DNAStringSet> <DNAStringSet>
##    spacer_1      chr12  25209794      + | CATAATTACACACTTTGTCTTTG           TTTA
##    spacer_2      chr12  25209811      + | TCTTTGACTTCTTTTTCTTCTTT           TTTG
##    spacer_3      chr12  25209817      + | ACTTCTTTTTCTTCTTTTTACCA           TTTG
##    spacer_4      chr12  25209828      + | TTCTTTTTACCATCTTTGCTCAT           TTTC
##    spacer_5      chr12  25209837      + | CCATCTTTGCTCATCTTTTCTTT           TTTA
##         ...        ...       ...    ... .                    ...            ...
##   spacer_30      chr12  25227376      + | TCCATCAATTACTACTTGCTTCC           TTTC
##   spacer_31      chr12  25227427      - | TCCCTTCTCAGGATTCCTACAGG           TTTC
##   spacer_32      chr12  25245269      + | CCTCTATTGTTGGATCATATTCG           TTTA
##   spacer_33      chr12  25245303      - | TGGACGAATATGATCCAACAATA           TTTG
##   spacer_34      chr12  25245406      - | TTATAAGGCCTGCTGAAAATGAC           TTTA
##                pam_site  cut_site      region percentGC     polyA     polyC
##               <numeric> <numeric> <character> <numeric> <logical> <logical>
##    spacer_1  25209794  25209818    region_8      30.4     FALSE     FALSE
##    spacer_2  25209811  25209835    region_8      26.1     FALSE     FALSE
##    spacer_3  25209817  25209841    region_8      26.1     FALSE     FALSE
##    spacer_4  25209828  25209852    region_8      30.4     FALSE     FALSE
##    spacer_5  25209837  25209861    region_8      34.8     FALSE     FALSE
```

```
##          ...        ...        ...          ...       ...       ...       ...
##   spacer_30  25227376  25227400    region_6      39.1     FALSE     FALSE
##   spacer_31  25227427  25227403    region_6      52.2     FALSE     FALSE
##   spacer_32  25245269  25245293    region_5      39.1     FALSE     FALSE
##   spacer_33  25245303  25245279    region_5      34.8     FALSE     FALSE
##   spacer_34  25245406  25245382    region_5      39.1      TRUE     FALSE
##                  polyG     polyT startingGGGGG        n0        n1        n2
##              <logical> <logical>     <logical> <numeric> <numeric> <numeric>
##    spacer_1     FALSE     FALSE         FALSE         1         1         0
##    spacer_2     FALSE      TRUE         FALSE         1         0         1
##    spacer_3     FALSE      TRUE         FALSE         1         0         1
##    spacer_4     FALSE      TRUE         FALSE         1         1         0
##    spacer_5     FALSE      TRUE         FALSE         1         0         0
##         ...       ...       ...           ...       ...       ...       ...
##   spacer_30     FALSE     FALSE         FALSE         2         0         0
##   spacer_31     FALSE     FALSE         FALSE         1         0         0
##   spacer_32     FALSE     FALSE         FALSE         1         0         0
##   spacer_33     FALSE     FALSE         FALSE         1         1         0
##   spacer_34     FALSE     FALSE         FALSE         1         0         0
##                   n0_c      n1_c      n2_c                        alignments
##              <numeric> <numeric> <numeric>                     <GRangesList>
##    spacer_1          1         0         0  chr12:25209794:+,chr6:54771134:-
##    spacer_2          1         0         0 chr12:25209811:+,chr6:117625992:-
##    spacer_3          1         0         0  chr12:25209817:+,chr5:54961047:-
##    spacer_4          1         0         0  chr12:25209828:+,chr6:54771104:-
##    spacer_5          1         0         0                  chr12:25209837:+
##         ...        ...       ...       ...                               ...
##   spacer_30          1         0         0  chr12:25227376:+,chr6:54770730:-
##   spacer_31          1         0         0                  chr12:25227427:-
##   spacer_32          1         0         0                  chr12:25245269:+
##   spacer_33          1         0         0  chr12:25245303:-,chr6:54770664:+
##   spacer_34          1         0         0                  chr12:25245406:-
##   -------
##   seqinfo: 640 sequences (1 circular) from hg38 genome
##   crisprNuclease: AsCas12a
```

Several columns were added to the `GuideSet` object that summarize the number of on- and off-targets for each spacer sequence, and take genomic context into account:

- **n0, n1, n2, n3**: specify the number of alignments with 0, 1, 2 and 3 mismatches, respectively.
- **n0_c, n1_c, n2_c, n3_c**: specify the number of alignments in a coding region, with 0, 1, 2 and 3 mismatches, respectively.
- **n0_p, n1_p, n2_p, n3_p**: specify the number of alignments in a promoter region of a coding gene, with 0, 1, 2 and 3 mismatches, respectively.

Our `guideSet` now has columns of the first two categories, up to 2 mismatches (the value passed to `n_mismatches`); had we also supplied a `GRanges` of TSS coordinates to the `tssObject` argument, our `guideSet` would include columns in the last category.

To inspect individual on- and off-targets and their context, one can use the `alignments` function, which returns a table of all genomic alignments stored in the `GuideSet` object:

```
alignments(guideSet)
```

```
## GRanges object with 50 ranges and 14 metadata columns:
##              seqnames    ranges strand |                  spacer
##                 <Rle> <IRanges>  <Rle> |          <DNAStringSet>
```

```
##    spacer_1    chr12  25209794      + | CATAATTACACACTTTGTCTTTG
##    spacer_1     chr6  54771134      - | CATAATTACACACTTTGTCTTTG
##    spacer_2    chr12  25209811      + | TCTTTGACTTCTTTTTCTTCTTT
##    spacer_2     chr6 117625992      - | TCTTTGACTTCTTTTTCTTCTTT
##    spacer_3    chr12  25209817      + | ACTTCTTTTTCTTCTTTTTACCA
##         ...      ...       ...    ... .                     ...
##   spacer_31    chr12  25227427      - | TCCCTTCTCAGGATTCCTACAGG
##   spacer_32    chr12  25245269      + | CCTCTATTGTTGGATCATATTCG
##   spacer_33    chr12  25245303      - | TGGACGAATATGATCCAACAATA
##   spacer_33     chr6  54770664      + | TGGACGAATATGATCCAACAATA
##   spacer_34    chr12  25245406      - | TTATAAGGCCTGCTGAAAATGAC
##                          protospacer           pam  pam_site n_mismatches
##                       <DNAStringSet> <DNAStringSet> <numeric>    <integer>
##    spacer_1 CATAATTACACACTTTGTCTTTG           TTTA  25209794            0
##    spacer_1 CATAATTACACACTTTGTCATTG           TTTA  54771134            1
##    spacer_2 TCTTTGACTTCTTTTTCTTCTTT           TTTG  25209811            0
##    spacer_2 TCTTTCCCTTCTTTTTCTTCTTT           TTTG 117625992            2
##    spacer_3 ACTTCTTTTTCTTCTTTTTACCA           TTTG  25209817            0
##         ...                     ...            ...       ...          ...
##   spacer_31 TCCCTTCTCAGGATTCCTACAGG           TTTC  25227427            0
##   spacer_32 CCTCTATTGTTGGATCATATTCG           TTTA  25245269            0
##   spacer_33 TGGACGAATATGATCCAACAATA           TTTG  25245303            0
##   spacer_33 TGGACCAATATGATCCAACAATA           TTTG  54770664            1
##   spacer_34 TTATAAGGCCTGCTGAAAATGAC           TTTA  25245406            0
##             canonical  cut_site         cds    fiveUTRs   threeUTRs       exons
##             <logical> <numeric> <character> <character> <character> <character>
##    spacer_1      TRUE  25209818        KRAS        <NA>        KRAS        KRAS
##    spacer_1      TRUE  54771110        <NA>        <NA>        <NA>      KRASP1
##    spacer_2      TRUE  25209835        KRAS        <NA>        KRAS        KRAS
##    spacer_2      TRUE 117625968        <NA>        <NA>        <NA>        <NA>
##    spacer_3      TRUE  25209841        KRAS        <NA>        KRAS        KRAS
##         ...       ...       ...         ...         ...         ...         ...
##   spacer_31      TRUE  25227403        KRAS        <NA>        <NA>        KRAS
##   spacer_32      TRUE  25245293        KRAS        <NA>        <NA>        KRAS
##   spacer_33      TRUE  25245279        KRAS        <NA>        <NA>        KRAS
##   spacer_33      TRUE  54770688        <NA>        <NA>        <NA>      KRASP1
##   spacer_34      TRUE  25245382        KRAS        <NA>        <NA>        KRAS
##                 introns  intergenic intergenic_distance
##             <character> <character>           <integer>
##    spacer_1        <NA>        <NA>                <NA>
##    spacer_1        <NA>        <NA>                <NA>
##    spacer_2        <NA>        <NA>                <NA>
##    spacer_2        <NA>       NEPNP                7737
##    spacer_3        <NA>        <NA>                <NA>
##         ...         ...         ...                 ...
##   spacer_31        KRAS        <NA>                <NA>
##   spacer_32        <NA>        <NA>                <NA>
##   spacer_33        <NA>        <NA>                <NA>
##   spacer_33        <NA>        <NA>                <NA>
##   spacer_34        <NA>        <NA>                <NA>
##   -------
##   seqinfo: 25 sequences (1 circular) from hg38 genome
```

Similarly, the functions `onTargets` and `offTargets` return on-target alignments (no mismatches) and off-target alignments (having at least one mismatch), respectively. See `?addSpacerAlignments` for more details about the different options.

We note that gRNAs that align to hundreds of different locations are highly unspecific and undesirable. This can also cause `addSpacerAlignments` to be slow. The function `addSpacerAlignmentsIterative` is an iterative version of `addSpacerAlignments` that curtails alignment searches for gRNAs having more hits than the user-defined threshold. See `?addSpacerAlignmentsIterative` for more information.

##### Removing repeat elements

Many promiscuous protospacer sequences occur in repeats or low-complexity DNA sequences (regions identified by RepeatMasker). These sequences are usually not of interest due to their low specificity, and can be easily removed with `removeRepeats`:

```
data("gr.repeats.hg38", package="crisprDesignData")
guideSet <- removeRepeats(guideSet,
                          gr.repeats=gr.repeats.hg38)
```

##### On-target scoring (gRNA efficiency)

`addOnTargetScores` adds scores from on-target efficiency algorithms specified by the `methods` argument (or all available methods if `NULL`) available in the R package crisprScore and appends them to the `GuideSet`:

```
guideSet <- addOnTargetScores(guideSet,
                              methods=c("deepcpf1"))
guideSet
```

```
## GuideSet object with 34 ranges and 20 metadata columns:
##              seqnames    ranges strand |            protospacer           pam
##                 <Rle> <IRanges>  <Rle> |         <DNAStringSet> <DNAStringSet>
##    spacer_1     chr12  25209794      + | CATAATTACACACTTTGTCTTTG          TTTA
##    spacer_2     chr12  25209811      + | TCTTTGACTTCTTTTTCTTCTTT          TTTG
##    spacer_3     chr12  25209817      + | ACTTCTTTTTCTTCTTTTTACCA          TTTG
##    spacer_4     chr12  25209828      + | TTCTTTTTACCATCTTTGCTCAT          TTTC
##    spacer_5     chr12  25209837      + | CCATCTTTGCTCATCTTTTCTTT          TTTA
##         ...       ...       ...    ... .                    ...           ...
##   spacer_30     chr12  25227376      + | TCCATCAATTACTACTTGCTTCC          TTTC
##   spacer_31     chr12  25227427      - | TCCCTTCTCAGGATTCCTACAGG          TTTC
##   spacer_32     chr12  25245269      + | CCTCTATTGTTGGATCATATTCG          TTTA
##   spacer_33     chr12  25245303      - | TGGACGAATATGATCCAACAATA          TTTG
##   spacer_34     chr12  25245406      - | TTATAAGGCCTGCTGAAAATGAC          TTTA
##               pam_site  cut_site      region percentGC     polyA     polyC
##              <numeric> <numeric> <character> <numeric> <logical> <logical>
##    spacer_1  25209794  25209818    region_8      30.4     FALSE     FALSE
##    spacer_2  25209811  25209835    region_8      26.1     FALSE     FALSE
##    spacer_3  25209817  25209841    region_8      26.1     FALSE     FALSE
##    spacer_4  25209828  25209852    region_8      30.4     FALSE     FALSE
##    spacer_5  25209837  25209861    region_8      34.8     FALSE     FALSE
##         ...       ...       ...         ...       ...       ...       ...
##   spacer_30  25227376  25227400    region_6      39.1     FALSE     FALSE
##   spacer_31  25227427  25227403    region_6      52.2     FALSE     FALSE
##   spacer_32  25245269  25245293    region_5      39.1     FALSE     FALSE
##   spacer_33  25245303  25245279    region_5      34.8     FALSE     FALSE
##   spacer_34  25245406  25245382    region_5      39.1      TRUE     FALSE
##                  polyG     polyT startingGGGGG        n0        n1        n2
##              <logical> <logical>     <logical> <numeric> <numeric> <numeric>
```

```
##    spacer_1     FALSE     FALSE         FALSE         1         1         0
##    spacer_2     FALSE      TRUE         FALSE         1         0         1
##    spacer_3     FALSE      TRUE         FALSE         1         0         1
##    spacer_4     FALSE      TRUE         FALSE         1         1         0
##    spacer_5     FALSE      TRUE         FALSE         1         0         0
##         ...       ...       ...           ...       ...       ...       ...
##   spacer_30     FALSE     FALSE         FALSE         2         0         0
##   spacer_31     FALSE     FALSE         FALSE         1         0         0
##   spacer_32     FALSE     FALSE         FALSE         1         0         0
##   spacer_33     FALSE     FALSE         FALSE         1         1         0
##   spacer_34     FALSE     FALSE         FALSE         1         0         0
##                  n0_c      n1_c      n2_c                       alignments
##             <numeric> <numeric> <numeric>                    <GRangesList>
##    spacer_1         1         0         0  chr12:25209794:+,chr6:54771134:-
##    spacer_2         1         0         0 chr12:25209811:+,chr6:117625992:-
##    spacer_3         1         0         0  chr12:25209817:+,chr5:54961047:-
##    spacer_4         1         0         0  chr12:25209828:+,chr6:54771104:-
##    spacer_5         1         0         0                  chr12:25209837:+
##         ...       ...       ...       ...                               ...
##   spacer_30         1         0         0  chr12:25227376:+,chr6:54770730:-
##   spacer_31         1         0         0                  chr12:25227427:-
##   spacer_32         1         0         0                  chr12:25245269:+
##   spacer_33         1         0         0  chr12:25245303:-,chr6:54770664:+
##   spacer_34         1         0         0                  chr12:25245406:-
##             inRepeats score_deepcpf1
##             <logical>      <numeric>
##    spacer_1     FALSE      0.4334809
##    spacer_2     FALSE      0.0121805
##    spacer_3     FALSE      0.0112045
##    spacer_4     FALSE      0.0116443
##    spacer_5     FALSE      0.3527995
##         ...       ...            ...
##   spacer_30     FALSE       0.600635
##   spacer_31     FALSE       0.586483
##   spacer_32     FALSE       0.609269
##   spacer_33     FALSE       0.596874
##   spacer_34     FALSE       0.604862
##   -------
##   seqinfo: 640 sequences (1 circular) from hg38 genome
##   crisprNuclease: AsCas12a
```

```
geneAnnotation(guideSet)
```

```
## DataFrame with 91 rows and 23 columns
##                 chr anchor_site   strand gene_symbol         gene_id
##            <factor>   <integer> <factor> <character>     <character>
## spacer_1      chr12    25209818        +        KRAS ENSG00000133703
## spacer_1      chr12    25209818        +        KRAS ENSG00000133703
## spacer_1      chr12    25209818        +        KRAS ENSG00000133703
## spacer_2      chr12    25209835        +        KRAS ENSG00000133703
## spacer_2      chr12    25209835        +        KRAS ENSG00000133703
## ...             ...         ...      ...         ...             ...
## spacer_33     chr12    25245279        -        KRAS ENSG00000133703
## spacer_34     chr12    25245382        -        KRAS ENSG00000133703
## spacer_34     chr12    25245382        -        KRAS ENSG00000133703
## spacer_34     chr12    25245382        -        KRAS ENSG00000133703
## spacer_34     chr12    25245382        -        KRAS ENSG00000133703
##                     tx_id      protein_id   cut_cds cut_fiveUTRs cut_threeUTRs
##               <character>     <character> <logical>    <logical>     <logical>
## spacer_1  ENST00000256078              NA     FALSE        FALSE          TRUE
## spacer_1  ENST00000311936 ENSP00000308495      TRUE        FALSE         FALSE
## spacer_1  ENST00000557334 ENSP00000452512      TRUE        FALSE         FALSE
## spacer_2  ENST00000256078              NA     FALSE        FALSE          TRUE
## spacer_2  ENST00000311936 ENSP00000308495      TRUE        FALSE         FALSE
## ...                   ...             ...       ...          ...           ...
## spacer_33 ENST00000556131 ENSP00000256078      TRUE        FALSE         FALSE
## spacer_34 ENST00000256078 ENSP00000256078      TRUE        FALSE         FALSE
## spacer_34 ENST00000311936 ENSP00000256078      TRUE        FALSE         FALSE
## spacer_34 ENST00000557334 ENSP00000256078      TRUE        FALSE         FALSE
## spacer_34 ENST00000556131 ENSP00000256078      TRUE        FALSE         FALSE
##           cut_introns percentCDS aminoAcidIndex downtreamATG percentTx
##             <logical>  <numeric>      <numeric>    <numeric> <numeric>
```

```
## spacer_1        FALSE        NA            NA          NA      15.8
## spacer_1        FALSE      95.9           182           1      13.8
## spacer_1        FALSE      89.9            69           1      38.2
## spacer_2        FALSE        NA            NA          NA      15.5
## spacer_2        FALSE      92.9           176           1      13.5
## ...               ...       ...           ...         ...       ...
## spacer_33       FALSE      80.3            36           1      16.7
## spacer_34       FALSE       0.5             1           2       3.6
## spacer_34       FALSE       0.5             1           2       3.6
## spacer_34       FALSE       1.3             1           2      19.0
## spacer_34       FALSE       2.3             1           1      10.6
##           nIsoforms totalIsoforms percentIsoforms isCommonExon nCodingIsoforms
##           <integer>     <numeric>       <numeric>    <logical>       <integer>
## spacer_1          3             4              75        FALSE               3
## spacer_1          3             4              75        FALSE               3
## spacer_1          3             4              75        FALSE               3
## spacer_2          3             4              75        FALSE               3
## spacer_2          3             4              75        FALSE               3
## ...             ...           ...             ...          ...             ...
## spacer_33         4             4             100         TRUE               4
## spacer_34         4             4             100         TRUE               4
## spacer_34         4             4             100         TRUE               4
## spacer_34         4             4             100         TRUE               4
## spacer_34         4             4             100         TRUE               4
##           totalCodingIsoforms percentCodingIsoforms isCommonCodingExon
##                     <numeric>             <numeric>          <logical>
## spacer_1                    4                    75              FALSE
## spacer_1                    4                    75              FALSE
## spacer_1                    4                    75              FALSE
## spacer_2                    4                    75              FALSE
## spacer_2                    4                    75              FALSE
## ...                       ...                   ...                ...
## spacer_33                   4                   100               TRUE
## spacer_34                   4                   100               TRUE
## spacer_34                   4                   100               TRUE
## spacer_34                   4                   100               TRUE
## spacer_34                   4                   100               TRUE
```

VCF files for common SNPs (dbSNPs) can be downloaded from NCBI on the dbSNP website. We will use one of those files, after having downloaded it to our local machine.

```
# Users need to change this path to their local file
vcf <- "/Users/fortinj2/crisprIndices/snps/dbsnp151.grch38/00-common_all_snps_only.vcf.gz"
```

and we add a SNP annotation using the following command:

```
guideSet <- addSNPAnnotation(guideSet, vcf=vcf)
snps(guideSet)
```

```
## DataFrame with 4 rows and 9 columns
##                     rs   rs_site rs_site_rel     allele_ref   allele_minor
##            <character> <integer>   <numeric> <DNAStringSet> <DNAStringSet>
## spacer_3     rs1137282  25209843          26              A              G
## spacer_4     rs1137282  25209843          15              A              G
## spacer_5     rs1137282  25209843           6              A              G
## spacer_12   rs12313763  25209920           5              C              T
##            MAF_1000G MAF_TOPMED        type    length
##            <numeric>  <numeric> <character> <integer>
## spacer_3     0.17550    0.19671         snp         1
## spacer_4     0.17550    0.19671         snp         1
## spacer_5     0.17550    0.19671         snp         1
## spacer_12    0.08367    0.08473         snp         1
```

The rs_site_rel gives the relative position of the SNP with respect to the pam_site. allele_ref and allele_minor report the nucleotide of the reference and minor alleles, respectively. MAF_1000G and

MAF_TOPMED report the minor allele frequency (MAF) in the 1000Genomes and TOPMED populations, respectively.

#### Filtering and ranking gRNAs

Once gRNAs are fully annotated, it is easy to filter out any unwanted gRNAs since GuideSet objects can be subsetted like regular vectors in R.

As an example, suppose that we only want to keep gRNAs that have percent GC between 20% and 80% and that do not contain a polyT stretch. This can be achieved using the following lines:

```
guideSet <- guideSet[guideSet$percentGC>=20]
guideSet <- guideSet[guideSet$percentGC<=80]
guideSet <- guideSet[!guideSet$polyT]
```

Similarly, it is easy to rank gRNAs based on a set of criteria using the regular order function.

For instance, let's sort gRNAs by the DeepCpf1 on-target score:

```
# Creating an ordering index based on the DeepCpf1 score:
# Using the negative values to make sure higher scores are ranked first:
o <- order(-guideSet$score_deepcpf1)
# Ordering the GuideSet:
guideSet <- guideSet[o]
head(guideSet)
```

```
## GuideSet object with 6 ranges and 25 metadata columns:
##             seqnames    ranges strand |            protospacer            pam
##                <Rle> <IRanges>  <Rle> |         <DNAStringSet> <DNAStringSet>
##   spacer_24    chr12  25227223      + | AACCCACCTATAATGGTGAATAT           TTTA
##   spacer_19    chr12  25225707      - | CCTTCTAGAACAGTAGACACAAA           TTTG
##   spacer_21    chr12  25225717      + | CTACTAGGACCATAGGTACATCT           TTTC
##   spacer_16    chr12  25225634      + | AATAAAAGGAATTCCATAACTTC           TTTC
##   spacer_27    chr12  25227280      - | CCATAAATAATACTAAATCATTT           TTTG
##   spacer_14    chr12  25225598      + | AGTGTTACTTACCTGTCTTGTCT           TTTC
##              pam_site  cut_site      region percentGC     polyA     polyC
##             <numeric> <numeric> <character> <numeric> <logical> <logical>
##   spacer_24  25227223  25227247    region_6      34.8     FALSE     FALSE
##   spacer_19  25225707  25225683    region_7      39.1     FALSE     FALSE
##   spacer_21  25225717  25225741    region_7      43.5     FALSE     FALSE
##   spacer_16  25225634  25225658    region_7      26.1      TRUE     FALSE
##   spacer_27  25227280  25227256    region_6      17.4     FALSE     FALSE
##   spacer_14  25225598  25225622    region_7      39.1     FALSE     FALSE
##                 polyG     polyT startingGGGGG        n0        n1        n2
##             <logical> <logical>     <logical> <numeric> <numeric> <numeric>
##   spacer_24     FALSE     FALSE         FALSE         1         0         0
##   spacer_19     FALSE     FALSE         FALSE         2         0         0
##   spacer_21     FALSE     FALSE         FALSE         1         1         0
##   spacer_16     FALSE     FALSE         FALSE         1         0         0
##   spacer_27     FALSE     FALSE         FALSE         1         1         0
##   spacer_14     FALSE     FALSE         FALSE         1         0         0
##                  n0_c      n1_c      n2_c                      alignments
##             <numeric> <numeric> <numeric>                   <GRangesList>
##   spacer_24         1         0         0                chr12:25227223:+
##   spacer_19         1         0         0 chr12:25225707:-,chr6:54770950:+
##   spacer_21         1         0         0 chr12:25225717:+,chr6:54770940:-
##   spacer_16         1         0         0                chr12:25225634:+
```

```
##   spacer_27         1         0         0 chr12:25227280:-,chr6:54770837:+
##   spacer_14         1         0         0                chr12:25225598:+
##            inRepeats score_deepcpf1      enzymeAnnotation
##            <logical>      <numeric>  <SplitDataFrameList>
##   spacer_24     FALSE       0.826290 FALSE:FALSE:FALSE:...
##   spacer_19     FALSE       0.811090 FALSE:FALSE:FALSE:...
##   spacer_21     FALSE       0.744766 FALSE:FALSE:FALSE:...
##   spacer_16     FALSE       0.685073  TRUE:FALSE:FALSE:...
##   spacer_27     FALSE       0.665761 FALSE:FALSE:FALSE:...
##   spacer_14     FALSE       0.665542 FALSE:FALSE:FALSE:...
##                                                             geneAnnotation
##                                                       <SplitDataFrameList>
##   spacer_24                     chr12:25227247:+:...,chr12:25227247:+:...,...
##   spacer_19                     chr12:25225683:-:...,chr12:25225683:-:...,...
##   spacer_21                     chr12:25225741:+:...,chr12:25225741:+:...,...
##   spacer_16 chr12:25225658:+:...,chr12:25225658:+:...,chr12:25225658:+:...,...
##   spacer_27                     chr12:25227256:-:...,chr12:25227256:-:...,...
##   spacer_14 chr12:25225622:+:...,chr12:25225622:+:...,chr12:25225622:+:...,...
##                   tssAnnotation    hasSNP                 snps
##            <SplitDataFrameList> <logical> <SplitDataFrameList>
##   spacer_24             :...,...     FALSE             :...,...
##   spacer_19             :...,...     FALSE             :...,...
##   spacer_21             :...,...     FALSE             :...,...
##   spacer_16             :...,...     FALSE             :...,...
##   spacer_27             :...,...     FALSE             :...,...
##   spacer_14             :...,...     FALSE             :...,...
##   -------
##   seqinfo: 640 sequences (1 circular) from hg38 genome
##   crisprNuclease: AsCas12a
```

One can also sort gRNAs using several annotation columns. For instance, let's sort gRNAs using the DeepCpf1 score, but also by prioritizing first gRNAs that have no 1-mismatch off-targets in coding regions:

```
o <- order(guideSet$n1_c, -guideSet$score_deepcpf1)
# Ordering the GuideSet:
guideSet <- guideSet[o]
head(guideSet)
```

```
## GuideSet object with 6 ranges and 25 metadata columns:
##            seqnames    ranges strand |            protospacer            pam
##               <Rle> <IRanges>  <Rle> |         <DNAStringSet> <DNAStringSet>
##   spacer_24    chr12  25227223      + | AACCCACCTATAATGGTGAATAT           TTTA
##   spacer_19    chr12  25225707      - | CCTTCTAGAACAGTAGACACAAA           TTTG
##   spacer_21    chr12  25225717      + | CTACTAGGACCATAGGTACATCT           TTTC
##   spacer_16    chr12  25225634      + | AATAAAAGGAATTCCATAACTTC           TTTC
##   spacer_27    chr12  25227280      - | CCATAAATAATACTAAATCATTT           TTTG
##   spacer_14    chr12  25225598      + | AGTGTTACTTACCTGTCTTGTCT           TTTC
##             pam_site  cut_site      region percentGC     polyA     polyC
##            <numeric> <numeric> <character> <numeric> <logical> <logical>
##   spacer_24  25227223  25227247    region_6      34.8     FALSE     FALSE
##   spacer_19  25225707  25225683    region_7      39.1     FALSE     FALSE
##   spacer_21  25225717  25225741    region_7      43.5     FALSE     FALSE
##   spacer_16  25225634  25225658    region_7      26.1      TRUE     FALSE
##   spacer_27  25227280  25227256    region_6      17.4     FALSE     FALSE
##   spacer_14  25225598  25225622    region_7      39.1     FALSE     FALSE
```

```
##                polyG     polyT startingGGGGG        n0        n1        n2
##            <logical> <logical>     <logical> <numeric> <numeric> <numeric>
##   spacer_24     FALSE     FALSE         FALSE         1         0         0
##   spacer_19     FALSE     FALSE         FALSE         2         0         0
##   spacer_21     FALSE     FALSE         FALSE         1         1         0
##   spacer_16     FALSE     FALSE         FALSE         1         0         0
##   spacer_27     FALSE     FALSE         FALSE         1         1         0
##   spacer_14     FALSE     FALSE         FALSE         1         0         0
##                 n0_c      n1_c      n2_c                      alignments
##            <numeric> <numeric> <numeric>                    <GRangesList>
##   spacer_24         1         0         0                 chr12:25227223:+
##   spacer_19         1         0         0 chr12:25225707:-,chr6:54770950:+
##   spacer_21         1         0         0 chr12:25225717:+,chr6:54770940:-
##   spacer_16         1         0         0                 chr12:25225634:+
##   spacer_27         1         0         0 chr12:25227280:-,chr6:54770837:+
##   spacer_14         1         0         0                 chr12:25225598:+
##            inRepeats score_deepcpf1      enzymeAnnotation
##            <logical>      <numeric>  <SplitDataFrameList>
##   spacer_24     FALSE       0.826290 FALSE:FALSE:FALSE:...
##   spacer_19     FALSE       0.811090 FALSE:FALSE:FALSE:...
##   spacer_21     FALSE       0.744766 FALSE:FALSE:FALSE:...
##   spacer_16     FALSE       0.685073  TRUE:FALSE:FALSE:...
##   spacer_27     FALSE       0.665761 FALSE:FALSE:FALSE:...
##   spacer_14     FALSE       0.665542 FALSE:FALSE:FALSE:...
##                                                          geneAnnotation
##                                                    <SplitDataFrameList>
##   spacer_24                     chr12:25227247:+:...,chr12:25227247:+:...,...
##   spacer_19                     chr12:25225683:-:...,chr12:25225683:-:...,...
##   spacer_21                     chr12:25225741:+:...,chr12:25225741:+:...,...
##   spacer_16 chr12:25225658:+:...,chr12:25225658:+:...,chr12:25225658:+:...,...
##   spacer_27                     chr12:25227256:-:...,chr12:25227256:-:...,...
##   spacer_14 chr12:25225622:+:...,chr12:25225622:+:...,chr12:25225622:+:...,...
##                   tssAnnotation    hasSNP                 snps
##            <SplitDataFrameList> <logical> <SplitDataFrameList>
##   spacer_24               :...,...     FALSE             :...,...
##   spacer_19               :...,...     FALSE             :...,...
##   spacer_21               :...,...     FALSE             :...,...
##   spacer_16               :...,...     FALSE             :...,...
##   spacer_27               :...,...     FALSE             :...,...
##   spacer_14               :...,...     FALSE             :...,...
##   -------
##   seqinfo: 640 sequences (1 circular) from hg38 genome
##   crisprNuclease: AsCas12a
```

The `rankSpacers` function is a convenience function that implements our recommended rankings for the SpCas9, enAsCas12a and CasRx nucleases. For a detailed description of our recommended rankings, see the documentation of `rankSpacers` by typing `?rankSpacers`.

If an Ensembl transcript ID is provided, the ranking function will also take into account the position of the gRNA within the target CDS of the transcript ID in the ranking procedure. Our recommendation is to specify the Ensembl canonical transcript as the representative transcript for the gene. In our example, ENST00000311936 is the canonical transcript for KRAS:

```
tx_id <- "ENST00000311936"
guideSet <- rankSpacers(guideSet,
```

```r
                        tx_id=tx_id)
head(guideSet)
```

### Session Info

```r
sessionInfo()
```

```
## R version 4.2.1 (2022-06-23)
## Platform: x86_64-apple-darwin17.0 (64-bit)
## Running under: macOS Catalina 10.15.7
##
## Matrix products: default
## BLAS:   /Library/Frameworks/R.framework/Versions/4.2/Resources/lib/libRblas.0.dylib
## LAPACK: /Library/Frameworks/R.framework/Versions/4.2/Resources/lib/libRlapack.dylib
##
## locale:
## [1] en_US.UTF-8/en_US.UTF-8/en_US.UTF-8/C/en_US.UTF-8/en_US.UTF-8
##
## attached base packages:
## [1] stats4    stats     graphics  grDevices utils     datasets  methods
## [8] base
##
## other attached packages:
##  [1] BSgenome.Hsapiens.UCSC.hg38_1.4.4 BSgenome_1.65.2
##  [3] rtracklayer_1.57.0                Biostrings_2.65.2
##  [5] XVector_0.37.0                    GenomicRanges_1.49.1
##  [7] GenomeInfoDb_1.33.5               IRanges_2.31.2
##  [9] S4Vectors_0.35.1                  crisprDesignData_0.99.17
## [11] crisprDesign_0.99.133             crisprScore_1.1.14
## [13] crisprScoreData_1.1.3             ExperimentHub_2.5.0
## [15] AnnotationHub_3.5.0               BiocFileCache_2.5.0
## [17] dbplyr_2.2.1                      BiocGenerics_0.43.1
## [19] crisprBowtie_1.1.1                crisprBase_1.1.5
## [21] crisprVerse_0.99.8                rmarkdown_2.15.2
##
## loaded via a namespace (and not attached):
##  [1] rjson_0.2.21                  ellipsis_0.3.2
##  [3] Rbowtie_1.37.0                bit64_4.0.5
##  [5] lubridate_1.8.0               interactiveDisplayBase_1.35.0
##  [7] AnnotationDbi_1.59.1          fansi_1.0.3
##  [9] xml2_1.3.3                    codetools_0.2-18
## [11] cachem_1.0.6                  knitr_1.40
## [13] jsonlite_1.8.0                Rsamtools_2.13.4
## [15] png_0.1-7                     shiny_1.7.2
## [17] BiocManager_1.30.18           readr_2.1.2
## [19] compiler_4.2.1                httr_1.4.4
## [21] basilisk_1.9.2                assertthat_0.2.1
## [23] Matrix_1.4-1                  fastmap_1.1.0
## [25] cli_3.3.0                     later_1.3.0
## [27] htmltools_0.5.3               prettyunits_1.1.1
## [29] tools_4.2.1                   glue_1.6.2
## [31] GenomeInfoDbData_1.2.8        dplyr_1.0.9
```

```
## [33] rappdirs_0.3.3              tinytex_0.41
## [35] Rcpp_1.0.9                  Biobase_2.57.1
## [37] vctrs_0.4.1                 crisprBwa_1.1.3
## [39] xfun_0.32                   stringr_1.4.1
## [41] mime_0.12                   lifecycle_1.0.1
## [43] restfulr_0.0.15             XML_3.99-0.10
## [45] zlibbioc_1.43.0             basilisk.utils_1.9.1
## [47] vroom_1.5.7                 VariantAnnotation_1.43.3
## [49] hms_1.1.2                   promises_1.2.0.1
## [51] MatrixGenerics_1.9.1        parallel_4.2.1
## [53] SummarizedExperiment_1.27.1 RMariaDB_1.2.2
## [55] yaml_2.3.5                  curl_4.3.2
## [57] memoise_2.0.1               reticulate_1.25
## [59] biomaRt_2.53.2              stringi_1.7.8
## [61] RSQLite_2.2.16              BiocVersion_3.16.0
## [63] highr_0.9                   BiocIO_1.7.1
## [65] randomForest_4.7-1.1        GenomicFeatures_1.49.6
## [67] filelock_1.0.2              BiocParallel_1.31.12
## [69] rlang_1.0.4                 pkgconfig_2.0.3
## [71] matrixStats_0.62.0          bitops_1.0-7
## [73] evaluate_0.16               lattice_0.20-45
## [75] purrr_0.3.4                 GenomicAlignments_1.33.1
## [77] bit_4.0.4                   tidyselect_1.1.2
## [79] magrittr_2.0.3              R6_2.5.1
## [81] generics_0.1.3              DelayedArray_0.23.1
## [83] DBI_1.1.3                   pillar_1.8.1
## [85] KEGGREST_1.37.3             RCurl_1.98-1.8
## [87] tibble_3.1.8                dir.expiry_1.5.0
## [89] crayon_1.5.1                utf8_1.2.2
## [91] tzdb_0.3.0                  progress_1.2.2
## [93] grid_4.2.1                  blob_1.2.3
## [95] digest_0.6.29               xtable_1.8-4
## [97] httpuv_1.6.5                Rbwa_1.1.0
```
