## Supplementary material for "The crisprVerse: a comprehensive Bioconductor ecosystem for the design of CRISPR guide RNAs across nucleases and technologies": Tutorial7_CRISPRko_Design_CasRx

### Using crisprDesign to design gRNAs for CRISPRkd with CasRx

#### Introduction

In this tutorial, we will design gRNAs for the RNA-targeting nuclease CasRx (RfxCas13d) (Konermann et al. 2018). We will design all gRNAs targeting the primary isoform of the human gene KRAS.

#### Installation

See the Installation tutorial to learn how to install the packages necessary for this tutorial: `crisprDesign`, `crisprDesignData`

#### Terminology

See the CRISPRko design vignette to get familiar with the terminology used throughout this tutorial.

#### End-to-end gRNA design workflow

We first start by loading the crisprVerse packages needed for this tutorial:

```
library(crisprBase)
library(crisprDesign)
library(crisprDesignData)
```

We will also load the `BSgenome` package containing DNA sequences for the hg38 genome:

```
library(BSgenome.Hsapiens.UCSC.hg38)
```

##### Creating the GuideSet

We begin by loading the CasRx `CrisprNuclease` object from the `crisprBase` package:

```
data(CasRx, package="crisprBase")
CasRx
```

```
## Class: CrisprNuclease
##   Name: CasRx
##   Target type: RNA
##   Metadata: list of length 2
##   PFS: N
##   Weights: 1
##   Spacer length: 23
##   PFS side: 3prime
##     Distance from PFS: 0
##   Prototype protospacers: 5'--SSSSSSSSSSSSSSSSSSSSSSS[N]--3'
```

The PFS sequence (the equivalent of a PAM sequence for RNA-targeting nucleases) for CasRx is `N`, meaning there is no specific PFS sequences preferred by CasRx.

Next, we will extract the mRNA sequence for the KRAS transcript ENST00000311936 with the function `getMrnaSequences` from `crisprDesign`. The function requires a gene annotation object. We will load the Ensembl model from the `crisprDesignData` package stored in the `GRangesList` object `txdb_human`:

```
data("txdb_human", package="crisprDesignData")
```

For more information on `txdb_human` and how to create similar gene annotation objects, see the Building a gene annotation object tutorial).

We also need a `BSgenome` object containing the DNA sequences:

```
bsgenome <- BSgenome.Hsapiens.UCSC.hg38
```

We are now ready to obtain our mRNA sequence:

```
txid <- "ENST00000311936"
mrna <- getMrnaSequences(txids=txid,
                         bsgenome=bsgenome,
                         txObject=txdb_human)
mrna
```

```
## DNAStringSet object of length 1:
##     width seq                                               names
## [1]  5306 CTAGGCGGCGGCCGCGGCGGCGG...GTAATAAAAATAGTTACAGTGAC ENST00000311936
```

Similar to the CRISPRko gRNA design, we use the function `findSpacers` to design our gRNAs:

```
gs <- findSpacers(mrna,
                  crisprNuclease=CasRx)
head(gs)
```

```
## GuideSet object with 6 ranges and 5 metadata columns:
##                   seqnames    ranges strand |             protospacer
##                      <Rle> <IRanges>  <Rle> |          <DNAStringSet>
##   spacer_1 ENST00000311936        24      + | CTAGGCGGCGGCCGCGGCGGCGG
##   spacer_2 ENST00000311936        25      + | TAGGCGGCGGCCGCGGCGGCGGA
##   spacer_3 ENST00000311936        26      + | AGGCGGCGGCCGCGGCGGCGGAG
##   spacer_4 ENST00000311936        27      + | GGCGGCGGCCGCGGCGGCGGAGG
##   spacer_5 ENST00000311936        28      + | GCGGCGGCCGCGGCGGCGGAGGC
##   spacer_6 ENST00000311936        29      + | CGGCGGCCGCGGCGGCGGAGGCA
##                        pam  pam_site  cut_site          region
##             <DNAStringSet> <numeric> <numeric>     <character>
##   spacer_1               A        24        NA ENST00000311936
##   spacer_2               G        25        NA ENST00000311936
##   spacer_3               G        26        NA ENST00000311936
##   spacer_4               C        27        NA ENST00000311936
##   spacer_5               A        28        NA ENST00000311936
##   spacer_6               G        29        NA ENST00000311936
##   -------
##   seqinfo: 1 sequence from custom genome
##   crisprNuclease: CasRx
```

Note that all protospacer sequences are located on the original strand of the mRNA sequence. For RNA-targeting nucleases, the spacer and protospacer sequences are the reverse complement of each other. (Compare the output of the code below with a `GuideSet` that uses a DNA-targeting nuclease–for such `GuideSet` pbjects, the output of `spacers` and `protospacers` are identical.)

```
head(spacers(gs))
```

```
## DNAStringSet object of length 6:
##     width seq                                               names
## [1]    23 CCGCCGCCGCGGCCGCCGCCTAG                           spacer_1
## [2]    23 TCCGCCGCCGCGGCCGCCGCCTA                           spacer_2
## [3]    23 CTCCGCCGCCGCGGCCGCCGCCT                           spacer_3
## [4]    23 CCTCCGCCGCCGCGGCCGCCGCC                           spacer_4
## [5]    23 GCCTCCGCCGCCGCGGCCGCCGC                           spacer_5
## [6]    23 TGCCTCCGCCGCCGCGGCCGCCG                           spacer_6
```

```
head(protospacers(gs))
```

```
## DNAStringSet object of length 6:
##     width seq                                               names
## [1]    23 CTAGGCGGCGGCCGCGGCGGCGG                           spacer_1
## [2]    23 TAGGCGGCGGCCGCGGCGGCGGA                           spacer_2
## [3]    23 AGGCGGCGGCCGCGGCGGCGGAG                           spacer_3
## [4]    23 GGCGGCGGCCGCGGCGGCGGAGG                           spacer_4
## [5]    23 GCGGCGGCCGCGGCGGCGGAGGC                           spacer_5
## [6]    23 CGGCGGCCGCGGCGGCGGAGGCA                           spacer_6
```

#### Annotating the GuideSet

Next, we annotate our candidate gRNAs to assess quality. There are several functions in `crisprDesign` that provide annotation for features that are nonspecific to CRISPRkd, for which we refer the reader to the CRISPRko design with Cas9 tutorial for more information. The sections below will cover annotation functions that are of particular interest to, or deserve extra care for CRISPRkd applications.

##### Adding spacer alignments

Since our CRISPR nuclease targets RNA rather than DNA, off-target searches should be restricted to the transcriptome. We can perform such a search using one of two methods.

**Adding spacer alignments with Biostrings** For the first method, we set the `aligner` argument to `"biostrings"` and pass a `DNAStringSet` representation of the transcriptome to the argument `custom_seq`. We can create this representation with `getMrnaSequences` and all transcript IDs found in `txdb_human`. The code below uses this method to search for off-targets having up to one mismatch and passes `txdb_human` to the `txObject` argument so that the alignments will be accompanied with gene annotation.

```
exon_ids <- unique(txdb_human$exons$tx_id)
mrnasHuman <- getMrnaSequences(exon_ids,
                               bsgenome=BSgenome.Hsapiens.UCSC.hg38,
                               txObject=txdb_human)
## long run time
results <- addSpacerAlignments(gs,
                               aligner="biostrings",
                               txObject=txdb_human,
                               n_mismatches=1,
                               custom_seq=mrnasHuman)
```

NOTE: since `mrnasHuman` contains many sequences (>100k), this method has a very long run time; for transcriptome-wide searches, or for searches against a large number of sequences, we recommend the following method instead.

**Adding spacer alignments with bowtie or BWA** The second method uses the `bowtie` (or `bwa`) aligner. This requires building a transcriptome bowtie (or BWA) index file first. See the Building genome indices for short read aligners tutorial for more information.

Here we set `aligner` to `"bowtie"` and pass a precomputed transcriptome bowtie index to `aligner_index` to find off-targets:

```
bowtie_index <- "/Users/fortinj2/crisprIndices/bowtie/ensembl_human_104/ensembl_human_104"
results <- addSpacerAlignments(gs,
                               aligner="bowtie",
                               aligner_index=bowtie_index,
                               txObject=txdb_human,
                               n_mismatches=1)
```

```
head(results)
```

```
## GuideSet object with 6 ranges and 10 metadata columns:
##                   seqnames    ranges strand |             protospacer
##                      <Rle> <IRanges>  <Rle> |          <DNAStringSet>
##   spacer_1 ENST00000311936        24      + | CTAGGCGGCGGCCGCGGCGGCGG
##   spacer_2 ENST00000311936        25      + | TAGGCGGCGGCCGCGGCGGCGGA
##   spacer_3 ENST00000311936        26      + | AGGCGGCGGCCGCGGCGGCGGAG
##   spacer_4 ENST00000311936        27      + | GGCGGCGGCCGCGGCGGCGGAGG
##   spacer_5 ENST00000311936        28      + | GCGGCGGCCGCGGCGGCGGAGGC
##   spacer_6 ENST00000311936        29      + | CGGCGGCCGCGGCGGCGGAGGCA
##                       pam  pam_site  cut_site          region     n0_tx
##            <DNAStringSet> <numeric> <numeric>     <character> <numeric>
##   spacer_1              A        24        NA ENST00000311936         4
##   spacer_2              G        25        NA ENST00000311936         4
##   spacer_3              G        26        NA ENST00000311936         4
##   spacer_4              C        27        NA ENST00000311936         4
##   spacer_5              A        28        NA ENST00000311936         4
##   spacer_6              G        29        NA ENST00000311936         4
##                n1_tx   n0_gene   n1_gene
##            <numeric> <numeric> <numeric>
##   spacer_1         0         1         0
##   spacer_2         0         1         0
##   spacer_3         9         1         6
##   spacer_4        81         1        26
##   spacer_5        67         1        18
##   spacer_6        44         1         5
##                                                                    alignments
##                                                                 <GRangesList>
##   spacer_1 ENST00000256078:24:+,ENST00000311936:24:+,ENST00000556131:24:+,...
##   spacer_2 ENST00000256078:25:+,ENST00000311936:25:+,ENST00000556131:25:+,...
##   spacer_3 ENST00000256078:26:+,ENST00000311936:26:+,ENST00000556131:26:+,...
##   spacer_4 ENST00000256078:27:+,ENST00000311936:27:+,ENST00000556131:27:+,...
##   spacer_5 ENST00000256078:28:+,ENST00000311936:28:+,ENST00000556131:28:+,...
##   spacer_6 ENST00000256078:29:+,ENST00000311936:29:+,ENST00000556131:29:+,...
##   -------
##   seqinfo: 1 sequence from custom genome
##   crisprNuclease: CasRx
```

The columns `n0_gene` and `n0_tx` report the number of on-targets at the gene- and transcript-level, respectively. For instance, each spacer shown above shows `n0_gene` equal to 1 and `n0_tx` equal to 4, meaning each spacer maps to all four isoforms of KRAS. We can retrieve information about each alignment with the `onTargets` function. Looking at the on-targets for the first spacer we can see where the target `pam_site` is relative to the start of the transcript with respect to each isoform of KRAS.

```
onTargets(results["spacer_1"])
```

```
## GRanges object with 4 ranges and 9 metadata columns:
##                  seqnames    ranges strand |                 spacer
##                     <Rle> <IRanges>  <Rle> |         <DNAStringSet>
##   spacer_1 ENST00000256078        24      + | CCGCCGCCGCGGCCGCCGCCTAG
##   spacer_1 ENST00000311936        24      + | CCGCCGCCGCGGCCGCCGCCTAG
##   spacer_1 ENST00000556131        24      + | CCGCCGCCGCGGCCGCCGCCTAG
##   spacer_1 ENST00000557334        31      + | CCGCCGCCGCGGCCGCCGCCTAG
##                       protospacer           pam  pam_site n_mismatches
##                    <DNAStringSet> <DNAStringSet> <numeric>    <integer>
##   spacer_1 CTAGGCGGCGGCCGCGGCGGCGG             A        24            0
##   spacer_1 CTAGGCGGCGGCCGCGGCGGCGG             A        24            0
##   spacer_1 CTAGGCGGCGGCCGCGGCGGCGG             A        24            0
##   spacer_1 CTAGGCGGCGGCCGCGGCGGCGG             A        31            0
##            canonical  cut_site         gene_id gene_symbol
##            <logical> <numeric>     <character> <character>
##   spacer_1      TRUE        NA ENSG00000133703        KRAS
##   spacer_1      TRUE        NA ENSG00000133703        KRAS
##   spacer_1      TRUE        NA ENSG00000133703        KRAS
##   spacer_1      TRUE        NA ENSG00000133703        KRAS
##   -------
##   seqinfo: 7514 sequences from an unspecified genome; no seqlengths
```

Note that each annotated alignment is specific to the transcript ID given under `seqnames`.

Below is a spacer that targets (with no mismatches) multiple genes:

```
results["spacer_244"]
```

```
## GuideSet object with 1 range and 10 metadata columns:
##                    seqnames    ranges strand |             protospacer
##                       <Rle> <IRanges>  <Rle> |          <DNAStringSet>
##   spacer_244 ENST00000311936       267      + | CTTGACGATACAGCTAATTCAGA
##                         pam  pam_site  cut_site          region     n0_tx
##              <DNAStringSet> <numeric> <numeric>     <character> <numeric>
##   spacer_244              A       267        NA ENST00000311936         5
##                   n1_tx   n0_gene   n1_gene
##              <numeric> <numeric> <numeric>
##   spacer_244         0         2         0
##                                                                   alignments
##                                                                <GRangesList>
##   spacer_244 ENST00000256078:267:+,ENST00000311936:267:+,ENST00000407852:77:+,...
##   -------
##   seqinfo: 1 sequence from custom genome
##   crisprNuclease: CasRx
```

Upon further inspection of this spacer's alignments, however, we can see that the off-target occurs in the pseudogene KRASP1, and should be harmless.

```
onTargets(results["spacer_244"])
```

```
## GRanges object with 5 ranges and 9 metadata columns:
##                    seqnames    ranges strand |                 spacer
##                       <Rle> <IRanges>  <Rle> |         <DNAStringSet>
```

```
##   spacer_244 ENST00000256078       267      + | TCTGAATTAGCTGTATCGTCAAG
##   spacer_244 ENST00000311936       267      + | TCTGAATTAGCTGTATCGTCAAG
##   spacer_244 ENST00000407852        77      + | TCTGAATTAGCTGTATCGTCAAG
##   spacer_244 ENST00000556131       254      + | TCTGAATTAGCTGTATCGTCAAG
##   spacer_244 ENST00000557334       274      + | TCTGAATTAGCTGTATCGTCAAG
##                          protospacer           pam  pam_site n_mismatches
##                       <DNAStringSet> <DNAStringSet> <numeric>    <integer>
##   spacer_244 CTTGACGATACAGCTAATTCAGA              A       267            0
##   spacer_244 CTTGACGATACAGCTAATTCAGA              A       267            0
##   spacer_244 CTTGACGATACAGCTAATTCAGA              A        77            0
##   spacer_244 CTTGACGATACAGCTAATTCAGA              A       254            0
##   spacer_244 CTTGACGATACAGCTAATTCAGA              A       274            0
##               canonical  cut_site         gene_id gene_symbol
##               <logical> <numeric>     <character> <character>
##   spacer_244      TRUE        NA ENSG00000133703        KRAS
##   spacer_244      TRUE        NA ENSG00000133703        KRAS
##   spacer_244      TRUE        NA ENSG00000220635      KRASP1
##   spacer_244      TRUE        NA ENSG00000133703        KRAS
##   spacer_244      TRUE        NA ENSG00000133703        KRAS
##   -------
##   seqinfo: 7514 sequences from an unspecified genome; no seqlengths
```

##### On-target scoring (gRNA efficiency)

Finally, we add an on-target activity score using the CasRx-RF method (Wessels et al. 2020) using the `addOnTargetScores` function from `crisprDesign` package:

```
gs <- addOnTargetScores(gs, methods=c("casrxrf"))
```

```
## [addCasRxScores] Calculating MFE features
## [addCasRxScores] Calculating accessibility features
## [addCasRxScores] Calculating hybridization features
## [addCasRxScores] Calculating nucleotide density features
## [getCasRxRFScores] Calculating scores
```

```
gs
```

```
## GuideSet object with 5283 ranges and 6 metadata columns:
##                     seqnames    ranges strand |             protospacer
##                        <Rle> <IRanges>  <Rle> |          <DNAStringSet>
##      spacer_1 ENST00000311936        24      + | CTAGGCGGCGGCCGCGGCGGCGG
##      spacer_2 ENST00000311936        25      + | TAGGCGGCGGCCGCGGCGGCGGA
##      spacer_3 ENST00000311936        26      + | AGGCGGCGGCCGCGGCGGCGGAG
##      spacer_4 ENST00000311936        27      + | GGCGGCGGCCGCGGCGGCGGAGG
##      spacer_5 ENST00000311936        28      + | GCGGCGGCCGCGGCGGCGGAGGC
##           ...             ...       ...    ... .                     ...
##   spacer_5279 ENST00000311936      5302      + | GTAATGTAATAAAAATAGTTACA
##   spacer_5280 ENST00000311936      5303      + | TAATGTAATAAAAATAGTTACAG
##   spacer_5281 ENST00000311936      5304      + | AATGTAATAAAAATAGTTACAGT
##   spacer_5282 ENST00000311936      5305      + | ATGTAATAAAAATAGTTACAGTG
##   spacer_5283 ENST00000311936      5306      + | TGTAATAAAAATAGTTACAGTGA
##                          pam  pam_site  cut_site          region score_casrxrf
##               <DNAStringSet> <numeric> <numeric>     <character>     <numeric>
##      spacer_1              A        24        NA ENST00000311936            NA
##      spacer_2              G        25        NA ENST00000311936      0.387198
##      spacer_3              G        26        NA ENST00000311936            NA
```

```
##      spacer_4             C        27        NA ENST00000311936            NA
##      spacer_5             A        28        NA ENST00000311936            NA
##           ...           ...       ...       ...             ...           ...
##   spacer_5279             G      5302        NA ENST00000311936            NA
##   spacer_5280             T      5303        NA ENST00000311936            NA
##   spacer_5281             G      5304        NA ENST00000311936            NA
##   spacer_5282             A      5305        NA ENST00000311936            NA
##   spacer_5283             C      5306        NA ENST00000311936            NA
##   -------
##   seqinfo: 1 sequence from custom genome
##   crisprNuclease: CasRx
```

### Session Info

```
sessionInfo()
```

```
## R version 4.2.1 (2022-06-23)
## Platform: x86_64-apple-darwin17.0 (64-bit)
## Running under: macOS Catalina 10.15.7
##
## Matrix products: default
## BLAS:   /Library/Frameworks/R.framework/Versions/4.2/Resources/lib/libRblas.0.dylib
## LAPACK: /Library/Frameworks/R.framework/Versions/4.2/Resources/lib/libRlapack.dylib
##
## locale:
## [1] en_US.UTF-8/en_US.UTF-8/en_US.UTF-8/C/en_US.UTF-8/en_US.UTF-8
##
## attached base packages:
## [1] stats4    stats     graphics  grDevices utils     datasets  methods
## [8] base
##
## other attached packages:
##  [1] BSgenome.Hsapiens.UCSC.hg38_1.4.4 BSgenome_1.65.2
##  [3] rtracklayer_1.57.0                Biostrings_2.65.2
##  [5] XVector_0.37.0                    GenomicRanges_1.49.1
##  [7] GenomeInfoDb_1.33.5               IRanges_2.31.2
##  [9] S4Vectors_0.35.1                  crisprDesignData_0.99.17
## [11] crisprDesign_0.99.133             crisprScore_1.1.14
## [13] crisprScoreData_1.1.3             ExperimentHub_2.5.0
## [15] AnnotationHub_3.5.0               BiocFileCache_2.5.0
## [17] dbplyr_2.2.1                      BiocGenerics_0.43.1
## [19] crisprBowtie_1.1.1                crisprBase_1.1.5
## [21] crisprVerse_0.99.8                rmarkdown_2.15.2
##
## loaded via a namespace (and not attached):
##  [1] rjson_0.2.21                  ellipsis_0.3.2
##  [3] Rbowtie_1.37.0                bit64_4.0.5
##  [5] lubridate_1.8.0               interactiveDisplayBase_1.35.0
##  [7] AnnotationDbi_1.59.1          fansi_1.0.3
##  [9] xml2_1.3.3                    codetools_0.2-18
## [11] cachem_1.0.6                  knitr_1.40
## [13] jsonlite_1.8.0                Rsamtools_2.13.4
## [15] png_0.1-7                     shiny_1.7.2
```

```
## [17] BiocManager_1.30.18          readr_2.1.2
## [19] compiler_4.2.1                httr_1.4.4
## [21] basilisk_1.9.2                assertthat_0.2.1
## [23] Matrix_1.4-1                  fastmap_1.1.0
## [25] cli_3.3.0                     later_1.3.0
## [27] htmltools_0.5.3               prettyunits_1.1.1
## [29] tools_4.2.1                   glue_1.6.2
## [31] GenomeInfoDbData_1.2.8        dplyr_1.0.9
## [33] rappdirs_0.3.3                tinytex_0.41
## [35] Rcpp_1.0.9                    Biobase_2.57.1
## [37] vctrs_0.4.1                   crisprBwa_1.1.3
## [39] xfun_0.32                     stringr_1.4.1
## [41] mime_0.12                     lifecycle_1.0.1
## [43] restfulr_0.0.15               XML_3.99-0.10
## [45] zlibbioc_1.43.0               basilisk.utils_1.9.1
## [47] vroom_1.5.7                   VariantAnnotation_1.43.3
## [49] hms_1.1.2                     promises_1.2.0.1
## [51] MatrixGenerics_1.9.1          parallel_4.2.1
## [53] SummarizedExperiment_1.27.1   RMariaDB_1.2.2
## [55] yaml_2.3.5                    curl_4.3.2
## [57] memoise_2.0.1                 reticulate_1.25
## [59] biomaRt_2.53.2                stringi_1.7.8
## [61] RSQLite_2.2.16                BiocVersion_3.16.0
## [63] highr_0.9                     BiocIO_1.7.1
## [65] randomForest_4.7-1.1          GenomicFeatures_1.49.6
## [67] filelock_1.0.2                BiocParallel_1.31.12
## [69] rlang_1.0.4                   pkgconfig_2.0.3
## [71] matrixStats_0.62.0            bitops_1.0-7
## [73] evaluate_0.16                 lattice_0.20-45
## [75] purrr_0.3.4                   GenomicAlignments_1.33.1
## [77] bit_4.0.4                     tidyselect_1.1.2
## [79] magrittr_2.0.3                R6_2.5.1
## [81] generics_0.1.3                DelayedArray_0.23.1
## [83] DBI_1.1.3                     pillar_1.8.1
## [85] KEGGREST_1.37.3               RCurl_1.98-1.8
## [87] tibble_3.1.8                  dir.expiry_1.5.0
## [89] crayon_1.5.1                  utf8_1.2.2
## [91] tzdb_0.3.0                    progress_1.2.2
## [93] grid_4.2.1                    blob_1.2.3
## [95] digest_0.6.29                 xtable_1.8-4
## [97] httpuv_1.6.5                  Rbwa_1.1.0
```

### References


Konermann, Silvana, Peter Lotfy, Nicholas J Brideau, Jennifer Oki, Maxim N Shokhirev, and Patrick D Hsu. 2018. "Transcriptome Engineering with RNA-Targeting Type VI-d CRISPR Effectors." *Cell* 173 (3): 665–76.

Wessels, Hans-Hermann, Alejandro Méndez-Mancilla, Xinyi Guo, Mateusz Legut, Zharko Daniloski, and Neville E Sanjana. 2020. "Massively Parallel Cas13 Screens Reveal Principles for Guide RNA Design." *Nature Biotechnology* 38 (6): 722–27.
