## Supplementary material for "The crisprVerse: a comprehensive Bioconductor ecosystem for the design of CRISPR guide RNAs across nucleases and technologies": Tutorial10_CRISPRi_Design

### gRNA design for CRISPR interference (CRISPRi)

#### Introduction

This tutorial will demonstrate how to use `crisprDesign` to design gRNAs for CRISPR interference (CRISPRi). Specifically, we will target the human KRAS gene and use the SpCas9 nuclease.

#### Installation

See the Installation tutorial to learn how to install the packages necessary for this tutorial: `crisprDesign`, `crisprDesignData`

#### Terminology

See the CRISPRko design tutorial to get familiar with the terminology used throughout this tutorial.

#### CRISPRi design

For CRISPR activation (CRISPRa) and interference (CRISPRi) applications, the CRISPR nuclease is engineered to lose its endonuclease activity, and should therefore not introduce double-stranded breaks (DSBs). We will use the dead SpCas9 (dSpCas9) nuclease as an example here. Note that users don't have to distinguish between dSpCas9 and SpCas9 when specifying the nuclease in the crisprVerse as they do not differ in terms of the characteristics stored in the `CrisprNuclease` object.

In CRISPRi, fusing dSpCas9 with a Krüppel-associated box (KRAB) domain has been shown to be effective at repressing transcription in mammalian cells (Gilbert et al. et al. 2013). The dSpCas9-KRAB fused protein is a commonly-used construct to conduct CRISPR inhibition (CRISPRi) experiments. To achieve optimal inhibition, gRNAs are usually designed targeting the region directly downstream of the gene transcription starting site (TSS).

`crisprDesign` provides functionalities to be able to take into account design rules that are specific to CRISPRi applications. The `queryTss` function allows for specifying genomic coordinates of promoter regions. The `addTssAnnotation` function annotates gRNAs for known TSSs, and includes a column `dist_to_tss` that gives the distance in nucleotides between the TSS position and the PAM site of the gRNA. For CRISPRi, we recommend targeting the region 25-75bp region downstream of the TSS for optimal inhibition; see Sanson et al. et al. (2018) for more information. Finally, the function `addCrispraiScores` adds CRISPRi-specific on-target activity scores based on the work of (Horlbeck et al. et al. 2016).

##### Creating the GuideSet

We first start by loading the required packages:

```
library(crisprBase)
library(crisprDesign)
```

```r
library(crisprDesignData)
library(BSgenome.Hsapiens.UCSC.hg38)
```

To demonstrate CRISPRi design, we will design gRNAs to inhibit expression of the human KRAS gene using the SpCas9 nuclease. To accomplish this, we want our gRNAs to target the region downstream of the KRAS TSS; let's consider the window containing 500bp immediately downstream of the TSS to explore candidate gRNAs.

We first need to retrieve the TSS coordinates for KRAS. These data are conveniently stored in the `crisprDesignData` package as the dataset `tss_human`. For more information on `tss_human` and how to create similar TSS annotation objects, see the Building a gene annotation object tutorial.

We load the TSS coordinates stored in the `tss_human` object

```r
data("tss_human", package="crisprDesignData")
```

and query for KRAS using the `queryTss` function from `crisprDesign`:

```r
target_window <- c(0, 500)
target_region <- queryTss(tss_human,
                          queryColumn="gene_symbol",
                          queryValue="KRAS",
                          tss_window=target_window)
```

```r
target_region
## GRanges object with 1 range and 9 metadata columns:
##            seqnames            ranges strand |     score peak_start   peak_end
##               <Rle>         <IRanges>  <Rle> | <numeric>  <integer> <integer>
##   region_1    chr12 25250429-25250928      - |   5.20187   25250928  25250928
##                      tx_id         gene_id      source    promoter
##                <character>     <character> <character> <character>
##   region_1 ENST00000256078 ENSG00000133703     fantom5          P1
##                         ID gene_symbol
##                <character> <character>
##   region_1 ENSG00000133703_P1        KRAS
##   -------
##   seqinfo: 25 sequences from an unspecified genome; no seqlengths
```

We load the `crisprNuclease` object storing information about the SpCas9 nuclease from the `crisprBase` package:

```r
data(SpCas9, package="crisprBase")
```

We then find all candidate protospacer sequences in our target region with `findSpacers`:

```r
gs <- findSpacers(target_region,
                  crisprNuclease=SpCas9,
                  bsgenome=BSgenome.Hsapiens.UCSC.hg38)
```

```r
gs
## GuideSet object with 160 ranges and 5 metadata columns:
##              seqnames    ranges strand |          protospacer            pam
##                 <Rle> <IRanges>  <Rle> |       <DNAStringSet> <DNAStringSet>
##     spacer_1    chr12  25250434      - | GCCGCGGCTGGAGGCTTCTG            GGG
##     spacer_2    chr12  25250435      - | AGCCGCGGCTGGAGGCTTCT            GGG
##     spacer_3    chr12  25250436      - | GAGCCGCGGCTGGAGGCTTC            TGG
##     spacer_4    chr12  25250443      - | TCCCCGAGAGCCGCGGCTGG            AGG
##     spacer_5    chr12  25250446      - | TCCTCCCCGAGAGCCGCGGC            TGG
```

```
##           ...       ...        ...    ... .                  ...          ...
##   spacer_156    chr12  25250915      - | ATTTTCCTAGGCGGCGGCCG          CGG
##   spacer_157    chr12  25250916      + | CGCTGCTGCCTCCGCCGCCG          CGG
##   spacer_158    chr12  25250921      - | AGCTCGATTTTCCTAGGCGG          CGG
##   spacer_159    chr12  25250924      - | CGGAGCTCGATTTTCCTAGG          CGG
##   spacer_160    chr12  25250928      + | CGCCGCCGCGGCCGCCGCCT          AGG
##                pam_site  cut_site      region
##               <numeric> <numeric> <character>
##     spacer_1  25250434  25250437    region_1
##     spacer_2  25250435  25250438    region_1
##     spacer_3  25250436  25250439    region_1
##     spacer_4  25250443  25250446    region_1
##     spacer_5  25250446  25250449    region_1
##          ...       ...       ...         ...
##   spacer_156  25250915  25250918    region_1
##   spacer_157  25250916  25250913    region_1
##   spacer_158  25250921  25250924    region_1
##   spacer_159  25250924  25250927    region_1
##   spacer_160  25250928  25250925    region_1
##   -------
##   seqinfo: 640 sequences (1 circular) from hg38 genome
##   crisprNuclease: SpCas9
```

#### Annotating the GuideSet

Next, we annotate our candidate gRNAs to assess quality. There are several functions in `crisprDesign` that provide annotation for features that are not specific to CRISPRi, for which we refer the reader to the CRISPRko design with Cas9 tutorial for more information. The sections below will cover annotation functions that are of particular interest to CRISPRi applications.

#### Adding TSS annotation

As the name implies, the `addTssAnnotation` function annotates gRNAs with TSS context such as the distance between the gRNA and the TSS, as well as which TSS is targeted (many genes contain different TSSs corresponding to different isoforms).

The function requires a `tssObject` object, and the `tss_window` values that we used earlier to define the target region. We can then retrieve the appended annotation with the accessor function `tssAnnotation`:

```
gs <- addTssAnnotation(gs,
                       tssObject=tss_human,
                       tss_window=target_window)
tssAnnotation(gs)
## DataFrame with 160 rows and 15 columns
##                  chr anchor_site   strand     score peak_start  peak_end
##             <factor>   <integer> <factor> <numeric>  <integer> <integer>
## spacer_1       chr12    25250437        -   5.20187   25250928  25250928
## spacer_2       chr12    25250438        -   5.20187   25250928  25250928
## spacer_3       chr12    25250439        -   5.20187   25250928  25250928
## spacer_4       chr12    25250446        -   5.20187   25250928  25250928
## spacer_5       chr12    25250449        -   5.20187   25250928  25250928
## ...              ...         ...      ...       ...        ...       ...
## spacer_156     chr12    25250918        -   5.20187   25250928  25250928
## spacer_157     chr12    25250913        +   5.20187   25250928  25250928
```

```
## spacer_158    chr12    25250924        -   5.20187   25250928  25250928
## spacer_159    chr12    25250927        -   5.20187   25250928  25250928
## spacer_160    chr12    25250925        +   5.20187   25250928  25250928
##                         tx_id         gene_id       source    promoter
##                   <character>     <character> <character> <character>
## spacer_1    ENST00000256078 ENSG00000133703      fantom5          P1
## spacer_2    ENST00000256078 ENSG00000133703      fantom5          P1
## spacer_3    ENST00000256078 ENSG00000133703      fantom5          P1
## spacer_4    ENST00000256078 ENSG00000133703      fantom5          P1
## spacer_5    ENST00000256078 ENSG00000133703      fantom5          P1
## ...                     ...             ...          ...         ...
## spacer_156 ENST00000256078 ENSG00000133703      fantom5          P1
## spacer_157 ENST00000256078 ENSG00000133703      fantom5          P1
## spacer_158 ENST00000256078 ENSG00000133703      fantom5          P1
## spacer_159 ENST00000256078 ENSG00000133703      fantom5          P1
## spacer_160 ENST00000256078 ENSG00000133703      fantom5          P1
##                         tss_id gene_symbol  tss_strand   tss_pos dist_to_tss
##                    <character> <character> <character> <integer>   <numeric>
## spacer_1    ENSG00000133703_P1        KRAS           -  25250928         491
## spacer_2    ENSG00000133703_P1        KRAS           -  25250928         490
## spacer_3    ENSG00000133703_P1        KRAS           -  25250928         489
## spacer_4    ENSG00000133703_P1        KRAS           -  25250928         482
## spacer_5    ENSG00000133703_P1        KRAS           -  25250928         479
## ...                        ...         ...         ...       ...         ...
## spacer_156 ENSG00000133703_P1        KRAS           -  25250928          10
## spacer_157 ENSG00000133703_P1        KRAS           -  25250928          15
## spacer_158 ENSG00000133703_P1        KRAS           -  25250928           4
## spacer_159 ENSG00000133703_P1        KRAS           -  25250928           1
## spacer_160 ENSG00000133703_P1        KRAS           -  25250928           3
```

#### Adding spacer alignments with TSS annotation

As with all CRISPR applications, potential off-targets effects are an important concern in assessing gRNA quality. While this concern is somewhat moderated for CRISPRi, since the dead CRISPR nuclease does not make DSBs, we should be aware of off-targetsoccuring in the promoter regions of other genes. This can be handled by passing our `tssObject` to the `addSpacerAlignments` function. We will search for up to 2 mismatches and increase the size of our `tss_window` (which defines the promoter region when searching for off-targets) to err on the safe side.

Similar to the CRISPRko design tutorial, we need to specify a Bowtie index of the human referenge genome; see the Building genome indices for short read aligners tutorial to learn how to create such an index.

Here, we specify the index that was available to us when generating this tutorial:

```
# Users need to specify the path of their bowtie index
index_path <- "/Users/fortinj2/crisprIndices/bowtie/hg38/hg38"
```

We are ready to add on- and off-target alignments:

```
gs <- addSpacerAlignments(gs,
                          aligner="bowtie",
                          aligner_index=index_path,
                          bsgenome=BSgenome.Hsapiens.UCSC.hg38,
                          n_mismatches=2,
                          tssObject=tss_human,
                          tss_window=c(-500, 2000))
```

```
gs
## GuideSet object with 160 ranges and 13 metadata columns:
##               seqnames    ranges strand |          protospacer           pam
##                  <Rle> <IRanges>  <Rle> |       <DNAStringSet> <DNAStringSet>
##      spacer_1    chr12  25250434      - | GCCGCGGCTGGAGGCTTCTG           GGG
##      spacer_2    chr12  25250435      - | AGCCGCGGCTGGAGGCTTCT           GGG
##      spacer_3    chr12  25250436      - | GAGCCGCGGCTGGAGGCTTC           TGG
##      spacer_4    chr12  25250443      - | TCCCCGAGAGCCGCGGCTGG           AGG
##      spacer_5    chr12  25250446      - | TCCTCCCCGAGAGCCGCGGC           TGG
##           ...      ...       ...    ... .                  ...           ...
##    spacer_156    chr12  25250915      - | ATTTTCCTAGGCGGCGGCCG           CGG
##    spacer_157    chr12  25250916      + | CGCTGCTGCCTCCGCCGCCG           CGG
##    spacer_158    chr12  25250921      - | AGCTCGATTTTCCTAGGCGG           CGG
##    spacer_159    chr12  25250924      - | CGGAGCTCGATTTTCCTAGG           CGG
##    spacer_160    chr12  25250928      + | CGCCGCCGCGGCCGCCGCCT           AGG
##                pam_site  cut_site      region        tssAnnotation        n0
##               <numeric> <numeric> <character> <SplitDataFrameList> <numeric>
##      spacer_1  25250434  25250437    region_1 chr12:25250437:-:...         1
##      spacer_2  25250435  25250438    region_1 chr12:25250438:-:...         1
##      spacer_3  25250436  25250439    region_1 chr12:25250439:-:...         1
##      spacer_4  25250443  25250446    region_1 chr12:25250446:-:...         1
##      spacer_5  25250446  25250449    region_1 chr12:25250449:-:...         1
##           ...       ...       ...         ...                  ...       ...
##    spacer_156  25250915  25250918    region_1 chr12:25250918:-:...         1
##    spacer_157  25250916  25250913    region_1 chr12:25250913:+:...         1
##    spacer_158  25250921  25250924    region_1 chr12:25250924:-:...         1
##    spacer_159  25250924  25250927    region_1 chr12:25250927:-:...         1
##    spacer_160  25250928  25250925    region_1 chr12:25250925:+:...         1
##                      n1        n2      n0_p      n1_p      n2_p
##               <numeric> <numeric> <numeric> <numeric> <numeric>
##      spacer_1         0         1         1         0         0
##      spacer_2         0         1         1         0         0
##      spacer_3         0         1         1         0         0
##      spacer_4         0         0         1         0         0
##      spacer_5         0         0         1         0         0
##           ...       ...       ...       ...       ...       ...
##    spacer_156         0         0         1         0         0
##    spacer_157         0        27         1         0        18
##    spacer_158         0         0         1         0         0
##    spacer_159         0         0         1         0         0
##    spacer_160        18       160         1        17       121
##                                                      alignments
##                                                   <GRangesList>
##      spacer_1                  chr12:25250434:-,chr7:155971918:-
##      spacer_2                  chr12:25250435:-,chr3:184602035:+
##      spacer_3                  chr12:25250436:-,chr3:184602034:+
##      spacer_4                                   chr12:25250443:-
##      spacer_5                                   chr12:25250446:-
##           ...                                                ...
##    spacer_156                                   chr12:25250915:-
##    spacer_157 chr12:25250916:+,chr2:55050346:+,chr20:21397165:+,...
##    spacer_158                                   chr12:25250921:-
##    spacer_159                                   chr12:25250924:-
```

```
##   spacer_160  chr12:25250928:+,chr17:49361951:+,chr1:3069055:-,...
##   -------
##   seqinfo: 640 sequences (1 circular) from hg38 genome
##   crisprNuclease: SpCas9
```

Including a `tssObject` parameter in the `addSpacerAlignments` function appends columns to the `GuideSet` that tallies the alignments restricted to the defined (via `tss_window`) promoter regions: `n0_p`, `n1_p`, and `n2_p` (the `_p` suffix denotes "promoter").

#### Adding CRISPRai scores

The CRISPRai algorithm was developed by the Weissman lab to score SpCas9 gRNAs for CRISPRa and CRISPRi applications for the human genome (Horlbeck et al. et al. 2016). The function `addCrispraiScores` implements this algorithm to add scores to the `GuideSet`. Compared to other on-target scoring algorithms, it requires several additional inputs:

- The `gr` argument is the `GRanges` object derived from the `queryTss` function and used to create the `GuideSet` object. In our example, this is the object named `target_region`.
- The `tssObject` argument is a `GRanges` object that contains TSS coordinates and annotation. It must also contain the following columns: `ID`, `promoter`, `tx_id`, and `gene_symbol`. Our `tssObject` in this instance is `tss_human`.
- `geneCol` indicates which column of `tssObject` should be used as the unique gene identifier.
- `modality` is the modality of the CRISPR application, in our case, `CRISPRi`.
- `fastaFile` is the path of a FASTA file containing the sequence of the human reference genome in hg38 coordinates. This file is available here.
- `chromatinFiles` is a vector of length 3 specifying the path of files containing the chromatin accessibility data needed for the algorithm in hg38 coordinates. The chromatin files can be downloaded from Zenodo here.

We first prepare all needed inputs for `addCrispraiScores`. We start by specifying the location of the FASTA file on our local machine:

```
fastaPath <- "/Users/fortinj2/crisprIndices/genomes/hg38/hg38.fa"
```

This corresponds to the path where the downloaded file from here is stored. Next, we specify the location of the chromatin files:

```
mnasePath <- "/Users/fortinj2/crisprIndices/chromatin/hg38/crispria_mnase_human_K562_hg38.bigWig"
dnasePath <- "/Users/fortinj2/crisprIndices/chromatin/hg38/crispria_dnase_human_K562_hg38.bigWig"
fairePath <- "/Users/fortinj2/crisprIndices/chromatin/hg38/crispria_faire_human_K562_hg38.bigWig"
chromatinFiles <- c(mnase=mnasePath,
                    dnase=dnasePath,
                    faire=fairePath)
```

This should correspond to the files that were downloaded from here.

We are now ready to add the scores:

```
results <- addCrispraiScores(gs,
                             gr=target_region,
                             tssObject=tss_human,
                             geneCol="gene_id",
                             modality="CRISPRi",
                             fastaFile=fastaPath,
                             chromatinFiles=chromatinFiles)
```

Let's look at the results:

```
results
## GuideSet object with 160 ranges and 14 metadata columns:
##               seqnames    ranges strand |          protospacer           pam
##                  <Rle> <IRanges>  <Rle> |       <DNAStringSet> <DNAStringSet>
##     spacer_1     chr12  25250434      - | GCCGCGGCTGGAGGCTTCTG           GGG
##     spacer_2     chr12  25250435      - | AGCCGCGGCTGGAGGCTTCT           GGG
##     spacer_3     chr12  25250436      - | GAGCCGCGGCTGGAGGCTTC           TGG
##     spacer_4     chr12  25250443      - | TCCCCGAGAGCCGCGGCTGG           AGG
##     spacer_5     chr12  25250446      - | TCCTCCCCGAGAGCCGCGGC           TGG
##          ...       ...       ...    ... .                  ...           ...
##   spacer_156     chr12  25250915      - | ATTTTCCTAGGCGGCGGCCG           CGG
##   spacer_157     chr12  25250916      + | CGCTGCTGCCTCCGCCGCCG           CGG
##   spacer_158     chr12  25250921      - | AGCTCGATTTTCCTAGGCGG           CGG
##   spacer_159     chr12  25250924      - | CGGAGCTCGATTTTCCTAGG           CGG
##   spacer_160     chr12  25250928      + | CGCCGCCGCGGCCGCCGCCT           AGG
##                 pam_site  cut_site      region        tssAnnotation        n0
##                <numeric> <numeric> <character> <SplitDataFrameList> <numeric>
##     spacer_1  25250434  25250437    region_1 chr12:25250437:-:...         1
##     spacer_2  25250435  25250438    region_1 chr12:25250438:-:...         1
##     spacer_3  25250436  25250439    region_1 chr12:25250439:-:...         1
##     spacer_4  25250443  25250446    region_1 chr12:25250446:-:...         1
##     spacer_5  25250446  25250449    region_1 chr12:25250449:-:...         1
##          ...       ...       ...         ...                  ...       ...
##   spacer_156  25250915  25250918    region_1 chr12:25250918:-:...         1
##   spacer_157  25250916  25250913    region_1 chr12:25250913:+:...         1
##   spacer_158  25250921  25250924    region_1 chr12:25250924:-:...         1
##   spacer_159  25250924  25250927    region_1 chr12:25250927:-:...         1
##   spacer_160  25250928  25250925    region_1 chr12:25250925:+:...         1
##                       n1        n2      n0_p      n1_p      n2_p
##                <numeric> <numeric> <numeric> <numeric> <numeric>
##     spacer_1         0         1         1         0         0
##     spacer_2         0         1         1         0         0
##     spacer_3         0         1         1         0         0
##     spacer_4         0         0         1         0         0
##     spacer_5         0         0         1         0         0
##          ...       ...       ...       ...       ...       ...
##   spacer_156         0         0         1         0         0
##   spacer_157         0        27         1         0        18
##   spacer_158         0         0         1         0         0
##   spacer_159         0         0         1         0         0
##   spacer_160        18       160         1        17       121
##                                                  alignments
##                                               <GRangesList>
##     spacer_1                  chr12:25250434:-,chr7:155971918:-
##     spacer_2                  chr12:25250435:-,chr3:184602035:+
##     spacer_3                  chr12:25250436:-,chr3:184602034:+
##     spacer_4                                   chr12:25250443:-
##     spacer_5                                   chr12:25250446:-
##          ...                                                ...
##   spacer_156                                   chr12:25250915:-
##   spacer_157 chr12:25250916:+,chr2:55050346:+,chr20:21397165:+,...
##   spacer_158                                   chr12:25250921:-
##   spacer_159                                   chr12:25250924:-
```

```
##   spacer_160  chr12:25250928:+,chr17:49361951:+,chr1:3069055:-,...
##              score_crispri
##                  <numeric>
##     spacer_1      0.372821
##     spacer_2      0.356982
##     spacer_3      0.390816
##     spacer_4      0.421704
##     spacer_5      0.408481
##          ...           ...
##   spacer_156      0.599555
##   spacer_157      0.666575
##   spacer_158      0.599259
##   spacer_159      0.565775
##   spacer_160      0.636552
##   -------
##   seqinfo: 640 sequences (1 circular) from hg38 genome
##   crisprNuclease: SpCas9
```

You can see that the column `score_crispri` was added to the `GuideSet`. Note that this function works identically for CRISPRa applications, with the `modality` argument replaced by `CRISPRa`.

### Session Info

```
sessionInfo()
```

```
## R version 4.2.1 (2022-06-23)
## Platform: x86_64-apple-darwin17.0 (64-bit)
## Running under: macOS Catalina 10.15.7
##
## Matrix products: default
## BLAS:   /Library/Frameworks/R.framework/Versions/4.2/Resources/lib/libRblas.0.dylib
## LAPACK: /Library/Frameworks/R.framework/Versions/4.2/Resources/lib/libRlapack.dylib
##
## locale:
## [1] en_US.UTF-8/en_US.UTF-8/en_US.UTF-8/C/en_US.UTF-8/en_US.UTF-8
##
## attached base packages:
## [1] stats4    stats     graphics  grDevices utils     datasets  methods
## [8] base
##
## other attached packages:
##  [1] BSgenome.Hsapiens.UCSC.hg38_1.4.4 BSgenome_1.65.2
##  [3] rtracklayer_1.57.0                Biostrings_2.65.2
##  [5] XVector_0.37.0                    GenomicRanges_1.49.1
##  [7] GenomeInfoDb_1.33.5               IRanges_2.31.2
##  [9] S4Vectors_0.35.1                  crisprDesignData_0.99.17
## [11] crisprDesign_0.99.133             crisprScore_1.1.14
## [13] crisprScoreData_1.1.3             ExperimentHub_2.5.0
## [15] AnnotationHub_3.5.0               BiocFileCache_2.5.0
## [17] dbplyr_2.2.1                      BiocGenerics_0.43.1
## [19] crisprBowtie_1.1.1                crisprBase_1.1.5
## [21] crisprVerse_0.99.8                rmarkdown_2.15.2
##
```

```
## loaded via a namespace (and not attached):
##  [1] rjson_0.2.21                  ellipsis_0.3.2
##  [3] Rbowtie_1.37.0                bit64_4.0.5
##  [5] lubridate_1.8.0               interactiveDisplayBase_1.35.0
##  [7] AnnotationDbi_1.59.1          fansi_1.0.3
##  [9] xml2_1.3.3                    codetools_0.2-18
## [11] cachem_1.0.6                  knitr_1.40
## [13] jsonlite_1.8.0                Rsamtools_2.13.4
## [15] png_0.1-7                     shiny_1.7.2
## [17] BiocManager_1.30.18           readr_2.1.2
## [19] compiler_4.2.1                httr_1.4.4
## [21] basilisk_1.9.2                assertthat_0.2.1
## [23] Matrix_1.4-1                  fastmap_1.1.0
## [25] cli_3.3.0                     later_1.3.0
## [27] htmltools_0.5.3               prettyunits_1.1.1
## [29] tools_4.2.1                   glue_1.6.2
## [31] GenomeInfoDbData_1.2.8        dplyr_1.0.9
## [33] rappdirs_0.3.3                tinytex_0.41
## [35] Rcpp_1.0.9                    Biobase_2.57.1
## [37] vctrs_0.4.1                   crisprBwa_1.1.3
## [39] xfun_0.32                     stringr_1.4.1
## [41] mime_0.12                     lifecycle_1.0.1
## [43] restfulr_0.0.15               XML_3.99-0.10
## [45] zlibbioc_1.43.0               basilisk.utils_1.9.1
## [47] vroom_1.5.7                   VariantAnnotation_1.43.3
## [49] hms_1.1.2                     promises_1.2.0.1
## [51] MatrixGenerics_1.9.1          parallel_4.2.1
## [53] SummarizedExperiment_1.27.1   RMariaDB_1.2.2
## [55] yaml_2.3.5                    curl_4.3.2
## [57] memoise_2.0.1                 reticulate_1.25
## [59] biomaRt_2.53.2                stringi_1.7.8
## [61] RSQLite_2.2.16                BiocVersion_3.16.0
## [63] highr_0.9                     BiocIO_1.7.1
## [65] randomForest_4.7-1.1          GenomicFeatures_1.49.6
## [67] filelock_1.0.2                BiocParallel_1.31.12
## [69] rlang_1.0.4                   pkgconfig_2.0.3
## [71] matrixStats_0.62.0            bitops_1.0-7
## [73] evaluate_0.16                 lattice_0.20-45
## [75] purrr_0.3.4                   GenomicAlignments_1.33.1
## [77] bit_4.0.4                     tidyselect_1.1.2
## [79] magrittr_2.0.3                R6_2.5.1
## [81] generics_0.1.3                DelayedArray_0.23.1
## [83] DBI_1.1.3                     pillar_1.8.1
## [85] KEGGREST_1.37.3               RCurl_1.98-1.8
## [87] tibble_3.1.8                  dir.expiry_1.5.0
## [89] crayon_1.5.1                  utf8_1.2.2
## [91] tzdb_0.3.0                    progress_1.2.2
## [93] grid_4.2.1                    blob_1.2.3
## [95] digest_0.6.29                 xtable_1.8-4
## [97] httpuv_1.6.5                  Rbwa_1.1.0
```

### References

Gilbert, Luke A, Matthew H Larson, Leonardo Morsut, Zairan Liu, Gloria A Brar, Sandra E Torres, Noam Stern-Ginossar, et al., et al. 2013. "CRISPR-Mediated Modular RNA-Guided Regulation of Transcription in Eukaryotes." *Cell* 154 (2): 442–51.

Horlbeck, Max A, Luke A Gilbert, Jacqueline E Villalta, Britt Adamson, Ryan A Pak, Yuwen Chen, Alexander P Fields, et al., et al. 2016. "Compact and Highly Active Next-Generation Libraries for CRISPR-Mediated Gene Repression and Activation." *Elife* 5.

Sanson, Kendall R, Ruth E Hanna, Mudra Hegde, Katherine F Donovan, Christine Strand, Meagan E Sullender, Emma W Vaimberg, et al., et al. 2018. "Optimized Libraries for CRISPR-Cas9 Genetic Screens with Multiple Modalities." *Nature Communications* 9 (1): 1–15.
