## Supplementary material for "The crisprVerse: a comprehensive Bioconductor ecosystem for the design of CRISPR guide RNAs across nucleases and technologies": Tutorial8_CRISPRbe_Design

### Using crisprDesign to design gRNAs for CRISPRbe

#### Introduction

In this tutorial, We illustrate the CRISPR base editing (CRISPRbe) functionalities of `crisprDesign` by designing and characterizing gRNAs targeting the human gene KRAS using the cytidine base editor BE4max (Koblan et al. 2018).

#### Installation

See the Installation tutorial to learn how to install the packages necessary for this tutorial: `crisprDesign`, `crisprDesignData`

#### Terminology

See the CRISPRko design tutorial to get familiar with the terminology used throughout this tutorial.

#### CRISPR base editing with BE4max

##### Loading packages

We first load the necessary packages for this tutorial:

```
library(crisprBase)
library(crisprDesign)
library(crisprDesignData)
library(BSgenome.Hsapiens.UCSC.hg38)
```

##### Creating the GuideSet

We first load the BE4max `BaseEditor` object from the `crisprBase` package:

```
data(BE4max, package="crisprBase")
BE4max
```

```
## Class: BaseEditor
##   CRISPR Nuclease name: SpCas9
##       Target type: DNA
##       Metadata: list of length 2
##       PAMs: NGG, NAG, NGA
##       Weights: 1, 0.2593, 0.0694
##       Spacer length: 20
##       PAM side: 3prime
##         Distance from PAM: 0
##       Prototype protospacers: 5'--SSSSSSSSSSSSSSSSSSSS[NGG]--3', 5'--SSSSSSSSSSSSSSSSSSSS[NAG]--3', 5
```

```
##   Base editor name: BE4max
##       Editing strand: original
##       Maximum editing weight: C2T at position -15
```

The editing probabilities of the base editor BE4max are stored in a matrix where rows correspond to the different nucleotide substitutions, and columns correspond to the genomic coordinate relative to the PAM site. The `editingWeights` function from `crisprBase` retrieves those probabilities. One can see that C to T editing is optimal around 15 nucleotides upstream of the PAM site for the BE4max base editor:

```
crisprBase::editingWeights(BE4max)["C2T",]
```

```
##   -36   -35   -34   -33   -32   -31   -30   -29   -28   -27   -26   -25   -24
## 0.007 0.007 0.008 0.018 0.010 0.020 0.014 0.012 0.023 0.013 0.024 0.022 0.034
##   -23   -22   -21   -20   -19   -18   -17   -16   -15   -14   -13   -12   -11
## 0.022 0.021 0.035 0.058 0.162 0.318 0.632 0.903 1.000 0.870 0.620 0.314 0.163
##   -10    -9    -8    -7    -6    -5    -4    -3    -2    -1
## 0.100 0.056 0.033 0.019 0.018 0.024 0.017 0.005 0.002 0.001
```

Let's create the `GuideSet` containing gRNAs targeting KRAS.

We first load the data containing gene regions for the human genome from `crisprDesignData`:

```
data("txdb_human", package="crisprDesignData")
```

For more information on `txdb_human` and how to create similar gene annotation objects, see the Building a gene annotation object tutorial.

We will also load the `BSgenome` package containing DNA sequences for the hg38 genome:

```
library(BSgenome.Hsapiens.UCSC.hg38)
```

We retrive the genomic coordinates of the KRAS CDS

```
gr <- queryTxObject(txObject=txdb_human,
                    featureType="cds",
                    queryColumn="gene_symbol",
                    queryValue="KRAS")
```

and design all possigle gRNAs using the function `findSpacers`:

```
bsgenome <- BSgenome.Hsapiens.UCSC.hg38
gs <- findSpacers(gr,
                  bsgenome=bsgenome,
                  crisprNuclease=BE4max)
```

#### Annotating the GuideSet

Next, we annotate our candidate gRNAs to assess quality. There are several functions in `crisprDesign` that provide annotation for features that are nonspecific to CRISPRbe, for which we refer the reader to the CRISPRko design with Cas9 tutorial for more information. The sections below will cover annotation functions that are of particular interest to, or deserve extra care for CRISPRbe applications.

##### Adding edited alleles

The function `addEditedAlleles` finds, characterizes, and scores predicted edited alleles for each gRNA and a chosen transcript. It requires a transcript-specific annotation that can be obtained with the `getTxInfoDataFrame` function. Here, we perform the analysis using the primary isoform of KRAS (Ensembl transcript ID: ENST00000311936).

We first get the transcript table for our transcript

```r
txid <- "ENST00000311936"
txTable <- getTxInfoDataFrame(tx_id=txid,
                              txObject=txdb_human,
                              bsgenome=bsgenome)
head(txTable)
```

```
## DataFrame with 6 rows and 10 columns
##           chr       pos         nuc          aa aa_number      exon  pos_plot
##   <character> <numeric> <character> <character> <integer> <integer> <integer>
## 1       chr12  25250929           C          NA        NA         1        31
## 2       chr12  25250928           T          NA        NA         1        32
## 3       chr12  25250927           A          NA        NA         1        33
## 4       chr12  25250926           G          NA        NA         1        34
## 5       chr12  25250925           G          NA        NA         1        35
## 6       chr12  25250924           C          NA        NA         1        36
##    pos_mrna   pos_cds      region
##   <integer> <integer> <character>
## 1         1        NA        5UTR
## 2         2        NA        5UTR
## 3         3        NA        5UTR
## 4         4        NA        5UTR
## 5         5        NA        5UTR
## 6         6        NA        5UTR
```

and then add the edited alleles annotation to the `GuideSet`:

```r
editingWindow <- c(-20,-8)
gs <- addEditedAlleles(gs,
                       baseEditor=BE4max,
                       txTable=txTable,
                       editingWindow=editingWindow)
```

```
## [addEditedAlleles] Obtaining edited alleles at each gRNA target site.

## [addEditedAlleles] Adding functional consequences to alleles.
```

The `editingWindow` argument specifies the window of editing that we are interested in. When not provided, it uses the default window provided in the `BaseEditor` object. Note that providing large windows can exponentially increase computing time as the number of possible alleles grows exponentially.

Let's retrieve the edited alleles for the first gRNA:

```r
alleles <- editedAlleles(gs)[[1]]
```

We get a `DataFrame` object with useful metadata:

```r
metadata(alleles)
```

```
## $wildtypeAllele
##        spacer_1
## "AAAGAAAAGATGA"
##
## $start
## [1] 25209851
##
## $end
## [1] 25209863
##
```

```
## $chr
## [1] "chr12"
##
## $strand
## [1] "-"
##
## $editingWindow
## [1] -20  -8
##
## $wildtypeAmino
## [1] "SMMMKKKEEEKKK"
```

The `wildtypeAllele` reports the unedited nucleotide sequence of the region specified by the editing window (with respect to the gRNA PAM site). It is always reported from the 5' to 3' direction on the strand corresponding to the gRNA strand. The `start` and `end` fields specify the corresponding coordinates on the transcript.

Let's look at the edited alleles:

```
head(alleles)
```

```
## DataFrame with 6 rows and 4 columns
##               seq       score     variant            aa
##    <DNAStringSet>   <numeric> <character>   <character>
## 1  AAAGAAAACATGA 0.000922509    missense SMMMNNNEEEKKK
## 2  AAAAAAAAAATAA 0.000000000    missense SIIIKKKKKKKKK
## 3  AAACAAAAAATAA 0.000000000    missense SIIIKKKQQQKKK
## 4  AAAGAAAAAATAA 0.000000000    missense SIIIKKKEEEKKK
## 5  AAAAAAAACATAA 0.000000000    missense SIIINNNKKKKKK
## 6  AAACAAAACATAA 0.000000000    missense SIIINNNQQQKKK
```

The `DataFrame` is ordered by descending values in the `score` column. This `score` represents the likelihood of the edited allele to occur relative to all possible edited alleles, and is calculated using the editing weights stored in the `BE4max` object. The `seq` column represents the edited nucleotide sequences. As with the `wildtypeAllele` in the metadata, they are always reported from the 5' to 3' direction on the strand corresponding to the gRNA strand.

The `variant` column describes the functional consequence of the editing event (silent, nonsense or missense mutation). If an edited allele results in multiple editing events, as can happen when multiple bases are edited, the most consequential mutation (nonsense over missense, missense over silent) is reported. Finally, the `aa` column reports the resulting edited amino acid sequence, with each single letter code mapping to its corresponding nucleotide (`*` for termination).

Note that `addEditedAlleles` also appended several gRNA-level aggregate scores to the `GuideSet` object:

```
head(gs)
```

```
## GuideSet object with 6 ranges and 11 metadata columns:
##             seqnames    ranges strand |          protospacer            pam
##                <Rle> <IRanges>  <Rle> |       <DNAStringSet> <DNAStringSet>
##   spacer_1    chr12  25209843      - | AAAGAAAAGATGAGCAAAGA            TGG
##   spacer_2    chr12  25209843      - | AAAGAAAAGATGAGCAAAGA            TGG
##   spacer_3    chr12  25209896      + | TTCTCGAACTAATGTATAGA            AGG
##   spacer_4    chr12  25209896      + | TTCTCGAACTAATGTATAGA            AGG
##   spacer_5    chr12  25215438      - | AAATGCATTATAATGTAATC            TGG
##   spacer_6    chr12  25215477      - | AGCAAAGAAGAAAAGACTCC            TGG
##              pam_site  cut_site      region
##             <numeric> <numeric> <character>
```

```
##   spacer_1  25209843  25209846    region_8
##   spacer_2  25209843  25209846   region_10
##   spacer_3  25209896  25209893    region_8
##   spacer_4  25209896  25209893   region_10
##   spacer_5  25215438  25215441    region_4
##   spacer_6  25215477  25215480    region_4
##
##
##   spacer_1 AAAGAAAACATGA:0.000922509:missense:...,AAAAAAAAAATAA:0.000000000:missense:...,AAACAAAAAAT
##   spacer_2 AAAGAAAACATGA:0.000922509:missense:...,AAAAAAAAAATAA:0.000000000:missense:...,AAACAAAAAAT
##   spacer_3          TTCTTGAACTAAT:0.392378:missense:...,TTTTTGAACTAAT:0.167936:missense:...,TTCTTGAA
##   spacer_4          TTCTTGAACTAAT:0.392378:missense:...,TTTTTGAACTAAT:0.167936:missense:...,TTCTTGAA
##   spacer_5        AAATGTATTATAA:0.9199562:not_targeting,AAATGGATTATAA:0.0496776:not_targeting,AAATGA
##   spacer_6     AGTAAAGAAGAAA:0.29281668:not_targeting,AGGAAAGAAGAAA:0.01197049:not_targeting,AGAAAAG
##            score_missense score_nonsense score_silent    maxVariant
##                 <numeric>      <numeric>    <numeric>   <character>
##   spacer_1    0.000922509     0.00000000            0      missense
##   spacer_2    0.000922509     0.00000000            0      missense
##   spacer_3    0.944195218     0.00931016            0      missense
##   spacer_4    0.944195218     0.00931016            0      missense
##   spacer_5    0.000000000     0.00000000            0 not_targeting
##   spacer_6    0.000000000     0.00000000            0 not_targeting
##            maxVariantScore
##                  <numeric>
##   spacer_1     0.000922509
##   spacer_2     0.000922509
##   spacer_3     0.944195218
##   spacer_4     0.944195218
##   spacer_5              NA
##   spacer_6              NA
##   -------
##   seqinfo: 640 sequences (1 circular) from hg38 genome
##   crisprNuclease: SpCas9
```

The `score_missense`, `score_nonsense` and `score_silent` columns report aggregated scores for each mutation type. They are calculated by summing all scores of a given mutation type across the set of edited alleles for a given gRNA. The `maxVariant` column indicates the most probable mutation type for the given gRNA based on the maximum aggregated score, which is stored in `maxVariantScore`. In our example, the highest score for `spacer_4` is `score_nonsense`, and so `maxVariant` is set to `nonsense`.

### References

Koblan, Luke W, Jordan L Doman, Christopher Wilson, Jonathan M Levy, Tristan Tay, Gregory A Newby, Juan Pablo Maianti, Aditya Raguram, and David R Liu. 2018. "Improving Cytidine and Adenine Base Editors by Expression Optimization and Ancestral Reconstruction." *Nature Biotechnology* 36 (9): 843–46.
