## Supplementary material for "The crisprVerse: a comprehensive Bioconductor ecosystem for the design of CRISPR guide RNAs across nucleases and technologies": Tutorial9_CRISPRa_Design

### gRNA design for CRISPR activation (CRISPRa)

#### Introduction

This tutorial will demonstrate how to use `crisprDesign` to design gRNAs for CRISPR activation (CRISPRa). Specifically, we will target the human KRAS gene and use the SpCas9 nuclease.

#### Installation

See the Installation tutorial to learn how to install the packages necessary for this tutorial: `crisprDesign`, `crisprDesignData`

#### Terminology

See the CRISPRko design tutorial to get familiar with the terminology used throughout this tutorial.

#### CRISPRa design

For CRISPR activation (CRISPRa) and interference (CRISPRi) applications, the CRISPR nuclease is engineered to lose its endonuclease activity, and should therefore not introduce double-stranded breaks (DSBs). We will use the dead SpCas9 (dSpCas9) nuclease as an example here. Note that users don't have to distinguish between dSpCas9 and SpCas9 when specifying the nuclease in the crisprVerse as they do not differ in terms of the characteristics stored in the `CrisprNuclease` object.

In CRISPRa, dSpCas9 is used to activate gene expression by coupling the dead nuclease with activation factors. Several CRISPRa systems have been developed (see Kampmann (2018) for a review). For optimal activation, gRNAs are usually designed to target the region directly upstream of the gene transcription start site (TSS).

`crisprDesign` provides functionalities to be able to take into account design rules that are specific to CRISPRa applications. The `queryTss` function allows for specifying genomic coordinates of promoter regions. The `addTssAnnotation` function annotates gRNAs for known TSSs, and includes a column `dist_to_tss` that gives the distance in nucleotides between the TSS position and the PAM site of the gRNA. For CRISPRa, we recommend targeting the region 75-150bp upstream of the TSS for optimal activation; see Sanson et al. et al. (2018) for more information. Finally, the function `addCrispraiScores` adds on-target activity scores based on the work of (Horlbeck et al. et al. 2016).

##### Creating the GuideSet

We first start by loading the required packages:

```
library(crisprBase)
library(crisprDesign)
library(crisprDesignData)
library(BSgenome.Hsapiens.UCSC.hg38)
```

To demonstrate CRISPRa design, we will design gRNAs to activate the human KRAS gene using the SpCas9 nuclease. To accomplish this, we want our gRNAs to target the region upstream of the KRAS TSS; let's consider the window containing 500bp immediately upstream of the TSS. We first need to retrieve the TSS coordinates for KRAS. These data are conveniently stored in the `crisprDesignData` package as the dataset `tss_human`. For more information on `tss_human` and how to create similar TSS annotation objects, see the Building a gene annotation object tutorial.

We load the TSS coordinates stored in the `tss_human` object

```
data("tss_human", package="crisprDesignData")
```

and query for KRAS using the `queryTss` function from `crisprDesign`:

```
target_window <- c(-500, 0)
target_region <- queryTss(tss_human,
                          queryColumn="gene_symbol",
                          queryValue="KRAS",
                          tss_window=target_window)
```

```
target_region
## GRanges object with 1 range and 9 metadata columns:
##            seqnames            ranges strand |     score peak_start  peak_end
##               <Rle>         <IRanges>  <Rle> | <numeric>  <integer> <integer>
##   region_1    chr12 25250929-25251428      - |   5.20187   25250928  25250928
##                       tx_id         gene_id      source    promoter
##                 <character>     <character> <character> <character>
##   region_1 ENST00000256078 ENSG00000133703     fantom5          P1
##                          ID gene_symbol
##                 <character> <character>
##   region_1 ENSG00000133703_P1        KRAS
##   -------
##   seqinfo: 25 sequences from an unspecified genome; no seqlengths
```

We load the `crisprNuclease` object storing information about the SpCas9 nuclease from the `crisprBase` package:

```
data(SpCas9, package="crisprBase")
```

We then find all candidate protospacer sequences in our target region with `findSpacers`:

```
gs <- findSpacers(target_region,
                  crisprNuclease=SpCas9,
                  bsgenome=BSgenome.Hsapiens.UCSC.hg38)
```

```
gs
## GuideSet object with 146 ranges and 5 metadata columns:
##             seqnames    ranges strand |          protospacer            pam
##                <Rle> <IRanges>  <Rle> |       <DNAStringSet> <DNAStringSet>
##     spacer_1    chr12  25250927      - | GCTCGGAGCTCGATTTTCCT            AGG
##     spacer_2    chr12  25250944      - | CCCGAACTCATCGGTGTGCT            CGG
##     spacer_3    chr12  25250953      - | CCGCCCGGCCCCGAACTCAT            CGG
##     spacer_4    chr12  25250961      + | TCCGAGCACACCGATGAGTT            CGG
##     spacer_5    chr12  25250962      + | CCGAGCACACCGATGAGTTC            GGG
##          ...      ...       ...    ... .                  ...            ...
##   spacer_142    chr12  25251419      - | AGGCCGACCCTGAGGGTGGC            GGG
##   spacer_143    chr12  25251420      - | TAGGCCGACCCTGAGGGTGG            CGG
##   spacer_144    chr12  25251423      - | GTATAGGCCGACCCTGAGGG            TGG
##   spacer_145    chr12  25251429      + | AAGAGCACCCCGCCACCCTC            AGG
```

```
##   spacer_146    chr12  25251430      + | AGAGCACCCCGCCACCCTCA            GGG
##              pam_site  cut_site      region
##             <numeric> <numeric> <character>
##     spacer_1  25250927  25250930    region_1
##     spacer_2  25250944  25250947    region_1
##     spacer_3  25250953  25250956    region_1
##     spacer_4  25250961  25250958    region_1
##     spacer_5  25250962  25250959    region_1
##          ...       ...       ...         ...
##   spacer_142  25251419  25251422    region_1
##   spacer_143  25251420  25251423    region_1
##   spacer_144  25251423  25251426    region_1
##   spacer_145  25251429  25251426    region_1
##   spacer_146  25251430  25251427    region_1
##   -------
##   seqinfo: 640 sequences (1 circular) from hg38 genome
##   crisprNuclease: SpCas9
```

#### Annotating the GuideSet

Next, we annotate our candidate gRNAs to assess quality. There are several functions in `crisprDesign` that provide annotation for features that are not specific to CRISPRa, for which we refer the reader to the CRISPRko design with Cas9 tutorial for more information. The sections below will cover annotation functions that are of particular interest to CRISPRa applications.

#### Adding TSS annotation

As the name implies, the `addTssAnnotation` function annotates gRNAs with TSS context such as the distance between the gRNA and the TSS, as well as which TSS is targeted (many genes contain different TSSs corresponding to different isoforms).

The function requires a `tssObject` object, and the `tss_window` values that we used earlier to define the target region. We can then retrieve the appended annotation with the accessor function `tssAnnotation`:

```
gs <- addTssAnnotation(gs,
                       tssObject=tss_human,
                       tss_window=target_window)
tssAnnotation(gs)
## DataFrame with 146 rows and 15 columns
##                  chr anchor_site   strand     score peak_start  peak_end
##             <factor>   <integer> <factor> <numeric>  <integer> <integer>
## spacer_1      chr12    25250930        -   5.20187   25250928  25250928
## spacer_2      chr12    25250947        -   5.20187   25250928  25250928
## spacer_3      chr12    25250956        -   5.20187   25250928  25250928
## spacer_4      chr12    25250958        +   5.20187   25250928  25250928
## spacer_5      chr12    25250959        +   5.20187   25250928  25250928
## ...             ...         ...      ...       ...        ...       ...
## spacer_142    chr12    25251422        -   5.20187   25250928  25250928
## spacer_143    chr12    25251423        -   5.20187   25250928  25250928
## spacer_144    chr12    25251426        -   5.20187   25250928  25250928
## spacer_145    chr12    25251426        +   5.20187   25250928  25250928
## spacer_146    chr12    25251427        +   5.20187   25250928  25250928
##                      tx_id         gene_id      source    promoter
##                <character>     <character> <character> <character>
```

```
## spacer_1   ENST00000256078 ENSG00000133703     fantom5          P1
## spacer_2   ENST00000256078 ENSG00000133703     fantom5          P1
## spacer_3   ENST00000256078 ENSG00000133703     fantom5          P1
## spacer_4   ENST00000256078 ENSG00000133703     fantom5          P1
## spacer_5   ENST00000256078 ENSG00000133703     fantom5          P1
## ...                    ...             ...         ...         ...
## spacer_142 ENST00000256078 ENSG00000133703     fantom5          P1
## spacer_143 ENST00000256078 ENSG00000133703     fantom5          P1
## spacer_144 ENST00000256078 ENSG00000133703     fantom5          P1
## spacer_145 ENST00000256078 ENSG00000133703     fantom5          P1
## spacer_146 ENST00000256078 ENSG00000133703     fantom5          P1
##                         tss_id gene_symbol  tss_strand   tss_pos dist_to_tss
##                    <character> <character> <character> <integer>   <numeric>
## spacer_1   ENSG00000133703_P1         KRAS           -  25250928          -2
## spacer_2   ENSG00000133703_P1         KRAS           -  25250928         -19
## spacer_3   ENSG00000133703_P1         KRAS           -  25250928         -28
## spacer_4   ENSG00000133703_P1         KRAS           -  25250928         -30
## spacer_5   ENSG00000133703_P1         KRAS           -  25250928         -31
## ...                       ...          ...         ...       ...         ...
## spacer_142 ENSG00000133703_P1         KRAS           -  25250928        -494
## spacer_143 ENSG00000133703_P1         KRAS           -  25250928        -495
## spacer_144 ENSG00000133703_P1         KRAS           -  25250928        -498
## spacer_145 ENSG00000133703_P1         KRAS           -  25250928        -498
## spacer_146 ENSG00000133703_P1         KRAS           -  25250928        -499
```

##### Adding spacer alignments with TSS annotation

As with all CRISPR applications, off-targets is an important concern in assessing gRNA quality. While this concern is somewhat moderated for CRISPRa, since the dead CRISPR nuclease does not make DSBs, we should be aware of off-targets occuring in the promoter regions of other genes. This can be handled by passing our `tssObject` to the `addSpacerAlignments` function. We will search for up to 2 mismatches and increase the size of our `tss_window` to err on the safe side.

Similar to the CRISPRko design tutorial, we need to specify a Bowtie index of the human referenge genome; see the Building genome indices for short read aligners tutorial to learn how to create such an index.

Here we specify the index that was available to us when generating this tutorial:

```
index_path <- "/Users/fortinj2/crisprIndices/bowtie/hg38/hg38"
```

(this needs to be changed by users). We are now ready to add on- and off-target alignments:

```
gs <- addSpacerAlignments(gs,
                          aligner="bowtie",
                          aligner_index=index_path,
                          bsgenome=BSgenome.Hsapiens.UCSC.hg38,
                          n_mismatches=2,
                          tssObject=tss_human,
                          tss_window=c(-2000, 500))
```

```
gs
## GuideSet object with 146 ranges and 13 metadata columns:
##              seqnames    ranges strand |          protospacer            pam
##                 <Rle> <IRanges>  <Rle> |       <DNAStringSet> <DNAStringSet>
##     spacer_1    chr12  25250927      - | GCTCGGAGCTCGATTTTCCT            AGG
##     spacer_2    chr12  25250944      - | CCCGAACTCATCGGTGTGCT            CGG
```

```
##     spacer_3    chr12  25250953      - | CCGCCCGGCCCCGAACTCAT           CGG
##     spacer_4    chr12  25250961      + | TCCGAGCACACCGATGAGTT           CGG
##     spacer_5    chr12  25250962      + | CCGAGCACACCGATGAGTTC           GGG
##          ...      ...        ...    ... .                  ...           ...
##   spacer_142    chr12  25251419      - | AGGCCGACCCTGAGGGTGGC           GGG
##   spacer_143    chr12  25251420      - | TAGGCCGACCCTGAGGGTGG           CGG
##   spacer_144    chr12  25251423      - | GTATAGGCCGACCCTGAGGG           TGG
##   spacer_145    chr12  25251429      + | AAGAGCACCCCGCCACCCTC           AGG
##   spacer_146    chr12  25251430      + | AGAGCACCCCGCCACCCTCA           GGG
##               pam_site  cut_site      region        tssAnnotation        n0
##              <numeric> <numeric> <character> <SplitDataFrameList> <numeric>
##     spacer_1  25250927  25250930    region_1 chr12:25250930:-:...         1
##     spacer_2  25250944  25250947    region_1 chr12:25250947:-:...         1
##     spacer_3  25250953  25250956    region_1 chr12:25250956:-:...         1
##     spacer_4  25250961  25250958    region_1 chr12:25250958:+:...         1
##     spacer_5  25250962  25250959    region_1 chr12:25250959:+:...         1
##          ...       ...       ...         ...                  ...       ...
##   spacer_142  25251419  25251422    region_1 chr12:25251422:-:...         1
##   spacer_143  25251420  25251423    region_1 chr12:25251423:-:...         1
##   spacer_144  25251423  25251426    region_1 chr12:25251426:-:...         1
##   spacer_145  25251429  25251426    region_1 chr12:25251426:+:...         1
##   spacer_146  25251430  25251427    region_1 chr12:25251427:+:...         1
##                     n1        n2      n0_p      n1_p      n2_p
##              <numeric> <numeric> <numeric> <numeric> <numeric>
##     spacer_1         0         0         1         0         0
##     spacer_2         0         0         1         0         0
##     spacer_3         0         0         1         0         0
##     spacer_4         0         0         1         0         0
##     spacer_5         0         0         1         0         0
##          ...       ...       ...       ...       ...       ...
##   spacer_142         0         1         1         0         0
##   spacer_143         0         0         1         0         0
##   spacer_144         0         0         1         0         0
##   spacer_145         0         0         1         0         0
##   spacer_146         0         0         1         0         0
##                                      alignments
##                                  <GRangesList>
##     spacer_1                chr12:25250927:-
##     spacer_2                chr12:25250944:-
##     spacer_3                chr12:25250953:-
##     spacer_4                chr12:25250961:+
##     spacer_5                chr12:25250962:+
##          ...                             ...
##   spacer_142 chr12:25251419:-,chr16:88925550:+
##   spacer_143                chr12:25251420:-
##   spacer_144                chr12:25251423:-
##   spacer_145                chr12:25251429:+
##   spacer_146                chr12:25251430:+
##   -------
##   seqinfo: 640 sequences (1 circular) from hg38 genome
##   crisprNuclease: SpCas9
```

This should correspond to the files that were downloaded from here.

We are now ready to add the scores:

```
results <- addCrispraiScores(gs,
                             gr=target_region,
                             tssObject=tss_human,
                             geneCol="gene_id",
                             modality="CRISPRa",
                             fastaFile=fastaPath,
                             chromatinFiles=chromatinFiles)
```

Let's look at the results:

```
results
## GuideSet object with 146 ranges and 14 metadata columns:
##            seqnames    ranges strand |          protospacer           pam
##               <Rle> <IRanges>  <Rle> |       <DNAStringSet> <DNAStringSet>
##     spacer_1    chr12  25250927      - | GCTCGGAGCTCGATTTTCCT           AGG
##     spacer_2    chr12  25250944      - | CCCGAACTCATCGGTGTGCT           CGG
##     spacer_3    chr12  25250953      - | CCGCCCGGCCCCGAACTCAT           CGG
```

```
##     spacer_4    chr12  25250961      + | TCCGAGCACACCGATGAGTT            CGG
##     spacer_5    chr12  25250962      + | CCGAGCACACCGATGAGTTC            GGG
##          ...      ...       ...    ... .                  ...            ...
##   spacer_142    chr12  25251419      - | AGGCCGACCCTGAGGGTGGC            GGG
##   spacer_143    chr12  25251420      - | TAGGCCGACCCTGAGGGTGG            CGG
##   spacer_144    chr12  25251423      - | GTATAGGCCGACCCTGAGGG            TGG
##   spacer_145    chr12  25251429      + | AAGAGCACCCCGCCACCCTC            AGG
##   spacer_146    chr12  25251430      + | AGAGCACCCCGCCACCCTCA            GGG
##              pam_site  cut_site      region        tssAnnotation        n0
##             <numeric> <numeric> <character> <SplitDataFrameList> <numeric>
##     spacer_1  25250927  25250930    region_1 chr12:25250930:-:...         1
##     spacer_2  25250944  25250947    region_1 chr12:25250947:-:...         1
##     spacer_3  25250953  25250956    region_1 chr12:25250956:-:...         1
##     spacer_4  25250961  25250958    region_1 chr12:25250958:+:...         1
##     spacer_5  25250962  25250959    region_1 chr12:25250959:+:...         1
##          ...       ...       ...         ...                  ...       ...
##   spacer_142  25251419  25251422    region_1 chr12:25251422:-:...         1
##   spacer_143  25251420  25251423    region_1 chr12:25251423:-:...         1
##   spacer_144  25251423  25251426    region_1 chr12:25251426:-:...         1
##   spacer_145  25251429  25251426    region_1 chr12:25251426:+:...         1
##   spacer_146  25251430  25251427    region_1 chr12:25251427:+:...         1
##                    n1        n2      n0_p      n1_p      n2_p
##             <numeric> <numeric> <numeric> <numeric> <numeric>
##     spacer_1         0         0         1         0         0
##     spacer_2         0         0         1         0         0
##     spacer_3         0         0         1         0         0
##     spacer_4         0         0         1         0         0
##     spacer_5         0         0         1         0         0
##          ...       ...       ...       ...       ...       ...
##   spacer_142         0         1         1         0         0
##   spacer_143         0         0         1         0         0
##   spacer_144         0         0         1         0         0
##   spacer_145         0         0         1         0         0
##   spacer_146         0         0         1         0         0
##                                    alignments score_crispra
##                                 <GRangesList>     <numeric>
##     spacer_1                 chr12:25250927:-      0.439319
##     spacer_2                 chr12:25250944:-      0.392932
##     spacer_3                 chr12:25250953:-      0.477453
##     spacer_4                 chr12:25250961:+      0.437693
##     spacer_5                 chr12:25250962:+      0.437368
##          ...                              ...           ...
##   spacer_142 chr12:25251419:-,chr16:88925550:+      0.339499
##   spacer_143                 chr12:25251420:-      0.377727
##   spacer_144                 chr12:25251423:-      0.387729
##   spacer_145                 chr12:25251429:+      0.362817
##   spacer_146                 chr12:25251430:+      0.363131
##   -------
##   seqinfo: 640 sequences (1 circular) from hg38 genome
##   crisprNuclease: SpCas9
```

You can see that the column `score_crispra` was added to the `GuideSet`. Note that this function works identically for CRISPRi applications, with the `modality` argument replaced by `CRISPRi`.

### Session Info

```
sessionInfo()
```

Kampmann, Martin. 2018. "CRISPRi and CRISPRa Screens in Mammalian Cells for Precision Biology and Medicine." *ACS Chemical Biology* 13 (2): 406–16.

Sanson, Kendall R, Ruth E Hanna, Mudra Hegde, Katherine F Donovan, Christine Strand, Meagan E Sullender, Emma W Vaimberg, et al., et al. 2018. "Optimized Libraries for CRISPR-Cas9 Genetic Screens with Multiple Modalities." *Nature Communications* 9 (1): 1–15.
