## Supplementary material for "The crisprVerse: a comprehensive Bioconductor ecosystem for the design of CRISPR guide RNAs across nucleases and technologies": Tutorial2_Building_Genome_Index

### Building genome indices off-target alignment

#### Introduction

This vignette demonstrates how to build genome indices for the purpose of performing on- and off-target alignment. In particular, we show how to build such indices for the short read aligners bowtie (Langmead et al. 2009), as used by the `Rbowtie` and `crisprBowtie` packages, and BWA-backtrack (Li and Durbin 2009), as used by the `Rbwa` and `crisprBwa` packages. Note that BWA is not available for Windows users.

Generating a genome index file is time consuming, but only needs to be done once for a given genome.

#### Installation

See the Installation tutorial to learn how to install the `crisprBowtie` and `crisprBwa` packages.

#### Building a bowtie index

In the following example, we build a bowtie index for the human genome using the hg38 build. First, users will need to donwload the FASTA file from the UCSC genome browser. Here's the link: https://hgdownload.soe.ucsc.edu/goldenPath/hg38/bigZips/hg38.fa.gz

Next, assuming the `hg38.fa.gz` is located in the current directory, we build the bowtie genome index using the function `bowtie_build` from the `Rbowtie` package (which is installed when `crisprBowtie` is installed):

```
library(Rbowtie)
fastaFile <- "./hg38.fa.gz"
bowtie_build(fastaFile,
             outdir="./",
             force=TRUE,
             prefix="hg38")
```

This should take a couple of hours to run, and the resulting bowtie index files will be located in the folder `./hg38` and can be used to run bowtie alignment. See the crisprBowtie package to learn how to perform a bowtie alignment within R.

#### Building a BWA index

Building a BWA index is similar to building a bowtie index. Assuming the `hg38.fa.gz` is located in the current directory, we build the BWA genome index using the function `bwa_build_index` from the `Rbwa` package (which is installed when `crisprBwa` is installed):

```
library(Rbwa)
fastaFile <- "./hg38.fa.gz"
bwa_build_index(fastaFile,
                index_prefix="hg38")
```

This should take a couple of hours to run, and the resulting BWA index files will be located in the folder `./hg38` and can be used to run BWA alignment. See the crisprBwa package to learn how to perform a BWA alignment within R.

### Building a transcriptome index

For applications using RNA-targeting nucleases such as CasRx, off-target search is performed against against transcriptomes rather than genomes. Building a transcriptome index works similar, except that we first need to generate a FASTA file containing the transcriptome sequences. This is easily accomplished with the function `getMrnaSequences` from the `crisprDesign` package, assuming that a gene model is provided, as well as a `BSgenome` object containing the DNA sequences for the hg38 genome (`BSgenome.Hsapiens.UCSC.hg38`).

We first load the necessary packages

```
library(BSgenome.Hsapiens.UCSC.hg38)
library(crisprDesign)
```

The `crisprDesignData` package (see Installation) contains a gene model annotation for the hg38 genome, and can be loaded using the following:

```
library(crisprDesignData)
data("txdb_human", package="crisprDesignData")
```

See the Gene annotation tutorial to learn more about how to build such gene annotation objects.

We will now extract mRNA sequences for all available transcripts:

```
txids <- unique(txdb_human$exons$tx_id)
mrnasHuman <- getMrnaSequences(txids,
                               bsgenome=BSgenome.Hsapiens.UCSC.hg38,
                               txObject=txdb_human)
```

This should take less than an hour to run. Once completed, we will write the extracted mRNA sequences to disk using the FASTA format. This can be accomplished using the `writeXStringSet` function from the `Biostrings` package:

```
library(Biostrings)
writeXStringSet(mrnasHuman,
                file="ensembl_human_104.fasta",
                format="fasta")
```

Note that the `seqnames` of this FASTA file are Ensembl transcript IDs instead of chromosomes. Once the FASTA file has been generated, the process for constructing either a bowtie or BWA index file is the same as described in the above sections.

### References


Langmead, Ben, Cole Trapnell, Mihai Pop, and Steven L. Salzberg. 2009. "Ultrafast and Memory-Efficient Alignment of Short DNA Sequences to the Human Genome." *Genome Biology* 10 (3): R25. https://doi.org/10.1186/gb-2009-10-3-r25.

Li, Heng, and Richard Durbin. 2009. "Fast and Accurate Short Read Alignment with Burrows–Wheeler Transform." *Bioinformatics* 25 (14): 1754–60.
