## Supplementary material for "The crisprVerse: a comprehensive Bioconductor ecosystem for the design of CRISPR guide RNAs across nucleases and technologies": Tutorial4_Building_Gene_Annotation

### Building a gene annotation object

#### Introduction

In this tutorial, we describe the process for making and using rich gene annotation objects to be used throughout the crisprVerse ecosystem. Such objects enable users to retrieve coordinates of transcripts, exons, etc. Those objects are also used by several functions in the crisprDesign package to add gene annotations to both gRNA on-targets and off-targets. This is what the `txObject` argument in many of the functions expect.

We will also describe the process for constructing and using a transcription start site (TSS) annotation object (`tssObject` argument in many of the functions).

#### Installation

See the Installation tutorial to learn how to install the packages `crisprDesign` and `crisprDesignData` required in this tutorial.

##### Getting started

The packages can be loaded into an R session in the usual way:

```
library(crisprDesign)
library(crisprDesignData)
```

#### Building gene annotation objects

In the crisprVerse, we represent gene annotations using `GRangesList` object, and this can be easily constructed using the commonly-used Bioconductor objects `TxDb` (see the GenomicFeatures package to learn more about `TxDb` objects). We will now show several ways of constructing such objects.

#### Building a GRangesList from Ensembl

We construct a gene annotation object for the human genome using the Ensembl release 104 (hg38). This can be done using the function `getTxDb` in `crisprDesign`:

```
txdb <- getTxDb(organism="Homo sapiens", release=104)
```

This may take several minutes, and note that this requires an internet connection. In case it times out, one can increase the timeout option using the following:

```
options(timeout = max(10000000, getOption("timeout")))
```

Once obtained, we can convert the object into a `GRangesList` using the function `TxDb2GRangesList` from `crisprDesign`:

```r
grList <- TxDb2GRangesList(txdb)
```

We will specify that the genome is hg38:

```r
GenomeInfoDb::genome(grList) <- "hg38"
```

And that's it! The `grList` object contains all of the information about the Ensembl release 104 gene model, and is ready to be used in the crisprVerse. Let's take a quick look at our gene annotation object:

```r
names(grList)
```

```
## [1] "transcripts" "exons"       "cds"         "fiveUTRs"    "threeUTRs"  
## [6] "introns"     "tss"
```

```r
grList$transcripts
```

```
## GRanges object with 111751 ranges and 14 metadata columns:
##     seqnames      ranges strand |           tx_id         gene_id
##        <Rle>   <IRanges>  <Rle> |     <character>     <character>
##            1 11869-14409      + | ENST00000456328 ENSG00000223972
##            1 12010-13670      + | ENST00000450305 ENSG00000223972
##            1 29554-31097      + | ENST00000473358 ENSG00000243485
##            1 30267-31109      + | ENST00000469289 ENSG00000243485
##            1 30366-30503      + | ENST00000607096 ENSG00000284332
##   .      ...         ...    ... .             ...             ...
##           MT   5826-5891      - | ENST00000387409 ENSG00000210144
##           MT   7446-7514      - | ENST00000387416 ENSG00000210151
##           MT 14149-14673      - | ENST00000361681 ENSG00000198695
##           MT 14674-14742      - | ENST00000387459 ENSG00000210194
##           MT 15956-16023      - | ENST00000387461 ENSG00000210196
##          protein_id                tx_type gene_symbol     exon_id exon_rank
##         <character>            <character> <character> <character> <integer>
##                <NA>   processed_transcript     DDX11L1        <NA>      <NA>
##                <NA> transcribed_unproces..     DDX11L1        <NA>      <NA>
##                <NA>                 lncRNA MIR1302-2HG        <NA>      <NA>
##                <NA>                 lncRNA MIR1302-2HG        <NA>      <NA>
##                <NA>                  miRNA   MIR1302-2        <NA>      <NA>
##   .             ...                    ...         ...         ...       ...
##                <NA>                Mt_tRNA       MT-TY        <NA>      <NA>
##                <NA>                Mt_tRNA      MT-TS1        <NA>      <NA>
##     ENSP00000354665         protein_coding      MT-ND6        <NA>      <NA>
##                <NA>                Mt_tRNA       MT-TE        <NA>      <NA>
##                <NA>                Mt_tRNA       MT-TP        <NA>      <NA>
##     cds_start   cds_end  tx_start    tx_end   cds_len exon_start  exon_end
##     <integer> <integer> <integer> <integer> <integer>  <integer> <integer>
##          <NA>      <NA>      <NA>      <NA>      <NA>       <NA>      <NA>
##          <NA>      <NA>      <NA>      <NA>      <NA>       <NA>      <NA>
##          <NA>      <NA>      <NA>      <NA>      <NA>       <NA>      <NA>
##          <NA>      <NA>      <NA>      <NA>      <NA>       <NA>      <NA>
##          <NA>      <NA>      <NA>      <NA>      <NA>       <NA>      <NA>
##   .       ...       ...       ...       ...       ...        ...       ...
##          <NA>      <NA>      <NA>      <NA>      <NA>       <NA>      <NA>
##          <NA>      <NA>      <NA>      <NA>      <NA>       <NA>      <NA>
##          <NA>      <NA>      <NA>      <NA>      <NA>       <NA>      <NA>
##          <NA>      <NA>      <NA>      <NA>      <NA>       <NA>      <NA>
##          <NA>      <NA>      <NA>      <NA>      <NA>       <NA>      <NA>
```

```
##   -------
##   seqinfo: 25 sequences (1 circular) from hg38 genome
```

##### Building a tssObject

Building a TSS annotation object requires only one additional step after constructing the `GRangesList` object described above. This can be obtained using the function `getTssObjectFromTxObject` in `crisprDesign`:

```
tssObject <- getTssObjectFromTxObject(grList)
tssObject
```

```
## GRanges object with 52547 ranges and 5 metadata columns:
##          seqnames    ranges strand |           tx_id         gene_id
##             <Rle> <IRanges>  <Rle> |     <character>     <character>
##    11402        1     65419      + | ENST00000641515 ENSG00000186092
##    11442        1    923923      + | ENST00000616016 ENSG00000187634
##    11444        1    925731      + | ENST00000342066 ENSG00000187634
##    11445        1    960584      + | ENST00000338591 ENSG00000187961
##    11446        1    960639      + | ENST00000622660 ENSG00000187961
##      ...      ...       ...    ... .             ...             ...
##   123058        Y  24047689      - | ENST00000382407 ENSG00000172352
##   123073        Y  24813186      - | ENST00000382365 ENSG00000187191
##   123074        Y  24813186      - | ENST00000315357 ENSG00000187191
##   123075        Y  24813186      - | ENST00000446723 ENSG00000187191
##   123080        Y  25052074      - | ENST00000382287 ENSG00000185894
##          gene_symbol        promoter                     ID
##          <character>     <character>            <character>
##    11402       OR4F5 ENST00000641515 ENSG00000186092_ENST..
##    11442      SAMD11 ENST00000616016 ENSG00000187634_ENST..
##    11444      SAMD11 ENST00000342066 ENSG00000187634_ENST..
##    11445      KLHL17 ENST00000338591 ENSG00000187961_ENST..
##    11446        <NA> ENST00000622660 ENSG00000187961_ENST..
##      ...         ...             ...                    ...
##   123058       CDY1B ENST00000382407 ENSG00000172352_ENST..
##   123073        DAZ3 ENST00000382365 ENSG00000187191_ENST..
##   123074        DAZ3 ENST00000315357 ENSG00000187191_ENST..
##   123075        DAZ3 ENST00000446723 ENSG00000187191_ENST..
##   123080       BPY2C ENST00000382287 ENSG00000185894_ENST..
##   -------
##   seqinfo: 25 sequences (1 circular) from hg38 genome
```

#### Using gene annotation objects

The gene (or TSS) annotation objects described above are often necessary for the full characterization of CRISPR gRNAs as they as inputs for several of the `crisprDesign` functions, including `queryTxObject`, `queryTssObject`, `addGeneAnnotation`, `addTssAnnotation`, and `addSpacerAlignments`.

For convenience, we provide in the crisprDesignData package precomputed gene annotation for human and mouse:

| Object name | Object class | Version | Description |
|---|---|---|---|
| `txdb_human` | `GRangesList` | Release 104 | Ensembl gene model for human (hg38/GRCh38) |
| `txdb_mouse` | `GRangesList` | Release 102 | Ensembl gene model for mouse (mm10/GRCm38) |

| Object name | Object class | Version | Description |
|---|---|---|---|
| `tss_human` | `GRanges` | Release 104 | Ensembl-based TSS coordinates for human (hg38/GRCh38) |
| `tss_mouse` | `GRanges` | Release 102 | Ensembl-based TSS coordinates for human (mm10/GRCm38) |

### Building a gene annotation object from a GFF file

If you have a General Feature Format (GFF) file from which you want to construct the gene annotation object, you can pass this to the `file` argument of the `crisprDesign` function `getTxDb`; this will create the `TxDb` object using the `GenomicFeatures` function `makeTxDbFromGFF`.
