## Supplementary material for "The crisprVerse: a comprehensive Bioconductor ecosystem for the design of CRISPR guide RNAs across nucleases and technologies": Tutorial17_Building_gRNA_Database

### Building a genome-wide gRNA database

#### Introduction

In this tutorial, we provide reproducible code to design and annotate gRNAs against all human protein-coding genes using the nuclease SpCas9.

#### Loading necessary packages

We first load the necessary packages:

```
library(crisprBase)
library(crisprScore)
library(crisprDesign)
library(crisprDesignData)
library(BSgenome.Hsapiens.UCSC.hg38)
```

##### Specifying the genome

We specify a `BSGenome` object that contains the DNA sequence of the human genome in hg38 coordinates:

```
bsgenome <- BSgenome.Hsapiens.UCSC.hg38
```

##### Specifying the genome index

We specify the file path of the Bowtie index that we will need for off-target alignment:

```
bowtie_index <- "/Users/fortinj2/crisprIndices/bowtie/hg38/hg38"
```

For instructions on how to build a Bowtie index from a given reference genome, see the genome index tutorial.

##### Specifying a SNP VCF file

To flag gRNAs overlapping common SNPs, we specify a VCF file obtained from the dbSNP website containing common SNPs from the dbSNP151 release:

```
vcf <- "/Users/fortinj2/crisprIndices/snps/dbsnp151.grch38/00-common_all.vcf.gz"
```

The VCF file was obtained from NCBI.

##### Specifying the nuclease

We load a `CrisprNuclease` object representing the SpCas9 nuclease from the `crisprBase` package:

```
data(SpCas9, package="crisprBase")
crisprNuclease <- SpCas9
```

To learn how to specify or build a custom nuclease, see the nuclease tutorial.

##### Specifying on-target scoring methods

We specify which on-target scoring methods should be used to score the gRNAs:

```
scoring_methods <- c("deephf", "deepspcas9")
```

One can see which scoring methods are available for a given nuclease using the following command:

```
crisprScore::scoringMethodsInfo
```

```
##        method   nuclease left right       type      label len
## 1    ruleset1      SpCas9  -24     5  On-target   RuleSet1  30
## 2     azimuth      SpCas9  -24     5  On-target    Azimuth  30
## 3      deephf      SpCas9  -20     2  On-target     DeepHF  23
## 4      lindel      SpCas9  -33    31  On-target     Lindel  65
## 5         mit      SpCas9  -20     2 Off-target        MIT  23
## 6         cfd      SpCas9  -20     2 Off-target        CFD  23
## 7    deepcpf1    AsCas12a   -4    29  On-target   DeepCpf1  34
## 8     enpamgb  enAsCas12a   -4    29  On-target    EnPAMGB  34
## 9  crisprscan      SpCas9  -26     8  On-target CRISPRscan  35
## 10    casrxrf       CasRx   NA    NA  On-target    CasRx-RF  NA
## 11   crisprai      SpCas9  -19     2  On-target   CRISPRai  22
## 12 crisprater      SpCas9  -20    -1  On-target CRISPRater  20
## 13 deepspcas9      SpCas9  -24     5  On-target DeepSpCas9  30
## 14   ruleset3      SpCas9  -24     5  On-target   RuleSet3  30
```

##### Specifying gene models and TSS annotations

To annotate gRNAs with a gene and TSS annotation, we need to specify a gene model formatted as a `GRangesList` object, as well as a TSS annotation with a `GRanges` object. The `crisprDesignData` contains such objects for both the human and mouse genomes, in GRCh38 (hg38) and GRCm38 (mm10) coordinates, respectively. Ensembl gene models were used to generate such objects. We load those objects:

```
data(txdb_human, package="crisprDesignData")
data(tss_human, package="crisprDesignData")
txObject <- txdb_human
tssObject <- tss_human
```

See the gene annotation tutorial to learn how to build such objects. The crisprDesignData also has tons of useful information.

##### Specifying repeat elements

To avoid designing gRNAs targeting repeat elements, we will specify a `GRanges` object containing repeats coordinates for the human genome. Here, we use the object `gr.repeats.hg38` in `crisprDesignData`. It contains genomic coordinates of the RepeatMasker UCSC track, for the hg38 reference genome:

```
data(gr.repeats.hg38, package="crisprDesignData")
grRepeats <- gr.repeats.hg38
```

#### Building a complete annotation for a given gene

The `designCompleteAnnotation` function in `crisprDesign` provides a one-step workflow to design and annotate all gRNAs targeting a given gene. The function was designed to be as comprehensive as possible to design and annotate gRNAs in one step. It does the following:

- Extract the DNA/RNA sequences with `queryTss`/`queryTxDB`

- Design gRNAs with `findSpacers`
- Remove gRNAs targeting repeat elements with `removeRepeats`
- Characterize spacer sequences with `addSequenceFeatures`
- Find on- and off-targets with `addSpacerAlignmentsIterative`
- Add gene annotation with `addGeneAnnotation`
- Add TSS annotation with `addTssAnnotation`
- Add on-target efficiency scores with `addOnTargetScores`
- Add off-target specificity scores with `addOffTargetScores`
- Add SNP annotation with `addSNPAnnotation`
- Add restriction enzymes information with `addRestrictionEnzymes`

Here, we design all CRISPRko gRNAs targeting the human KRAS gene (ENSG00000133703):

```
gs <- designCompleteAnnotation(queryValue="ENSG00000133703",
                               queryColumn="gene_id",
                               modality="CRISPRko",
                               bsgenome=bsgenome,
                               bowtie_index=bowtie_index,
                               crisprNuclease=SpCas9,
                               txObject=txObject,
                               tssObject=tssObject,
                               grRepeats=grRepeats,
                               vcf=vcf,
                               n_mismatches=1,
                               scoring_methods=scoring_methods)
```

```
## [designCompleteAnnotation] Adding sequence statistics
## [designCompleteAnnotation] Adding spacer alignments

## Loading required namespace: crisprBwa

## [runCrisprBowtie] Using BSgenome.Hsapiens.UCSC.hg38
## [runCrisprBowtie] Searching for SpCas9 protospacers
## [runCrisprBowtie] Using BSgenome.Hsapiens.UCSC.hg38
## [runCrisprBowtie] Searching for SpCas9 protospacers
## [designCompleteAnnotation] Adding gene annotation
## [designCompleteAnnotation] Adding on-target scores

## [addOnTargetScores] Adding deephf scores.

## snapshotDate(): 2022-08-23

## see ?crisprScoreData and browseVignettes('crisprScoreData') for documentation

## loading from cache

## [addOnTargetScores] Adding deepspcas9 scores.

## [designCompleteAnnotation] Adding CFD scores annotation
## [designCompleteAnnotation] Adding SNP annotation
## [designCompleteAnnotation] Adding composite scores
```

The resulting object is a `GuideSet` object. To learn more about what are `GuideSet` objects, and how to interact with them, see the CRISPRko gRNA design tutorial.

```
gs
```

```
## GuideSet object with 56 ranges and 28 metadata columns:
##                      seqnames    ranges strand |          protospacer
##                         <Rle> <IRanges>  <Rle> |       <DNAStringSet>
##    ENSG00000133703_1    chr12  25209843      - | AAAGAAAAGATGAGCAAAGA
```

```
##    ENSG00000133703_2    chr12  25209896      + | TTCTCGAACTAATGTATAGA
##    ENSG00000133703_3    chr12  25215438      - | AAATGCATTATAATGTAATC
##    ENSG00000133703_4    chr12  25215477      - | AGCAAAGAAGAAAAGACTCC
##    ENSG00000133703_5    chr12  25215477      + | TTTTTAATTTTCACACAGCC
##                  ...      ...       ...    ... .                  ...
##   ENSG00000133703_52    chr12  25245349      - | CTTGTGGTAGTTGGAGCTGG
##   ENSG00000133703_53    chr12  25245352      - | AAACTTGTGGTAGTTGGAGC
##   ENSG00000133703_54    chr12  25245358      - | GAATATAAACTTGTGGTAGT
##   ENSG00000133703_55    chr12  25245365      - | AATGACTGAATATAAACTTG
##   ENSG00000133703_56    chr12  25245392      + | TATATTCAGTCATTTTCAGC
##                                  pam  pam_site  cut_site      region inRepeats
##                       <DNAStringSet> <numeric> <numeric> <character> <logical>
##    ENSG00000133703_1             TGG  25209843  25209846    region_8     FALSE
##    ENSG00000133703_2             AGG  25209896  25209893    region_8     FALSE
##    ENSG00000133703_3             TGG  25215438  25215441    region_4     FALSE
##    ENSG00000133703_4             TGG  25215477  25215480    region_4     FALSE
##    ENSG00000133703_5             AGG  25215477  25215474    region_4     FALSE
##                  ...             ...       ...       ...         ...       ...
##   ENSG00000133703_52             TGG  25245349  25245352    region_1     FALSE
##   ENSG00000133703_53             TGG  25245352  25245355    region_1     FALSE
##   ENSG00000133703_54             TGG  25245358  25245361    region_1     FALSE
##   ENSG00000133703_55             TGG  25245365  25245368    region_1     FALSE
##   ENSG00000133703_56             AGG  25245392  25245389    region_1     FALSE
##                        percentGC     polyA     polyC     polyG     polyT
##                        <numeric> <logical> <logical> <logical> <logical>
##    ENSG00000133703_1          30      TRUE     FALSE     FALSE     FALSE
##    ENSG00000133703_2          30     FALSE     FALSE     FALSE     FALSE
##    ENSG00000133703_3          20     FALSE     FALSE     FALSE     FALSE
##    ENSG00000133703_4          40      TRUE     FALSE     FALSE     FALSE
##    ENSG00000133703_5          30     FALSE     FALSE     FALSE      TRUE
##                  ...         ...       ...       ...       ...       ...
##   ENSG00000133703_52          55     FALSE     FALSE     FALSE     FALSE
##   ENSG00000133703_53          45     FALSE     FALSE     FALSE     FALSE
##   ENSG00000133703_54          30     FALSE     FALSE     FALSE     FALSE
##   ENSG00000133703_55          25     FALSE     FALSE     FALSE     FALSE
##   ENSG00000133703_56          30     FALSE     FALSE     FALSE      TRUE
##                        startingGGGGG        n0      n0_c      n0_p        n1
##                            <logical> <numeric> <numeric> <numeric> <numeric>
##    ENSG00000133703_1           FALSE         1         1         0         4
##    ENSG00000133703_2           FALSE         1         1         0         1
##    ENSG00000133703_3           FALSE         1         1         0         0
##    ENSG00000133703_4           FALSE         1         1         0         0
##    ENSG00000133703_5           FALSE         1         1         0         0
##                  ...             ...       ...       ...       ...       ...
##   ENSG00000133703_52           FALSE         1         1         0         1
##   ENSG00000133703_53           FALSE         1         1         0         1
##   ENSG00000133703_54           FALSE         1         1         0         1
##   ENSG00000133703_55           FALSE         2         1         0         2
##   ENSG00000133703_56           FALSE         2         0         0         1
##                             n1_c      n1_p
##                        <numeric> <numeric>
##    ENSG00000133703_1           0         0
##    ENSG00000133703_2           0         0
##    ENSG00000133703_3           0         0
```

```
##    ENSG00000133703_4         0         0
##    ENSG00000133703_5         0         0
##                  ...       ...       ...
##   ENSG00000133703_52         0         0
##   ENSG00000133703_53         0         0
##   ENSG00000133703_54         0         0
##   ENSG00000133703_55         0         0
##   ENSG00000133703_56         0         0
##                                                                alignments
##                                                             <GRangesList>
##    ENSG00000133703_1  chr12:25209843:-,chr8:68551391:-,chr6:54771089:+,...
##    ENSG00000133703_2                     chr12:25209896:+,chr6:54771050:-
##    ENSG00000133703_3                                     chr12:25215438:-
##    ENSG00000133703_4                                     chr12:25215477:-
##    ENSG00000133703_5                                     chr12:25215477:+
##                  ...                                                   ...
##   ENSG00000133703_52                     chr12:25245349:-,chr6:54770618:+
##   ENSG00000133703_53                     chr12:25245352:-,chr6:54770615:+
##   ENSG00000133703_54                     chr12:25245358:-,chr6:54770609:+
##   ENSG00000133703_55 chr12:25245365:-,chr6:54770602:+,chr13:60822020:-,...
##   ENSG00000133703_56     chr12:25245392:+,chr6:54770575:-,chr1:210618123:-
##                                                                         geneAnnotation
##                                                                  <SplitDataFrameList>
##    ENSG00000133703_1 chr12:25209846:-:...,chr12:25209846:-:...,chr12:25209846:-:...,...
##    ENSG00000133703_2 chr12:25209893:+:...,chr12:25209893:+:...,chr12:25209893:+:...,...
##    ENSG00000133703_3                                                  chr12:25215441:-:...
##    ENSG00000133703_4                                                  chr12:25215480:-:...
##    ENSG00000133703_5                                                  chr12:25215474:+:...
##                  ...                                                                   ...
##   ENSG00000133703_52 chr12:25245352:-:...,chr12:25245352:-:...,chr12:25245352:-:...,...
##   ENSG00000133703_53 chr12:25245355:-:...,chr12:25245355:-:...,chr12:25245355:-:...,...
##   ENSG00000133703_54 chr12:25245361:-:...,chr12:25245361:-:...,chr12:25245361:-:...,...
##   ENSG00000133703_55 chr12:25245368:-:...,chr12:25245368:-:...,chr12:25245368:-:...,...
##   ENSG00000133703_56 chr12:25245389:+:...,chr12:25245389:+:...,chr12:25245389:+:...,...
##                          enzymeAnnotation score_deephf score_deepspcas9
##                      <SplitDataFrameList>    <numeric>        <numeric>
##    ENSG00000133703_1 FALSE:FALSE:FALSE:...     0.450868      0.427276688
##    ENSG00000133703_2 FALSE:FALSE:FALSE:...     0.428607      0.204131565
##    ENSG00000133703_3 FALSE:FALSE:FALSE:...     0.292229      0.029736991
##    ENSG00000133703_4 FALSE:FALSE:FALSE:...     0.612286      0.477413216
##    ENSG00000133703_5 FALSE:FALSE:FALSE:...     0.183310      0.000671324
##                  ...                   ...          ...              ...
##   ENSG00000133703_52 FALSE:FALSE:FALSE:...     0.644286        0.5256023
##   ENSG00000133703_53 FALSE:FALSE:FALSE:...     0.439317        0.3657698
##   ENSG00000133703_54 FALSE:FALSE:FALSE:...     0.433265        0.2556772
##   ENSG00000133703_55 FALSE:FALSE:FALSE:...     0.671397        0.6270906
##   ENSG00000133703_56 FALSE:FALSE:FALSE:...     0.320574        0.0444068
##                      score_cfd score_mit    hasSNP                   snps
##                      <numeric> <numeric> <logical>   <SplitDataFrameList>
##    ENSG00000133703_1  0.425027  0.426600      TRUE rs1137282:25209843:0:...
##    ENSG00000133703_2  0.500000  0.577367     FALSE                 :...,...
##    ENSG00000133703_3  1.000000  1.000000     FALSE                 :...,...
##    ENSG00000133703_4  1.000000  1.000000     FALSE                 :...,...
##    ENSG00000133703_5  1.000000  1.000000     FALSE                 :...,...
```

```
##                 ...       ...       ...       ...                  ...
##   ENSG00000133703_52  0.500000  0.547046     FALSE                :...,...
##   ENSG00000133703_53  0.500000  0.619963     FALSE                :...,...
##   ENSG00000133703_54  0.777778  0.759301     FALSE                :...,...
##   ENSG00000133703_55  0.458599  0.489579     FALSE                :...,...
##   ENSG00000133703_56  0.442623  0.464868     FALSE                :...,...
##                     score_composite
##                           <numeric>
##    ENSG00000133703_1           25.5
##    ENSG00000133703_2           16.0
##    ENSG00000133703_3            7.0
##    ENSG00000133703_4           37.0
##    ENSG00000133703_5            3.0
##                 ...             ...
##   ENSG00000133703_52           43.0
##   ENSG00000133703_53           19.5
##   ENSG00000133703_54           17.5
##   ENSG00000133703_55           51.5
##   ENSG00000133703_56            8.0
##   -------
##   seqinfo: 640 sequences (1 circular) from hg38 genome
##   crisprNuclease: SpCas9
```

###### Converting the `GuideSet` object to a list of data.frames

The `flattenGuideSet` function in `crisprDesign` is a convenience function to convert a `GuideSet` object into a set of `data.frames` that can be saved as plain text files:

```
dfs <- flattenGuideSet(gs)
```

We can look at the names of the data.frames:

```
names(dfs)
```

```
## [1] "primary"          "alignments"       "geneAnnotation"   "enzymeAnnotation"
## [5] "snps"
```

As an example, let's look at the first rows of the primary data.frame:

```
head(dfs$primary)
```

```
##                   ID               spacer          protospacer   chr    start
## 1 ENSG00000133703_1 AAAGAAAAGATGAGCAAAGA AAAGAAAAGATGAGCAAAGA chr12 25209844
## 2 ENSG00000133703_2 TTCTCGAACTAATGTATAGA TTCTCGAACTAATGTATAGA chr12 25209876
## 3 ENSG00000133703_3 AAATGCATTATAATGTAATC AAATGCATTATAATGTAATC chr12 25215439
## 4 ENSG00000133703_4 AGCAAAGAAGAAAAGACTCC AGCAAAGAAGAAAAGACTCC chr12 25215478
## 5 ENSG00000133703_5 TTTTTAATTTTCACACAGCC TTTTTAATTTTCACACAGCC chr12 25215457
## 6 ENSG00000133703_6 TTTTTTTCAATCTGTATTGT TTTTTTTCAATCTGTATTGT chr12 25215500
##        end strand pam pam_site cut_site   region inRepeats percentGC polyA
## 1 25209863      - TGG 25209843 25209846 region_8     FALSE        30  TRUE
## 2 25209895      + AGG 25209896 25209893 region_8     FALSE        30 FALSE
## 3 25215458      - TGG 25215438 25215441 region_4     FALSE        20 FALSE
## 4 25215497      - TGG 25215477 25215480 region_4     FALSE        40  TRUE
## 5 25215476      + AGG 25215477 25215474 region_4     FALSE        30 FALSE
## 6 25215519      + CGG 25215520 25215517 region_4     FALSE        20 FALSE
##   polyC polyG polyT startingGGGGG n0 n0_c n0_p n1 n1_c n1_p score_deephf
## 1 FALSE FALSE FALSE         FALSE  1    1    0  4    0    0    0.4508680
```

```
## 2 FALSE FALSE FALSE         FALSE  1    1    0  1    0    0    0.4286066
## 3 FALSE FALSE FALSE         FALSE  1    1    0  0    0    0    0.2922295
## 4 FALSE FALSE FALSE         FALSE  1    1    0  0    0    0    0.6122858
## 5 FALSE FALSE  TRUE         FALSE  1    1    0  0    0    0    0.1833103
## 6 FALSE FALSE  TRUE         FALSE  1    1    0  4    0    0    0.1669266
##   score_deepspcas9 score_cfd score_mit hasSNP score_composite
## 1     0.4272766876 0.4250273 0.4266001   TRUE            25.5
## 2     0.2041315651 0.5000000 0.5773672  FALSE            16.0
## 3     0.0297369909 1.0000000 1.0000000  FALSE             7.0
## 4     0.4774132156 1.0000000 1.0000000  FALSE            37.0
## 5     0.0006713235 1.0000000 1.0000000  FALSE             3.0
## 6     0.0166297376 0.5212645 0.8835838  FALSE             3.5
```

#### Building a complete gRNA database across all protein-coding genes

We first get all possibles genes from our gene model:

```
gene_ids <- unique(txObject$cds$gene_id)
head(gene_ids)
```

```
## [1] "ENSG00000186092" "ENSG00000187634" "ENSG00000187961" "ENSG00000187583"
## [5] "ENSG00000187608" "ENSG00000188157"
```

and specify where to save the `GuideSet` objects:

```
dir <- "./crisprko_cas9_hg38"
if (!dir.exists(dir)){
    dir.create(dir, recursive=TRUE)
}
```

We are now looping over all genes to generate the data:

```
lapply(gene_index, function(gene){
    gs <- designCompleteAnnotation(queryValue=gene,
                                   queryColumn="gene_id",
                                   modality="CRISPRko",
                                   bsgenome=bsgenome,
                                   bowtie_index=bowtie_index,
                                   crisprNuclease=SpCas9,
                                   txObject=txObject,
                                   tssObject=tssObject,
                                   grRepeats=grRepeats,
                                   vcf=vcf,
                                   n_mismatches=3,
                                   scoring_methods=scoring_methods)
    write.rds(gs, file=file.path(dir, paste0(gene, ".rds")))
})
```

This loop can be modified by the user to use an embarrassingly-parallel approach, using the BiocParallel package, for instance.

Building a database for CRISPRa and CRISPRi applications works similarly See `?designCompleteAnnotation` for more information.

### Reproducibility

```
sessionInfo()
```

```
## R version 4.2.1 (2022-06-23)
## Platform: x86_64-apple-darwin17.0 (64-bit)
## Running under: macOS Catalina 10.15.7
##
## Matrix products: default
## BLAS:   /Library/Frameworks/R.framework/Versions/4.2/Resources/lib/libRblas.0.dylib
## LAPACK: /Library/Frameworks/R.framework/Versions/4.2/Resources/lib/libRlapack.dylib
##
## locale:
## [1] en_US.UTF-8/en_US.UTF-8/en_US.UTF-8/C/en_US.UTF-8/en_US.UTF-8
##
## attached base packages:
## [1] stats4    stats     graphics  grDevices utils     datasets  methods
## [8] base
##
## other attached packages:
##  [1] BSgenome.Hsapiens.UCSC.hg38_1.4.4 BSgenome_1.65.2
##  [3] rtracklayer_1.57.0                Biostrings_2.65.2
##  [5] XVector_0.37.0                    GenomicRanges_1.49.1
##  [7] GenomeInfoDb_1.33.5               IRanges_2.31.2
##  [9] S4Vectors_0.35.1                  crisprDesignData_0.99.17
## [11] crisprDesign_0.99.133             crisprScore_1.1.14
## [13] crisprScoreData_1.1.3             ExperimentHub_2.5.0
## [15] AnnotationHub_3.5.0               BiocFileCache_2.5.0
## [17] dbplyr_2.2.1                      BiocGenerics_0.43.1
## [19] crisprBowtie_1.1.1                crisprBase_1.1.5
## [21] crisprVerse_0.99.8                rmarkdown_2.15.2
##
## loaded via a namespace (and not attached):
##  [1] rjson_0.2.21                  ellipsis_0.3.2
##  [3] Rbowtie_1.37.0                bit64_4.0.5
##  [5] interactiveDisplayBase_1.35.0 AnnotationDbi_1.59.1
##  [7] fansi_1.0.3                   xml2_1.3.3
##  [9] codetools_0.2-18              cachem_1.0.6
## [11] knitr_1.40                    jsonlite_1.8.0
## [13] Rsamtools_2.13.4              png_0.1-7
## [15] shiny_1.7.2                   BiocManager_1.30.18
## [17] readr_2.1.2                   compiler_4.2.1
## [19] httr_1.4.4                    basilisk_1.9.2
## [21] assertthat_0.2.1              Matrix_1.4-1
## [23] fastmap_1.1.0                 cli_3.3.0
## [25] later_1.3.0                   htmltools_0.5.3
## [27] prettyunits_1.1.1             tools_4.2.1
## [29] glue_1.6.2                    GenomeInfoDbData_1.2.8
## [31] dplyr_1.0.9                   rappdirs_0.3.3
## [33] tinytex_0.41                  Rcpp_1.0.9
## [35] Biobase_2.57.1                vctrs_0.4.1
## [37] crisprBwa_1.1.3               xfun_0.32
## [39] stringr_1.4.1                 mime_0.12
## [41] lifecycle_1.0.1               restfulr_0.0.15
```

```
## [43] XML_3.99-0.10                zlibbioc_1.43.0
## [45] basilisk.utils_1.9.1         vroom_1.5.7
## [47] VariantAnnotation_1.43.3     hms_1.1.2
## [49] promises_1.2.0.1             MatrixGenerics_1.9.1
## [51] parallel_4.2.1               SummarizedExperiment_1.27.1
## [53] yaml_2.3.5                   curl_4.3.2
## [55] memoise_2.0.1                reticulate_1.25
## [57] biomaRt_2.53.2               stringi_1.7.8
## [59] RSQLite_2.2.16               BiocVersion_3.16.0
## [61] highr_0.9                    BiocIO_1.7.1
## [63] randomForest_4.7-1.1         GenomicFeatures_1.49.6
## [65] filelock_1.0.2               BiocParallel_1.31.12
## [67] rlang_1.0.4                  pkgconfig_2.0.3
## [69] matrixStats_0.62.0           bitops_1.0-7
## [71] evaluate_0.16                lattice_0.20-45
## [73] purrr_0.3.4                  GenomicAlignments_1.33.1
## [75] bit_4.0.4                    tidyselect_1.1.2
## [77] magrittr_2.0.3               R6_2.5.1
## [79] generics_0.1.3               DelayedArray_0.23.1
## [81] DBI_1.1.3                    pillar_1.8.1
## [83] KEGGREST_1.37.3              RCurl_1.98-1.8
## [85] tibble_3.1.8                 dir.expiry_1.5.0
## [87] crayon_1.5.1                 utf8_1.2.2
## [89] tzdb_0.3.0                   progress_1.2.2
## [91] grid_4.2.1                   blob_1.2.3
## [93] digest_0.6.29                xtable_1.8-4
## [95] httpuv_1.6.5                 Rbwa_1.1.0
```
