## Supplementary material for "The crisprVerse: a comprehensive Bioconductor ecosystem for the design of CRISPR guide RNAs across nucleases and technologies": Tutorial3_Specifying_CRISPR_Nuclease

### Building custom CRISPR nuclease objects

#### Introduction

The `crisprBase` package provides functionalities to represent and specify custom CRISPR nucleases to be used in the crisprVerse ecosystem, as well as other enzymes such as base editors and nickases. Commonly-used CRISPR nucleases, such as SpCas9, AsCas12a, enAsCas12a, and CasRx are readily available in the package, and users do not need to reconstruct those nucleases.

The package also provides arithmetic functions to extract genomic ranges to help with the design and manipulation of CRISPR guide RNAs (gRNAs). The classes and functions are designed to work with a broad spectrum of nucleases and applications, including PAM-free CRISPR nucleases, RNA-targeting nucleases, and the more general class of restriction enzymes. It also includes functionalities for CRISPR nickases.

In this tutorial we show how to construct objects for the following enzymes: `Nuclease`, `CrisprNuclease`, `BaseEditor`, `CrisprNickase`. We also discuss some important details of these objects, how to easily retrieve key information with accessor functions, and how to use arithmetic functions on these objects in facilitating gRNA design.

#### Installation

`crisprBase` is part of the crisprVerse, and can be installed by installing the `crisprVerse` package; see the Installation tutorial.

##### Getting started

`crisprBase` can be loaded by loading the `crisprVerse` package:

```
library(crisprVerse)
```

#### Nuclease class

The `Nuclease` class is designed to store minimal information about the recognition sites of general nucleases, such as restriction enzymes. The `Nuclease` class has 5 fields:

- `nucleaseName`: a string giving the name of the nuclease
- `targetType`: a string specifying whether the nuclease targets `"DNA"` (deoxyribonucleases) or `"RNA"` (ribonucleases)
- `metadata`: a list of arbitrary length that stores additional information about the nuclease
- `motifs`: a character vector that describes the sequence motifs that are recognized by the nuclease for cleavage (always given in the 5' to 3' direction)
- `weights`: an optional numeric vector that gives the relative cleavage probabilities corresponding to the motifs in `motifs`

Note that we use DNA to represent motifs irrespective of the target type for the sake of simplicity.

We use the Rebase convention in representing motif sequences (Roberts et al. 2010). For enzymes that cleave within the recognition site, we add the symbol `^` within the recognition sequence to specify the cleavage site, always in the 5' to 3' direction. For enzymes that cleave away from the recognition site, we specify the distance of the cleavage site using a `(x/y)` notation where `x` represents the number of nucleotides away from the recognition sequence on the original strand, and `y` represents the number of nucleotides away from the recognition sequence on the reverse strand.

#### Examples

The following examples use the `Nuclease` class constructor function from `crisprBase` to create `Nuclease` objects representing common restriction enzymes.

The EcoRI enzyme recognizes the palindromic motif `GAATTC`, and cuts after the first nucleotide. We can create this nuclease with the following function:

```
EcoRI <- Nuclease("EcoRI",
                  targetType="DNA",
                  motifs=c("G^AATTC"),
                  metadata=list(description="EcoRI restriction enzyme"))
EcoRI
```

```
## Class: Nuclease
##   Name: EcoRI
##   Target type: DNA
##   Metadata: list of length 1
##   Motifs: GAATTC
##   Weights: 1
```

The HgaI enzyme recognizes the motif `GACGC`, and cleaves the DNA at 5 nucleotides downstream of the recognition sequence on the original strand, and at 10 nucleotides downstream of the recognition sequence on the reverse strand:

```
HgaI <- Nuclease("HgaI",
                 targetType="DNA",
                 motifs=c("GACGC(5/10)"),
                 metadata=list(description="HgaI restriction enzyme"))
HgaI
```

```
## Class: Nuclease
##   Name: HgaI
##   Target type: DNA
##   Metadata: list of length 1
##   Motifs: GACGC
##   Weights: 1
```

If the cleavage site was upstream of the recognition sequence, we would instead specify `(5/10)GACGC`.

Any nucleotide letter that is part of the extended IUPAC nucleic acid code can be used to represent recognition motifs. For instance, we use `Y` and `R` (pyrimidine and purine, respectively) to specify the possible recognition sequences for PfaAI:

```
PfaAI <- Nuclease("PfaAI",
                  targetType="DNA",
                  motifs=c("G^GYRCC"),
                  metadata=list(description="PfaAI restriction enzyme"))
PfaAI
```

```
## Class: Nuclease
##   Name: PfaAI
##   Target type: DNA
##   Metadata: list of length 1
```

```
##   Motifs: GGYRCC
##   Weights: 1
```

##### Accessor functions

The accessor function `motifs` retrieve the motif sequences:

```
motifs(PfaAI)
```

```
## DNAStringSet object of length 1:
##     width seq
## [1]     6 GGYRCC
```

To expand the motif sequence into all possible combinations of valid sequences with only those nucleotides in the set {A, C, G, T}, pass the argument `expand=TRUE`.

```
motifs(PfaAI, expand=TRUE)
```

```
## DNAStringSet object of length 4:
##     width seq                                                    names
## [1]     6 GGCACC                                                 GGYRCC
## [2]     6 GGTACC                                                 GGYRCC
## [3]     6 GGCGCC                                                 GGYRCC
## [4]     6 GGTGCC                                                 GGYRCC
```

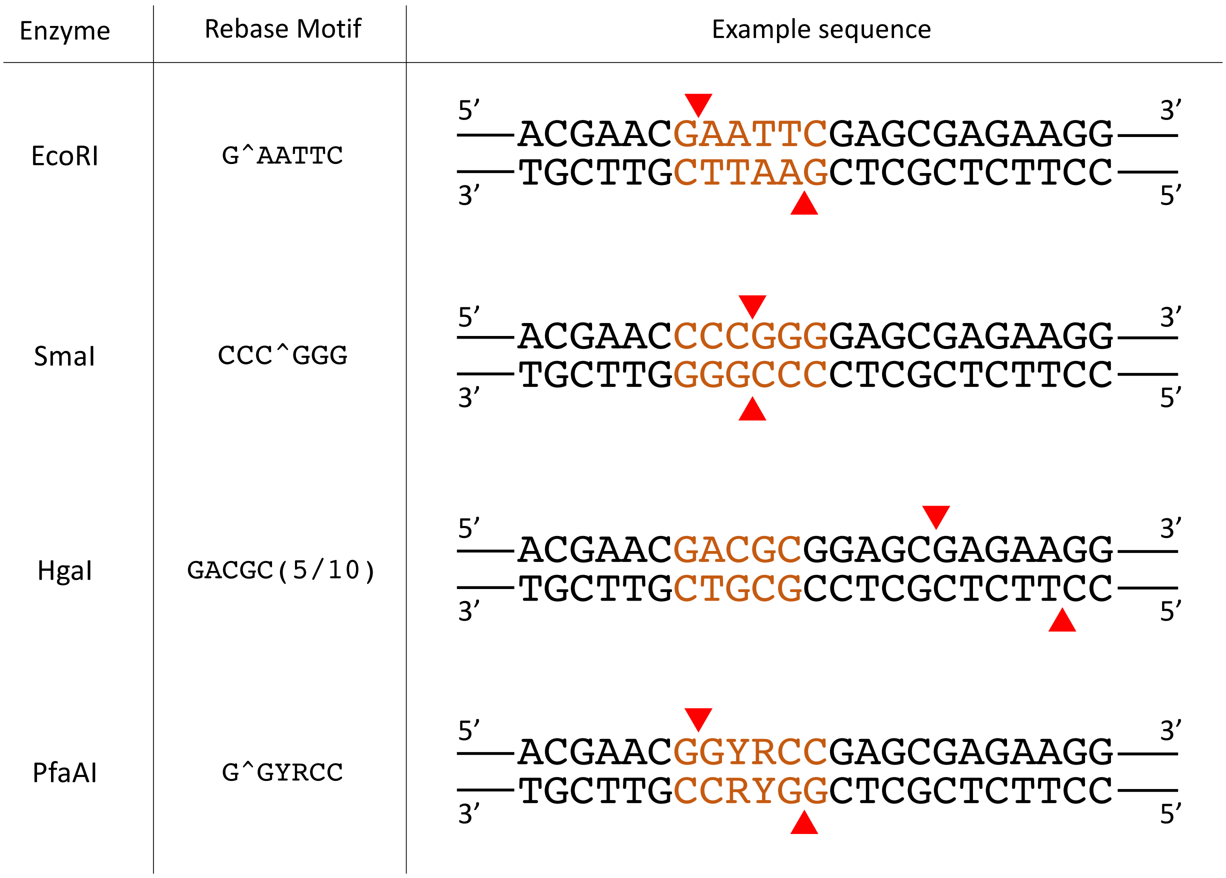


Figure 1: Examples of restriction enzymes

### CrisprNuclease class

CRISPR nucleases are examples of RNA-guided nucleases. They require two binding components for cleavage. For DNA-targeting CRISPR nucleases, the nuclease must first recognize a constant nucleotide motif in the target DNA called the protospacer adjacent motif (PAM) sequence. Second, the guide-RNA (gRNA), which guides the nuclease to the target sequence, needs to bind to a complementary sequence adjacent to the PAM sequence (protospacer sequence). The latter can be thought of a variable binding motif that can be specified by designing corresponding gRNA sequences. For CRISPR nucleases targeting RNA, the equivalent of the PAM sequence is called the Protospacer Flanking Sequence (PFS). We use the terms PAM and PFS interchangeably as it should be clear from context.

The `CrisprNuclease` class allows for characterization of both binding components by extending the `Nuclease` class to contain information about gRNA sequences. The PAM sequence characteristics, and the cleavage distance with respect to the PAM sequence, are specified using the motif nomenclature described in the Nuclease section above.

Three additional fields are required:

- `pam_side`: either `"5prime"` or `"3prime"`, which specifies on which side the PAM sequence is located with repsect to the protospacer sequence (while it would be more appropriate to use the term `pfs_side` for RNA-targeting nucleases, we still use `pam_side` for simplicity)
- `spacer_length`: a numeric value that specifies the default spacer length in nucleotides
- `spacer_gap`: a numeric value that gives the distance in nucleotides between the PAM (or PFS) sequence and spacer sequence (for most nucleases, `spacer_gap=0` as the spacer sequence is immediately adjacent to the PAM/PFS sequence)

#### Examples

These examples use the `CrisprNuclease` class constructor function from `crisprBase` to create `CrisprNuclease` objects that represent some of the more common CRISPR nucleases.

Here we construct a `CrisprNuclease` object for the Cas9 nuclease (Streptococcus pyogenes Cas9):

```
SpCas9 <- CrisprNuclease("SpCas9",
                         targetType="DNA",
                         pams=c("(3/3)NGG", "(3/3)NAG", "(3/3)NGA"),
                         weights=c(1, 0.2593, 0.0694),
                         metadata=list(description="Wildtype Streptococcus pyogenes Cas9 (SpCas9) nucle
                         pam_side="3prime",
                         spacer_length=20)
SpCas9
```

```
## Class: CrisprNuclease
##   Name: SpCas9
##   Target type: DNA
##   Metadata: list of length 1
##   PAMs: NGG, NAG, NGA
##   Weights: 1, 0.2593, 0.0694
##   Spacer length: 20
##   PAM side: 3prime
##     Distance from PAM: 0
##   Prototype protospacers: 5'--SSSSSSSSSSSSSSSSSSSS[NGG]--3', 5'--SSSSSSSSSSSSSSSSSSSS[NAG]--3', 5'--S
```

As with the `Nuclease` class, we can specify PAM sequences using the extended nucleotide code. SaCas9 serves as a good example:

```
SaCas9 <- CrisprNuclease("SaCas9",
                         targetType="DNA",
```

```
                          pams=c("(3/3)NNGRRT"),
                          metadata=list(description="Wildtype Staphylococcus
                          aureus Cas9 (SaCas9) nuclease"),
                          pam_side="3prime",
                          spacer_length=21)
SaCas9
```

```
## Class: CrisprNuclease
##   Name: SaCas9
##   Target type: DNA
##   Metadata: list of length 1
##   PAMs: NNGRRT
##   Weights: 1
##   Spacer length: 21
##   PAM side: 3prime
##     Distance from PAM: 0
##   Prototype protospacers: 5'--SSSSSSSSSSSSSSSSSSSSS[NNGRRT]--3'
```

Here is another example where we construct a `CrisprNuclease` object for the Cas12a nuclease (AsCas12a):

```
AsCas12a <- CrisprNuclease("AsCas12a",
                            targetType="DNA",
                            pams="TTTV(18/23)",
                            metadata=list(description="Wildtype Acidaminococcus
                            Cas12a (AsCas12a) nuclease."),
                            pam_side="5prime",
                            spacer_length=23)
AsCas12a
```

```
## Class: CrisprNuclease
##   Name: AsCas12a
##   Target type: DNA
##   Metadata: list of length 1
##   PAMs: TTTV
##   Weights: 1
##   Spacer length: 23
##   PAM side: 5prime
##     Distance from PAM: 0
##   Prototype protospacers: 5'--[TTTV]SSSSSSSSSSSSSSSSSSSSSSS--3'
```

##### CrisprNuclease objects provided in CrisprBase

Several already-constructed `crisprNuclease` objects for some of the most popular CRISPR nucleases are available in `crisprBase` for your convenience. The list of available nucleases can be accessed by typing the following:

```
data(package="crisprBase")
```

#### CRISPR arithmetics

##### CRISPR terminology

The terms **spacer** and **protospacer** are not interchangeable. **spacer** refers to the sequence used in the gRNA construct to guide the Cas nuclease to the target **protospacer** sequence in the host genome or transcriptome. The **protospacer** sequence is adjacent to the PAM sequence or PFS sequence. We use the terminology **target** sequence to refer to the protospacer and PAM sequence taken together. For DNA-targeting nucleases such as Cas9 and Cas12a, the sequences that make up the spacer and protospacer identical, while for RNA-targeting nucleases such as Cas13d, the spacer and protospacer sequences are reverse complements.

A given gRNA spacer sequence may not uniquely target the host genome–it can map to multiple protospacers in the genome. However, for a given reference genome we can uniquely identify protospacer sequences using a combination of 3 attributes:

- **chr**: chromosome name
- **strand**: forward (+) or reverse (-)
- **pam_site**: genomic coordinate of the first nucleotide of the nuclease-specific PAM sequence. For SpCas9, this corresponds to the genomic coordinate of N in the NGG PAM sequence. For AsCas12a, this corresponds to the genomic coordinate of the first T nucleotide in the TTTV PAM sequence. For RNA-targeting nucleases, this corresponds to the first nucleotide of the PFS (we do not use `pfs_site` for simplicity).

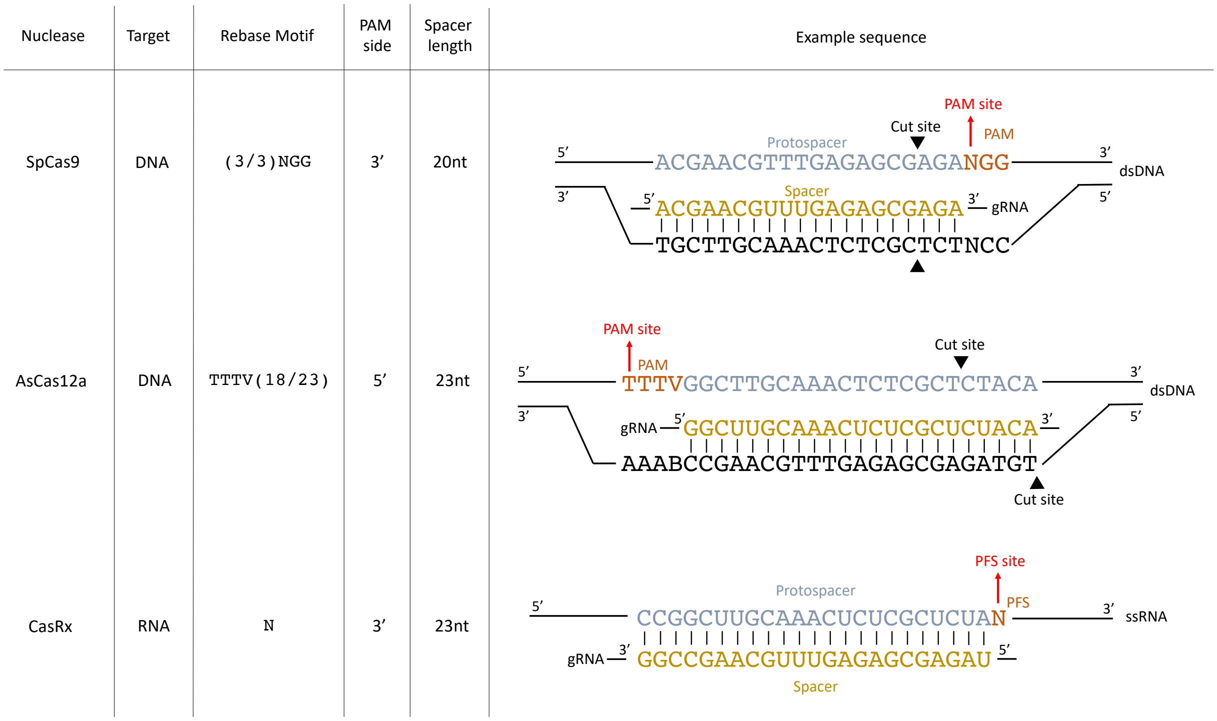


Figure 2: Examples of CRISPR nucleases

#### Cut site

By convention, we used the nucleotide directly downstream of the DNA cut to represent the cut site nucleotide position. For instance, for SpCas9 (blunt-ended dsDNA break), the cut site occurs at position `-3` with respect to the PAM site. For AsCas12a, the 5nt overhang dsDNA break occurs at 18 nucleotides after the PAM sequence on the targeted strand. With a PAM sequence of 4 nucleotides (TTTV), the cut site on the forward strand occurs at position `22` with respect to the PAM site, and at position `27` on the reverse strand.

The convenience function `cutSites` extracts the cut site coordinates relative to the PAM site:

```
data(SpCas9, package="crisprBase")
data(AsCas12a, package="crisprBase")
cutSites(SpCas9)
```

```
## [1] -3
```

```r
cutSites(SpCas9, strand="-")
```

```
## [1] -3
```

```r
cutSites(AsCas12a)
```

```
## [1] 22
```

```r
cutSites(AsCas12a, strand="-")
```

```
## [1] 27
```

Below is an illustration of how different motif sequences and cut patterns translate into cut site coordinates with respect to a PAM sequence NGG:

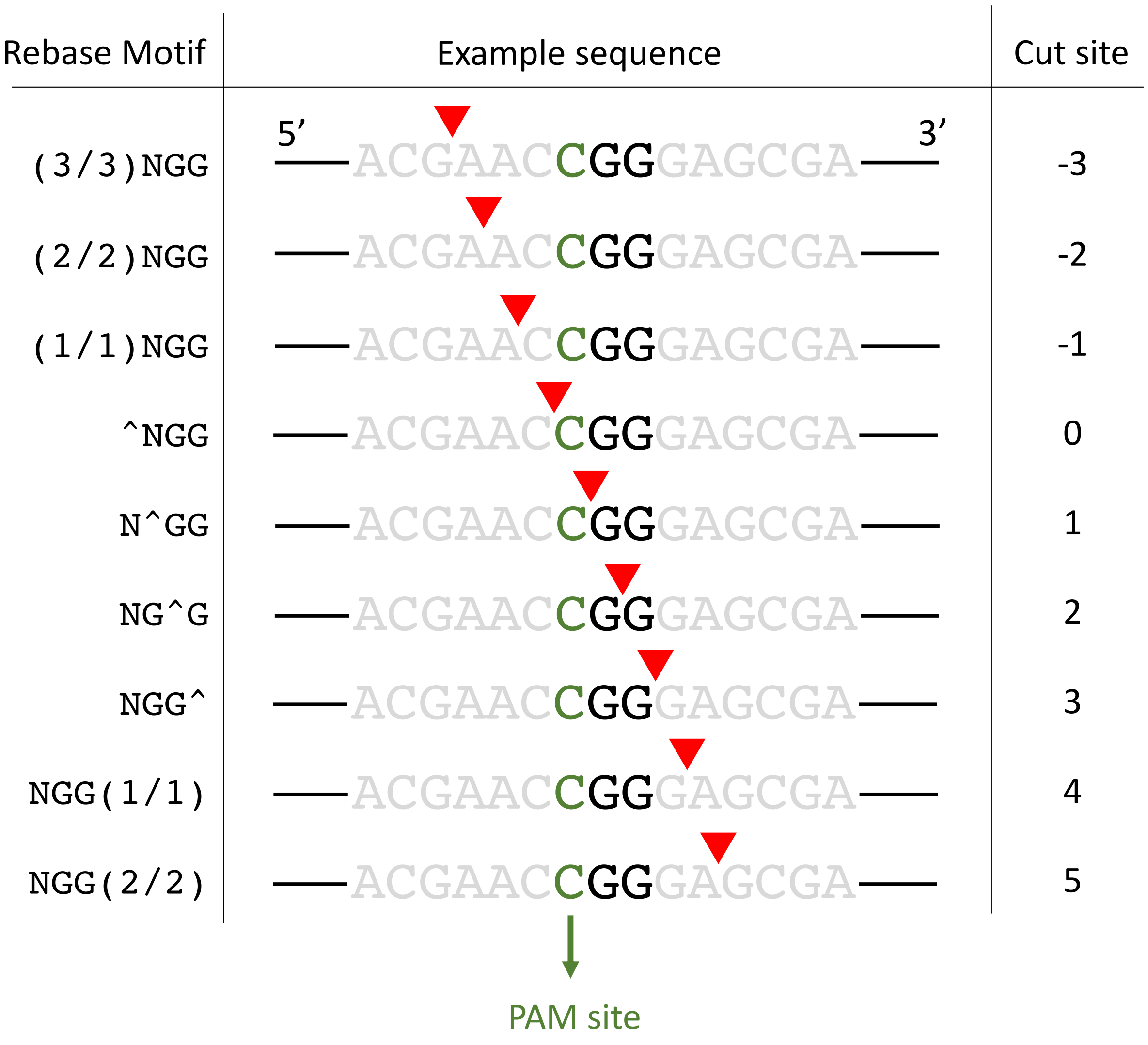


Figure 3: Examples of cut site coordinates

#### Obtaining spacer and PAM sequences from target sequences

Given a list of target sequences (protospacer + PAM) and a `CrisprNuclease` object, one can extract the protospacer and PAM sequences with the functions `extractProtospacerFromTarget` and `extractPamFromTarget`, respectively.

```
targets <- c("AGGTGCTGATTGTAGTGCTGCGG",
             "AGGTGCTGATTGTAGTGCTGAGG")
extractPamFromTarget(targets, SpCas9)
```

```
## [1] "CGG" "AGG"
```

```
extractProtospacerFromTarget(targets, SpCas9)
```

```
## [1] "AGGTGCTGATTGTAGTGCTG" "AGGTGCTGATTGTAGTGCTG"
```

#### Obtaining genomic coordinates of protospacer sequences using PAM site coordinates

Given a PAM coordinate, there are several functions in `crisprBase` that allows us to obtain the coordinates of the full PAM sequence, protospacer sequence, and target sequence: `getPamRanges`, `getTargetRanges`, and `getProtospacerRanges`, respectively. The output objects are `GRanges`:

```
chr      <- rep("chr7",2)
pam_site <- rep(200,2)
strand   <- c("+", "-")
gr_pam <- getPamRanges(seqnames=chr,
                       pam_site=pam_site,
                       strand=strand,
                       nuclease=SpCas9)
gr_protospacer <- getProtospacerRanges(seqnames=chr,
                                       pam_site=pam_site,
                                       strand=strand,
                                       nuclease=SpCas9)
gr_target <- getTargetRanges(seqnames=chr,
                             pam_site=pam_site,
                             strand=strand,
                             nuclease=SpCas9)
gr_pam
```

```
## GRanges object with 2 ranges and 0 metadata columns:
##       seqnames    ranges strand
##          <Rle> <IRanges>  <Rle>
##   [1]     chr7   200-202      +
##   [2]     chr7   198-200      -
##   -------
##   seqinfo: 1 sequence from an unspecified genome; no seqlengths
```

```
gr_protospacer
```

```
## GRanges object with 2 ranges and 0 metadata columns:
##       seqnames    ranges strand
##          <Rle> <IRanges>  <Rle>
##   [1]     chr7   180-199      +
##   [2]     chr7   201-220      -
##   -------
##   seqinfo: 1 sequence from an unspecified genome; no seqlengths
```

```r
gr_target
```

```
## GRanges object with 2 ranges and 0 metadata columns:
##       seqnames    ranges strand
##          <Rle> <IRanges>  <Rle>
##   [1]     chr7   180-202      +
##   [2]     chr7   198-220      -
##   -------
##   seqinfo: 1 sequence from an unspecified genome; no seqlengths
```

and for AsCas12a:

```r
gr_pam <- getPamRanges(seqnames=chr,
                       pam_site=pam_site,
                       strand=strand,
                       nuclease=AsCas12a)
gr_protospacer <- getProtospacerRanges(seqnames=chr,
                                       pam_site=pam_site,
                                       strand=strand,
                                       nuclease=AsCas12a)
gr_target <- getTargetRanges(seqnames=chr,
                             pam_site=pam_site,
                             strand=strand,
                             nuclease=AsCas12a)
gr_pam
```

```
## GRanges object with 2 ranges and 0 metadata columns:
##       seqnames    ranges strand
##          <Rle> <IRanges>  <Rle>
##   [1]     chr7   200-203      +
##   [2]     chr7   197-200      -
##   -------
##   seqinfo: 1 sequence from an unspecified genome; no seqlengths
```

```r
gr_protospacer
```

```
## GRanges object with 2 ranges and 0 metadata columns:
##       seqnames    ranges strand
##          <Rle> <IRanges>  <Rle>
##   [1]     chr7   204-226      +
##   [2]     chr7   174-196      -
##   -------
##   seqinfo: 1 sequence from an unspecified genome; no seqlengths
```

```r
gr_target
```

```
## GRanges object with 2 ranges and 0 metadata columns:
##       seqnames    ranges strand
##          <Rle> <IRanges>  <Rle>
##   [1]     chr7   200-226      +
##   [2]     chr7   174-200      -
##   -------
##   seqinfo: 1 sequence from an unspecified genome; no seqlengths
```

### BaseEditor class

Base editors are inactive Cas nucleases coupled with a specific deaminase. For instance, the first cytosine base editor (CBE) was obtained by coupling a cytidine deaminase with dCas9 to convert Cs to Ts (Komor et al. 2016).

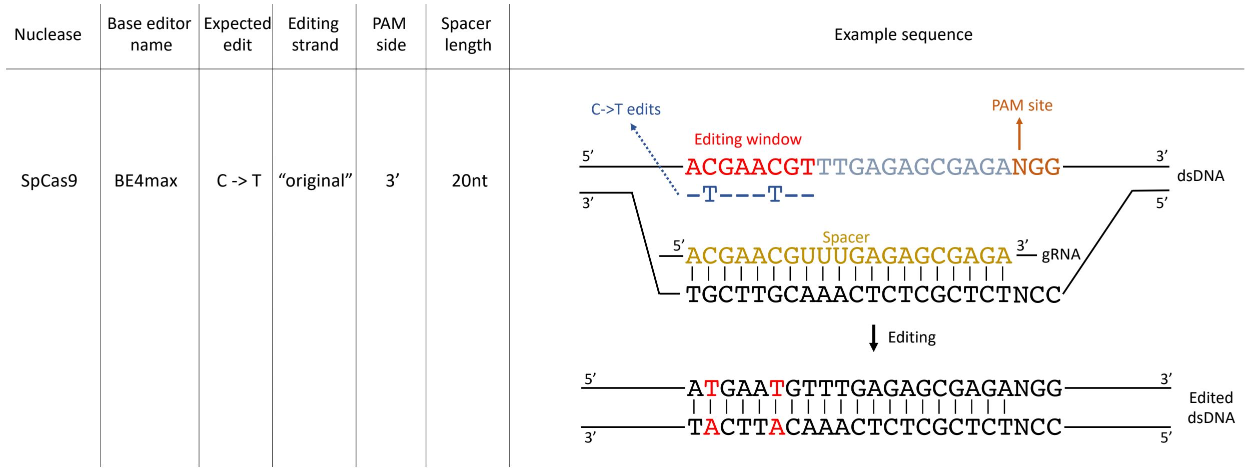


Figure 4: Examples of base editors.

We provide in `crisprBase` an S4 class, `BaseEditor`, to represent base editors. It extends the `CrisprNuclase` class with 3 additional fields:

- `baseEditorName`: string specifying the name of the base editor.
- `editingStrand`: strand where the editing happens with respect to the target protospacer sequence ("original" or "opposite").
- `editingWeights`: a matrix of experimentally-derived editing weights.

We show below how to build a `BaseEditor` object with the CBE base editor BE4max with weights obtained from Arbab et al. (2020).

We first obtain a matrix of weights for the BE4max editor stored in the package `crisprBase`:

```
# Creating weight matrix
weightsFile <- system.file("be/b4max.csv",
                           package="crisprBase",
                           mustWork=TRUE)
ws <- t(read.csv(weightsFile))
ws <- as.data.frame(ws)
```

The row names of the matrix must correspond to the nucleotide substitutions. Nucleotide substitutions that are not present in the matrix will have weight assigned to `0`.

```
rownames(ws)
```

```
## [1] "Position" "C2A"      "C2G"      "C2T"      "G2A"      "G2C"
```

The column names must correspond to the relative position with respect to the PAM site.

```
colnames(ws) <- ws["Position",]
ws <- ws[-c(match("Position", rownames(ws))),,drop=FALSE]
ws <- as.matrix(ws)
head(ws)
```

```
##     -36 -35 -34 -33 -32 -31 -30 -29 -28 -27 -26 -25 -24 -23 -22 -21 -20  -19
```

```
## C2A 0.0 0.0 0.0 0.7 0.1 0.2 0.0 0.2 0.3 0.0 0.2 0.0 0.9 0.0 0.1 0.2 0.1  0.3
## C2G 0.9 0.1 0.1 0.0 0.3 0.7 0.1 0.1 0.7 0.0 0.4 0.1 0.1 0.1 0.1 0.1 0.0  0.5
## C2T 0.7 0.7 0.8 1.8 1.0 2.0 1.4 1.2 2.3 1.3 2.4 2.2 3.4 2.2 2.1 3.5 5.8 16.2
## G2A 0.0 0.0 0.5 0.0 0.0 0.3 0.4 1.1 0.9 0.6 0.3 1.7 0.7 0.8 0.1 0.3 0.1  0.0
## G2C 0.1 0.0 0.0 0.0 0.6 2.8 0.0 0.0 0.3 0.2 0.2 0.1 0.0 0.3 0.0 0.0 0.0  0.0
##      -18  -17  -16    -15  -14  -13  -12  -11  -10  -9  -8  -7  -6  -5  -4  -3
## C2A  1.0  2.0  2.7   3.00  2.7  1.9  0.8  0.6  0.3 0.0 0.1 0.1 0.1 0.0 0.0 0.0
## C2G  1.3  2.7  4.7   5.40  5.6  3.9  1.7  0.6  0.6 0.4 0.5 0.1 0.0 0.1 0.0 0.0
## C2T 31.8 63.2 90.3 100.00 87.0 62.0 31.4 16.3 10.0 5.6 3.3 1.9 1.8 2.4 1.7 0.5
## G2A  0.0  0.0  0.1   0.01  0.0  0.0  0.0  0.0  0.0 0.0 0.0 0.0 0.0 0.2 0.2 0.0
## G2C  0.0  0.0  0.2   0.00  0.0  0.1  0.1  0.2  0.2 0.0 0.0 0.0 0.1 0.0 0.0 0.0
##      -2  -1
## C2A 0.0 0.0
## C2G 0.0 0.0
## C2T 0.2 0.1
## G2A 0.0 0.1
## G2C 0.0 0.0
```

Since BE4max uses Cas9, we can use the SpCas9 `CrisprNuclease` object available in `crisprBase` to build the `BaseEditor` object:

```
data(SpCas9, package="crisprBase")
BE4max <- BaseEditor(SpCas9,
                     baseEditorName="BE4max",
                     editingStrand="original",
                     editingWeights=ws)
metadata(BE4max)$description_base_editor <- "BE4max cytosine base editor."
BE4max
```

```
## Class: BaseEditor
##   CRISPR Nuclease name: SpCas9
##       Target type: DNA
##       Metadata: list of length 2
##       PAMs: NGG, NAG, NGA
##       Weights: 1, 0.2593, 0.0694
##       Spacer length: 20
##       PAM side: 3prime
##         Distance from PAM: 0
##       Prototype protospacers: 5'--SSSSSSSSSSSSSSSSSSSS[NGG]--3', 5'--SSSSSSSSSSSSSSSSSSSS[NAG]--3', 5
##   Base editor name: BE4max
##       Editing strand: original
##       Maximum editing weight: C2T at position -15
```

One can quickly visualize the editing weights using the function `plotEditingWeights`:

```
plotEditingWeights(BE4max)
```

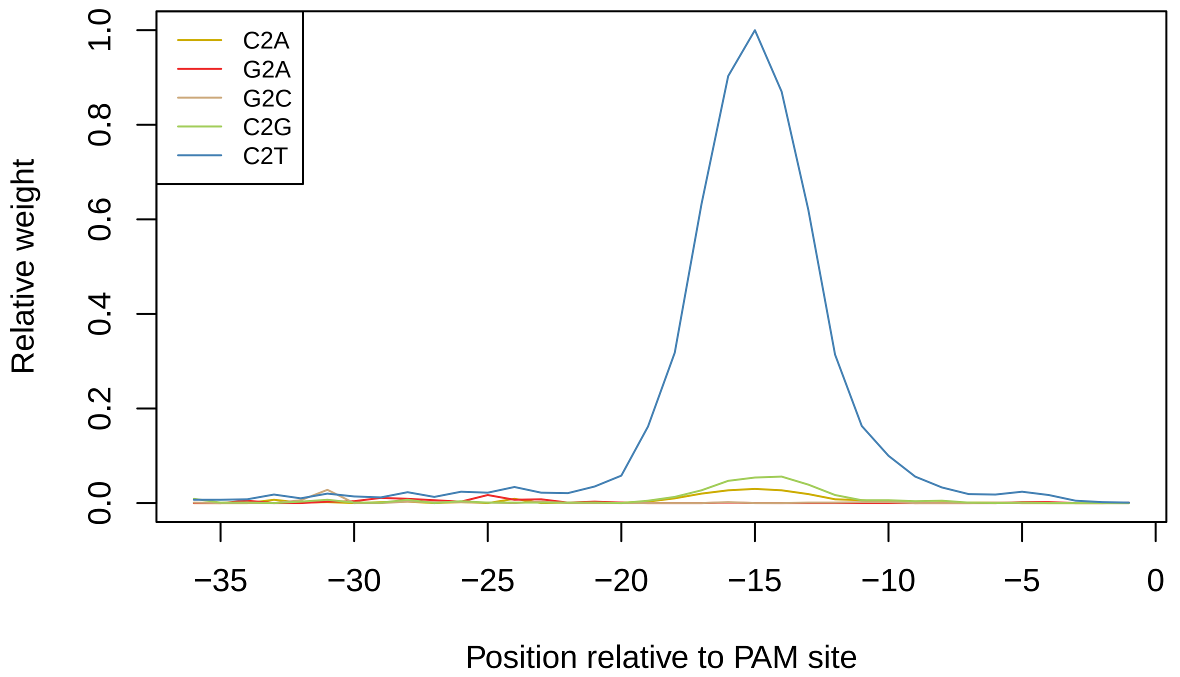


#### CrisprNickase class

CRISPR nickases can be created by mutating one of the two nuclease domains of a CRISPR nuclease. They create single-strand breaks instead of double-strand breaks.

For instance, the D10A mutation of SpCas9 inactivates the RuvC domain, and the resulting CRISPR nickase (Cas9D10A) cleaves only the strand opposite to the protospacer sequence. The H840A mutation of SpCas9 inactivates the HNN domain, and the resulting CRISPR nickase (Cas9H840A) cleaves only the strand that contains the protospacer sequence. See Figure below.

| CRISPR Nickase | Nicking strand | Rebase Motif | PAM side | Spacer length | Example sequence |
|---|---|---|---|---|---|
| SpCas9 D10A | Opposite | (3)NGG | 3′ | 20nt | 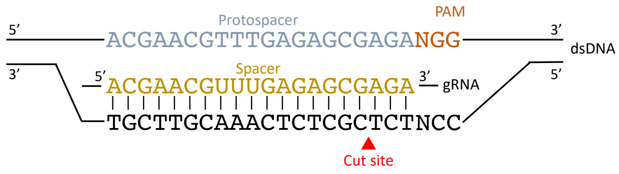 |
| SpCas9 H840A | Original | (3)NGG | 3′ | 20nt | 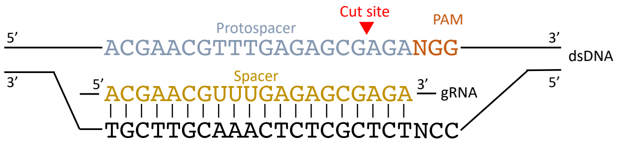 |

Figure 5: Examples of CRISPR nickases.

The `CrisprNickase` class in `crisprBase` works similar to the `CrisprNuclease` class:

```r
Cas9D10A <- CrisprNickase("Cas9D10A",
                          nickingStrand="opposite",
                          pams=c("(3)NGG", "(3)NAG", "(3)NGA"),
                          weights=c(1, 0.2593, 0.0694),
                          metadata=list(description="D10A-mutated Streptococcus
                                        pyogenes Cas9 (SpCas9) nickase"),
                          pam_side="3prime",
                          spacer_length=20)

Cas9H840A <- CrisprNickase("Cas9H840A",
                           nickingStrand="original",
                           pams=c("(3)NGG", "(3)NAG", "(3)NGA"),
                           weights=c(1, 0.2593, 0.0694),
                           metadata=list(description="H840A-mutated Streptococcus
                                         pyogenes Cas9 (SpCas9) nickase"),
                           pam_side="3prime",
                           spacer_length=20)
```

The `nickingStrand` field indicates which strand is cleaved by the nickase.

#### RNA-targeting nucleases

RNA-targeting CRISPR nucleases, such as the Cas13 family of nucleases, target single-stranded RNA (ssRNA) instead of dsDNA, as the name suggests. The equivalent of the PAM sequence is called Protospacer Flanking Sequence (PFS).

For RNA-targeting CRISPR nucleases, the spacer sequence is the reverse complement of the protospacer sequence. This differs from DNA-targeting CRISPR nucleases, for which the spacer and protospacer sequences are identical.

We can construct an RNA-targeting nuclease in way similar to a DNA-targeting nuclease by specifying `target="RNA"`. As an example, we construct a CrisprNuclease object for the CasRx nuclease (Cas13d from Ruminococcus flavefaciens strain XPD3002):

```r
CasRx <- CrisprNuclease("CasRx",
                        targetType="RNA",
                        pams="N",
                        metadata=list(description="CasRx nuclease"),
                        pam_side="3prime",
                        spacer_length=23)
CasRx
```

```
## Class: CrisprNuclease
##   Name: CasRx
##   Target type: RNA
##   Metadata: list of length 1
##   PFS: N
##   Weights: 1
##   Spacer length: 23
##   PFS side: 3prime
##     Distance from PFS: 0
##   Prototype protospacers: 5'--SSSSSSSSSSSSSSSSSSSSSSS[N]--3'
```

#### Additional notes

##### dCas9 and other "dead" nucleases

The CRISPR inhibition (CRISPRi) and CRISPR activation (CRISPRa) technologies uses modified versions of CRISPR nucleases that lack endonuclease activity, often referred to as "dead Cas" nucleases, such as the dCas9.

While fully-active Cas nucleases and dCas nucleases differ in terms of applications and type of genomic perturbations, the gRNA design remains unchanged in terms of spacer sequence search and genomic coordinates. Therefore, it is convenient to use the fully-active version of the nuclease throughout `crisprBase`.

#### Reproducibility

```
sessionInfo()
```

```
## R version 4.2.1 (2022-06-23)
## Platform: x86_64-apple-darwin17.0 (64-bit)
## Running under: macOS Catalina 10.15.7
##
## Matrix products: default
## BLAS:   /Library/Frameworks/R.framework/Versions/4.2/Resources/lib/libRblas.0.dylib
## LAPACK: /Library/Frameworks/R.framework/Versions/4.2/Resources/lib/libRlapack.dylib
##
## locale:
## [1] en_US.UTF-8/en_US.UTF-8/en_US.UTF-8/C/en_US.UTF-8/en_US.UTF-8
##
## attached base packages:
## [1] stats     graphics  grDevices utils     datasets  methods   base
##
## other attached packages:
##  [1] crisprDesign_0.99.133 crisprScore_1.1.14    crisprScoreData_1.1.3
##  [4] ExperimentHub_2.5.0   AnnotationHub_3.5.0   BiocFileCache_2.5.0
##  [7] dbplyr_2.2.1          BiocGenerics_0.43.1   crisprBowtie_1.1.1
## [10] crisprBase_1.1.5      crisprVerse_0.99.8    rmarkdown_2.15.2
##
## loaded via a namespace (and not attached):
##   [1] bitops_1.0-7                  matrixStats_0.62.0
##   [3] bit64_4.0.5                   filelock_1.0.2
##   [5] progress_1.2.2                httr_1.4.4
##   [7] GenomeInfoDb_1.33.5           tools_4.2.1
##   [9] utf8_1.2.2                    R6_2.5.1
##  [11] DBI_1.1.3                     tidyselect_1.1.2
##  [13] prettyunits_1.1.1             bit_4.0.4
##  [15] curl_4.3.2                    compiler_4.2.1
##  [17] cli_3.3.0                     Biobase_2.57.1
##  [19] basilisk.utils_1.9.1          xml2_1.3.3
##  [21] DelayedArray_0.23.1           rtracklayer_1.57.0
##  [23] randomForest_4.7-1.1          readr_2.1.2
##  [25] rappdirs_0.3.3                stringr_1.4.1
##  [27] digest_0.6.29                 Rsamtools_2.13.4
##  [29] basilisk_1.9.2                XVector_0.37.0
##  [31] pkgconfig_2.0.3               htmltools_0.5.3
##  [33] MatrixGenerics_1.9.1          highr_0.9
##  [35] fastmap_1.1.0                 BSgenome_1.65.2
##  [37] rlang_1.0.4                   RSQLite_2.2.16
```

```
##  [39] shiny_1.7.2                  BiocIO_1.7.1
##  [41] generics_0.1.3               jsonlite_1.8.0
##  [43] BiocParallel_1.31.12         dplyr_1.0.9
##  [45] VariantAnnotation_1.43.3     RCurl_1.98-1.8
##  [47] magrittr_2.0.3               GenomeInfoDbData_1.2.8
##  [49] Matrix_1.4-1                 Rcpp_1.0.9
##  [51] S4Vectors_0.35.1             fansi_1.0.3
##  [53] reticulate_1.25              Rbowtie_1.37.0
##  [55] lifecycle_1.0.1              stringi_1.7.8
##  [57] yaml_2.3.5                   SummarizedExperiment_1.27.1
##  [59] zlibbioc_1.43.0              grid_4.2.1
##  [61] blob_1.2.3                   promises_1.2.0.1
##  [63] parallel_4.2.1               crayon_1.5.1
##  [65] dir.expiry_1.5.0             lattice_0.20-45
##  [67] Biostrings_2.65.2            GenomicFeatures_1.49.6
##  [69] hms_1.1.2                    KEGGREST_1.37.3
##  [71] knitr_1.40                   pillar_1.8.1
##  [73] GenomicRanges_1.49.1         rjson_0.2.21
##  [75] codetools_0.2-18             biomaRt_2.53.2
##  [77] stats4_4.2.1                 BiocVersion_3.16.0
##  [79] XML_3.99-0.10                glue_1.6.2
##  [81] evaluate_0.16                BiocManager_1.30.18
##  [83] httpuv_1.6.5                 png_0.1-7
##  [85] vctrs_0.4.1                  tzdb_0.3.0
##  [87] purrr_0.3.4                  assertthat_0.2.1
##  [89] cachem_1.0.6                 xfun_0.32
##  [91] mime_0.12                    xtable_1.8-4
##  [93] restfulr_0.0.15              later_1.3.0
##  [95] tibble_3.1.8                 GenomicAlignments_1.33.1
##  [97] AnnotationDbi_1.59.1         memoise_2.0.1
##  [99] IRanges_2.31.2               interactiveDisplayBase_1.35.0
## [101] ellipsis_0.3.2
```

### References


Arbab, Mandana, Max W Shen, Beverly Mok, Christopher Wilson, Żaneta Matuszek, Christopher A Cassa, and David R Liu. 2020. "Determinants of Base Editing Outcomes from Target Library Analysis and Machine Learning." *Cell* 182 (2): 463–80.

Komor, Alexis C, Yongjoo B Kim, Michael S Packer, John A Zuris, and David R Liu. 2016. "Programmable Editing of a Target Base in Genomic DNA Without Double-Stranded DNA Cleavage." *Nature* 533 (7603): 420–24.

Roberts, Richard J, Tamas Vincze, Janos Posfai, and Dana Macelis. 2010. "REBASE—a Database for DNA Restriction and Modification: Enzymes, Genes and Genomes." *Nucleic Acids Research* 38 (suppl_1): D234–36.
