## Supplementary material for "The crisprVerse: a comprehensive Bioconductor ecosystem for the design of CRISPR guide RNAs across nucleases and technologies": Vignette_crisprBase

### Base functions and classes for CRISPR gRNA design

Jean-Philippe Fortin

#### 1 Overview

`crisprBase` provides S4 classes to represent nucleases, and more specifically CRISPR nucleases. It also provides arithmetic functions to extract genomic ranges to help with the design and manipulation of CRISPR guide-RNAs (gRNAs). The classes and functions are designed to work with a broad spectrum of nucleases and applications, including PAM-free CRISPR nucleases, RNA-targeting nucleases, and the more general class of restriction enzymes. It also includes functionalities for CRISPR nickases.

It provides a language and convention for our gRNA design ecosystem described in our recent bioRxiv preprint: “A comprehensive Bioconductor ecosystem for the design of CRISPR guide RNAs across nucleases and technologies”

#### 2 Installation

##### 2.1 Software requirements

###### 2.1.1 OS Requirements

This package is supported for macOS, Linux and Windows machines. It was developed and tested on R version 4.2.

##### 2.2 Installation

`crisprBase` can be installed from the Bioconductor devel branch by typing the following commands inside of an R session:

```
if (!require("BiocManager", quietly = TRUE))
    install.packages("BiocManager")

BiocManager::install(version="devel")
BiocManager::install("crisprBase")
```

###### 2.2.1 Getting started

We load `crisprBase` in the usual way:

```
library(crisprBase)
```

#### 3 Nuclease class

The `Nuclease` class is designed to store minimal information about the recognition sites of general nucleases, such as restriction enzymes. The `Nuclease` class has 5 fields: `nucleaseName`, `targetType`, `metadata`, `motifs` and `weights`. The `nucleaseName` field is a string specifying a name for the nuclease. The `targetType` specifies if the nuclease targets “DNA” (deoxyribonucleases) or “RNA” (ribonucleases). The `metadata` field is a `list` of arbitrary length to store additional information about the nuclease.

The `motifs` field is a character vector that specify one of several DNA sequence motifs that are recognized by the nuclease for cleavage (always in the 5' to 3' direction). The optional `weights` field is a numeric vector specifying relative cleavage probabilities corresponding to the motifs specified by `motifs`. Note that we use DNA to represent motifs irrespectively of the target type for simplicity.

We use the Rebase convention to represent motif sequences (Roberts et al. 2010). For enzymes that cleave within the recognition site, we add the symbol `^` within the recognition sequence to specify the cleavage site, always in the 5' to 3' direction. For enzymes that cleave away from the recognition site, we specify the distance of the cleavage site using a `(x/y)` notation where `x` represents the number of nucleotides away from the recognition sequence on the original strand, and `y` represents the number of nucleotides away from the recognition sequence on the reverse strand.

#### 3.1 Examples

The EcoRI enzyme recognizes the palindromic motif `GAATTC`, and cuts after the first nucleotide, which is specified using the `^` below:

```
library(crisprBase)

EcoRI <- Nuclease("EcoRI",
                  targetType="DNA",
                  motifs=c("G^AATTC"),
                  metadata=list(description="EcoRI restriction enzyme"))
```

The HgaI enzyme recognizes the motif `GACGC`, and cleaves DNA at 5 nucleotides downstream of the recognition sequence on the original strand, and at 10 nucleotides downstream of the recognition sequence on the reverse strand:

```
HgaI <- Nuclease("HgaI",
                 targetType="DNA",
                 motifs=c("GACGC(5/10)"),
                 metadata=list(description="HgaI restriction enzyme"))
```

In case the cleavage site was upstream of the recognition sequence, we would instead specify `(5/10)GACGC`.

Note that any nucleotide letter that is part of the extended IUPAC nucleic acid code can be used to represent recognition motifs. For instance, we use `Y` and `R` (pyrimidine and purine, respectively) to specify the possible recognition sequences for PfaAI:

```
PfaAI <- Nuclease("PfaAI",
                  targetType="DNA",
                  motifs=c("G^GYRCC"),
                  metadata=list(description="PfaAI restriction enzyme"))
```

#### 3.2 Accessor functions

The accessor function `motifs` retrieve the motif sequences:

```
motifs(PfaAI)
```

```
## DNAStringSet object of length 1:
##     width seq
## [1]     6 GGYRCC
```

To expand the motif sequence into all combinations of valid sequences with only A/C/T/G nucleotides, users can use `expand=TRUE`.

```
motifs(PfaAI, expand=TRUE)
```

```
## DNAStringSet object of length 4:
##     width seq                                               names
## [1]     6 GGCACC                                            GGYRCC
## [2]     6 GGTACC                                            GGYRCC
## [3]     6 GGCGCC                                            GGYRCC
## [4]     6 GGTGCC                                            GGYRCC
```

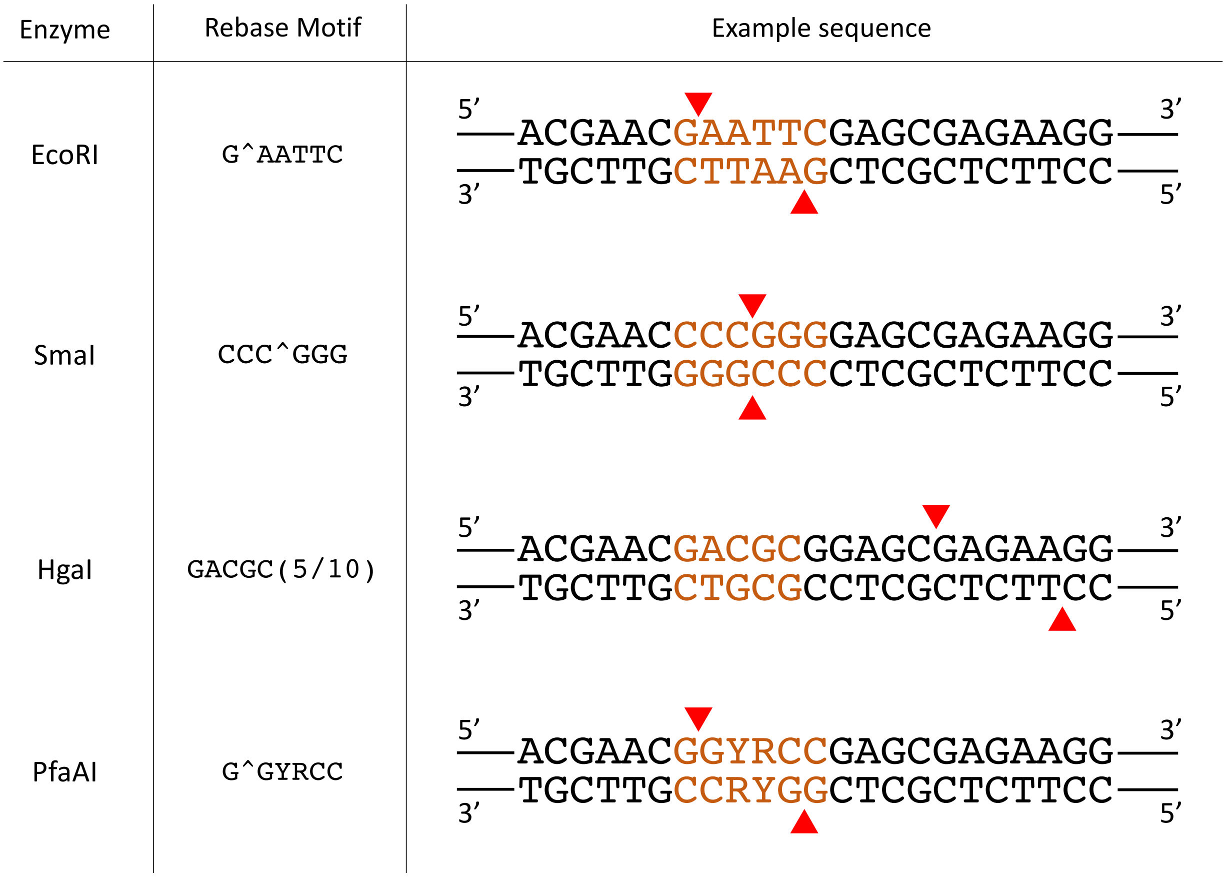


Figure 1: Examples of restriction enzymes

### 4 CrisprNuclease class

CRISPR nucleases are examples of RNA-guided nucleases. For cleavage, it requires two binding components. For CRISPR nucleases targeting DNA, the nuclease needs to first recognize a constant nucleotide motif in the target DNA called the protospacer adjacent motif (PAM) sequence. Second, the guide-RNA (gRNA), which guides the nuclease to the target sequence, needs to bind to a complementary sequence adjacent to the PAM sequence (protospacer sequence). The latter can be thought of a variable binding motif that can be specified by designing corresponding gRNA sequences. For CRISPR nucleases targeting RNA, the equivalent of the PAM sequence is called the Protospacer Flanking Sequence (PFS). We use the terms PAM and PFS interchangeably as it should be clear from context.

The `CrisprNuclease` class allows to characterize both binding components by extending the `Nuclease` class to contain information about the gRNA sequences.The PAM sequence characteristics, and the cleavage distance with respect to the PAM sequence, are specified using the motif nomenclature described in the Nuclease section above.

3 additional fields are required: `pam_side`, `spacer_length` and `spacer_gap`. The `pam_side` field can only take 2 values, `5prime` and `3prime`, and specifies on which side the PAM sequence is located with respect to the protospacer sequence. While it would be more appropriate to use the terminology `pfs_side` for RNA-targeting nucleases, we still use the term `pam_side` for simplicity.

The `spacer_length` specifies a default spacer length, and the `spacer_gap` specifies a distance (in nucleotides) between the PAM (or PFS) sequence and spacer sequence. For most nucleases,`spacer_gap=0` as the spacer sequence is located directly next to the PAM/PFS sequence.

We show how we construct a `CrisprNuclease` object for the commonly-used Cas9 nuclease (Streptococcus pyogenes Cas9):

```
SpCas9 <- CrisprNuclease("SpCas9",
                         targetType="DNA",
                         pams=c("(3/3)NGG", "(3/3)NAG", "(3/3)NGA"),
                         weights=c(1, 0.2593, 0.0694),
                         metadata=list(description="Wildtype Streptococcus pyogenes Cas9 (SpCas9) nucle
                         pam_side="3prime",
                         spacer_length=20)
SpCas9
```

```
## Class: CrisprNuclease
##   Name: SpCas9
##   Target type: DNA
##   Metadata: list of length 1
##   PAMs: NGG, NAG, NGA
##   Weights: 1, 0.2593, 0.0694
##   Spacer length: 20
##   PAM side: 3prime
##     Distance from PAM: 0
##   Prototype protospacers: 5'--SSSSSSSSSSSSSSSSSSSS[NGG]--3', 5'--SSSSSSSSSSSSSSSSSSSS[NAG]--3', 5'--S
```

Similar to the `Nuclease` class, we can specify PAM sequences using the extended nucleotide code. SaCas9 serves as a good example:

```
SaCas9 <- CrisprNuclease("SaCas9",
                         targetType="DNA",
                         pams=c("(3/3)NNGRRT"),
                         metadata=list(description="Wildtype Staphylococcus
                         aureus Cas9 (SaCas9) nuclease"),
                         pam_side="3prime",
                         spacer_length=21)
SaCas9
```

```
## Class: CrisprNuclease
##   Name: SaCas9
##   Target type: DNA
##   Metadata: list of length 1
##   PAMs: NNGRRT
##   Weights: 1
##   Spacer length: 21
##   PAM side: 3prime
##     Distance from PAM: 0
##   Prototype protospacers: 5'--SSSSSSSSSSSSSSSSSSSSS[NNGRRT]--3'
```

Here is another example where we construct a `CrisprNuclease` object for the commonly-used Cas12a nuclease (AsCas12a):

```r
AsCas12a <- CrisprNuclease("AsCas12a",
                           targetType="DNA",
                           pams="TTTV(18/23)",
                           metadata=list(description="Wildtype Acidaminococcus
                           Cas12a (AsCas12a) nuclease."),
                           pam_side="5prime",
                           spacer_length=23)
AsCas12a
```

```
## Class: CrisprNuclease
##   Name: AsCas12a
##   Target type: DNA
##   Metadata: list of length 1
##   PAMs: TTTV
##   Weights: 1
##   Spacer length: 23
##   PAM side: 5prime
##     Distance from PAM: 0
##   Prototype protospacers: 5'--[TTTV]SSSSSSSSSSSSSSSSSSSSSSS--3'
```

##### 4.1 CrisprNuclease objects provided in CrisprBase

Several already-constructed `crisprNuclease` objects are available in `crisprBase`, see `data(package="crisprBase")`.

#### 5 CRISPR arithmetics

##### 5.1 CRISPR terminology

The terms **spacer** and **protospacer** are not interchangeable. **spacer** refers to the sequence used in the gRNA construct to guide the Cas nuclease to the target **protospacer** sequence in the host genome / transcriptome. The **protospacer** sequence is adjacent to the PAM sequence / PFS sequence. We use the terminology **target** sequence to refer to the protospacer and PAM sequence taken together. For DNA-targeting nucleases such as Cas9 and Cas12a, the spacer and protospacer sequences are identical from a nucleotide point of view. For RNA-targeting nucleases such as Cas13d, the spacer and protospacer sequences are the reverse complement of each other.

An gRNA spacer sequence does not always uniquely target the host genome (a given sgRNA spacer can map to multiple protospacers in the genome). However, for a given reference genome, protospacer sequences can be uniquely identified using a combination of 3 attributes:

- **chr**: chromosome name
- **strand**: forward (+) or reverse (-)
- **pam_site**: genomic coordinate of the first nucleotide of the nuclease-specific PAM sequence. For SpCas9, this corresponds to the genomic coordinate of N in the NGG PAM sequence. For AsCas12a, this corresponds to the genomic coordinate of the first T nucleotide in the TTTV PAM sequence. For RNA-targeting nucleases, this corresponds to the first nucleotide of the PFS (we do not use `pfs_site` for simplicity).

##### 5.2 Cut site

For convention, we used the nucleotide directly downstream of the DNA cut to represent the cut site nucleotide position. For instance, for SpCas9 (blunt-ended dsDNA break), the cut site occurs at position -3 with respect to the PAM site. For AsCas12a, the 5nt overhang dsDNA break occurs at 18 nucleotides after the PAM sequence on the targeted strand. Therefore the cute site on the forward strand occurs at position 22 with respect to the PAM site, and at position 27 on the reverse strand.

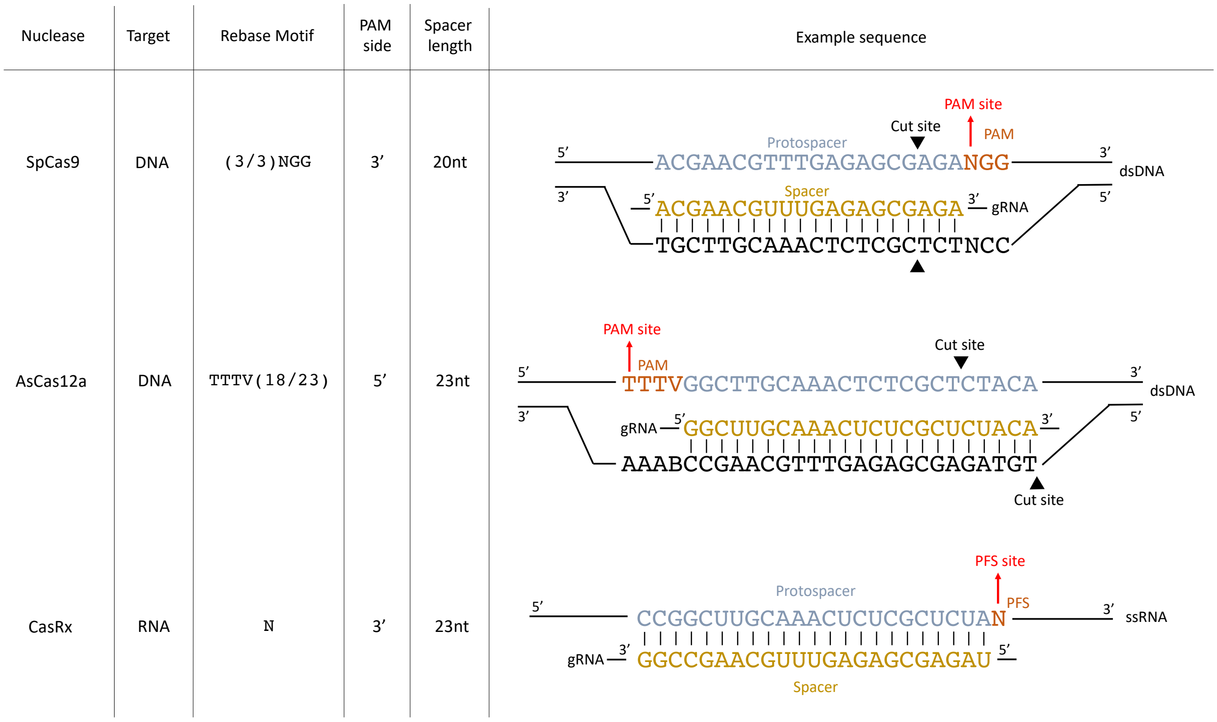


Figure 2: Examples of CRISPR nucleases

The convenience function `cutSites` extracts the cut site coordinates relative to the PAM site:

```
data(SpCas9, package="crisprBase")
data(AsCas12a, package="crisprBase")
cutSites(SpCas9)
```

```
## [1] -3
```

```
cutSites(SpCas9, strand="-")
```

```
## [1] -3
```

```
cutSites(AsCas12a)
```

```
## [1] 22
```

```
cutSites(AsCas12a, strand="-")
```

```
## [1] 27
```

Below is an illustration of how different motif sequences and cut patterns translate into cut site coordinates with respect to a PAM sequence NGG:

#### 5.3 Obtaining spacer and PAM sequences from target sequences

Given a list of target sequences (protospacer + PAM) and a `CrisprNuclease` object, one can extract protospacer and PAM sequences using the functions `extractProtospacerFromTarget` and `extractPamFromTarget`, respectively.

```
targets <- c("AGGTGCTGATTGTAGTGCTGCGG",
             "AGGTGCTGATTGTAGTGCTGAGG")
extractPamFromTarget(targets, SpCas9)
```

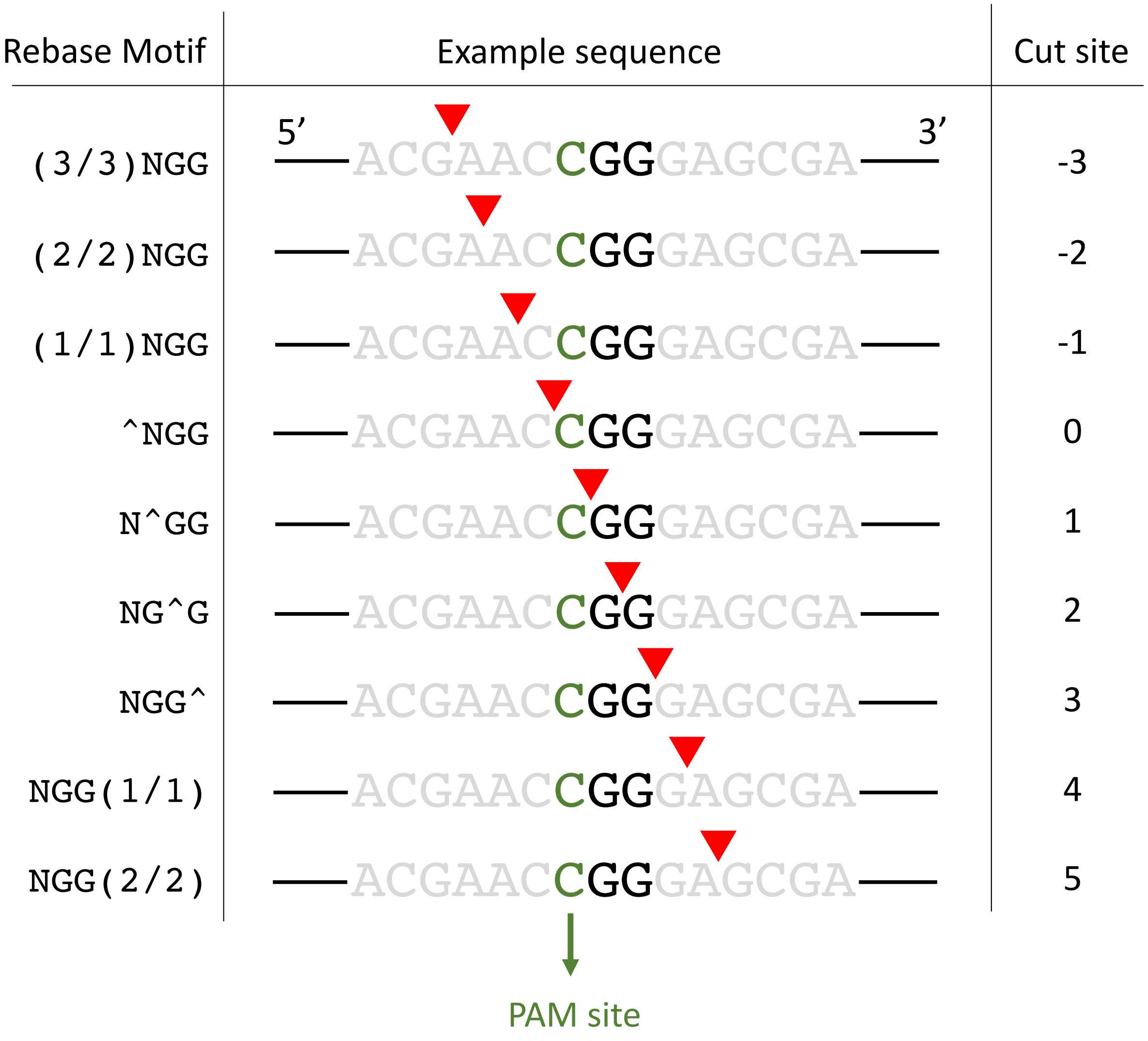


Figure 3: Examples of cut site coordinates

```
## [1] "CGG" "AGG"
```

```
extractProtospacerFromTarget(targets, SpCas9)
```

```
## [1] "AGGTGCTGATTGTAGTGCTG" "AGGTGCTGATTGTAGTGCTG"
```

We provide in `crisprBase` a S4 class, `BaseEditor`, to represent base editors. It extends the `CrisprNuclase` class with 3 additional fields:

- `baseEditorName`: string specifying the name of the base editor.

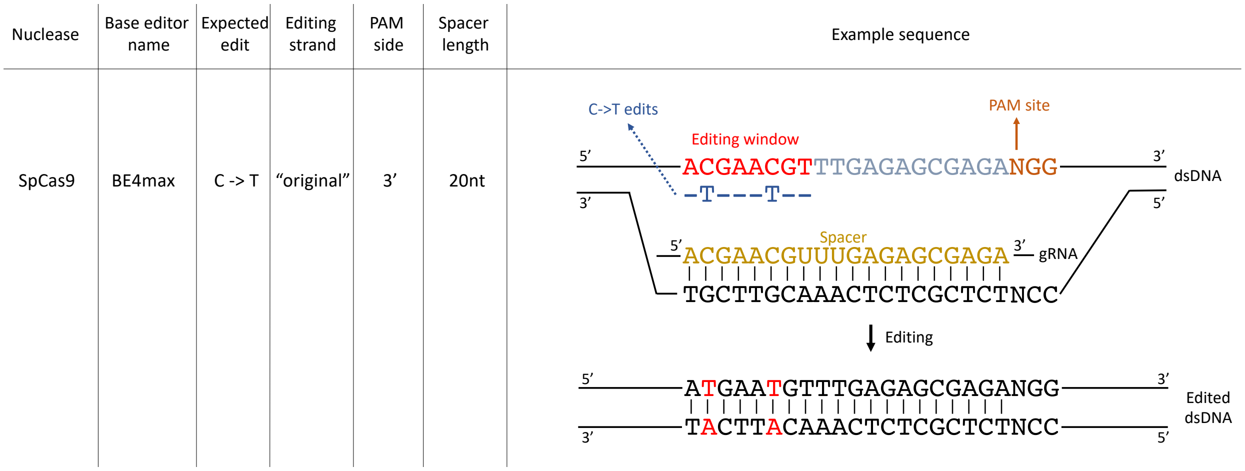


Figure 4: Examples of base editors.

One can quickly visualize the editing weights using the function `plotEditingWeights`:

```
plotEditingWeights(BE4max)
```

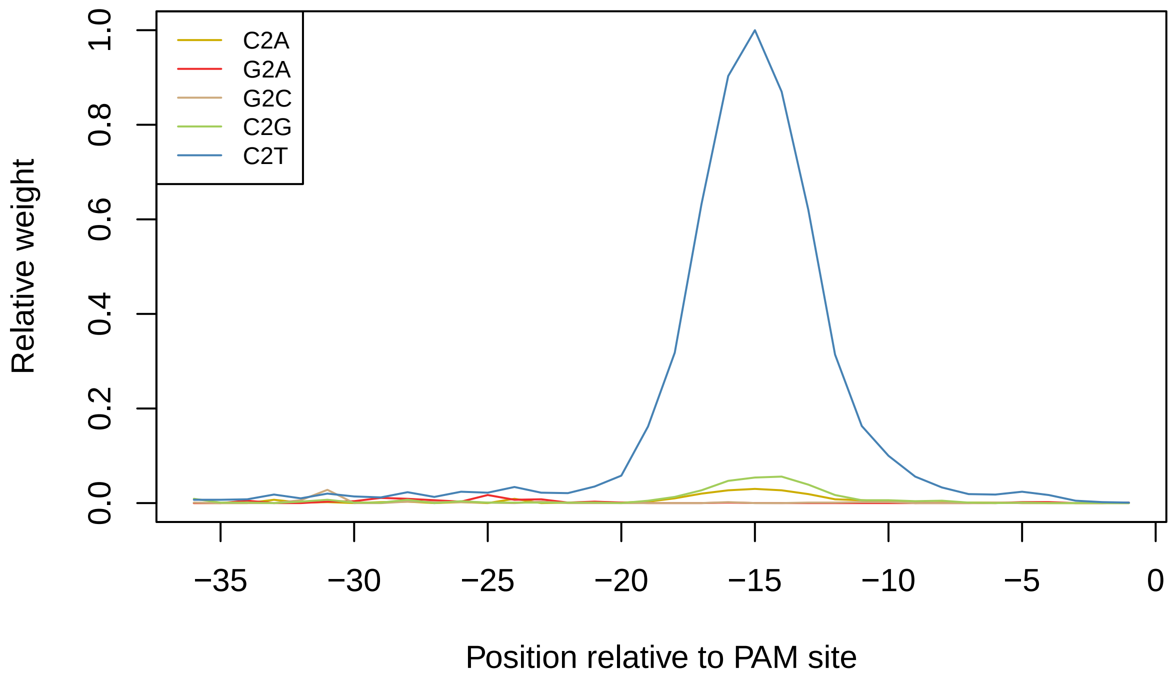


### 7 CrisprNickase class

CRISPR nickases can be created by mutating one of the two nuclease domains of a CRISPR nuclease. They create single-strand breaks instead of double-strand breaks.

For instance, the D10A mutation of SpCas9 inactivates the RuvC domain, and the resulting CRISPR nickase (Cas9D10A) cleaves only the strand opposite to the protospacer sequence. The H840A mutation of SpCas9 inactivates the HNN domain, and the resulting CRISPR nickase (Cas9H840A) cleaves only the strand that contains the protospacer sequence. See Figure below.

| CRISPR Nickase | Nicking strand | Rebase Motif | PAM side | Spacer length | Example sequence |
|---|---|---|---|---|---|
| SpCas9 D10A | Opposite | (3)NGG | 3′ | 20nt | 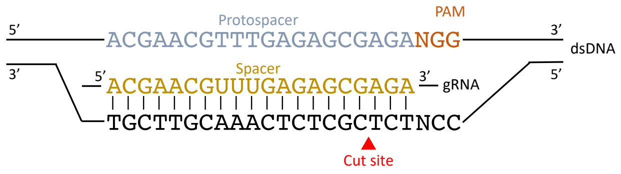 |
| SpCas9 H840A | Original | (3)NGG | 3′ | 20nt | 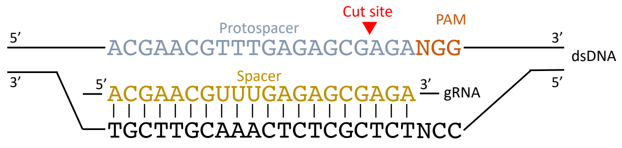 |

Figure 5: Examples of CRISPR nickases.

The `CrisprNickase` class in `crisprBase` works similar to the `CrisprNuclease` class:

```r
Cas9D10A <- CrisprNickase("Cas9D10A",
                          nickingStrand="opposite",
                          pams=c("(3)NGG", "(3)NAG", "(3)NGA"),
                          weights=c(1, 0.2593, 0.0694),
                          metadata=list(description="D10A-mutated Streptococcus
                                        pyogenes Cas9 (SpCas9) nickase"),
                          pam_side="3prime",
                          spacer_length=20)

Cas9H840A <- CrisprNickase("Cas9H840A",
                           nickingStrand="original",
                           pams=c("(3)NGG", "(3)NAG", "(3)NGA"),
                           weights=c(1, 0.2593, 0.0694),
                           metadata=list(description="H840A-mutated Streptococcus
                                         pyogenes Cas9 (SpCas9) nickase"),
                           pam_side="3prime",
                           spacer_length=20)
```

The `nickingStrand` field indicates which strand is being cleaved by the nickase.

### 8 RNA-targeting nucleases

RNA-targeting CRISPR nucleases, such as the Cas13 family of nucleases, target single-stranded RNA (ssRNA) instead of dsDNA as the name suggests. The equivalent of the PAM sequence is called Protospacer Flanking Sequence (PFS).

For RNA-targeting CRISPR nucleases, the spacer sequence is the reverse complement of the protospacer sequence. This differs from DNA-targeting CRISPR nucleases, for which the spacer and protospacer sequences are identical.

We can construct an RNA-targeting nuclease in way similar to a DNA-targeting nuclease by specifying `target="RNA"`. As an example, we construct below a CrisprNuclease object for the CasRx nuclease (Cas13d from Ruminococcus flavefaciens strain XPD3002):

```r
CasRx <- CrisprNuclease("CasRx",
                        targetType="RNA",
                        pams="N",
                        metadata=list(description="CasRx nuclease"),
                        pam_side="3prime",
                        spacer_length=23)
CasRx
```

```
## Class: CrisprNuclease
##   Name: CasRx
##   Target type: RNA
##   Metadata: list of length 1
##   PFS: N
##   Weights: 1
##   Spacer length: 23
##   PFS side: 3prime
##     Distance from PFS: 0
##   Prototype protospacers: 5'--SSSSSSSSSSSSSSSSSSSSSSS[N]--3'
```

### 9 Additional notes

#### 9.1 dCas9 and other "dead" nucleases

The CRISPR inhibition (CRISPRi) and CRISPR activation (CRISPRa) technologies uses modified versions of CRISPR nucleases that lack endonuclease activity, often referred to as "dead Cas" nucleases, such as the dCas9.

While fully-active Cas nucleases and dCas nucleases differ in terms of applications and type of genomic perturbations, the gRNA design remains unchanged in terms of spacer sequence search and genomic coordinates. Therefore it is convenient to use the fully-active version of the nuclease throughout `crisprBase`.

### 10 License

The project as a whole is covered by the MIT license.

### 11 Reproducibility

```
sessionInfo()
```

```
## R version 4.2.1 (2022-06-23)
## Platform: x86_64-apple-darwin17.0 (64-bit)
## Running under: macOS Catalina 10.15.7
##
## Matrix products: default
## BLAS:   /Library/Frameworks/R.framework/Versions/4.2/Resources/lib/libRblas.0.dylib
## LAPACK: /Library/Frameworks/R.framework/Versions/4.2/Resources/lib/libRlapack.dylib
##
## locale:
## [1] en_US.UTF-8/en_US.UTF-8/en_US.UTF-8/C/en_US.UTF-8/en_US.UTF-8
##
## attached base packages:
## [1] stats     graphics  grDevices utils     datasets  methods   base
##
## other attached packages:
## [1] crisprBase_1.1.5
##
## loaded via a namespace (and not attached):
##  [1] rstudioapi_0.14        knitr_1.40             XVector_0.37.0
##  [4] magrittr_2.0.3         GenomicRanges_1.49.1   BiocGenerics_0.43.1
##  [7] zlibbioc_1.43.0        IRanges_2.31.2         rlang_1.0.4
## [10] fastmap_1.1.0          highr_0.9              stringr_1.4.1
## [13] GenomeInfoDb_1.33.5    tools_4.2.1            xfun_0.32
## [16] cli_3.3.0              htmltools_0.5.3        yaml_2.3.5
## [19] digest_0.6.29          crayon_1.5.1           GenomeInfoDbData_1.2.8
## [22] S4Vectors_0.35.1       bitops_1.0-7           RCurl_1.98-1.8
## [25] evaluate_0.16          rmarkdown_2.15.2       stringi_1.7.8
## [28] compiler_4.2.1         Biostrings_2.65.2      stats4_4.2.1
```

### References


Arbab, Mandana, Max W Shen, Beverly Mok, Christopher Wilson, Żaneta Matuszek, Christopher A Cassa, and David R Liu. 2020. "Determinants of Base Editing Outcomes from Target Library Analysis and Machine Learning." *Cell* 182 (2): 463–80.

Komor, Alexis C, Yongjoo B Kim, Michael S Packer, John A Zuris, and David R Liu. 2016. "Programmable Editing of a Target Base in Genomic DNA Without Double-Stranded DNA Cleavage." *Nature* 533 (7603):

420–24.

Roberts, Richard J, Tamas Vincze, Janos Posfai, and Dana Macelis. 2010. "REBASE—a Database for DNA Restriction and Modification: Enzymes, Genes and Genomes." *Nucleic Acids Research* 38 (suppl_1): D234–36.
