## Supplementary material for "The crisprVerse: a comprehensive Bioconductor ecosystem for the design of CRISPR guide RNAs across nucleases and technologies": Vignette_crisprBowtie

### crisprBowtie: alignment of gRNA spacer sequences using bowtie

Jean-Philippe Fortin

2022-08-30

### 1 Overview of crisprBowtie

`crisprBowtie` provides two main functions to align short DNA sequences to a reference genome using the short read aligner bowtie (Langmead et al. 2009) and return the alignments as R objects: `runBowtie` and `runCrisprBowtie`. It utilizes the Bioconductor package `Rbowtie` to access the bowtie program in a platform-independent manner. This means that users do not need to install bowtie prior to using `crisprBowtie`.

The latter function (`runCrisprBowtie`) is specifically designed to map and annotate CRISPR guide RNA (gRNA) spacer sequences using CRISPR nuclease objects and CRISPR genomic arithmetics defined in the Bioconductor `crisprBase` package. This enables a fast and accurate on-target and off-target search of gRNA spacer sequences for virtually any type of CRISPR nucleases.

### 2 Installation and getting started

#### 2.1 Software requirements

##### 2.1.1 OS Requirements

This package is supported for macOS, Linux and Windows machines. Package was developed and tested on R version 4.2.

##### 2.1.2 R Dependencies

- crisprBase: https://github.com/Jfortin1/crisprBase
- RBowtie: https://bioconductor.org/packages/release/bioc/html/Rbowtie.html

#### 2.2 Installation from Bioconductor

`crisprBowtie` can be installed from from the Bioconductor devel branch using the following commands in a fresh R session:

```
if (!require("BiocManager", quietly = TRUE))
    install.packages("BiocManager")

BiocManager::install(version="devel")
BiocManager::install("crisprBowtie")
```

### 3 Building a bowtie index

To use `runBowtie` or `runCrisprBowtie`, users need to first build a bowtie genome index. For a given genome, this step has to be done only once. The `Rbowtie` package convenitently provides the function `bowtie_build` to build a bowtie index from any custom genome from a FASTA file.

As an example, we build a bowtie index for a small portion of the human chromosome 1 (`chr1.fa` file provided in the `crisprBowtie` package) and save the index file as `myIndex` to a temporary directory:

```
library(Rbowtie)
fasta <- file.path(find.package("crisprBowtie"), "example/chr1.fa")
tempDir <- tempdir()
Rbowtie::bowtie_build(fasta,
                      outdir=tempDir,
                      force=TRUE,
                      prefix="myIndex")
```

### 4 Alignment using `runCrisprBowtie`

As an example, we align 6 spacer sequences (of length 20bp) to the custom genome built above, allowing a maximum of 3 mismatches between the spacer and protospacer sequences.

We specify that the search is for the wildtype Cas9 (SpCas9) nuclease by providing the `CrisprNuclease` object `SpCas9` available through the `crisprBase` package. The argument `canonical=FALSE` specifies that non-canonical PAM sequences are also considered (NAG and NGA for SpCas9). The function `getAvailableCrisprNucleases` in `crisprBase` returns a character vector of available `crisprNuclease` objects found in `crisprBase`.

```
library(crisprBowtie)
data(SpCas9, package="crisprBase")
crisprNuclease <- SpCas9
spacers <- c("TCCGCGGGCGACAATGGCAT",
             "TGATCCCGCGCTCCCCGATG",
             "CCGGGAGCCGGGGCTGGACG",
             "CCACCCTCAGGTGTGCGGCC",
             "CGGAGGGCTGCAGAAAGCCT",
             "GGTGATGGCGCGGGCCGGGC")
runCrisprBowtie(spacers,
                crisprNuclease=crisprNuclease,
                n_mismatches=3,
                canonical=FALSE,
                bowtie_index=file.path(tempDir, "myIndex"))
```

```
## [runCrisprBowtie] Searching for SpCas9 protospacers

##                 spacer          protospacer pam  chr pam_site strand
## 1 CCACCCTCAGGTGTGCGGCC CCACCCTCAGGTGTGCGGCC TGG chr1      679      +
## 2 CCGGGAGCCGGGGCTGGACG CCGGGAGCCGGGGCTGGACG GAG chr1      466      +
## 3 CGGAGGGCTGCAGAAAGCCT CGGAGGGCTGCAGAAAGCCT TGG chr1      706      +
## 4 GGTGATGGCGCGGGCCGGGC GGTGATGGCGCGGGCCGGGC CGG chr1      831      +
## 5 TGATCCCGCGCTCCCCGATG TGATCCCGCGCTCCCCGATG CAG chr1      341      +
##   n_mismatches canonical
## 1            0      TRUE
## 2            0     FALSE
## 3            0      TRUE
## 4            0      TRUE
## 5            0     FALSE
```

### 5 Applications beyond CRISPR

The function `runBowtie` is similar to `runCrisprBowtie`, but does not impose constraints on PAM sequences. It can be used to search for any short read sequence in a genome.

#### 5.1 Example using RNAi (siRNA design)

Seed-related off-targets caused by mismatch tolerance outside of the seed region is a well-studied and characterized problem observed in RNA interference (RNA) experiments. `runBowtie` can be used to map shRNA/siRNA seed sequences to reference genomes to predict putative off-targets:

```
seeds <- c("GTAAAGGT", "AAGGATTG")
runBowtie(seeds,
          n_mismatches=2,
          bowtie_index=file.path(tempDir, "myIndex"))
```

```
##      query   target  chr pos strand n_mismatches
## 1 AAGGATTG AAAGAATG chr1 163      -            2
## 2 AAGGATTG AAGCCTTG chr1 700      +            2
## 3 AAGGATTG AAGGCTTT chr1 699      -            2
## 4 AAGGATTG CAGGCTTG chr1 905      -            2
## 5 GTAAAGGT GGGAAGGT chr1 724      +            2
```

### 6 Reproducibility

```
sessionInfo()
```

```
## R version 4.2.1 (2022-06-23)
## Platform: x86_64-apple-darwin17.0 (64-bit)
## Running under: macOS Catalina 10.15.7
##
## Matrix products: default
## BLAS:   /Library/Frameworks/R.framework/Versions/4.2/Resources/lib/libRblas.0.dylib
## LAPACK: /Library/Frameworks/R.framework/Versions/4.2/Resources/lib/libRlapack.dylib
##
## locale:
## [1] en_US.UTF-8/en_US.UTF-8/en_US.UTF-8/C/en_US.UTF-8/en_US.UTF-8
##
## attached base packages:
## [1] stats     graphics  grDevices utils     datasets  methods   base
##
## other attached packages:
## [1] crisprBowtie_1.1.1 Rbowtie_1.37.0
##
## loaded via a namespace (and not attached):
##  [1] SummarizedExperiment_1.27.1 tidyselect_1.1.2
##  [3] xfun_0.32                   purrr_0.3.4
##  [5] lattice_0.20-45             vctrs_0.4.1
##  [7] htmltools_0.5.3             stats4_4.2.1
##  [9] rtracklayer_1.57.0          yaml_2.3.5
## [11] utf8_1.2.2                  XML_3.99-0.10
## [13] rlang_1.0.4                 pillar_1.8.1
## [15] glue_1.6.2                  BiocParallel_1.31.12
## [17] bit64_4.0.5                 BiocGenerics_0.43.1
```

```
## [19] matrixStats_0.62.0         GenomeInfoDbData_1.2.8
## [21] lifecycle_1.0.1            stringr_1.4.1
## [23] zlibbioc_1.43.0            MatrixGenerics_1.9.1
## [25] Biostrings_2.65.2          codetools_0.2-18
## [27] evaluate_0.16              restfulr_0.0.15
## [29] Biobase_2.57.1             knitr_1.40
## [31] tzdb_0.3.0                 IRanges_2.31.2
## [33] fastmap_1.1.0              GenomeInfoDb_1.33.5
## [35] parallel_4.2.1             fansi_1.0.3
## [37] crisprBase_1.1.5           readr_2.1.2
## [39] BSgenome_1.65.2            DelayedArray_0.23.1
## [41] S4Vectors_0.35.1           vroom_1.5.7
## [43] XVector_0.37.0             bit_4.0.4
## [45] Rsamtools_2.13.4           rjson_0.2.21
## [47] hms_1.1.2                  digest_0.6.29
## [49] stringi_1.7.8              BiocIO_1.7.1
## [51] GenomicRanges_1.49.1       grid_4.2.1
## [53] cli_3.3.0                  tools_4.2.1
## [55] bitops_1.0-7               magrittr_2.0.3
## [57] RCurl_1.98-1.8             tibble_3.1.8
## [59] crayon_1.5.1               pkgconfig_2.0.3
## [61] ellipsis_0.3.2             Matrix_1.4-1
## [63] rmarkdown_2.15.2           rstudioapi_0.14
## [65] R6_2.5.1                   GenomicAlignments_1.33.1
## [67] compiler_4.2.1
```

### References

Langmead, Ben, Cole Trapnell, Mihai Pop, and Steven L. Salzberg. 2009. "Ultrafast and Memory-Efficient Alignment of Short DNA Sequences to the Human Genome." *Genome Biology* 10 (3): R25. https://doi.org/10.1186/gb-2009-10-3-r25.
