## Supplementary material for "The crisprVerse: a comprehensive Bioconductor ecosystem for the design of CRISPR guide RNAs across nucleases and technologies": Vignette_crisprBwa

### crisprBwa: alignment of gRNA spacer sequences using BWA

Jean-Philippe Fortin

2022-08-30

### 1 Overview of crisprBwa

`crisprBwa` provides two main functions to align short DNA sequences to a reference genome using the short read aligner BWA-backtrack (Li and Durbin 2009) and return the alignments as R objects: `runBwa` and `runCrisprBwa`. It utilizes the Bioconductor package `Rbwa` to access the BWA program in a platform-independent manner. This means that users do not need to install BWA prior to using `crisprBwa`.

The latter function (`runCrisprBwa`) is specifically designed to map and annotate CRISPR guide RNA (gRNA) spacer sequences using CRISPR nuclease objects and CRISPR genomic arithmetics defined in the Bioconductor `crisprBase` package. This enables a fast and accurate on-target and off-target search of gRNA spacer sequences for virtually any type of CRISPR nucleases.

### 2 Installation and getting started

#### 2.1 Software requirements

##### 2.1.1 OS Requirements

This package is supported for macOS and Linux only. Package was developed and tested on R version 4.2.

##### 2.1.2 R Dependencies

- crisprBase: https://github.com/Jfortin1/crisprBase
- RBwa: https://github.com/Jfortin1/Rbwa

#### 2.2 Installation from Bioconductor

`crisprBwa` can be installed from from the Bioconductor devel branch using the following commands in a fresh R session:

```
if (!require("BiocManager", quietly = TRUE))
    install.packages("BiocManager")

BiocManager::install(version="devel")
BiocManager::install("crisprBwa")
```

### 3 Building a bwa index

To use `runBwa` or `runCrisprBwa`, users need to first build a BWA genome index. For a given genome, this step has to be done only once. The `Rbwa` package conveniently provides the function `bwa_build_index` to build a BWA index from any custom genome from a FASTA file.

As an example, we build a BWA index for a small portion of the human chromosome 12 (`chr12.fa` file provided in the `crisprBwa` package) and save the index file as `myIndex` to a temporary directory:

```
library(Rbwa)
fasta <- system.file(package="crisprBwa", "example/chr12.fa")
outdir <- tempdir()
index <- file.path(outdir, "chr12")
Rbwa::bwa_build_index(fasta,
                      index_prefix=index)
```

### 4 Alignment using runCrisprBwa

As an example, we align 5 spacer sequences (of length 20bp) to the custom genome built above, allowing a maximum of 3 mismatches between the spacer and protospacer sequences.

We specify that the search is for the wildtype Cas9 (SpCas9) nuclease by providing the `CrisprNuclease` object `SpCas9` available through the `crisprBase` package. The argument `canonical=FALSE` specifies that non-canonical PAM sequences are also considered (NAG and NGA for SpCas9). The function `getAvailableCrisprNucleases` in `crisprBase` returns a character vector of available `crisprNuclease` objects found in `crisprBase`.

We also need to provide a `BSgenome` object corresponding to the reference genome used for alignment to extract protospacer and PAM sequences of the target sequences.

```
library(crisprBwa)
library(BSgenome.Hsapiens.UCSC.hg38)
```

```
## Loading required package: BSgenome

## Loading required package: BiocGenerics

##
## Attaching package: 'BiocGenerics'

## The following objects are masked from 'package:stats':
##
##     IQR, mad, sd, var, xtabs

## The following objects are masked from 'package:base':
##
##     anyDuplicated, append, as.data.frame, basename, cbind, colnames,
##     dirname, do.call, duplicated, eval, evalq, Filter, Find, get, grep,
##     grepl, intersect, is.unsorted, lapply, Map, mapply, match, mget,
##     order, paste, pmax, pmax.int, pmin, pmin.int, Position, rank,
##     rbind, Reduce, rownames, sapply, setdiff, sort, table, tapply,
##     union, unique, unsplit, which.max, which.min

## Loading required package: S4Vectors

## Loading required package: stats4

##
## Attaching package: 'S4Vectors'

## The following objects are masked from 'package:base':
##
##     expand.grid, I, unname

## Loading required package: IRanges

## Loading required package: GenomeInfoDb

## Loading required package: GenomicRanges
```

```
## Loading required package: Biostrings

## Loading required package: XVector

##
## Attaching package: 'Biostrings'

## The following object is masked from 'package:base':
##
##     strsplit

## Loading required package: rtracklayer
```

```
data(SpCas9, package="crisprBase")
crisprNuclease <- SpCas9
bsgenome <- BSgenome.Hsapiens.UCSC.hg38
spacers <- c("AGCTGTCCGTGGGGGTCCGC",
             "CCCCTGCTGCTGTGCCAGGC",
             "ACGAACTGTAAAAGGCTTGG",
             "ACGAACTGTAACAGGCTTGG",
             "AAGGCCCTCAGAGTAATTAC")
runCrisprBwa(spacers,
             bsgenome=bsgenome,
             crisprNuclease=crisprNuclease,
             n_mismatches=3,
             canonical=FALSE,
             bwa_index=index)
```

```
## [runCrisprBwa] Using BSgenome.Hsapiens.UCSC.hg38
## [runCrisprBwa] Searching for SpCas9 protospacers

##                 spacer          protospacer pam   chr pam_site strand
## 1 AAGGCCCTCAGAGTAATTAC AAGGCCCTCAGAGTAATTAC AGA chr12   170636      +
## 2 ACGAACTGTAAAAGGCTTGG ACGAACTGTAAAAGGCTTGG AGG chr12   170815      -
## 3 ACGAACTGTAACAGGCTTGG ACGAACTGTAAAAGGCTTGG AGG chr12   170815      -
## 4 AGCTGTCCGTGGGGGTCCGC AGCTGTCCGTGGGGGTCCGC AGG chr12   170585      +
## 5 CCCCTGCTGCTGTGCCAGGC CCCCTGCTGCTGTGCCAGGC CGG chr12   170609      +
##   n_mismatches canonical
## 1            0     FALSE
## 2            0      TRUE
## 3            1      TRUE
## 4            0      TRUE
## 5            0      TRUE
```

### 5 Applications beyond CRISPR

The function `runBwa` is similar to `runCrisprBwa`, but does not impose constraints on PAM sequences. It can be used to search for any short read sequence in a genome.

#### 5.1 Example using RNAi (siRNA design)

Seed-related off-targets caused by mismatch tolerance outside of the seed region is a well-studied and characterized problem observed in RNA interference (RNAi) experiments. `runBWa` can be used to map shRNA/siRNA seed sequences to reference genomes to predict putative off-targets:

```
seeds <- c("GTAAGCGGAGTGT", "AACGGGGAGATTG")
runBwa(seeds,
```

```r
      n_mismatches=2,
      bwa_index=index)
```

```
##           query   chr    pos strand n_mismatches
## 1 AACGGGGAGATTG chr12  68337      -            2
## 2 AACGGGGAGATTG chr12   1666      -            2
## 3 AACGGGGAGATTG chr12 123863      +            2
## 4 AACGGGGAGATTG chr12 151731      -            2
## 5 AACGGGGAGATTG chr12 110901      +            2
## 6 GTAAGCGGAGTGT chr12 101550      -            2
```

### 6 Reproducibility

```r
sessionInfo()
```

```
## R version 4.2.1 (2022-06-23)
## Platform: x86_64-apple-darwin17.0 (64-bit)
## Running under: macOS Catalina 10.15.7
##
## Matrix products: default
## BLAS:   /Library/Frameworks/R.framework/Versions/4.2/Resources/lib/libRblas.0.dylib
## LAPACK: /Library/Frameworks/R.framework/Versions/4.2/Resources/lib/libRlapack.dylib
##
## locale:
## [1] en_US.UTF-8/en_US.UTF-8/en_US.UTF-8/C/en_US.UTF-8/en_US.UTF-8
##
## attached base packages:
## [1] stats4    stats     graphics  grDevices utils     datasets  methods
## [8] base
##
## other attached packages:
##  [1] BSgenome.Hsapiens.UCSC.hg38_1.4.4 BSgenome_1.65.2
##  [3] rtracklayer_1.57.0                Biostrings_2.65.2
##  [5] XVector_0.37.0                    GenomicRanges_1.49.1
##  [7] GenomeInfoDb_1.33.5               IRanges_2.31.2
##  [9] S4Vectors_0.35.1                  BiocGenerics_0.43.1
## [11] crisprBwa_1.1.3                   Rbwa_1.1.0
##
## loaded via a namespace (and not attached):
##  [1] SummarizedExperiment_1.27.1 tidyselect_1.1.2
##  [3] xfun_0.32                   purrr_0.3.4
##  [5] lattice_0.20-45             vctrs_0.4.1
##  [7] htmltools_0.5.3             yaml_2.3.5
##  [9] utf8_1.2.2                  XML_3.99-0.10
## [11] rlang_1.0.4                 pillar_1.8.1
## [13] glue_1.6.2                  BiocParallel_1.31.12
## [15] bit64_4.0.5                 matrixStats_0.62.0
## [17] GenomeInfoDbData_1.2.8      lifecycle_1.0.1
## [19] stringr_1.4.1               zlibbioc_1.43.0
## [21] MatrixGenerics_1.9.1        codetools_0.2-18
## [23] evaluate_0.16               restfulr_0.0.15
## [25] Biobase_2.57.1              knitr_1.40
## [27] tzdb_0.3.0                  fastmap_1.1.0
```

```
## [29] parallel_4.2.1            fansi_1.0.3
## [31] crisprBase_1.1.5          readr_2.1.2
## [33] DelayedArray_0.23.1       vroom_1.5.7
## [35] bit_4.0.4                 Rsamtools_2.13.4
## [37] rjson_0.2.21              hms_1.1.2
## [39] digest_0.6.29             stringi_1.7.8
## [41] BiocIO_1.7.1              grid_4.2.1
## [43] cli_3.3.0                 tools_4.2.1
## [45] bitops_1.0-7              magrittr_2.0.3
## [47] RCurl_1.98-1.8            tibble_3.1.8
## [49] crayon_1.5.1              pkgconfig_2.0.3
## [51] ellipsis_0.3.2            Matrix_1.4-1
## [53] rmarkdown_2.15.2          rstudioapi_0.14
## [55] R6_2.5.1                  GenomicAlignments_1.33.1
## [57] compiler_4.2.1
```

### References

Li, Heng, and Richard Durbin. 2009. "Fast and Accurate Short Read Alignment with Burrows–Wheeler Transform." *Bioinformatics* 25 (14): 1754–60.
