## Supplementary material for "The crisprVerse: a comprehensive Bioconductor ecosystem for the design of CRISPR guide RNAs across nucleases and technologies": Vignette_crisprDesign

### Introduction to crisprDesign

2022-08-30

#### 1 Introduction

`crisprDesign` is the core package of the crisprVerse, and plays the role of a one-stop shop for designing and annotating CRISPR guide RNA (gRNA) sequences. This includes the characterization of on-targets and off-targets using different aligners, on- and off-target scoring, gene context annotation, SNP annotation, sequence feature characterization, repeat annotation, and many more.
The software was developed to be as applicable and generalizable as possible.

It currently support five types of CRISPR modalities (modes of perturbations): CRISPR knockout (CRISPRko), CRISPR activation (CRISPRa), CRISPR interference (CRISPRi), CRISPR base editing (CRISPRbe), and CRISPR knockdown (CRISPRkd) (see Kampmann (2018) for a review of CRISPR modalities).

It utilizes the `crisprBase` package to enable gRNA design for any CRISPR nuclease and base editor via the `CrisprNuclease` and `BaseEditor` classes, respectively. Nucleases that are commonly used in the field are provided, including DNA-targeting nucleases (e.g. SpCas9, AsCas12a) and RNA-targeting nucleases (e.g. CasRx (RfxCas13d)).

`crisprDesign` is fully developed to work with the genome of any organism, and can also be used to design gRNAs targeting custom DNA sequences.

Finally, more specialized gRNA design functionalities are also available, including design for optical pooled screening (OPS), paired gRNA design, and gRNA filtering and ranking functionalities.

This vignette is meant to be an overview of the main features included in the package, using toy examples for the sake of time (the vignette has to compile within a few minutes, as required by Bioconductor). For detailed and comprehensive tutorials, please visit our crisprVerse tutorials page.

#### 2 Installation

`crisprDesign` can be installed from from the Bioconductor devel branch using the following commands in a fresh R session:

```
if (!require("BiocManager", quietly = TRUE))
    install.packages("BiocManager")

BiocManager::install(version="devel")
BiocManager::install("crisprDesign")
```

Users interested in contributing to `crisprDesign` might want to look at the following CRISPR-related package dependencies:

- crisprBase: core CRISPR functions and S4 objects
- crisprBowtie: aligns gRNA spacers to genomes using the ungapped aligner `bowtie`
- crisprBwa: aligns gRNA spacers to genomes using the ungapped aligner `BWA`

- crisprScore: implements state-of-the-art on- and off-target scoring algorithms
- crisprViz: gRNA visualization using genomic tracks

You can contribute to the package by submitting pull requests to our GitHub repo.

### 3 Terminology

CRISPR nucleases are examples of RNA-guided endonucleases. They require two binding components for cleavage. First, the nuclease needs to recognize a constant nucleotide motif in the target DNA called the protospacer adjacent motif (PAM) sequence. Second, the gRNA, which guides the nuclease to the target sequence, needs to bind to a complementary sequence adjacent to the PAM sequence, called the **protospacer** sequence. The latter can be thought of as a variable binding motif that can be specified by designing corresponding gRNA sequences.

The **spacer** sequence is used in the gRNA construct to guide the CRISPR nuclease to the target **protospacer** sequence in the host genome.

For DNA-targeting nucleases, the nucleotide sequence of the spacer and protospacer are identical. For RNA-targeting nucleases, they are the reverse complement of each other.

While a gRNA spacer sequence may not always uniquely target the host genome (i.e. it may map to multiple protospacers in the host genome), we can, for a given reference genome, uniquely identify a protospacer sequence with a combination of 3 attributes:

- `chr`: chromosome name
- `strand`: forward (+) or reverse (-)
- `pam_site`: genomic coordinate of the first nucleotide of the nuclease-specific PAM sequence (e.g. for SpCas9, the "N" in the NGG PAM sequence; for AsCas12a, the first "T" of the TTTV PAM sequence)

For CRISPRko, we use an additional genomic coordinate, called `cut_site`, to represent where the double-stranded break (DSB) occurs. For SpCas9, the cut site (blunt-ended dsDNA break) is located 4nt upstream of the pam_site (PAM-proximal editing). For AsCas12a, the 5nt 5' overhang dsDNA break will cause a cut 19nt after the PAM sequence on the targeted strand, and 23nt after the PAM sequence on the opposite strand (PAM-distal editing).

### 4 CRISPRko design

We will illustrate the main functionalities of `crisprDesign` by performing a common task: designing gRNAs to knock out a coding gene. In our example, we will design gRNAs for the wildtype SpCas9 nuclease, with spacers having a length of 20nt.

```
library(crisprDesign)
```

#### 4.1 Nuclease specification

The `crisprBase` package provides functionalities to create objects that store information about CRISPR nucleases, and functions to interact with those objects (see the `crisprBase` vignette). It also provides commonly-used CRISPR nucleases. Let's look at the `SpCas9` nuclease object:

```
library(crisprBase)
data(SpCas9, package="crisprBase")
SpCas9
```

```
## Class: CrisprNuclease
##   Name: SpCas9
##   Target type: DNA
##   Metadata: list of length 1
```

```
##   PAMs: NGG, NAG, NGA
##   Weights: 1, 0.2593, 0.0694
##   Spacer length: 20
##   PAM side: 3prime
##     Distance from PAM: 0
##   Prototype protospacers: 5'--SSSSSSSSSSSSSSSSSSSS[NGG]--3', 5'--SSSSSSSSSSSSSSSSSSSS[NAG]--3', 5'--S
```

The three motifs (NGG, NAG and NGA) represent the recognized PAM sequences by SpCas9, and the weights indicate a recognition score. The canonical PAM sequence NGG is fully recognized (weight of 1), while the two non-canonical PAM sequences NAG and NGA are much less tolerated.

The spacer sequence is located on the 5-prime end with respect to the PAM sequence, and the default spacer sequence length is 20 nucleotides. If necessary, we can change the spacer length using the function `crisprBase::spacerLength`. Let's see what the protospacer construct looks like by using `prototypeSequence`:

```
prototypeSequence(SpCas9)
```

```
## [1] "5'--SSSSSSSSSSSSSSSSSSSS[NGG]--3'"
```

#### 4.2 Target DNA specification

As an example, we will design gRNAs that knockout the human gene IQSEC3 by finding all protospacer sequences located in the coding region (CDS) of IQSEC3.

To do so, we need to create a `GRanges` object that defines the genomic coordinates of the CDS of IQSEC3 in a reference genome.

The toy dataset `grListExample` object in `crisprDesign` contains gene coordinates in hg38 for exons of all human IQSEC3 isoforms, and was obtained by converting an Ensembl `TxDb` object into a `GRangesList` object using the `TxDb2GRangesList` convenience function in `crisprDesign`.

```
data(grListExample, package="crisprDesign")
```

The `queryTxObject` function allows us to query such objects for a specific gene and feature. Here, we obtain a `GRanges` object containing the CDS coordinates of IQSEC3:

```
gr <- queryTxObject(txObject=grListExample,
                    featureType="cds",
                    queryColumn="gene_symbol",
                    queryValue="IQSEC3")
```

We will only consider the first exon to speed up design:

```
gr <- gr[1]
```

#### 4.3 Designing spacer sequences

`findSpacers` is the main function to obtain a list of all possible spacer sequences targeting protospacers located in the target DNA sequence(s). If a `GRanges` object is provided as input, a `BSgenome` object (object containing sequences of a reference genome) will need to be provided as well:

```
library(BSgenome.Hsapiens.UCSC.hg38)
bsgenome <- BSgenome.Hsapiens.UCSC.hg38
guideSet <- findSpacers(gr,
                        bsgenome=bsgenome,
                        crisprNuclease=SpCas9)
guideSet
```

```
## GuideSet object with 123 ranges and 5 metadata columns:
##               seqnames    ranges strand |          protospacer            pam
##                  <Rle> <IRanges>  <Rle> |       <DNAStringSet> <DNAStringSet>
##     spacer_1    chr12     66893      - | CGCGCACCGGATTCTCCAGC            AGG
##     spacer_2    chr12     66896      + | GGGCGGCATGGAGAGCCTGC            TGG
##     spacer_3    chr12     66905      + | GGAGAGCCTGCTGGAGAATC            CGG
##     spacer_4    chr12     66906      - | AGGTAGAGCACGGCGCGCAC            CGG
##     spacer_5    chr12     66916      - | GAGCTCCTTGAGGTAGAGCA            CGG
##          ...      ...       ...    ... .                  ...            ...
##   spacer_119    chr12     67407      + | CACAAATCCCCCTCCGCCCT            CGG
##   spacer_120    chr12     67412      + | ATCCCCCTCCGCCCTCGGCA            AGG
##   spacer_121    chr12     67413      + | TCCCCCTCCGCCCTCGGCAA            GGG
##   spacer_122    chr12     67421      - | CTCACTCAGGTCTCCTGCTC            AGG
##   spacer_123    chr12     67426      + | TCGGCAAGGGCGTCCTGAGC            AGG
##               pam_site  cut_site      region
##              <numeric> <numeric> <character>
##     spacer_1     66893     66896    region_1
##     spacer_2     66896     66893    region_1
##     spacer_3     66905     66902    region_1
##     spacer_4     66906     66909    region_1
##     spacer_5     66916     66919    region_1
##          ...       ...       ...         ...
##   spacer_119     67407     67404    region_1
##   spacer_120     67412     67409    region_1
##   spacer_121     67413     67410    region_1
##   spacer_122     67421     67424    region_1
##   spacer_123     67426     67423    region_1
##   -------
##   seqinfo: 640 sequences (1 circular) from hg38 genome
##   crisprNuclease: SpCas9
```

This returns a `GuideSet` object that stores genomic coordinates for all spacer sequences found in the regions provided by `gr`. The `GuideSet` object is an extension of a `GenomicRanges` object that stores additional information about gRNAs.

For the subsequent sections, we will only work with a random subset of 20 spacer sequences:

```
set.seed(10)
guideSet <- guideSet[sample(seq_along((guideSet)),20)]
```

Several accessor functions are provided to extract information about the spacer sequences:

```
spacers(guideSet)
```

```
## DNAStringSet object of length 20:
##      width seq                                              names
##  [1]    20 CCGAGTTGCTGCGCTGCTGC                             spacer_107
##  [2]    20 GCTCTGCTGGTTCTGCACGA                             spacer_9
##  [3]    20 CGGCCGCCGCGTCAGCACCA                             spacer_74
##  [4]    20 GCCCTTGCCGAGGGCGGAGG                             spacer_112
##  [5]    20 GGCCCCGCTGGGGCTGCTCC                             spacer_76
##  ...   ... ...
## [16]    20 TCCCCCTCCGCCCTCGGCAA                             spacer_121
## [17]    20 CGGCAGCGGGGCCGATGACG                             spacer_34
## [18]    20 GACGAGCCCGGGCGGAGGCT                             spacer_24
## [19]    20 CTCGTCGATACGCTCTCGCT                             spacer_13
```

```
## [20]    20 CAGTCGCCCCACAAGCATCT                               spacer_95
```

```r
protospacers(guideSet)
```

```
## DNAStringSet object of length 20:
##      width seq                                                names
##  [1]    20 CCGAGTTGCTGCGCTGCTGC                               spacer_107
##  [2]    20 GCTCTGCTGGTTCTGCACGA                               spacer_9
##  [3]    20 CGGCCGCCGCGTCAGCACCA                               spacer_74
##  [4]    20 GCCCTTGCCGAGGGCGGAGG                               spacer_112
##  [5]    20 GGCCCCGCTGGGGCTGCTCC                               spacer_76
##  ...   ... ...
## [16]    20 TCCCCCTCCGCCCTCGGCAA                               spacer_121
## [17]    20 CGGCAGCGGGGCCGATGACG                               spacer_34
## [18]    20 GACGAGCCCGGGCGGAGGCT                               spacer_24
## [19]    20 CTCGTCGATACGCTCTCGCT                               spacer_13
## [20]    20 CAGTCGCCCCACAAGCATCT                               spacer_95
```

```r
pams(guideSet)
```

```
## DNAStringSet object of length 20:
##      width seq                                                names
##  [1]     3 CGG                                                spacer_107
##  [2]     3 TGG                                                spacer_9
##  [3]     3 CGG                                                spacer_74
##  [4]     3 GGG                                                spacer_112
##  [5]     3 AGG                                                spacer_76
##  ...   ... ...
## [16]     3 GGG                                                spacer_121
## [17]     3 GGG                                                spacer_34
## [18]     3 GGG                                                spacer_24
## [19]     3 GGG                                                spacer_13
## [20]     3 GGG                                                spacer_95
```

```r
head(pamSites(guideSet))
```

```
## spacer_107   spacer_9  spacer_74 spacer_112  spacer_76  spacer_55
##      67371      66943      67233      67396      67244      67153
```

```r
head(cutSites(guideSet))
```

```
## spacer_107   spacer_9  spacer_74 spacer_112  spacer_76  spacer_55
##      67368      66946      67230      67399      67247      67156
```

The genomic locations stored in the IRanges represent the PAM site locations in the reference genome.

#### 4.4 Sequence features characterization

There are specific spacer sequence features, independent of the genomic context of the protospacer sequence, that can reduce or even eliminate gRNA activity:

- **Poly-T stretches**: four or more consecutive T nucleotides in the spacer sequence may act as a transcriptional termination signal for the U6 promoter.
- **Self-complementarity**: complementary sites with the gRNA backbone can compete with the targeted genomic sequence.
- **Percent GC**: gRNAs with GC content between 20% and 80% are preferred.

Use the function `addSequenceFeatures` to adds these spacer sequence characteristics to the `GuideSet` object:

```
guideSet <- addSequenceFeatures(guideSet)
head(guideSet)
```

```
## GuideSet object with 6 ranges and 11 metadata columns:
##              seqnames    ranges strand |          protospacer            pam
##                 <Rle> <IRanges>  <Rle> |       <DNAStringSet> <DNAStringSet>
##   spacer_107    chr12     67371      + | CCGAGTTGCTGCGCTGCTGC            CGG
##     spacer_9    chr12     66943      - | GCTCTGCTGGTTCTGCACGA            TGG
##    spacer_74    chr12     67233      + | CGGCCGCCGCGTCAGCACCA            CGG
##   spacer_112    chr12     67396      - | GCCCTTGCCGAGGGCGGAGG            GGG
##    spacer_76    chr12     67244      - | GGCCCCGCTGGGGCTGCTCC            AGG
##    spacer_55    chr12     67153      - | CTGGTCCTGGAGAGGTTCCT            GGG
##               pam_site  cut_site      region percentGC     polyA     polyC
##              <numeric> <numeric> <character> <numeric> <logical> <logical>
##   spacer_107     67371     67368    region_1        70     FALSE     FALSE
##     spacer_9     66943     66946    region_1        60     FALSE     FALSE
##    spacer_74     67233     67230    region_1        80     FALSE     FALSE
##   spacer_112     67396     67399    region_1        80     FALSE     FALSE
##    spacer_76     67244     67247    region_1        85     FALSE      TRUE
##    spacer_55     67153     67156    region_1        60     FALSE     FALSE
##                  polyG     polyT startingGGGGG
##              <logical> <logical>     <logical>
##   spacer_107     FALSE     FALSE         FALSE
##     spacer_9     FALSE     FALSE         FALSE
##    spacer_74     FALSE     FALSE         FALSE
##   spacer_112     FALSE     FALSE         FALSE
##    spacer_76      TRUE     FALSE         FALSE
##    spacer_55     FALSE     FALSE         FALSE
##   -------
##   seqinfo: 640 sequences (1 circular) from hg38 genome
##   crisprNuclease: SpCas9
```

##### 4.5 Off-target search

In order to select gRNAs that are most specific to our target of interest, it is important to avoid gRNAs that target additional loci in the genome with either perfect sequence complementarity (multiple on-targets), or imperfect complementarity through tolerated mismatches (off-targets).

For instance, both the SpCas9 and AsCas12a nucleases can be tolerant to mismatches between the gRNA spacer sequence (RNA) and the protospacer sequence (DNA), thereby making it critical to characterize off-targets to minimize the introduction of double-stranded breaks (DSBs) beyond our intended target.

The `addSpacerAlignments` function appends a list of putative on- and off-targets to a `GuideSet` object using one of three methods. The first method uses the fast aligner bowtie (Langmead et al. 2009) via the `crisprBowtie` package to map spacer sequences to a specified reference genome. This can be done by specifying `aligner="bowtie"` in `addSpacerAlignments`.

The second method uses the fast aligner BWA via the `crisprBwa` package to map spacer sequences to a specified reference genome. This can be done by specifying `aligner="bwa"` in `addSpacerAlignments`. Note that this is not available for Windows machines.

The third method uses the package `Biostrings` to search for similar sequences in a set of DNA coordinates sequences, usually provided through a `BSGenome` object. This can be done by specifying `aligner="biostrings"` in `addSpacerAlignments`. This is extremely slow, but can be useful when searching for off-targets in custom short DNA sequences.

We can control the alignment parameters and output using several function arguments. `n_mismatches` sets the maximum number of permitted gRNA:DNA mismatches (up to 3 mismatches). `n_max_alignments` specifies the maximum number of alignments for a given gRNA spacer sequence (1000 by default). The `n_max_alignments` parameter may be overruled by setting `all_Possible_alignments=TRUE`, which returns all possible alignments. `canonical=TRUE` filters out protospacer sequences that do not have a canonical PAM sequence.

Finally, the `txObject` argument in `addSpacerAlignmentsused` allows users to provide a `TxDb` object, or a `TxDb` object converted in a `GRangesList` using the `TxDb2GRangesList` function, to annotate genomic alignments with a gene model annotation. This is useful to understand whether or not off-targets are located in the CDS of another gene, for instance.

For the sake of time here, we will search only for on- and off-targets located in the beginning of human chr12 where IQSEC3 is located. We note note that users should always perform a genome-wide search as shown in the [CRISPRko design tutorial](https://github.com/crisprVerse/Tutorials/tree/master/Design_CRISPRko_Cas9].

We will use the bowtie method, with a maximum of 1 mismatch. First, we need to build a bowtie index sequence using the fasta file provided in `crisprDesign`. We use the `RBowtie` package to build the index:

```
library(Rbowtie)
fasta <- system.file(package="crisprDesign", "fasta/chr12.fa")
outdir <- tempdir()
Rbowtie::bowtie_build(fasta,
                      outdir=outdir,
                      force=TRUE,
                      prefix="chr12")
bowtie_index <- file.path(outdir, "chr12")
```

For genome-wide off-target search, users will need to create a bowtie index on the whole genome. This is explained in this tutorial.

Finally, we also need to specify a `BSgenome` object storing DNA sequences of the human reference genome:

```
library(BSgenome.Hsapiens.UCSC.hg38)
bsgenome <- BSgenome.Hsapiens.UCSC.hg38
```

We are now ready to search for on- and off-targets:

```
guideSet <- addSpacerAlignments(guideSet,
                                txObject=grListExample,
                                aligner_index=bowtie_index,
                                bsgenome=bsgenome,
                                n_mismatches=1)
```

```
## Loading required namespace: crisprBwa
```

Let's look at what was added to the `GuideSet`:

```
guideSet
```

```
## GuideSet object with 20 ranges and 16 metadata columns:
##               seqnames    ranges strand |          protospacer            pam
##                  <Rle> <IRanges>  <Rle> |       <DNAStringSet> <DNAStringSet>
##   spacer_107    chr12     67371      + | CCGAGTTGCTGCGCTGCTGC            CGG
##     spacer_9    chr12     66943      - | GCTCTGCTGGTTCTGCACGA            TGG
##    spacer_74    chr12     67233      + | CGGCCGCCGCGTCAGCACCA            CGG
##   spacer_112    chr12     67396      - | GCCCTTGCCGAGGGCGGAGG            GGG
##    spacer_76    chr12     67244      - | GGCCCCGCTGGGGCTGCTCC            AGG
```

```
##          ...      ...       ...    ... .                   ...           ...
##   spacer_121    chr12     67413      + | TCCCCCTCCGCCCTCGGCAA           GGG
##    spacer_34    chr12     67093      - | CGGCAGCGGGGCCGATGACG           GGG
##    spacer_24    chr12     67069      - | GACGAGCCCGGGCGGAGGCT           GGG
##    spacer_13    chr12     66976      - | CTCGTCGATACGCTCTCGCT           GGG
##    spacer_95    chr12     67308      + | CAGTCGCCCCACAAGCATCT           GGG
##                 pam_site  cut_site      region percentGC     polyA     polyC
##                <numeric> <numeric> <character> <numeric> <logical> <logical>
##   spacer_107      67371     67368    region_1        70     FALSE     FALSE
##     spacer_9      66943     66946    region_1        60     FALSE     FALSE
##    spacer_74      67233     67230    region_1        80     FALSE     FALSE
##   spacer_112      67396     67399    region_1        80     FALSE     FALSE
##    spacer_76      67244     67247    region_1        85     FALSE      TRUE
##          ...        ...       ...         ...       ...       ...       ...
##   spacer_121      67413     67410    region_1        75     FALSE      TRUE
##    spacer_34      67093     67096    region_1        80     FALSE     FALSE
##    spacer_24      67069     67072    region_1        80     FALSE     FALSE
##    spacer_13      66976     66979    region_1        60     FALSE     FALSE
##    spacer_95      67308     67305    region_1        60     FALSE      TRUE
##                    polyG     polyT startingGGGGG        n0        n1      n0_c
##                <logical> <logical>     <logical> <numeric> <numeric> <numeric>
##   spacer_107      FALSE     FALSE         FALSE         1         0         1
##     spacer_9      FALSE     FALSE         FALSE         1         0         1
##    spacer_74      FALSE     FALSE         FALSE         1         0         1
##   spacer_112      FALSE     FALSE         FALSE         1         0         1
##    spacer_76       TRUE     FALSE         FALSE         1         0         1
##          ...        ...       ...           ...       ...       ...       ...
##   spacer_121      FALSE     FALSE         FALSE         1         0         1
##    spacer_34       TRUE     FALSE         FALSE         1         0         1
##    spacer_24      FALSE     FALSE         FALSE         1         0         1
##    spacer_13      FALSE     FALSE         FALSE         1         0         1
##    spacer_95      FALSE     FALSE         FALSE         1         0         1
##                     n1_c    alignments
##                <numeric> <GRangesList>
##   spacer_107          0 chr12:67371:+
##     spacer_9          0 chr12:66943:-
##    spacer_74          0 chr12:67233:+
##   spacer_112          0 chr12:67396:-
##    spacer_76          0 chr12:67244:-
##          ...        ...           ...
##   spacer_121          0 chr12:67413:+
##    spacer_34          0 chr12:67093:-
##    spacer_24          0 chr12:67069:-
##    spacer_13          0 chr12:66976:-
##    spacer_95          0 chr12:67308:+
##   -------
##   seqinfo: 640 sequences (1 circular) from hg38 genome
##   crisprNuclease: SpCas9
```

A few columns were added to the `GuideSet` object to summarize the number of on- and off-targets for each spacer sequence, taking into account genomic context:

- **n0, n1, n2, n3**: specify number of alignments with 0, 1, 2 and 3 mismatches, respectively.
- **n0_c, n1_c, n2_c, n3_c**: specify number of alignments in a coding region, with 0, 1, 2 and 3 mismatches, respectively.

- **n0__p, n1__p, n2__p, n3__p**: specify number of alignments in a promoter region of a coding gene, with 0, 1, 2 and 3 mismatches, respectively.

To look at the individual on- and off-targets and their context, use the `alignments` function to retrieve a table of all genomic alignments stored in the `GuideSet` object:

```
alignments(guideSet)
```

```
## GRanges object with 20 ranges and 14 metadata columns:
##             seqnames    ranges strand |               spacer
##                <Rle> <IRanges>  <Rle> |       <DNAStringSet>
##   spacer_107    chr12     67371      + | CCGAGTTGCTGCGCTGCTGC
##     spacer_9    chr12     66943      - | GCTCTGCTGGTTCTGCACGA
##    spacer_74    chr12     67233      + | CGGCCGCCGCGTCAGCACCA
##   spacer_112    chr12     67396      - | GCCCTTGCCGAGGGCGGAGG
##    spacer_76    chr12     67244      - | GGCCCCGCTGGGGCTGCTCC
##          ...      ...       ...    ... .                  ...
##   spacer_121    chr12     67413      + | TCCCCCTCCGCCCTCGGCAA
##    spacer_34    chr12     67093      - | CGGCAGCGGGGCCGATGACG
##    spacer_24    chr12     67069      - | GACGAGCCCGGGCGGAGGCT
##    spacer_13    chr12     66976      - | CTCGTCGATACGCTCTCGCT
##    spacer_95    chr12     67308      + | CAGTCGCCCCACAAGCATCT
##                      protospacer            pam  pam_site n_mismatches
##                   <DNAStringSet> <DNAStringSet> <numeric>    <integer>
##   spacer_107 CCGAGTTGCTGCGCTGCTGC            CGG     67371            0
##     spacer_9 GCTCTGCTGGTTCTGCACGA            TGG     66943            0
##    spacer_74 CGGCCGCCGCGTCAGCACCA            CGG     67233            0
##   spacer_112 GCCCTTGCCGAGGGCGGAGG            GGG     67396            0
##    spacer_76 GGCCCCGCTGGGGCTGCTCC            AGG     67244            0
##          ...                  ...            ...       ...          ...
##   spacer_121 TCCCCCTCCGCCCTCGGCAA            GGG     67413            0
##    spacer_34 CGGCAGCGGGGCCGATGACG            GGG     67093            0
##    spacer_24 GACGAGCCCGGGCGGAGGCT            GGG     67069            0
##    spacer_13 CTCGTCGATACGCTCTCGCT            GGG     66976            0
##    spacer_95 CAGTCGCCCCACAAGCATCT            GGG     67308            0
##             canonical  cut_site         cds    fiveUTRs   threeUTRs
##             <logical> <numeric> <character> <character> <character>
##   spacer_107      TRUE     67368      IQSEC3        <NA>        <NA>
##     spacer_9      TRUE     66946      IQSEC3        <NA>        <NA>
##    spacer_74      TRUE     67230      IQSEC3        <NA>        <NA>
##   spacer_112      TRUE     67399      IQSEC3        <NA>        <NA>
##    spacer_76      TRUE     67247      IQSEC3        <NA>        <NA>
##          ...       ...       ...         ...         ...         ...
##   spacer_121      TRUE     67410      IQSEC3        <NA>        <NA>
##    spacer_34      TRUE     67096      IQSEC3        <NA>        <NA>
##    spacer_24      TRUE     67072      IQSEC3        <NA>        <NA>
##    spacer_13      TRUE     66979      IQSEC3        <NA>        <NA>
##    spacer_95      TRUE     67305      IQSEC3        <NA>        <NA>
##                   exons     introns  intergenic intergenic_distance
##             <character> <character> <character>           <integer>
##   spacer_107      IQSEC3        <NA>        <NA>                <NA>
##     spacer_9      IQSEC3        <NA>        <NA>                <NA>
##    spacer_74      IQSEC3        <NA>        <NA>                <NA>
##   spacer_112      IQSEC3        <NA>        <NA>                <NA>
##    spacer_76      IQSEC3        <NA>        <NA>                <NA>
```

```
##         ...        ...        ...        ...                ...
##   spacer_121      IQSEC3        <NA>       <NA>               <NA>
##    spacer_34      IQSEC3        <NA>       <NA>               <NA>
##    spacer_24      IQSEC3        <NA>       <NA>               <NA>
##    spacer_13      IQSEC3        <NA>       <NA>               <NA>
##    spacer_95      IQSEC3        <NA>       <NA>               <NA>
##   -------
##   seqinfo: 25 sequences (1 circular) from hg38 genome
```

The functions `onTargets` and `offTargets` will return on-target alignments (no mismatch) and off-target alignment (with at least one mismatch), respectively. See `?addSpacerAlignments` for more details about the different options.

###### 4.5.1 Iterative spacer alignments

gRNAs that align to hundreds of different locations are highly unspecific and undesirable. This can also cause `addSpacerAlignments` to be slow. To mitigate this, we provide `addSpacerAlignmentsIterative`, an iterative version of `addSpacerAlignments` that curtails alignment searches for gRNAs having more hits than the user-defined threshold (see `?addSpacerAlignmentsIterative`).

###### 4.5.2 Faster alignment by removing repeat elements

To remove protospacer sequences located in repeats or low-complexity DNA sequences (regions identified by RepeatMasker), which are usually not of interest due to their low specificity, we provide the convenience function `removeRepeats`:

```
data(grRepeatsExample, package="crisprDesign")
guideSet <- removeRepeats(guideSet,
                          gr.repeats=grRepeatsExample)
```

##### 4.6 Off-target scoring

After retrieving a list of putative off-targets and on-targets for a given spacer sequence, we can use `addOffTargetScores` to predict the likelihood of the nuclease to cut at the off-targets based on mismatch tolerance. Currently, only off-target scoring for the SpCas9 nuclease are available (MIT and CFD algorithms):

```
guideSet <- addOffTargetScores(guideSet)
guideSet
```

```
## GuideSet object with 17 ranges and 19 metadata columns:
##              seqnames    ranges strand |          protospacer           pam
##                 <Rle> <IRanges>  <Rle> |       <DNAStringSet> <DNAStringSet>
##   spacer_107    chr12     67371      + | CCGAGTTGCTGCGCTGCTGC           CGG
##     spacer_9    chr12     66943      - | GCTCTGCTGGTTCTGCACGA           TGG
##    spacer_74    chr12     67233      + | CGGCCGCCGCGTCAGCACCA           CGG
##   spacer_112    chr12     67396      - | GCCCTTGCCGAGGGCGGAGG           GGG
##    spacer_76    chr12     67244      - | GGCCCCGCTGGGGCTGCTCC           AGG
##          ...      ...       ...    ... .                  ...           ...
##    spacer_71    chr12     67218      - | TGTCCGTGGTGCTGACGCGG           CGG
##   spacer_121    chr12     67413      + | TCCCCCTCCGCCCTCGGCAA           GGG
##    spacer_24    chr12     67069      - | GACGAGCCCGGGCGGAGGCT           GGG
##    spacer_13    chr12     66976      - | CTCGTCGATACGCTCTCGCT           GGG
##    spacer_95    chr12     67308      + | CAGTCGCCCCACAAGCATCT           GGG
##               pam_site  cut_site      region percentGC     polyA     polyC
##              <numeric> <numeric> <character> <numeric> <logical> <logical>
##   spacer_107     67371     67368    region_1        70     FALSE     FALSE
```

```
##     spacer_9     66943     66946    region_1        60     FALSE     FALSE
##    spacer_74     67233     67230    region_1        80     FALSE     FALSE
##   spacer_112     67396     67399    region_1        80     FALSE     FALSE
##    spacer_76     67244     67247    region_1        85     FALSE      TRUE
##          ...       ...       ...         ...       ...       ...       ...
##    spacer_71     67218     67221    region_1        70     FALSE     FALSE
##   spacer_121     67413     67410    region_1        75     FALSE      TRUE
##    spacer_24     67069     67072    region_1        80     FALSE     FALSE
##    spacer_13     66976     66979    region_1        60     FALSE     FALSE
##    spacer_95     67308     67305    region_1        60     FALSE      TRUE
##                  polyG     polyT startingGGGGG        n0        n1      n0_c
##              <logical> <logical>     <logical> <numeric> <numeric> <numeric>
##   spacer_107     FALSE     FALSE         FALSE         1         0         1
##     spacer_9     FALSE     FALSE         FALSE         1         0         1
##    spacer_74     FALSE     FALSE         FALSE         1         0         1
##   spacer_112     FALSE     FALSE         FALSE         1         0         1
##    spacer_76      TRUE     FALSE         FALSE         1         0         1
##          ...       ...       ...           ...       ...       ...       ...
##    spacer_71     FALSE     FALSE         FALSE         1         0         1
##   spacer_121     FALSE     FALSE         FALSE         1         0         1
##    spacer_24     FALSE     FALSE         FALSE         1         0         1
##    spacer_13     FALSE     FALSE         FALSE         1         0         1
##    spacer_95     FALSE     FALSE         FALSE         1         0         1
##                   n1_c    alignments inRepeats score_cfd score_mit
##              <numeric> <GRangesList> <logical> <numeric> <numeric>
##   spacer_107         0 chr12:67371:+     FALSE         1         1
##     spacer_9         0 chr12:66943:-     FALSE         1         1
##    spacer_74         0 chr12:67233:+     FALSE         1         1
##   spacer_112         0 chr12:67396:-     FALSE         1         1
##    spacer_76         0 chr12:67244:-     FALSE         1         1
##          ...       ...           ...       ...       ...       ...
##    spacer_71         0 chr12:67218:-     FALSE         1         1
##   spacer_121         0 chr12:67413:+     FALSE         1         1
##    spacer_24         0 chr12:67069:-     FALSE         1         1
##    spacer_13         0 chr12:66976:-     FALSE         1         1
##    spacer_95         0 chr12:67308:+     FALSE         1         1
##   -------
##   seqinfo: 640 sequences (1 circular) from hg38 genome
##   crisprNuclease: SpCas9
```

Note that this will only work after calling `addSpacerAlignments`, as it requires a list of off-targets for each gRNA entry.

##### 4.7 On-target scoring

`addOnTargetScores` adds scores from all on-target efficiency algorithms available in the R package `crisprScore` and appends them to the `GuideSet`. By default, scores for all available methods for a given nuclease will be computed. Here, for the sake of time, let's add only the CRISPRater score:

```
guideSet <- addOnTargetScores(guideSet, methods="crisprater")
head(guideSet)
```

```
## GuideSet object with 6 ranges and 20 metadata columns:
##              seqnames    ranges strand |        protospacer            pam
##                 <Rle> <IRanges>  <Rle> |     <DNAStringSet> <DNAStringSet>
```

```
##   spacer_107    chr12     67371      + | CCGAGTTGCTGCGCTGCTGC           CGG
##     spacer_9    chr12     66943      - | GCTCTGCTGGTTCTGCACGA           TGG
##    spacer_74    chr12     67233      + | CGGCCGCCGCGTCAGCACCA           CGG
##   spacer_112    chr12     67396      - | GCCCTTGCCGAGGGCGGAGG           GGG
##    spacer_76    chr12     67244      - | GGCCCCGCTGGGGCTGCTCC           AGG
##    spacer_55    chr12     67153      - | CTGGTCCTGGAGAGGTTCCT           GGG
##                pam_site  cut_site      region percentGC     polyA     polyC
##               <numeric> <numeric> <character> <numeric> <logical> <logical>
##   spacer_107     67371     67368    region_1        70     FALSE     FALSE
##     spacer_9     66943     66946    region_1        60     FALSE     FALSE
##    spacer_74     67233     67230    region_1        80     FALSE     FALSE
##   spacer_112     67396     67399    region_1        80     FALSE     FALSE
##    spacer_76     67244     67247    region_1        85     FALSE      TRUE
##    spacer_55     67153     67156    region_1        60     FALSE     FALSE
##                   polyG     polyT startingGGGGG        n0        n1      n0_c
##               <logical> <logical>     <logical> <numeric> <numeric> <numeric>
##   spacer_107     FALSE     FALSE         FALSE         1         0         1
##     spacer_9     FALSE     FALSE         FALSE         1         0         1
##    spacer_74     FALSE     FALSE         FALSE         1         0         1
##   spacer_112     FALSE     FALSE         FALSE         1         0         1
##    spacer_76      TRUE     FALSE         FALSE         1         0         1
##    spacer_55     FALSE     FALSE         FALSE         1         0         1
##                    n1_c    alignments inRepeats score_cfd score_mit
##               <numeric> <GRangesList> <logical> <numeric> <numeric>
##   spacer_107         0 chr12:67371:+     FALSE         1         1
##     spacer_9         0 chr12:66943:-     FALSE         1         1
##    spacer_74         0 chr12:67233:+     FALSE         1         1
##   spacer_112         0 chr12:67396:-     FALSE         1         1
##    spacer_76         0 chr12:67244:-     FALSE         1         1
##    spacer_55         0 chr12:67153:-     FALSE         1         1
##               score_crisprater
##                      <numeric>
##   spacer_107          0.782780
##     spacer_9          0.834319
##    spacer_74          0.764870
##   spacer_112          0.795745
##    spacer_76          0.755493
##    spacer_55          0.711902
##   -------
##   seqinfo: 640 sequences (1 circular) from hg38 genome
##   crisprNuclease: SpCas9
```

See the `crisprScore` vignette for a full description of the different scores.

#### 4.8 Restriction enzymes

Restriction enzymes are usually involved in the gRNA library synthesis process. Removing gRNAs that contain specific restriction sites is often necessary. We provide the function `addRestrictionEnzymes` to indicate whether or not gRNAs contain restriction sites for a user-defined set of enzymes:

```
guideSet <- addRestrictionEnzymes(guideSet)
```

When no enzymes are specified, the function adds annotation for the following default enzymes: EcoRI, KpnI, BsmBI, BsaI, BbsI, PacI, ISceI and MluI. The function also has two additional arguments, `flanking5` and `flanking3`, to specify nucleotide sequences flanking the spacer sequence (5' and 3', respectively) in the lentiviral cassette that will be used for gRNA delivery. The function will effectively search for restriction sites in the full sequence `[flanking5][spacer][flanking3]`.

The `enzymeAnnotation` function can be used to retrieve the added annotation:

```
head(enzymeAnnotation(guideSet))
```

```
## DataFrame with 6 rows and 7 columns
##                 EcoRI      KpnI     BsmBI      BsaI      BbsI      PacI
##             <logical> <logical> <logical> <logical> <logical> <logical>
## spacer_107      FALSE     FALSE     FALSE     FALSE     FALSE     FALSE
## spacer_9        FALSE     FALSE     FALSE     FALSE     FALSE     FALSE
## spacer_74       FALSE     FALSE     FALSE     FALSE     FALSE     FALSE
## spacer_112      FALSE     FALSE     FALSE     FALSE     FALSE     FALSE
## spacer_76       FALSE     FALSE     FALSE     FALSE     FALSE     FALSE
## spacer_55       FALSE     FALSE     FALSE     FALSE     FALSE     FALSE
##                  MluI
##             <logical>
## spacer_107      FALSE
## spacer_9        FALSE
## spacer_74       FALSE
## spacer_112      FALSE
## spacer_76       FALSE
## spacer_55       FALSE
```

#### 4.9 Gene annotation

The function `addGeneAnnotation` adds transcript- and gene-level contextual information to gRNAs from a `TxDb`-like object:

```
guideSet <- addGeneAnnotation(guideSet,
                              txObject=grListExample)
```

The gene annotation can be retrieved using the function `geneAnnotation`:

```
geneAnnotation(guideSet)
```

```
## DataFrame with 17 rows and 23 columns
##                  chr anchor_site   strand gene_symbol         gene_id
##             <factor>   <integer> <factor> <character>     <character>
## spacer_107     chr12       67368        +      IQSEC3 ENSG00000120645
## spacer_9       chr12       66946        -      IQSEC3 ENSG00000120645
## spacer_74      chr12       67230        +      IQSEC3 ENSG00000120645
## spacer_112     chr12       67399        -      IQSEC3 ENSG00000120645
## spacer_76      chr12       67247        -      IQSEC3 ENSG00000120645
## ...              ...         ...      ...         ...             ...
## spacer_71      chr12       67221        -      IQSEC3 ENSG00000120645
## spacer_121     chr12       67410        +      IQSEC3 ENSG00000120645
## spacer_24      chr12       67072        -      IQSEC3 ENSG00000120645
## spacer_13      chr12       66979        -      IQSEC3 ENSG00000120645
## spacer_95      chr12       67305        +      IQSEC3 ENSG00000120645
##                      tx_id      protein_id   cut_cds cut_fiveUTRs cut_threeUTRs
##                <character>     <character> <logical>    <logical>     <logical>
## spacer_107 ENST00000538872 ENSP00000437554      TRUE        FALSE         FALSE
## spacer_9   ENST00000538872 ENSP00000437554      TRUE        FALSE         FALSE
## spacer_74  ENST00000538872 ENSP00000437554      TRUE        FALSE         FALSE
## spacer_112 ENST00000538872 ENSP00000437554      TRUE        FALSE         FALSE
```

```
## spacer_76  ENST00000538872 ENSP00000437554      TRUE        FALSE         FALSE
## ...                    ...             ...       ...          ...           ...
## spacer_71  ENST00000538872 ENSP00000437554      TRUE        FALSE         FALSE
## spacer_121 ENST00000538872 ENSP00000437554      TRUE        FALSE         FALSE
## spacer_24  ENST00000538872 ENSP00000437554      TRUE        FALSE         FALSE
## spacer_13  ENST00000538872 ENSP00000437554      TRUE        FALSE         FALSE
## spacer_95  ENST00000538872 ENSP00000437554      TRUE        FALSE         FALSE
##            cut_introns percentCDS aminoAcidIndex downtreamATG percentTx
##              <logical>  <numeric>      <numeric>    <numeric> <numeric>
## spacer_107       FALSE       13.7            162            1       8.5
## spacer_9         FALSE        1.8             22            0       2.5
## spacer_74        FALSE        9.8            116            0       6.5
## spacer_112       FALSE       14.6            173            1       8.9
## spacer_76        FALSE       10.3            122            0       6.8
## ...                ...        ...            ...          ...       ...
## spacer_71        FALSE        9.6            113            0       6.4
## spacer_121       FALSE       14.9            176            1       9.1
## spacer_24        FALSE        5.4             64            0       4.3
## spacer_13        FALSE        2.7             33            0       3.0
## spacer_95        FALSE       11.9            141            1       7.6
##            nIsoforms totalIsoforms percentIsoforms isCommonExon nCodingIsoforms
##            <integer>     <numeric>       <numeric>    <logical>       <integer>
## spacer_107         1             2              50        FALSE               1
## spacer_9           1             2              50        FALSE               1
## spacer_74          1             2              50        FALSE               1
## spacer_112         1             2              50        FALSE               1
## spacer_76          1             2              50        FALSE               1
## ...              ...           ...             ...          ...             ...
## spacer_71          1             2              50        FALSE               1
## spacer_121         1             2              50        FALSE               1
## spacer_24          1             2              50        FALSE               1
## spacer_13          1             2              50        FALSE               1
## spacer_95          1             2              50        FALSE               1
##            totalCodingIsoforms percentCodingIsoforms isCommonCodingExon
##                      <numeric>             <numeric>          <logical>
## spacer_107                   2                    50              FALSE
## spacer_9                     2                    50              FALSE
## spacer_74                    2                    50              FALSE
## spacer_112                   2                    50              FALSE
## spacer_76                    2                    50              FALSE
## ...                        ...                   ...                ...
## spacer_71                    2                    50              FALSE
## spacer_121                   2                    50              FALSE
## spacer_24                    2                    50              FALSE
## spacer_13                    2                    50              FALSE
## spacer_95                    2                    50              FALSE
```

It contains a lot of information that contextualizes the genomic location of the protospacer sequences.

The ID columns (`tx_id`, `gene_id`, `protein_id`, `exon_id`) give Ensembl IDs. The `exon_rank` gives the order of the exon for the transcript, for example "2" indicates it is the second exon (from the 5' end) in the mature transcript.

The columns `cut_cds`, `cut_fiveUTRs`, `cut_threeUTRs` and `cut_introns` indicate whether the guide sequence overlaps with CDS, 5' UTR, 3' UTR, or an intron, respectively.

`percentCDS` gives the location of the `cut_site` within the transcript as a percent from the 5' end to the 3' end. `aminoAcidIndex` gives the number of the specific amino acid in the protein where the cut is predicted to occur. `downstreamATG` shows how many in-frame ATGs are downstream of the `cut_site` (and upstream from the defined percent transcript cutoff, `met_cutoff`), indicating a potential alternative translation initiation site that may preserve protein function.

For more information about the other columns, type `?addGeneAnnotation`.

#### 4.10 TSS annotation

Similarly, one might want to know which protospacer sequences are located within promoter regions of known genes:

```
data(tssObjectExample, package="crisprDesign")
guideSet <- addTssAnnotation(guideSet,
                             tssObject=tssObjectExample)
tssAnnotation(guideSet)
```

```
## DataFrame with 10 rows and 11 columns
##                 chr anchor_site   strand           tx_id         gene_id
##            <factor>   <integer> <factor>     <character>     <character>
## spacer_9      chr12       66946        - ENST00000538872 ENSG00000120645
## spacer_74     chr12       67230        + ENST00000538872 ENSG00000120645
## spacer_76     chr12       67247        - ENST00000538872 ENSG00000120645
## spacer_55     chr12       67156        - ENST00000538872 ENSG00000120645
## spacer_72     chr12       67224        - ENST00000538872 ENSG00000120645
## spacer_54     chr12       67145        + ENST00000538872 ENSG00000120645
## spacer_15     chr12       66995        + ENST00000538872 ENSG00000120645
## spacer_71     chr12       67221        - ENST00000538872 ENSG00000120645
## spacer_24     chr12       67072        - ENST00000538872 ENSG00000120645
## spacer_13     chr12       66979        - ENST00000538872 ENSG00000120645
##            gene_symbol    promoter      tss_id  tss_strand   tss_pos dist_to_tss
##            <character> <character> <character> <character> <integer>   <numeric>
## spacer_9        IQSEC3          P1   IQSEC3_P1           +     66767         179
## spacer_74       IQSEC3          P1   IQSEC3_P1           +     66767         463
## spacer_76       IQSEC3          P1   IQSEC3_P1           +     66767         480
## spacer_55       IQSEC3          P1   IQSEC3_P1           +     66767         389
## spacer_72       IQSEC3          P1   IQSEC3_P1           +     66767         457
## spacer_54       IQSEC3          P1   IQSEC3_P1           +     66767         378
## spacer_15       IQSEC3          P1   IQSEC3_P1           +     66767         228
## spacer_71       IQSEC3          P1   IQSEC3_P1           +     66767         454
## spacer_24       IQSEC3          P1   IQSEC3_P1           +     66767         305
## spacer_13       IQSEC3          P1   IQSEC3_P1           +     66767         212
```

For more information, type `?addTssAnnotation`.

#### 4.11 SNP information

Common single-nucleotide polymorphisms (SNPs) can change the on-target and off-target properties of gRNAs by altering the binding. The function `addSNPAnnotation` annotates gRNAs with respect to a reference database of SNPs (stored in a VCF file), specified by the `vcf` argument.

VCF files for common SNPs (dbSNPs) can be downloaded from NCBI on the dbSNP website. We include in this package an example VCF file for common SNPs located in the proximity of human gene IQSEC3. This was obtained using the dbSNP151 RefSNP database obtained by subsetting around IQSEC.

```r
vcf <- system.file("extdata",
                   file="common_snps_dbsnp151_example.vcf.gz",
                   package="crisprDesign")
guideSet <- addSNPAnnotation(guideSet, vcf=vcf)
snps(guideSet)
```

```
## DataFrame with 0 rows and 9 columns
```

The `rs_site_rel` gives the relative position of the SNP with respect to the `pam_site`. `allele_ref` and `allele_minor` report the nucleotide of the reference and minor alleles, respectively. `MAF_1000G` and `MAF_TOPMED` report the minor allele frequency (MAF) in the 1000Genomes and TOPMED populations.

#### 4.12 Filtering and ranking gRNAs

Once gRNAs are fully annotated, it is easy to filter out any unwanted gRNAs since `GuideSet` objects can be subsetted like regular vectors in R.

As an example, suppose that we only want to keep gRNAs that have percent GC between 20% and 80% and that do not contain a polyT stretch. This can be achieved using the following lines:

```r
guideSet <- guideSet[guideSet$percentGC>=20]
guideSet <- guideSet[guideSet$percentGC<=80]
guideSet <- guideSet[!guideSet$polyT]
```

Similarly, it is easy to rank gRNAs based on a set of criteria using the regular `order` function.

For instance, let's sort gRNAs by the CRISPRater on-target score:

```r
# Creating an ordering index based on the CRISPRater score:
# Using the negative values to make sure higher scores are ranked first:
o <- order(-guideSet$score_crisprater)
# Ordering the GuideSet:
guideSet <- guideSet[o]
head(guideSet)
```

```
## GuideSet object with 6 ranges and 25 metadata columns:
##               seqnames    ranges strand |          protospacer           pam
##                  <Rle> <IRanges>  <Rle> |       <DNAStringSet> <DNAStringSet>
##     spacer_9     chr12     66943      - | GCTCTGCTGGTTCTGCACGA           TGG
##   spacer_112     chr12     67396      - | GCCCTTGCCGAGGGCGGAGG           GGG
##   spacer_107     chr12     67371      + | CCGAGTTGCTGCGCTGCTGC           CGG
##    spacer_74     chr12     67233      + | CGGCCGCCGCGTCAGCACCA           CGG
##    spacer_76     chr12     67244      - | GGCCCCGCTGGGGCTGCTCC           AGG
##   spacer_121     chr12     67413      + | TCCCCCTCCGCCCTCGGCAA           GGG
##                pam_site  cut_site      region percentGC     polyA     polyC
##               <numeric> <numeric> <character> <numeric> <logical> <logical>
##     spacer_9      66943     66946    region_1        60     FALSE     FALSE
##   spacer_112      67396     67399    region_1        80     FALSE     FALSE
##   spacer_107      67371     67368    region_1        70     FALSE     FALSE
##    spacer_74      67233     67230    region_1        80     FALSE     FALSE
##    spacer_76      67244     67247    region_1        85     FALSE      TRUE
##   spacer_121      67413     67410    region_1        75     FALSE      TRUE
##                   polyG     polyT startingGGGGG        n0        n1      n0_c
##               <logical> <logical>     <logical> <numeric> <numeric> <numeric>
##     spacer_9      FALSE     FALSE         FALSE         1         0         1
##   spacer_112      FALSE     FALSE         FALSE         1         0         1
##   spacer_107      FALSE     FALSE         FALSE         1         0         1
```

```
##    spacer_74     FALSE     FALSE         FALSE         1         0         1
##    spacer_76      TRUE     FALSE         FALSE         1         0         1
##   spacer_121     FALSE     FALSE         FALSE         1         0         1
##                    n1_c    alignments inRepeats score_cfd score_mit
##               <numeric> <GRangesList> <logical> <numeric> <numeric>
##     spacer_9          0 chr12:66943:-     FALSE         1         1
##   spacer_112          0 chr12:67396:-     FALSE         1         1
##   spacer_107          0 chr12:67371:+     FALSE         1         1
##    spacer_74          0 chr12:67233:+     FALSE         1         1
##    spacer_76          0 chr12:67244:-     FALSE         1         1
##   spacer_121          0 chr12:67413:+     FALSE         1         1
##               score_crisprater      enzymeAnnotation       geneAnnotation
##                      <numeric>  <SplitDataFrameList> <SplitDataFrameList>
##     spacer_9          0.834319 FALSE:FALSE:FALSE:...    chr12:66946:-:...
##   spacer_112          0.795745 FALSE:FALSE:FALSE:...    chr12:67399:-:...
##   spacer_107          0.782780 FALSE:FALSE:FALSE:...    chr12:67368:+:...
##    spacer_74          0.764870 FALSE:FALSE:FALSE:...    chr12:67230:+:...
##    spacer_76          0.755493 FALSE:FALSE:FALSE:...    chr12:67247:-:...
##   spacer_121          0.741315 FALSE:FALSE:FALSE:...    chr12:67410:+:...
##                      tssAnnotation    hasSNP                 snps
##               <SplitDataFrameList> <logical> <SplitDataFrameList>
##     spacer_9     chr12:66946:-:...     FALSE              :...,...
##   spacer_112              :...,...     FALSE              :...,...
##   spacer_107              :...,...     FALSE              :...,...
##    spacer_74     chr12:67230:+:...     FALSE              :...,...
##    spacer_76     chr12:67247:-:...     FALSE              :...,...
##   spacer_121              :...,...     FALSE              :...,...
##   -------
##   seqinfo: 640 sequences (1 circular) from hg38 genome
##   crisprNuclease: SpCas9
```

One can also sort gRNAs using several annotation columns. For instance, let's sort gRNAs using the CRISPRrater score, but also by prioritizing first gRNAs that have no 1-mismatch off-targets:

```
o <- order(guideSet$n1, -guideSet$score_crisprater)
# Ordering the GuideSet:
guideSet <- guideSet[o]
head(guideSet)
```

```
## GuideSet object with 6 ranges and 25 metadata columns:
##               seqnames    ranges strand |          protospacer            pam
##                  <Rle> <IRanges>  <Rle> |       <DNAStringSet> <DNAStringSet>
##     spacer_9     chr12     66943      - | GCTCTGCTGGTTCTGCACGA            TGG
##   spacer_112     chr12     67396      - | GCCCTTGCCGAGGGCGGAGG            GGG
##   spacer_107     chr12     67371      + | CCGAGTTGCTGCGCTGCTGC            CGG
##    spacer_74     chr12     67233      + | CGGCCGCCGCGTCAGCACCA            CGG
##    spacer_76     chr12     67244      - | GGCCCCGCTGGGGCTGCTCC            AGG
##   spacer_121     chr12     67413      + | TCCCCCTCCGCCCTCGGCAA            GGG
##                 pam_site  cut_site      region percentGC     polyA     polyC
##                <numeric> <numeric> <character> <numeric> <logical> <logical>
##     spacer_9       66943     66946    region_1        60     FALSE     FALSE
##   spacer_112       67396     67399    region_1        80     FALSE     FALSE
##   spacer_107       67371     67368    region_1        70     FALSE     FALSE
##    spacer_74       67233     67230    region_1        80     FALSE     FALSE
##    spacer_76       67244     67247    region_1        85     FALSE      TRUE
```

```
##   spacer_121     67413     67410    region_1        75     FALSE      TRUE
##                  polyG     polyT startingGGGGG        n0        n1      n0_c
##              <logical> <logical>     <logical> <numeric> <numeric> <numeric>
##     spacer_9     FALSE     FALSE         FALSE         1         0         1
##   spacer_112     FALSE     FALSE         FALSE         1         0         1
##   spacer_107     FALSE     FALSE         FALSE         1         0         1
##    spacer_74     FALSE     FALSE         FALSE         1         0         1
##    spacer_76      TRUE     FALSE         FALSE         1         0         1
##   spacer_121     FALSE     FALSE         FALSE         1         0         1
##                   n1_c    alignments inRepeats score_cfd score_mit
##              <numeric> <GRangesList> <logical> <numeric> <numeric>
##     spacer_9         0 chr12:66943:-     FALSE         1         1
##   spacer_112         0 chr12:67396:-     FALSE         1         1
##   spacer_107         0 chr12:67371:+     FALSE         1         1
##    spacer_74         0 chr12:67233:+     FALSE         1         1
##    spacer_76         0 chr12:67244:-     FALSE         1         1
##   spacer_121         0 chr12:67413:+     FALSE         1         1
##              score_crisprater      enzymeAnnotation       geneAnnotation
##                     <numeric>  <SplitDataFrameList> <SplitDataFrameList>
##     spacer_9         0.834319 FALSE:FALSE:FALSE:...    chr12:66946:-:...
##   spacer_112         0.795745 FALSE:FALSE:FALSE:...    chr12:67399:-:...
##   spacer_107         0.782780 FALSE:FALSE:FALSE:...    chr12:67368:+:...
##    spacer_74         0.764870 FALSE:FALSE:FALSE:...    chr12:67230:+:...
##    spacer_76         0.755493 FALSE:FALSE:FALSE:...    chr12:67247:-:...
##   spacer_121         0.741315 FALSE:FALSE:FALSE:...    chr12:67410:+:...
##                     tssAnnotation    hasSNP                 snps
##              <SplitDataFrameList> <logical> <SplitDataFrameList>
##     spacer_9    chr12:66946:-:...     FALSE              :...,...
##   spacer_112             :...,...     FALSE              :...,...
##   spacer_107             :...,...     FALSE              :...,...
##    spacer_74    chr12:67230:+:...     FALSE              :...,...
##    spacer_76    chr12:67247:-:...     FALSE              :...,...
##   spacer_121             :...,...     FALSE              :...,...
##   -------
##   seqinfo: 640 sequences (1 circular) from hg38 genome
##   crisprNuclease: SpCas9
```

The `rankSpacers` function is a convenience function that implements our recommended rankings for the SpCas9, enAsCas12a and CasRx nucleases. For a detailed description of our recommended rankings, see the documentation of `rankSpacers` by typing `?rankSpacers`.

If an Ensembl transcript ID is provided, the ranking function will also take into account the position of the gRNA within the target CDS of the transcript ID in the ranking procedure. Our recommendation is to specify the Ensembl canonical transcript as the representative transcript for the gene. In our example, ENST00000538872 is the canonical transcript for IQSEC3:

```
tx_id <- "ENST00000538872"
guideSet <- rankSpacers(guideSet,
                        tx_id=tx_id)
```

### 5 CRISPRa/CRISPRi design

For CRISPRa and CRISPRi applications, the CRISPR nuclease is engineered to lose its endonuclease activity, therefore should not introduce double-stranded breaks (DSBs). We will use the dead SpCas9 (dSpCas9) nuclease as an example here. Note that users don't have to distinguish between dSpCas9 and SpCas9 when specifying the nuclease in `crisprDesign` and `crisprBase` as they do not differ in terms of the characteristics stored in the `CrisprNuclease` object.

*CRISPRi*: Fusing dSpCas9 with a Krüppel-associated box (KRAB) domain has been shown to be effective at repressing transcription in mammalian cells (Gilbert et al. 2013). The dSpCas9-KRAB fused protein is a commonly-used construct to conduct CRISPR inhibition (CRISPRi) experiments. To achieve optimal inhibition, gRNAs are usually designed targeting the region directly downstream of the gene transcription starting site (TSS).

*CRISPRa*: dSpCas9 can also be used to activate gene expression by coupling the dead nuclease with activation factors. The technology is termed CRISPR activation (CRISPRa), and several CRISPRa systems have been developed (see Kampmann (2018) for a review). For optimal activation, gRNAs are usually designed to target the region directly upstream of the gene TSS.

`crisprDesign` provides functionalities to be able to take into account design rules that are specific to CRISPRa and CRISPRi applications. The `queryTss` function allows to specify genomic coordinates of promoter regions. The `addTssAnnotation` annotates gRNAs for known TSSs, and includes a column named `dist_to_tss` that indicates the distance between the TSS position and the PAM site of the gRNA. For CRISPRi, we recommend targeting the 25-75bp region downstream of the TSS for optimal inhibition. For CRISPRa, we recommend targeting the region 75-150bp upstream of the TSS for optimal activation; see (Sanson et al. 2018) for more information.

For more information, please see the following two tutorials:

- CRISPR activation (CRISPRa) design
- CRISPR interference (CRISPRi) design

### 6 CRISPR base editing with BE4max

We illustrate the CRISPR base editing (CRISPRbe) functionalities of `crisprDesign` by designing and characterizing gRNAs targeting IQSEC3 using the cytidine base editor BE4max (Koblan et al. 2018).

We first load the BE4max `BaseEditor` object available in `crisprBase`:

```
data(BE4max, package="crisprBase")
BE4max
```

```
## Class: BaseEditor
##   CRISPR Nuclease name: SpCas9
##       Target type: DNA
##       Metadata: list of length 2
##       PAMs: NGG, NAG, NGA
##       Weights: 1, 0.2593, 0.0694
##       Spacer length: 20
##       PAM side: 3prime
##         Distance from PAM: 0
##       Prototype protospacers: 5'--SSSSSSSSSSSSSSSSSSSS[NGG]--3', 5'--SSSSSSSSSSSSSSSSSSSS[NAG]--3', 5
##   Base editor name: BE4max
##       Editing strand: original
##       Maximum editing weight: C2T at position -15
```

The editing probabilities of the base editor BE4max are stored in a matrix where rows correspond to the different nucleotide substitutions, and columns correspond to the genomic coordinate relative to the PAM site. The `editingWeights` function from `crisprBase` allows to retrieve those probabilities. One can see that C to T editing is optimal around 15 nucleotides upstream of the PAM site for the BE4max base editor:

```
crisprBase::editingWeights(BE4max)["C2T",]
```

```
##   -36   -35   -34   -33   -32   -31   -30   -29   -28   -27   -26   -25   -24
## 0.007 0.007 0.008 0.018 0.010 0.020 0.014 0.012 0.023 0.013 0.024 0.022 0.034
##   -23   -22   -21   -20   -19   -18   -17   -16   -15   -14   -13   -12   -11
## 0.022 0.021 0.035 0.058 0.162 0.318 0.632 0.903 1.000 0.870 0.620 0.314 0.163
##   -10    -9    -8    -7    -6    -5    -4    -3    -2    -1
## 0.100 0.056 0.033 0.019 0.018 0.024 0.017 0.005 0.002 0.001
```

We obtain a `GuideSet` object using the first exon of the IQSEC3 gene and retain only the first 2 gRNAs for the sake of time:

```
gr <- queryTxObject(txObject=grListExample,
                    featureType="cds",
                    queryColumn="gene_symbol",
                    queryValue="IQSEC3")
gs <- findSpacers(gr[1],
                  bsgenome=bsgenome,
                  crisprNuclease=BE4max)
gs <- gs[1:2]
```

The function `addEditedAlleles` finds, characterizes, and scores predicted edited alleles for each gRNA, for a chosen transcript. It requires a transcript-specific annotation that can be obtained using the function `getTxInfoDataFrame`. Here, we will perform the analysis using the main isoform of IQSEC3 (transcript id ENST00000538872).

We first get the transcript table for ENST00000538872,

```
txid <- "ENST00000538872"
txTable <- getTxInfoDataFrame(tx_id=txid,
                              txObject=grListExample,
                              bsgenome=bsgenome)
head(txTable)
```

```
## DataFrame with 6 rows and 10 columns
##           chr       pos         nuc          aa aa_number      exon  pos_plot
##   <character> <numeric> <character> <character> <integer> <integer> <integer>
## 1       chr12     66767           A          NA        NA         1        31
## 2       chr12     66768           G          NA        NA         1        32
## 3       chr12     66769           G          NA        NA         1        33
## 4       chr12     66770           C          NA        NA         1        34
## 5       chr12     66771           T          NA        NA         1        35
## 6       chr12     66772           G          NA        NA         1        36
##    pos_mrna   pos_cds      region
##   <integer> <integer> <character>
## 1         1        NA        5UTR
## 2         2        NA        5UTR
## 3         3        NA        5UTR
## 4         4        NA        5UTR
## 5         5        NA        5UTR
## 6         6        NA        5UTR
```

and then add the edited alleles annotation to the `GuideSet`:

```
editingWindow <- c(-20,-8)
gs <- addEditedAlleles(gs,
                       baseEditor=BE4max,
                       txTable=txTable,
                       editingWindow=editingWindow)
```

```
## [addEditedAlleles] Obtaining edited alleles at each gRNA target site.
```

```
## [addEditedAlleles] Adding functional consequences to alleles.
```

The `editingWindow` argument specifies the window of editing that we are interested in. When not provided, it uses the default window provided in the `BaseEditor` object. Note that providing large windows can exponentially increase computing time as the number of possible alleles grows exponentially.Let's retrieve the edited alleles for the first gRNA:

```
alleles <- editedAlleles(gs)[[1]]
```

It is a `DataFrame` object that contains useful metadata information:

```
metadata(alleles)
```

```
## $wildtypeAllele
##        spacer_1
## "CGCGCACCGGATT"
##
## $start
## [1] 66901
##
## $end
## [1] 66913
##
## $chr
## [1] "chr12"
##
## $strand
## [1] "-"
##
## $editingWindow
## [1] -20  -8
##
## $wildtypeAmino
## [1] "NNNPPPVVVRRRA"
```

The `wildtypeAllele` reports the unedited nucleotide sequence of the region specified by the editing window (with respect to the gRNA PAM site). It is always reported from the 5' to 3' direction on the strand corresponding to the gRNA strand. The `start` and `end` specify the corresponding coordinates on the transcript.

Let's look at the edited alleles:

```
head(alleles)
```

```
## DataFrame with 6 rows and 4 columns
##              seq     score     variant            aa
##   <DNAStringSet> <numeric> <character>   <character>
## 1  CGCGTATTGGATT 0.2471509    missense NNNPPPIIIRRRA
## 2  CGCGTATCGGATT 0.1618439    missense NNNPPPIIIRRRA
## 3  CGTGTATTGGATT 0.1057792    missense NNNPPPIIIHHHA
## 4  CGTGTATCGGATT 0.0692683    missense NNNPPPIIIHHHA
## 5  CGCGTACTGGATT 0.0372147      silent NNNPPPVVVRRRA
## 6  CGCGCATTGGATT 0.0292859    missense NNNPPPMMMRRRA
```

The `DataFrame` is ordered so that the top predicted alleles (based on the `score` column) are shown first. The `score` represents the likelihood of the edited allele to occur relative to all possible edited alleles, and is calculated using the editing weights stored in the `BE4max` object. The `seq` column represents the edited nucleotide sequences. Similar to the `wildtypeAllele` above, they are always reported from the 5' to 3' direction on the strand corresponding to the gRNA strand. The `variant` column indicates the functional consequence of the editing event (silent, nonsense or missense mutation). In case an edited allele leads to multiple editing events, the most detrimental mutation (nonsense over missense, missense over silent) is reported. The `aa` column reports the result edited amino acid sequence.

Note that several gRNA-level aggregate scores have also been added to the `GuideSet` object when calling `addEditedAlleles`:

```
head(gs)
```

```
## GuideSet object with 2 ranges and 11 metadata columns:
##            seqnames    ranges strand |         protospacer            pam
##               <Rle> <IRanges>  <Rle> |      <DNAStringSet> <DNAStringSet>
##   spacer_1    chr12     66893      - | CGCGCACCGGATTCTCCAGC            AGG
##   spacer_2    chr12     66896      + | GGGCGGCATGGAGAGCCTGC            TGG
##             pam_site  cut_site      region
##            <numeric> <numeric> <character>
##   spacer_1     66893     66896    region_1
##   spacer_2     66896     66893    region_1
##
##
##   spacer_1 CGCGTATTGGATT:0.247151:missense:...,CGCGTATCGGATT:0.161844:missense:...,CGTGTATTGGATT:0.1
##   spacer_2    GGGTGGTATGGAG:0.4644396:silent:...,GGGCGGTATGGAG:0.2976235:silent:...,GGGTGGCATGGAG:0.
##            score_missense score_nonsense score_silent  maxVariant
##                 <numeric>      <numeric>    <numeric> <character>
##   spacer_1      0.9020188              0    0.0745221    missense
##   spacer_2      0.0036734              0    0.9514897      silent
##            maxVariantScore
##                  <numeric>
##   spacer_1        0.902019
##   spacer_2        0.951490
##   -------
##   seqinfo: 640 sequences (1 circular) from hg38 genome
##   crisprNuclease: SpCas9
```

The `score_missense`, `score_nonsense` and `score_silent` columns represent aggregated scores for each of the mutation type. They were obtained by summing adding up all scores for a given mutation type across the set of edited alleles for a given gRNA. The `maxVariant` column indicates the most likely to occur mutation type for a given gRNA, and is based on the maximum aggregated score, which is stored in `maxVariantScore`. For instance, for spacer_1, the higher score is the `score_missense`, and therefore `maxVariant` is set to missense.

For more information, please see the following tutorial:

- CRISPR base editing (CRISPRbe) design

### 7 CRISPR knockdown with Cas13d

It is also possible to design gRNAs for RNA-targeting nucleases using `crisprDesign`. In contrast to DNA-targeting nucleases, the target spacer is composed of mRNA sequences instead of DNA genomic sequences.

We illustrate the functionalities of `crisprDesign` for RNA-targeting nucleases by designing gRNAs targeting IQSEC3 using the CasRx (RfxCas13d) nuclease (Konermann et al. 2018).

We first load the CasRx `CrisprNuclease` object from `crisprBase`:

```
data(CasRx, package="crisprBase")
CasRx
```

```
## Class: CrisprNuclease
##   Name: CasRx
##   Target type: RNA
##   Metadata: list of length 2
##   PFS: N
##   Weights: 1
##   Spacer length: 23
##   PFS side: 3prime
##     Distance from PFS: 0
##   Prototype protospacers: 5'--SSSSSSSSSSSSSSSSSSSSSSS[N]--3'
```

The PFS sequence (the equivalent of a PAM sequence for RNA-targeting nucleases) for CasRx is `N`, meaning that there is no specific PFS sequences preferred by CasRx.

We will now design CasRx gRNAs for the transcript ENST00000538872 of IQSEC3.

Let's first extract all mRNA sequences for IQSEC3:

```
txid <- c("ENST00000538872","ENST00000382841")
mrnas <- getMrnaSequences(txid=txid,
                          bsgenome=bsgenome,
                          txObject=grListExample)
mrnas
```

```
## DNAStringSet object of length 2:
##     width seq                                               names
## [1]  2701 AAGCCCCTCCCCTTCTCTGGGCC...AAAGTTACTGCTAGCATGGGTAA ENST00000382841
## [2]  7087 AGGCTGGGCCGGTGGGAGAGGGA...TTATATTGAAAGATGTCACTTGA ENST00000538872
```

We can use the usual function `findSpacers` to design gRNAs, and we only consider a random subset of 100 gRNAs for the sake of time:

```
gs <- findSpacers(mrnas[["ENST00000538872"]],
                  crisprNuclease=CasRx)
gs <- gs[1000:1100]
head(gs)
```

```
## GuideSet object with 6 ranges and 5 metadata columns:
##               seqnames    ranges strand |             protospacer
##                  <Rle> <IRanges>  <Rle> |          <DNAStringSet>
##   spacer_1000 region_1      1023      + | TTGACCTAAAGAATAAACAGATT
##   spacer_1001 region_1      1024      + | TGACCTAAAGAATAAACAGATTG
##   spacer_1002 region_1      1025      + | GACCTAAAGAATAAACAGATTGA
##   spacer_1003 region_1      1026      + | ACCTAAAGAATAAACAGATTGAA
##   spacer_1004 region_1      1027      + | CCTAAAGAATAAACAGATTGAAA
##   spacer_1005 region_1      1028      + | CTAAAGAATAAACAGATTGAAAT
##                         pam  pam_site  cut_site      region
##               <DNAStringSet> <numeric> <numeric> <character>
##   spacer_1000              G      1023        NA    region_1
##   spacer_1001              A      1024        NA    region_1
##   spacer_1002              A      1025        NA    region_1
##   spacer_1003              A      1026        NA    region_1
##   spacer_1004              T      1027        NA    region_1
##   spacer_1005              G      1028        NA    region_1
```

```
##   -------
##   seqinfo: 1 sequence from custom genome
##   crisprNuclease: CasRx
```

Note that all protospacer sequences are located on the original strand of the mRNA sequence. For RNA-targeting nucleases, the spacer and protospacer sequences are the reverse complement of each other:

```
head(spacers(gs))
```

```
## DNAStringSet object of length 6:
##     width seq                                               names
## [1]    23 AATCTGTTTATTCTTTAGGTCAA                           spacer_1000
## [2]    23 CAATCTGTTTATTCTTTAGGTCA                           spacer_1001
## [3]    23 TCAATCTGTTTATTCTTTAGGTC                           spacer_1002
## [4]    23 TTCAATCTGTTTATTCTTTAGGT                           spacer_1003
## [5]    23 TTTCAATCTGTTTATTCTTTAGG                           spacer_1004
## [6]    23 ATTTCAATCTGTTTATTCTTTAG                           spacer_1005
```

```
head(protospacers(gs))
```

```
## DNAStringSet object of length 6:
##     width seq                                               names
## [1]    23 TTGACCTAAAGAATAAACAGATT                           spacer_1000
## [2]    23 TGACCTAAAGAATAAACAGATTG                           spacer_1001
## [3]    23 GACCTAAAGAATAAACAGATTGA                           spacer_1002
## [4]    23 ACCTAAAGAATAAACAGATTGAA                           spacer_1003
## [5]    23 CCTAAAGAATAAACAGATTGAAA                           spacer_1004
## [6]    23 CTAAAGAATAAACAGATTGAAAT                           spacer_1005
```

The `addSpacerAlignments` can be used to perform an off-target search across all mRNA sequences using the argument `custom_seq`. Here, for the sake of time, we only perform an off-target search to the 2 isoforms of IQSEC3 specified by the `mRNAs` object:

```
gs <- addSpacerAlignments(gs,
                          aligner="biostrings",
                          txObject=grListExample,
                          n_mismatches=1,
                          custom_seq=mrnas)
tail(gs)
```

```
## GuideSet object with 6 ranges and 10 metadata columns:
##               seqnames    ranges strand |             protospacer
##                  <Rle> <IRanges>  <Rle> |          <DNAStringSet>
##   spacer_1095 region_1      1118      + | CGCCAATACCAGCTCAGCAAGAA
##   spacer_1096 region_1      1119      + | GCCAATACCAGCTCAGCAAGAAC
##   spacer_1097 region_1      1120      + | CCAATACCAGCTCAGCAAGAACT
##   spacer_1098 region_1      1121      + | CAATACCAGCTCAGCAAGAACTT
##   spacer_1099 region_1      1122      + | AATACCAGCTCAGCAAGAACTTC
##   spacer_1100 region_1      1123      + | ATACCAGCTCAGCAAGAACTTCG
##                          pam  pam_site  cut_site      region     n0_tx
##               <DNAStringSet> <numeric> <numeric> <character> <numeric>
##   spacer_1095              C      1118        NA    region_1         2
##   spacer_1096              T      1119        NA    region_1         2
##   spacer_1097              T      1120        NA    region_1         2
##   spacer_1098              C      1121        NA    region_1         2
##   spacer_1099              G      1122        NA    region_1         2
##   spacer_1100              A      1123        NA    region_1         2
```

```
##                   n1_tx   n0_gene   n1_gene
##               <numeric> <numeric> <numeric>
##   spacer_1095         0         1         0
##   spacer_1096         0         1         0
##   spacer_1097         0         1         0
##   spacer_1098         0         1         0
##   spacer_1099         0         1         0
##   spacer_1100         0         1         0
##                                                 alignments
##                                              <GRangesList>
##   spacer_1095 ENST00000382841:505:+,ENST00000538872:1118:+
##   spacer_1096 ENST00000382841:506:+,ENST00000538872:1119:+
##   spacer_1097 ENST00000382841:507:+,ENST00000538872:1120:+
##   spacer_1098 ENST00000382841:508:+,ENST00000538872:1121:+
##   spacer_1099 ENST00000382841:509:+,ENST00000538872:1122:+
##   spacer_1100 ENST00000382841:510:+,ENST00000538872:1123:+
##   -------
##   seqinfo: 1 sequence from custom genome
##   crisprNuclease: CasRx
```

The columns `n0_gene` and `n0_tx` report the number of on-targets at the gene- and transcript-level, respectively. For instance, `spacer_1095` maps to the two isoforms of IQSEC3 has `n0_tx` is equal to 2:

```
onTargets(gs["spacer_1095"])
```

```
## GRanges object with 2 ranges and 9 metadata columns:
##                      seqnames    ranges strand |                 spacer
##                         <Rle> <IRanges>  <Rle> |            <character>
##   spacer_1095 ENST00000382841       505      + | TTCTTGCTGAGCTGGTATTG..
##   spacer_1095 ENST00000538872      1118      + | TTCTTGCTGAGCTGGTATTG..
##                          protospacer            pam  pam_site n_mismatches
##                       <DNAStringSet> <DNAStringSet> <numeric>    <numeric>
##   spacer_1095 CGCCAATACCAGCTCAGCAAGAA              C       505            0
##   spacer_1095 CGCCAATACCAGCTCAGCAAGAA              C      1118            0
##                canonical  cut_site         gene_id gene_symbol
##                <logical> <numeric>     <character> <character>
##   spacer_1095      TRUE        NA ENSG00000120645      IQSEC3
##   spacer_1095      TRUE        NA ENSG00000120645      IQSEC3
##   -------
##   seqinfo: 2 sequences from custom genome
```

Note that one can also use the `bowtie` aligner to perform an off-target search to a set of mRNA sequences. This requires building a transcriptome bowtie index first instead of building a genome index. See the `crisprBowtie` vignette for more detail.

For more information, please see the following tutorial:

- CRISPR knockdown (CRISPRkd) design with CasRxdesign

### 8 Design for optical pooled screening (OPS)

Optical pooled screening (OPS) combines image-based sequencing (in situ sequencing) of gRNAs and optical phenotyping on the same physical wells (Feldman et al. 2019). In such experiments, gRNA spacer sequences are partially sequenced from the 5 prime end. From a gRNA design perspective, additional gRNA design constraints are needed to ensure sufficient dissimilarity of the truncated spacer sequences. The length of the truncated sequences, which corresponds to the number of sequencing cycles, is fixed and chosen by the experimentalist.

To illustrate the functionalities of `crisprDesign` for designing OPS libraries, we use the `guideSetExample`. We will design an OPS library with 8 cycles.

```
n_cycles=8
```

We add the 8nt OPS barcodes to the GuideSet using the `addOpsBarcodes` function:

```
data(guideSetExample, package="crisprDesign")
guideSetExample <- addOpsBarcodes(guideSetExample,
                                  n_cycles=n_cycles)
head(guideSetExample$opsBarcode)
```

```
## DNAStringSet object of length 6:
##     width seq                                               names
## [1]     8 CGCGCACC                                          spacer_1
## [2]     8 GGGCGGCA                                          spacer_2
## [3]     8 GGAGAGCC                                          spacer_3
## [4]     8 AGGTAGAG                                          spacer_4
## [5]     8 GAGCTCCT                                          spacer_5
## [6]     8 CGATGGCC                                          spacer_6
```

The function `getBarcodeDistanceMatrix` calculates the nucleotide distance between a set of query barcodes and a set of target barcodes. The type of distance (hamming or levenstein) can be specified using the `dist_method` argument. The Hamming distance (default) only considers substitutions when calculating distances, while the Levenstein distance allows insertions and deletions.

When the argument `binnarize` is set to `FALSE`, the return object is a matrix of pairwise distances between query and target barcodes:

```
barcodes <- guideSetExample$opsBarcode
dist <- getBarcodeDistanceMatrix(barcodes[1:5],
                                 barcodes[6:10],
                                 binnarize=FALSE)
print(dist)
```

```
## 5 x 5 sparse Matrix of class "dgCMatrix"
##          CGATGGCC GCGCGCCG GCTCTACC GCTCTGCT GGGTGTGG
## CGCGCACC        4        7        5        7        7
## GGGCGGCA        4        3        5        4        4
## GGAGAGCC        3        6        5        5        6
## AGGTAGAG        5        6        8        7        4
## GAGCTCCT        7        3        4        3        6
```

When `binnarize` is set to `TRUE` (default), the matrix of distances is binnarized so that 1 indicates similar barcodes, and 0 indicates dissimilar barcodes. The `min_dist_edit` argument specifies the minimal distance between two barcodes to be considered dissimilar:

```
dist <- getBarcodeDistanceMatrix(barcodes[1:5],
                                 barcodes[6:10],
                                 binnarize=TRUE,
                                 min_dist_edit=4)
print(dist)
```

```
## 5 x 5 sparse Matrix of class "dtCMatrix"
##          CGATGGCC GCGCGCCG GCTCTACC GCTCTGCT GGGTGTGG
## CGCGCACC        .        .        .        .        .
## GGGCGGCA        .        1        .        .        .
```

```
## GGAGAGCC        1        .        .        .        .
## AGGTAGAG        .        .        .        .        .
## GAGCTCCT        .        1        .        1        .
```

The `designOpsLibrary` allows users to perform a complete end-to-end library design; see `?designOpsLibrary` for documentation.

For more information, please see the following tutorial:

- Design for OPS

### 9 Design of gRNA pairs with the PairedGuideSet object

The `findSpacerPairs` function in `crisprDesign` enables the design of pairs of gRNAs and works similar to `findSpacers`. As an example, we will design candidate pairs of gRNAs that target a small locus located on chr12 in the human genome:

```
library(GenomicRanges)
library(BSgenome.Hsapiens.UCSC.hg38)
library(crisprBase)
bsgenome <- BSgenome.Hsapiens.UCSC.hg38
```

We first specify the genomic locus:

```
gr <- GRanges(c("chr12"),
              IRanges(start=22224014, end=22225007))
```

and find all pairs using the function `findSpacerPairs`:

```
pairs <- findSpacerPairs(gr, gr, bsgenome=bsgenome)
```

The first and second arguments of the function specify the which genomic region the first and second gRNA should target, respectively. In our case, we are targeting the same region with both gRNAs. The other arguments of the function are similar to the `findSpacers` function described below.

The output object is a `PairedGuideSet`, which can be thought of a list of two `GuideSet`:

```
pairs
```

```
## PairedGuideSet object with 2626 pairs and 4 metadata columns:
##                     first           second | pamOrientation pamDistance
##                <GuideSet>       <GuideSet> |    <character>   <numeric>
##      [1] chr12:22224025:- chr12:22224033:+ |            out           8
##      [2] chr12:22224025:- chr12:22224055:- |            rev          30
##      [3] chr12:22224033:+ chr12:22224055:- |             in          22
##      [4] chr12:22224025:- chr12:22224056:- |            rev          31
##      [5] chr12:22224033:+ chr12:22224056:- |             in          23
##      ...              ...              ... .            ...         ...
##   [2622] chr12:22224937:- chr12:22224994:+ |            out          57
##   [2623] chr12:22224938:- chr12:22224994:+ |            out          56
##   [2624] chr12:22224944:- chr12:22224994:+ |            out          50
##   [2625] chr12:22224950:+ chr12:22224994:+ |            fwd          44
##   [2626] chr12:22224958:- chr12:22224994:+ |            out          36
##          spacerDistance cutLength
##               <integer> <numeric>
##      [1]            -32         2
##      [2]             11        30
##      [3]             24        28
```

```
##      [4]             12        31
##      [5]             25        29
##      ...            ...       ...
##   [2622]             17        51
##   [2623]             16        50
##   [2624]             10        44
##   [2625]             25        44
##   [2626]             -4        30
```

The first and second `GuideSet` store information about gRNAs at position 1 and position 2, respectively. They can be accessed using the `first` and `second` functions:

```
grnas1 <- first(pairs)
grnas2 <- second(pairs)
grnas1
```

```
## GuideSet object with 2626 ranges and 5 metadata columns:
##              seqnames    ranges strand |          protospacer            pam
##                 <Rle> <IRanges>  <Rle> |       <DNAStringSet> <DNAStringSet>
##    spacer_1    chr12  22224025      - | ATTAGTACAACCTTTCTTTT            AGG
##    spacer_1    chr12  22224025      - | ATTAGTACAACCTTTCTTTT            AGG
##    spacer_2    chr12  22224033      + | CTTTTGTTTTCCTAAAAGAA            AGG
##    spacer_1    chr12  22224025      - | ATTAGTACAACCTTTCTTTT            AGG
##    spacer_2    chr12  22224033      + | CTTTTGTTTTCCTAAAAGAA            AGG
##         ...      ...       ...    ... .                  ...            ...
##   spacer_68    chr12  22224937      - | GGCTGCCAGTCATTGGATCA            GGG
##   spacer_69    chr12  22224938      - | AGGCTGCCAGTCATTGGATC            AGG
##   spacer_70    chr12  22224944      - | TTTATAAGGCTGCCAGTCAT            TGG
##   spacer_71    chr12  22224950      + | GTGAGCCCTGATCCAATGAC            TGG
##   spacer_72    chr12  22224958      - | CACTGTTTTTTCTTTTTATA            AGG
##               pam_site  cut_site      region
##              <numeric> <numeric> <character>
##    spacer_1  22224025  22224028    region_1
##    spacer_1  22224025  22224028    region_1
##    spacer_2  22224033  22224030    region_1
##    spacer_1  22224025  22224028    region_1
##    spacer_2  22224033  22224030    region_1
##         ...       ...       ...         ...
##   spacer_68  22224937  22224940    region_1
##   spacer_69  22224938  22224941    region_1
##   spacer_70  22224944  22224947    region_1
##   spacer_71  22224950  22224947    region_1
##   spacer_72  22224958  22224961    region_1
##   -------
##   seqinfo: 640 sequences (1 circular) from hg38 genome
##   crisprNuclease: SpCas9
```

```
grnas2
```

```
## GuideSet object with 2626 ranges and 5 metadata columns:
##              seqnames    ranges strand |          protospacer            pam
##                 <Rle> <IRanges>  <Rle> |       <DNAStringSet> <DNAStringSet>
##    spacer_2    chr12  22224033      + | CTTTTGTTTTCCTAAAAGAA            AGG
##    spacer_3    chr12  22224055      - | TATTCTCATGCACTGCTAGT            GGG
##    spacer_3    chr12  22224055      - | TATTCTCATGCACTGCTAGT            GGG
##    spacer_4    chr12  22224056      - | ATATTCTCATGCACTGCTAG            TGG
```

```
##    spacer_4    chr12  22224056      - | ATATTCTCATGCACTGCTAG            TGG
##         ...      ...       ...    ... .                  ...            ...
##   spacer_73    chr12  22224994      + | CAGTGACATAGATCATACAT            AGG
##   spacer_73    chr12  22224994      + | CAGTGACATAGATCATACAT            AGG
##   spacer_73    chr12  22224994      + | CAGTGACATAGATCATACAT            AGG
##   spacer_73    chr12  22224994      + | CAGTGACATAGATCATACAT            AGG
##   spacer_73    chr12  22224994      + | CAGTGACATAGATCATACAT            AGG
##              pam_site  cut_site      region
##             <numeric> <numeric> <character>
##    spacer_2  22224033  22224030    region_1
##    spacer_3  22224055  22224058    region_1
##    spacer_3  22224055  22224058    region_1
##    spacer_4  22224056  22224059    region_1
##    spacer_4  22224056  22224059    region_1
##         ...       ...       ...         ...
##   spacer_73  22224994  22224991    region_1
##   spacer_73  22224994  22224991    region_1
##   spacer_73  22224994  22224991    region_1
##   spacer_73  22224994  22224991    region_1
##   spacer_73  22224994  22224991    region_1
##   -------
##   seqinfo: 640 sequences (1 circular) from hg38 genome
##   crisprNuclease: SpCas9
```

The `pamOrientation` function returns the PAM orientation of the pairs:

```
head(pamOrientation(pairs))
```

```
## [1] "out" "rev" "in"  "rev" "in"  "rev"
```

and takes 4 different values: `in` (for PAM-in configuration) `out` (for PAM-out configuration), `fwd` (both gRNAs target the forward strand) and `rev` (both gRNAs target the reverse strand).

The function `pamDistance` returns the distance between the PAM sites of the two gRNAs. The function `cutLength` returns the distance between the cut sites of the two gRNAs. The function `spacerDistance` returns the distance between the two spacer sequences of the gRNAs.

For more information, please see the following tutorial:

- Paired gRNA design

### 10 Miscellaneous design use cases

#### 10.1 Design with custom sequences

`crisprDesign` also allows gRNA design for DNA sequences without genomic context (such as a synthesized DNA construct). See `?findSpacers` for more information, and here's an example:

```
seqs <- c(seq1="AGGCGGAGGCCCGACCCGGGCGCGGGGCGGCGC",
          seq2="AGGCGGAGGCCCGACCCGGGCGCGGGAAAAAAAAGGC")
gs <- findSpacers(seqs)
head(gs)
```

```
## GuideSet object with 6 ranges and 5 metadata columns:
##            seqnames    ranges strand |          protospacer             pam
##               <Rle> <IRanges>  <Rle> |       <DNAStringSet> <DNAStringSet>
##   spacer_1     seq1        12      - | CGCCGCCCCGCGCCCGGGTC            GGG
##   spacer_2     seq1        13      - | GCGCCGCCCCGCGCCCGGGT            CGG
```

```
##   spacer_3     seq1        23      + | GCGGAGGCCCGACCCGGGCG            CGG
##   spacer_4     seq1        24      + | CGGAGGCCCGACCCGGGCGC            GGG
##   spacer_5     seq1        25      + | GGAGGCCCGACCCGGGCGCG            GGG
##   spacer_6     seq1        28      + | GGCCCGACCCGGGCGCGGGG            CGG
##              pam_site  cut_site      region
##             <numeric> <numeric> <character>
##   spacer_1        12        15        seq1
##   spacer_2        13        16        seq1
##   spacer_3        23        20        seq1
##   spacer_4        24        21        seq1
##   spacer_5        25        22        seq1
##   spacer_6        28        25        seq1
##   -------
##   seqinfo: 2 sequences from custom genome
##   crisprNuclease: SpCas9
```

##### 10.2 Off-target search in custom sequences

One can also search for off-targets in a custom sequence as follows:

```
ontarget <- "AAGACCCGGGCGCGGGGCGGGGG"
offtarget <- "TTGACCCGGGCGCGGGGCGGGGG"
gs <- findSpacers(ontarget)
gs <- addSpacerAlignments(gs,
                          aligner="biostrings",
                          n_mismatches=2,
                          custom_seq=offtarget)
head(alignments(gs))
```

```
## GRanges object with 1 range and 7 metadata columns:
##                seqnames    ranges strand |                spacer
##                   <Rle> <IRanges>  <Rle> |        <DNAStringSet>
##   spacer_1 custom_seq1        21      + | AAGACCCGGGCGCGGGGCGG
##                         protospacer            pam  pam_site n_mismatches canonical
##                      <DNAStringSet> <DNAStringSet> <numeric>    <numeric> <logical>
##   spacer_1 TTGACCCGGGCGCGGGGCGG            GGG        21            2      TRUE
##              cut_site
##             <numeric>
##   spacer_1        18
##   -------
##   seqinfo: 1 sequence from custom genome
```

For more information, please see the following tutorial:

- Working with custom DNA sequences

### 11 Session Info

```
sessionInfo()
```

```
## R version 4.2.1 (2022-06-23)
## Platform: x86_64-apple-darwin17.0 (64-bit)
## Running under: macOS Catalina 10.15.7
##
## Matrix products: default
```

```
## BLAS:   /Library/Frameworks/R.framework/Versions/4.2/Resources/lib/libRblas.0.dylib
## LAPACK: /Library/Frameworks/R.framework/Versions/4.2/Resources/lib/libRlapack.dylib
##
## locale:
## [1] en_US.UTF-8/en_US.UTF-8/en_US.UTF-8/C/en_US.UTF-8/en_US.UTF-8
##
## attached base packages:
## [1] stats4    stats     graphics  grDevices utils     datasets  methods
## [8] base
##
## other attached packages:
##  [1] Rbowtie_1.37.0                    BSgenome.Hsapiens.UCSC.hg38_1.4.4
##  [3] BSgenome_1.65.2                   rtracklayer_1.57.0
##  [5] Biostrings_2.65.2                 XVector_0.37.0
##  [7] GenomicRanges_1.49.1              GenomeInfoDb_1.33.5
##  [9] IRanges_2.31.2                    S4Vectors_0.35.1
## [11] BiocGenerics_0.43.1               crisprDesign_0.99.134
## [13] crisprBase_1.1.5
##
## loaded via a namespace (and not attached):
##   [1] bitops_1.0-7                  matrixStats_0.62.0
##   [3] bit64_4.0.5                   filelock_1.0.2
##   [5] progress_1.2.2                httr_1.4.4
##   [7] tools_4.2.1                   utf8_1.2.2
##   [9] R6_2.5.1                      DBI_1.1.3
##  [11] tidyselect_1.1.2              prettyunits_1.1.1
##  [13] bit_4.0.4                     curl_4.3.2
##  [15] compiler_4.2.1                crisprBowtie_1.1.1
##  [17] cli_3.3.0                     Biobase_2.57.1
##  [19] basilisk.utils_1.9.1          crisprScoreData_1.1.3
##  [21] xml2_1.3.3                    DelayedArray_0.23.1
##  [23] randomForest_4.7-1.1          readr_2.1.2
##  [25] rappdirs_0.3.3                stringr_1.4.1
##  [27] digest_0.6.29                 Rsamtools_2.13.4
##  [29] rmarkdown_2.15.2              crisprScore_1.1.14
##  [31] basilisk_1.9.3                pkgconfig_2.0.3
##  [33] htmltools_0.5.3               MatrixGenerics_1.9.1
##  [35] dbplyr_2.2.1                  fastmap_1.1.0
##  [37] rlang_1.0.4                   rstudioapi_0.14
##  [39] RSQLite_2.2.16                shiny_1.7.2
##  [41] BiocIO_1.7.1                  generics_0.1.3
##  [43] jsonlite_1.8.0                vroom_1.5.7
##  [45] BiocParallel_1.31.12          dplyr_1.0.9
##  [47] VariantAnnotation_1.43.3      RCurl_1.98-1.8
##  [49] magrittr_2.0.3                GenomeInfoDbData_1.2.8
##  [51] Matrix_1.4-1                  Rcpp_1.0.9
##  [53] fansi_1.0.3                   reticulate_1.25
##  [55] lifecycle_1.0.1               stringi_1.7.8
##  [57] yaml_2.3.5                    SummarizedExperiment_1.27.1
##  [59] zlibbioc_1.43.0               BiocFileCache_2.5.0
##  [61] AnnotationHub_3.5.0           grid_4.2.1
##  [63] blob_1.2.3                    promises_1.2.0.1
##  [65] parallel_4.2.1                ExperimentHub_2.5.0
##  [67] crayon_1.5.1                  crisprBwa_1.1.3
```

```
## [69] dir.expiry_1.5.0              lattice_0.20-45
## [71] GenomicFeatures_1.49.6        hms_1.1.2
## [73] KEGGREST_1.37.3               knitr_1.40
## [75] pillar_1.8.1                  rjson_0.2.21
## [77] codetools_0.2-18              biomaRt_2.53.2
## [79] BiocVersion_3.16.0            XML_3.99-0.10
## [81] glue_1.6.2                    evaluate_0.16
## [83] BiocManager_1.30.18           httpuv_1.6.5
## [85] png_0.1-7                     vctrs_0.4.1
## [87] tzdb_0.3.0                    purrr_0.3.4
## [89] assertthat_0.2.1              cachem_1.0.6
## [91] xfun_0.32                     mime_0.12
## [93] Rbwa_1.1.0                    xtable_1.8-4
## [95] restfulr_0.0.15               later_1.3.0
## [97] tibble_3.1.8                  GenomicAlignments_1.33.1
## [99] AnnotationDbi_1.59.1          memoise_2.0.1
## [101] interactiveDisplayBase_1.35.0 ellipsis_0.3.2
```

### References


Feldman, David, Avtar Singh, Jonathan L Schmid-Burgk, Rebecca J Carlson, Anja Mezger, Anthony J Garrity, Feng Zhang, and Paul C Blainey. 2019. "Optical Pooled Screens in Human Cells." *Cell* 179 (3): 787–99.

Gilbert, Luke A, Matthew H Larson, Leonardo Morsut, Zairan Liu, Gloria A Brar, Sandra E Torres, Noam Stern-Ginossar, et al. 2013. "CRISPR-Mediated Modular RNA-Guided Regulation of Transcription in Eukaryotes." *Cell* 154 (2): 442–51.

Kampmann, Martin. 2018. "CRISPRi and CRISPRa Screens in Mammalian Cells for Precision Biology and Medicine." *ACS Chemical Biology* 13 (2): 406–16.

Koblan, Luke W, Jordan L Doman, Christopher Wilson, Jonathan M Levy, Tristan Tay, Gregory A Newby, Juan Pablo Maianti, Aditya Raguram, and David R Liu. 2018. "Improving Cytidine and Adenine Base Editors by Expression Optimization and Ancestral Reconstruction." *Nature Biotechnology* 36 (9): 843–46.

Konermann, Silvana, Peter Lotfy, Nicholas J Brideau, Jennifer Oki, Maxim N Shokhirev, and Patrick D Hsu. 2018. "Transcriptome Engineering with RNA-Targeting Type VI-d CRISPR Effectors." *Cell* 173 (3): 665–76.

Langmead, Ben, Cole Trapnell, Mihai Pop, and Steven L. Salzberg. 2009. "Ultrafast and Memory-Efficient Alignment of Short DNA Sequences to the Human Genome." *Genome Biology* 10 (3): R25. https://doi.org/10.1186/gb-2009-10-3-r25.

Sanson, Kendall R, Ruth E Hanna, Mudra Hegde, Katherine F Donovan, Christine Strand, Meagan E Sullender, Emma W Vaimberg, et al. 2018. "Optimized Libraries for CRISPR-Cas9 Genetic Screens with Multiple Modalities." *Nature Communications* 9 (1): 1–15.
