## Supplementary material for "The crisprVerse: a comprehensive Bioconductor ecosystem for the design of CRISPR guide RNAs across nucleases and technologies": Vignette_crisprDesignData

### crisprDesignData: useful data for the crisprVerse ecosystem

2022-08-24

#### 1 Overview

The `crisprDesignData` package provides ready-to-use annotation data needed needed for the `crisprVerse` ecosystem, for both human and human.

#### 2 Installation

##### 2.1 Software requirements

###### 2.1.1 OS Requirements

This package is supported for macOS, Linux and Windows machines. It was developed and tested on R version 4.2.1

##### 2.2 Installation

`crisprDesignData` can be installed by typing the following commands inside of an R session:

```
install.packages("devtools")
devtools::install_github("crisprVerse/crisprDesignData")
```

###### 2.2.1 Getting started

`crisprDesignData` can be loaded into an R session in the usual way:

```
library(crisprDesignData)
```

#### 3 Datasets

| Object name | Object class | Version | Description |
|---|---|---|---|
| `txdb_human` | `GRangesList` | Release 104 | Ensembl gene model for human (hg38/GRCh38) |
| `txdb_mouse` | `GRangesList` | Release 102 | Ensembl gene model for mouse (mm10/GRCm38) |
| `tss_human` | `GRanges` | Release 104 | Ensembl-based TSS coordinates for human (hg38/GRCh38) |
| `tss_mouse` | `GRanges` | Release 102 | Ensembl-based TSS coordinates for human (mm10/GRCm38) |

| Object name | Object class | Version | Description |
|---|---|---|---|
| `gr.repeats.hg38` | `GRanges` | | RepeatMasker data from UCSC genome browser (hg38/GRCh38) |
| `gr.repeats.mm10` | `GRanges` | | RepeatMasker data from UCSC genome browser (mm10/GRCm38) |
| `canonicalHuman` | `data.frame` | Release 104 | Canonical Ensembl transcripts for human |
| `canonicalMouse` | `data.frame` | Release 102 | Canonical Ensembl transcripts for mouse |

### 4 TxDb datasets

The `txdb_human` and `txdb_mouse` objects are `GRangesList` representing gene models for human and mouse, respectively, from Ensembl. They were constructed using the function `getTxDb` in `crisprDesign`. See the script `generateTxDbData.R`in the `inst` folder to see how to generate such data for other organisms (internet connection needed).

Let's look at the `txdb_human` object. We first load the data:

```
data(txdb_human, package="crisprDesignData")
```

We can look at metadata information about the gene model by using the `metadata` function from the `S4Vectors` package:

```
head(S4Vectors::metadata(txdb_human))
```

```
##                  name                      value
## 1             Db type                      TxDb
## 2 Supporting package          GenomicFeatures
## 3         Data source                   Ensembl
## 4            Organism              Homo sapiens
## 5     Ensembl release                       104
## 6    Ensembl database homo_sapiens_core_104_38
```

The object is a `GRangesList` with 7 elements that contain genomic coordinates for different levels of the gene model:

```
names(txdb_human)
```

```
## [1] "transcripts" "exons"       "cds"         "fiveUTRs"    "threeUTRs"
## [6] "introns"     "tss"
```

As an example, let's look at the `GRanges` containing genomic coordinates for all exons represented in the gene model:

```
txdb_human$exons
```

```
## GRanges object with 796644 ranges and 14 metadata columns:
##      seqnames      ranges strand |           tx_id         gene_id
##         <Rle>   <IRanges>  <Rle> |     <character>     <character>
##          chr1 11869-12227      + | ENST00000456328 ENSG00000223972
##          chr1 12613-12721      + | ENST00000456328 ENSG00000223972
##          chr1 13221-14409      + | ENST00000456328 ENSG00000223972
##          chr1 12010-12057      + | ENST00000450305 ENSG00000223972
##          chr1 12179-12227      + | ENST00000450305 ENSG00000223972
```

```
##   .       ...         ...     ... .             ...             ...
##         chrM   5826-5891      - | ENST00000387409 ENSG00000210144
##         chrM   7446-7514      - | ENST00000387416 ENSG00000210151
##         chrM 14149-14673      - | ENST00000361681 ENSG00000198695
##         chrM 14674-14742      - | ENST00000387459 ENSG00000210194
##         chrM 15956-16023      - | ENST00000387461 ENSG00000210196
##          protein_id                tx_type gene_symbol         exon_id
##         <character>            <character> <character>     <character>
##                <NA>   processed_transcript     DDX11L1 ENSE00002234944
##                <NA>   processed_transcript     DDX11L1 ENSE00003582793
##                <NA>   processed_transcript     DDX11L1 ENSE00002312635
##                <NA> transcribed_unproces..     DDX11L1 ENSE00001948541
##                <NA> transcribed_unproces..     DDX11L1 ENSE00001671638
##   .             ...                    ...         ...             ...
##                <NA>                Mt_tRNA       MT-TY ENSE00001544488
##                <NA>                Mt_tRNA      MT-TS1 ENSE00001544487
##     ENSP00000354665         protein_coding      MT-ND6 ENSE00001434974
##                <NA>                Mt_tRNA       MT-TE ENSE00001544476
##                <NA>                Mt_tRNA       MT-TP ENSE00001544473
##     exon_rank cds_start   cds_end  tx_start    tx_end   cds_len exon_start
##     <integer> <integer> <integer> <integer> <integer> <integer>  <integer>
##             1      <NA>      <NA>     11869     14409         0       <NA>
##             2      <NA>      <NA>     11869     14409         0       <NA>
##             3      <NA>      <NA>     11869     14409         0       <NA>
##             1      <NA>      <NA>     12010     13670         0       <NA>
##             2      <NA>      <NA>     12010     13670         0       <NA>
##   .       ...       ...       ...       ...       ...       ...        ...
##             1      <NA>      <NA>      5826      5891         0       <NA>
##             1      <NA>      <NA>      7446      7514         0       <NA>
##             1     14149     14673     14149     14673       525       <NA>
##             1      <NA>      <NA>     14674     14742         0       <NA>
##             1      <NA>      <NA>     15956     16023         0       <NA>
##      exon_end
##     <integer>
##          <NA>
##          <NA>
##          <NA>
##          <NA>
##          <NA>
##   .       ...
##          <NA>
##          <NA>
##          <NA>
##          <NA>
##          <NA>
##   -------
##   seqinfo: 25 sequences (1 circular) from hg38 genome
```

The function `queryTxObject` in `crisprDesign` is a user-friendly function to work with such objects, for instance once can return the CDS coordinates for the KRAS transcripts using the following lines of code:

```
library(crisprDesign)
cds <- queryTxObject(txdb_human,
                     featureType="cds",
                     queryColumn="gene_symbol",
```

```r
                      queryValue="KRAS")
head(cds)
```

```
## GRanges object with 6 ranges and 14 metadata columns:
##            seqnames            ranges strand |           tx_id         gene_id
##               <Rle>         <IRanges>  <Rle> |     <character>     <character>
##   region_1    chr12 25245274-25245384      - | ENST00000256078 ENSG00000133703
##   region_2    chr12 25227234-25227412      - | ENST00000256078 ENSG00000133703
##   region_3    chr12 25225614-25225773      - | ENST00000256078 ENSG00000133703
##   region_4    chr12 25215441-25215560      - | ENST00000256078 ENSG00000133703
##   region_5    chr12 25245274-25245384      - | ENST00000311936 ENSG00000133703
##   region_6    chr12 25227234-25227412      - | ENST00000311936 ENSG00000133703
##                 protein_id        tx_type gene_symbol         exon_id exon_rank
##                <character>    <character> <character>     <character> <integer>
##   region_1 ENSP00000256078 protein_coding        KRAS ENSE00000936617         2
##   region_2 ENSP00000256078 protein_coding        KRAS ENSE00001719809         3
##   region_3 ENSP00000256078 protein_coding        KRAS ENSE00001644818         4
##   region_4 ENSP00000256078 protein_coding        KRAS ENSE00001189807         5
##   region_5 ENSP00000256078 protein_coding        KRAS ENSE00000936617         2
##   region_6 ENSP00000256078 protein_coding        KRAS ENSE00001719809         3
##            cds_start   cds_end  tx_start    tx_end   cds_len exon_start
##            <integer> <integer> <integer> <integer> <integer>  <integer>
##   region_1      <NA>      <NA>  25205246  25250929       570   25245274
##   region_2      <NA>      <NA>  25205246  25250929       570   25227234
##   region_3      <NA>      <NA>  25205246  25250929       570   25225614
##   region_4      <NA>      <NA>  25205246  25250929       570   25215437
##   region_5      <NA>      <NA>  25205246  25250929       567   25245274
##   region_6      <NA>      <NA>  25205246  25250929       567   25227234
##             exon_end
##            <integer>
##   region_1  25245395
##   region_2  25227412
##   region_3  25225773
##   region_4  25215560
##   region_5  25245395
##   region_6  25227412
##   -------
##   seqinfo: 25 sequences (1 circular) from hg38 genome
```

### 5 TSS datasets

The `tss_human` and `tss_mouse` objects are `GRanges` representing the transcription starting sites (TSSs) coordinates for human and mouse, respectively. The coordinates were extracted from the transcripts stored in the Ensembl-based models `txdb_human` and `txdb_mouse` using the function `getTssObjectFromTxObject` from `crisprDesign`. See the script `generateTssObjects.R`in the `inst` folder to see how to generate such data.

Let's take a look at `tss_human`:

```r
data(tss_human, package="crisprDesignData")
head(tss_human)
```

```
## GRanges object with 6 ranges and 9 metadata columns:
##                         seqnames    ranges strand |     score peak_start  peak_end
```

```
##                          <Rle> <IRanges>  <Rle> | <numeric>  <integer> <integer>
##   ENSG00000000003_P1      chrX 100636805      - |   4.35417  100636805 100636805
##   ENSG00000000005_P1      chrX 100584935      + |   3.29137  100584935 100584935
##   ENSG00000000419_P1     chr20  50958531      - |   5.74747   50958531  50958531
##   ENSG00000000457_P1      chr1 169893895      - |   4.75432  169893895 169893895
##   ENSG00000000460_P1      chr1 169795044      + |   4.92777  169795044 169795044
##   ENSG00000000938_P1      chr1  27635184      - |   4.61214   27635184  27635184
##                                 tx_id         gene_id      source    promoter
##                           <character>     <character> <character> <character>
##   ENSG00000000003_P1 ENST00000373020 ENSG00000000003     fantom5          P1
##   ENSG00000000005_P1 ENST00000373031 ENSG00000000005     fantom5          P1
##   ENSG00000000419_P1 ENST00000371588 ENSG00000000419     fantom5          P1
##   ENSG00000000457_P1 ENST00000367771 ENSG00000000457     fantom5          P1
##   ENSG00000000460_P1 ENST00000359326 ENSG00000000460     fantom5          P1
##   ENSG00000000938_P1 ENST00000374005 ENSG00000000938     fantom5          P1
##                                      ID gene_symbol
##                             <character> <character>
##   ENSG00000000003_P1 ENSG00000000003_P1      TSPAN6
##   ENSG00000000005_P1 ENSG00000000005_P1        TNMD
##   ENSG00000000419_P1 ENSG00000000419_P1        DPM1
##   ENSG00000000457_P1 ENSG00000000457_P1       SCYL3
##   ENSG00000000460_P1 ENSG00000000460_P1    C1orf112
##   ENSG00000000938_P1 ENSG00000000938_P1         FGR
##   -------
##   seqinfo: 25 sequences from an unspecified genome; no seqlengths
```

The function `queryTss` in `crisprDesign` is a user-friendly function to work with such objects, accepting an argument called `tss_window` to specify a number of nucleotides upstream and downstream of the TSS. This is particularly useful to return genomic regions to target for CRISPRa and CRISPRi.

For instance, if we want to target the region 500 nucleotides upstream of any of the KRAS TSSs, one can use the following lines of code:

```
library(crisprDesign)
tss <- queryTss(tss_human,
                queryColumn="gene_symbol",
                queryValue="KRAS",
                tss_window=c(-500,0))
tss
```

```
## GRanges object with 1 range and 9 metadata columns:
##            seqnames            ranges strand |     score peak_start  peak_end
##               <Rle>         <IRanges>  <Rle> | <numeric>  <integer> <integer>
##   region_1    chr12 25250929-25251428      - |   5.20187   25250928  25250928
##                     tx_id         gene_id      source    promoter
##               <character>     <character> <character> <character>
##   region_1 ENST00000256078 ENSG00000133703     fantom5          P1
##                          ID gene_symbol
##                 <character> <character>
##   region_1 ENSG00000133703_P1        KRAS
##   -------
##   seqinfo: 25 sequences from an unspecified genome; no seqlengths
```

### 6 Repeats datasets

The objects `gr.repeats.hg38` and `gr.repeats.mm10` objects are `GRanges` representing the genomic coordinates of repeat elements in the human and mouse genomes, as defined by the RepeatMasker tracks in the UCSC genome browser.

Let's look at the repeats elements in the human genome:

```
data(gr.repeats.hg38, package="crisprDesignData")
head(gr.repeats.hg38)
```

```
## GRanges object with 6 ranges and 2 metadata columns:
##       seqnames            ranges strand |        type     score
##          <Rle>         <IRanges>  <Rle> | <character> <numeric>
##   [1]     chr1 67108753-67109046      + |        L1P5      1892
##   [2]     chr1   8388315-8388618      - |        AluY      2582
##   [3]     chr1 25165803-25166380      + |       L1MB5      4085
##   [4]     chr1 33554185-33554483      - |       AluSc      2285
##   [5]     chr1 41942894-41943205      - |        AluY      2451
##   [6]     chr1 50331336-50332274      + |        HAL1      1587
##   -------
##   seqinfo: 25 sequences (1 circular) from hg38 genome
```

### 7 Canonical transcripts

The data.frames `canonicalHuman` and `canonicalMouse` contains information about Ensembl canonical transcripts for human and mouse respectively. The Ensembl canonical transcript is the best well-supported, biologically representative, highly expressed, and highly conserved transcript for a given gene. MANE Select is used as the canonical transcript for human protein coding genes where available.

```
data(canonicalHuman, package="crisprDesignData")
head(canonicalHuman)
```

```
##             tx_id         gene_id
## 1 ENST00000272065 ENSG00000143727
## 2 ENST00000329066 ENSG00000115705
## 3 ENST00000252505 ENSG00000151360
## 4 ENST00000256509 ENSG00000134121
## 5 ENST00000349077 ENSG00000118004
## 6 ENST00000273130 ENSG00000144635
```

### 8 License

The package is licensed under the MIT license.

### 9 Reproducibility

```
sessionInfo()
```

```
## R version 4.2.1 (2022-06-23)
## Platform: x86_64-apple-darwin17.0 (64-bit)
## Running under: macOS Catalina 10.15.7
##
## Matrix products: default
```

```
## BLAS:   /Library/Frameworks/R.framework/Versions/4.2/Resources/lib/libRblas.0.dylib
## LAPACK: /Library/Frameworks/R.framework/Versions/4.2/Resources/lib/libRlapack.dylib
##
## locale:
## [1] en_US.UTF-8/en_US.UTF-8/en_US.UTF-8/C/en_US.UTF-8/en_US.UTF-8
##
## attached base packages:
## [1] stats4    stats     graphics  grDevices utils     datasets  methods
## [8] base
##
## other attached packages:
## [1] crisprDesign_0.99.132    crisprBase_1.1.5         GenomicRanges_1.49.1
## [4] GenomeInfoDb_1.33.5      IRanges_2.31.2           S4Vectors_0.35.1
## [7] BiocGenerics_0.43.1      crisprDesignData_0.99.17
##
## loaded via a namespace (and not attached):
##   [1] bitops_1.0-7                  matrixStats_0.62.0
##   [3] bit64_4.0.5                   filelock_1.0.2
##   [5] progress_1.2.2                httr_1.4.4
##   [7] tools_4.2.1                   utf8_1.2.2
##   [9] R6_2.5.1                      DBI_1.1.3
##  [11] tidyselect_1.1.2              prettyunits_1.1.1
##  [13] bit_4.0.4                     curl_4.3.2
##  [15] compiler_4.2.1                crisprBowtie_1.1.1
##  [17] cli_3.3.0                     Biobase_2.57.1
##  [19] basilisk.utils_1.9.1          crisprScoreData_1.1.3
##  [21] xml2_1.3.3                    DelayedArray_0.23.1
##  [23] rtracklayer_1.57.0            randomForest_4.7-1.1
##  [25] readr_2.1.2                   rappdirs_0.3.3
##  [27] stringr_1.4.1                 digest_0.6.29
##  [29] Rsamtools_2.13.4              rmarkdown_2.15.2
##  [31] crisprScore_1.1.14            basilisk_1.9.2
##  [33] XVector_0.37.0                pkgconfig_2.0.3
##  [35] htmltools_0.5.3               MatrixGenerics_1.9.1
##  [37] dbplyr_2.2.1                  fastmap_1.1.0
##  [39] BSgenome_1.65.2               rlang_1.0.4
##  [41] rstudioapi_0.14               RSQLite_2.2.16
##  [43] shiny_1.7.2                   BiocIO_1.7.1
##  [45] generics_0.1.3                jsonlite_1.8.0
##  [47] BiocParallel_1.31.12          dplyr_1.0.9
##  [49] VariantAnnotation_1.43.3      RCurl_1.98-1.8
##  [51] magrittr_2.0.3                GenomeInfoDbData_1.2.8
##  [53] Matrix_1.4-1                  Rcpp_1.0.9
##  [55] fansi_1.0.3                   reticulate_1.25
##  [57] Rbowtie_1.37.0                lifecycle_1.0.1
##  [59] stringi_1.7.8                 yaml_2.3.5
##  [61] SummarizedExperiment_1.27.1   zlibbioc_1.43.0
##  [63] AnnotationHub_3.5.0           BiocFileCache_2.5.0
##  [65] grid_4.2.1                    blob_1.2.3
##  [67] promises_1.2.0.1              parallel_4.2.1
##  [69] ExperimentHub_2.5.0           crayon_1.5.1
##  [71] dir.expiry_1.5.0              lattice_0.20-45
##  [73] Biostrings_2.65.2             GenomicFeatures_1.49.6
##  [75] hms_1.1.2                     KEGGREST_1.37.3
```

```
##  [77] knitr_1.40                    pillar_1.8.1
##  [79] rjson_0.2.21                  codetools_0.2-18
##  [81] biomaRt_2.53.2                BiocVersion_3.16.0
##  [83] XML_3.99-0.10                 glue_1.6.2
##  [85] evaluate_0.16                 BiocManager_1.30.18
##  [87] httpuv_1.6.5                  png_0.1-7
##  [89] vctrs_0.4.1                   tzdb_0.3.0
##  [91] purrr_0.3.4                   assertthat_0.2.1
##  [93] cachem_1.0.6                  xfun_0.32
##  [95] mime_0.12                     xtable_1.8-4
##  [97] restfulr_0.0.15               later_1.3.0
##  [99] tibble_3.1.8                  GenomicAlignments_1.33.1
## [101] AnnotationDbi_1.59.1          memoise_2.0.1
## [103] interactiveDisplayBase_1.35.0 ellipsis_0.3.2
```
