## Supplementary material for "The crisprVerse: a comprehensive Bioconductor ecosystem for the design of CRISPR guide RNAs across nucleases and technologies": Vignette_crisprDesignScore

### On-target and off-target scoring for CRISPR gRNAs

2022-08-30

### 1 Overview

crisprScore provides R wrappers of several on-target and off-target scoring methods for CRISPR guide RNAs (gRNAs). The following nucleases are supported: SpCas9, AsCas12a, enAsCas12a, and RfxCas13d (CasRx). The available on-target cutting efficiency scoring methods are RuleSet1, RuleSet3, Azimuth, DeepHF, DeepSpCas9, DeepCpf1, enPAM+GB, CRISPRscan and CRISPRater. Both the CFD and MIT scoring methods are available for off-target specificity prediction. The package also provides a Lindel-derived score to predict the probability of a gRNA to produce indels inducing a frameshift for the Cas9 nuclease. Note that DeepHF, DeepCpf1 and enPAM+GB are not available on Windows machines.

Our work is described in a recent bioRxiv preprint: "A comprehensive Bioconductor ecosystem for the design of CRISPR guide RNAs across nucleases and technologies"

### 2 Installation and getting started

#### 2.1 Software requirements

##### 2.1.1 OS Requirements

This package is supported for macOS, Linux and Windows machines. Some functionalities are not supported for Windows machines. Packages were developed and tested on R version 4.2.1.

#### 2.2 Installation from Bioconductor

`crisprScore` can be installed from from the Bioconductor devel branch using the following commands in a fresh R session:

```
if (!require("BiocManager", quietly = TRUE))
    install.packages("BiocManager")

BiocManager::install(version="devel")
BiocManager::install("crisprScore")
```

#### 2.3 Installation from GitHub

Alternatively, the development version of `crisprScore` and its dependencies can be installed by typing the following commands inside of an R session:

```
install.packages("devtools")
library(devtools)
install_github("crisprVerse/crisprScoreData")
install_github("crisprVerse/crisprScore")
```

When calling one of the scoring methods for the first time after package installation, the underlying python module and conda environment will be automatically downloaded and installed without the need for user intervention. This may take several minutes, but this is a one-time installation. the first time after package installation.

Note that RStudio users will need to add the following line to their `.Rprofile` file in order for `crisprScore` to work properly:

```
options(reticulate.useImportHook=FALSE)
```

### 3 Getting started

We load `crisprScore` in the usual way:

```
library(crisprScore)
```

The `scoringMethodsInfo` data.frame contains a succinct summary of scoring methods available in `crisprScore`:

```
data(scoringMethodsInfo)
print(scoringMethodsInfo)
```

```
##        method   nuclease left right       type      label len
## 1    ruleset1     SpCas9  -24     5  On-target   RuleSet1  30
## 2     azimuth     SpCas9  -24     5  On-target    Azimuth  30
## 3      deephf     SpCas9  -20     2  On-target     DeepHF  23
## 4      lindel     SpCas9  -33    31  On-target     Lindel  65
## 5         mit     SpCas9  -20     2 Off-target        MIT  23
## 6         cfd     SpCas9  -20     2 Off-target        CFD  23
## 7    deepcpf1   AsCas12a   -4    29  On-target   DeepCpf1  34
## 8     enpamgb enAsCas12a   -4    29  On-target     EnPAMGB  34
## 9  crisprscan     SpCas9  -26     8  On-target CRISPRscan  35
## 10    casrxrf      CasRx   NA    NA  On-target    CasRx-RF  NA
## 11   crisprai     SpCas9  -19     2  On-target    CRISPRai  22
## 12 crisprater     SpCas9  -20    -1  On-target CRISPRater  20
## 13 deepspcas9     SpCas9  -24     5  On-target DeepSpCas9  30
## 14   ruleset3     SpCas9  -24     5  On-target   RuleSet3  30
```

Each scoring algorithm requires a different contextual nucleotide sequence. The `left` and `right` columns indicates how many nucleotides upstream and downstream of the first nucleotide of the PAM sequence are needed for input, and the `len` column indicates the total number of nucleotides needed for input. The `crisprDesign` (GitHub link) package provides user-friendly functionalities to extract and score those sequences automatically via the `addOnTargetScores` function.

### 4 On-targeting efficiency scores

Predicting on-target cutting efficiency is an extensive area of research, and we try to provide in `crisprScore` the latest state-of-the-art algorithms as they become available.

#### 4.1 Cas9 methods

Different algorithms require different input nucleotide sequences to predict cutting efficiency as illustrated in the figure below.

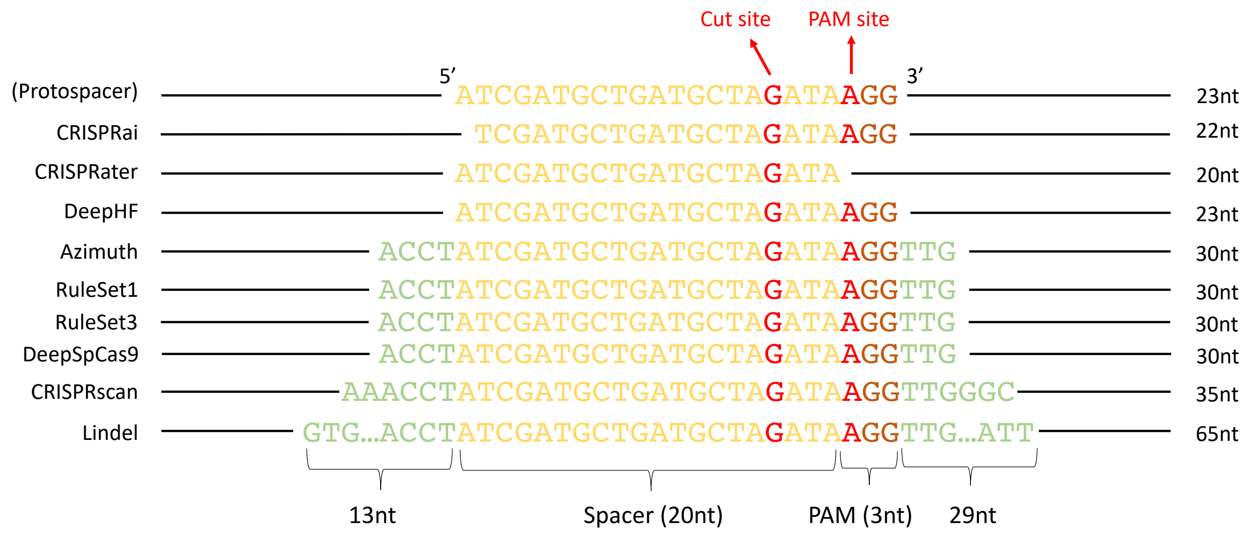


Figure 1: Sequence inputs for Cas9 scoring methods

###### 4.1.1 Rule Set 1

The Rule Set 1 algorithm is one of the first on-target efficiency methods developed for the Cas9 nuclease (Doench et al. 2014). It generates a probability (therefore a score between 0 and 1) that a given sgRNA will cut at its intended target. 4 nucleotides upstream and 3 nucleotides downstream of the PAM sequence are needed for scoring:

```
flank5 <- "ACCT" #4bp
spacer <- "ATCGATGCTGATGCTAGATA" #20bp
pam    <- "AGG" #3bp
flank3 <- "TTG" #3bp
input  <- paste0(flank5, spacer, pam, flank3)
results <- getRuleSet1Scores(input)
```

The Azimuth score described below is an improvement over Rule Set 1 from the same lab.

###### 4.1.2 Azimuth

The Azimuth algorithm is an improved version of the popular Rule Set 2 score for the Cas9 nuclease (Doench et al. 2016). It generates a probability (therefore a score between 0 and 1) that a given sgRNA will cut at its intended target. 4 nucleotides upstream and 3 nucleotides downstream of the PAM sequence are needed for scoring:

```
flank5 <- "ACCT" #4bp
spacer <- "ATCGATGCTGATGCTAGATA" #20bp
pam    <- "AGG" #3bp
flank3 <- "TTG" #3bp
input  <- paste0(flank5, spacer, pam, flank3)
results <- getAzimuthScores(input)
```

###### 4.1.3 Rule Set 3

The Rule Set 3 is an improvement over Rule Set 1 and Rule Set 2/Azimuth developed for the SpCas9 nuclease, taking into account the type of tracrRNAs (DeWeirdt et al. 2022). Two types of tracrRNAs are currently offered:

```
GTTTTAGAGCTA-----GAAA-----TAGCAAGTTAAAAT... --> Hsu2013 tracrRNA
GTTTAAGAGCTATGCTGGAAACAGCATAGCAAGTTTAAAT... --> Chen2013 tracrRNA
```

Similar to Rule Set 1 and Azimuth, the input sequence requires 4 nucleotides upstream of the protospacer sequence, the protospacer sequence itself (20nt spacersequence and PAM sequence), and 3 nucleotides downstream of the PAM sequence:

```
flank5 <- "ACCT" #4bp
spacer <- "ATCGATGCTGATGCTAGATA" #20bp
pam    <- "AGG" #3bp
flank3 <- "TTG" #3bp
input  <- paste0(flank5, spacer, pam, flank3)
results <- getRuleSet3Scores(input, tracrRNA="Hsu2013")
```

A more involved version of the algorithm takes into account gene context of the target protospacer sequence (Rule Set 3 Target) and will be soon implemented in `crisprScore`.

###### 4.1.4 DeepHF

The DeepHF algorithm is an on-target cutting efficiency prediction algorithm for several variants of the Cas9 nuclease (Wang et al. 2019) using a recurrent neural network (RNN) framework. Similar to the Azimuth score, it generates a probability of cutting at the intended on-target. The algorithm only needs the protospacer and PAM sequences as inputs:

```
spacer  <- "ATCGATGCTGATGCTAGATA" #20bp
pam     <- "AGG" #3bp
input   <- paste0(spacer, pam)
results <- getDeepHFScores(input)
```

Users can specify for which Cas9 they wish to score sgRNAs by using the argument `enzyme`: “WT” for Wildtype Cas9 (WT-SpCas9), “HF” for high-fidelity Cas9 (SpCas9-HF), or “ESP” for enhancedCas9 (eSpCas9). For wildtype Cas9, users can also specify the promoter used for expressing sgRNAs using the argument `promoter` (“U6” by default). See `?getDeepHFScores` for more details.

###### 4.1.5 DeepSpCas9

The DeepSpCas9 algorithm is an on-target cutting efficiency prediction algorithm for the SpCas9 nuclease (Kim et al. 2019). Similar to the Azimuth score, it generates a probability of cutting at the intended on-target. 4 nucleotides upstream of the protospacer sequence, and 3 nucleotides downstream of the PAM sequence are needed in top of the protospacer sequence for scoring:

```
flank5 <- "ACCT" #4bp
spacer <- "ATCGATGCTGATGCTAGATA" #20bp
pam    <- "AGG" #3bp
flank3 <- "TTG" #3bp
input  <- paste0(flank5, spacer, pam, flank3)
results <- getDeepSpCas9Scores(input)
```


```
spacer  <- "ATCGATGCTGATGCTAGATA" #20bp
pam     <- "AGG" #3bp
input   <- paste0(spacer, pam)
results <- getDeepHFScores(input)
```

Users can specify for which Cas9 they wish to score sgRNAs by using the argument `enzyme`: “WT” for Wildtype Cas9 (WT-SpCas9), “HF” for high-fidelity Cas9 (SpCas9-HF), or “ESP” for enhancedCas9 (eSpCas9). For wildtype Cas9, users can also specify the promoter used for expressing sgRNAs using the argument `promoter` (“U6” by default). See `?getDeepHFScores` for more details.

##### 4.1.6 CRISPRscan

The CRISPRscan algorithm, also known as the Moreno-Mateos score, is an on-target efficiency method for the SpCas9 nuclease developed for sgRNAs expressed from a T7 promoter, and trained on zebrafish data (Moreno-Mateos et al. 2015). It generates a probability (therefore a score between 0 and 1) that a given sgRNA will cut at its intended target. 6 nucleotides upstream of the protospacer sequence and 6 nucleotides downstream of the PAM sequence are needed for scoring:

```
flank5 <- "ACCTAA" #6bp
spacer <- "ATCGATGCTGATGCTAGATA" #20bp
pam    <- "AGG" #3bp
flank3 <- "TTGAAT" #6bp
input  <- paste0(flank5, spacer, pam, flank3)
results <- getCRISPRscanScores(input)
```

##### 4.1.7 CRISPRater

The CRISPRater algorithm is an on-target efficiency method for the SpCas9 nuclease (Labuhn et al. 2018). It generates a probability (therefore a score between 0 and 1) that a given sgRNA will cut at its intended target. Only the 20bp spacer sequence is required.

```
spacer <- "ATCGATGCTGATGCTAGATA" #20bp
results <- getCRISPRaterScores(spacer)
```

##### 4.1.8 CRISPRai

The CRISPRai algorithm was developed by the Weissman lab to score SpCas9 gRNAs for CRISPRa and CRISPRi applications (Horlbeck et al. 2016), for the human genome. The function `getCrispraiScores` requires several inputs.

First, it requires a data.frame specifying the genomic coordinates of the transcription starting sites (TSSs). An example of such a data.frame is provided in the crisprScore package:

```
head(tssExampleCrispri)
```

```
##       tss_id gene_symbol promoter     transcripts position strand   chr
## 1    A1BG_P1        A1BG       P1 ENST00000596924 58347625      - chr19
## 2    A1BG_P2        A1BG       P2 ENST00000263100 58353463      - chr19
## 3    KRAS_P1        KRAS       P1 ENST00000311936 25250929      - chr12
## 4 SMARCA2_P1     SMARCA2       P1 ENST00000357248  2015347      +  chr9
## 5 SMARCA2_P2     SMARCA2       P2 ENST00000382194  2017615      +  chr9
## 6 SMARCA2_P3     SMARCA2       P3 ENST00000635133  2158470      +  chr9
```

It also requires a data.frame specifying the genomic coordinates of the gRNA sequences to score. An example of such a data.frame is provided in the crisprScore package:

```
head(sgrnaExampleCrispri)
```

```
##     grna_id  tss_id pam_site strand        spacer_19mer
## 1 A1BG_P1_1 A1BG_P1 58347601      - CTCCGGGCGACGTGGAGTG
## 2 A1BG_P1_2 A1BG_P1 58347421      - GGGCACCCAGGAGCGGTAG
## 3 A1BG_P1_3 A1BG_P1 58347624      - TCCACGTCGCCCGGAGCTG
## 4 A1BG_P1_4 A1BG_P1 58347583      - GCAGCGCAGGACGGCATCT
## 5 A1BG_P1_5 A1BG_P1 58347548      - AGCAGCTCGAAGGTGACGT
## 6 A1BG_P2_1 A1BG_P2 58353455      - ATGATGGTCGCGCTCACTC
```

All columns present in `tssExampleCrispri` and `sgrnaExampleCrispri` are mandatory for `getCrispraiScores` to work.

Two additional arguments are required: `fastaFile`, to specify the path of the fasta file of the human reference genome, and `chromatinFiles`, which is a list of length 3 specifying the path of files containing the chromatin accessibility data needed for the algorithm in hg38 coordinates. The chromatin files can be downloaded from Zenodo here. The fasta file for the human genome (hg38) can be downloaded directly from here: https://hgdownload.soe.ucsc.edu/goldenPath/hg38/bigZips/hg38.fa.gz

One can obtain the CRISPRai scores using the following command:

```
results <- getCrispraiScores(tss_df=tssExampleCrispri,
                             sgrna_df=sgrnaExampleCrispri,
                             modality="CRISPRi",
                             fastaFile="your/path/hg38.fa",
                             chromatinFiles=list(mnase="path/to/mnaseFile.bw",
                                                 dnase="path/to/dnaseFile.bw",
                                                 faire="oath/to/faireFile.bw"))
```

The function works identically for CRISPRa applications, with modality replaced by `CRISPRa`.

#### 4.2 Cas12a methods

Different algorithms require different input nucleotide sequences to predict cutting efficiency as illustrated in the figure below.

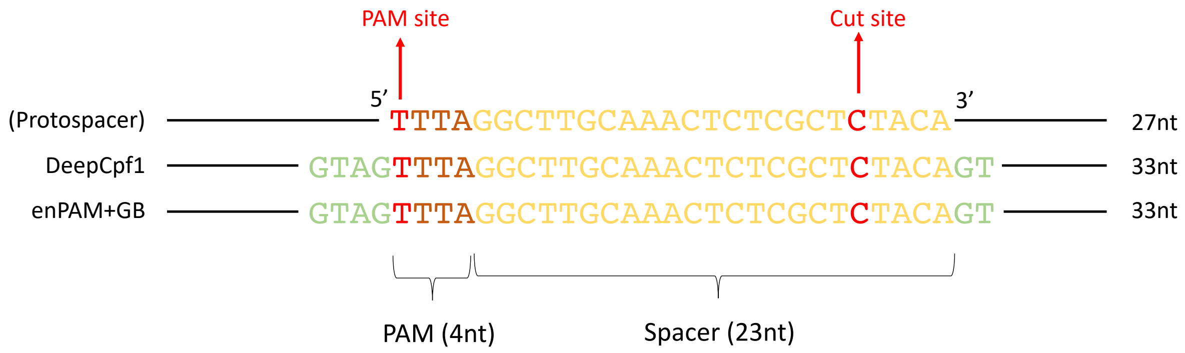


Figure 2: Sequence inputs for Cas12a scoring methods

##### 4.2.1 DeepCpf1 score

The DeepCpf1 algorithm is an on-target cutting efficiency prediction algorithm for the Cas12a nuclease (Kim et al. 2018) using a convolutional neural network (CNN) framework. It generates a score between 0 and 1 to quantify the likelihood of Cas12a to cut for a given sgRNA. 3 nucleotides upstream and 4 nucleotides downstream of the PAM sequence are needed for scoring:

```
flank5 <- "ACC" #3bp
pam    <- "TTTT" #4bp
spacer <- "AATCGATGCTGATGCTAGATATT" #23bp
flank3 <- "AAGT" #4bp
input  <- paste0(flank5, pam, spacer, flank3)
results <- getDeepCpf1Scores(input)
```

##### 4.2.2 enPAM+GB score

The enPAM+GB algorithm is an on-target cutting efficiency prediction algorithm for the enhanced Cas12a (enCas12a) nuclease (DeWeirdt et al. 2020) using a gradient-booster (GB) model. The enCas12a nuclease as an extended set of active PAM sequences in comparison to the wildtype Cas12 nuclease (Kleinstiver et al.

2019), and the enPAM+GB algorithm takes PAM activity into account in the calculation of the final score. It generates a probability (therefore a score between 0 and 1) of a given sgRNA to cut at the intended target. 3 nucleotides upstream of the PAM sequence and 4 nucleotides downstream of the protospacer sequence are needed for scoring:

```
flank5 <- "ACC" #3bp
pam    <- "TTTT" #4bp
spacer <- "AATCGATGCTGATGCTAGATATT" #23bp
flank3 <- "AAGT" #4bp
input  <- paste0(flank5, pam, spacer, flank3)
results <- getEnPAMGBScores(input)
```

#### 4.3 Cas13d methods

##### 4.3.1 CasRxRF

The CasRxRF method was developed to characterize on-target efficiency of the RNA-targeting nuclease RfxCas13d, abbreviated as CasRx (Wessels et al. 2020).

It requires as an input the mRNA sequence targeted by the gRNAs, and returns as an output on-target efficiency scores for all gRNAs targeting the mRNA sequence.

As an example, we predict on-target efficiency for gRNAs targeting the mRNA sequence stored in the file `test.fa`:

```
fasta <- file.path(system.file(package="crisprScore"),
                   "casrxrf/test.fa")
mrnaSequence <- Biostrings::readDNAStringSet(filepath=fasta
                                             format="fasta",
                                             use.names=TRUE)
results <- getCasRxRFScores(mrnaSequence)
```

Note that the function has a default argument `directRepeat` set to `aacccctaccaactggtcggggtttgaaac`, specifying the direct repeat used in the CasRx construct (see (Wessels et al. 2020).) The function also has an argument `binaries` that specifies the file path of the binaries for three programs necessary by the CasRxRF algorithm:

- `RNAfold`: available as part of the ViennaRNA package
- `RNAplfold`: available as part of the ViennaRNA package
- `RNAhybrid`: available as part of the RNAhybrid package

Those programs can be installed from their respective websites: VienneRNA and RNAhybrid.

If the argument is `NULL`, the binaries are assumed to be available on the PATH.

### 5 Off-target specificity scores

For CRISPR knockout systems, off-targeting effects can occur when the CRISPR nuclease tolerates some levels of imperfect complementarity between gRNA spacer sequences and protospacer sequences of the targeted genome. Generally, a greater number of mismatches between spacer and protospacer sequences decreases the likelihood of cleavage by a nuclease, but the nature of the nucleotide substitution can module the likelihood as well. Several off-target specificity scores were developed to predict the likelihood of a nuclease to cut at an unintended off-target site given a position-specific set of nucleotide mismatches.

We provide in `crisprScore` two popular off-target specificity scoring methods for CRISPR/Cas9 knockout systems: the MIT score (Hsu et al. 2013) and the cutting frequency determination (CFD) score (Doench et al. 2016).

#### 5.1 MIT score

The MIT score was an early off-target specificity prediction algorithm developed for the CRISPR/Cas9 system (Hsu et al. 2013). It predicts the likelihood that the Cas9 nuclease will cut at an off-target site using position-specific mismatch tolerance weights. It also takes into consideration the total number of mismatches, as well as the average distance between mismatches. However, it does not take into account the nature of the nucleotide substitutions. The exact formula used to estimate the cutting likelihood is

$$\text{MIT} = \left(\prod_{p \in M} w_p\right) \times \frac{1}{\frac{19-d}{19} \times 4 + 1} \times \frac{1}{m^2}$$

where $M$ is the set of positions for which there is a mismatch between the sgRNA spacer sequence and the off-target sequence, $w_p$ is an experimentally-derived mismatch tolerance weight at position $p$, $d$ is the average distance between mismatches, and $m$ is the total number of mismatches. As the number of mismatches increases, the cutting likelihood decreases. In addition, off-targets with more adjacent mismatches will have a lower cutting likelihood.

The `getMITScores` function takes as argument a character vector of 20bp sequences specifying the spacer sequences of sgRNAs (`spacers` argument), as well as a vector of 20bp sequences representing the protospacer sequences of the putative off-targets in the targeted genome (`protospacers` argument). PAM sequences (`pams`) must also be provided. If only one spacer sequence is provided, it will reused for all provided protospacers.

The following code will generate MIT scores for 3 off-targets with respect to the sgRNA `ATCGATGCTGATGCTAGATA`:

```
spacer   <- "ATCGATGCTGATGCTAGATA"
protospacers  <- c("ACCGATGCTGATGCTAGATA",
                   "ATCGATGCTGATGCTAGATT",
                   "ATCGATGCTGATGCTAGATA")
pams <- c("AGG", "AGG", "AGA")
getMITScores(spacers=spacer,
             protospacers=protospacers,
             pams=pams)
```

```
##                 spacer          protospacer      score
## 1 ATCGATGCTGATGCTAGATA ACCGATGCTGATGCTAGATA 0.68500000
## 2 ATCGATGCTGATGCTAGATA ATCGATGCTGATGCTAGATT 0.00000000
## 3 ATCGATGCTGATGCTAGATA ATCGATGCTGATGCTAGATA 0.06944444
```

#### 5.2 CFD score

The CFD off-target specificity prediction algorithm was initially developed for the CRISPR/Cas9 system, and was shown to be superior to the MIT score (Doench et al. 2016). Unlike the MIT score, position-specific mismatch weights vary according to the nature of the nucleotide substitution (e.g. an A->G mismatch at position 15 has a different weight than an A->T mismatch at position 15).

Similar to the `getMITScores` function, the `getCFDScores` function takes as argument a character vector of 20bp sequences specifying the spacer sequences of sgRNAs (`spacers` argument), as well as a vector of 20bp sequences representing the protospacer sequences of the putative off-targets in the targeted genome (`protospacers` argument). `pams` must also be provided. If only one spacer sequence is provided, it will be used for all provided protospacers.

The following code will generate CFD scores for 3 off-targets with respect to the sgRNA `ATCGATGCTGATGCTAGATA`:

```
spacer   <- "ATCGATGCTGATGCTAGATA"
protospacers  <- c("ACCGATGCTGATGCTAGATA",
                   "ATCGATGCTGATGCTAGATT",
                   "ATCGATGCTGATGCTAGATA")
```

```r
pams <- c("AGG", "AGG", "AGA")
getCFDScores(spacers=spacer,
             protospacers=protospacers,
             pams=pams)
```

```
##                 spacer          protospacer      score
## 1 ATCGATGCTGATGCTAGATA ACCGATGCTGATGCTAGATA 0.85714286
## 2 ATCGATGCTGATGCTAGATA ATCGATGCTGATGCTAGATT 0.60000000
## 3 ATCGATGCTGATGCTAGATA ATCGATGCTGATGCTAGATA 0.06944444
```

### 6 Indel prediction score

#### 6.1 Lindel score (Cas9)

Non-homologous end-joining (NHEJ) plays an important role in double-strand break (DSB) repair of DNA. Error patterns of NHEJ can be strongly biased by sequence context, and several studies have shown that microhomology can be used to predict indels resulting from CRISPR/Cas9-mediated cleavage. Among other useful metrics, the frequency of frameshift-causing indels can be estimated for a given sgRNA.

Lindel (Chen et al. 2019) is a logistic regression model that was trained to use local sequence context to predict the distribution of mutational outcomes. In `crisprScore`, the function `getLindelScores` return the proportion of "frameshifting" indels estimated by Lindel. By chance, assuming a random distribution of indel lengths, frameshifting proportions should be roughly around 0.66. A Lindel score higher than 0.66 indicates a higher than by chance probability that a sgRNA induces a frameshift mutation.

The Lindel algorithm requires nucleotide context around the protospacer sequence; the following full sequence is needed: [13bp upstream flanking sequence][23bp protospacer sequence] [29bp downstream flanking sequence], for a total of 65bp. The function `getLindelScores` takes as inputs such 65bp sequences:

```r
flank5 <- "ACCTTTTAATCGA" #13bp
spacer <- "TGCTGATGCTAGATATTAAG" #20bp
pam    <- "TGG" #3bp
flank3 <- "CTTTTAATCGATGCTGATGCTAGATATTA" #29bp
input <- paste0(flank5, spacer, pam, flank3)
results <- getLindelScores(input)
```

### 7 License

The project as a whole is covered by the MIT license. The code for all underlying Python packages, with their original licenses, can be found in `inst/python`. We made sure that all licenses are compatible with the MIT license and to indicate changes that we have made to the original code.

### 8 Reproducibility

```r
sessionInfo()
```

```
## R version 4.2.1 (2022-06-23)
## Platform: x86_64-apple-darwin17.0 (64-bit)
## Running under: macOS Catalina 10.15.7
##
## Matrix products: default
## BLAS:   /Library/Frameworks/R.framework/Versions/4.2/Resources/lib/libRblas.0.dylib
## LAPACK: /Library/Frameworks/R.framework/Versions/4.2/Resources/lib/libRlapack.dylib
```

```
##
## locale:
## [1] en_US.UTF-8/en_US.UTF-8/en_US.UTF-8/C/en_US.UTF-8/en_US.UTF-8
##
## attached base packages:
## [1] stats     graphics  grDevices utils     datasets  methods   base
##
## other attached packages:
## [1] crisprScore_1.1.14    crisprScoreData_1.1.3 ExperimentHub_2.5.0
## [4] AnnotationHub_3.5.0   BiocFileCache_2.5.0   dbplyr_2.2.1
## [7] BiocGenerics_0.43.1
##
## loaded via a namespace (and not attached):
##  [1] Rcpp_1.0.9                    lattice_0.20-45
##  [3] dir.expiry_1.5.0              png_0.1-7
##  [5] Biostrings_2.65.2             assertthat_0.2.1
##  [7] digest_0.6.29                 utf8_1.2.2
##  [9] mime_0.12                     R6_2.5.1
## [11] GenomeInfoDb_1.33.5           stats4_4.2.1
## [13] RSQLite_2.2.16                evaluate_0.16
## [15] highr_0.9                     httr_1.4.4
## [17] pillar_1.8.1                  basilisk_1.9.3
## [19] zlibbioc_1.43.0               rlang_1.0.4
## [21] curl_4.3.2                    rstudioapi_0.14
## [23] blob_1.2.3                    S4Vectors_0.35.1
## [25] Matrix_1.4-1                  reticulate_1.25
## [27] rmarkdown_2.15.2              stringr_1.4.1
## [29] RCurl_1.98-1.8                bit_4.0.4
## [31] shiny_1.7.2                   compiler_4.2.1
## [33] httpuv_1.6.5                  xfun_0.32
## [35] pkgconfig_2.0.3               htmltools_0.5.3
## [37] tidyselect_1.1.2              KEGGREST_1.37.3
## [39] tibble_3.1.8                  GenomeInfoDbData_1.2.8
## [41] interactiveDisplayBase_1.35.0 IRanges_2.31.2
## [43] randomForest_4.7-1.1          fansi_1.0.3
## [45] crayon_1.5.1                  dplyr_1.0.9
## [47] later_1.3.0                   basilisk.utils_1.9.1
## [49] bitops_1.0-7                  rappdirs_0.3.3
## [51] grid_4.2.1                    jsonlite_1.8.0
## [53] xtable_1.8-4                  lifecycle_1.0.1
## [55] DBI_1.1.3                     magrittr_2.0.3
## [57] cli_3.3.0                     stringi_1.7.8
## [59] cachem_1.0.6                  XVector_0.37.0
## [61] promises_1.2.0.1              ellipsis_0.3.2
## [63] filelock_1.0.2                generics_0.1.3
## [65] vctrs_0.4.1                   tools_4.2.1
## [67] bit64_4.0.5                   Biobase_2.57.1
## [69] glue_1.6.2                    purrr_0.3.4
## [71] BiocVersion_3.16.0            parallel_4.2.1
## [73] fastmap_1.1.0                 yaml_2.3.5
## [75] AnnotationDbi_1.59.1          BiocManager_1.30.18
## [77] memoise_2.0.1                 knitr_1.40
```

### References

Chen, Wei, Aaron McKenna, Jacob Schreiber, Maximilian Haeussler, Yi Yin, Vikram Agarwal, William Stafford Noble, and Jay Shendure. 2019. "Massively Parallel Profiling and Predictive Modeling of the Outcomes of CRISPR/Cas9-Mediated Double-Strand Break Repair." *Nucleic Acids Research* 47 (15): 7989–8003.

DeWeirdt, Peter C, Abby V McGee, Fengyi Zheng, Ifunanya Nwolah, Mudra Hegde, and John G Doench. 2022. "Accounting for Small Variations in the tracrRNA Sequence Improves sgRNA Activity Predictions for CRISPR Screening." *bioRxiv*. https://doi.org/10.1101/2022.06.27.497780.

DeWeirdt, Peter C, Kendall R Sanson, Annabel K Sangree, Mudra Hegde, Ruth E Hanna, Marissa N Feeley, Audrey L Griffith, et al. 2020. "Optimization of AsCas12a for Combinatorial Genetic Screens in Human Cells." *Nature Biotechnology*, 1–11.

Doench, John G, Nicolo Fusi, Meagan Sullender, Mudra Hegde, Emma W Vaimberg, Katherine F Donovan, Ian Smith, et al. 2016. "Optimized sgRNA Design to Maximize Activity and Minimize Off-Target Effects of CRISPR-Cas9." *Nature Biotechnology* 34 (2): 184.

Doench, John G, Ella Hartenian, Daniel B Graham, Zuzana Tothova, Mudra Hegde, Ian Smith, Meagan Sullender, Benjamin L Ebert, Ramnik J Xavier, and David E Root. 2014. "Rational Design of Highly Active sgRNAs for CRISPR-Cas9–Mediated Gene Inactivation." *Nature Biotechnology* 32 (12): 1262–67.

Horlbeck, Max A, Luke A Gilbert, Jacqueline E Villalta, Britt Adamson, Ryan A Pak, Yuwen Chen, Alexander P Fields, et al. 2016. "Compact and Highly Active Next-Generation Libraries for CRISPR-Mediated Gene Repression and Activation." *Elife* 5.

Hsu, Patrick D, David A Scott, Joshua A Weinstein, F Ann Ran, Silvana Konermann, Vineeta Agarwala, Yinqing Li, et al. 2013. "DNA Targeting Specificity of RNA-Guided Cas9 Nucleases." *Nature Biotechnology* 31 (9): 827.

Kim, Hui Kwon, Younggwang Kim, Sungtae Lee, Seonwoo Min, Jung Yoon Bae, Jae Woo Choi, Jinman Park, Dongmin Jung, Sungroh Yoon, and Hyongbum Henry Kim. 2019. "SpCas9 Activity Prediction by DeepSpCas9, a Deep Learning–Based Model with High Generalization Performance." *Science Advances* 5 (11): eaax9249.

Kim, Hui Kwon, Seonwoo Min, Myungjae Song, Soobin Jung, Jae Woo Choi, Younggwang Kim, Sangeun Lee, Sungroh Yoon, and Hyongbum Henry Kim. 2018. "Deep Learning Improves Prediction of CRISPR–Cpf1 Guide RNA Activity." *Nature Biotechnology* 36 (3): 239.

Kleinstiver, Benjamin P, Alexander A Sousa, Russell T Walton, Y Esther Tak, Jonathan Y Hsu, Kendell Clement, Moira M Welch, et al. 2019. "Engineered CRISPR–Cas12a Variants with Increased Activities and Improved Targeting Ranges for Gene, Epigenetic and Base Editing." *Nature Biotechnology* 37 (3): 276–82.

Labuhn, Maurice, Felix F Adams, Michelle Ng, Sabine Knoess, Axel Schambach, Emmanuelle M Charpentier, Adrian Schwarzer, Juan L Mateo, Jan-Henning Klusmann, and Dirk Heckl. 2018. "Refined sgRNA Efficacy Prediction Improves Large-and Small-Scale CRISPR–Cas9 Applications." *Nucleic Acids Research* 46 (3): 1375–85.

Moreno-Mateos, Miguel A, Charles E Vejnar, Jean-Denis Beaudoin, Juan P Fernandez, Emily K Mis, Mustafa K Khokha, and Antonio J Giraldez. 2015. "CRISPRscan: Designing Highly Efficient sgRNAs for CRISPR-Cas9 Targeting in Vivo." *Nature Methods* 12 (10): 982–88.

Wang, Daqi, Chengdong Zhang, Bei Wang, Bin Li, Qiang Wang, Dong Liu, Hongyan Wang, et al. 2019. "Optimized CRISPR Guide RNA Design for Two High-Fidelity Cas9 Variants by Deep Learning." *Nature Communications* 10 (1): 1–14.

Wessels, Hans-Hermann, Alejandro Méndez-Mancilla, Xinyi Guo, Mateusz Legut, Zharko Daniloski, and Neville E Sanjana. 2020. "Massively Parallel Cas13 Screens Reveal Principles for Guide RNA Design." *Nature Biotechnology* 38 (6): 722–27.
