## Supplementary material for "The crisprVerse: a comprehensive Bioconductor ecosystem for the design of CRISPR guide RNAs across nucleases and technologies": Vignette_crisprDesignVerse

2022-08-30

### 1 Installation and getting started

The crisprVerse is a collection of packages for CRISPR guide RNA (gRNA) design that can easily be installed with the `crisprVerse` package. This provides a convenient way of downloading and installing all crisprVerse packages with a single R command.

The package can be installed from the Bioconductor devel branch using the following commands in an R session:

```
if (!requireNamespace("BiocManager", quietly = TRUE))
    install.packages("BiocManager")

BiocManager::install(version="devel")
BiocManager::install("crisprVerse")
```

The core crisprVerse includes the packages that are commonly used for gRNA design, and are attached when you attach the `crisprVerse` package:

```
library(crisprVerse)
```

You can check that all crisprVerse packages are up-to-date with the function `crisprVerse_update()`.

### 2 Components

The following packages are installed and loaded with the `crisprVerse` package:

- *crisprBase* to specify and manipulate CRISPR nucleases.
- *crisprBowtie* to perform gRNA spacer sequence alignment with Bowtie.
- *crisprScore* to annotate gRNAs with on-target and off-target scores.
- *crisprDesign* to design and manipulate gRNAs with `GuideSet` objects.

### 3 Reproducibility

```
sessionInfo()
```

```
## R version 4.2.1 (2022-06-23)
## Platform: x86_64-apple-darwin17.0 (64-bit)
## Running under: macOS Catalina 10.15.7
##
## Matrix products: default
## BLAS:   /Library/Frameworks/R.framework/Versions/4.2/Resources/lib/libRblas.0.dylib
## LAPACK: /Library/Frameworks/R.framework/Versions/4.2/Resources/lib/libRlapack.dylib
##
```

```
## locale:
## [1] en_US.UTF-8/en_US.UTF-8/en_US.UTF-8/C/en_US.UTF-8/en_US.UTF-8
##
## attached base packages:
## [1] stats     graphics  grDevices utils     datasets  methods   base
##
## other attached packages:
##  [1] crisprDesign_0.99.134 crisprScore_1.1.14    crisprScoreData_1.1.3
##  [4] ExperimentHub_2.5.0   AnnotationHub_3.5.0   BiocFileCache_2.5.0
##  [7] dbplyr_2.2.1          BiocGenerics_0.43.1   crisprBowtie_1.1.1
## [10] crisprBase_1.1.5      crisprVerse_0.99.8
##
## loaded via a namespace (and not attached):
##   [1] bitops_1.0-7                  matrixStats_0.62.0
##   [3] bit64_4.0.5                   filelock_1.0.2
##   [5] progress_1.2.2                httr_1.4.4
##   [7] GenomeInfoDb_1.33.5           tools_4.2.1
##   [9] utf8_1.2.2                    R6_2.5.1
##  [11] DBI_1.1.3                     tidyselect_1.1.2
##  [13] prettyunits_1.1.1             bit_4.0.4
##  [15] curl_4.3.2                    compiler_4.2.1
##  [17] cli_3.3.0                     Biobase_2.57.1
##  [19] basilisk.utils_1.9.1          xml2_1.3.3
##  [21] DelayedArray_0.23.1           rtracklayer_1.57.0
##  [23] randomForest_4.7-1.1          readr_2.1.2
##  [25] rappdirs_0.3.3                stringr_1.4.1
##  [27] digest_0.6.29                 Rsamtools_2.13.4
##  [29] rmarkdown_2.15.2              basilisk_1.9.3
##  [31] XVector_0.37.0                pkgconfig_2.0.3
##  [33] htmltools_0.5.3               MatrixGenerics_1.9.1
##  [35] fastmap_1.1.0                 BSgenome_1.65.2
##  [37] rlang_1.0.4                   rstudioapi_0.14
##  [39] RSQLite_2.2.16                shiny_1.7.2
##  [41] BiocIO_1.7.1                  generics_0.1.3
##  [43] jsonlite_1.8.0                BiocParallel_1.31.12
##  [45] dplyr_1.0.9                   VariantAnnotation_1.43.3
##  [47] RCurl_1.98-1.8                magrittr_2.0.3
##  [49] GenomeInfoDbData_1.2.8        Matrix_1.4-1
##  [51] Rcpp_1.0.9                    S4Vectors_0.35.1
##  [53] fansi_1.0.3                   reticulate_1.25
##  [55] Rbowtie_1.37.0                lifecycle_1.0.1
##  [57] stringi_1.7.8                 yaml_2.3.5
##  [59] SummarizedExperiment_1.27.1   zlibbioc_1.43.0
##  [61] grid_4.2.1                    blob_1.2.3
##  [63] promises_1.2.0.1              parallel_4.2.1
##  [65] crayon_1.5.1                  dir.expiry_1.5.0
##  [67] lattice_0.20-45               Biostrings_2.65.2
##  [69] GenomicFeatures_1.49.6        hms_1.1.2
##  [71] KEGGREST_1.37.3               knitr_1.40
##  [73] pillar_1.8.1                  GenomicRanges_1.49.1
##  [75] rjson_0.2.21                  codetools_0.2-18
##  [77] biomaRt_2.53.2                stats4_4.2.1
##  [79] BiocVersion_3.16.0            XML_3.99-0.10
##  [81] glue_1.6.2                    evaluate_0.16
```

```
##  [83] BiocManager_1.30.18           httpuv_1.6.5
##  [85] png_0.1-7                     vctrs_0.4.1
##  [87] tzdb_0.3.0                    purrr_0.3.4
##  [89] assertthat_0.2.1              cachem_1.0.6
##  [91] xfun_0.32                     mime_0.12
##  [93] xtable_1.8-4                  restfulr_0.0.15
##  [95] later_1.3.0                   tibble_3.1.8
##  [97] GenomicAlignments_1.33.1      AnnotationDbi_1.59.1
##  [99] memoise_2.0.1                 IRanges_2.31.2
## [101] interactiveDisplayBase_1.35.0 ellipsis_0.3.2
## [103] BiocStyle_2.25.0
```
