## Supplementary material for "The crisprVerse: a comprehensive Bioconductor ecosystem for the design of CRISPR guide RNAs across nucleases and technologies": Vignette_crisprViz

### Introduction to crisprViz

Luke Hoberecht

2022-08-27

### 1 Introduction

The `crisprViz` package enables the graphical interpretation of `GuideSet` objects from the `crisprDesign` package by plotting guide RNA (gRNA) cutting locations against their target gene or other genomic region. These genomic plots are constructed using the `Gviz` package from Bioconductor.

This vignette walks through several use cases that demonstrate the range of and how to use plotting functions in the `crisprViz` package. This vignette also uses the `crisprDesign` package to manipulate `GuideSet` objects in conjunction with plotting in the process of gRNA design. For more information about the `crisprDesign` package see [vignettes].

### 2 Installation and getting started

#### 2.1 Software requirements

##### 2.1.1 OS Requirements

This package is supported for macOS, Linux and Windows machines. Packages were developed and tested on R version 4.2.1.

#### 2.2 Installation from Bioconductor

`crisprViz` can be installed from Bioconductor using the following commands in a fresh R session:

```
install.packages("BiocManager")
BiocManager::install("crisprViz")
```

### 3 Use cases

All examples in this vignette will use human genome assembly `hg38` from the `BSgenome.Hsapiens.UCSC.hg38` package and gene model coordinates from Ensembl release 104. We begin by loading the necessary packages.

```
library(BSgenome.Hsapiens.UCSC.hg38)
library(crisprDesign)
library(crisprViz)
```

#### 3.1 Visualizing the best gRNAs for a given gene

Suppose we want to design the four best gRNAs using the SpCas9 CRISPR nuclease to knockout the human KRAS gene. To have the greatest impact on gene function we want to prioritize gRNAs that have greater isoform coverage, and target closer to the 5' end of the CDS.

Let's load a precomputed `GuideSet` object containing all possible gRNAs targeting the the CDS of KRAS, and a `GRangesList` object describing the gene model for KRAS.

```r
data("krasGuideSet", package="crisprViz")
data("krasGeneModel", package="crisprViz")
length(krasGuideSet) # number of candidate gRNAs
```

```
## [1] 52
```

For how to design such gRNAs, see the `crisprDesign` package. Before we plot all of our candidate gRNAs, let's first generate a simple plot with a few gRNAs to familiarize ourselves with some plot components and options.

```r
plotGuideSet(krasGuideSet[1:4],
             geneModel=krasGeneModel,
             targetGene="KRAS")
```

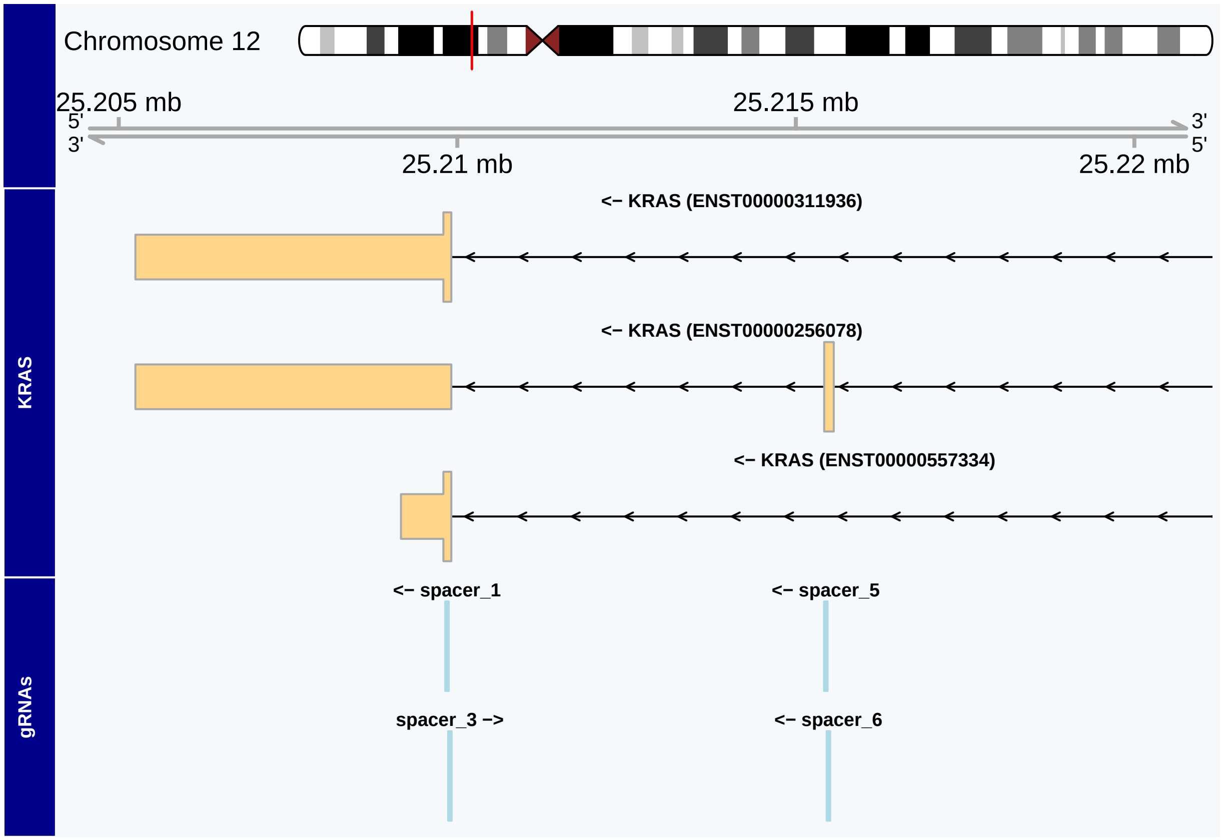


There are a few things to note here.

- The ideogram track and genome axis track are at the top of our plot and give us coordinate information.
- Our `targetGene` KRAS is plotted next, using coordinates from the provided gene model `krasGeneModel`, followed by our spacer subset. The name of each track is given on the left.
- The strand information for each track is included in the label: `<-` for reverse strand and `->` for forward strand.
- While we can identify which exon each spacer targets (which may be sufficient), the plot window is too large to provide further information.
- The plot only shows the 3' end of KRAS, rather than the entire gene.

This last point is important: the default plot window is set by the spacers' ranges in the input `GuideSet` object. We can manually adjust this window by using the `from`, `to`, `extend.left`, and `extend.right` arguments. Here is the same plot adjusted to show the whole KRAS gene, which also reveals an additional isoform that is not targeted by any spacer in this example subset.

```r
from <- min(start(krasGeneModel$transcripts))
to <- max(end(krasGeneModel$transcripts))
plotGuideSet(krasGuideSet[1:4],
             geneModel=krasGeneModel,
             targetGene="KRAS",
             from=from,
             to=to,
             extend.left=1000,
             extend.right=1000)
```

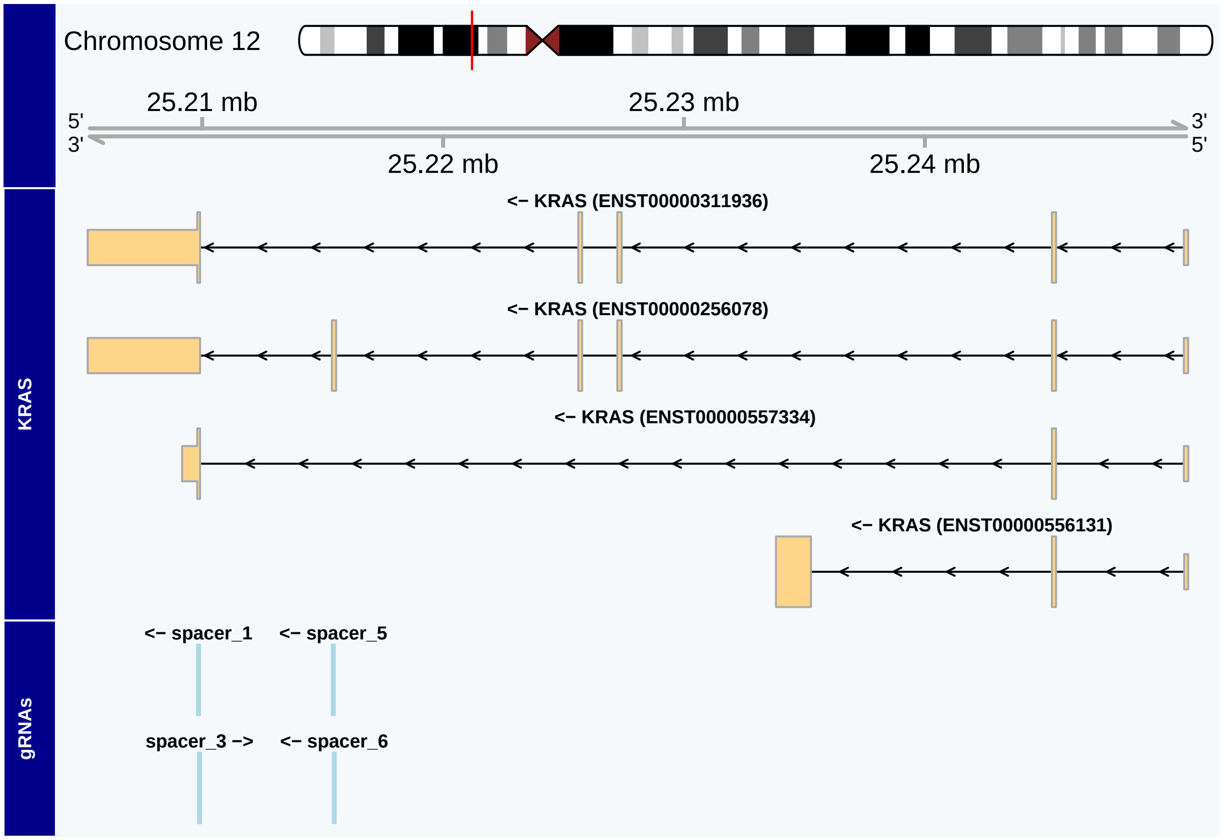


As calculated above, there are a total of 52 candidate gRNAs targeting the CDS of KRAS. Including all of them could crowd the plot space, making it difficult to interpret. To alleviate this we can hide the gRNA labels by setting the `showGuideLabels` argument to `FALSE`.

```r
plotGuideSet(krasGuideSet,
             geneModel=krasGeneModel,
             targetGene="KRAS",
             showGuideLabels=FALSE,
             from=from,
             to=to,
             extend.left=1000,
             extend.right=1000)
```

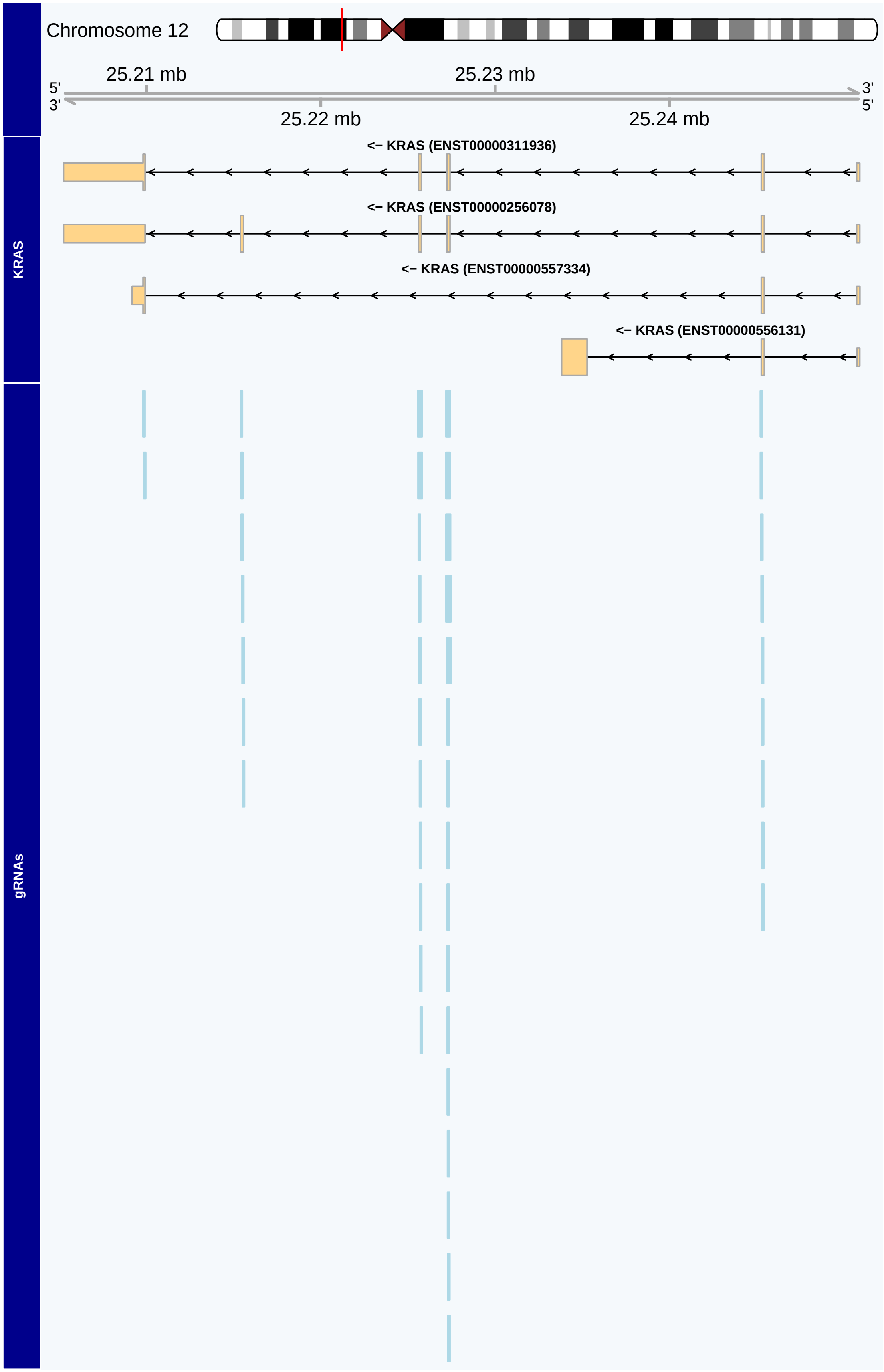
Chromosome 12
25.21 mb
25.23 mb
5'
3'
3'
5'
25.22 mb
25.24 mb
<– KRAS (ENST00000311936)
<– KRAS (ENST00000256078)
<– KRAS (ENST00000557334)
<– KRAS (ENST00000556131)
KRAS
gRNAs

At the gene level, the plot window is too large to discern details for each spacer target. However, we can see five distinct clusters of spacer target locations that cover the CDS of KRAS. The spacers in the 5'-most cluster (on the reverse strand) target the only coding region of the gene that is expressed by all isoforms, making it an ideal target for our scenario.

We can see which gRNAs target this region by returning `showGuideLabels` to its default value of `TRUE`, and by adjusting the plot window to focus on our exon of interest.

```
# new window range around target exon
targetExon <- queryTxObject(krasGeneModel,
                            featureType="cds",
                            queryColumn="exon_id",
                            queryValue="ENSE00000936617")
targetExon <- unique(targetExon)
from <- start(targetExon)
to <- end(targetExon)
plotGuideSet(krasGuideSet,
             geneModel=krasGeneModel,
             targetGene="KRAS",
             from=from,
             to=to,
             extend.left=20,
             extend.right=20)
```

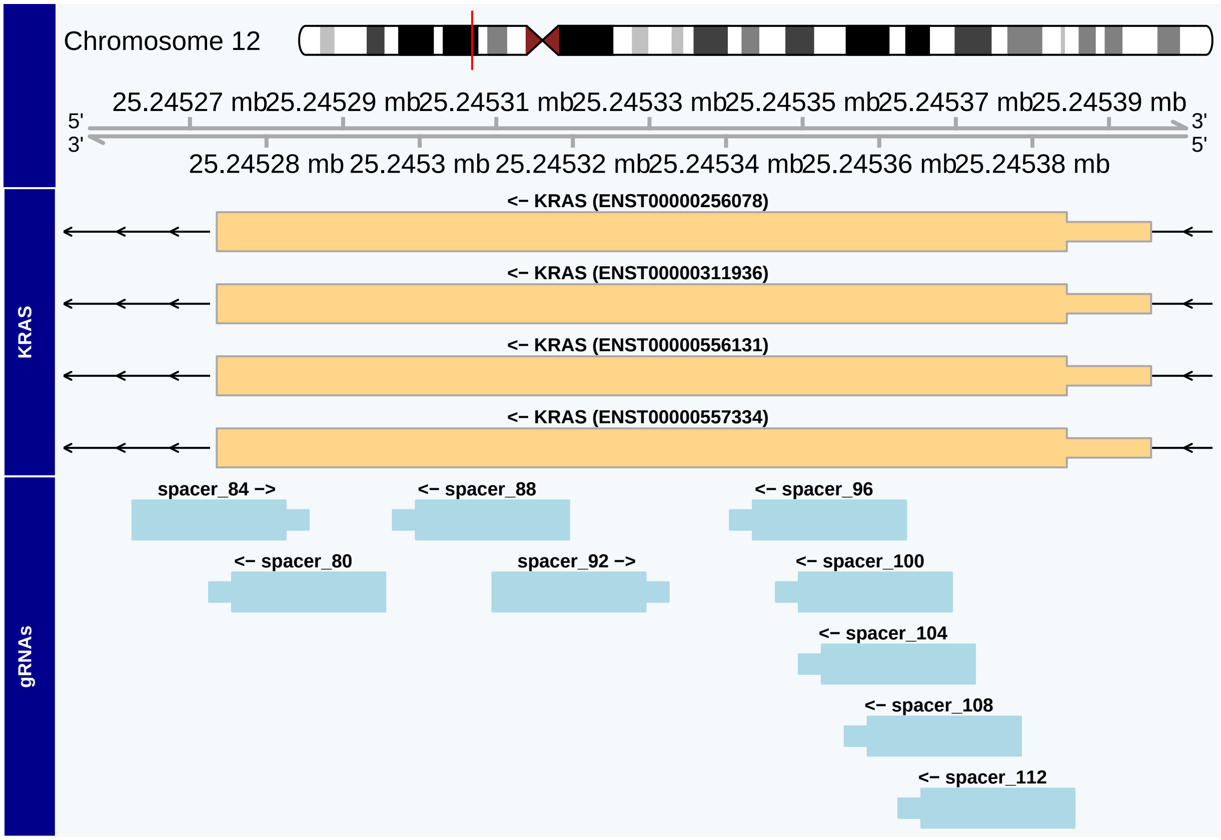


At this resolution we can get a much better idea of spacer location and orientation. In particular, the PAM sequence is visible as a narrow box on the 3' side of our protospacer sequences. We can also distinctly see which spacer targets overlap each other–it may be best to avoid pairing such spacers in some applications lest they sterically interfere with each other.

If we have many gRNA targets in a smaller window and are not concerned with overlaps, we can configure the plot to only show the `pam_site`, rather than the entire protospacer and PAM sequence, by setting `pamSiteOnly` to `TRUE`.

```
plotGuideSet(krasGuideSet,
             geneModel=krasGeneModel,
             targetGene="KRAS",
             from=from,
             to=to,
             extend.left=20,
             extend.right=20,
             pamSiteOnly=TRUE)
```

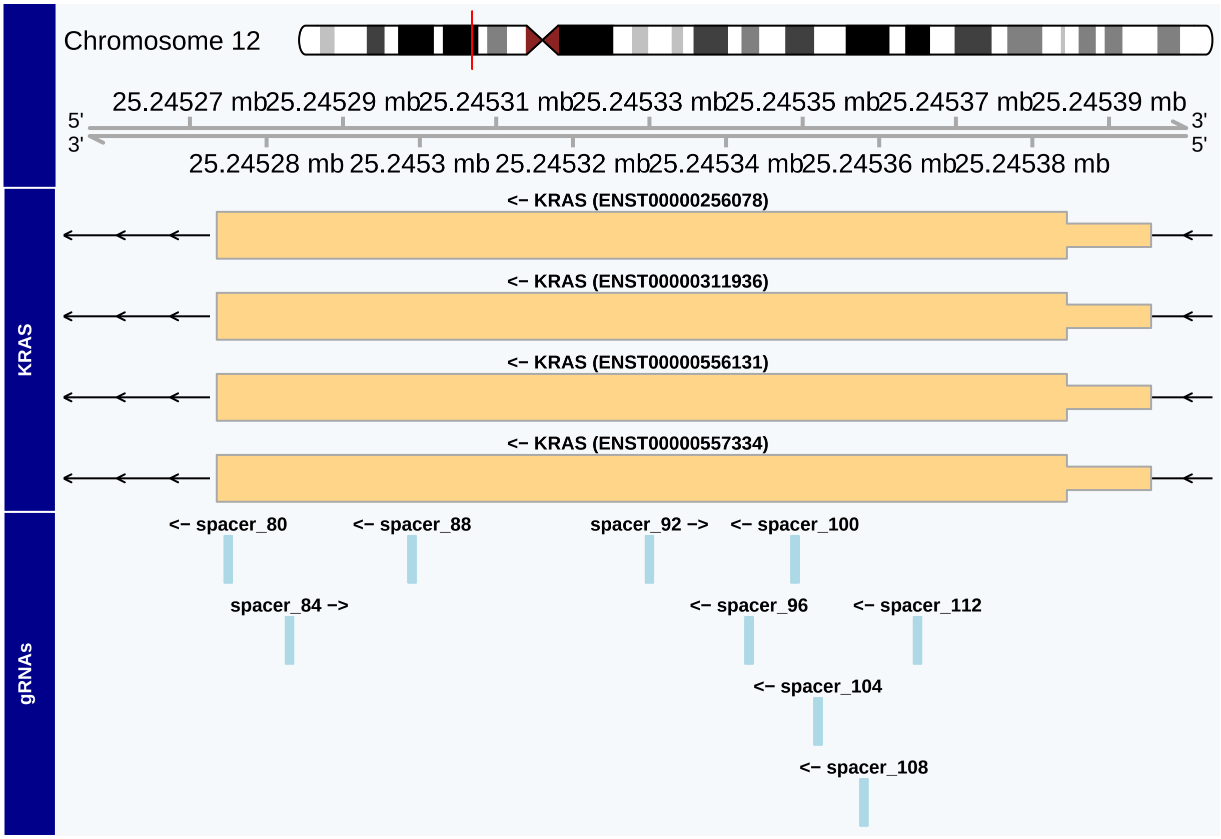


Let's filter our `GuideSet` by the spacer names in the plot then pass an on-target score column in our `GuideSet` to `onTargetScores` to color the spacers according to that score, with darker blue colors indicating higher scores. Note that for this plot we need not provide values for `from` and `to`, as the plot window adjusts to our filtered `GuideSet`.

```
selectedGuides <- c("spacer_80", "spacer_84", "spacer_88", "spacer_92",
                    "spacer_96", "spacer_100", "spacer_104", "spacer_108",
                    "spacer_112")
candidateGuides <- krasGuideSet[selectedGuides]
plotGuideSet(candidateGuides,
             geneModel=krasGeneModel,
             targetGene="KRAS",
             onTargetScore="score_deephf")
```

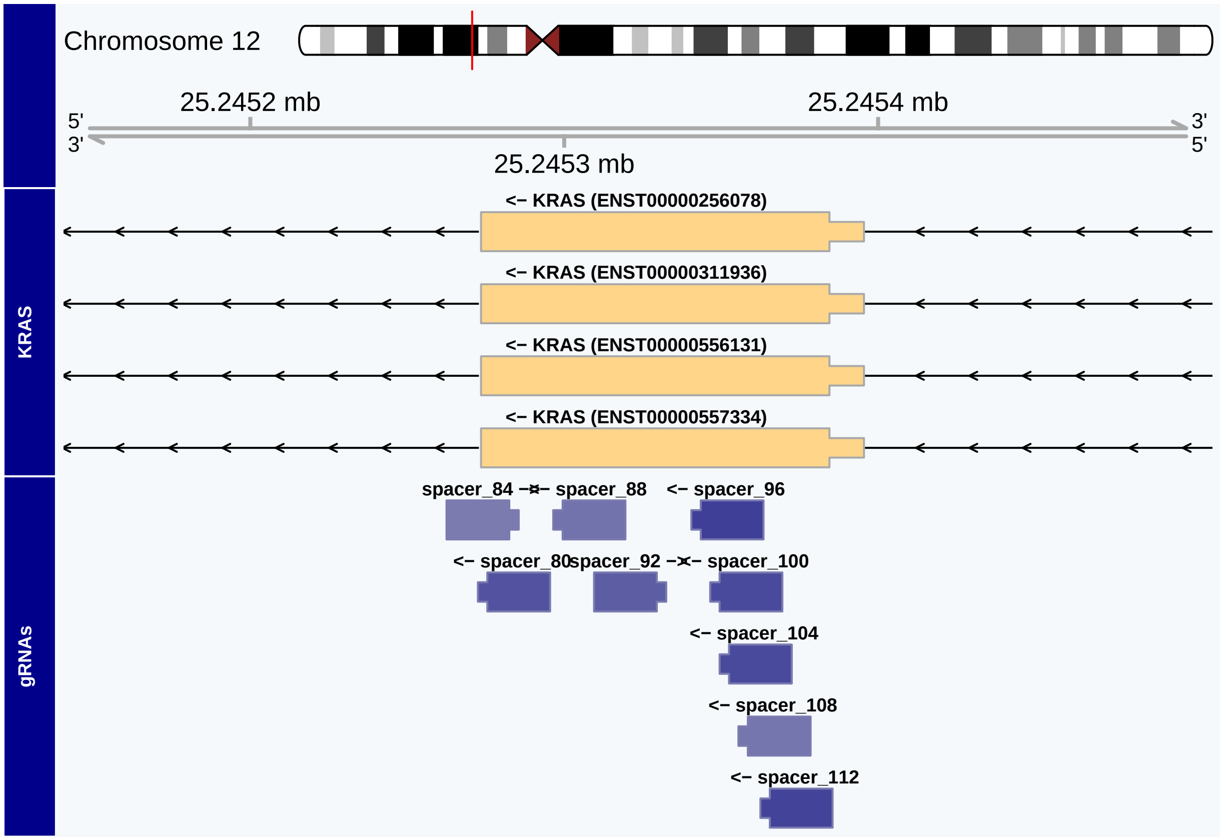


#### 3.2 Plotting for precision targeting

For a given CRISPR application, the target region may consist of only several base pairs rather than an exon or entire gene CDS. In these instances it may be important to know exactly where the gRNAs target, and plots of gRNAs must be at a resolution capable of distinguishing individual bases. This is often the case for CRISPR base editor (CRISPRbe) applications, as the editing window for each gRNA is narrow and the results are specific to each target sequence.

In this example, we will zoom in on a few gRNAs targeting the 5' end of the human GPR21 gene. We want our plot to include genomic sequence information so we will set the `bsgenome` argument to the same `BSgenome` object we used to create our `GuideSet`.

First, we load the precomputed `GuideSet` and gene model objects for GPR21,

```
data("gpr21GuideSet", package="crisprViz")
data("gpr21GeneModel", package="crisprViz")
```

and then plot the gRNAs.

```
plotGuideSet(gpr21GuideSet,
             geneModel=gpr21GeneModel,
             targetGene="GPR21",
             bsgenome=BSgenome.Hsapiens.UCSC.hg38,
             margin=0.3)
```

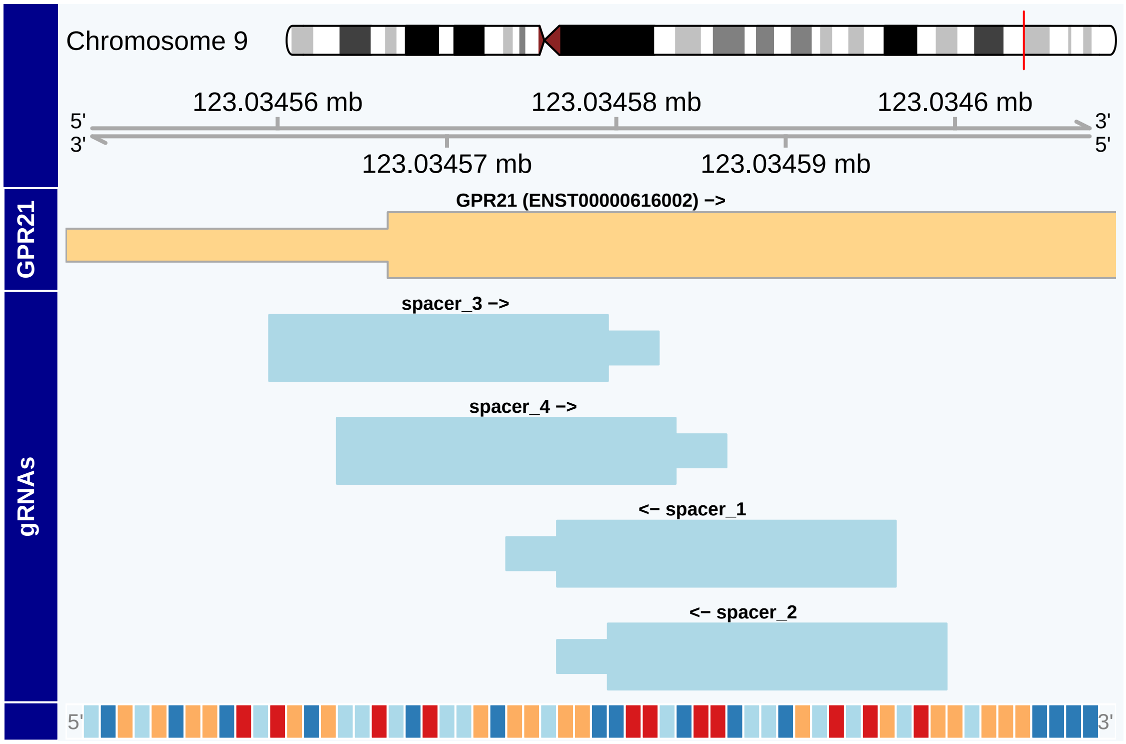


The genomic sequence is given at the bottom of the plot as color-coded boxes. The color scheme for describing the nucleotides is given in the **biovizBase** package. If the plot has sufficient space, it will display nucleotide symbols rather than boxes. We can accomplish this by plotting a narrower range or by increasing the width of our plot space (see "Setting plot size" section).

The plot above was generated with a plot space width of 6 inches; here's the same plot after we increase the width to 10 inches:

```
# increase plot width from 6" to 10"
plotGuideSet(gpr21GuideSet,
             geneModel=gpr21GeneModel,
             targetGene="GPR21",
             bsgenome=BSgenome.Hsapiens.UCSC.hg38,
             margin=0.3)
```

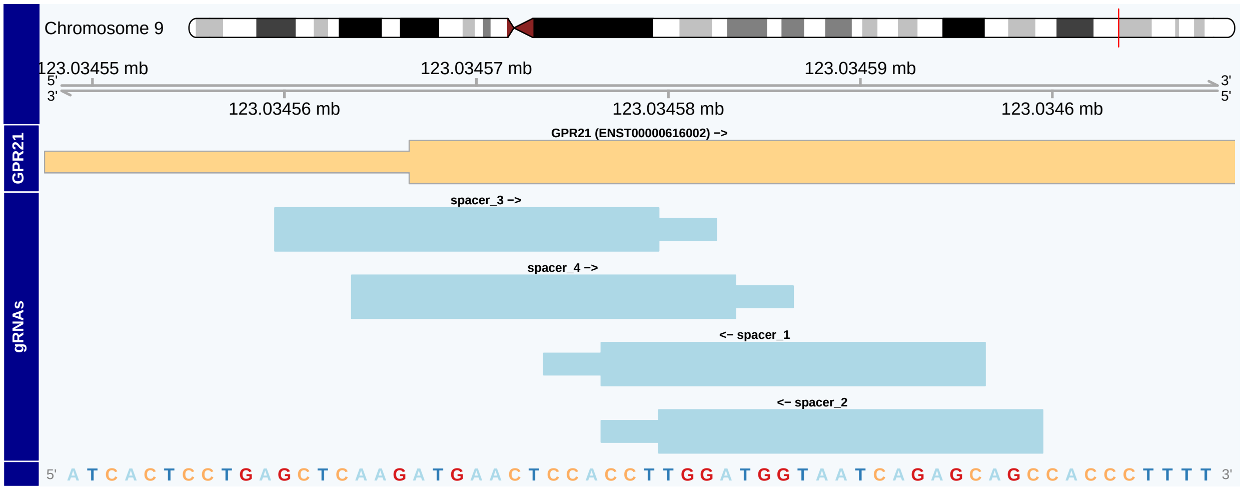

#### 3.3 CRISPRa and adding genomic annotations

In this scenario we want to increase expression of the human MMP7 gene via CRISPR activation (CRISPRa). We will use the SpCas9 CRISPR nuclease.

```
data("mmp7GuideSet", package="crisprViz")
data("mmp7GeneModel", package="crisprViz")
```

The `GuideSet` contains candidate gRNAs in the 2kb window immediately upstream of the TSS of MMP7. We will also use a `GRanges` object containing repeat elements in this region:

```
data("repeats", package="crisprViz")
```

Let's begin by plotting our `GuideSet`, and adding a track of repeat elements using the `annotations` argument. Our `guideSet` also contains SNP annotation, which we would also prefer our gRNAs to not overlap. To include a SNP annotation track, we will set `includeSNPTrack=TRUE` (default).

```
from <- min(start(mmp7GuideSet))
to <- max(end(mmp7GuideSet))
plotGuideSet(mmp7GuideSet,
             geneModel=mmp7GeneModel,
             targetGene="MMP7",
             guideStacking="dense",
             annotations=list(Repeats=repeats),
             pamSiteOnly=TRUE,
             from=from,
             to=to,
             extend.left=600,
             extend.right=100,
             includeSNPTrack=TRUE)
```

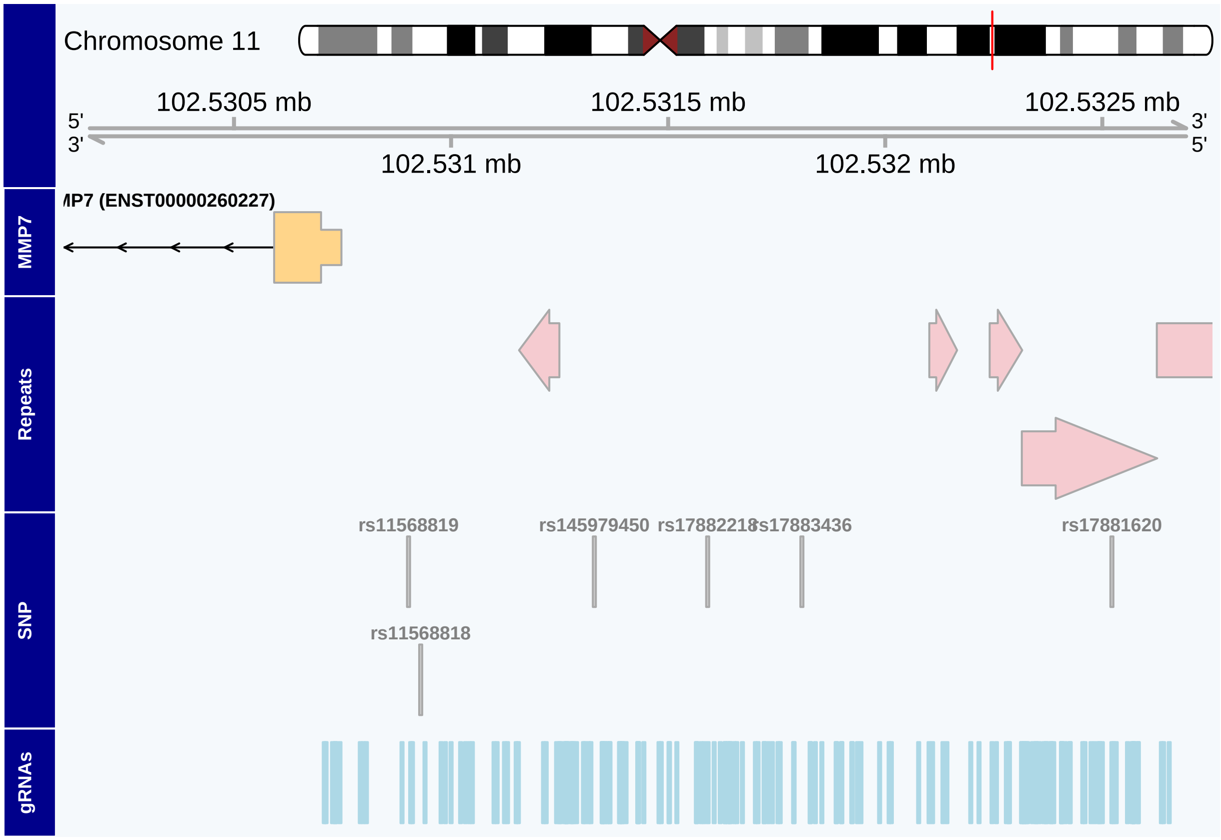


Some of our candidate gRNAs target repeat elements and likely target a large number of loci in the genome, potentially causing unintended effects, or overlap with SNPs, which can reduce its activity. Let's remove these gRNAs and regenerate the plot.

```
filteredGuideSet <- crisprDesign::removeRepeats(mmp7GuideSet,
                                                gr.repeats=repeats)
filteredGuideSet <- filteredGuideSet[!filteredGuideSet$hasSNP]
plotGuideSet(filteredGuideSet,
             geneModel=mmp7GeneModel,
             targetGene="MMP7",
             guideStacking="dense",
             annotations=list(Repeats=repeats),
             pamSiteOnly=TRUE,
             from=from,
             to=to,
             extend.left=600,
             extend.right=100,
             includeSNPTrack=TRUE)
```

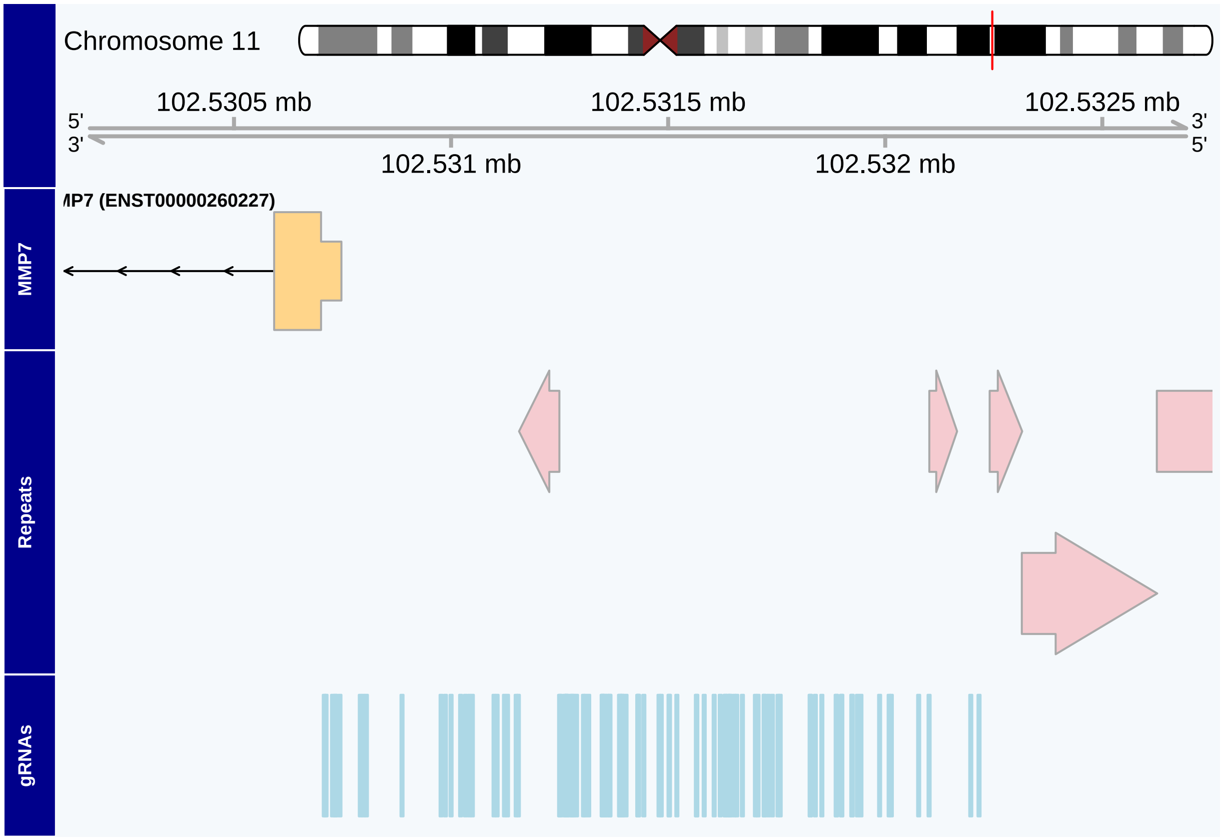


Note how removing gRNAs that overlap SNPs from our `GuideSet` also removed the SNP track. To prevent plotting an empty track, `plotGuideSet` will only include a SNPs track if at least one gRNA includes SNP annotation (i.e. overlaps a SNP).

Conversely, there are specific genomic regions that would be beneficial to target, such as CAGE peaks and DNase I Hypersensitivity tracks. We show in the `inst\scripts` folder how to obtain such data from the Bioconductor package `AnnotationHub`, but for the sake of time, we have precomputed those objects and they can be loaded from the `crisprViz` package directly:

```
data("cage", package="crisprViz")
data("dnase", package="crisprViz")
```

We now plot gRNAs alongside with those two tracks:

```
plotGuideSet(filteredGuideSet,
             geneModel=mmp7GeneModel,
             targetGene="MMP7",
             guideStacking="dense",
             annotations=list(CAGE=cage, DNase=dnase),
             pamSiteOnly=TRUE,
             from=from,
             to=to,
             extend.left=600,
             extend.right=100)
```

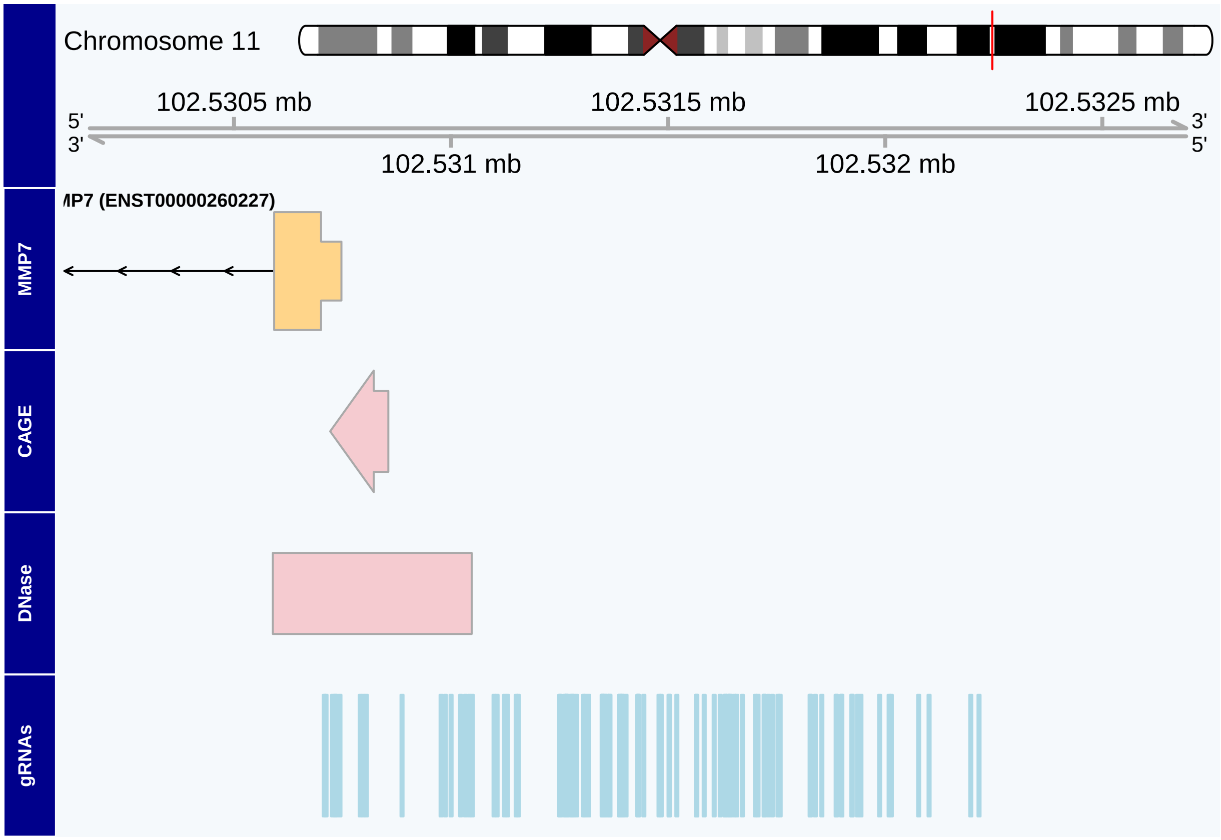


Let's filter our `GuideSet` for guides overlapping the plotted DNase site then regenerate the plot.

```
# filter GuideSet for gRNAs overlapping DNase track
overlaps <- findOverlaps(filteredGuideSet, dnase, ignore.strand=TRUE)
finalGuideSet <- filteredGuideSet[queryHits(overlaps)]
plotGuideSet(finalGuideSet,
             geneModel=mmp7GeneModel,
             targetGene="MMP7",
             guideStacking="dense",
             annotations=list(CAGE=cage, DNase=dnase),
             pamSiteOnly=TRUE,
             margin=0.4)
```

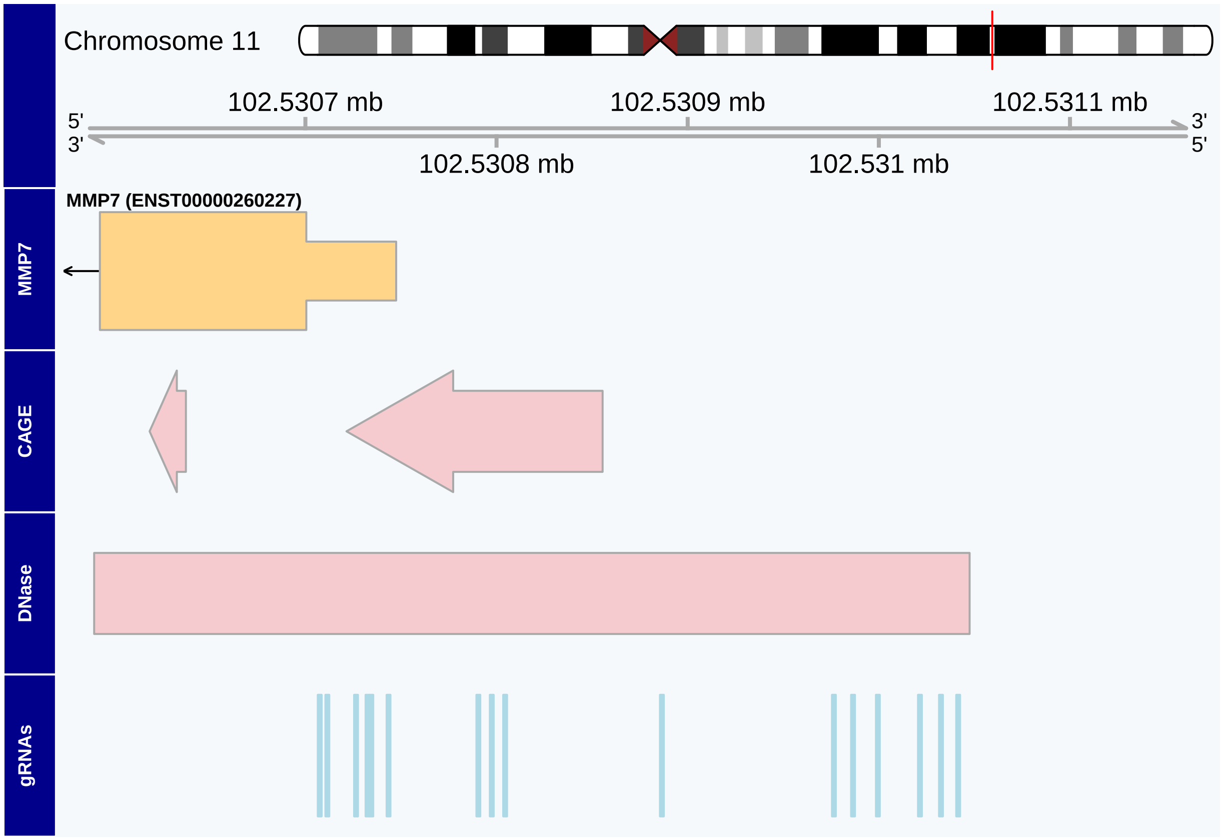


#### 3.4 Comparing multiple GuideSets targeting the same region

The choice of the CRISPR nuclease can be influenced by the abundance of PAM sequences recognized by a given nuclease in the target region. For example, we would expect AT-rich regions to have fewer possible targets for the SpCas9 nuclease, whose PAM is NGG. In these regions, the CRISPR nuclease AsCas12a, whose PAM is TTTV, may prove more appropriate. Given multiple `GuideSets` targeting the same region, we can compare the gRNAs of each in the same plot using `plotMultipleGuideSets`.

Here, we pass our `GuideSets` targeting an exon in the human gene LTN1 in a named list. Note that there are no available options for displaying guide labels or guide stacking, and only the PAM sites are plotted. We will also add a track to monitor the percent GC content (using a window roughly the length of our protospacers). Not surprisingly, this AT-rich region has fewer targets for SpCas9 compared to AsCas12a. (Note: when plotting several GuideSets you may need to increase the height of the plot space in order for the track names to appear on the left side; see “Setting plot size” below.)

We first load the precomputed `GuideSet` objects:

```
data("cas9GuideSet", package="crisprViz")
data("cas12aGuideSet", package="crisprViz")
data("ltn1GeneModel", package="crisprViz")
```

```
plotMultipleGuideSets(list(SpCas9=cas9GuideSet, AsCas12a=cas12aGuideSet),
                      geneModel=ltn1GeneModel,
                      targetGene="LTN1",
                      bsgenome=BSgenome.Hsapiens.UCSC.hg38,
                      margin=0.2,
                      gcWindow=10)
```

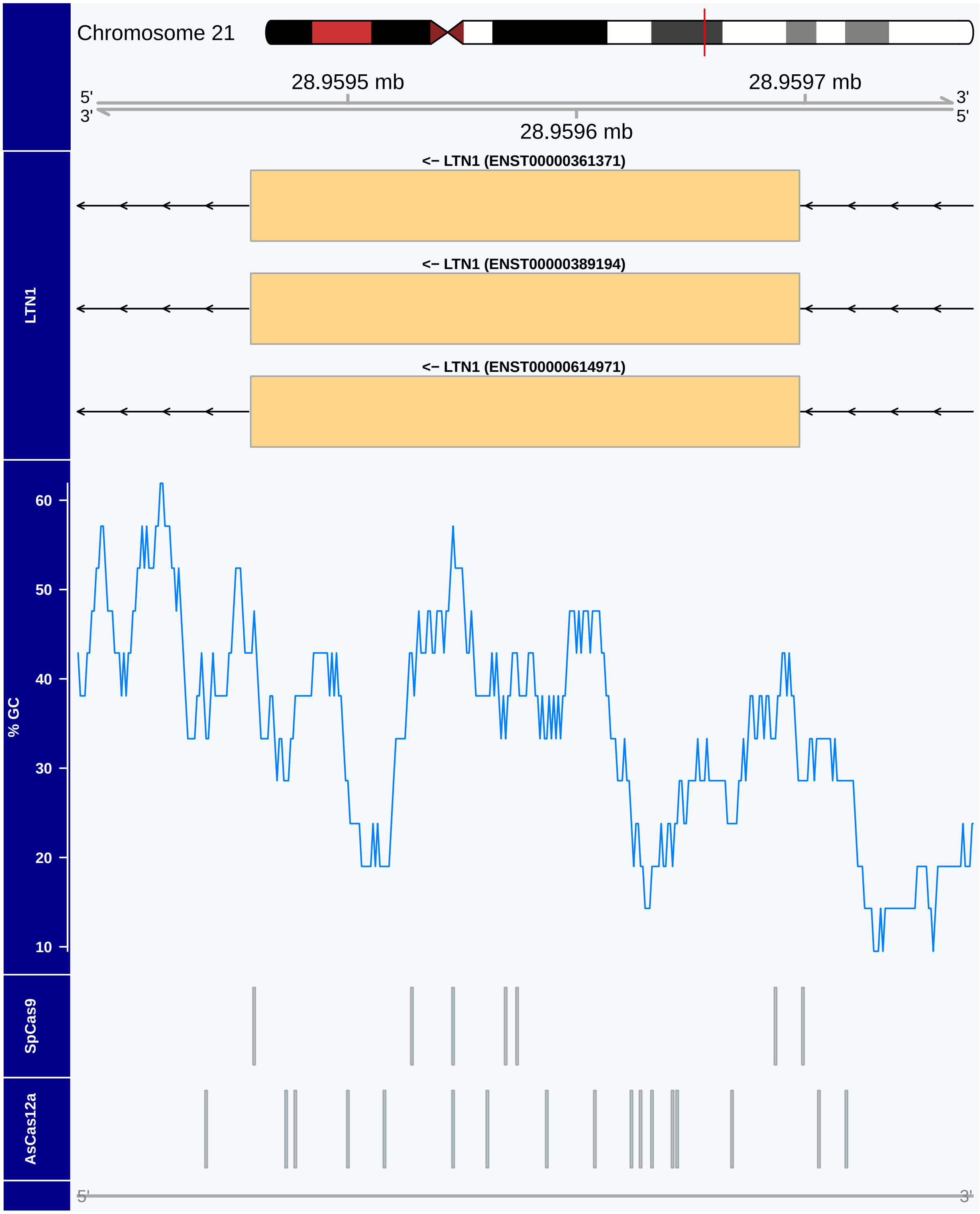


### 4 Setting plot size

Plots with many gene isoforms and/or gRNAs may require more space to render than is allotted by your graphical device's default settings, resulting in an error. One solution, depending on your graphical device, is offered by the **grDevices** package.

Here is an example using macOS Quartz device:

```
grDevices::quartz("Example plot", width=6, height=7)
# plot function
```

### 5 Session Info

```
sessionInfo()
```

```
## R version 4.2.1 (2022-06-23)
## Platform: x86_64-apple-darwin17.0 (64-bit)
## Running under: macOS Catalina 10.15.7
##
## Matrix products: default
## BLAS:   /Library/Frameworks/R.framework/Versions/4.2/Resources/lib/libRblas.0.dylib
## LAPACK: /Library/Frameworks/R.framework/Versions/4.2/Resources/lib/libRlapack.dylib
##
## locale:
## [1] en_US.UTF-8/en_US.UTF-8/en_US.UTF-8/C/en_US.UTF-8/en_US.UTF-8
##
## attached base packages:
## [1] stats4    stats     graphics  grDevices utils     datasets  methods
## [8] base
##
## other attached packages:
##  [1] crisprViz_0.99.18                 crisprDesign_0.99.134
##  [3] crisprBase_1.1.5                  BSgenome.Hsapiens.UCSC.hg38_1.4.4
##  [5] BSgenome_1.65.2                   rtracklayer_1.57.0
##  [7] Biostrings_2.65.2                 XVector_0.37.0
##  [9] GenomicRanges_1.49.1              GenomeInfoDb_1.33.5
## [11] IRanges_2.31.2                    S4Vectors_0.35.1
## [13] BiocGenerics_0.43.1
##
## loaded via a namespace (and not attached):
##   [1] backports_1.4.1              Hmisc_4.7-1
##   [3] AnnotationHub_3.5.0          BiocFileCache_2.5.0
##   [5] lazyeval_0.2.2               splines_4.2.1
##   [7] BiocParallel_1.31.12         ggplot2_3.3.6
##   [9] digest_0.6.29                ensembldb_2.21.3
##  [11] htmltools_0.5.3              fansi_1.0.3
##  [13] checkmate_2.1.0              magrittr_2.0.3
##  [15] memoise_2.0.1                cluster_2.1.4
##  [17] tzdb_0.3.0                   readr_2.1.2
##  [19] matrixStats_0.62.0           prettyunits_1.1.1
##  [21] jpeg_0.1-9                   colorspace_2.0-3
##  [23] blob_1.2.3                   rappdirs_0.3.3
##  [25] crisprScoreData_1.1.3        xfun_0.32
##  [27] dplyr_1.0.9                  crayon_1.5.1
##  [29] RCurl_1.98-1.8               jsonlite_1.8.0
##  [31] survival_3.4-0               VariantAnnotation_1.43.3
##  [33] glue_1.6.2                   gtable_0.3.0
##  [35] zlibbioc_1.43.0              DelayedArray_0.23.1
```

```
##  [37] scales_1.2.1                  DBI_1.1.3
##  [39] Rcpp_1.0.9                    htmlTable_2.4.1
##  [41] xtable_1.8-4                  progress_1.2.2
##  [43] reticulate_1.25               foreign_0.8-82
##  [45] bit_4.0.4                     Formula_1.2-4
##  [47] htmlwidgets_1.5.4             httr_1.4.4
##  [49] dir.expiry_1.5.0              RColorBrewer_1.1-3
##  [51] ellipsis_0.3.2                pkgconfig_2.0.3
##  [53] XML_3.99-0.10                 Gviz_1.41.1
##  [55] nnet_7.3-17                   dbplyr_2.2.1
##  [57] deldir_1.0-6                  utf8_1.2.2
##  [59] tidyselect_1.1.2              rlang_1.0.4
##  [61] later_1.3.0                   AnnotationDbi_1.59.1
##  [63] munsell_0.5.0                 BiocVersion_3.16.0
##  [65] tools_4.2.1                   cachem_1.0.6
##  [67] cli_3.3.0                     generics_0.1.3
##  [69] RSQLite_2.2.16                ExperimentHub_2.5.0
##  [71] evaluate_0.16                 stringr_1.4.1
##  [73] fastmap_1.1.0                 yaml_2.3.5
##  [75] knitr_1.40                    bit64_4.0.5
##  [77] purrr_0.3.4                   randomForest_4.7-1.1
##  [79] AnnotationFilter_1.21.0       KEGGREST_1.37.3
##  [81] Rbowtie_1.37.0                mime_0.12
##  [83] xml2_1.3.3                    biomaRt_2.53.2
##  [85] compiler_4.2.1                rstudioapi_0.14
##  [87] filelock_1.0.2                curl_4.3.2
##  [89] png_0.1-7                     interactiveDisplayBase_1.35.0
##  [91] tibble_3.1.8                  stringi_1.7.8
##  [93] crisprScore_1.1.14            highr_0.9
##  [95] basilisk.utils_1.9.1          GenomicFeatures_1.49.6
##  [97] lattice_0.20-45               ProtGenerics_1.29.0
##  [99] Matrix_1.4-1                  vctrs_0.4.1
## [101] pillar_1.8.1                  lifecycle_1.0.1
## [103] BiocManager_1.30.18           data.table_1.14.2
## [105] bitops_1.0-7                  httpuv_1.6.5
## [107] R6_2.5.1                      BiocIO_1.7.1
## [109] latticeExtra_0.6-30           promises_1.2.0.1
## [111] gridExtra_2.3                 codetools_0.2-18
## [113] dichromat_2.0-0.1             crisprBowtie_1.1.1
## [115] assertthat_0.2.1              SummarizedExperiment_1.27.1
## [117] rjson_0.2.21                  GenomicAlignments_1.33.1
## [119] Rsamtools_2.13.4              GenomeInfoDbData_1.2.8
## [121] parallel_4.2.1                hms_1.1.2
## [123] rpart_4.1.16                  grid_4.2.1
## [125] basilisk_1.9.2                rmarkdown_2.15.2
## [127] MatrixGenerics_1.9.1          biovizBase_1.45.0
## [129] Biobase_2.57.1                shiny_1.7.2
## [131] base64enc_0.1-3               interp_1.1-3
## [133] restfulr_0.0.15
```
